# Impacts of genomic networks governed by human-specific regulatory sequences and genetic loci harboring fixed human-specific neuro-regulatory single nucleotide mutations on phenotypic traits of Modern Humans

**DOI:** 10.1101/848762

**Authors:** Gennadi V. Glinsky

## Abstract

Recent advances in identification and characterization of human-specific regulatory DNA sequences set the stage for the assessment of their global impact on physiology and pathology of Modern Humans. Gene set enrichment analyses (GSEA) of 8,405 genes linked with 35,074 human-specific neuro-regulatory single-nucleotide changes (hsSNCs) revealed a staggering breadth of significant associations with morphological structures, physiological processes, and pathological conditions of Modern Humans. Significantly enriched traits include more than 1,000 anatomically-distinct regions of the adult human brain, many different types of cells and tissues, more than 200 common human disorders and more than 1,000 records of rare diseases. Thousands of genes connected with neuro-regulatory hsSNCs have been identified, which represent essential genetic elements of the autosomal inheritance and offspring survival phenotypes. A total of 1,494 hsSNC- linked genes are associated with either autosomal dominant or recessive inheritance and 2,273 hsSNC-linked genes have been associated with premature death, embryonic lethality, as well as pre-, peri-, neo-, and post-natal lethality phenotypes of both complete and incomplete penetrance. Differential GSEA implemented on hsSNC-linked loci and associated genes identify 7,990 genes linked to evolutionary distinct classes of human-specific regulatory sequences (HSRS), expression of a majority of which (5,389 genes; 67%) is regulated by stem cell-associated retroviral sequences (SCARS). Interrogations of the MGI database revealed readily available mouse models tailored for precise experimental definitions of functional effects of hsSNCs and SCARS on genes causally affecting thousands of mammalian phenotypes and implicated in hundreds of common and rare human disorders. These observations suggest that a preponderance of human-specific traits evolved under a combinatorial regulatory control of HSRS and neuro-regulatory loci harboring hsSNCs that are fixed in humans, distinct from other primates, and located in differentially-accessible chromatin regions during brain development.

## Introduction

DNA sequences of coding genes defining the structure of macromolecules comprising the essential building blocks of life at the cellular and organismal levels remain highly conserved during the evolution of humans and other Great Apes (Chimpanzee Sequencing and Analysis Consortium, 2005; Kronenberg et al., 2018). In striking contrast, a compendium of nearly hundred thousand candidate human-specific regulatory sequences (HSRS) has been assembled in recent years (Glinsky et al., 2015-2020; Kanton et al., 2019), thus providing further genetic and molecular evidence supporting the idea that unique to human phenotypes may result from human-specific changes to genomic regulatory sequences defined as “regulatory mutations” (King and Wilson, 1975). Structurally, functionally, and evolutionary distinct classes of HSRS appear to cooperate in shaping developmentally and physiologically diverse human-specific genomic regulatory networks (GRNs) impacting preimplantation embryogenesis, pluripotency, and development and functions of human brain (Glinsky, 2020). The best evidence of the exquisite degree of accuracy of the contemporary molecular definition of human-specific regulatory sequences is exemplified by the identification of 35,074 single nucleotide changes (SNCs) that are fixed in humans, distinct from other primates, and located within differentially-accessible (DA) chromatin regions during the human brain development in cerebral organoids (Kanton et al., 2019). Therefore, this type of mutations could be defined as fixed neuro-regulatory human-specific single nucleotide changes (hsSNCs). However, only a small fraction of identified DA chromatin peaks (600 of 17,935 DA peaks; 3.3%) manifest associations with differential expression in human versus chimpanzee cerebral organoids model of brain development, consistent with the hypothesis that regulatory effects on gene expression of these DA chromatin regions are not restricted to the early stages of brain development. Annotation of SNCs derived and fixed in modern humans that overlap DA chromatin regions during brain development revealed that essentially all candidate regulatory human-specific SNCs are shared with the archaic humans (35,010 SNCs; 99.8%) and only 64 SNCs are unique to modern humans (Kanton et al., 2019). This remarkable conservation on the human lineage of human-specific SNCs associated with human brain development sows the seed of interest for in-depth exploration of coding genes expression of which may be affected by genetic regulatory loci harboring human-specific SNCs.

In this contribution, the GREAT algorithm (McLean et al., 2010, 2011) was utilized to identify 8,405 hsSNCs-linked genes expression of which might be affected by 35,074 human-specific SNCs located in DA chromatin regions during brain development. Comprehensive gene set enrichment analyses (GSEA) of these genes revealed the staggering breadth of associations with physiological processes and pathological conditions of *H. sapiens*, including more than 1,000 anatomically-distinct regions of the adult human brain, many human tissues and cell types, more than 200 common human disorders and more than 1,000 rare diseases. It has been concluded that hsSNCs-linked genes appear contributing to development and functions of the adult human brain and other components of the central nervous system; they were defined as genetic markers of many tissues across human body and were implicated in the extensive range of human physiological and pathological conditions, thus supporting the hypothesis that phenotype-altering effects of neuro-regulatory hsSNCs are not restricted to the early-stages of human brain development. Differential GSEA implemented on hsSNC-linked loci and associated genes identify 7,990 genes linked to evolutionary distinct classes of human-specific regulatory sequences (HSRS). Notably, expression of a majority of this common set of genes (5,389 genes; 67%) is regulated by stem cell-associated retroviral sequences (SCARS). Collectively, observations reported in this contribution indicate that structurally, functionally and evolutionary diverse classes of HSRS, neuro-regulatory hsSNCs, and associated elite set of 7,990 genes affect wide spectra of traits defining both physiology and pathology of Modern Humans by asserting human-specific regulatory impacts on thousands essential mammalian phenotypes.

## Results

### Identification and characterization of putative genetic regulatory targets associated with human-specific single nucleotide changes (SNCs) in in differentially accessible (DA) chromatin regions during brain development

To identify and characterize human genes associated with 35,074 human-specific single nucleotide changes (SNCs) in differentially accessible (DA) chromatin regions during human and chimpanzee neurogenesis in cerebral organoids (Kanton et al., 2019), the GREAT algorithm (McLean et al., 2011) have been employed. These analyses identified 8,405 genes with putative regulatory connections to human-specific SNCs (Figure 1) and revealed a remarkable breadth of highly significant associations with a multitude of biological processes, molecular functions, genetic and metabolic pathways, cellular compartments, and gene expression perturbations (Supplemental Table Set S1).

**Figure 1.**
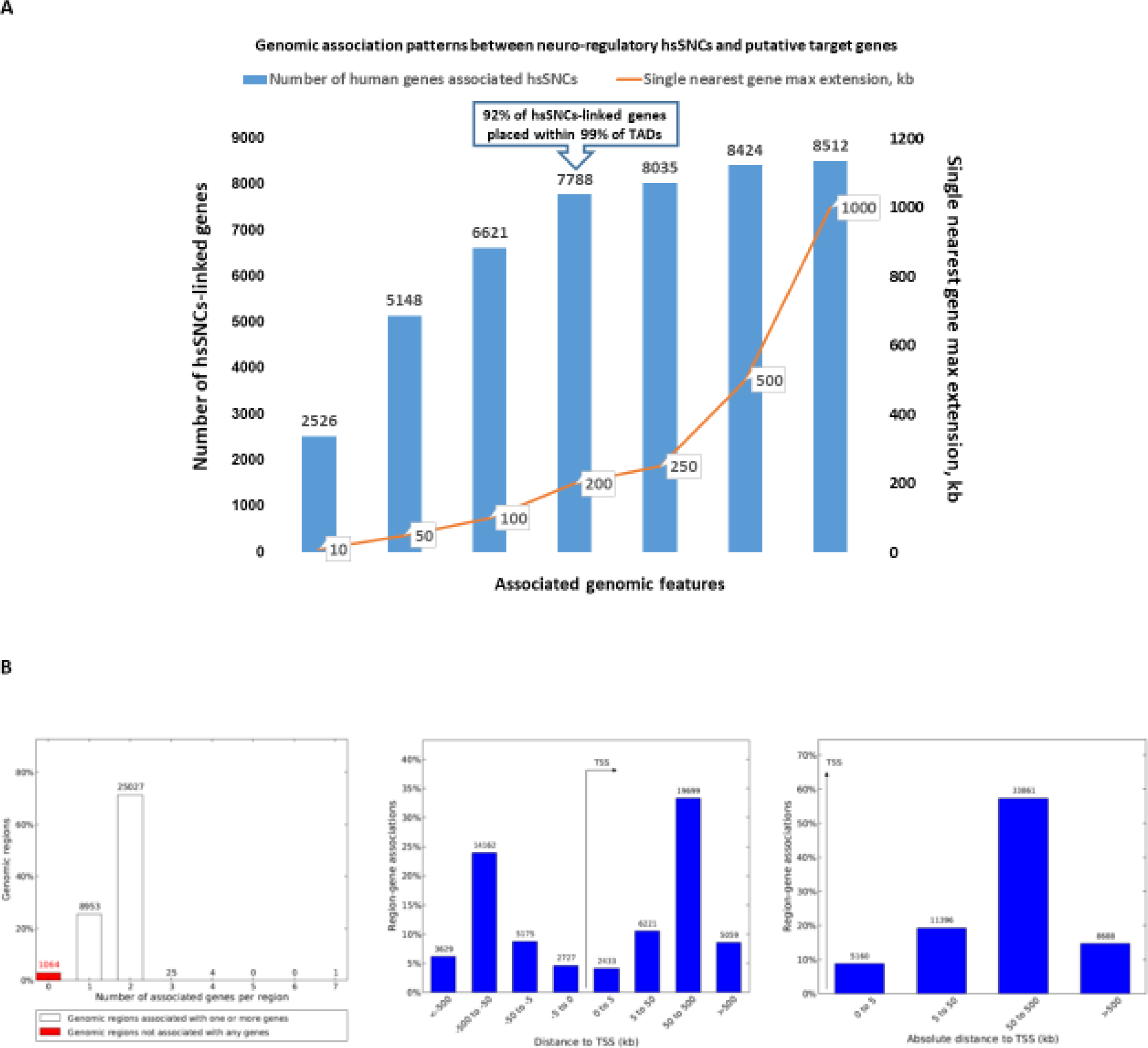
GREAT analysis identifies 8,405 human genes associated with 35,074 neuro-regulatory human-specific single nucleotide changes (hsSNCs) identified in differentially accessible (DA) chromatin regions during human and chimpanzee brain development in cerebral organoids. A. Patterns of genomic associations between neuro-regulatory hsSNCs and putative target genes defined at different single nearest gene maximum extensions. GREAT algorithm version 4.0.4. B. A total of 1,064 of all 35,074 SNCs (3%) are not associated with any genes in the human genome, while a total of 34,010 (97%) human-specific SNCs in DA regions appear associated with 8,405 human genes. GREAT algorithm version 3.0.0.

To ascertain patterns of genomic associations between neuro-regulatory human-specific SNCs and putative target genes, the GREAT analyses were performed at different proximity placement distances defined by the single nearest gene maximum extension ranging from 10 Kb to 1 Mb (Figure 1). It has been observed that from 92% of all hsSNCs-linked genes are located within 200 Kb distances from their putative regulatory loci (Figure 1A). Since the size of more than 99% of topologically-associating domains (TADs) in human genomes is 200 kb or more (Dixon et al., 2012), these findings indicate that a marked majority of neuro-regulatory hsSNCs and their putative target genes would be located in human genomes within the boundaries of the same TAD.

It has been noted that particularly striking numbers of significant associations were uncovered by the GREAT algorithm during the analyses of two databases:

1. The Human Phenotype Ontology containing over 13,000 terms describing clinical phenotypic abnormalities that have been observed in human diseases, including hereditary disorders (326 significant records with binominal FDR Q-Value < 0.05);
2. The MGI Expression Detected ontology referencing genes expressed in specific anatomical structures at specific developmental stages (Theiler stages) in the mouse (370 significant records with binominal FDR Q-Value < 0.05).

These observations support the hypothesis that biological functions of genes under the putative regulatory control of human-specific SNCs in DA chromatin regions during brain development are not limited to the contribution to the early stages of neuro- and corticogenesis. Collectively, findings reported in the Supplemental Table Set S1 argue that genes expression of which is affected by human-specific SNCs may represent a genomic dominion of putative regulatory dependency from HSRS that is likely to play an important role in a broad spectrum of pathological conditions of Modern Humans.

### Identification of hsSNCs-linked genes expression of which distinguishes thousands of anatomically distinct areas of the adult human brain, various regions of the central nervous system, and many different cell types and tissues in the human body

To validate and extend these observations, next the comprehensive gene set enrichment analyses were performed employing the web-based Enrichr API bioinformatics platform (Chen et al., 2013; Kuleshov et al., 2016), which interrogated nearly 200,000 gene sets from more than 100 gene set libraries. The results of these analyses are summarized in the Table 1 and reported in details in the Supplemental Table Set S2. Genes expression of which were placed during evolution under the regulatory control of ∼ 35,000 neuro-regulatory human-specific SNCs demonstrate a staggering breadth of significant associations with a broad spectrum of anatomically distinct regions, various cell and tissue types, a multitude of physiological processes, and a numerous pathological conditions of *H. sapiens*.

**Table 1.**
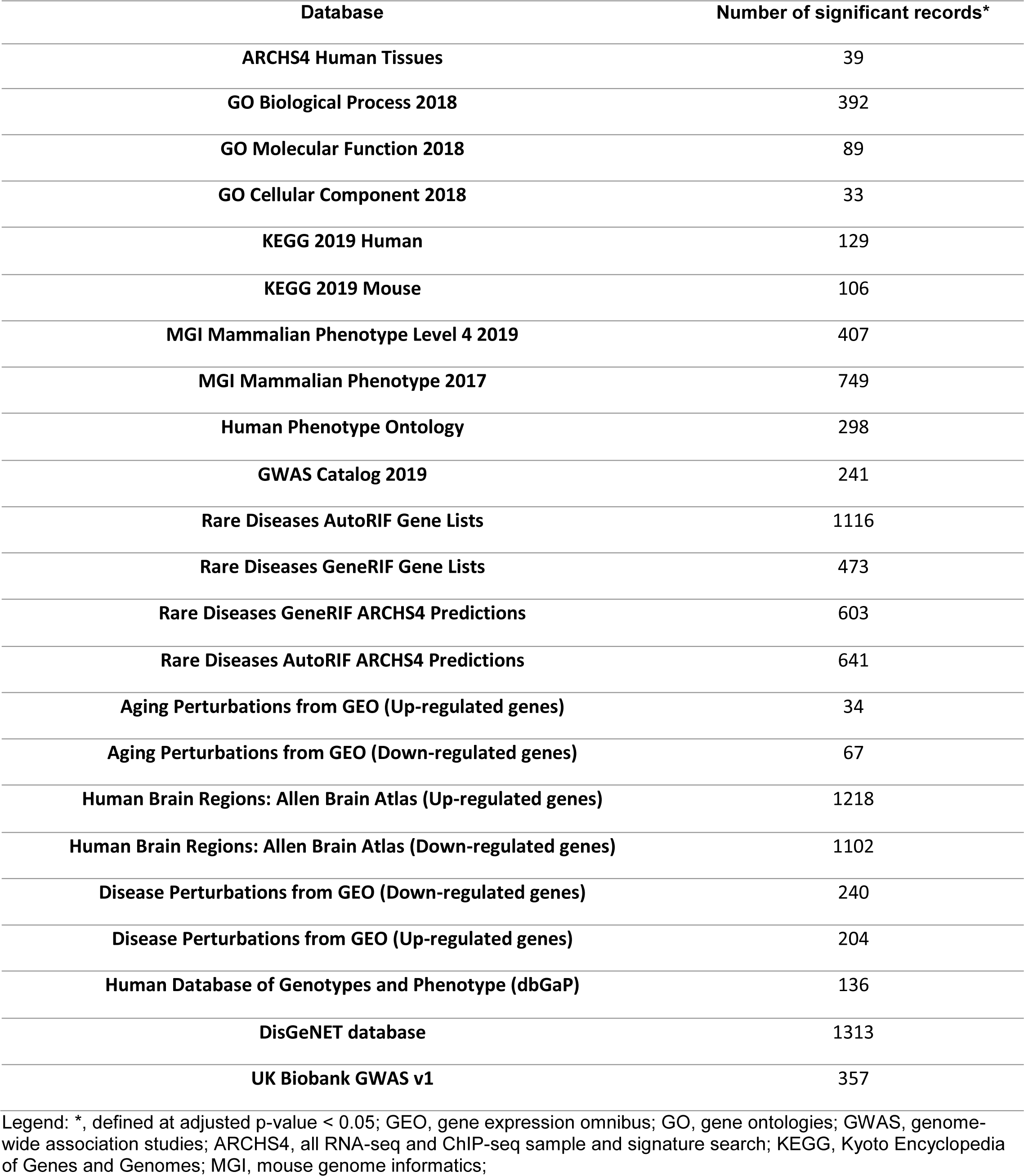
Associations with human physiological processes and pathological conditions of 8,405 genes linked with 35,074 human-specific single nucleotide changes (SNC) within differentially-accessible (DA) chromatin regions identified during human and chimpanzee brain development in cerebral organoids.

| Database | Number of significant records* |
| --- | --- |
| ARCHS4 Human Tissues | 39 |
| GO Biological Process 2018 | 392 |
| GO Molecular Function 2018 | 89 |
| GO Cellular Component 2018 | 33 |
| KEGG 2019 Human | 129 |
| KEGG 2019 Mouse | 106 |
| MGI Mammalian Phenotype Level 4 2019 | 407 |
| MGI Mammalian Phenotype 2017 | 749 |
| Human Phenotype Ontology | 298 |
| GWAS Catalog 2019 | 241 |
| Rare Diseases AutoRIF Gene Lists | 1116 |
| Rare Diseases GeneRIF Gene Lists | 473 |
| Rare Diseases GeneRIF ARCHS4 Predictions | 603 |
| Rare Diseases AutoRIF ARCHS4 Predictions | 641 |
| Aging Perturbations from GEO (Up-regulated genes) | 34 |
| Aging Perturbations from GEO (Down-regulated genes) | 67 |
| Human Brain Regions: Allen Brain Atlas (Up-regulated genes) | 1218 |
| Human Brain Regions: Allen Brain Atlas (Down-regulated genes) | 1102 |
| Disease Perturbations from GEO (Down-regulated genes) | 240 |
| Disease Perturbations from GEO (Up-regulated genes) | 204 |
| Human Database of Genotypes and Phenotype (dbGaP) | 136 |
| DisGeNET database | 1313 |
| UK Biobank GWAS v1 | 357 |

Legend: \*, defined at adjusted p-value < 0.05; GEO, gene expression omnibus; GO, gene ontologies; GWAS, genome-wide association studies; ARCHS4, all RNA-seq and ChIP-seq sample and signature search; KEGG, Kyoto Encyclopedia of Genes and Genomes; MGI, mouse genome informatics;

Of particular interest is the apparent significant enrichment of human-specific SNCs-associated genes among both up-regulated and down-regulated genes, expression of which discriminates thousands of anatomically distinct areas of the adult human brain defined in the Allen Brain Atlas (Supplemental Figure S1; Supplemental Table Set S2). Notably, genes expressed in various thalamus regions appear frequently among the top-scored anatomical areas of the human brain (Supplemental Figure S1; Supplemental Table Set S2). These findings were further corroborated by the identification of hsSNCs-linked genes among genetic markers of 26 human brain regions using the Allen Human Brain Atlas database (Figure 2). Notably, a significant majority of hsSNCs-linked genes (6,640 of 8,405 genes; 79%) represents genetic markers of 26 human brain regions examined in this study (Figure 2). In agreement with the hypothesis that neuro-regulatory hsSNCs may exert the human-specific regulatory effects on target genes, a significant fraction of hsSNCs-linked genes (3,212 genes; 38%) manifest significant expression changes in human versus chimpanzee adult brains (Figure 2B).

**Figure 2.**
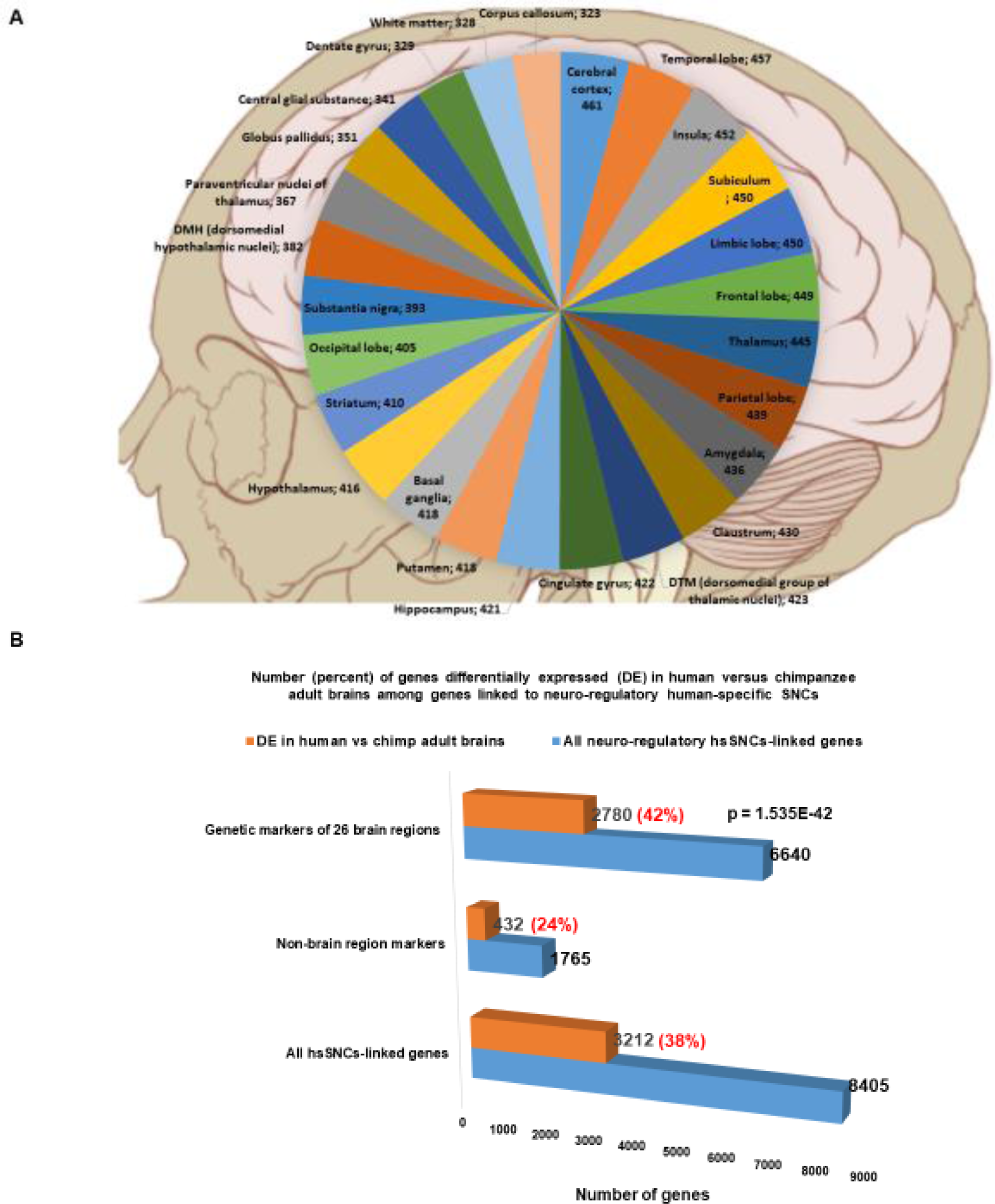
A dominant majority (6,640 of 8,405 genes; 79%) of genes linked to 35,074 human-specific single nucleotide changes (hsSNCs) in chromatin’s differentially accessible (DA) regions during human and chimpanzee brain development in cerebral organoids represents genetic markers of 26 human brain regions. A. Number of brain regions’ marker genes linked to 35,074 neuro-regulatory hsSNCs in specified human brain regions (the normalized values calculated per 1,000 region-specific marker genes are shown). Genes linked to hsSNCs were identified among genes significantly up-regulated in specified human brain regions using the Allen Brain Atlas database (records manifesting increased expression at 1.5-fold cut-off were identified and selected for analyses). B. Number (percent) of genes differentially expressed (DE) in human versus chimpanzee adult brains among genes linked to neuro-regulatory human-specific SNCs. Genes linked to hsSNCs were identified among genes differentially expressed in eight regions of human versus chimpanzee adult brains (Xu et al., 2018).

A dominant majority of hsSNCs-linked genes manifesting differential expression in human versus chimpanzee adult brains represents genetic markers of human brain regions (2,780 of 3,212 genes; 87%). The fraction of hsSNCs-linked genes differentially expressed in human versus chimpanzee brains is significantly higher among human brain regions’ marker genes compared to non-markers (Figure 2B; p = 1,535E-42; 2-tail Fisher’s exact test). These observations support the hypothesis that genetic loci harboring human-specific SNCs may exert regulatory effects on structural and functional features of the adult human brain, thus, likely affecting the development and functions of the central nervous system in Modern Humans. Consistent with this idea, the examination of the enrichment patterns of human-specific SNCs-associated genes in the ARCHS4 Human Tissues’ gene expression database revealed that top 10 most significantly enriched records overlapping a majority of region-specific marker genes constitute various anatomically-distinct regions of the central nervous system (Supplemental Figure 1; Supplemental Table Set S2). However, results of gene set enrichment analyses convincingly demonstrate that inferred regulatory effects of genetic loci harboring human-specific SNCs are not restricted only to the various regions of the central nervous system, they appear to affect gene expression profiles of many different cell types and tissues in the human body (Table 1; Supplemental Table Set S2).

### Identification and characterization of hsSNCs-linked genes expression of which is altered during aging of humans, rats, and mice

Genes altered expression of which is implicated in the aging of various tissues and organs of humans, rats, and mice are significantly enriched among 8,405 genes associated with human-specific regulatory SNCs (Supplemental Figure S2; Supplemental Table Set S2).

Aging of the hippocampus was implicated most frequently among genes manifesting increased expression with age, while among genes exhibiting aging-associated decreased expression the hippocampus and frontal cortex were identified repeatedly (Supplemental Figure S2). Overall, twice as many significant association records were observed among aging-associated down-regulated genes compared to up-regulated genes (Table 1). Collectively, these observations indicate that genes changes in expression of which were associated with aging in mammals, in particular, hippocampal and frontal cortex aging, represent important elements of a genomic dominion that was placed under regulatory control of genetic loci harboring human-specific SNCs.

### Identification of hsSNCs-linked genes implicated in development and manifestations of hundreds physiological and pathological phenotypes and autosomal inheritance in Modern Humans

Interrogations of the Human Phenotype Ontology database (298 significantly enriched records identified), the Genome-Wide Association Study (GWAS) Catalogue (241 significantly enriched records identified), and the database of Human Genotypes and Phenotypes (136 significantly enriched records identified) revealed several hundred physiological and pathological phenotypes and thousands of genes manifesting significant enrichment patterns defined at the adjusted p value < 0.05 (Supplemental Figure S3; Table 1; Supplemental Table Set S2). Interestingly, 645 and 849 genes implicated in the autosomal dominant (HP:0000006) and recessive (HP:0000007) inheritance were identified amongst genes associated with human-specific regulatory SNCs (Supplemental Figure S3; Supplemental Table Set S2). Notable pathological conditions among top-scored records identified in the database of Human Genotypes and Phenotypes are stroke, myocardial infarction, coronary artery disease, and heart failure (Supplemental Figure S3).

A total of 241 significantly enriched records (Table 1) were documented by gene set enrichment analyses of the GWAS catalogue (2019), among which a highly diverse spectrum of pathological conditions linked to genes associated with human-specific regulatory SNCs was identified, including obesity, type 2 diabetes, amyotrophic lateral sclerosis, autism spectrum disorders, attention deficit hyperactivity disorder, bipolar disorder, major depressive disorder, schizophrenia, Alzheimer’s disease, malignant melanoma, diverticular disease, asthma, coronary artery disease, glaucoma, as well as breast, prostate and colorectal cancers (Supplemental Figure S3; Supplemental Table Set S2). These observations indicate that thousands of genes putatively associated with genetic regulatory loci harboring human-specific SNCs affect risk of developing numerous pathological conditions in Modern Humans.

### Identification of hsSNCs-linked genes expression of which is altered in several hundred common human disorders

Gene set enrichment analyses-guided interrogation of the Gene Expression Omnibus (GEO) database revealed the remarkably diverse spectrum of human diseases with the etiologic origins in multiple organs and tissues and highly heterogeneous pathophysiological trajectories of their pathogenesis (Supplemental Figure S4; Supplemental Table Set S2). Overlapping gene sets between disease-associated genes and human-specific SNCs-linked genes comprise of hundreds of genes that were either up-regulated (204 significant disease records) or down-regulated (240 significant disease records) in specific pathological conditions, including schizophrenia, bipolar disorder, various types of malignant tumors, Crohn’s disease, ulcerative colitis, Down syndrome, Alzheimer’s disease, spinal muscular atrophy, multiple sclerosis, autism spectrum disorders, type 2 diabetes mellitus, morbid obesity, cardiomyopathy (Supplemental Figure S4; Supplemental Table Set S2). These observations demonstrate that thousands of genes expression of which is altered in a myriad of human diseases appear associated with genetic regulatory loci harboring human-specific SNCs. Consistent with the hypothesis that genes linked to human-specific neuro-regulatory SNCs represent a network of essential genetic loci implicated in a broad spectrum of physiological and pathological traits of Modern Humans, nearly half of human genes (162 of 332 genes; 49%) encoding prey proteins for SARS-CoV-2 coronavirus are members of this gene set (data not shown).

### Identification of hsSNCs-linked genes implicated in more than 1,000 records classified as human rare diseases

Present analyses demonstrate that thousands of genes associated with human-specific regulatory SNCs have been previously identified as genetic elements affecting the likelihood of development a multitude of common human disorders. Similarly, thousands of genes expression of which is altered during development and manifestation of multiple common human disorders appear linked to genetic regulatory loci harboring human-specific SNCs. Remarkably, interrogations of the Enrichr’s libraries of genes associated with Modern Humans’ rare diseases identified 473, 603, 641, and 1,116 significantly enriched records of various rare disorders employing the Rare Diseases GeneRIF gene lists library, the Rare Diseases GeneRIF ARCHS4 predictions library, the Rare Diseases AutoRIF ARCHS4 predictions library, and the Rare Diseases AutoRIF Gene lists library, respectively (Supplemental Figure S5; Supplemental Table Set S2). Taken together, these observations demonstrate that thousands of genes associated with hundreds of human rare disorders appear linked with human-specific regulatory SNCs.

### Gene ontology analyses of putative regulatory targets of genetic loci harboring human-specific SNCs

Gene Ontology (GO) analyses identified a constellation of biological processes (GO Biological Process: 308 significant records) supplemented with a multitude of molecular functions (GO Molecular Function: 81 significant records) that appear under the regulatory control of human-specific SNCs (Supplemental Figure S6; Supplemental Table Set 2). Consistently, both databases identified frequently the components of transcriptional regulation and protein kinase activities among most significant records. Other significantly enriched records of interest are regulation of apoptosis, cell proliferation, migration, and various binding properties (cadherin binding; sequence-specific DNA binding; protein-kinase binding; amyloid-beta binding; actin binding; tubulin binding; microtubule binding; PDZ domain binding) which are often supplemented by references to the corresponding activity among the enriched records, for example, enriched records of both binding and activity of protein kinases.

Interrogation of GO Cellular Component database identified 29 significantly enriched records, among which nuclear chromatin as well as various cytoskeleton and membrane components appear noteworthy (Supplemental Figure S6). Both GO Biological Process and GO Cellular Component database identified significantly enriched records associated with the central nervous system development and functions such as axonogenesis and axon guidance; generation of neurons, neuron differentiation, and neuron projection morphogenesis; cellular components of dendrites and dendrite’s membrane; ionotropic glutamate receptor complex. In several instances biologically highly consistent enrichment records have been identified in different GO databases: cadherin binding (GO Molecular Function) and catenin complex (GO Cellular Component); actin binding (GO Molecular Function) and actin cytoskeleton, cortical actin cytoskeleton, actin-based cell projections (GO Cellular Component); microtubule motor activity, tubulin binding, microtubule binding (GO Molecular Function) and microtubule organizing center, microtubule cytoskeleton (GO Cellular Component).

Analyses of human and mouse databases of the Kyoto Encyclopedia of Genes and Genomes (KEGG; Supplemental Figure S7) identified more than 100 significantly enriched records in each database (KEGG 2019 Human: 129 significant records; KEGG 2019 Mouse: 106 significant records). Genes associated with human-specific regulatory SNCs were implicated in a remarkably broad spectrum of signaling pathways ranging from pathways regulating the pluripotency of stem cells to cell type-specific morphogenesis and differentiation pathways, for example, melanogenesis and adrenergic signaling in cardiomyocytes (Supplemental Figure S7). Genes under putative regulatory control of human-specific SNCs include hundreds of genes contributing to specific functions of specialized differentiated cells (gastric acid secretion; insulin secretion; aldosterone synthesis and secretion), multiple receptor/ligand-specific signaling pathways, as well as genetic constituents of pathways commonly deregulated in cancer and linked to the organ-specific malignancies, for example, breast, colorectal, and small cell lung cancers (Supplemental Figure S7). Other notable entries among most significantly enriched records include pathways of the axon guidance; dopaminergic, glutamatergic, and cholinergic synapses; neuroactive receptor-ligand interactions; and AGE-RAGE signaling pathway in diabetic complications (Supplemental Figure S7; Supplemental Table Set 2).

### Identification of 2,273 genes associated with human-specific SNCs and implicated in premature death and embryonic, prenatal, perinatal, neonatal, and postnatal lethality phenotypes

Interrogation of MGI Mammalian Phenotype databases revealed several hundred mammalian phenotypes affected by thousands of genes associated with genomic regulatory regions harboring human-specific SNCs: the MGI Mammalian Phenotype (2017) database identified 749 significant enrichment records, while the MGI Mammalian Phenotype Level 4 (2019) database identified 407 significant enrichment records (Table 2; Supplemental Figure S8; Supplemental Table Set S2). Strikingly, among the records of mammalian phenotypes identified by the gene set enrichment analyses of neuro-regulatory hsSNCs-linked genes, there are 2,273 genes mutations of which result in phenotypes of premature death, embryonic lethality, as well as prenatal, perinatal, neonatal, and postnatal lethality of both complete and incomplete penetrance. A significant fraction of these 2,273 genes, which collectively could be defined based on patterns of phenotypes caused by their mutations as an offspring survival genomic dominion, was implicated in the autosomal dominant (389 genes) and recessive (426 genes) inheritance in Modern Humans. Based on these observations, it has been concluded that thousands of genes within the genomic dominions of putative regulatory dependencies from human-specific SNCs represent the essential genetic elements of the mammalian offspring survival phenotypes.

**Table 2.**
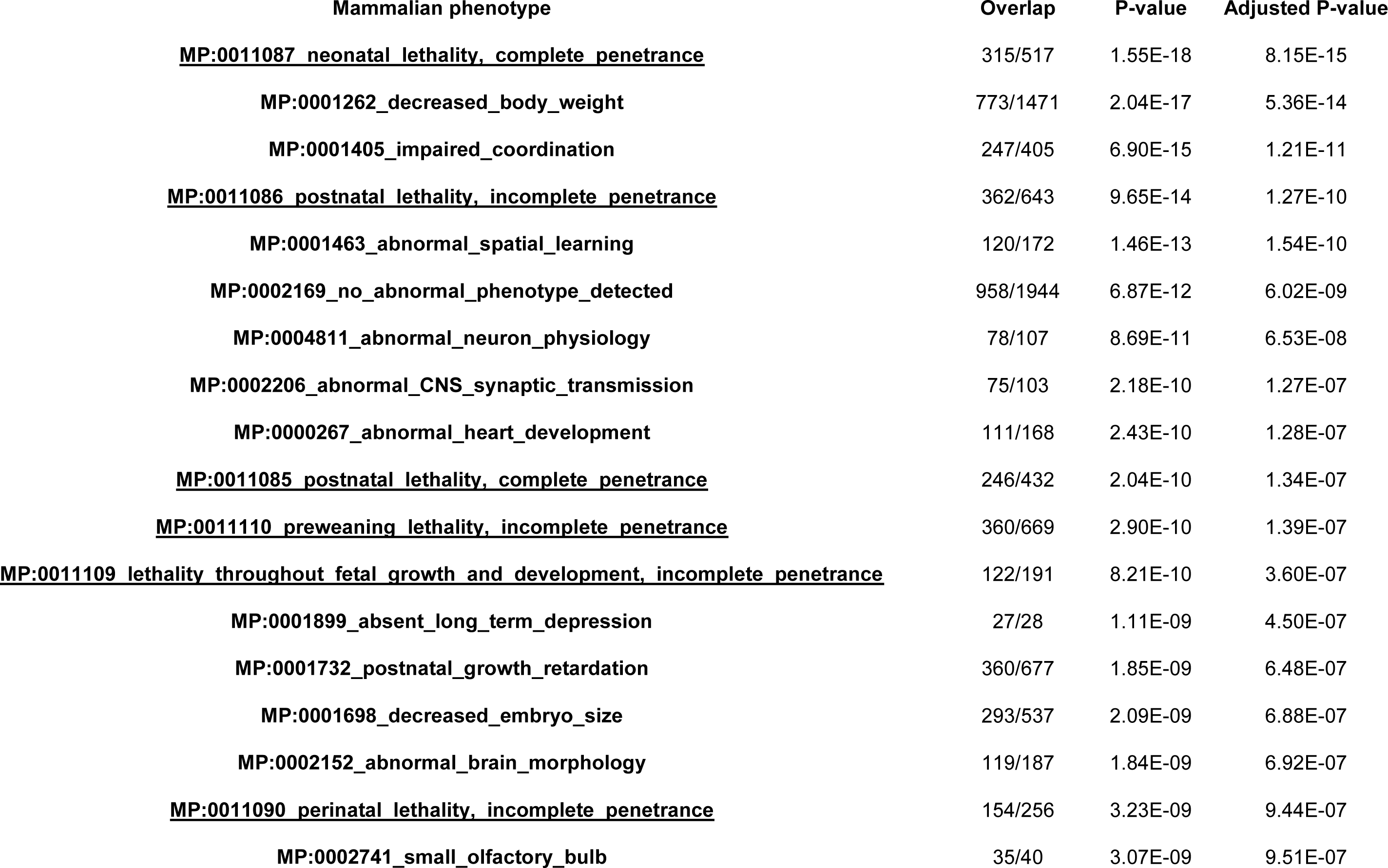
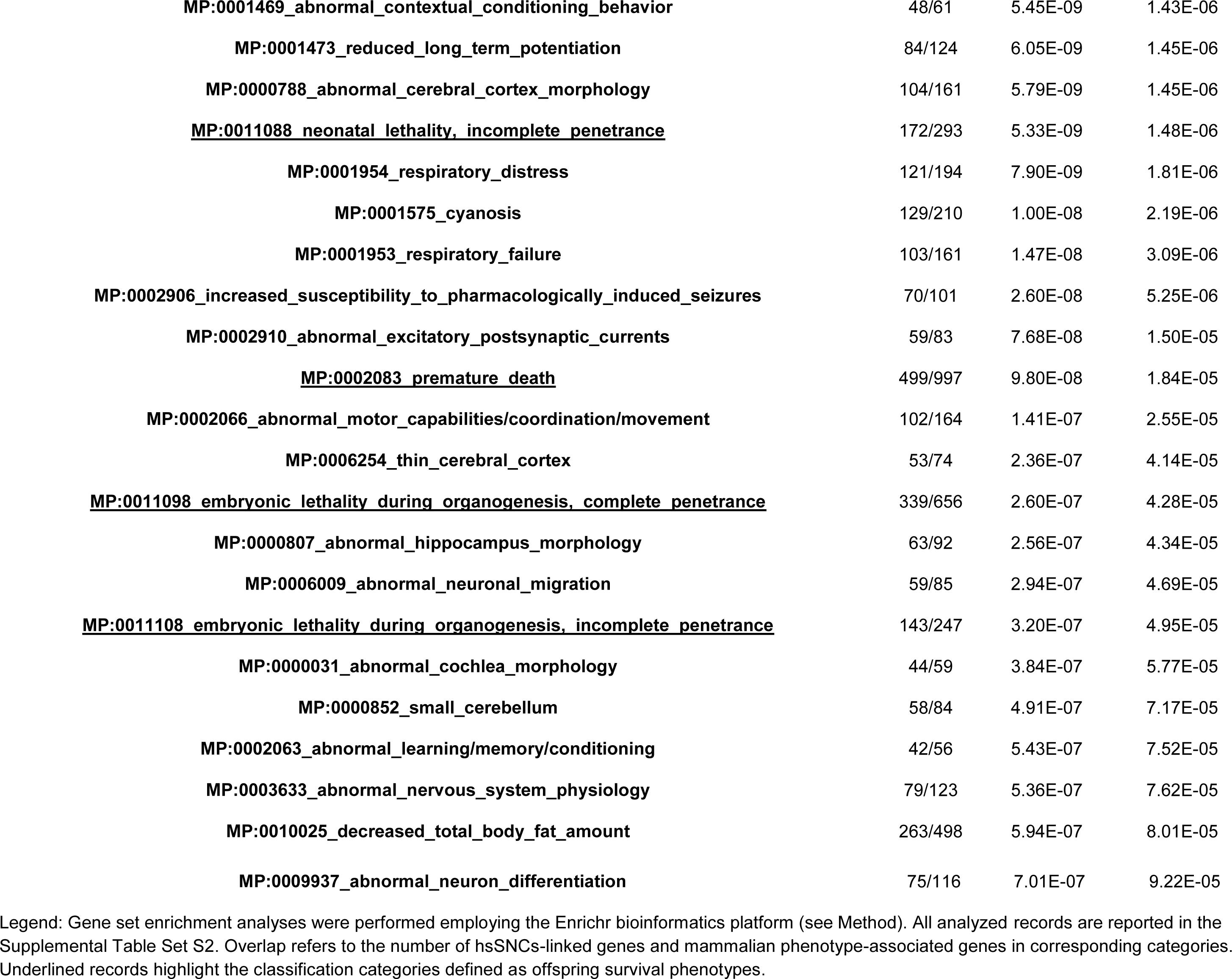
Gene set enrichment analyses of the MGI Mammalian Phenotype Level 4 (2019) database identify mammalian phenotypes manifesting significant associations with neuro-regulatory human-specific SNCs-linked genes. Top 40 of the 407 significant records are reported.

### Identification of the experimentally tractable models for molecular definitions of regulatory effects of human-specific SNCs on expression of genes associated with thousands of mammalian phenotypes and human diseases

To identify all genes linked with human-specific neuro-regulatory SNCs that are associated with defined mammalian phenotypes and human diseases with one or more mouse models, the analyses have been carried out utilizing the Mouse Genome Informatics (MGI) database (http://www.informatics.jax.org/). These analyses identified 125,938 Mammalian Phenotype Ontology records and 1,807 Human Disease Ontology records associated with 5,730 and 1,162 human-specific regulatory SNCs-linked genes, respectively (Supplemental Table Sets S4 and S5). Remarkably, genes linked with human-specific regulatory SNCs have been associated with a majority (61%) of all human diseases with one or more mouse models (967 of 1,584 human disease ontology terms; Supplemental Table Set S4). Similarly, human-specific SNCs-linked genes have been associated with 71% of all Mammalian Phenotype Ontology terms (9,190 of 12,936 records; Supplemental Table Set S5). These observations identify readily available mouse models for experimental interrogations of regulatory effects of human-specific SNCs and other types of HSRS on genes causally affecting thousands of defined mammalian phenotypes and hundreds of common and rare human disorders.

### Structurally, functionally, and evolutionary distinct classes of human-specific regulatory sequences (HSRS) share the relatively restricted elite set of common genetic targets

It has been suggested that unified activities of thousands candidate HSRS comprising a coherent compendium of genomic regulatory elements markedly distinct in their structure, function, and evolutionary origin may have contributed to development and manifestation of human-specific phenotypic traits (Glinsky, 2020). It was interest to determine whether genes previously linked to other classes of HSRS, which were identified without considerations of human-specific neuro-regulatory SNCs, overlap with genes associated in this contribution with genomic regulatory loci harboring human-specific SNCs. To this end, genes associated with different classes of HSRS were identified using the GREAT algorithm, subjected to the GSEA, and compared with the set of 8,405 genes linked with neuro-regulatory hsSNCs. Notably, all classes of HSRS appear to share common sub-sets of putative genetic regulatory targets with neuro-regulatory hsSNCs (Table 3; Figure 3; Supplemental Figure S9; Supplemental Table Set S3). GSEA of genes linked with different classes of HSRS revealed strikingly similar patterns of associations with human phenotypic traits (Supplemental Notes 1 & 2), which recapitulate, in part, the patterns of phenotypic associations of neuro-regulatory hsSNC-linked genes.

**Table 3.**
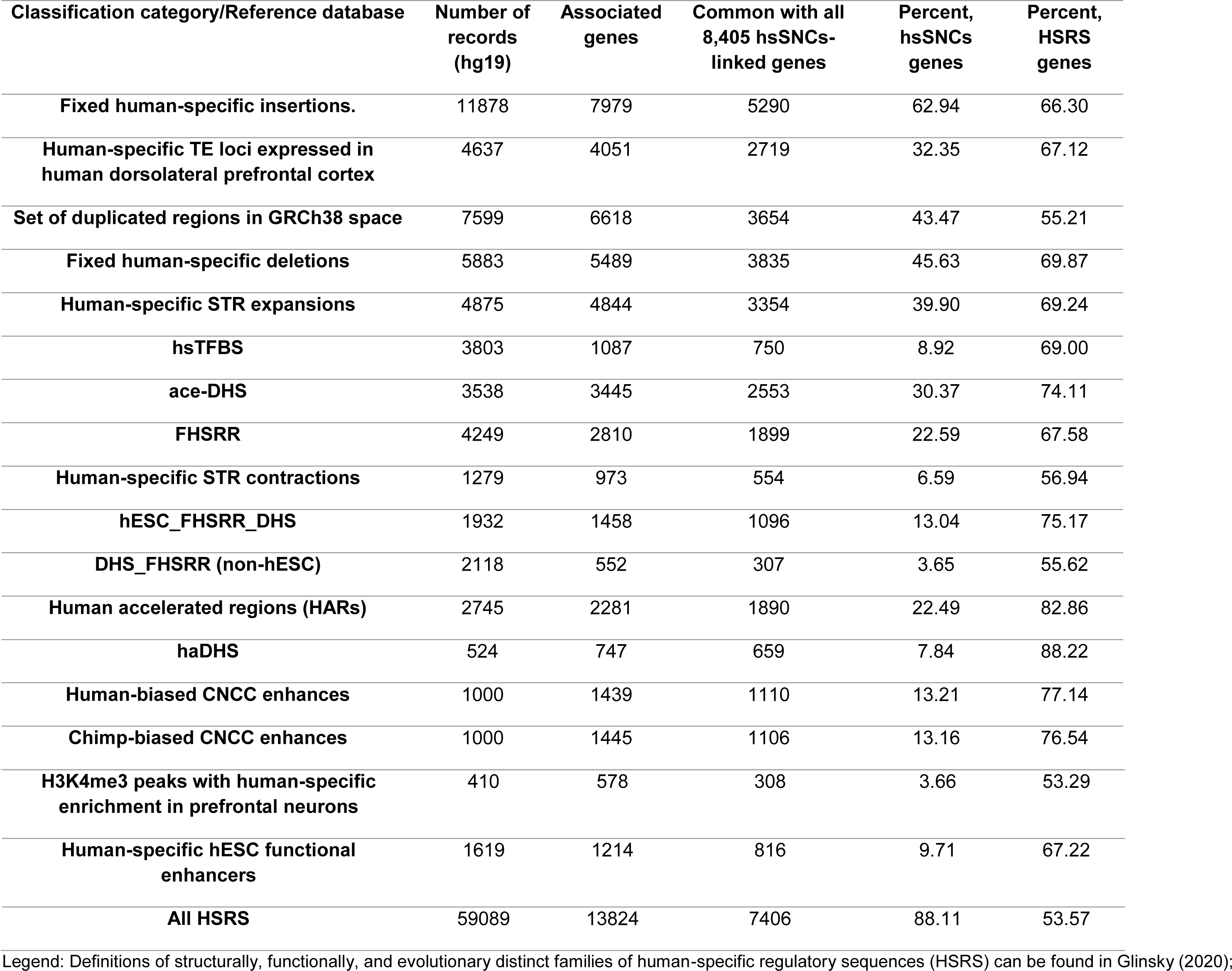
Structurally, functionally, and evolutionary distinct families of human-specific regulatory sequences (HSRS) manifest common enrichment patterns of associations with 8,405 hsSNCs-linked genes.

| **Classification category/Reference database** | **Number of records (hg19)** | **Associated genes** | **Common with all 8,405 hsSNCs-linked genes** | **Percent, hsSNCs genes** | **Percent, HSRS genes** |
| --- | --- | --- | --- | --- | --- |
| **Fixed human-specific insertions.** | 11878 | 7979 | 5290 | 62.94 | 66.30 |
| **Human-specific TE loci expressed in human dorsolateral prefrontal cortex** | 4637 | 4051 | 2719 | 32.35 | 67.12 |
| **Set of duplicated regions in GRCh38 space** | 7599 | 6618 | 3654 | 43.47 | 55.21 |
| **Fixed human-specific deletions** | 5883 | 5489 | 3835 | 45.63 | 69.87 |
| **Human-specific STR expansions** | 4875 | 4844 | 3354 | 39.90 | 69.24 |
| **hsTFBS** | 3803 | 1087 | 750 | 8.92 | 69.00 |
| **ace-DHS** | 3538 | 3445 | 2553 | 30.37 | 74.11 |
| **FHSRR** | 4249 | 2810 | 1899 | 22.59 | 67.58 |
| **Human-specific STR contractions** | 1279 | 973 | 554 | 6.59 | 56.94 |
| **hESC_FHSRR_DHS** | 1932 | 1458 | 1096 | 13.04 | 75.17 |
| **DHS_FHSRR (non-hESC)** | 2118 | 552 | 307 | 3.65 | 55.62 |
| **Human accelerated regions (HARs)** | 2745 | 2281 | 1890 | 22.49 | 82.86 |
| **haDHS** | 524 | 747 | 659 | 7.84 | 88.22 |
| **Human-biased CNCC enhances** | 1000 | 1439 | 1110 | 13.21 | 77.14 |
| **Chimp-biased CNCC enhances** | 1000 | 1445 | 1106 | 13.16 | 76.54 |
| **H3K4me3 peaks with human-specific enrichment in prefrontal neurons** | 410 | 578 | 308 | 3.66 | 53.29 |
| **Human-specific hESC functional enhancers** | 1619 | 1214 | 816 | 9.71 | 67.22 |
| **All HSRS** | 59089 | 13824 | 7406 | 88.11 | 53.57 |

Legend: Definitions of structurally, functionally, and evolutionary distinct families of human-specific regulatory sequences (HSRS) can be found in Glinsky (2020);

**Figure 3.**
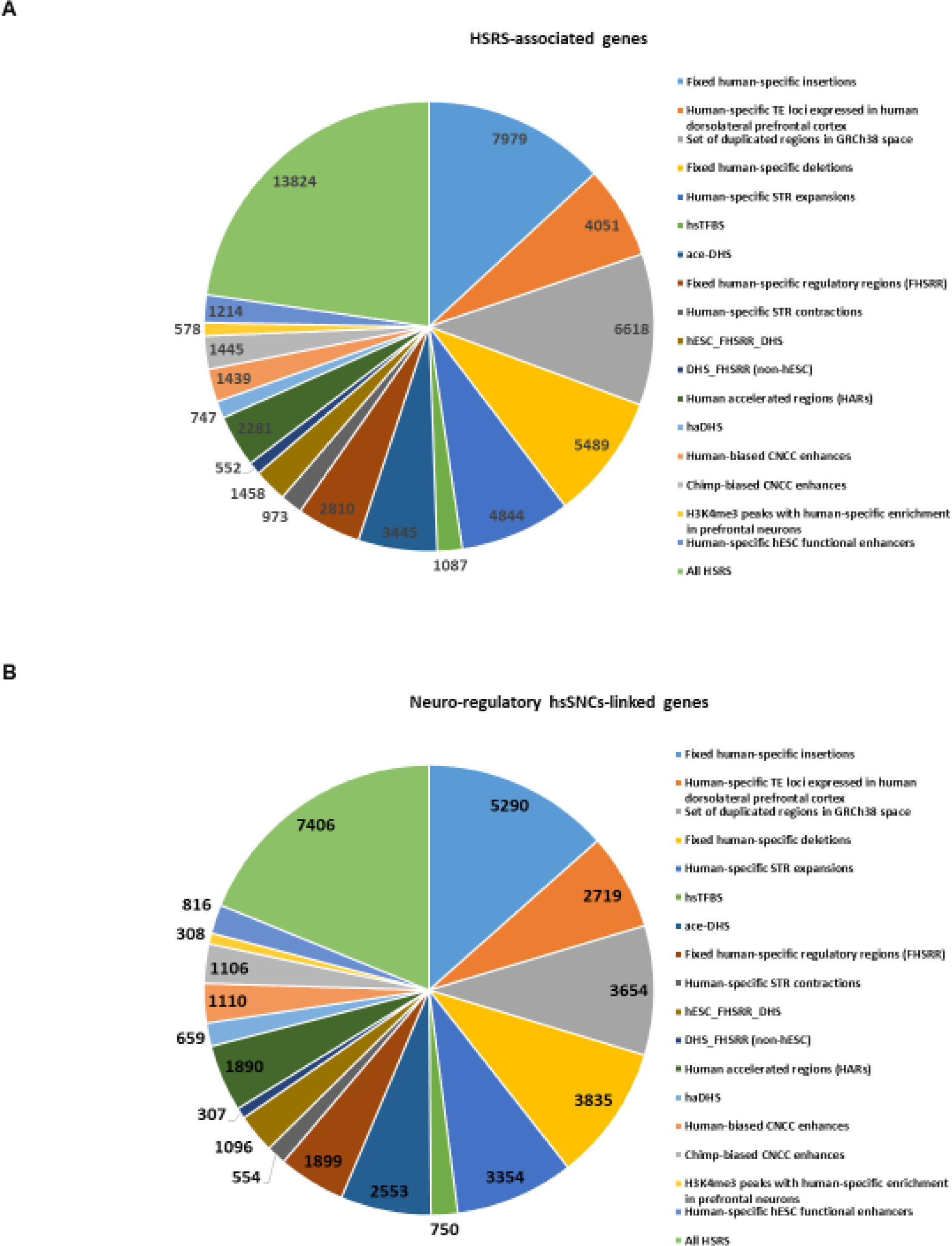
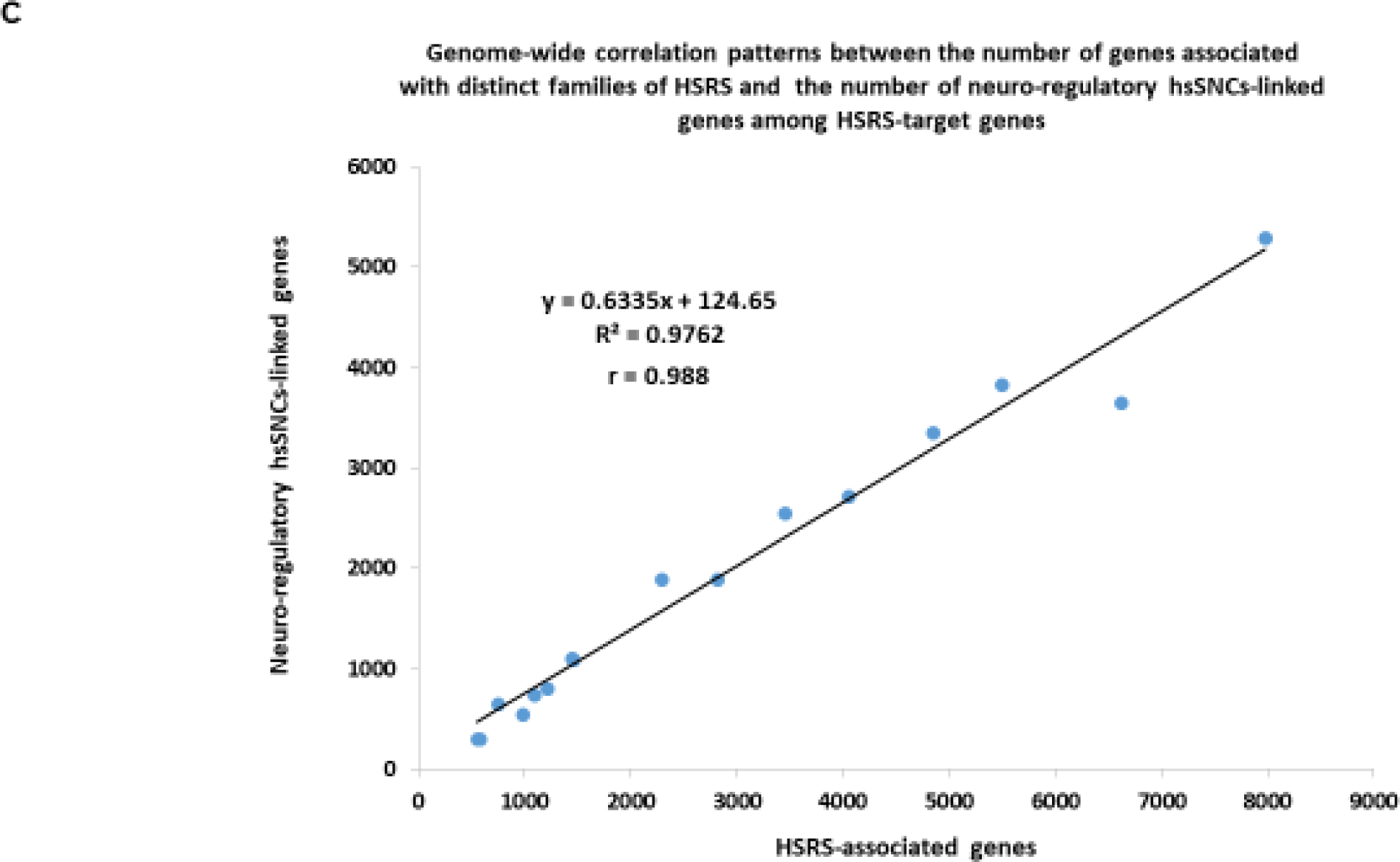
Distinct families of regulatory DNA sequences comprising a compendium of 59,089 human-specific regulatory sequences (HSRS) manifest common enrichment patterns of associations with sub-sets of 8,405 neuro-regulatory hsSNCs-linked genes. A. Number of genes identified by the GREAT algorithm as putative regulatory targets of distinct families of HSRS. B. Number of hsSNCs-linked genes among genes comprising putative regulatory targets of distinct families of HSRS. C. Genome-wide correlation patterns between the number of genes comprising the putative regulatory targets of distinct families of HSRS and the number of hsSNCs-linked genes among the HSRS-target genes.

To determine whether the patterns of significant phenotypic associations observed for genes linked with HSRS are specific and not related to the size effects of relatively large gene sets subjected to the GSEA, 42,847 human genes not linked by the GREAT algorithm with HSRS were randomly split into 21 control gene sets of various sizes ranging from 2,847 to 6,847 genes and subjected to the GSEA (Supplemental Notes 1 & 2). Importantly, no significant phenotypic associations were observed for 21 control gene sets, consistent with the conclusion that significant phenotypic associations documented for genes linked with HSRS and neuro-regulatory hsSNCs are not likely due to non-specific size effects captured by the GSEA.

In agreement with this conclusion, it was observed that the common gene set of putative regulatory targets shared by HSRS and neuro-regulatory hsSNCs comprises of 7,406 coding genes (88% of all human-specific SNCs-associated genes), indicating that structurally and functionally diverse HSRS, the evolutionary origin of which has been driven by mechanistically-distinct processes, appear to favor the genomic regulatory alignment with the relatively restricted elite set of genetic targets (Figure 3; Table 3; Supplemental Figure S9; Supplemental Table Set S3). The estimated timeline of the evolutionary origin of 59,089 HSRS is likely to encompass many thousands, perhaps, hundred thousand years. Therefore, it seems remarkable that the patterns of their genomic placement appear uniformly associated with sub-sets of genes comprising the putative regulatory targets of human-specific neuro-regulatory SNCs (Figure 3; Table 3; Supplemental Table Set S3).

Previous studies have identified stem cell-associated retroviral sequences (SCARS) encoded by human endogenous retroviruses LTR7/HERVH and LTR5_Hs/HERVK as one of the significant sources of the evolutionary origin of HSRS (Glinsky, 2015-2019), including human-specific transcription factor binding sites (TFBS) for NANOG, OCT4, and CTCF (Glinsky, 2015).

Next, the common sets of genetic regulatory targets were identified for genes expression of which is regulated by SCARS and genes associated in this study with human-specific regulatory SNCs (Supplemental Figure S9). It has been determined that each of the structurally-distinct families of SCARS appears to share a common set of genetic regulatory targets with human-specific SNCs (Supplemental Figure S8). Overall, expression of nearly two thirds (5,389 genes; 64%) of all genes identified as putative regulatory targets of human-specific SNCs is regulated by SCARS (Supplemental Figure S9; Supplemental Table Set S3). Consistent with the idea that structurally-diverse HSRS may favor the relatively restricted elite set of genetic targets, the common gene set of regulatory targets for HSRS, SCARS, and SNCs comprises of 7,990 coding genes or 95% of all genes associated in this contribution with human-specific neuro-regulatory SNCs (Supplemental Figure S9; Supplemental Table Set S3).

To gain insights into mechanisms of SCARS-mediated effects on expression of 5,389 genes linked to human-specific regulatory SNCs, the numbers of genes expression of which was either activated (down-regulated following SCARS silencing) or inhibited (up-regulated following SCARS silencing) by SCARS have been determined. It was observed that SCARS exert the predominantly inhibitory effect on expression of genes associated with human-specific regulatory SNCs, which is exemplified by activated expression of as many as 87% of genes affected by SCARS silencing (Supplemental Figure S9; Supplemental Table Set S3). These findings indicate that when SCARS-associated networks are active during the human preimplantation embryogenesis, they exert a dominant effect on gene expression, whereas when SCARS are silenced during the postimplantation embryonic development and in the adulthood, regulatory impact of human-specific neuro-regulatory SNCs may be prevalent.

### Genes linked with neuro-regulatory hsSNCs represents intrinsic genetic elements of developmentally and physiologically distinct human-specific genomic regulatory networks (GRNs)

Since distinct families of HSRS, including neuro-regulatory hsSNCs, share common sets of genetic targets, it was of interest to determine whether hsSNCs-linked genes are represented among genes previously identified as components of human-specific GRNs operating in developmentally and physiologically distinct human tissues and cells. Importantly, human-specific GRNs selected for these analyses were defined employing vastly different experimental, analytical, and computational approaches that were applied within the broad range of experimental settings (Glinsky, 2019). Specifically, the interrogated human-specific GRNs include the following data sets: i) Great Apes’ whole-genome sequencing-guided human-specific insertions and deletions (Kronenberg et al., 2018); ii) genome-wide analysis of retrotransposon’s transcriptome in postmortem samples of human dorsolateral prefrontal cortex (Guffanti et al., 2018); iii) shRNA-mediated silencing of LTR7/HERVH retrovirus-derived long non-coding RNAs in hESC (Wang et al., 2014); iv) single-cell expression profiling analyses of human preimplantation embryos (Glinsky et al., 2018); v) network of genes associated with regulatory transposable elements (TE) operating in naïve and primed hESC (Theunissen et al., 2016; Pontis et al., 2019); vi) pluripotency-related network of genes manifesting concordant expression changes in human fetal brain and adult neocortex (Glinsky, 2017); vii) network of genes governing human neurogenesis in vivo (Nowakowski et al., 2017); viii) network of genes differentially expressed during human corticogenesis in vitro (van de Leemput et al., 2014). Thus, selected for these analyses human-specific GRNs appear to function in a developmentally and physiologically diverse spectrum of human cells that are biologically and anatomically highly relevant to manifestations of human-specific phenotypes ranging from preimplantation embryos to adult dorsolateral prefrontal cortex (Table 4; Supplemental Table Set S3).

**Table 4.**
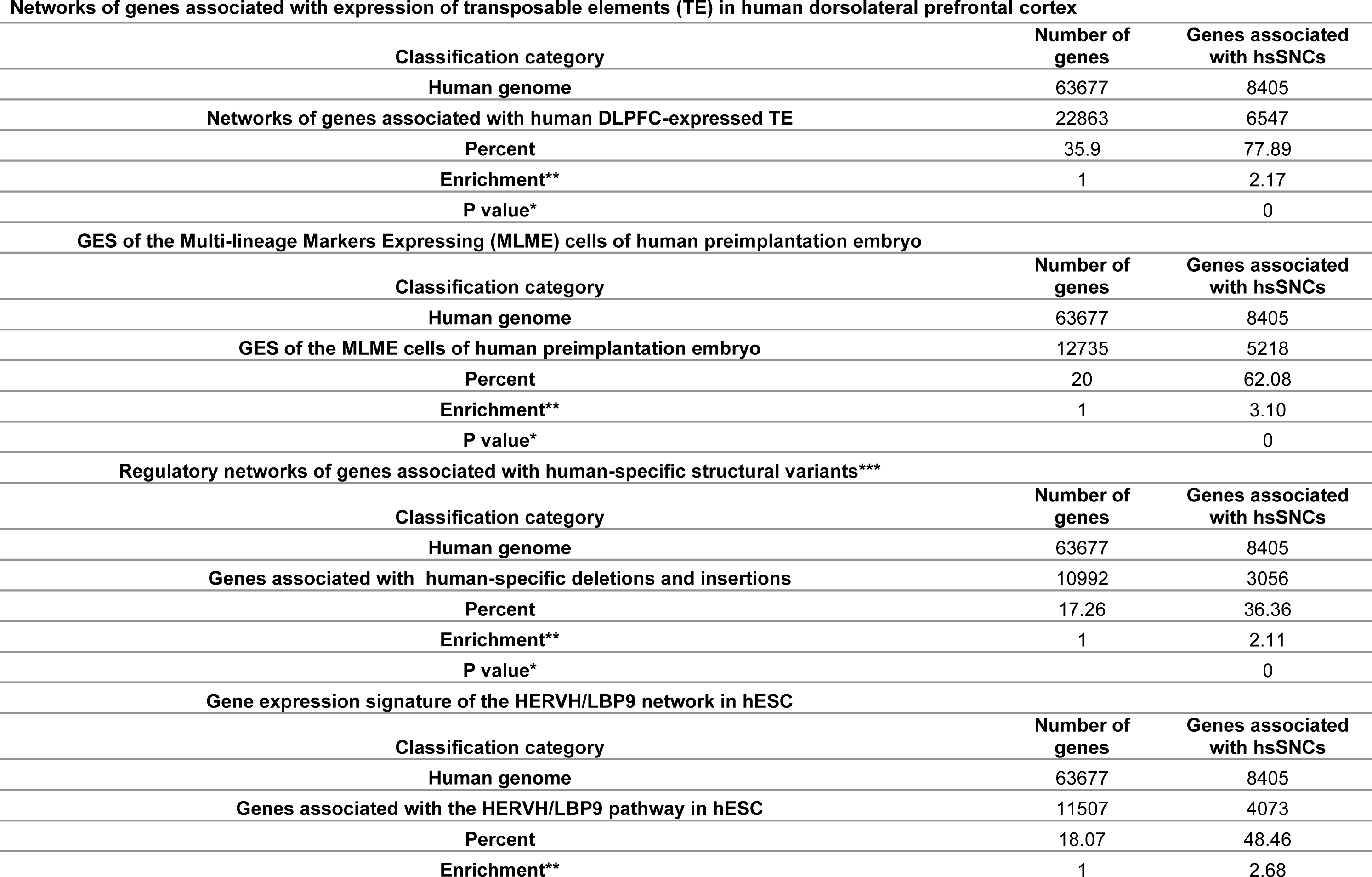
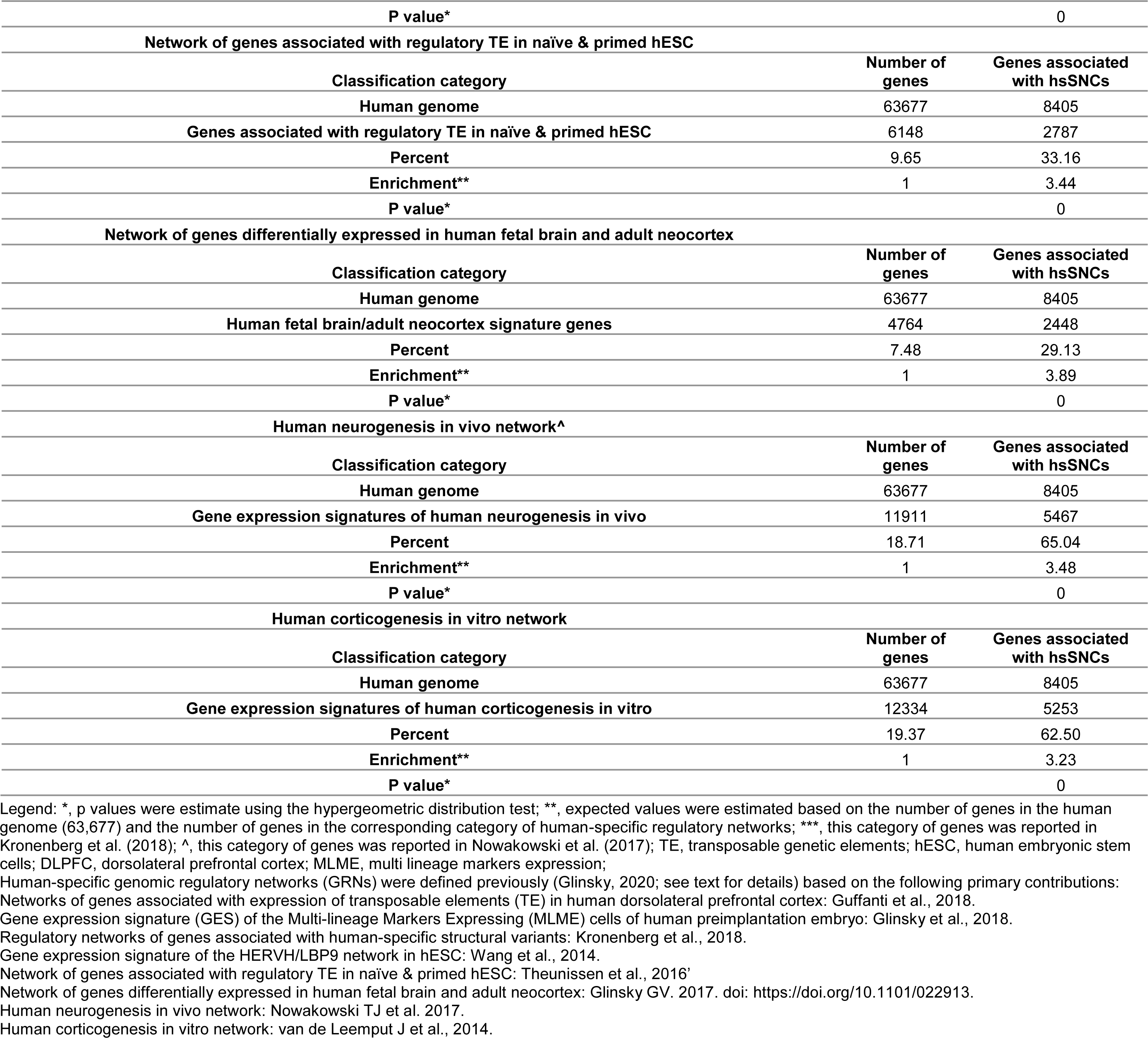
Enrichment within human-specific genomic regulatory networks (GRNs) of 8,405 genes associated with neuro-regulatory hsSNCs

Importantly, in all instances a highly significant enrichment of hsSNCs-linked genes has been observed (Table 4; Supplemental Table Set S3). These observations are consistent with the hypothesis that neuro-regulatory hsSNCs and associated genes represent principal components of the exceptionally broad range of human-specific GRNs operating in the wide spectra of developmental and physiological contexts reflecting species-defining human-specific phenotypes.

## Discussion

In recent years, elucidation of genetic and molecular mechanisms defining the phenotypic uniqueness of Modern Humans attained a significant progress in illuminating the potentially broad role of thousands human-specific regulatory sequences (HSRS) in contrast to the relatively modest impact of human-specific changes of a limited number of coding genes (Kronenberg et al., 2018; Glinsky, 2019; Kanton et al., 2019). The macromolecules comprising the essential building blocks of life at the cellular and organismal levels remain highly conserved during the evolution of humans and other Great Apes. Identification and initial structural-functional characterization of nearly hundred thousand candidate HSRS (Kronenberg et al., 2018; Glinsky, 2019; Kanton et al., 2019; this contribution) validate the idea that unique to human phenotypes may result from human-specific changes to genomic regulatory sequences defined as “regulatory mutations” (King and Wilson, 1975). Technological advances enabled the exquisite degree of accuracy of molecular definition of 35,074 SNCs that are fixed in humans, distinct from other primates, and located in DA chromatin regions during human brain development (Kanton et al., 2019). Notably, 99.8% of candidate regulatory hsSNCs that overlap DA chromatin regions during brain development are shared with the archaic humans while only 64 hsSNCs are unique to Modern Humans. The conservation on the human lineage of a vast majority of regulatory hsSNCs associated with early stages of human brain development suggest that coding genes expression of which is regulated by hsSNCs may have a broad effect on human-specific traits beyond embryonic development. This concept has been substantiated by the multiple lines of evidence acquired and reported in the present contribution.

Employing the GREAT algorithm (McLean et al., 2010, 2011), 8,405 genes have been identified that are linked to 35,074 hsSNCs via genomic proximity co-localization analysis, indicating that expression of these hsSNCs- linked genes might be affected by hsSNCs located in DA chromatin regions during brain development. Comprehensive gene set enrichment analyses (GSEA) of these 8,405 genes revealed the staggering breadth of associations with physiological processes, morphological features, and pathological conditions of *H. sapiens*. Significantly enriched records include more than 1,000 anatomically-distinct regions of the adult human brain, many human tissues and cell types, more than 200 common human disorders and more than 1,000 rare diseases. Strikingly similar patterns of phenotypic associations have been observed for genes linked to 59,089 previously defined structurally, functionally, and evolutionary distinct classes of HSRS.

Based on the reported above observations, it has been concluded that genes linked to neuro-regulatory hsSNCs appear contributing to development, morphological architecture, and biological functions of the adult human brain, other components of the central nervous system, and many tissues and organs across human body. They were implicated in the extensive range of human physiological and pathological conditions, thus supporting the hypothesis that phenotype-altering effects of neuro-regulatory hsSNCs are not restricted to the early-stages of human brain development. Results of the analyses utilizing the Mouse Genome Informatics (MGI) database (http://www.informatics.jax.org/) revealed that neuro-regulatory hsSNCs-associated genes affect wide spectra of traits defining both physiology and pathology of Modern Humans, perhaps, reflecting the global scale of human-specific regulatory impacts on thousands essential mammalian phenotypes.

Significantly, outlined herein analytical approaches and reported end-points provide readily available access to mouse models for precise molecular definitions of unique to humans regulatory effects of neuro-regulatory human-specific SNCs and other types of HSRS on genes causally affecting thousands of defined mammalian phenotypes and hundreds of common and rare human disorders.

## Methods

### Data source and analytical protocols

#### Candidate human-specific regulatory sequences and African Apes-specific retroviral insertions

A total of 94,806 candidate HSRS, including 35,074 neuro-regulatory human-specific SNCs, detailed descriptions of which and corresponding references of primary original contributions are reported elsewhere (Glinsky et al., 2015-2019; Kanton et al., 2019). Solely publicly available datasets and resources were used in this contribution. The significance of the differences in the expected and observed numbers of events was calculated using two-tailed Fisher’s exact test. Additional placement enrichment tests were performed for individual classes of HSRS taking into account the size in bp of corresponding genomic regions.

## Data analysis

### Categories of DNA sequence conservation

Identification of highly-conserved in primates (pan-primate), primate-specific, and human-specific sequences was performed as previously described (Glinsky, 2015-2019). In brief, all categories were defined by direct and reciprocal mapping using LiftOver. Specifically, the following categories of candidate regulatory sequences were distinguished:

- <u>Highly conserved in primates’ sequences</u>: DNA sequences that have at least 95% of bases remapped during conversion from/to human (Homo sapiens, hg38), chimp (Pan troglodytes, v5), and bonobo (Pan paniscus, v2; in specifically designated instances, Pan paniscus, v1 was utilized for comparisons). Similarly, highly-conserved sequences were defined for hg38 and latest releases of genomes of Gorilla, Orangutan, Gibbon, and Rhesus.
- <u>Primate-specific</u>: DNA sequences that failed to map to the mouse genome (mm10).
- <u>Human-specific</u>: DNA sequences that failed to map at least 10% of bases from human to both chimpanzee and bonobo. All candidate HSRS identified based on the sequence alignments failures to genomes of both chimpanzee and bonobo were subjected to more stringent additional analyses requiring the mapping failures to genomes of Gorilla, Orangutan, Gibbon, and Rhesus. These loci were considered created *de novo* human-specific regulatory sequences (HSRS).

To infer the putative evolutionary origins, each evolutionary classification was defined independently by running the corresponding analyses on all candidate HSRS representing the specific category. For example, human-rodent conversion identify sequences that are absent in the mouse genome based on the sequence identity threshold of 10%). Additional comparisons were performed using the same methodology and exactly as stated in the manuscript text. Human brain regions’ marker genes were identified among genes linked to hsSNCs by analyzing genes significantly up-regulated in specified human brain regions using the Allen Brain Atlas database (brain region-specific records manifesting significantly increased expression at 1.5-fold cut-off were selected for analyses). Genes differentially expressed in human versus chimpanzee adult brains were identified among hsSNCs-linked genes by analyzing genes differentially expressed in eight regions of human versus chimpanzee adult brains (Xu et al., 2018).

### Gene set enrichment and genome-wide proximity placement analyses

Gene set enrichment analyses were carried-out using the Enrichr bioinformatics platform, which enables the interrogation of nearly 200,000 gene sets from more than 100 gene set libraries. The Enrichr API (January 2018 through January 2020 releases) (Chen et al., 2013; Kuleshov et al., 2016) was used to test genes linked to HSRS of interest for significant enrichment in numerous functional categories. In all tables and plots (unless stated otherwise), in addition to the nominal p values and adjusted p values, the “combined score” calculated by Enrichr is reported, which is a product of the significance estimate and the magnitude of enrichment (combined score *c = log(p) * z*, where *p* is the Fisher’s exact test p-value and *z* is the z-score deviation from the expected rank). When technically feasible, larger sets of genes comprising several thousand entries were analyzed. Regulatory connectivity maps between HSRS and coding genes and additional functional enrichment analyses were performed with the GREAT algorithm (McLean et al., 2010; 2011) at default settings. The reproducibility of the results was validated by implementing two releases of the GREAT algorithm: GREAT version 3.0.0 (2/15/2015 to 08/18/2019) and GREAT version 4.0.4 (08/19/2019). Genome-wide Proximity Placement Analysis (GPPA) of distinct genomic features co-localizing with HSRS was carried out as described previously and originally implemented for human-specific transcription factor binding sites (Glinsky, 2015-2019).

### Mammalian Phenotype Ontology and Human Disease Ontology analyses

To validate and extend findings afforded by the gene set enrichment analyses and to identify all genes linked with human-specific regulatory SNCs that are associated with defined mammalian phenotypes as well as implicated in development of human diseases with one or more mouse models, the additional analyses have been carried out utilizing the Mouse Genome Informatics (MGI) database (http://www.informatics.jax.org/).

#### Statistical Analyses of the Publicly Available Datasets

All statistical analyses of the publicly available genomic datasets, including error rate estimates, background and technical noise measurements and filtering, feature peak calling, feature selection, assignments of genomic coordinates to the corresponding builds of the reference human genome, and data visualization, were performed exactly as reported in the original publications and associated references linked to the corresponding data visualization tracks (http://genome.ucsc.edu/). Any modifications or new elements of statistical analyses are described in the corresponding sections of the Results. Statistical significance of the Pearson correlation coefficients was determined using GraphPad Prism version 6.00 software. Both nominal and Bonferroni adjusted p values were estimated. The significance of the differences in the numbers of events between the groups was calculated using two-sided Fisher’s exact and Chi-square test, and the significance of the overlap between the events was determined using the hypergeometric distribution test (Tavazoie et al., 1999).

## Supplemental Information

Supplemental information includes Supplemental Table Sets S1 – S5, Supplemental Text, Supplemental Notes, and Supplemental Figures S1-S9.

## Author Contributions

This is a single author contribution. All elements of this work, including the conception of ideas, formulation, and development of concepts, execution of experiments, analysis of data, and writing of the paper, were performed by the author.

## Acknowledgements

This work was made possible by the open public access policies of major grant funding agencies and international genomic databases and the willingness of many investigators worldwide to share their primary research data. This work was supported, in part, by OncoScar, Inc.

