## Supplemental Figure for "Impacts of genomic networks governed by human-specific regulatory sequences and genetic loci harboring fixed human-specific neuro-regulatory single nucleotide mutations on phenotypic traits of Modern Humans"

Gennadi V. Glinsky[1]

[1] Institute of Engineering in Medicine

University of California, San Diego

9500 Gilman Dr. MC 0435

La Jolla, CA 92093-0435, USA

Web: http://iem.ucsd.edu/people/profiles/guennadi-v-glinskii.html

**Running title:** Global impact of fixed human-specific neuro-regulatory mutations in Modern Humans

**Key words:** human phenotypic uniqueness; human-specific regulatory sequences; stem cell-associated retroviral sequences; human-specific traits; fixed neuro-regulatory human-specific single nucleotide mutations.

### SUPPLEMENTAL FIGURES

ARCHS4 Human Tissues: 8,045 genes

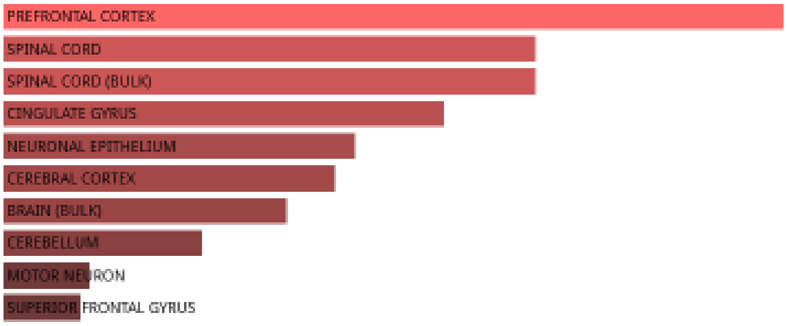


ARCHS4 Human Tissues: 8,045 genes
Top 10 of 39 significant records

| Term | Overlap | P-value | Adjusted P-value |
| --- | --- | --- | --- |
| PREFRONTAL CORTEX | 1348/2316 | 2.06686E-62 | 2.23E-60 |
| SPINAL CORD | 1299/2316 | 1.17222E-47 | 4.22E-46 |
| SPINAL CORD (BULK) | 1299/2316 | 1.17222E-47 | 4.22E-46 |
| CINGULATE GYRUS | 1279/2316 | 3.13703E-42 | 8.47E-41 |
| NEURONAL EPITHELIUM | 1258/2316 | 6.71385E-37 | 1.45E-35 |
| CEREBRAL CORTEX | 1253/2316 | 1.09815E-35 | 1.98E-34 |
| BRAIN (BULK) | 1241/2316 | 7.35673E-33 | 1.14E-31 |
| CEREBELLUM | 1218/2316 | 8.73598E-28 | 1.18E-26 |
| MOTOR NEURON | 1184/2316 | 4.19448E-21 | 5.03E-20 |
| SUPERIOR FRONTAL GYRUS | 1181/2316 | 1.4634E-20 | 1.58E-19 |

Allen Brain Atlas Up: 8,045 genes

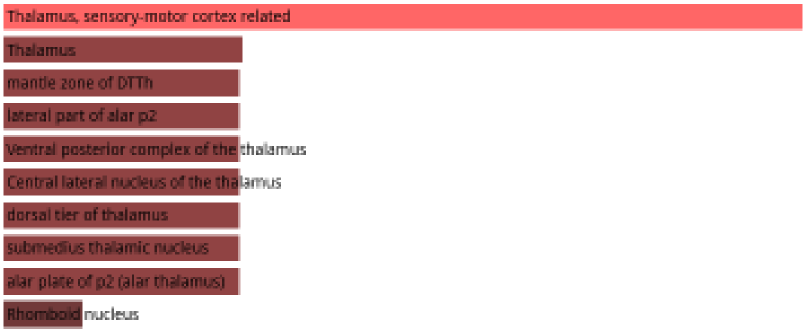


Allen Brain Atlas Up: 8,045 genes
Networks

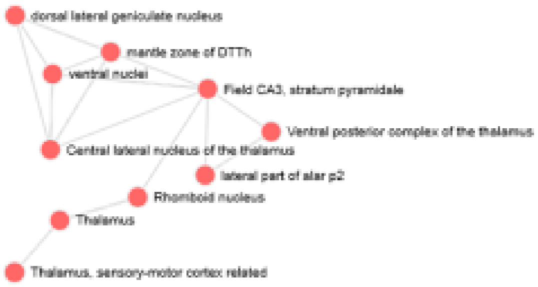

### Allen Brain Atlas Up: 8,045 genes

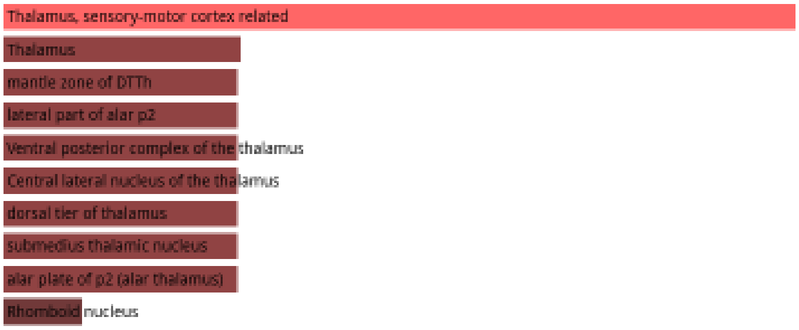


### Allen Brain Atlas Up: 8,045 genes
### Networks

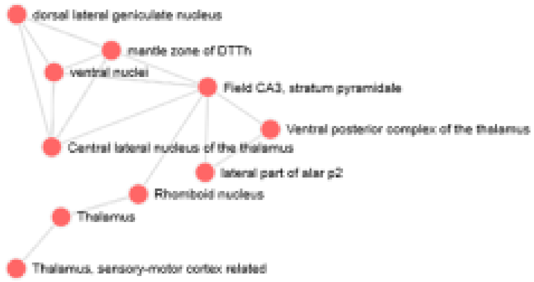


### Allen Brain Atlas Up: 8,045 genes
### Top 30 of 1,200 significant records

| Term | Overlap | P-value | Adjusted P-value |
|---|---|---|---|
| Thalamus, sensory-motor cortex related | 207/301 | 3.57E-21 | 7.82E-18 |
| Thalamus | 222/334 | 9.71E-20 | 2.42E-17 |
| mantle zone of DTTh | 204/301 | 9.94E-20 | 2.42E-17 |
| lateral part of alar p2 | 204/301 | 9.94E-20 | 2.42E-17 |
| Ventral posterior complex of the thalamus | 204/301 | 9.94E-20 | 2.42E-17 |
| Central lateral nucleus of the thalamus | 204/301 | 9.94E-20 | 2.42E-17 |
| alar plate of p2 (alar thalamus) | 204/301 | 9.94E-20 | 2.42E-17 |
| submedius thalamic nucleus | 204/301 | 9.94E-20 | 2.42E-17 |
| dorsal tier of thalamus | 204/301 | 9.94E-20 | 2.42E-17 |
| Rhomboid nucleus | 227/345 | 2.51E-19 | 5.34E-17 |
| ventral nuclei | 203/301 | 2.92E-19 | 5.34E-17 |
| Ventral group of the dorsal thalamus | 203/301 | 2.92E-19 | 5.34E-17 |
| Field CA3, stratum pyramidale | 202/301 | 8.47E-19 | 1.24E-16 |
| Ventral posteromedial nucleus of the thalamus | 202/301 | 8.47E-19 | 1.24E-16 |
| ventral posteromedial nucleus | 202/301 | 8.47E-19 | 1.24E-16 |
| intermediate stratum of DTTh | 201/301 | 2.42E-18 | 2.94E-16 |
| prosomere 2 | 201/301 | 2.42E-18 | 2.94E-16 |
| central lateral nucleus | 201/301 | 2.42E-18 | 2.94E-16 |
| dorsal lateral geniculate nucleus | 200/301 | 6.79E-18 | 7.84E-16 |
| hilus of the DG | 199/301 | 1.88E-17 | 2.06E-15 |
| lateral (parvicellular) part of MD | 198/301 | 5.13E-17 | 4.33E-15 |
| Dorsal part of the lateral geniculate complex | 198/301 | 5.13E-17 | 4.33E-15 |
| Ventral posterolateral nucleus of the thalamus | 198/301 | 5.13E-17 | 4.33E-15 |
| posterior (ventral) nucleus | 198/301 | 5.13E-17 | 4.33E-15 |
| Hippocampal formation | 198/301 | 5.13E-17 | 4.33E-15 |
| intralaminar nuclei | 198/301 | 5.13E-17 | 4.33E-15 |
| intermediate stratum of DG | 353/597 | 9.09E-17 | 7.38E-15 |
| Field CA3, stratum radiatum | 219/342 | 1.17E-16 | 9.12E-15 |
| mantle zone of CA | 197/301 | 1.38E-16 | 9.76E-15 |
| Thalamus, polymodal association cortex related | 197/301 | 1.38E-16 | 9.76E-15 |

Allen Brain Atlas Down: 8,045 genes

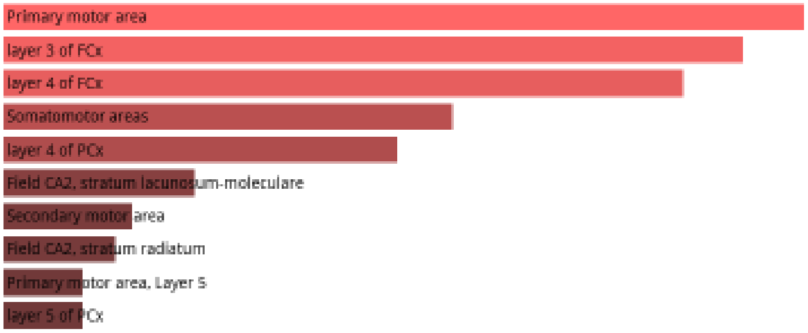


Allen Brain Atlas Down: 8,045 genes
Networks

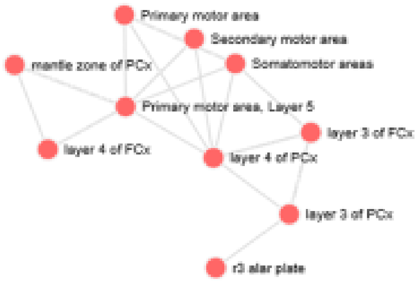


Allen Brain Atlas Down: 8,045 genes

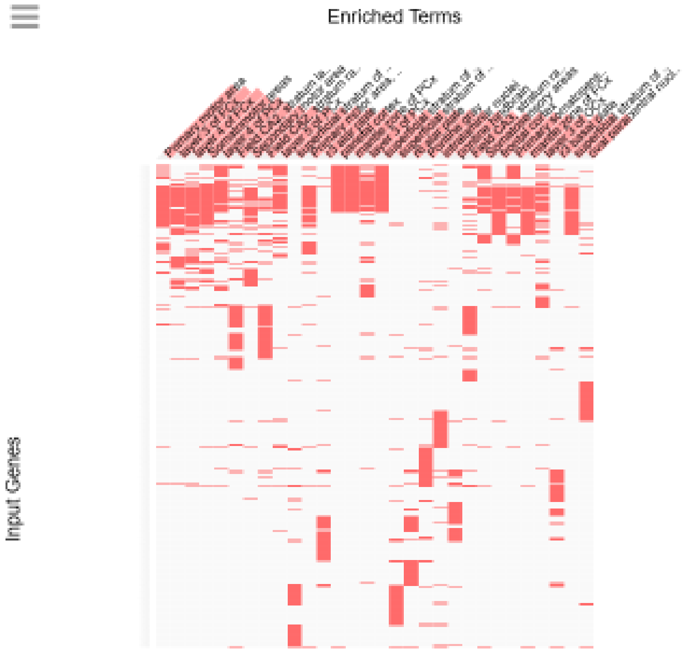

### Allen Brain Atlas Down: 8,045 genes
### Top 31 of 1,062 significant records

| Term | Overlap | P-value | Adjusted P-value |
|---|---|---|---|
| Primary motor area | 195/300 | 5.73E-16 | 1.26E-12 |
| layer 3 of FCx | 194/300 | 1.49E-15 | 1.63E-12 |
| layer 4 of FCx | 193/300 | 3.81E-15 | 2.79E-12 |
| Somatomotor areas | 189/300 | 1.42E-13 | 7.77E-11 |
| layer 4 of PCx | 188/300 | 3.38E-13 | 1.48E-10 |
| Field CA2, stratum lacunosum-moleculare | 248/426 | 8.01E-12 | 2.93E-09 |
| Secondary motor area | 183/300 | 2.1E-11 | 6.57E-09 |
| Field CA2, stratum radiatum | 262/458 | 2.75E-11 | 7.54E-09 |
| Primary motor area, Layer 5 | 182/300 | 4.59E-11 | 9.16E-09 |
| layer 5 of PCx | 182/300 | 4.59E-11 | 9.16E-09 |
| superficial stratum of m1B | 182/300 | 4.59E-11 | 9.16E-09 |
| r3 alar plate | 181/300 | 9.92E-11 | 1.67E-08 |
| parietal cortex | 181/300 | 9.92E-11 | 1.67E-08 |
| mantle zone of PCx | 180/300 | 2.11E-10 | 2.44E-08 |
| layer 3 of PCx | 180/300 | 2.11E-10 | 2.44E-08 |
| Pontine gray | 180/300 | 2.11E-10 | 2.44E-08 |
| r6 alar plate | 183/300 | 2.11E-10 | 2.44E-08 |
| superficial stratum of PCx (cortical plate/marginal zone) | 180/300 | 2.11E-10 | 2.44E-08 |
| superficial stratum of p2B | 180/300 | 2.11E-10 | 2.44E-08 |
| intralaminar nuclei | 179/300 | 4.44E-10 | 4.64E-08 |
| pontine hindbrain | 182/300 | 4.44E-10 | 4.64E-08 |
| Field CA3, stratum radiatum | 211/364 | 4.83E-10 | 4.81E-08 |
| Primary somatosensory area | 178/300 | 9.2E-10 | 8.07E-08 |
| frontal cortex | 179/300 | 9.2E-10 | 8.07E-08 |
| Somatosensory areas | 178/300 | 9.2E-10 | 8.07E-08 |
| mantle zone of FCx | 178/300 | 1.88E-09 | 1.37E-07 |
| layer 3 of OCx | 177/300 | 1.88E-09 | 1.37E-07 |
| r3 basal plate | 177/300 | 1.88E-09 | 1.37E-07 |
| superficial stratum of FCx (cortical plate/marginal zone) | 178/300 | 1.88E-09 | 1.37E-07 |
| oval paracentral nucleus | 177/300 | 1.88E-09 | 1.37E-07 |
| Primary motor area, Layer 2/3 | 176/300 | 3.79E-09 | 2.6E-07 |

**Supplemental Figure S1.** Identification of genes expression of which distinguishes thousands of anatomically distinct areas of the adult human brain, various regions of the central nervous system, and many different cell types and tissues in the human body.

### Aging Perturbations from GEO up: 8,405 genes

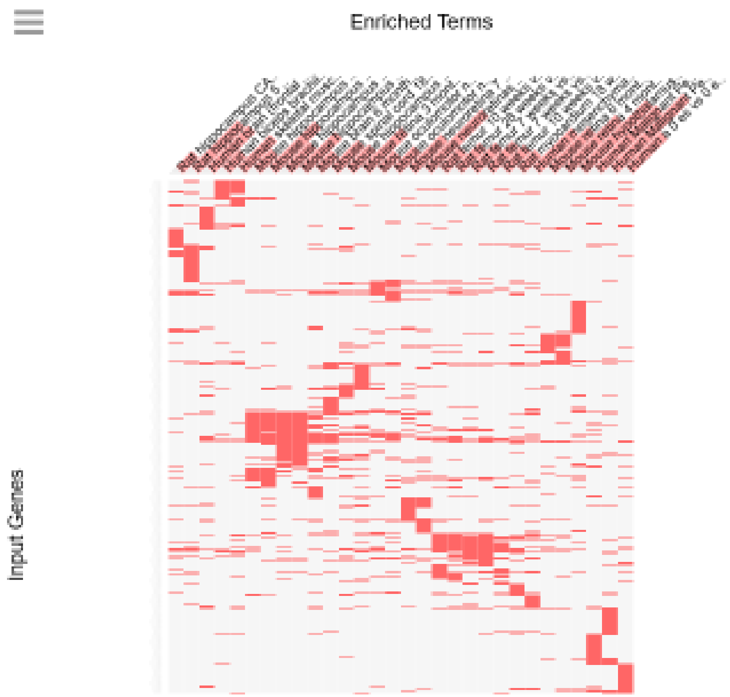


### Aging Perturbations from GEO up: 8,405 genes
### Top 30 significant records

| Term | Overlap | P-value | Adjusted P-value |
|---|---|---|---|
| Human_Malignant Peripheral Nerve Sheath Tumour_24 years vs 53 years_GSE17118_aging:364 | 197/337 | 6.9E-10 | 1.97E-07 |
| Mouse_hypothalamus_42 days vs 182 days_GDS3895_aging:107 | 201/349 | 2.59E-09 | 3.7E-07 |
| Human_Malignant Peripheral Nerve Sheath Tumour_27 years vs 61 years_GSE17118_aging:363 | 195/343 | 1.78E-08 | 1.5E-06 |
| Mouse_neuroretinas_7 weeks vs 64 weeks_GSE38671_aging:211 | 168/289 | 2.1E-08 | 1.5E-06 |
| Mouse_retina_4 months vs 10 months_GSE33674_aging:304 | 148/254 | 1.16E-07 | 6.63E-06 |
| Rat_hippocampus_5 months vs 24 months_GSE14505_aging:346 | 230/429 | 6.82E-07 | 3.25E-05 |
| Mouse_hippocampus_9 months vs 20 months_GSE48911_aging:391 | 243/459 | 1.23E-06 | 5.02E-05 |
| Mouse_retina_6 months vs 10 months_GSE33674_aging:305 | 125/218 | 3.39E-06 | 0.000121 |
| Human_brain (frontal cortex)_55 years vs 82 years_GSE53890_aging:229 | 171/313 | 4.05E-06 | 0.000129 |
| Mouse_retina_3 months vs 16 months_GDS2654_aging:66 | 199/372 | 4.53E-06 | 0.00013 |
| Human_mesenchymal stem cells (from bone marrow)_42 years vs 79 years_GSE35955_aging:293 | 133/238 | 1.03E-05 | 0.000255 |
| Human_a_0 es vs 0 es_GDS5077_aging:106 | 172/319 | 1.07E-05 | 0.000255 |
| Mouse_hippocampus_9 months vs 14 months_GSE48911_aging:390 | 254/494 | 1.28E-05 | 0.000283 |
| Mouse_neuroretina_7 weeks vs 64 weeks_GSE38671_aging:210 | 171/319 | 1.76E-05 | 0.00036 |
| Human_skeletal muscle_19 years vs 65 years_GDS4858_aging:10 | 140/259 | 5.77E-05 | 0.0011 |
| Mouse_spinal cord_18 months vs 30 months_GDS1280_aging:1 | 172/331 | 0.000151 | 0.002697 |
| Mouse_hippocampus_9 months vs 14 months_GSE48911_aging:384 | 246/494 | 0.000252 | 0.004247 |
| Rat_femur_7 weeks vs 53 weeks_GDS509_aging:264 | 197/389 | 0.000331 | 0.005257 |
| Rat_femur_28 weeks vs 54 weeks_GDS509_aging:271 | 193/383 | 0.000523 | 0.007575 |
| Rat_femur_7 weeks vs 27 weeks_GDS509_aging:258 | 155/301 | 0.00053 | 0.007575 |
| Mouse_hippocampus_2 months vs 15 months_GSE5078_aging:398 | 151/293 | 0.000593 | 0.008072 |
| Rat_femur_10 weeks vs 30 weeks_GDS509_aging:260 | 123/236 | 0.001057 | 0.013739 |
| Mouse_oculomotor nucleus_6 months vs 30 months_GDS1280_aging:6 | 140/273 | 0.001185 | 0.01473 |
| Rat_femur_10 weeks vs 56 weeks_GDS509_aging:266 | 191/384 | 0.001248 | 0.014873 |
| Mouse_hippocampus_9 months vs 20 months_GSE48911_aging:385 | 216/442 | 0.001959 | 0.022408 |
| Human_biceps brachii muscles_24 years vs 70 years_GDS4858_aging:33 | 119/233 | 0.003152 | 0.034649 |
| Rat_myocardium_18 weeks vs 22 weeks_GDS4025_aging:142 | 124/244 | 0.003271 | 0.034649 |
| Rat_hippocampus CA3 region_6 months vs 25 months_GSE14724_aging:347 | 170/345 | 0.003642 | 0.036658 |
| Mouse_brain_6 months vs 14 months_GSE15129_aging:313 | 189/387 | 0.003717 | 0.036658 |
| Mouse_spinal cord_6 months vs 30 months_GDS1280_aging:3 | 197/405 | 0.003882 | 0.037008 |

Aging Perturbations from GEO down: 8,405 genes

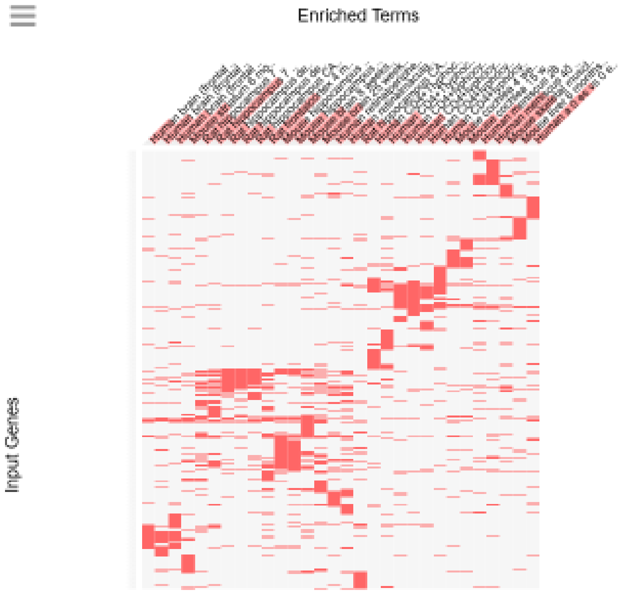


Aging Perturbations from GEO down: 8,405 genes
Top 30 significant records

| Term | Overlap | P-value | Adjusted P-value |
|---|---|---|---|
| Rat_hippocampus_7 months vs 21 months_GDS4019_aging:98 | 196/314 | 1.85E-13 | 5.28E-11 |
| Human_a_0 es vs 0 es_GDS5077_aging:106 | 175/281 | 4.78E-12 | 6.83E-10 |
| Human_meniscus_40 years vs 55 years_GSE45233_aging:209 | 189/320 | 4.76E-10 | 4.54E-08 |
| Rat_frontal cortex_4 months vs 22 months_GDS3939_aging:99 | 192/327 | 6.86E-10 | 4.9E-08 |
| Mouse_skeletal muscle precursor_12 months vs 24 months_GDS4892_aging:9 | 137/224 | 5.29E-09 | 3.02E-07 |
| Mouse_brain_25 weeks vs 100 weeks_GSE41018_aging:400 | 178/311 | 3.83E-08 | 1.68E-06 |
| Mouse_striatum_6 months vs 21 months_GDS4153_aging:78 | 164/283 | 4.12E-08 | 1.68E-06 |
| Human_meniscus_28 years vs 47 years_GSE45233_aging:208 | 165/286 | 5.7E-08 | 2.04E-06 |
| Mouse_brain_8 weeks vs 104 weeks_GSE20411_aging:311 | 224/408 | 8.36E-08 | 2.66E-06 |
| Human_brain (frontal cortex)_28 years vs 100 years_GSE53890_aging:228 | 197/356 | 2.32E-07 | 6.64E-06 |
| Human_CD4+Tlyphocytes_61 years vs 81 years_GSE62373_aging:185 | 193/349 | 3.26E-07 | 8.48E-06 |
| Human_CD4+lyphocytes_40 years vs 72 years_GSE62373_aging:163 | 203/373 | 7.65E-07 | 1.82E-05 |
| Mouse_hippocampus_4 months vs 9 months_GSE48911_aging:387 | 230/430 | 8.6E-07 | 1.89E-05 |
| Rat_hippocampus_7 months vs 21 months_GDS4019_aging:97 | 156/277 | 9.99E-07 | 2.04E-05 |
| Mouse_cochlea_15 weeks vs 45 weeks_GSE35234_aging:164 | 142/250 | 1.58E-06 | 3.01E-05 |
| Human_brain (frontal cortex)_55 years vs 82 years_GSE53890_aging:229 | 159/287 | 2.97E-06 | 5.31E-05 |
| Human_retinalmacula_18 years vs 74 years_GSE32614_aging:150 | 182/336 | 4.14E-06 | 6.96E-05 |
| Mouse_hippocampus_4 months vs 9 months_GSE48911_aging:381 | 226/429 | 4.52E-06 | 7.18E-05 |
| Human_CD4+lyphocytes_40 years vs 61 years_GSE62373_aging:160 | 192/358 | 5.34E-06 | 7.77E-05 |
| Human_CD4+Tlyphocytes_29 years vs 81 years_GSE62373_aging:176 | 201/377 | 5.44E-06 | 7.77E-05 |
| Human_CD4+lyphocytes_29 years vs 61 years_GSE62373_aging:154 | 184/342 | 6.43E-06 | 8.75E-05 |
| Human_retinalperiphery_32 years vs 74 years_GSE32614_aging:151 | 160/293 | 8.42E-06 | 0.000109 |
| Human_brain (frontal cortex)_28 years vs 82 years_GSE53890_aging:227 | 154/281 | 9.47E-06 | 0.000118 |
| Rat_hippocampus CA3 region_6 months vs 25 months_GSE14724_aging:347 | 141/255 | 1.2E-05 | 0.000143 |
| Mouse_cochlea_4 weeks vs 45 weeks_GSE35234_aging:161 | 145/268 | 4.04E-05 | 0.000463 |
| Rat_hippocampus dentate gyrus_18 months vs 28 months_GSE21681_aging:357 | 124/225 | 4.78E-05 | 0.000522 |
| Rat_hippocampus dentate gyrus_18 months vs 28 months_GSE21681_aging:358 | 147/273 | 4.93E-05 | 0.000522 |
| Human_CD4+lyphocytes_40 years vs 81 years_GSE62373_aging:165 | 197/380 | 6.24E-05 | 0.000638 |
| Human_CD4+lyphocytes_29 years vs 81 years_GSE62373_aging:156 | 215/419 | 6.73E-05 | 0.000664 |
| Rat_hippocampus CA1_18 months vs 28 months_GSE21681_aging:353 | 144/270 | 0.000107 | 0.001004 |

**Supplemental Figure S2.** Identification and characterization of genes expression of which is altered during aging of humans, rats, and mice.

Human Phenotype Ontology: 8,405 genes

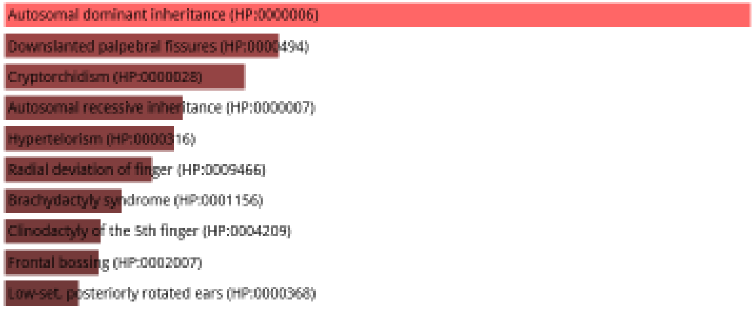


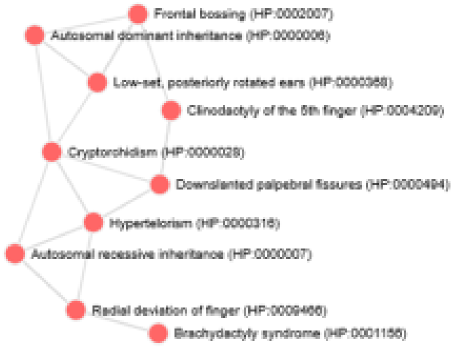


Human Phenotype Ontology: 8,405 genes

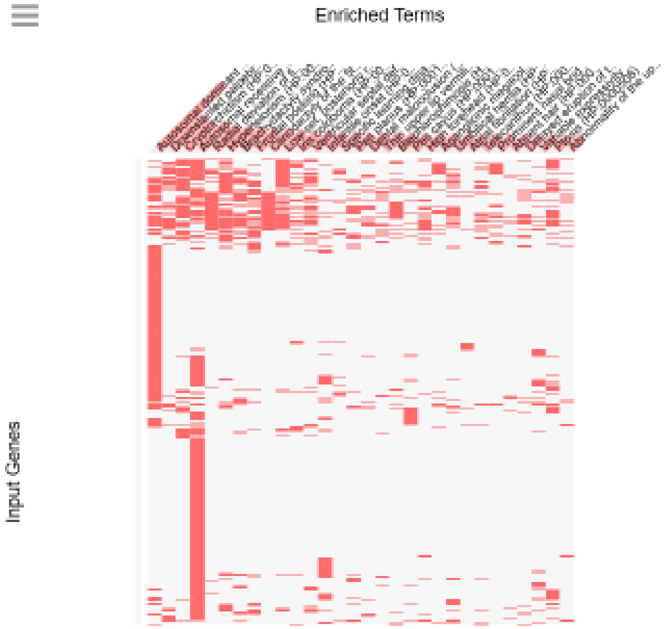

### Human Phenotype Ontology (8,405 genes): Top 35 of 298 significant records

| Term | Overlap | P-value | Adjusted P-value |
|---|---|---|---|
| Autosomal dominant inheritance (HP:0000006) | 645/1134 | 2.85E-25 | 5.07E-22 |
| Downslanted palpebral fissures (HP:0000494) | 127/188 | 1.17E-12 | 1.04E-09 |
| Cryptorchidism (HP:0000028) | 222/379 | 4.46E-11 | 2.64E-08 |
| Autosomal recessive inheritance (HP:0000007) | 849/1722 | 1.14E-10 | 5.07E-08 |
| Radial deviation of finger (HP:0009466) | 140/229 | 3.75E-09 | 1.11E-06 |
| Hypertelorism (HP:0000316) | 190/328 | 3.76E-09 | 1.11E-06 |
| Brachydactyly syndrome (HP:0001156) | 119/192 | 1.77E-08 | 4.5E-06 |
| Frontal bossing (HP:0002007) | 129/213 | 3.38E-08 | 7.51E-06 |
| Clinodactyly of the 5th finger (HP:0004209) | 118/192 | 4.03E-08 | 7.96E-06 |
| Low-set, posteriorly rotated ears (HP:0000368) | 158/276 | 2.14E-07 | 3.8E-05 |
| Iris coloboma (HP:0000612) | 72/110 | 5.77E-07 | 9.34E-05 |
| Ventricular septal defect (HP:0001629) | 106/176 | 7.99E-07 | 0.000113 |
| Infantile onset (HP:0003593) | 145/254 | 8.28E-07 | 0.000113 |
| Specific learning disability (HP:0001328) | 36/47 | 1.49E-06 | 0.000189 |
| Pes planus (HP:0001763) | 68/105 | 2.09E-06 | 0.000246 |
| Dental malocclusion (HP:0000689) | 55/81 | 2.21E-06 | 0.000246 |
| Thin upper lip vermilion (HP:0000219) | 38/51 | 2.5E-06 | 0.000261 |
| Blepharophimosis (HP:0000581) | 67/104 | 3.26E-06 | 0.000323 |
| Pes cavus (HP:0001761) | 80/129 | 3.55E-06 | 0.000323 |
| High forehead (HP:0000348) | 69/108 | 3.63E-06 | 0.000323 |
| Aganglionic megacolon (HP:0002251) | 63/97 | 4.26E-06 | 0.000361 |
| Umbilical hernia (HP:0001537) | 91/151 | 4.5E-06 | 0.000364 |
| Atrial fibrillation (HP:0005110) | 26/32 | 6.52E-06 | 0.000505 |
| Telecanthus (HP:0000506) | 55/83 | 6.96E-06 | 0.000516 |
| Prominent nasal bridge (HP:0000426) | 64/100 | 7.43E-06 | 0.000529 |
| Absent hair (HP:0002298) | 18/20 | 1.14E-05 | 0.000781 |
| Delayed eruption of teeth (HP:0000684) | 56/86 | 1.28E-05 | 0.000841 |
| Variable expressivity (HP:0003828) | 86/144 | 1.34E-05 | 0.00085 |
| Ptosis (HP:0000508) | 180/338 | 1.79E-05 | 0.001074 |
| Abnormality of the upper arm (HP:0001454) | 29/38 | 1.81E-05 | 0.001074 |
| Progressive disorder (HP:0003676) | 86/145 | 1.94E-05 | 0.001103 |
| Depressed nasal bridge (HP:0005280) | 127/228 | 1.99E-05 | 0.001103 |
| Sporadic (HP:0003745) | 42/61 | 2.05E-05 | 0.001103 |
| Left ventricular hypertrophy (HP:0001712) | 31/42 | 2.93E-05 | 0.0015 |
| Postaxial hand polydactyly (HP:0001162) | 53/82 | 2.95E-05 | 0.0015 |

### Database of Human Genotypes and Phenotypes (dbGaP): 8,405 genes

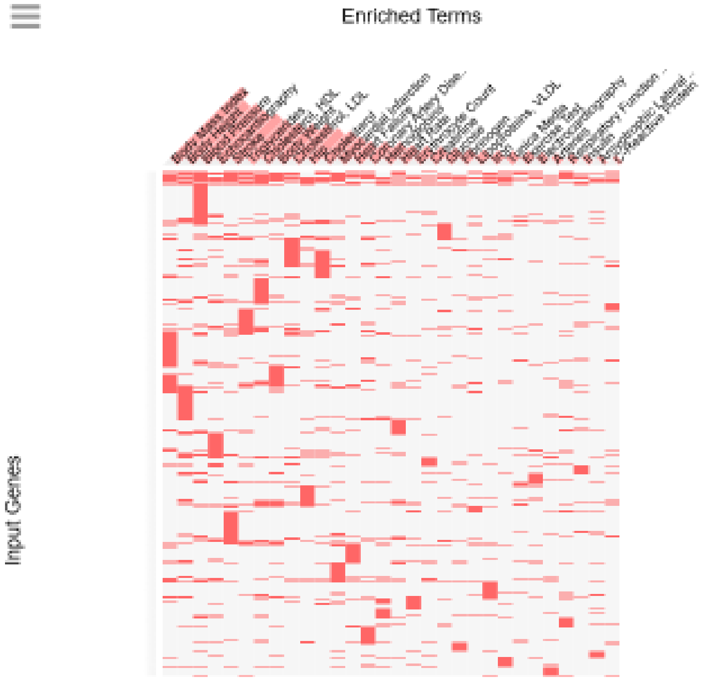

### Database of Human Genotypes and Phenotypes (dbGaP): 8,405 genes
### Top 20 of 136 significant records

| Term | Overlap | P-value | Adjusted P-value |
|---|---|---|---|
| Body Mass Index | 313/437 | 1.02E-36 | 3.46E-34 |
| Body Height | 276/385 | 1.32E-32 | 2.23E-30 |
| Blood Pressure | 310/454 | 3.19E-30 | 3.59E-28 |
| Echocardiography | 204/273 | 2.47E-28 | 2.08E-26 |
| Stroke | 211/289 | 6.02E-27 | 4.07E-25 |
| Triglycerides | 180/244 | 4.77E-24 | 2.68E-22 |
| Cholesterol, HDL | 242/357 | 3.57E-23 | 1.72E-21 |
| Body Weight | 161/216 | 1.91E-22 | 8.05E-21 |
| Cholesterol, LDL | 210/304 | 7.78E-22 | 2.92E-20 |
| Hip | 151/202 | 2.39E-21 | 8.06E-20 |
| Cholesterol | 183/268 | 2.28E-18 | 7.02E-17 |
| Myocardial Infarction | 157/229 | 3.32E-16 | 9.36E-15 |
| Coronary Artery Disease | 140/205 | 2.16E-14 | 5.31E-13 |
| Heart Failure | 130/187 | 2.2E-14 | 5.31E-13 |
| Hemoglobins | 113/157 | 2.38E-14 | 5.37E-13 |
| Erythrocyte Count | 87/115 | 2.09E-13 | 4.41E-12 |
| Heart Rate | 110/155 | 2.47E-13 | 4.86E-12 |
| Creatinine | 73/92 | 2.59E-13 | 4.86E-12 |
| Fibrinogen | 69/86 | 4.29E-13 | 7.05E-12 |
| Lipoproteins, VLDL | 69/86 | 4.29E-13 | 7.05E-12 |

### GWAS Catalog 2019: 8,405 genes

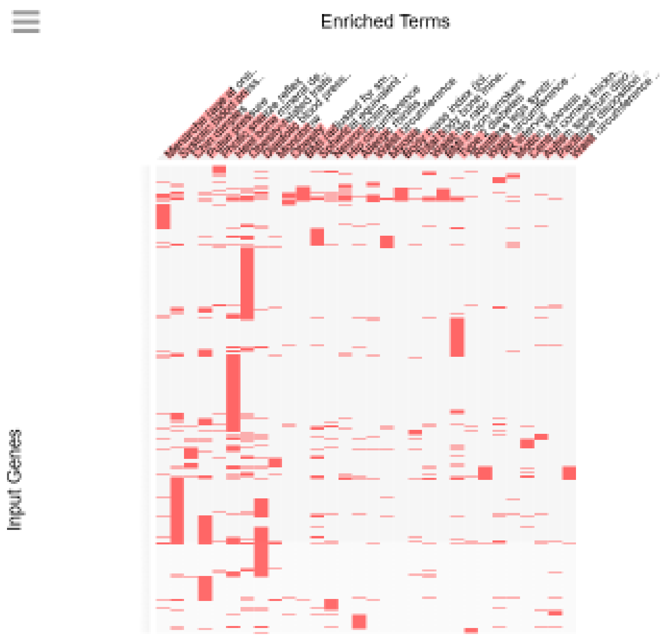

GWAS Catalog 2019 (8,405 genes): Top 40 of 241 significant records

| Term | Overlap | P-value | Adjusted P-value |
|---|---|---|---|
| Menarche (age at onset) | 154/212 | 1.12E-19 | 1.4E-16 |
| Systolic blood pressure | 389/657 | 1.62E-19 | 1.4E-16 |
| Chin dimples | 62/74 | 1.51E-13 | 8.68E-11 |
| Pulse pressure | 321/567 | 9.66E-13 | 4.16E-10 |
| Photic sneeze reflex | 55/65 | 1.6E-12 | 5.51E-10 |
| Heel bone mineral density | 478/898 | 3.09E-12 | 8.88E-10 |
| Obesity-related traits | 427/804 | 6.98E-11 | 1.72E-08 |
| Diastolic blood pressure | 348/646 | 4.87E-10 | 1.05E-07 |
| Monobrow | 58/76 | 1.19E-09 | 2.28E-07 |
| Obesity | 45/55 | 1.61E-09 | 2.78E-07 |
| Height | 288/527 | 2.32E-09 | 3.63E-07 |
| Atrial fibrillation | 146/240 | 2.83E-09 | 3.78E-07 |
| BMI (adjusted for smoking behaviour) | 58/77 | 2.85E-09 | 3.78E-07 |
| Spherical equivalent or myopia (age of diagnosis) | 120/191 | 4.87E-09 | 5.99E-07 |
| Neuroticism | 69/97 | 5.87E-09 | 6.74E-07 |
| Hip circumference | 52/69 | 1.83E-08 | 1.97E-06 |
| Diverticular disease | 100/156 | 2.04E-08 | 2.06E-06 |
| Allergic rhinitis | 77/114 | 3.13E-08 | 3E-06 |
| Waist circumference | 55/75 | 3.67E-08 | 3.33E-06 |
| Myopia | 52/70 | 4.2E-08 | 3.62E-06 |
| Body mass index (joint analysis main effects and smoking interaction) | 56/77 | 4.5E-08 | 3.69E-06 |
| Hand grip strength | 99/156 | 5.05E-08 | 3.96E-06 |
| Total body bone mineral density | 31/36 | 5.77E-08 | 4.32E-06 |
| Waist-hip ratio | 44/58 | 1.65E-07 | 1.14E-05 |
| BMI in non-smokers | 44/58 | 1.65E-07 | 1.14E-05 |
| Type 2 diabetes | 217/397 | 2.08E-07 | 1.38E-05 |
| Restless legs syndrome | 32/39 | 3.48E-07 | 2.22E-05 |
| Blond vs. brown/black hair color | 98/160 | 6.9E-07 | 4.25E-05 |
| Waist circumference adjusted for BMI (adjusted for smoking behaviour) | 56/81 | 7.21E-07 | 4.29E-05 |
| Amyotrophic lateral sclerosis (sporadic) | 109/182 | 8.3E-07 | 4.77E-05 |
| PR interval | 48/67 | 8.6E-07 | 4.78E-05 |
| Motion sickness | 27/32 | 1.01E-06 | 5.45E-05 |
| Male-pattern baldness | 143/251 | 1.15E-06 | 6E-05 |
| Central corneal thickness | 47/66 | 1.48E-06 | 7.14E-05 |
| Paclitaxel disposition in epithelial ovarian cancer | 36/47 | 1.49E-06 | 7.14E-05 |
| Autism spectrum disorder, ADHD, bipolar disorder, MDD, and schizophrenia (combined) | 36/47 | 1.49E-06 | 7.14E-05 |
| Rosacea symptom severity | 87/141 | 1.84E-06 | 8.57E-05 |
| Waist circumference adjusted for BMI (joint analysis main effects and smoking interaction) | 54/79 | 1.98E-06 | 8.99E-05 |
| Waist circumference adjusted for body mass index | 80/128 | 2.29E-06 | 0.000101 |
| Glaucoma (primary open-angle) | 52/76 | 2.92E-06 | 0.000124 |

**Supplemental Figure S3.** Identification of genes implicated in development and manifestations of hundreds physiological and pathological phenotypes and autosomal inheritance in Modern Humans.

**Disease Perturbations from GEO down: 8,405 genes**

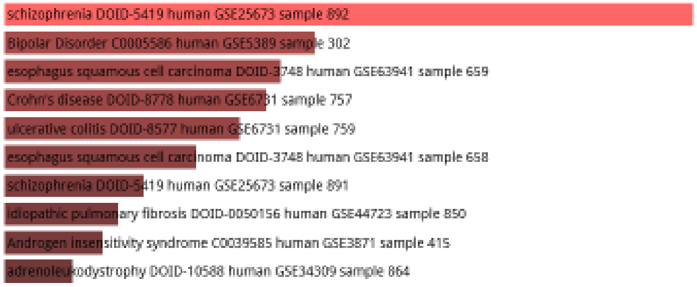


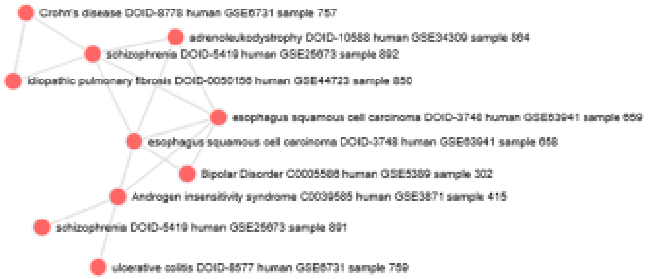


**Disease Perturbations from GEO down: 8,405 genes**

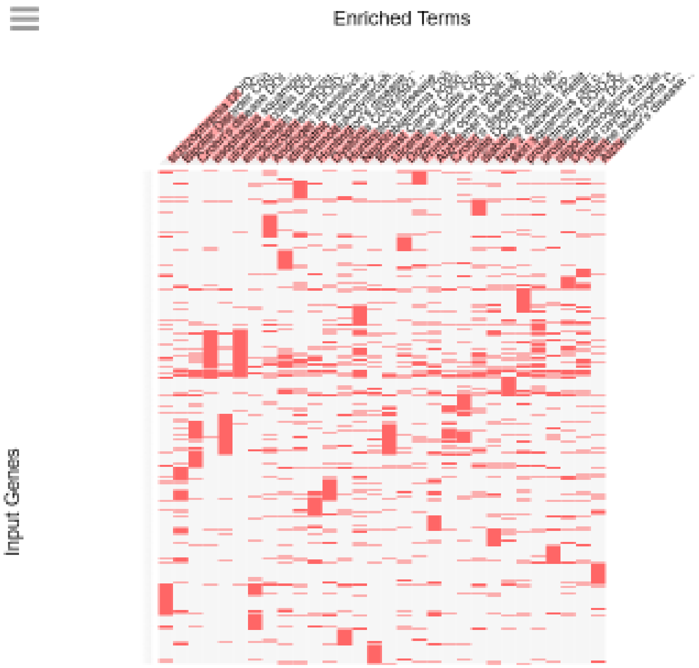

#### Disease Perturbations from GEO down (8,405 genes): Top 30 of 240 significant records

| Term | Overlap | P-value | Adjusted P-value |
|---|---|---|---|
| schizophrenia DOID-5419 human GSE25673 sample 892 | 242/337 | 6.49E-29 | 5.44E-26 |
| Bipolar Disorder C0005586 human GSE5389 sample 302 | 256/395 | 2.78E-20 | 1.17E-17 |
| esophagus squamous cell carcinoma DOID-3748 human GSE63941 sample 659 | 252/391 | 1.65E-19 | 4.6E-17 |
| Crohn's disease DOID-8778 human GSE6731 sample 757 | 240/370 | 3.64E-19 | 7.63E-17 |
| ulcerative colitis DOID-8577 human GSE6731 sample 759 | 246/384 | 1.4E-18 | 2.34E-16 |
| esophagus squamous cell carcinoma DOID-3748 human GSE63941 sample 658 | 256/408 | 1.43E-17 | 2E-15 |
| schizophrenia DOID-5419 human GSE25673 sample 891 | 182/274 | 2.18E-16 | 2.61E-14 |
| idiopathic pulmonary fibrosis DOID-0050156 human GSE44723 sample 850 | 201/312 | 8.28E-16 | 8.68E-14 |
| Androgen insensitivity syndrome C0039585 human GSE3871 sample 415 | 208/327 | 1.94E-15 | 1.81E-13 |
| adrenoleukodystrophy DOID-10588 human GSE34309 sample 864 | 212/338 | 9.23E-15 | 7.75E-13 |
| Huntington's disease DOID-12858 mouse GSE3621 sample 704 | 219/356 | 6.55E-14 | 5E-12 |
| Dystonia C0393593 human GSE3064 sample 329 | 198/317 | 1.27E-13 | 8.89E-12 |
| Nephroblastoma C0027708 human GSE2712 sample 418 | 248/419 | 6.95E-13 | 4.48E-11 |
| Huntington's disease DOID-12858 mouse GSE3583 sample 929 | 183/293 | 1.08E-12 | 6.44E-11 |
| Ulcerative Colitis C0009324 human GSE6731 sample 249 | 213/354 | 3.26E-12 | 1.82E-10 |
| Breast Cancer C0006142 human GSE1378 sample 52 | 184/299 | 6.23E-12 | 3.27E-10 |
| Down syndrome DOID-14250 human GSE42956 sample 1060 | 156/247 | 1.41E-11 | 6.95E-10 |
| Primary open angle glaucoma C0339573 human GSE2705 sample 257 | 156/249 | 3.5E-11 | 1.63E-09 |
| colitis DOID-0060180 human GSE6731 sample 761 | 211/359 | 8.78E-11 | 3.71E-09 |
| Alzheimer's disease DOID-10652 human GSE4757 sample 592 | 216/369 | 8.85E-11 | 3.71E-09 |
| Crohn's disease DOID-8778 human GSE6731 sample 758 | 162/263 | 1.01E-10 | 4.01E-09 |
| prolactinoma DOID-5394 human GSE36314 sample 636 | 251/440 | 1.05E-10 | 4.01E-09 |
| diabetes mellitus type 2 DOID-9352 human GSE12643 sample 766 | 204/346 | 1.22E-10 | 4.34E-09 |
| type 2 diabetes mellitus DOID-9352 human GSE13760 sample 882 | 221/380 | 1.24E-10 | 4.34E-09 |
| skin squamous cell carcinoma DOID-3151 human GSE45164 sample 657 | 177/295 | 2.98E-10 | 1E-08 |
| prostate cancer DOID-10283 human GSE3868 sample 638 | 201/344 | 4.9E-10 | 1.58E-08 |
| Dental cavity, complex C0399396 human GSE1629 sample 175 | 228/401 | 1.12E-09 | 3.49E-08 |
| Alzheimer's disease DOID-10652 human GSE36980 sample 520 | 190/325 | 1.37E-09 | 4.1E-08 |
| oligodendroglioma DOID-3181 human GSE15824 sample 858 | 183/313 | 2.71E-09 | 7.83E-08 |
| breast cancer DOID-1612 human GSE3744 sample 978 | 247/443 | 2.84E-09 | 7.95E-08 |

#### Disease Perturbations from GEO up: 8,405 genes

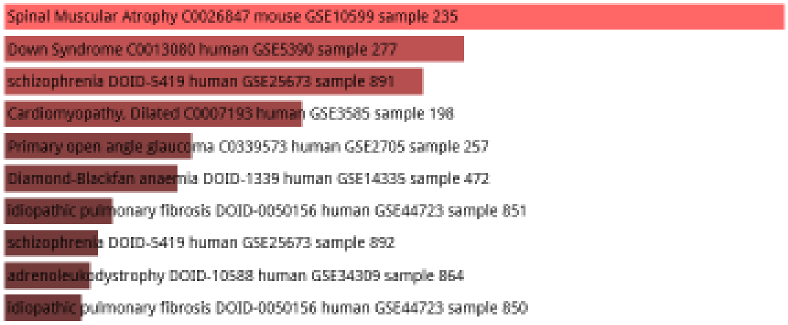


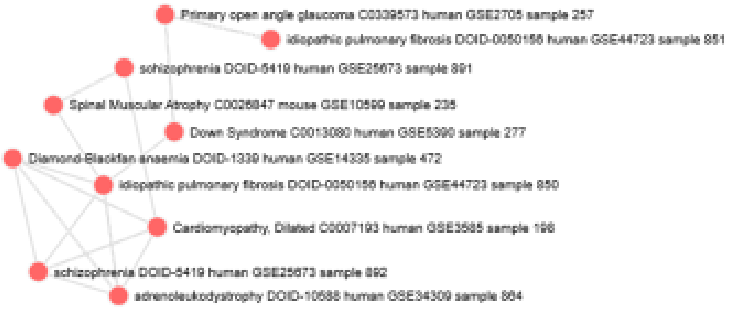

Disease Perturbations from GEO up: 8,405 genes

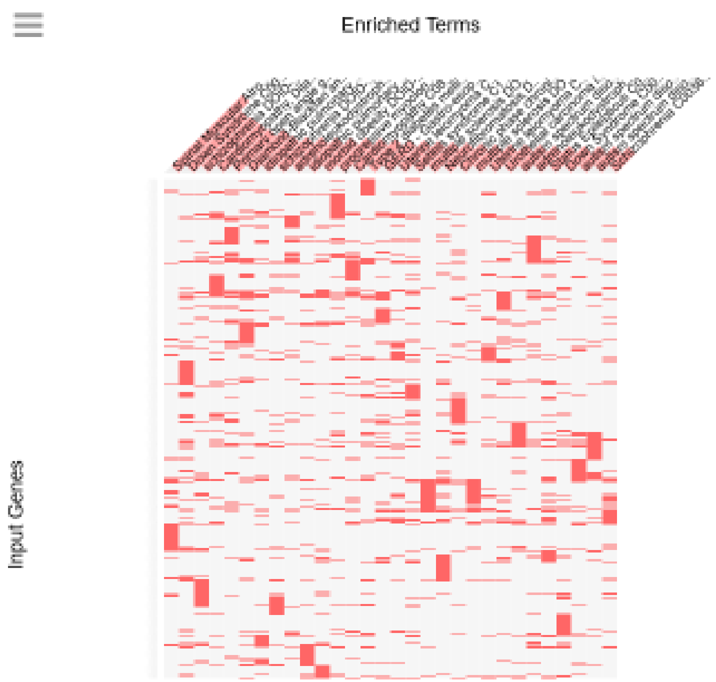


Disease Perturbations from GEO up (8,405 genes): Top 30 of 204 significant records

| Term | Overlap | P-value | Adjusted P-value |
|---|---|---|---|
| Spinal Muscular Atrophy C0026847 mouse GSE10599 sample 235 | 254/368 | 4.69E-26 | 3.93E-23 |
| Down Syndrome C0013080 human GSE5390 sample 277 | 288/460 | 2.08E-19 | 8.72E-17 |
| schizophrenia DOID-5419 human GSE25673 sample 891 | 215/326 | 1.48E-18 | 4.14E-16 |
| Cardiomyopathy, Dilated C0007193 human GSE3585 sample 198 | 209/326 | 4.82E-16 | 1.01E-13 |
| Primary open angle glaucoma C0339573 human GSE2705 sample 257 | 216/351 | 9.23E-14 | 1.55E-11 |
| Diamond-Blackfan anaemia DOID-1339 human GSE14335 sample 472 | 231/382 | 1.88E-13 | 2.64E-11 |
| idiopathic pulmonary fibrosis DOID-0050156 human GSE44723 sample 851 | 185/300 | 4.2E-12 | 5.03E-10 |
| schizophrenia DOID-5419 human GSE25673 sample 892 | 165/263 | 8.13E-12 | 8.53E-10 |
| adrenoleukodystrophy DOID-10588 human GSE34309 sample 864 | 164/262 | 1.23E-11 | 1.14E-09 |
| idiopathic pulmonary fibrosis DOID-0050156 human GSE44723 sample 850 | 177/288 | 1.83E-11 | 1.5E-09 |
| morbid obesity DOID-11981 human GSE48964 sample 583 | 180/294 | 1.96E-11 | 1.5E-09 |
| Cardiomyopathy C0878544 human GSE1869 sample 79 | 197/330 | 5.63E-11 | 3.87E-09 |
| Type 2 diabetes mellitus C0011860 human GSE12643 sample 274 | 205/346 | 5.99E-11 | 3.87E-09 |
| GERD - Gastro-esophageal reflux disease C0017168 human GSE2144 sample 27 | 203/346 | 2.46E-10 | 1.48E-08 |
| Uterine leiomyoma C0042133 human GSE2725 sample 399 | 158/258 | 3.19E-10 | 1.78E-08 |
| multiple sclerosis DOID-2377 human GSE38010 sample 737 | 183/310 | 9.47E-10 | 4.96E-08 |
| Chronic phase chronic myelogenous leukemia DOID-8552 human GSE5550 sample 456 | 192/330 | 1.92E-09 | 9.47E-08 |
| autism spectrum disorder DOID-0060041 human GSE28521 sample 1041 | 185/317 | 2.71E-09 | 1.26E-07 |
| Setleis syndrome C1744559 human GSE16524 sample 285 | 222/393 | 4E-09 | 1.77E-07 |
| Neurofibromatosis DOID-8712 mouse GSE1482 sample 667 | 210/369 | 4.54E-09 | 1.9E-07 |
| multiple sclerosis DOID-2377 human GSE38010 sample 738 | 180/309 | 5.36E-09 | 2.14E-07 |
| Uterine leiomyoma C0042133 human GSE593 sample 16 | 160/270 | 7.23E-09 | 2.76E-07 |
| autism spectrum disorder DOID-0060041 human GSE28521 sample 1040 | 194/341 | 1.8E-08 | 6.56E-07 |
| Urothelial carcinoma in situ C0334267 human GSE3167 sample 229 | 213/380 | 1.94E-08 | 6.77E-07 |
| Status Epilepticus C0038220 rat GSE4236 sample 391 | 191/336 | 2.51E-08 | 8.11E-07 |
| autism spectrum disorder DOID-0060041 human GSE28521 sample 1039 | 191/336 | 2.51E-08 | 8.11E-07 |
| adrenoleukodystrophy DOID-10588 human GSE34308 sample 709 | 205/365 | 2.88E-08 | 8.95E-07 |
| Down Syndrome C0013080 human GSE10758 sample 310 | 210/376 | 3.57E-08 | 1.07E-06 |
| chronic lymphocytic leukemia DOID-1040 human GSE6691 sample 786 | 180/315 | 3.77E-08 | 1.09E-06 |
| Oligodendroglioma C0028945 human GSE2223 sample 116 | 167/289 | 4.08E-08 | 1.14E-06 |

**Supplemental Figure S4.** Identification of genes expression of which is altered in several hundred common human disorders.

Rare Diseases GeneRIF Gene Lists: 8,405 genes

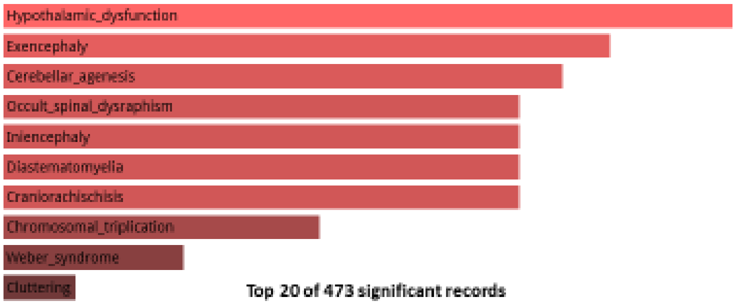


Top 20 of 473 significant records

| Term | Overlap | P-value | Adjusted P-value |
|---|---|---|---|
| Hypothalamic_dysfunction | 90/128 | 8.12E-11 | 6.55E-08 |
| Exencephaly | 158/256 | 1.38E-10 | 6.55E-08 |
| Cerebellar_agenesis | 198/335 | 1.7E-10 | 6.55E-08 |
| Occult_spinal_dysraphism | 157/255 | 2.05E-10 | 6.55E-08 |
| Diastematomyelia | 157/255 | 2.05E-10 | 6.55E-08 |
| Craniorachischisis | 157/255 | 2.05E-10 | 6.55E-08 |
| Iniencephaly | 157/255 | 2.05E-10 | 6.55E-08 |
| Chromosomal_triplication | 201/344 | 4.9E-10 | 1.37E-07 |
| Weber_syndrome | 94/139 | 8.85E-10 | 2.2E-07 |
| Cluttering | 106/162 | 1.41E-09 | 3.16E-07 |
| Mental_retardation_epilepsy | 190/326 | 1.92E-09 | 3.9E-07 |
| Single_ventricular_heart | 73/104 | 5.49E-09 | 1.02E-06 |
| Sydenham's_chorea | 209/369 | 8.42E-09 | 1.34E-06 |
| Chorea_minor | 209/369 | 8.42E-09 | 1.34E-06 |
| Antisocial_personality_disorder | 46/59 | 1.93E-08 | 2.87E-06 |
| Sudden_infant_death_syndrome | 103/162 | 2.37E-08 | 3.31E-06 |
| Stress_cardiomyopathy | 110/176 | 3.16E-08 | 4.10E-06 |
| N_syndrome | 222/401 | 3.89E-08 | 4.83E-06 |
| Basilar_migraine | 127/210 | 4.85E-08 | 5.7E-06 |
| Corpus_callosum_agenesis | 89/138 | 7.92E-08 | 8.85E-06 |

Rare Diseases GeneRIF Gene Lists: 8,405 genes

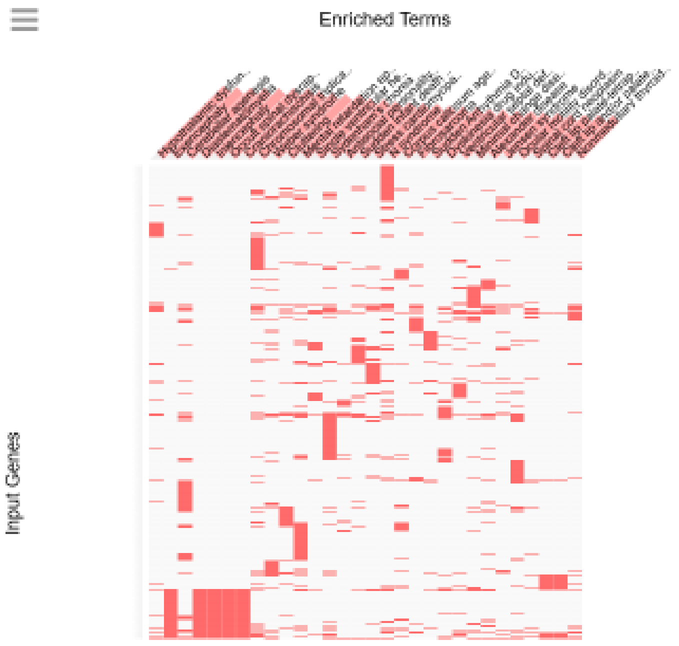

Rare Diseases GeneRIF ARCHS4 Predictions: 8,405 genes

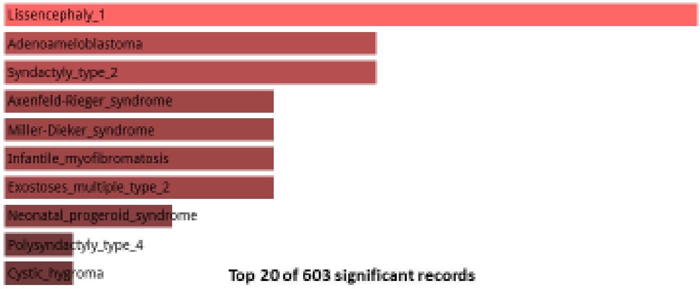


Top 20 of 603 significant records

| Term | Overlap | P-value | Adjusted P-value |
|---|---|---|---|
| Lissencephaly_1 | 154/200 | 4.79E-24 | 1.08E-20 |
| Adenoameloblastoma | 151/200 | 4.35E-22 | 3.26E-19 |
| Syndactyly_type_2 | 151/200 | 4.35E-22 | 3.26E-19 |
| Axenfeld-Rieger_syndrome | 150/200 | 1.85E-21 | 5.94E-19 |
| Miller-Dieker_syndrome | 150/200 | 1.85E-21 | 5.94E-19 |
| Exostoses_multiple_type_2 | 150/200 | 1.85E-21 | 5.94E-19 |
| Infantile_myofibromatosis | 150/200 | 1.85E-21 | 5.94E-19 |
| Neonatal_progeroid_syndrome | 149/200 | 7.69E-21 | 2.16E-18 |
| Polysyndactyly_type_4 | 148/200 | 3.11E-20 | 6.97E-18 |
| Cystic_hygroma | 148/200 | 3.11E-20 | 6.97E-18 |
| Acromesomelic_dysplasia_Hunter_Thompson_type | 147/200 | 1.22E-19 | 2.11E-17 |
| Acromesomelic_dysplasia | 147/200 | 1.22E-19 | 2.11E-17 |
| Exostoses_multiple_type_1 | 147/200 | 1.22E-19 | 2.11E-17 |
| Scholte_syndrome | 146/200 | 4.69E-19 | 6.58E-17 |
| Congenital_diaphragmatic_hernia | 146/200 | 4.69E-19 | 6.58E-17 |
| Hereditary_multiple_osteochondromas | 146/200 | 4.69E-19 | 6.58E-17 |
| Aortopulmonary_window | 145/200 | 1.76E-18 | 1.88E-16 |
| Corpus_callosum_agenesis | 145/200 | 1.76E-18 | 1.88E-16 |
| Apert_syndrome | 145/200 | 1.76E-18 | 1.88E-16 |
| Subependymoma | 145/200 | 1.76E-18 | 1.88E-16 |

Rare Diseases GeneRIF ARCHS4 Predictions: 8,405 genes

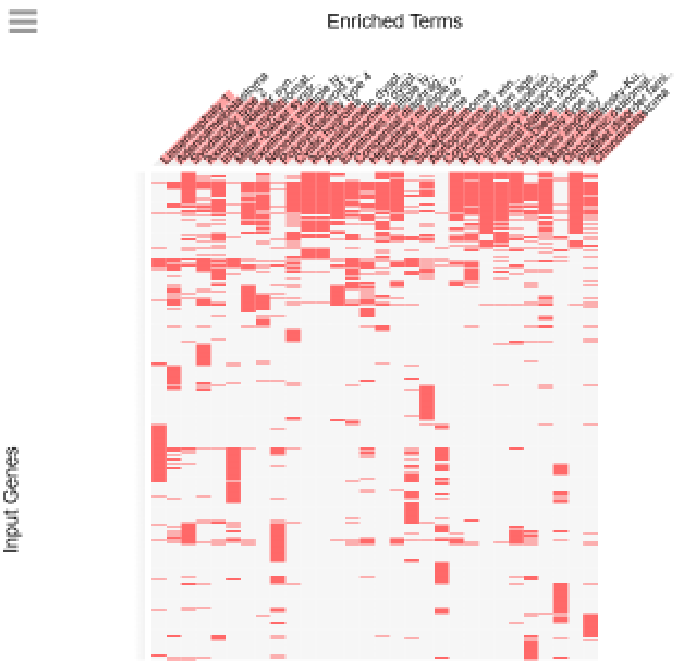

### Rare Diseases AutoRIF ARCHS4 Predictions: 8,405 genes

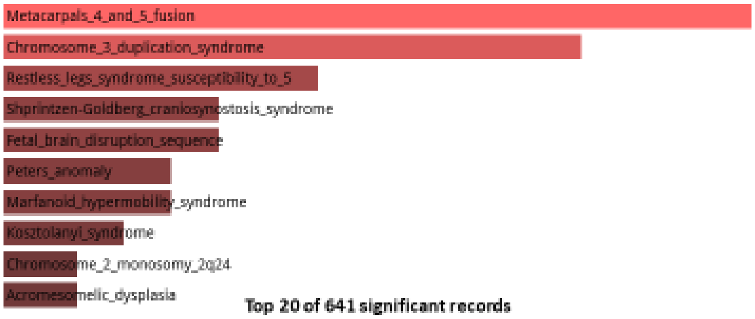


**Top 20 of 641 significant records**

| Term | Overlap | P-value | Adjusted P-value |
|---|---|---|---|
| Metacarpals_4_and_5_fusion | 163/200 | 1.33E-30 | 4.97E-27 |
| Chromosome_3_duplication_syndrome | 160/200 | 2.69E-28 | 5.01E-25 |
| Restless_legs_syndrome_susceptibility_to_5 | 155/200 | 1.01E-24 | 1.25E-21 |
| Shprintzen-Goldberg_craniosynostosis_syndrome | 153/200 | 2.21E-23 | 1.65E-20 |
| Fetal_brain_disruption_sequence | 153/200 | 2.21E-23 | 1.65E-20 |
| Peters_anomaly | 152/200 | 9.95E-23 | 5.3E-20 |
| Marfanoid_hypermobility_syndrome | 152/200 | 9.95E-23 | 5.3E-20 |
| Kosztolanyi_syndrome | 151/200 | 4.35E-22 | 2.03E-19 |
| Chromosome_2_monosomy_2q24 | 150/200 | 1.85E-21 | 6.28E-19 |
| Acromesomelic_dysplasia_Hunter_Thompson_type | 150/200 | 1.85E-21 | 6.28E-19 |
| Acromesomelic_dysplasia | 150/200 | 1.85E-21 | 6.28E-19 |
| Chromosome_3_trisomy_3p | 149/200 | 7.69E-21 | 2.2E-18 |
| Buschke_Ollendorff_syndrome | 149/200 | 7.69E-21 | 2.2E-18 |
| Fraser_like_syndrome | 148/200 | 3.11E-20 | 6.81E-18 |
| Crandall_syndrome | 148/200 | 3.11E-20 | 6.81E-18 |
| Orofaciodigital_syndrome_11 | 148/200 | 3.11E-20 | 6.81E-18 |
| Postaxial_polydactyly_mental_retardation | 148/200 | 3.11E-20 | 6.81E-18 |
| Short_rib-polydactyly_syndrome_type_4 | 147/200 | 1.22E-19 | 2.28E-17 |
| Trichorhinophalangeal_syndrome_type_3 | 147/200 | 1.22E-19 | 2.28E-17 |
| Duane_syndrome | 147/200 | 1.22E-19 | 2.28E-17 |

### Rare Diseases AutoRIF ARCHS4 Predictions: 8,405 genes

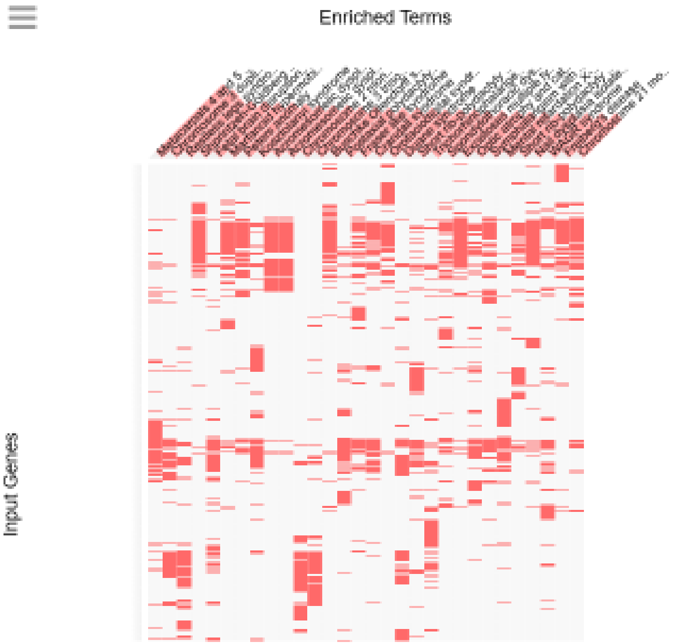

Rare Diseases AutoRIF Gene Lists: 8,405 genes

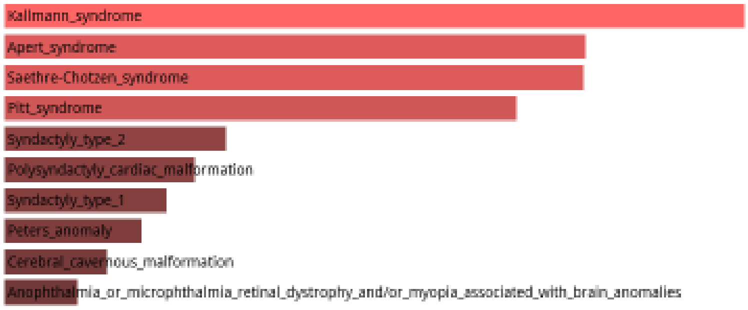


Top 20 of 1,116 significant records

| Term | Overlap | P-value | Adjusted P-value |
|---|---|---|---|
| Kallmann_syndrome | 159/247 | 9.25E-13 | 3.25E-09 |
| Apert_syndrome | 143/219 | 2.58E-12 | 3.25E-09 |
| Saethre-Chotzen_syndrome | 136/206 | 2.62E-12 | 3.25E-09 |
| Pitt_syndrome | 241/410 | 4.01E-12 | 3.73E-09 |
| Syndactyly_type_2 | 124/188 | 2.62E-11 | 1.95E-08 |
| Polysyndactyly_cardiac_malformation | 77/104 | 3.2E-11 | 1.98E-08 |
| Syndactyly_type_1 | 122/185 | 3.85E-11 | 2.04E-08 |
| Peters_anomaly | 75/101 | 4.53E-11 | 2.1E-08 |
| Cerebral_cavernous_malformation | 157/252 | 5.65E-11 | 2.33E-08 |
| Anophthalmia_or_microphthalmia_retinal_dystrophy_and/or_myopia_associated_with_brain_anomalies | 182/301 | 6.82E-11 | 2.36E-08 |
| X-linked_periventricular_heterotopia | 89/126 | 6.98E-11 | 2.36E-08 |
| Aniridia | 171/282 | 1.85E-10 | 5.73E-08 |
| Dominant_cleft_palate | 163/267 | 2.37E-10 | 6.76E-08 |
| Craniofacial_and_skeletal_defects | 94/137 | 2.64E-10 | 7.01E-08 |
| Anodontia | 140/223 | 2.84E-10 | 7.03E-08 |
| Glaucoma_congenital | 202/345 | 3.48E-10 | 8.08E-08 |
| Childhood-Onset_Schizophrenia | 168/278 | 3.87E-10 | 8.46E-08 |
| Hennekam_syndrome | 139/223 | 6.68E-10 | 1.38E-07 |
| Kurczynski_Casperson_syndrome | 114/176 | 8.57E-10 | 1.68E-07 |
| Osteochondroma | 177/299 | 1.31E-09 | 2.44E-07 |

Rare Diseases AutoRIF Gene Lists: 8,405 genes

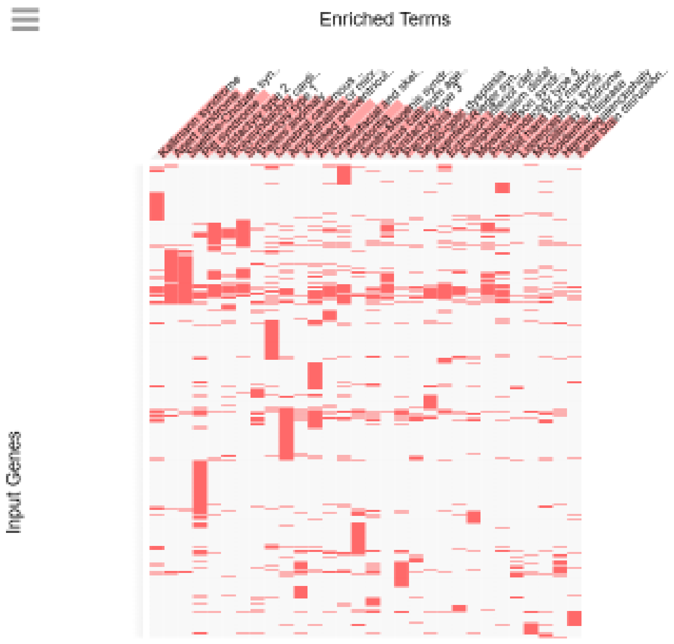


**Supplemental Figure S5.** Identification of genes implicated in more than 1,000 records classified as human rare diseases.

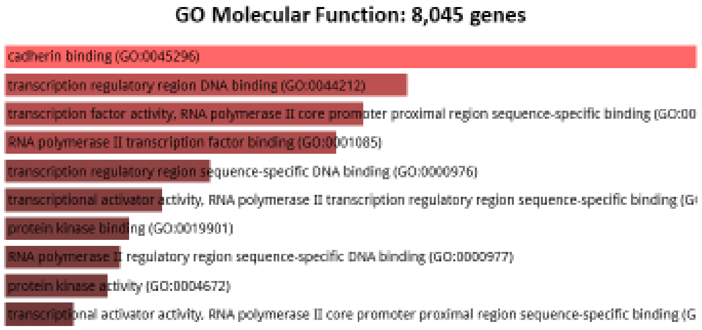
GO Molecular Function: 8,045 genes
cadherin binding (GO:0045296)
transcription regulatory region DNA binding (GO:0044212)
transcription factor activity, RNA polymerase II core promoter proximal region sequence-specific binding (GO:00
RNA polymerase II transcription factor binding (GO:0001085)
transcription regulatory region sequence-specific DNA binding (GO:0000976)
transcriptional activator activity, RNA polymerase II transcription regulatory region sequence-specific binding (G
protein kinase binding (GO:0019901)
RNA polymerase II regulatory region sequence-specific DNA binding (GO:0000977)
protein kinase activity (GO:0004672)
transcriptional activator activity, RNA polymerase II core promoter proximal region sequence-specific binding (G

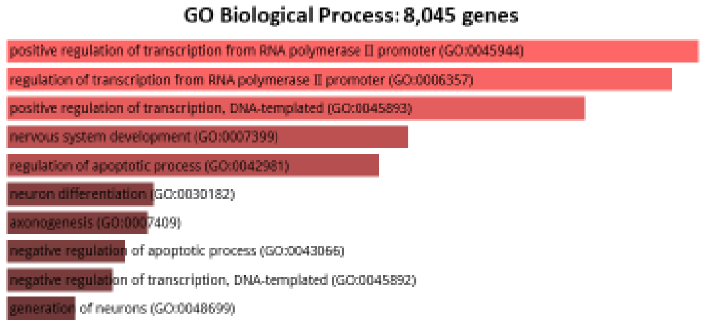
GO Biological Process: 8,045 genes
positive regulation of transcription from RNA polymerase II promoter (GO:0045944)
regulation of transcription from RNA polymerase II promoter (GO:0006357)
positive regulation of transcription, DNA-templated (GO:0045893)
nervous system development (GO:0007399)
regulation of apoptotic process (GO:0042981)
neuron differentiation (GO:0030182)
axonogenesis (GO:0007409)
negative regulation of apoptotic process (GO:0043066)
negative regulation of transcription, DNA-templated (GO:0045892)
generation of neurons (GO:0048699)

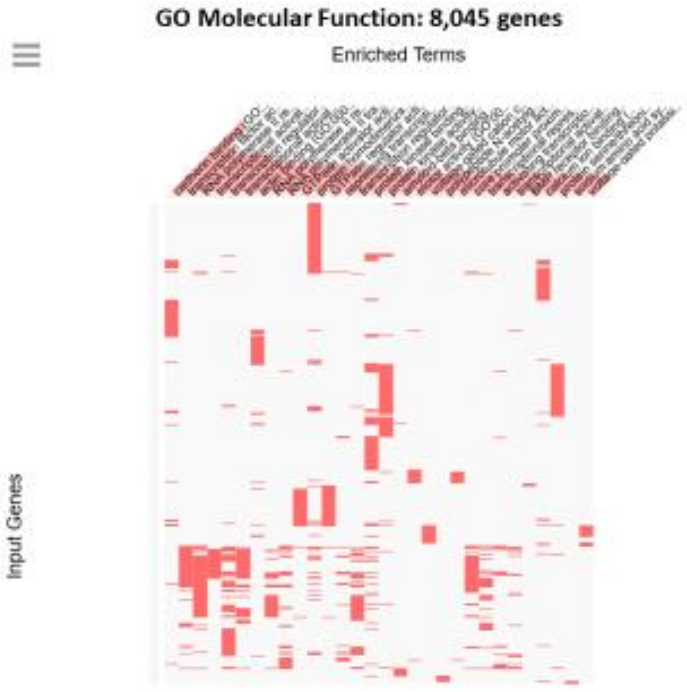
GO Molecular Function: 8,045 genes
Enriched Terms
Input Genes

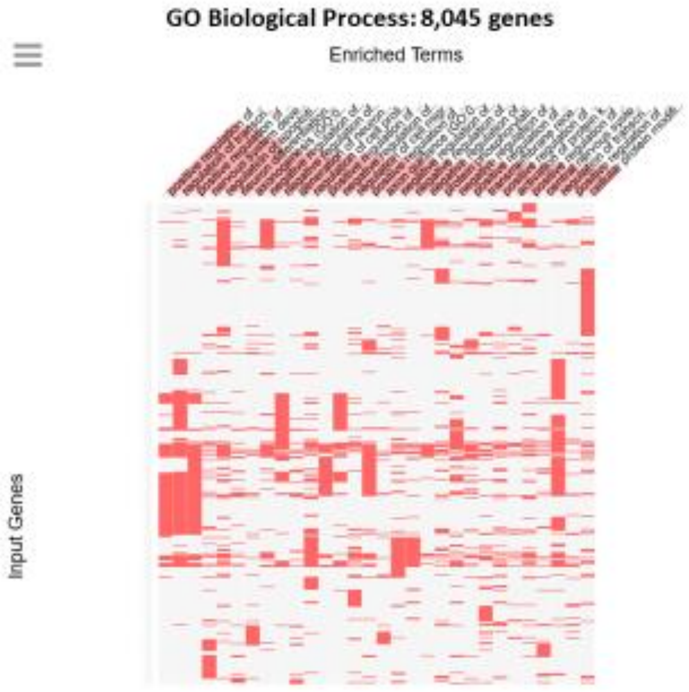
GO Biological Process: 8,045 genes
Enriched Terms
Input Genes

### GO Biological Process (8,045 genes): Top 35 of 308 significant records

| Term | Overlap | P-value | Adjusted P-value |
|---|---|---|---|
| positive regulation of transcription from RNA polymerase II promoter (GO:0045944) | 485/849 | 1.12E-19 | 5.67E-16 |
| regulation of transcription from RNA polymerase II promoter (GO:0006357) | 786/1479 | 2.51E-19 | 6.37E-16 |
| positive regulation of transcription, DNA-templated (GO:0045893) | 611/1121 | 3.46E-18 | 5.85E-15 |
| nervous system development (GO:0007399) | 276/456 | 7.07E-16 | 8.98E-13 |
| regulation of apoptotic process (GO:0042981) | 453/816 | 1.74E-15 | 1.77E-12 |
| neuron differentiation (GO:0030182) | 100/140 | 1.57E-12 | 1.33E-09 |
| axonogenesis (GO:0007409) | 146/224 | 1.88E-12 | 1.37E-09 |
| negative regulation of apoptotic process (GO:0043066) | 279/486 | 3.63E-12 | 2.31E-09 |
| negative regulation of transcription, DNA-templated (GO:0045892) | 437/814 | 5.44E-12 | 3.07E-09 |
| generation of neurons (GO:0048699) | 93/131 | 1.66E-11 | 7.69E-09 |
| regulation of cell proliferation (GO:0042127) | 400/741 | 1.67E-11 | 7.69E-09 |
| positive regulation of nucleic acid-templated transcription (GO:1903508) | 284/503 | 2.98E-11 | 1.26E-08 |
| negative regulation of transcription from RNA polymerase II promoter (GO:0000122) | 314/566 | 4.51E-11 | 1.76E-08 |
| regulation of cell migration (GO:0030334) | 190/317 | 7.61E-11 | 2.76E-08 |
| positive regulation of gene expression (GO:0010628) | 410/772 | 1.69E-10 | 5.73E-08 |
| axon guidance (GO:0007411) | 106/159 | 2.78E-10 | 8.82E-08 |
| negative regulation of cellular process (GO:0048523) | 295/535 | 4.21E-10 | 1.26E-07 |
| negative regulation of cell proliferation (GO:0008285) | 210/364 | 9.35E-10 | 2.64E-07 |
| negative regulation of programmed cell death (GO:0043069) | 232/409 | 1.08E-09 | 2.88E-07 |
| protein phosphorylation (GO:0006468) | 261/471 | 2.26E-09 | 5.62E-07 |
| negative regulation of gene expression (GO:0010629) | 332/619 | 2.32E-09 | 5.62E-07 |
| positive regulation of macromolecule metabolic process (GO:0010604) | 165/277 | 2.5E-09 | 5.78E-07 |
| transmembrane receptor protein tyrosine kinase signaling pathway (GO:0007169) | 224/397 | 3.91E-09 | 8.63E-07 |
| positive regulation of cell differentiation (GO:0045597) | 122/195 | 5.25E-09 | 1.11E-06 |
| activation of protein kinase activity (GO:0032147) | 142/234 | 5.85E-09 | 1.19E-06 |
| positive regulation of protein phosphorylation (GO:0001934) | 231/413 | 6.36E-09 | 1.24E-06 |
| central nervous system development (GO:0007417) | 133/218 | 1.1E-08 | 2.06E-06 |
| negative regulation of cellular macromolecule biosynthetic process (GO:2000113) | 278/513 | 1.29E-08 | 2.34E-06 |
| regulation of transcription, DNA-templated (GO:0006355) | 776/1599 | 2.65E-08 | 4.64E-06 |
| positive regulation of epithelial cell migration (GO:0010634) | 55/75 | 3.67E-08 | 6.13E-06 |
| cellular protein modification process (GO:0006464) | 504/1002 | 3.74E-08 | 6.13E-06 |
| positive regulation of cell migration (GO:0030335) | 133/222 | 5.27E-08 | 8.35E-06 |
| positive regulation of cell proliferation (GO:0008284) | 233/425 | 5.42E-08 | 8.35E-06 |
| neuron projection morphogenesis (GO:0048812) | 103/164 | 5.98E-08 | 8.94E-06 |
| positive regulation of multicellular organismal process (GO:0051240) | 123/203 | 6.72E-08 | 9.66E-06 |

### GO Molecular Function (8,045 genes): Top 30 of 81 significant records

| Term | Overlap | P-value | Adjusted P-value |
|---|---|---|---|
| cadherin binding (GO:0045296) | 191/314 | 1.1E-11 | 1.26E-08 |
| transcription regulatory region DNA binding (GO:0044212) | 216/375 | 6.56E-10 | 3.76E-07 |
| transcription factor activity, RNA polymerase II core promoter proximal region sequence-specific binding (GO:0000982) | 168/281 | 1.22E-09 | 4.66E-07 |
| RNA polymerase II transcription factor binding (GO:0001085) | 84/122 | 1.79E-09 | 5.11E-07 |
| transcription regulatory region sequence-specific DNA binding (GO:0000976) | 171/293 | 1.06E-08 | 2.43E-06 |
| transcriptional activator activity, RNA polymerase II transcription regulatory region sequence-specific binding (GO:0001228) | 166/285 | 2.1E-08 | 4.01E-06 |
| protein kinase binding (GO:0019901) | 268/496 | 3.35E-08 | 5.4E-06 |
| RNA polymerase II regulatory region sequence-specific DNA binding (GO:0000977) | 251/461 | 3.78E-08 | 5.4E-06 |
| protein kinase activity (GO:0004672) | 276/514 | 4.5E-08 | 5.72E-06 |
| transcriptional activator activity, RNA polymerase II core promoter proximal region sequence-specific binding (GO:0001077) | 109/176 | 7.35E-08 | 8.4E-06 |
| GTPase regulator activity (GO:0030695) | 154/276 | 2.44E-06 | 0.000254 |
| amyloid-beta binding (GO:0001540) | 37/50 | 4.47E-06 | 0.000426 |
| protein serine/threonine kinase activity (GO:0004674) | 197/369 | 5.99E-06 | 0.000518 |
| repressing transcription factor binding (GO:0070491) | 39/54 | 6.67E-06 | 0.000518 |
| RNA polymerase II regulatory region DNA binding (GO:0001012) | 116/202 | 6.8E-06 | 0.000518 |
| GTPase activator activity (GO:0005096) | 139/250 | 9.43E-06 | 0.000675 |
| voltage-gated cation channel activity (GO:0022843) | 64/101 | 1.2E-05 | 0.000807 |
| motor activity (GO:0003774) | 56/86 | 1.28E-05 | 0.000811 |
| regulatory region DNA binding (GO:0000975) | 126/225 | 1.52E-05 | 0.000916 |
| core promoter proximal region sequence-specific DNA binding (GO:0000987) | 152/279 | 1.65E-05 | 0.000943 |
| protein homodimerization activity (GO:0042803) | 332/665 | 1.79E-05 | 0.000973 |
| actin binding (GO:0003779) | 140/255 | 2.09E-05 | 0.001086 |
| PDZ domain binding (GO:0030165) | 43/63 | 2.29E-05 | 0.001141 |
| RNA polymerase II core promoter proximal region sequence-specific DNA binding (GO:0000978) | 143/263 | 3.29E-05 | 0.001569 |
| transcriptional repressor activity, RNA polymerase II transcription regulatory region sequence-specific binding (GO:0001227) | 92/160 | 5.43E-05 | 0.002452 |
| protein tyrosine kinase activity (GO:0004713) | 86/148 | 5.57E-05 | 0.002452 |
| microtubule motor activity (GO:0003777) | 41/61 | 6.16E-05 | 0.002608 |
| tubulin binding (GO:0015631) | 138/256 | 7.7E-05 | 0.003145 |
| microtubule binding (GO:0008017) | 109/196 | 8.13E-05 | 0.003208 |
| acetylgalactosaminyltransferase activity (GO:0008376) | 34/49 | 9.81E-05 | 0.003742 |

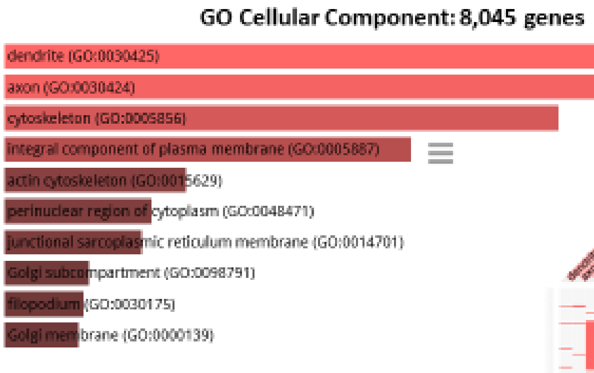


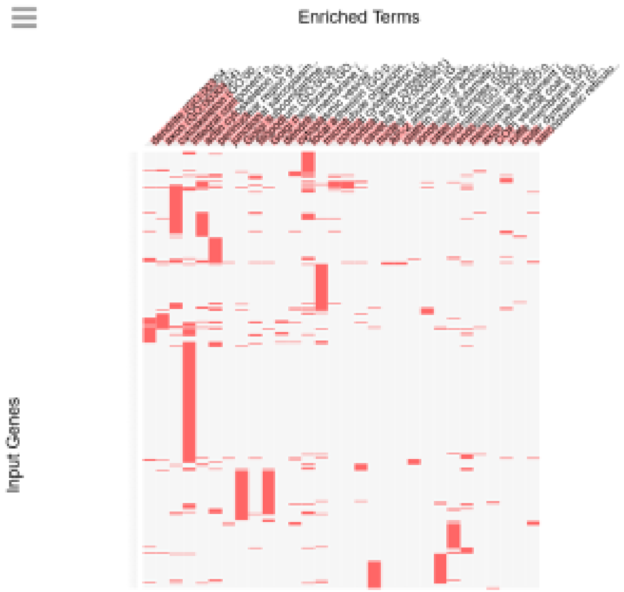


GO Cellular Component (8,045 genes): 29 significant records

| Term | Overlap | P-value | Adjusted P-value |
|---|---|---|---|
| cytoskeleton (GO:0005856) | 284/521 | 4.18E-09 | 1.85E-06 |
| dendrite (GO:0030425) | 131/216 | 2.35E-08 | 5.12E-06 |
| axon (GO:0030424) | 92/142 | 3.47E-08 | 5.12E-06 |
| actin cytoskeleton (GO:0015629) | 169/295 | 7.67E-08 | 8.23E-06 |
| integral component of plasma membrane (GO:0005887) | 711/1464 | 9.29E-08 | 8.23E-06 |
| focal adhesion (GO:0005925) | 194/357 | 1.52E-06 | 0.000112 |
| Golgi subcompartment (GO:0098791) | 250/480 | 4.46E-06 | 0.000282 |
| perinuclear region of cytoplasm (GO:0048471) | 198/379 | 3.32E-05 | 0.001839 |
| cytoplasmic vesicle membrane (GO:0030659) | 38/55 | 4.48E-05 | 0.002203 |
| Golgi membrane (GO:0000139) | 225/443 | 0.000104 | 0.004546 |
| cortical cytoskeleton (GO:0030863) | 36/53 | 0.000123 | 0.004546 |
| cortical actin cytoskeleton (GO:0030864) | 36/53 | 0.000123 | 0.004546 |
| caveola (GO:0005901) | 38/57 | 0.000148 | 0.00505 |
| filopodium (GO:0030175) | 40/61 | 0.000173 | 0.005461 |
| membrane raft (GO:0045121) | 70/120 | 0.000224 | 0.006622 |
| endoplasmic reticulum lumen (GO:0005788) | 142/271 | 0.000339 | 0.009382 |
| microtubule organizing center (GO:0005815) | 251/508 | 0.000401 | 0.010456 |
| cytoplasmic vesicle (GO:0031410) | 115/216 | 0.000546 | 0.013065 |
| chromatin (GO:0000785) | 153/297 | 0.00056 | 0.013065 |
| nuclear chromatin (GO:0000790) | 132/254 | 0.000826 | 0.018285 |
| catenin complex (GO:0016342) | 21/29 | 0.0009 | 0.018978 |
| ionotropic glutamate receptor complex (GO:0008328) | 27/40 | 0.001001 | 0.020165 |
| dendrite membrane (GO:0032590) | 16/21 | 0.001571 | 0.030261 |
| cation channel complex (GO:0034703) | 40/66 | 0.00177 | 0.032667 |
| centrosome (GO:0005813) | 225/462 | 0.00199 | 0.035259 |
| main axon (GO:0044304) | 23/34 | 0.002257 | 0.038461 |
| junctional sarcoplasmic reticulum membrane (GO:0014701) | 10-Sep | 0.002537 | 0.040494 |
| microtubule cytoskeleton (GO:0015630) | 191/389 | 0.002642 | 0.040494 |
| actin-based cell projection (GO:0098858) | 42/71 | 0.002651 | 0.040494 |
| ruffle membrane (GO:0032587) | 33/54 | 0.003598 | 0.053124 |

**Supplemental Figure S6.** Gene ontology analyses of putative regulatory targets of genetic loci harboring human-specific SNCs.

**KEGG 2019 Human: 8,405 genes**

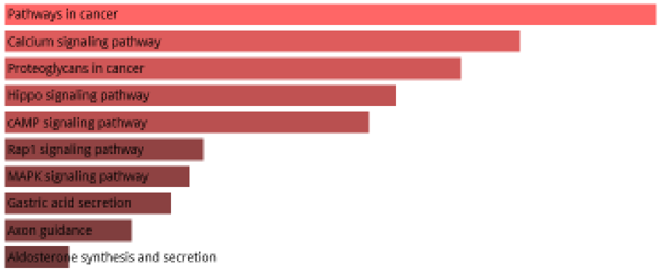


**KEGG 2019 Mouse: 8,405 genes**

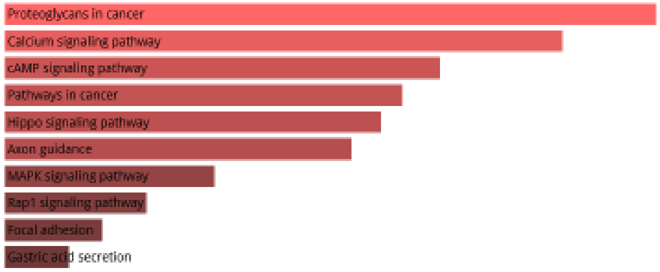


**KEGG 2019 Human: 8,405 genes**

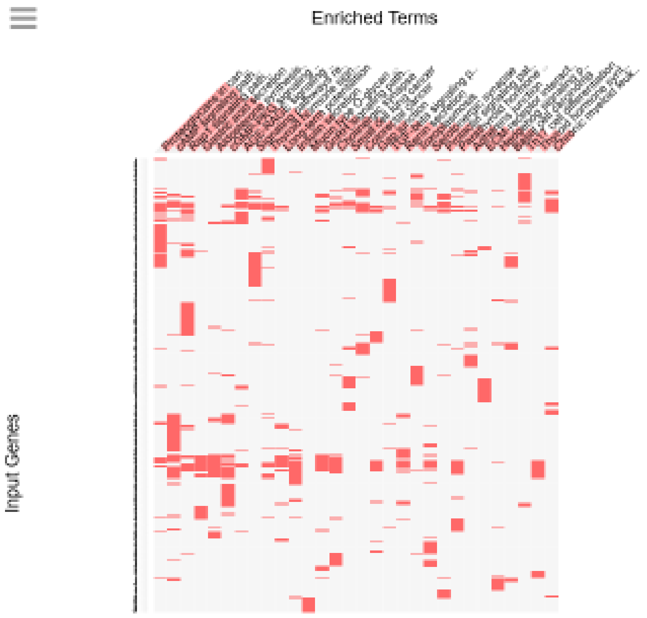

#### KEGG 2019 Human (8,405 genes): Top 40 of 129 significant records

| Term | Overlap | P-value | Adjusted P-value |
|---|---|---|---|
| Pathways in cancer | 312/530 | 2.04E-15 | 6.28E-13 |
| Calcium signaling pathway | 130/188 | 4.19E-14 | 6.45E-12 |
| Proteoglycans in cancer | 136/201 | 1.55E-13 | 1.59E-11 |
| Hippo signaling pathway | 112/160 | 6.61E-13 | 5.09E-11 |
| cAMP signaling pathway | 140/212 | 1.21E-12 | 7.48E-11 |
| Rap1 signaling pathway | 133/206 | 4.85E-11 | 2.49E-09 |
| MAPK signaling pathway | 179/295 | 6.48E-11 | 2.85E-09 |
| Gastric acid secretion | 59/75 | 9.75E-11 | 3.75E-09 |
| Axon guidance | 118/181 | 2.39E-10 | 8.17E-09 |
| Aldosterone synthesis and secretion | 71/98 | 9.54E-10 | 2.94E-08 |
| cGMP-PKG signaling pathway | 108/166 | 1.62E-09 | 4.54E-08 |
| Focal adhesion | 125/199 | 2.39E-09 | 6.14E-08 |
| AGE-RAGE signaling pathway in diabetic complications | 70/100 | 1.35E-08 | 3.13E-07 |
| Dopaminergic synapse | 87/131 | 1.42E-08 | 3.13E-07 |
| Signaling pathways regulating pluripotency of stem cells | 91/139 | 1.94E-08 | 3.98E-07 |
| Wnt signaling pathway | 101/158 | 2.15E-08 | 4.13E-07 |
| Adrenergic signaling in cardiomyocytes | 94/145 | 2.36E-08 | 4.28E-07 |
| Gastric cancer | 96/149 | 2.67E-08 | 4.56E-07 |
| Amoebiasis | 67/96 | 3.33E-08 | 5.32E-07 |
| Thyroid hormone signaling pathway | 78/116 | 3.45E-08 | 5.32E-07 |
| PI3K-Akt signaling pathway | 199/354 | 4.17E-08 | 6.11E-07 |
| Long-term potentiation | 50/67 | 5.95E-08 | 8.32E-07 |
| Melanogenesis | 69/101 | 8.04E-08 | 1.08E-06 |
| Breast cancer | 93/147 | 1.57E-07 | 2.02E-06 |
| Insulin secretion | 60/86 | 1.75E-07 | 2.16E-06 |
| Cushing syndrome | 97/155 | 1.84E-07 | 2.18E-06 |
| Circadian entrainment | 66/97 | 1.94E-07 | 2.22E-06 |
| Glutamatergic synapse | 75/114 | 2.48E-07 | 2.73E-06 |
| Mucin type O-glycan biosynthesis | 27/31 | 2.65E-07 | 2.81E-06 |
| Relaxin signaling pathway | 83/130 | 4.06E-07 | 4.17E-06 |
| GnRH signaling pathway | 63/93 | 4.68E-07 | 4.65E-06 |
| Cholinergic synapse | 73/112 | 6.18E-07 | 5.95E-06 |
| Oxytocin signaling pathway | 94/153 | 9.5E-07 | 8.87E-06 |
| Inflammatory mediator regulation of TRP channels | 66/100 | 1.08E-06 | 9.78E-06 |
| Small cell lung cancer | 62/93 | 1.35E-06 | 1.19E-05 |
| Human papillomavirus infection | 181/330 | 1.53E-06 | 1.31E-05 |
| Neuroactive ligand-receptor interaction | 184/338 | 2.41E-06 | 2.01E-05 |
| Colorectal cancer | 57/86 | 4.72E-06 | 3.82E-05 |
| Gap junction | 58/88 | 5.1E-06 | 4.03E-05 |
| TGF-beta signaling pathway | 59/90 | 5.49E-06 | 4.22E-05 |

#### KEGG 2019 Mouse: 8,405 genes

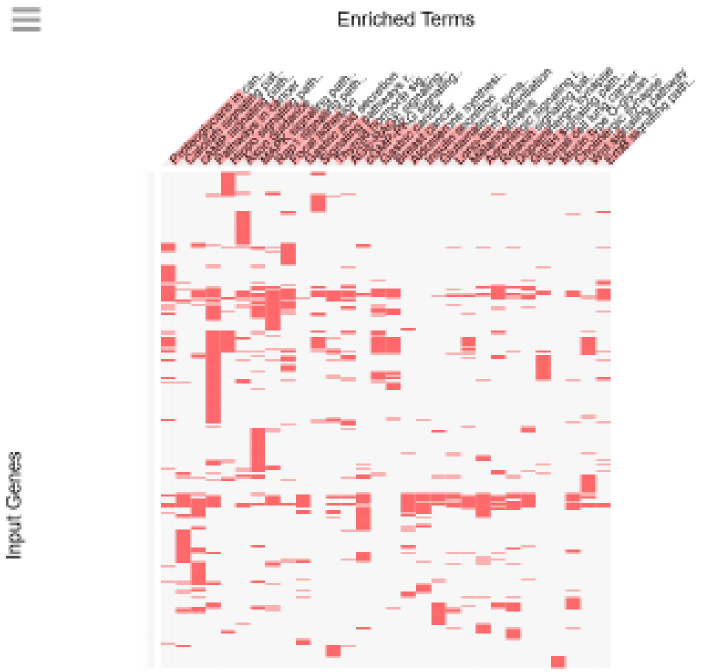

KEGG 2019 Mouse (8,405 genes): Top 35 of 106 significant records

| Term | Overlap | P-value | Adjusted P-value |
|---|---|---|---|
| Proteoglycans in cancer | 136/203 | 4.94E-13 | 1.5E-10 |
| Calcium signaling pathway | 127/189 | 2.09E-12 | 3.16E-10 |
| cAMP signaling pathway | 137/211 | 1.36E-11 | 1.38E-09 |
| Pathways in cancer | 300/535 | 2.41E-11 | 1.83E-09 |
| Hippo signaling pathway | 108/159 | 3.32E-11 | 2.01E-09 |
| Axon guidance | 119/180 | 5.22E-11 | 2.64E-09 |
| MAPK signaling pathway | 176/294 | 4.31E-10 | 1.86E-08 |
| Rap1 signaling pathway | 131/209 | 1.22E-09 | 4.62E-08 |
| Focal adhesion | 125/199 | 2.39E-09 | 8.05E-08 |
| Gastric acid secretion | 56/74 | 3.97E-09 | 1.2E-07 |
| Signaling pathways regulating pluripotency of stem cells | 90/137 | 1.8E-08 | 4.96E-07 |
| Thyroid hormone signaling pathway | 77/115 | 5.58E-08 | 1.41E-06 |
| AGE-RAGE signaling pathway in diabetic complications | 69/101 | 8.04E-08 | 1.88E-06 |
| cGMP-PKG signaling pathway | 106/172 | 1.58E-07 | 3.43E-06 |
| Gastric cancer | 94/150 | 2.59E-07 | 5.22E-06 |
| Breast cancer | 92/147 | 3.76E-07 | 7.12E-06 |
| Aldosterone synthesis and secretion | 68/102 | 4.22E-07 | 7.52E-06 |
| Insulin secretion | 59/86 | 5.53E-07 | 9.32E-06 |
| Glutamatergic synapse | 74/114 | 6.6E-07 | 1.05E-05 |
| Long-term potentiation | 48/67 | 8.6E-07 | 1.3E-05 |
| Melanogenesis | 66/100 | 1.08E-06 | 1.56E-05 |
| Adrenergic signaling in cardiomyocytes | 91/148 | 1.34E-06 | 1.84E-05 |
| PI3K-Akt signaling pathway | 194/357 | 1.52E-06 | 1.98E-05 |
| Relaxin signaling pathway | 82/131 | 1.57E-06 | 1.98E-05 |
| Dopaminergic synapse | 84/135 | 1.68E-06 | 2.04E-05 |
| GnRH signaling pathway | 60/90 | 1.99E-06 | 2.32E-05 |
| Small cell lung cancer | 61/92 | 2.17E-06 | 2.43E-05 |
| Mucin type O-glycan biosynthesis | 24/28 | 2.36E-06 | 2.56E-05 |
| Cholinergic synapse | 72/113 | 2.6E-06 | 2.72E-05 |
| Wnt signaling pathway | 96/160 | 3.27E-06 | 3.3E-05 |
| ErbB signaling pathway | 56/84 | 4.34E-06 | 4.24E-05 |
| Circadian entrainment | 64/99 | 4.53E-06 | 4.29E-05 |
| Oxytocin signaling pathway | 92/154 | 6.68E-06 | 6.13E-05 |
| Cushing syndrome | 94/159 | 9.8E-06 | 8.74E-05 |
| Gap junction | 56/86 | 1.28E-05 | 0.000111 |

**Supplemental Figure S7.** KEGG analyses of putative regulatory targets of genetic loci harboring human-specific SNCs.

**MGI Mammalian Phenotype 2017: 8,405 genes**

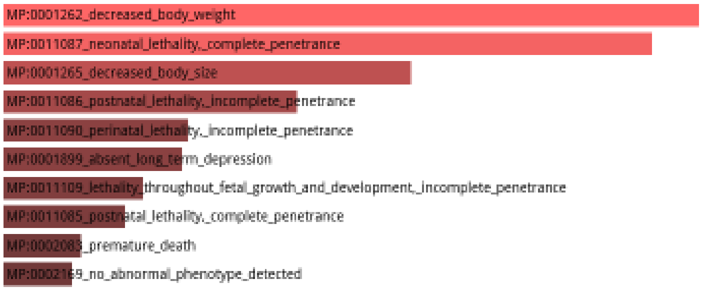


**MGI Mammalian Phenotype 2017: 8,405 genes**

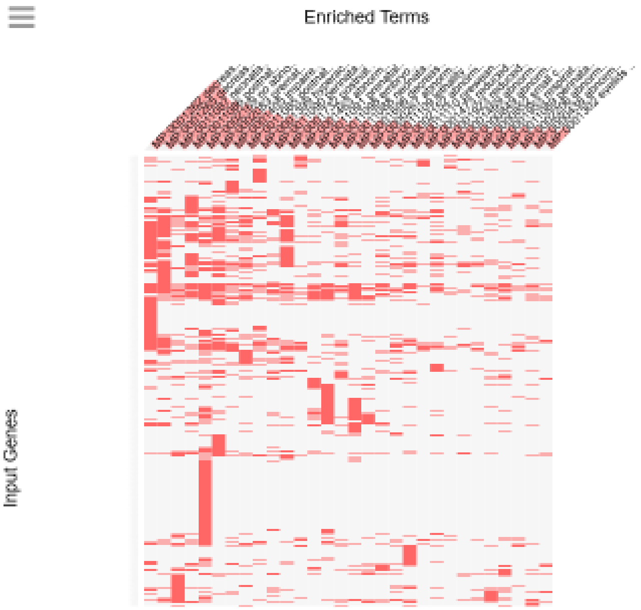

### MGI Mammalian Phenotype 2017 (8,405 genes): Top 40 of 749 significant records

| Term | Overlap | P-value | Adjusted P-value |
|---|---|---|---|
| MP:0001262_decreased_body_weight | 692/1189 | 4.71E-31 | 2.44E-27 |
| MP:0001265_decreased_body_size | 472/774 | 2.25E-27 | 5.82E-24 |
| MP:0011087_neonatal_lethality,_complete_penetrance | 299/462 | 2.47E-23 | 4.25E-20 |
| MP:0011086_postnatal_lethality,_incomplete_penetrance | 346/563 | 4.06E-21 | 5.26E-18 |
| MP:0002169_no_abnormal_phenotype_detected | 882/1674 | 2.98E-20 | 3.08E-17 |
| MP:0002083_premature_death | 474/834 | 1.12E-18 | 9.65E-16 |
| MP:0001405_impaired_coordination | 215/332 | 3.43E-17 | 2.54E-14 |
| MP:0011085_postnatal_lethality,_complete_penetrance | 238/384 | 1.57E-15 | 1.02E-12 |
| MP:0001399_hyperactivity | 216/344 | 4.36E-15 | 2.51E-12 |
| MP:0011090_perinatal_lethality,_incomplete_penetrance | 152/226 | 1.28E-14 | 6.65E-12 |
| MP:0001732_postnatal_growth_retardation | 359/590 | 1.43E-14 | 6.75E-12 |
| MP:0001463_abnormal_spatial_learning | 115/162 | 6.87E-14 | 2.96E-11 |
| MP:0011091_prenatal_lethality,_complete_penetrance | 174/272 | 1.81E-13 | 7.22E-11 |
| MP:0011098_embryonic_lethality_during_organogenesis,_complete_penetrance | 319/559 | 2.9E-13 | 1.07E-10 |
| MP:0011088_neonatal_lethality,_incomplete_penetrance | 163/255 | 1.15E-12 | 3.97E-10 |
| MP:0001698_decreased_embryo_size | 273/472 | 1.96E-12 | 6.36E-10 |
| MP:0000267_abnormal_heart_development | 109/157 | 3.11E-12 | 9.48E-10 |
| MP:0011109_lethality_throughout_fetal_growth_and_development,_incomplete_penetrance | 116/170 | 3.92E-12 | 1.13E-09 |
| MP:0002152_abnormal_brain_morphology | 104/152 | 3.9E-11 | 1.06E-08 |
| MP:0011110_preweaning_lethality,_incomplete_penetrance | 222/381 | 8.85E-11 | 2.29E-08 |
| MP:0001473_reduced_long_term_potentiation | 77/106 | 1.52E-10 | 3.75E-08 |
| MP:0001923_reduced_female_fertility | 142/227 | 3.06E-10 | 7.2E-08 |
| MP:0004811_abnormal_neuron_physiology | 67/90 | 4.06E-10 | 8.85E-08 |
| MP:0000788_abnormal_cerebral_cortex_morphology | 98/145 | 4.1E-10 | 8.85E-08 |
| MP:0001406_abnormal_gait | 180/302 | 4.48E-10 | 9.28E-08 |
| MP:0001469_abnormal_contextual_conditioning_behavior | 45/54 | 4.96E-10 | 9.82E-08 |
| MP:0000849_abnormal_cerebellum_morphology | 68/92 | 5.12E-10 | 9.82E-08 |
| MP:0000266_abnormal_heart_morphology | 139/224 | 1.03E-09 | 1.91E-07 |
| MP:0000852_small_cerebellum | 52/66 | 1.15E-09 | 2.05E-07 |
| MP:0001953_respiratory_failure | 98/147 | 1.29E-09 | 2.23E-07 |
| MP:0011089_perinatal_lethality,_complete_penetrance | 135/217 | 1.41E-09 | 2.35E-07 |
| MP:0002206_abnormal_CNS_synaptic_transmission | 66/90 | 1.6E-09 | 2.59E-07 |
| MP:0000438_abnormal_cranium_morphology | 90/133 | 1.91E-09 | 2.99E-07 |
| MP:0001302_eyelids_open_at_birth | 43/52 | 1.97E-09 | 3E-07 |
| MP:0000807_abnormal_hippocampus_morphology | 63/86 | 4.04E-09 | 5.97E-07 |
| MP:0006009_abnormal_neuronal_migration | 57/76 | 5.14E-09 | 7.28E-07 |
| MP:0001954_respiratory_distress | 111/174 | 5.2E-09 | 7.28E-07 |
| MP:0001899_absent_long_term_depression | 25/26 | 5.88E-09 | 7.95E-07 |
| MP:0002906_increased_susceptibility_to_pharmacologically_induced_seizures | 65/90 | 5.99E-09 | 7.95E-07 |
| MP:0003633_abnormal_nervous_system_physiology | 74/106 | 6.33E-09 | 8.12E-07 |

### MGI Mammalian Phenotype Level 4 2019: 8.405 genes

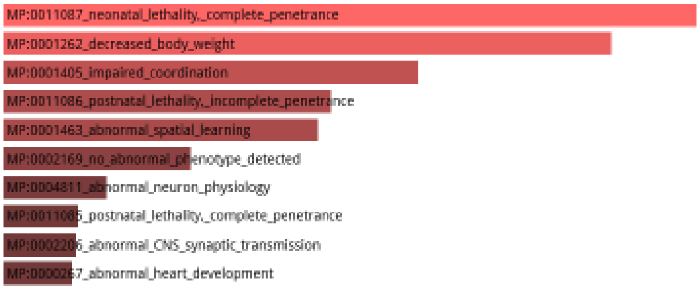

### MGI Mammalian Phenotype Level 4 2019: 8.405 genes

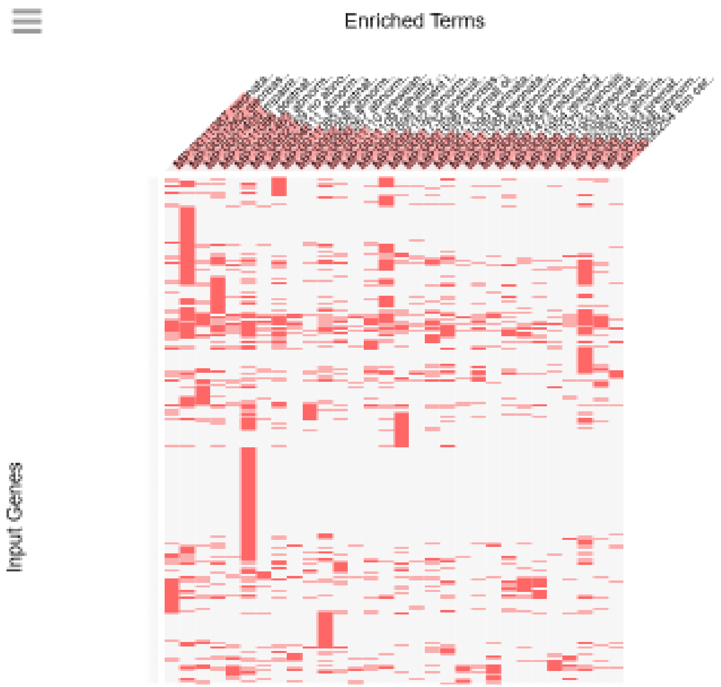


### MGI Mammalian Phenotype Level 4 2019 (8,405 genes): top 40 of 407 significant records

| Term | Overlap | P-value | Adjusted P-value |
|---|---|---|---|
| MP:0011087_neonatal_lethality,_complete_penetrance | 315/517 | 1.55E-18 | 8.15E-15 |
| MP:0001262_decreased_body_weight | 773/1471 | 2.04E-17 | 5.36E-14 |
| MP:0001405_impaired_coordination | 247/405 | 6.90E-15 | 1.21E-11 |
| MP:0011086_postnatal_lethality,_incomplete_penetrance | 362/643 | 9.65E-14 | 1.27E-10 |
| MP:0001463_abnormal_spatial_learning | 120/172 | 1.46E-13 | 1.54E-10 |
| MP:0002169_no_abnormal_phenotype_detected | 958/1944 | 6.87E-12 | 6.02E-09 |
| MP:0004811_abnormal_neuron_physiology | 78/107 | 8.69E-11 | 6.53E-08 |
| MP:0002206_abnormal_CNS_synaptic_transmission | 75/103 | 2.18E-10 | 1.27E-07 |
| MP:0000267_abnormal_heart_development | 111/168 | 2.43E-10 | 1.28E-07 |
| MP:0011085_postnatal_lethality,_complete_penetrance | 246/432 | 2.04E-10 | 1.34E-07 |
| MP:0011110_preweaning_lethality,_incomplete_penetrance | 360/669 | 2.90E-10 | 1.39E-07 |
| MP:0011109_lethality_throughout_fetal_growth_and_development,_incomplete_penetrance | 122/191 | 8.21E-10 | 3.60E-07 |
| MP:0001899_absent_long_term_depression | 27/28 | 1.11E-09 | 4.50E-07 |
| MP:0001732_postnatal_growth_retardation | 360/677 | 1.85E-09 | 6.48E-07 |
| MP:0001698_decreased_embryo_size | 293/537 | 2.09E-09 | 6.88E-07 |
| MP:0002152_abnormal_brain_morphology | 119/187 | 1.84E-09 | 6.92E-07 |
| MP:0011090_perinatal_lethality,_incomplete_penetrance | 154/256 | 3.23E-09 | 9.44E-07 |
| MP:0002741_small_olfactory_bulb | 35/40 | 3.07E-09 | 9.51E-07 |
| MP:0001469_abnormal_contextual_conditioning_behavior | 48/61 | 5.45E-09 | 1.43E-06 |
| MP:0001473_reduced_long_term_potentiation | 84/124 | 6.05E-09 | 1.45E-06 |
| MP:0000788_abnormal_cerebral_cortex_morphology | 104/161 | 5.79E-09 | 1.45E-06 |
| MP:0011088_neonatal_lethality,_incomplete_penetrance | 172/293 | 5.33E-09 | 1.48E-06 |
| MP:0001954_respiratory_distress | 121/194 | 7.90E-09 | 1.81E-06 |
| MP:0001575_cyanosis | 129/210 | 1.00E-08 | 2.19E-06 |
| MP:0001953_respiratory_failure | 103/161 | 1.47E-08 | 3.09E-06 |
| MP:0002906_increased_susceptibility_to_pharmacologically_induced_seizures | 70/101 | 2.60E-08 | 5.25E-06 |
| MP:0002910_abnormal_excitatory_postsynaptic_currents | 59/83 | 7.68E-08 | 1.50E-05 |
| MP:0002083_premature_death | 499/997 | 9.80E-08 | 1.84E-05 |
| MP:0002066_abnormal_motor_capabilities/coordination/movement | 102/164 | 1.41E-07 | 2.55E-05 |
| MP:0006254_thin_cerebral_cortex | 53/74 | 2.36E-07 | 4.14E-05 |
| MP:0011098_embryonic_lethality_during_organogenesis,_complete_penetrance | 339/656 | 2.60E-07 | 4.28E-05 |
| MP:0000807_abnormal_hippocampus_morphology | 65/92 | 2.56E-07 | 4.34E-05 |
| MP:0006009_abnormal_neuronal_migration | 59/85 | 2.94E-07 | 4.69E-05 |
| MP:0011108_embryonic_lethality_during_organogenesis,_incomplete_penetrance | 143/247 | 3.20E-07 | 4.95E-05 |
| MP:0000031_abnormal_cochlea_morphology | 44/59 | 3.84E-07 | 5.77E-05 |
| MP:0000852_small_cerebellum | 58/84 | 4.91E-07 | 7.17E-05 |
| MP:0002063_abnormal_learning/memory/conditioning | 42/56 | 5.43E-07 | 7.52E-05 |
| MP:0003633_abnormal_nervous_system_physiology | 79/123 | 5.36E-07 | 7.62E-05 |
| MP:0010025_decreased_total_body_fat_amount | 263/498 | 5.94E-07 | 8.01E-05 |
| MP:0009937_abnormal_neuron_differentiation | 75/116 | 7.01E-07 | 9.22E-05 |

**Supplemental Figure S8.** Interrogation of MGI Mammalian Phenotype databases identifies genes associated with human-specific SNCs and implicated in premature death and embryonic, perinatal, neonatal, and postnatal lethality phenotypes.

**Association with networks of human-specific regulatory sequences (HSGRS) and stem cell-associated retroviral sequences (SCARS) of 8,405 genes associated with 35,074 fixed human-specific single nucleotide changes located in differentially-accessible chromatin regions during human neurogenesis in cerebral organoids**

| Classification category | Number of genes | Perent |
|---|---|---|
| Unique genes | 8405 | 100.00 |
| In network of human-specific genomic regulatory sequences (HSGRS) | 7406 | 88.11 |
| LTR5_Hs/SVA_D enhancers-regulated genes | 1387 | 16.50 |
| HERVH lncRNA-regulated genes | 3191 | 37.97 |
| LTR7Y/B enhancers-regulated genes | 3306 | 39.33 |
| In network of stem cell-associated retroviral sequences (SCARS) | 5389 | 64.12 |
| Both HSRGS & SCARS-regulated genes | 4805 | 57.17 |
| All HSGRS & SCARS-regulated genes | 7990 | 95.06 |

**Effect of CRISPR-guided epigentic silencing of LRT5_Hs/SVA_D enhancers on expression of 1,387 genes associated with human-specific SNC in DA chromatin regions**

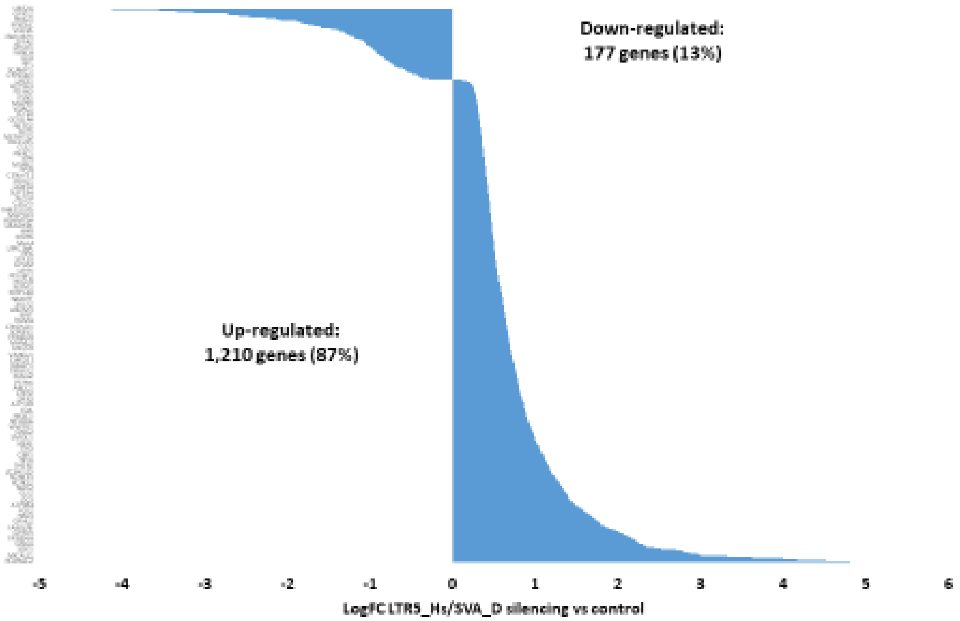

**Effects of stem cell-associated retroviral sequences (SCARS) on expression of 5,389 genes associated with human-specific neuro-regulatory SNC located in DA chromatin regions during brain development in cerebral organoids**

| Classification category | Number of genes | Down-regulated | Percent | Up-regulated | Percent |
|---|---|---|---|---|---|
| LTR5_Hs/SVA_D enhancers-regulated genes | 1387 | 1210 | 87.24 | 177 | 12.76 |
| HERVH lncRNA-regulated genes | 3191 | 1733 | 54.31 | 1458 | 45.69 |
| LTR7Y/B enhancers-regulated genes | 3306 | 2494 | 75.44 | 812 | 24.56 |

**Supplemental Figure S9.** Structurally, functionally, and evolutionary distinct classes of HSRS share the relatively restricted elite set of common genetic targets.
