## Supplemental Note for "Impacts of genomic networks governed by human-specific regulatory sequences and genetic loci harboring fixed human-specific neuro-regulatory single nucleotide mutations on phenotypic traits of Modern Humans"

The principal element of our analytical approach is to define specific sets of genomic regulatory loci
WITHOUT ANY PRIOR KNOWLEDGE
of what genes (if any) they may (or may not) control.

This is in striking contrast with the approaches that being utilized in the prior art: to identify genes that are differentially regulated between different states and/or conditions, thus introducing the confounders and biases associated with the multiple hypothesis testing, overfitting, and random co-occurrence due to the very large excess of analyzed features (genes) over the relatively small numbers of samples (100K range or more of analyzed features versus just a few hundred analyzed samples at best). Attempts to address these issues using statistical approaches (corrections for multiple hypothesis testing) and experimental design improvements (multiple replications, validation in independent data sets and using independent analytical techniques) did not adequately resolve these problems.

Importantly, our analyses revealed that this approach identifies not only genes altered expression of which implicated in the broad spectrum of human physiological and pathological conditions. It also identifies genes defined by the independent studies as genetic loci comprising mutation signatures associated with development and progression of multiple common human disorders, including cancer, neurodevelopmental, neuropsychiatric, and neurodegenerative disorders, as well autoimmune and immuno-inflammatory diseases.

Supporting the concept that many major human disorders are driven by aberrant functions of primate-specific genomic regulatory networks with prominent unique-to-human (human-specific) components, this approach identifies genetic loci implicated in inter-individual genetic mosaicism (somatic mosaicism) of cells, tissues and organs in the human body.

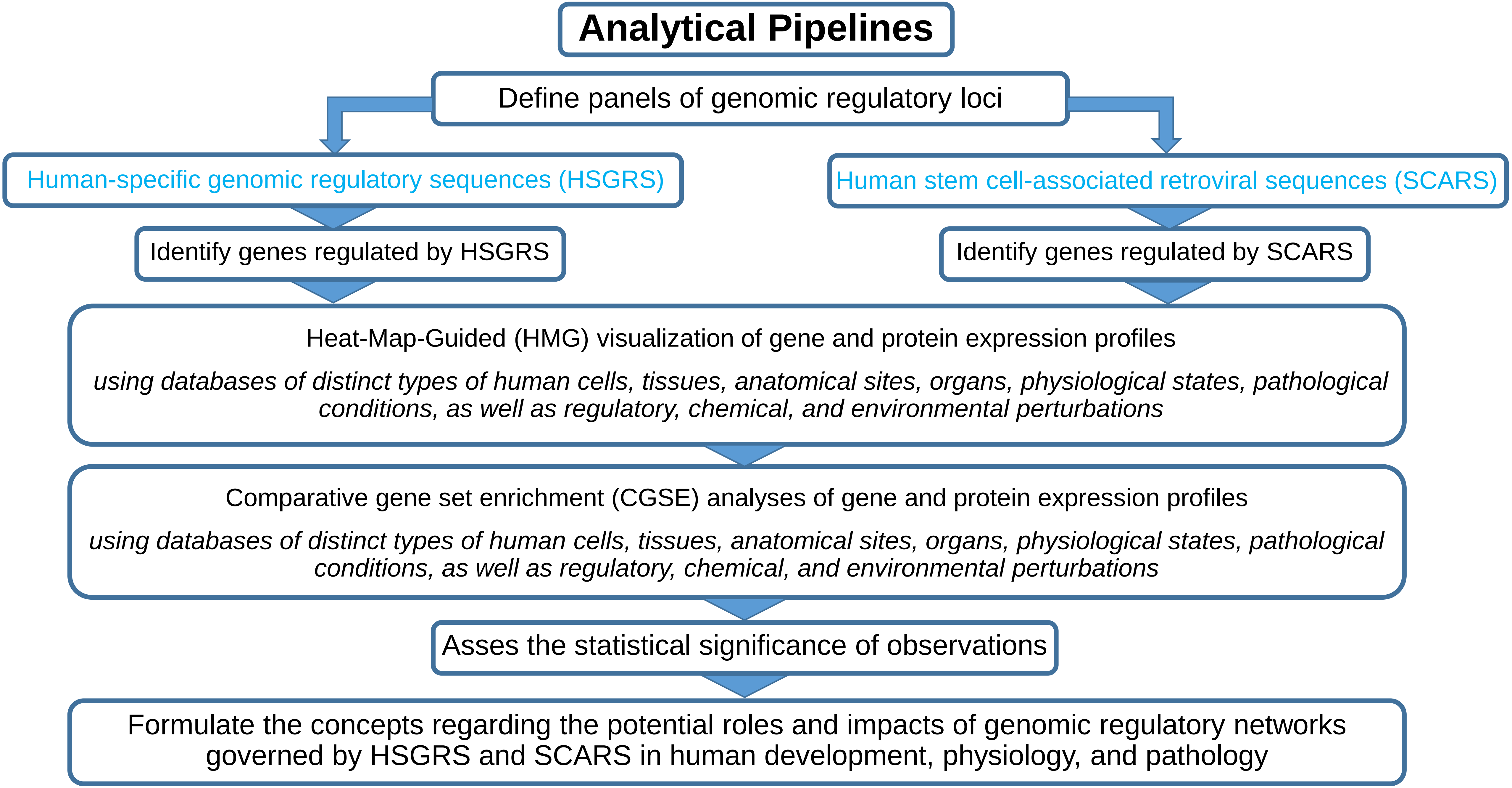
Analytical Pipelines
Define panels of genomic regulatory loci
Human-specific genomic regulatory sequences (HSGRS)
Human stem cell-associated retroviral sequences (SCARS)
Identify genes regulated by HSGRS
Identify genes regulated by SCARS
Heat-Map-Guided (HMG) visualization of gene and protein expression profiles
using databases of distinct types of human cells, tissues, anatomical sites, organs, physiological states, pathological conditions, as well as regulatory, chemical, and environmental perturbations
Comparative gene set enrichment (CGSE) analyses of gene and protein expression profiles
using databases of distinct types of human cells, tissues, anatomical sites, organs, physiological states, pathological conditions, as well as regulatory, chemical, and environmental perturbations
Asses the statistical significance of observations
Formulate the concepts regarding the potential roles and impacts of genomic regulatory networks governed by HSGRS and SCARS in human development, physiology, and pathology

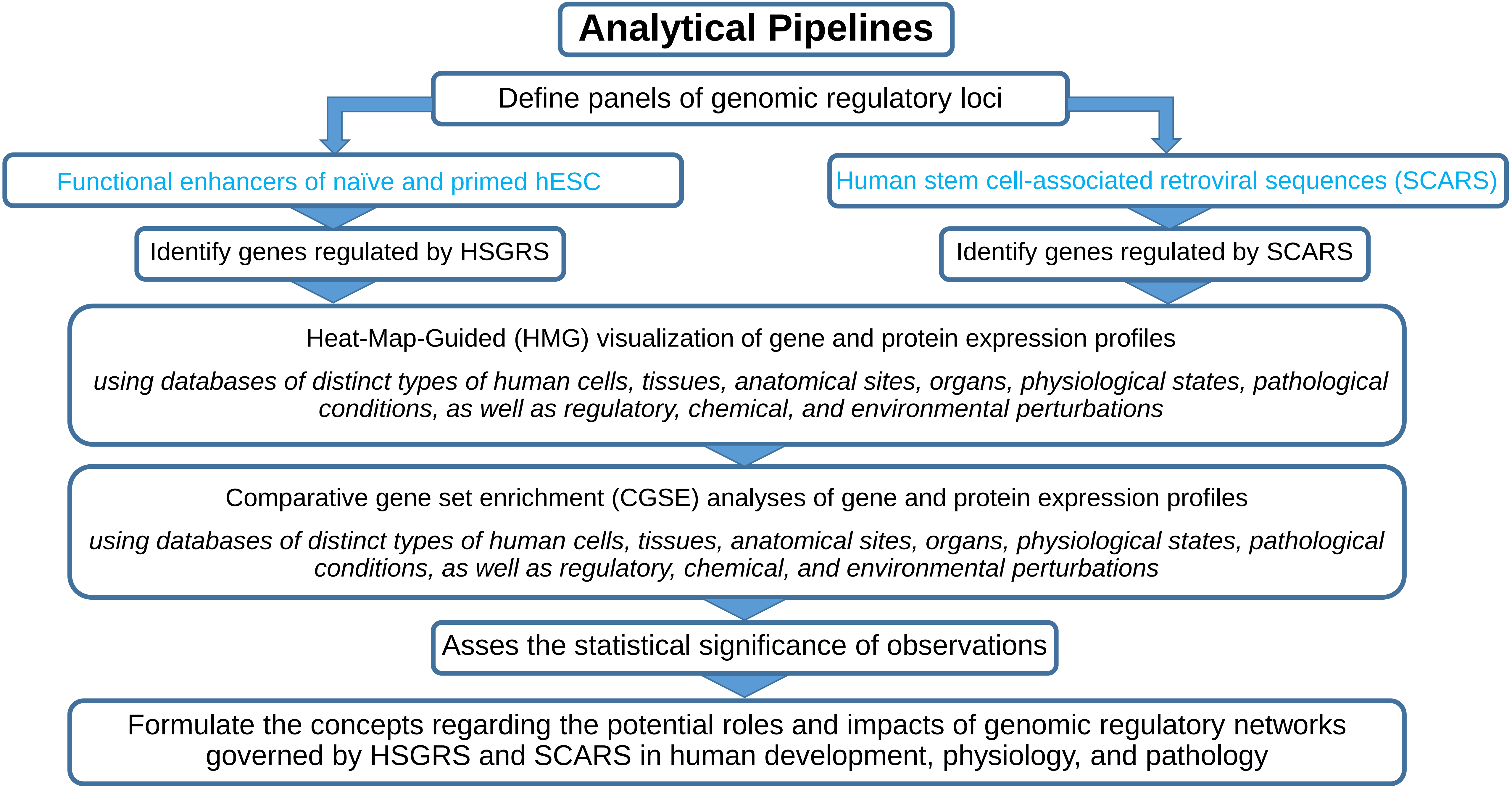
Analytical Pipelines
Define panels of genomic regulatory loci
Functional enhancers of naïve and primed hESC
Human stem cell-associated retroviral sequences (SCARS)
Identify genes regulated by HSGRS
Identify genes regulated by SCARS
Heat-Map-Guided (HMG) visualization of gene and protein expression profiles
using databases of distinct types of human cells, tissues, anatomical sites, organs, physiological states, pathological conditions, as well as regulatory, chemical, and environmental perturbations
Comparative gene set enrichment (CGSE) analyses of gene and protein expression profiles
using databases of distinct types of human cells, tissues, anatomical sites, organs, physiological states, pathological conditions, as well as regulatory, chemical, and environmental perturbations
Asses the statistical significance of observations
Formulate the concepts regarding the potential roles and impacts of genomic regulatory networks governed by HSGRS and SCARS in human development, physiology, and pathology

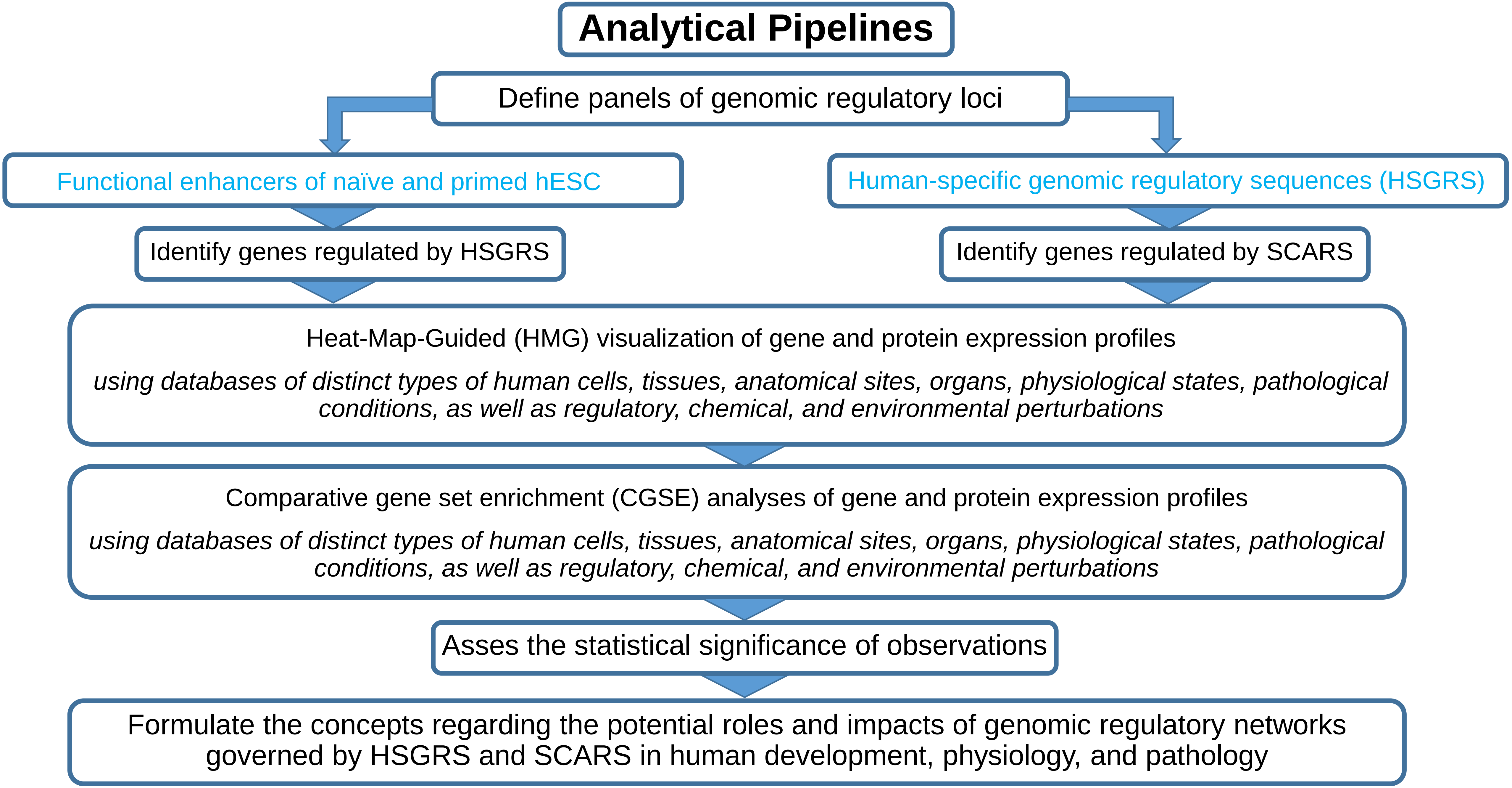
Analytical Pipelines
Define panels of genomic regulatory loci
Functional enhancers of naïve and primed hESC
Human-specific genomic regulatory sequences (HSGRS)
Identify genes regulated by HSGRS
Identify genes regulated by SCARS
Heat-Map-Guided (HMG) visualization of gene and protein expression profiles
using databases of distinct types of human cells, tissues, anatomical sites, organs, physiological states, pathological conditions, as well as regulatory, chemical, and environmental perturbations
Comparative gene set enrichment (CGSE) analyses of gene and protein expression profiles
using databases of distinct types of human cells, tissues, anatomical sites, organs, physiological states, pathological conditions, as well as regulatory, chemical, and environmental perturbations
Asses the statistical significance of observations
Formulate the concepts regarding the potential roles and impacts of genomic regulatory networks governed by HSGRS and SCARS in human development, physiology, and pathology

- **MERGE THE RESULTS OF THE ANALYSES TO EXPLORE THE CONTRIBUTION OF SCARS TO FUNCTIONS OF HUMAN-SPECIFIC GENOMIC REGULATORY NETWORKS IN HUMAN DEVELOPMENT AS WELL AS HEALTH AND DISEASE STATES**

- **IDENTIFY THE HIGH-VALUE GENETIC TARGETS FOR MECHANISTIC AND FUNCTIONAL STUDIES IN THE EXPERIMENTAL MODELS OF HUMAN NEUROGENSIS AND CORTICOGENESIS**

### 59K human-specific regulatory sequences

Analysis of genes representing the putative regulatory targets of human-specific regulatory sequences

**Structurally, functionally, and evolutionary distinct families of human-specific regulatory sequences (HSRS) and associated putative regulatory target genes defined by the GREAT algorithm.**

| Classification category/Reference database | Number of records (hg19) | Associated genes |
|---|---|---|
| Fixed human-specific insertions. | 11878 | 7979 |
| Human-specific TE loci expressed in human dorsolateral prefrontal cortex | 4637 | 4051 |
| Set of duplicated regions in GRCh38 space | 7599 | 6618 |
| Fixed human-specific deletions | 5883 | 5489 |
| Human-specific STR expansions | 4875 | 4844 |
| hsTFBS | 3803 | 1087 |
| ace-DHS | 3538 | 3445 |
| FHSRR | 4249 | 2810 |
| Human-specific STR contractions | 1279 | 973 |
| hESC_FHSRR_DHS | 1932 | 1458 |
| DHS_FHSRR (non-hESC) | 2118 | 552 |
| HARs | 2745 | 2281 |
| haDHS | 524 | 747 |
| Human-biased CNCC enhances | 1000 | 1439 |
| Chimp-biased CNCC enhances | 1000 | 1445 |
| H3K4me3 peaks with human-specific enrichment in prefrontal neurons | 410 | 578 |
| Human-specific hESC functional enhancers | 1619 | 1214 |
| All HSRS | 59089 | 13824 |

Legend: Definitions of structurally, functionally, and evolutionary distinct families of human-specific regulatory sequences (HSRS) can be found in Glinsky (2020);

Figure 1. Heat-Map-Guided Visualization (HMGV) analysis of expression profiles of mRNAs encoded by 90 genes associated with human-specific genomic regulatory loci derived from transcriptionally active in human dorsolateral prefrontal cortex (DLPFC) transposable elements (A; B; E) and mRNAs encoded by 326 genes associated with human-specific genomic regulatory loci derived from fixed human-specific insertions (C; D; F). Results of the HMGV analysis employing the following databases are shown: HPA Human Tissues' Gene Expression Profiles (A; C); GTEx Human Tissues' Gene Expression Profiles (B; D); BioGPS Human Cell Type and Tissue Gene Expression Profiles (E; F). In (A; B; C; E; F), tissues (columns) are sorted by the rank order function from left to right; genes (rows) are sorted by the cluster order function from top to bottom. In (D), both columns and rows are sorted by the cluster order function.

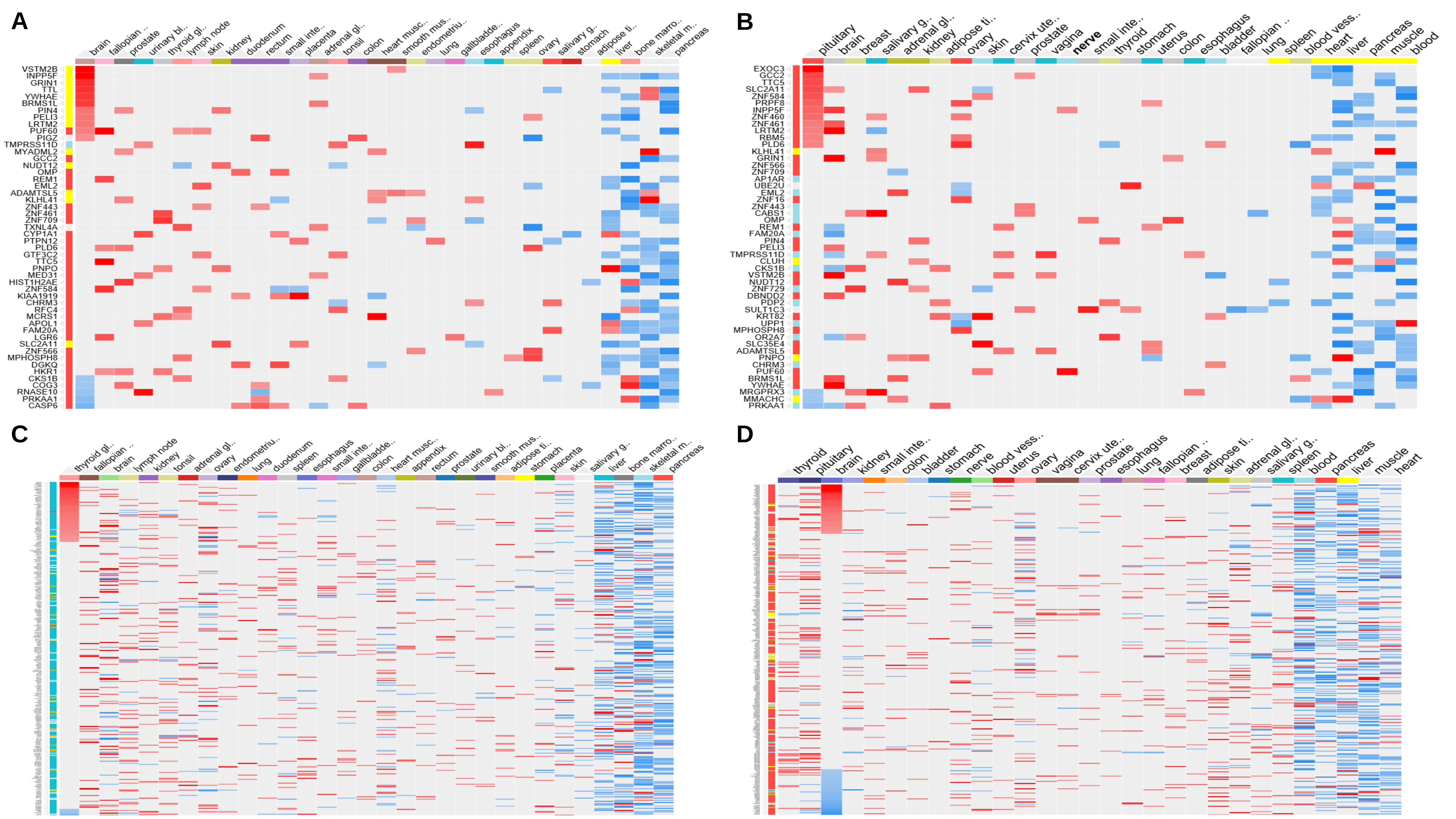
A
B
C
D
brain
fallopian ..
prostate
urinary bl..
thyroid gl..
lymph node
skin
kidney
duodenum
rectum
small inte..
placenta
adrenal gl..
tonsil
colon
heart musc..
smooth mus..
endometriu..
lung
gallbladde..
esophagus
appendix
spleen
ovary
salivary g..
stomach
adipose ti..
liver
bone marro..
skeletal m..
pancreas
VSTM2B
INPP5F
GRIN1
TTL
YWHAE
BRMS1L
PIN4
PELI3
LRTM2
PUF60
PIGZ
TMPRSS11D
MYADML2
GCC2
NUDT12
OMP
REM1
EML2
ADAMTSL5
KLHL41
ZNF443
ZNF461
ZNF709
TXNL4A
CYP1A1
PTPN12
PLD6
GTF3C2
TTC5
PNPO
MED31
HIST1H2AE
ZNF584
KIAA1919
CHRM3
RFC4
MCRS1
APOL1
FAM20A
LGR6
SLC2A11
ZNF566
MPHOSPH8
DGKQ
HKR1
CKS1B
COG3
RNASE10
PRKAA1
CASP6
pituitary
brain
breast
salivary g..
adrenal gl..
kidney
adipose ti..
ovary
skin
cervix ute..
prostate
vagina
nerve
small inte..
thyroid
stomach
uterus
colon
esophagus
bladder
fallopian ..
lung
spleen
blood vess..
heart
liver
pancreas
muscle
blood
EXOC3
GCC2
TTC5
SLC2A11
ZNF584
PRPF8
INPP5F
ZNF460
ZNF461
LRTM2
RBM5
PLD6
KLHL41
GRIN1
ZNF566
ZNF709
AP1AR
UBE2U
EML2
ZNF16
ZNF443
CABS1
OMP
REM1
FAM20A
PIN4
PELI3
TMPRSS11D
CLUH
CKS1B
VSTM2B
NUDT12
ZNF729
DBNDD2
PDP2
SULT1C3
KRT82
UPP1
MPHOSPH8
OR2A7
SLC35E4
ADAMTSL5
PNPO
CHRM3
PUF60
BRMS1L
YWHAE
MRGPRX3
MMACHC
PRKAA1
thyroid gl..
fallopian ..
brain
lymph node
kidney
tonsil
adrenal gl..
ovary
endometriu..
lung
duodenum
spleen
esophagus
small inte..
gallbladde..
colon
heart musc..
appendix
rectum
prostate
urinary bl..
smooth mus..
adipose ti..
stomach
placenta
skin
salivary g..
liver
bone marro..
skeletal m..
pancreas
thyroid
pituitary
brain
kidney
small inte..
colon
bladder
stomach
nerve
blood vess..
uterus
ovary
vagina
cervix ute..
prostate
esophagus
lung
fallopian ..
breast
adipose ti..
skin
adrenal gl..
salivary g..
spleen
blood
pancreas
liver
muscle
heart

E

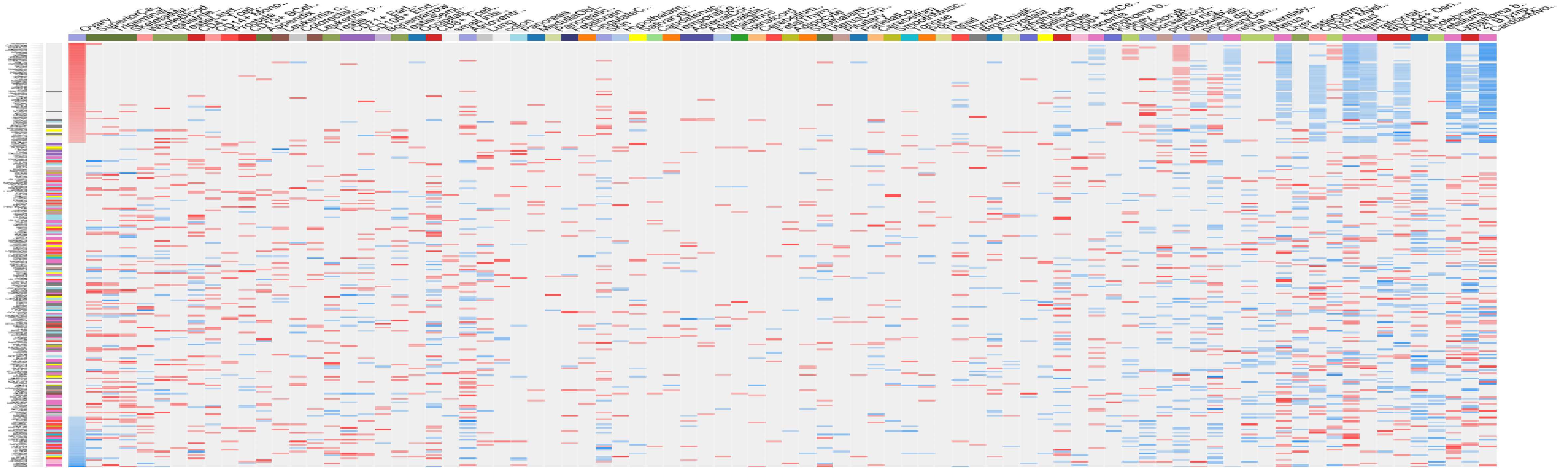

F

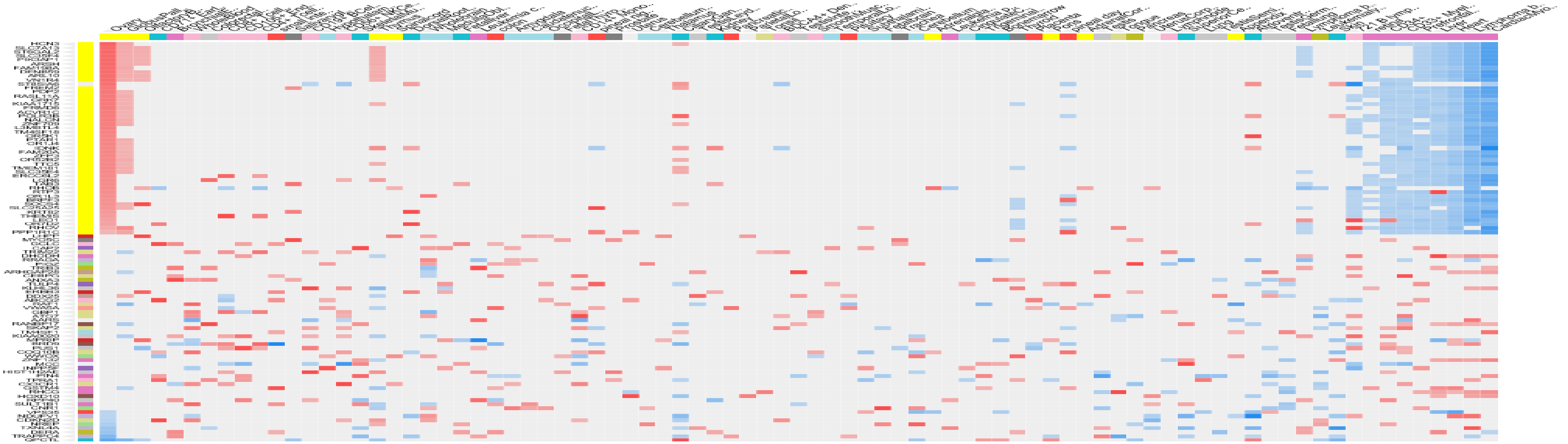

HeatMaps Visualization of expression profiles of mRNAs encoded by 90 genes associated with human-specific genomic regulatory loci derived from transcriptionally active in human DLPFC transposable elements

### HPA Human Tissues' Gene Expression Profiles

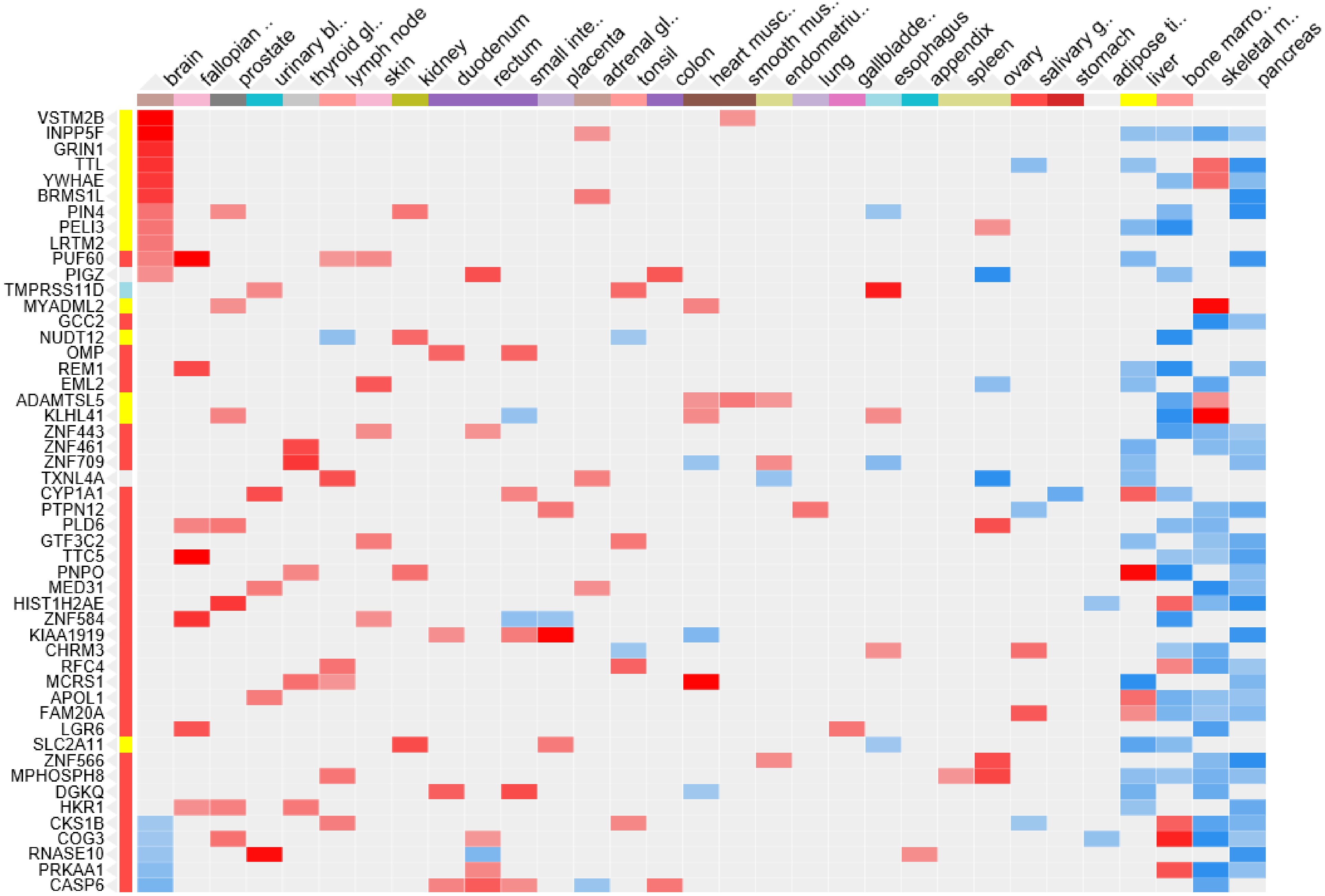


hsTE_DLPFC_1kb
90 genes
top 50 targets

### GTEx Human Tissues' Gene Expression Profiles

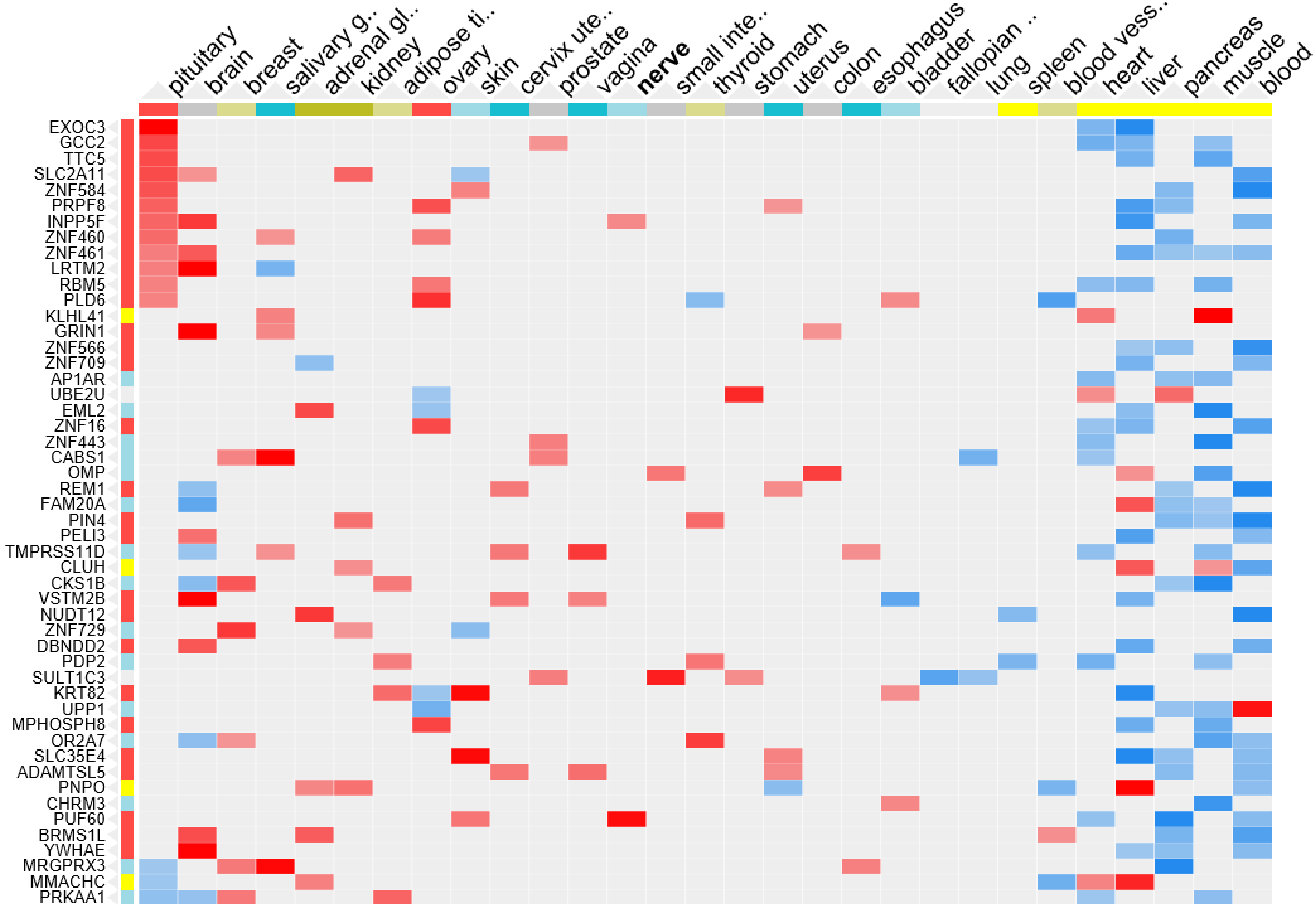


hsTE_DLPFC_1kb
90 genes
top 50 targets

### HeatMaps Visualization of expression profiles of mRNAs encoded by 326 genes associated with human-specific genomic regulatory loci derived from fixed human-specific insertions

### HPA Human Tissues' Gene Expression Profiles

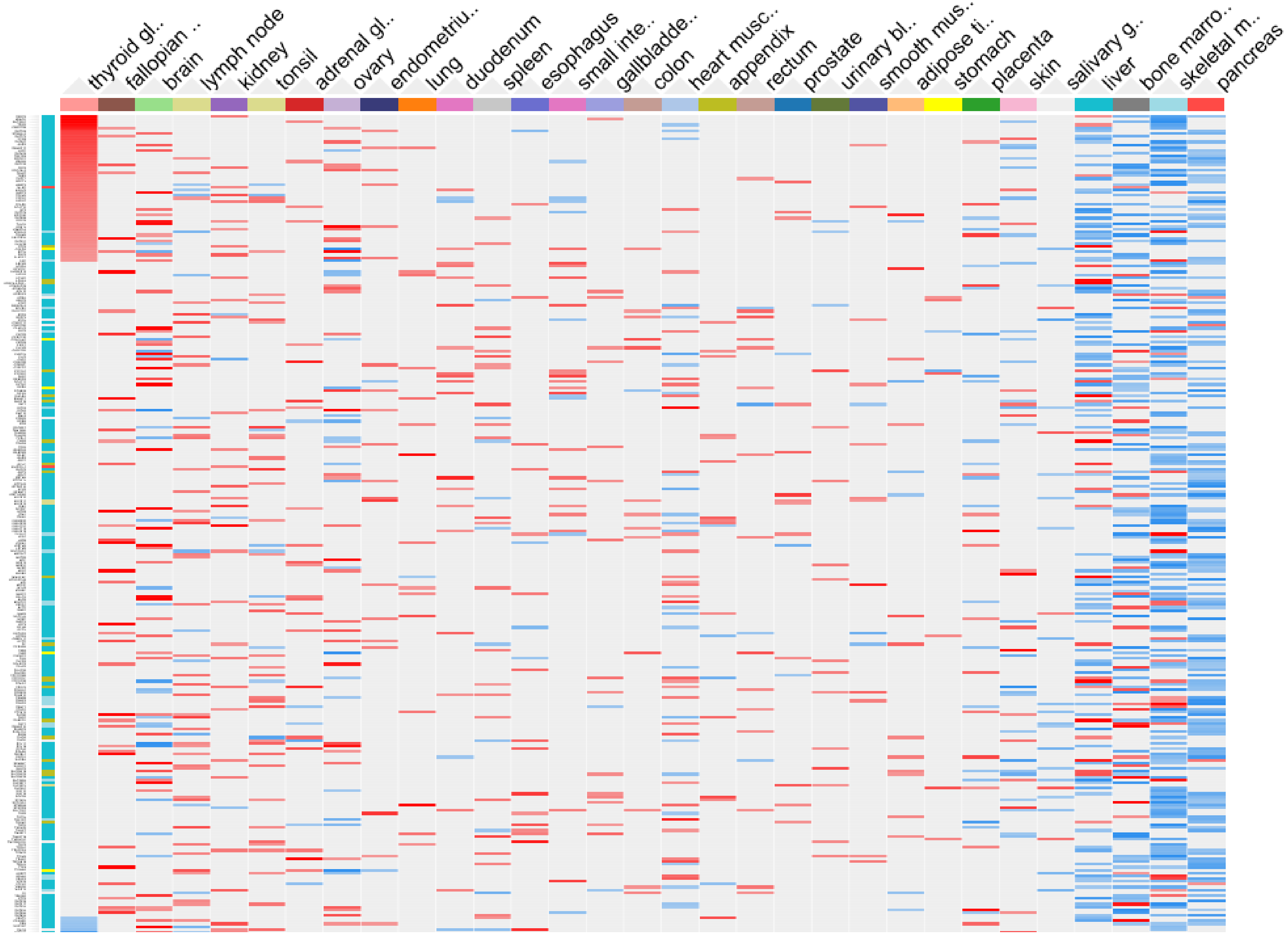


Fixed_hs_Insertions
1kb_326 genes
All targets

### GTEx Human Tissues' Gene Expression Profiles

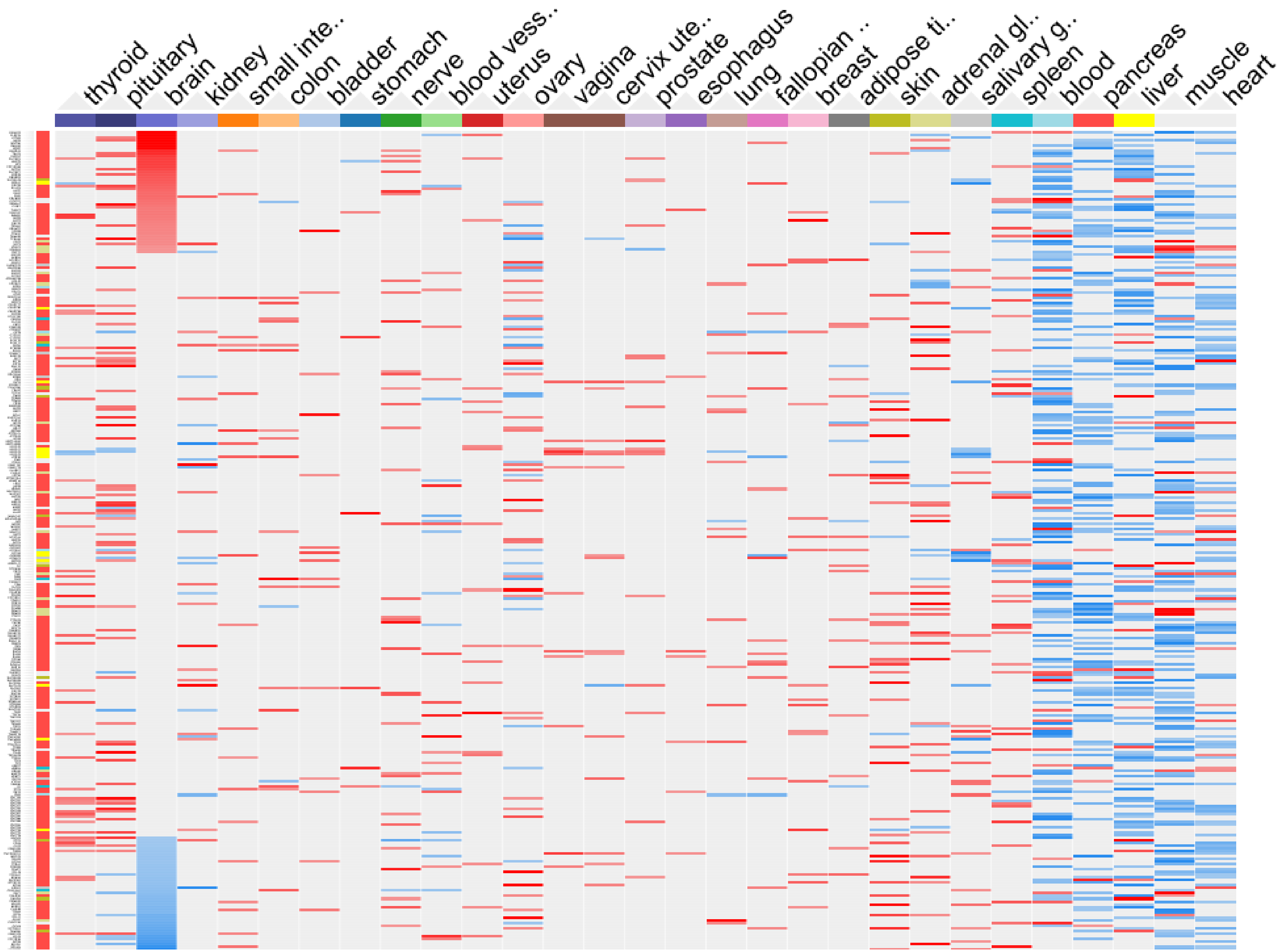


Fixed_hs_Insertions
1kb_326 genes
All targets

### GTEx Human Tissues' Gene Expression Profiles

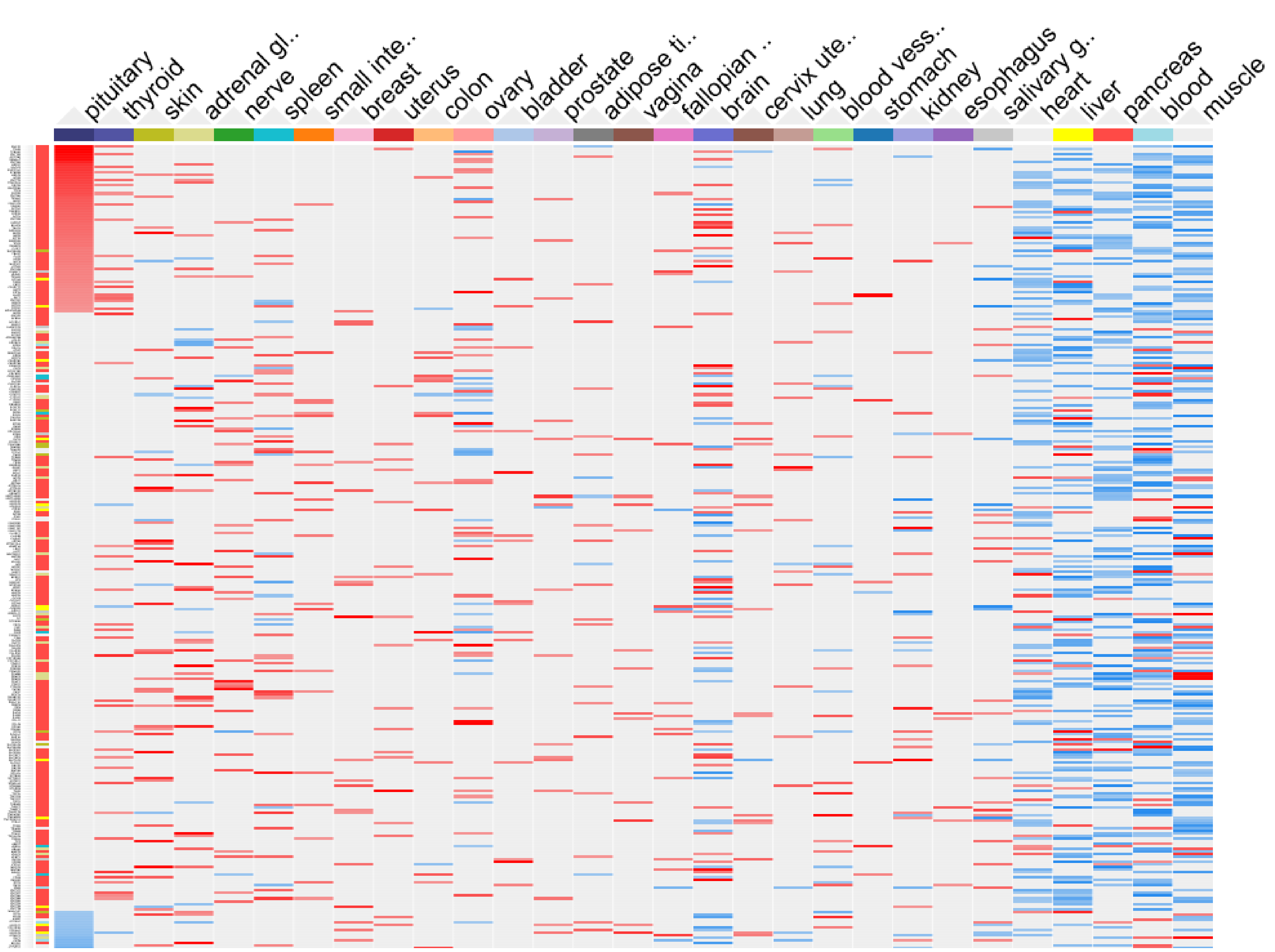


Fixed_hs_Insertions
1kb_326 genes
All targets

### BioGPS Human Cell Type and Tissue Gene Expression Profiles

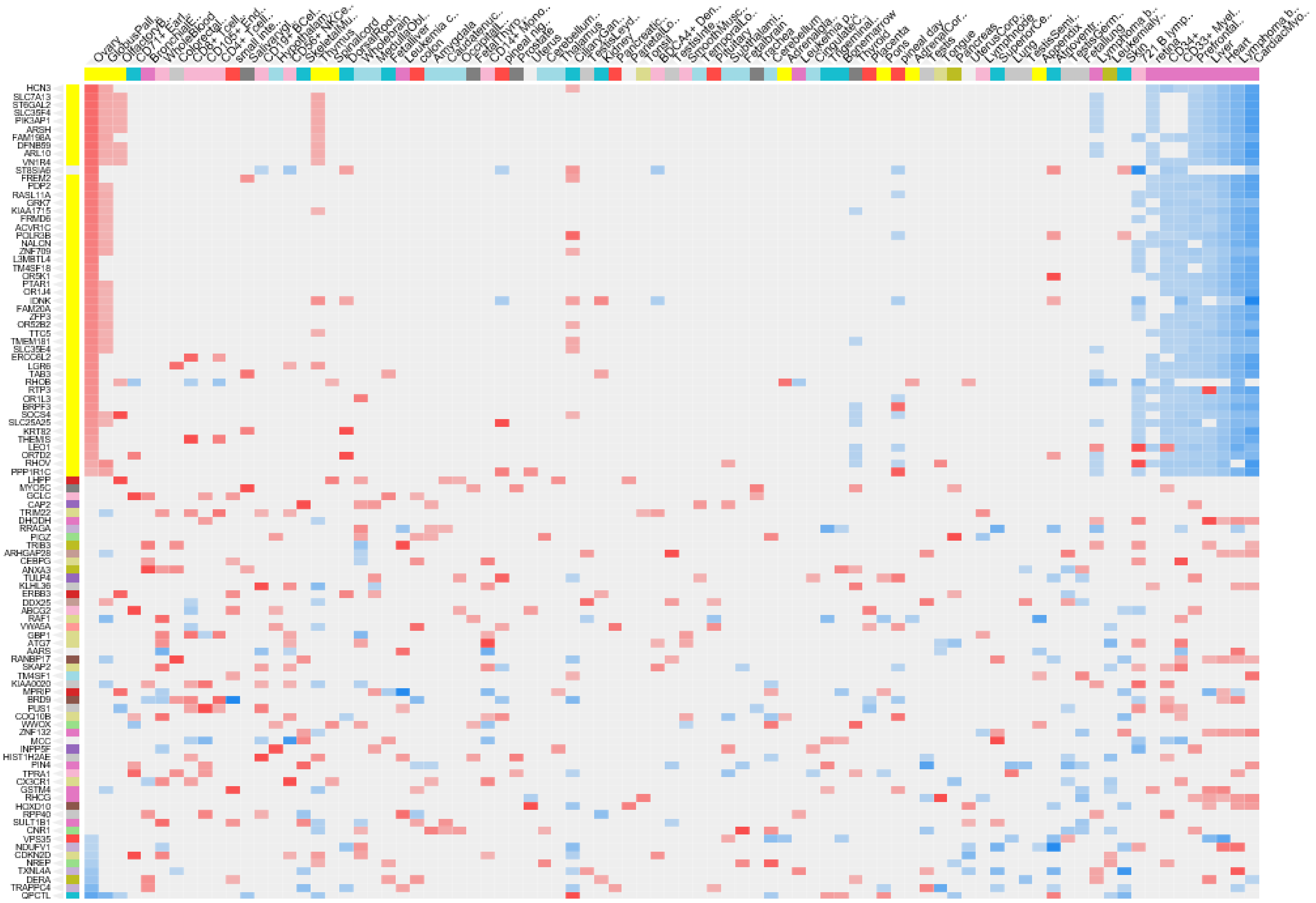

Fixed_hs_Insertions
1kb_326 genes
Top 100 targets

### BioGPS Human Cell Type and Tissue Gene Expression Profiles

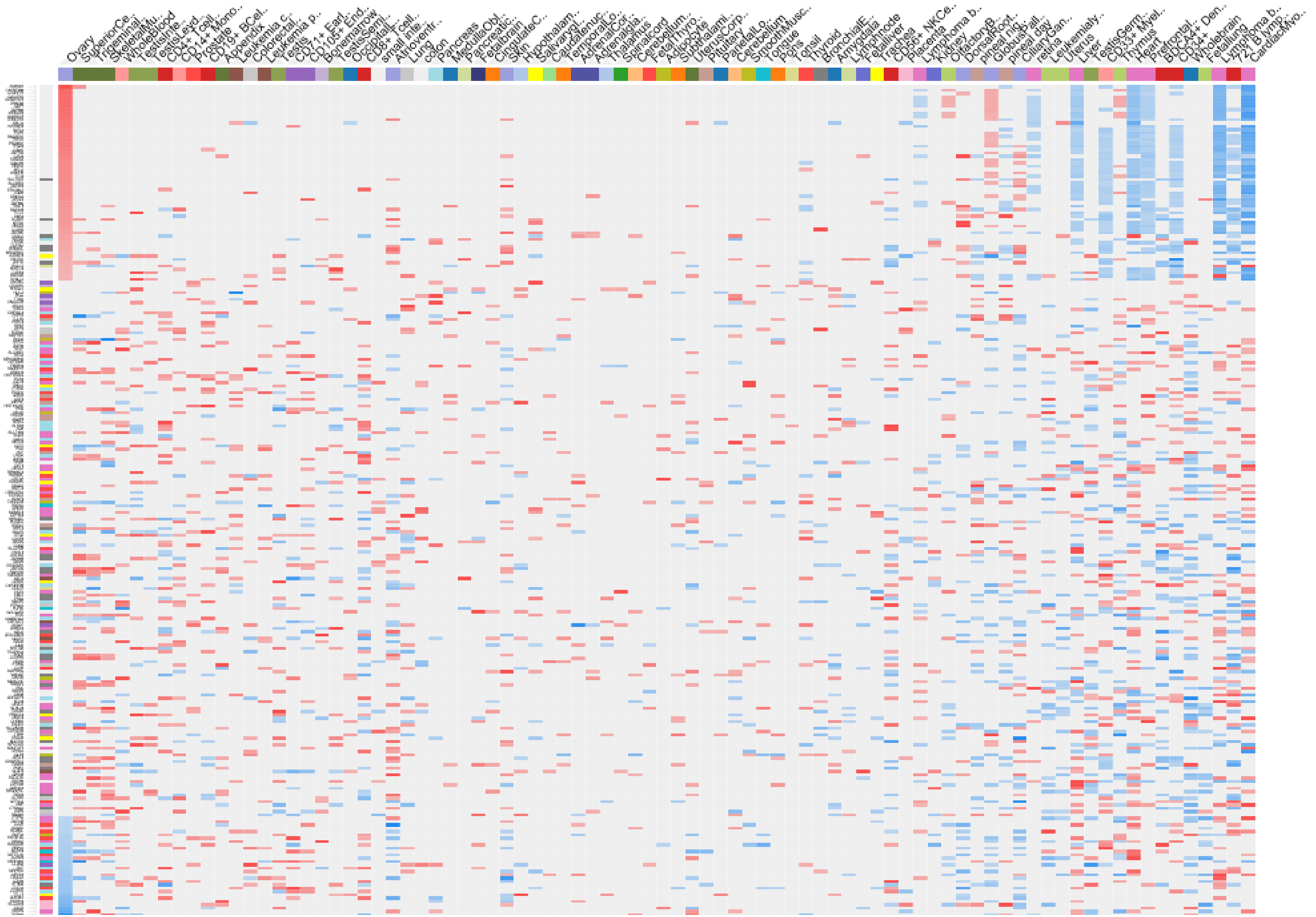

hsTE_DLPFC_10kb
336 genes
top 250 targets

**11,878 fixed human-specific insertions**

### 11,878 fixed human-specific insertions

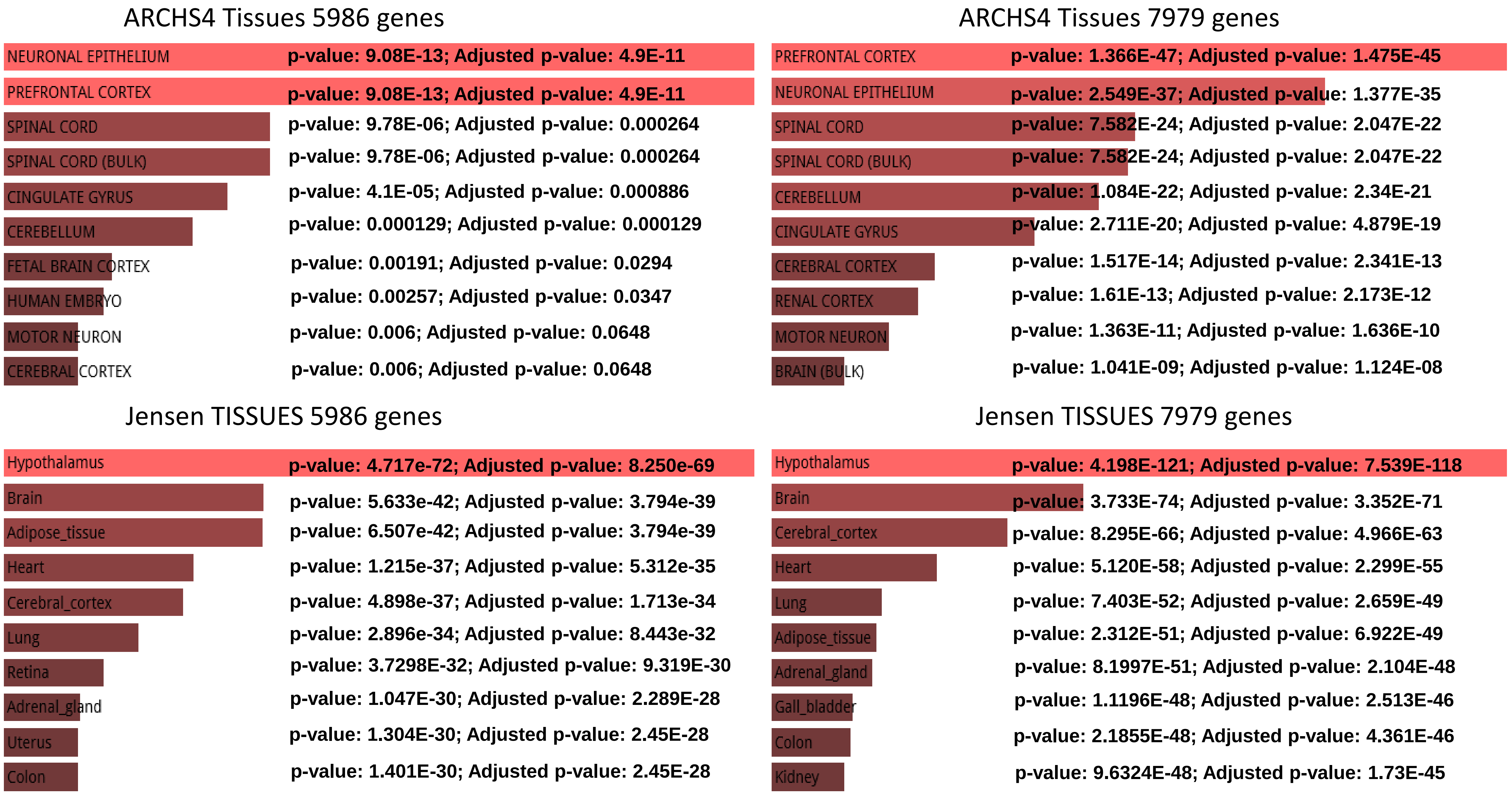

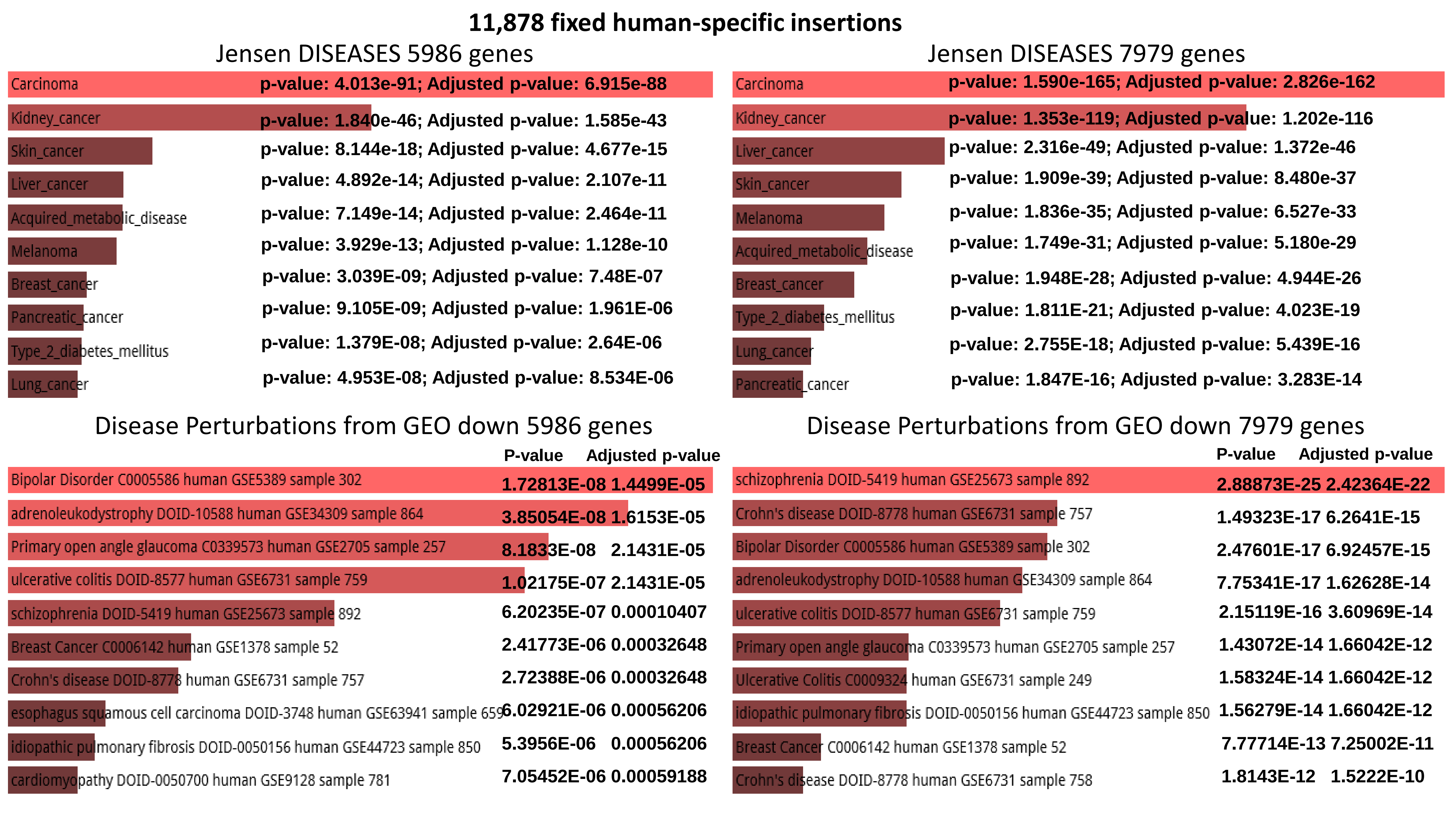

11,878 fixed human-specific insertions
Jensen DISEASES 5986 genes
Carcinoma p-value: 4.013e-91; Adjusted p-value: 6.915e-88
Kidney_cancer p-value: 1.840e-46; Adjusted p-value: 1.585e-43
Skin_cancer p-value: 8.144e-18; Adjusted p-value: 4.677e-15
Liver_cancer p-value: 4.892e-14; Adjusted p-value: 2.107e-11
Acquired_metabolic_disease p-value: 7.149e-14; Adjusted p-value: 2.464e-11
Melanoma p-value: 3.929e-13; Adjusted p-value: 1.128e-10
Breast_cancer p-value: 3.039E-09; Adjusted p-value: 7.48E-07
Pancreatic_cancer p-value: 9.105E-09; Adjusted p-value: 1.961E-06
Type_2_diabetes_mellitus p-value: 1.379E-08; Adjusted p-value: 2.64E-06
Lung_cancer p-value: 4.953E-08; Adjusted p-value: 8.534E-06
Jensen DISEASES 7979 genes
Carcinoma p-value: 1.590e-165; Adjusted p-value: 2.826e-162
Kidney_cancer p-value: 1.353e-119; Adjusted p-value: 1.202e-116
Liver_cancer p-value: 2.316e-49; Adjusted p-value: 1.372e-46
Skin_cancer p-value: 1.909e-39; Adjusted p-value: 8.480e-37
Melanoma p-value: 1.836e-35; Adjusted p-value: 6.527e-33
Acquired_metabolic_disease p-value: 1.749e-31; Adjusted p-value: 5.180e-29
Breast_cancer p-value: 1.948E-28; Adjusted p-value: 4.944E-26
Type_2_diabetes_mellitus p-value: 1.811E-21; Adjusted p-value: 4.023E-19
Lung_cancer p-value: 2.755E-18; Adjusted p-value: 5.439E-16
Pancreatic_cancer p-value: 1.847E-16; Adjusted p-value: 3.283E-14
Disease Perturbations from GEO down 5986 genes
P-value Adjusted p-value
Bipolar Disorder C0005586 human GSE5389 sample 302 1.72813E-08 1.4499E-05
adrenoleukodystrophy DOID-10588 human GSE34309 sample 864 3.85054E-08 1.6153E-05
Primary open angle glaucoma C0339573 human GSE2705 sample 257 8.1833E-08 2.1431E-05
ulcerative colitis DOID-8577 human GSE6731 sample 759 1.02175E-07 2.1431E-05
schizophrenia DOID-5419 human GSE25673 sample 892 6.20235E-07 0.00010407
Breast Cancer C0006142 human GSE1378 sample 52 2.41773E-06 0.00032648
Crohn's disease DOID-8778 human GSE6731 sample 757 2.72388E-06 0.00032648
esophagus squamous cell carcinoma DOID-3748 human GSE63941 sample 659 6.02921E-06 0.00056206
idiopathic pulmonary fibrosis DOID-0050156 human GSE44723 sample 850 5.3956E-06 0.00056206
cardiomyopathy DOID-0050700 human GSE9128 sample 781 7.05452E-06 0.00059188
Disease Perturbations from GEO down 7979 genes
P-value Adjusted p-value
schizophrenia DOID-5419 human GSE25673 sample 892 2.88873E-25 2.42364E-22
Crohn's disease DOID-8778 human GSE6731 sample 757 1.49323E-17 6.2641E-15
Bipolar Disorder C0005586 human GSE5389 sample 302 2.47601E-17 6.92457E-15
adrenoleukodystrophy DOID-10588 human GSE34309 sample 864 7.75341E-17 1.62628E-14
ulcerative colitis DOID-8577 human GSE6731 sample 759 2.15119E-16 3.60969E-14
Primary open angle glaucoma C0339573 human GSE2705 sample 257 1.43072E-14 1.66042E-12
Ulcerative Colitis C0009324 human GSE6731 sample 249 1.58324E-14 1.66042E-12
idiopathic pulmonary fibrosis DOID-0050156 human GSE44723 sample 850 1.56279E-14 1.66042E-12
Breast Cancer C0006142 human GSE1378 sample 52 7.77714E-13 7.25002E-11
Crohn's disease DOID-8778 human GSE6731 sample 758 1.8143E-12 1.5222E-10

### 11,878 fixed human-specific insertions

#### Disease Perturbations from GEO up 5986 genes

| Term | P-value | Adjusted p-value |
|---|---|---|
| cardiomyopathy DOID-0050700 human GSE9128 sample 780 | 1.87765E-06 | 0.001446337 |
| schizophrenia DOID-5419 human GSE25673 sample 891 | 3.44776E-06 | 0.001446337 |
| Spinal Muscular Atrophy C0026847 mouse GSE10599 sample 235 | 8.69631E-06 | 0.001889865 |
| idiopathic pulmonary fibrosis DOID-0050156 human GSE44723 sample 851 | 9.01008E-06 | 0.001889865 |
| Primary open angle glaucoma C0339573 human GSE2705 sample 257 | 3.83914E-05 | 0.006442083 |
| idiopathic urticaria DOID-1555 human GSE57178 sample 815 | 7.75193E-05 | 0.009776282 |
| Diamond-Blackfan anaemia DOID-1339 human GSE14335 sample 472 | 8.15661E-05 | 0.009776282 |
| Neurofibromatosis DOID-8712 mouse GSE1482 sample 667 | 0.000103508 | 0.010653102 |
| schizophrenia DOID-5419 human GSE25673 sample 892 | 0.000132556 | 0.010653102 |
| hepatitis C DOID-1883 human GSE20948 sample 598 | 0.000139671 | 0.010653102 |

#### Disease Perturbations from GEO up 7979 genes

| Term | P-value | Adjusted p-value |
|---|---|---|
| schizophrenia DOID-5419 human GSE25673 sample 891 | 1.8927E-19 | 1.58798E-16 |
| Spinal Muscular Atrophy C0026847 mouse GSE10599 sample 235 | 5.92303E-18 | 2.48471E-15 |
| schizophrenia DOID-5419 human GSE25673 sample 892 | 1.75756E-14 | 4.9153E-12 |
| idiopathic pulmonary fibrosis DOID-0050156 human GSE44723 sample 850 | 5.0913E-13 | 1.0679E-10 |
| morbid obesity DOID-11981 human GSE48964 sample 583 | 3.15E-11 | 5.28055E-09 |
| idiopathic pulmonary fibrosis DOID-0050156 human GSE44723 sample 851 | 6.54233E-11 | 9.14836E-09 |
| Primary open angle glaucoma C0339573 human GSE2705 sample 257 | 8.55002E-10 | 1.02478E-07 |
| cardiomyopathy DOID-0050700 human GSE9128 sample 780 | 3.23739E-09 | 3.39521E-07 |
| adrenoleukodystrophy DOID-10588 human GSE34309 sample 864 | 4.13498E-09 | 3.85472E-07 |
| Cardiomyopathy, Dilated C0007193 human GSE3585 sample 198 | 6.70904E-09 | 5.62888E-07 |

#### ENCODE and ChEA Consensus TFs from ChIP-X 5986 genes

| Term | P-value | Adjusted p-value |
|---|---|---|
| AR_CHEA | 3.841E-28 | 3.9946E-26 |
| SUZ12_CHEA | 1.09013E-22 | 5.66869E-21 |
| SMAD4_CHEA | 3.08881E-18 | 1.07079E-16 |
| REST_CHEA | 1.13969E-16 | 2.9632E-15 |
| NFE2L2_CHEA | 9.57565E-14 | 1.99174E-12 |
| TP63_CHEA | 3.48038E-10 | 6.03266E-09 |
| STAT3_CHEA | 2.40449E-08 | 3.57239E-07 |
| GATA1_CHEA | 1.41526E-07 | 1.48683E-06 |
| GATA2_CHEA | 8.89814E-08 | 1.15676E-06 |
| SALL4_CHEA | 1.42965E-07 | 1.48683E-06 |

#### ENCODE and ChEA Consensus TFs from ChIP-X 7979 genes

| Term | P-value | Adjusted p-value |
|---|---|---|
| AR_CHEA | 4.679E-85 | 4.866E-83 |
| SUZ12_CHEA | 1.084E-80 | 5.635E-79 |
| SMAD4_CHEA | 2.824E-61 | 9.789E-60 |
| NFE2L2_CHEA | 1.509E-60 | 3.923E-59 |
| REST_CHEA | 7.5683E-41 | 1.5742E-39 |
| SOX2_CHEA | 8.5218E-26 | 1.4771E-24 |
| TRIM28_CHEA | 2.2682E-22 | 3.3699E-21 |
| SALL4_CHEA | 1.8672E-19 | 2.4274E-18 |
| GATA1_CHEA | 6.2282E-19 | 7.1970E-18 |
| TP63_CHEA | 1.3467E-17 | 1.2732E-16 |

### 11,878 fixed human-specific insertions

#### ESCAPE 5986 genes

| Term | P-value | Adjusted p-value |
|---|---|---|
| hESC_H3K27me3_20682450 | 5.89784E-14 | 1.79884E-11 |
| mESC_H3K27me3_17603471 | 1.78285E-13 | 2.71885E-11 |
| mESC_H3K9me3_19884255 | 1.91796E-09 | 1.94993E-07 |
| CHiP_SUZ12-18974828 | 3.79545E-08 | 2.89403E-06 |
| mMEF_K27me3_17603471 | 1.44703E-07 | 8.82688E-06 |
| CHiP_MTF2-20144788 | 8.63335E-07 | 4.38862E-05 |
| CHiP_SUZ12-18692474 | 1.42594E-06 | 6.21303E-05 |
| CHiP_EZH2-18974828 | 1.21833E-05 | 0.000464488 |
| mNPC_K27me3_17603471 | 5.49037E-05 | 0.001674563 |
| CHiP_SUZ12-18555785 | 5.27687E-05 | 0.001674563 |

#### ESCAPE 7979 genes

| Term | P-value | Adjusted p-value |
|---|---|---|
| mESC_H3K27me3_17603471 | 4.44942E-51 | 1.37932E-48 |
| CHiP_SUZ12-18974828 | 2.39781E-38 | 3.71661E-36 |
| hESC_H3K27me3_20682450 | 2.69286E-37 | 2.78263E-35 |
| CHiP_SUZ12-18692474 | 3.02417E-32 | 2.34373E-30 |
| CHiP_MTF2-20144788 | 5.57499E-32 | 3.4565E-30 |
| CHiP_EZH2-18974828 | 1.0298E-31 | 5.32065E-30 |
| mESC_H3K9me3_19884255 | 4.45745E-27 | 1.97402E-25 |
| CHiP_JARID2-20064375 | 4.72639E-26 | 1.83148E-24 |
| CHiP_RNF2-22325148 | 4.92907E-25 | 1.69779E-23 |
| CHiP_RNF2-18974828 | 3.7711E-24 | 1.16904E-22 |

#### GTEx Tissue Sample Gene Expression Profiles up 5986 genes

| Term | P-value | Adjusted p-value |
|---|---|---|
| GTEX-T2IS-0011-R5A-SM-32QP4_brain_female_20-29_years | 7.39447E-21 | 2.15771E-17 |
| GTEX-QDT8-0011-R10A-SM-32PKG_brain_female_30-39_years | 5.34561E-20 | 7.79924E-17 |
| GTEX-OIZI-0008-SM-2XCFD_skin_male_40-49_years | 6.25612E-19 | 6.08512E-16 |
| GTEX-SNMC-1526-SM-2XCFN_blood vessel_male_20-29_years | 1.49898E-16 | 1.0935E-13 |
| GTEX-QDT8-0011-R2A-SM-32PKQ_brain_female_30-39_years | 6.57826E-16 | 3.83907E-13 |
| GTEX-OIZI-0526-SM-2XCEG_blood vessel_male_40-49_years | 1.52296E-15 | 7.40664E-13 |
| GTEX-QMR6-0011-R8A-SM-32PKJ_brain_male_50-59_years | 2.96092E-15 | 1.23428E-12 |
| GTEX-NPJ8-0011-R8a-SM-2HMLG_brain_male_40-49_years | 4.29977E-15 | 1.56834E-12 |
| GTEX-SN8G-0526-SM-32PLE_blood vessel_female_50-59_years | 5.27021E-14 | 1.70872E-11 |
| GTEX-PVOW-0011-R5A-SM-32PL7_brain_male_40-49_years | 7.63633E-14 | 2.22828E-11 |

#### GTEx Tissue Sample Gene Expression Profiles up 7979 genes

| Term | P-value | Adjusted p-value |
|---|---|---|
| GTEX-SNMC-1526-SM-2XCFN_blood vessel_male_20-29_years | 1.00794E-41 | 2.94116E-38 |
| GTEX-T2IS-0011-R5A-SM-32QP4_brain_female_20-29_years | 1.1941E-40 | 1.74219E-37 |
| GTEX-QDT8-0011-R10A-SM-32PKG_brain_female_30-39_years | 1.21874E-39 | 1.18543E-36 |
| GTEX-OIZI-0008-SM-2XCFD_skin_male_40-49_years | 3.80382E-34 | 2.77489E-31 |
| GTEX-PVOW-0011-R5A-SM-32PL7_brain_male_40-49_years | 7.93741E-31 | 4.63227E-28 |
| GTEX-PVOW-0011-R3A-SM-32PKX_brain_male_40-49_years | 6.28057E-30 | 3.05445E-27 |
| GTEX-OIZI-0526-SM-2XCEG_blood vessel_male_40-49_years | 8.54238E-30 | 3.56095E-27 |
| GTEX-QDT8-0011-R2A-SM-32PKQ_brain_female_30-39_years | 9.93569E-29 | 3.62404E-26 |
| GTEX-TMMY-0626-SM-33HBD_blood vessel_female_40-49_years | 1.71193E-28 | 5.55046E-26 |
| GTEX-NPJ8-0011-R8a-SM-2HMLG_brain_male_40-49_years | 1.91286E-28 | 5.58173E-26 |

### 11,878 fixed human-specific insertions

#### GTEx Tissue Sample Gene Expression Profiles down 7979 genes

| | P-value | Adjusted p-value |
|---|---|---|
| GTEX-X638-0005-SM-47JX6_blood_female_70-79_years | 1.04434E-63 | 3.04737E-60 |
| GTEX-XQ3S-0006-SM-4BOQ4_blood_male_20-29_years | 3.90159E-53 | 5.69242E-50 |
| GTEX-XMK1-0005-SM-4B665_blood_male_40-49_years | 1.28663E-47 | 1.25146E-44 |
| GTEX-XMD3-0006-SM-4AT5X_blood_female_50-59_years | 3.40004E-45 | 2.48033E-42 |
| GTEX-NPJ7-0006-SM-3GACR_blood_female_60-69_years | 3.02613E-42 | 1.76605E-39 |
| GTEX-XLM4-0005-SM-4AT4P_blood_male_60-69_years | 1.75487E-38 | 8.53453E-36 |
| GTEX-WCDI-0005-SM-3NB2M_blood_male_50-59_years | 7.88578E-38 | 3.28725E-35 |
| GTEX-XGQ4-0004-SM-4AT5S_blood_male_50-59_years | 5.61964E-37 | 2.04977E-34 |
| GTEX-X15G-0005-SM-3NMDA_blood_female_50-59_years | 5.55996E-36 | 1.80266E-33 |
| GTEX-XUYS-0002-SM-47JXL_blood_male_50-59_years | 9.02958E-35 | 2.3953E-32 |

#### GTEx Tissue Sample Gene Expression Profiles down 5986 genes

| | P-value | Adjusted p-value |
|---|---|---|
| GTEX-XQ3S-0006-SM-4BOQ4_blood_male_20-29_years | 4.60421E-31 | 1.34351E-27 |
| GTEX-X638-0005-SM-47JX6_blood_female_70-79_years | 2.69265E-30 | 3.92858E-27 |
| GTEX-XMK1-0005-SM-4B665_blood_male_40-49_years | 1.1337E-25 | 1.10271E-22 |
| GTEX-XMD3-0006-SM-4AT5X_blood_female_50-59_years | 5.63683E-22 | 4.11207E-19 |
| GTEX-NPJ7-0006-SM-3GACR_blood_female_60-69_years | 7.72356E-21 | 4.50747E-18 |
| GTEX-VJYA-0005-SM-3P5ZD_blood_male_60-69_years | 5.96893E-19 | 2.90289E-16 |
| GTEX-X4XX-0005-SM-3NMCS_blood_male_60-69_years | 8.15686E-19 | 3.40024E-16 |
| GTEX-XOT4-0005-SM-4B64S_blood_female_60-69_years | 4.54574E-18 | 1.65806E-15 |
| GTEX-WCDI-0005-SM-3NB2M_blood_male_50-59_years | 1.97267E-17 | 6.39584E-15 |
| GTEX-X88G-0006-SM-47JX5_blood_male_30-39_years | 4.52769E-17 | 1.32118E-14 |

### 4,637 human-specific TE-encoded loci expressed in human DLPFC

**4,637 human-specific TE-encoded loci expressed in human DLPFC**

| ARCHS4 Tissues 2323 genes | P-value | Adjusted p-value |
|---|---|---|
| PREFRONTAL CORTEX | 9.8570E-06 | 0.00107 |
| CINGULATE GYRUS | 0.000652 | 0.0295 |
| NEURONAL EPITHELIUM | 0.000820 | 0.0295 |
| FETAL BRAIN CORTEX | 0.004439 | 0.1199 |
| CEREBELLUM | | |
| BRAIN (BULK) | | |
| PANCREATIC ISLET | | |
| CEREBRAL CORTEX | | |
| DENTATE GRANULE CELL | | |
| DORSAL STRIATUM | | |

| ARCHS4 Tissues 4051 genes | P-value | Adjusted p-value |
|---|---|---|
| PREFRONTAL CORTEX | 7.00452E-50 | 7.56488E-48 |
| CINGULATE GYRUS | 6.71991E-34 | 3.62875E-32 |
| NEURONAL EPITHELIUM | 8.68697E-32 | 3.12731E-30 |
| CEREBELLUM | 9.55784E-29 | 2.58062E-27 |
| CEREBRAL CORTEX | 3.61112E-21 | 7.80002E-20 |
| SPINAL CORD | 2.49377E-20 | 3.84754E-19 |
| SPINAL CORD (BULK) | 2.49377E-20 | 3.84754E-19 |
| DORSAL STRIATUM | 1.66653E-18 | 2.24982E-17 |
| MOTOR NEURON | 6.09151E-15 | 7.30981E-14 |
| FETAL BRAIN CORTEX | 9.14708E-15 | 9.87885E-14 |

| Jensen TISSUES 2323 genes | P-value | Adjusted p-value |
|---|---|---|
| Hypothalamus | 6.32274E-21 | 8.46615E-18 |
| Occipital_lobe | 4.24993E-14 | 2.84533E-11 |
| Brain | 2.96068E-13 | 1.32145E-10 |
| Parietal_lobe | 6.54347E-12 | 2.19043E-09 |
| Adipose_tissue | 1.12994E-11 | 3.02598E-09 |
| Frontal_lobe | 2.30117E-11 | 5.13545E-09 |
| Cerebral_cortex | 1.36673E-10 | 2.61436E-08 |
| Thyroid_gland | 2.08353E-10 | 3.48731E-08 |
| Lymph_node | 2.7611E-10 | 4.1079E-08 |
| Ovary | 3.53351E-10 | 4.73136E-08 |

| Jensen TISSUES 4051 genes | P-value | Adjusted p-value |
|---|---|---|
| Hypothalamus | 1.42728E-50 | 2.3279E-47 |
| Brain | 1.03046E-34 | 8.40342E-32 |
| Cerebral_cortex | 2.82988E-25 | 1.53851E-22 |
| Frontal_lobe | 6.36197E-23 | 2.59409E-20 |
| Occipital_lobe | 1.39104E-20 | 4.53756E-18 |
| Heart | 4.28896E-18 | 1.16588E-15 |
| Ovary | 6.32042E-17 | 1.28858E-14 |
| Cerebellum | 5.93135E-17 | 1.28858E-14 |
| Parietal_lobe | 2.07605E-16 | 3.76226E-14 |
| Gall_bladder | 1.09267E-15 | 1.78215E-13 |

### 4,637 human-specific TE-encoded loci expressed in human DLPFC

#### Jensen DISEASES 2323 genes

| Term | P-value | Adjusted p-value |
|---|---|---|
| Carcinoma | 1.7817E-14 | 2.295E-11 |
| Kidney_cancer | 1.324E-05 | 0.0085 |
| Takayasu's_arteritis | 0.00191 | 0.615 |
| Liver_cancer | 0.00475 | 0.615 |
| Essential_hypertension | 0.00481 | 0.615 |
| Major_depressive_disorder | 0.00572 | 0.615 |
| Aicardi-Goutieres_syndrome | 0.00548 | 0.615 |
| Sclerocornea | 0.00503 | 0.615 |
| Schizophrenia | 0.00311 | 0.615 |
| Melanoma | 0.00360 | 0.615 |

#### Jensen DISEASES 4051 genes

| Term | P-value | Adjusted p-value |
|---|---|---|
| Carcinoma | 1.28507E-65 | 2.0317E-62 |
| Kidney_cancer | 9.99971E-54 | 7.90477E-51 |
| Liver_cancer | 7.45524E-35 | 3.92891E-32 |
| Melanoma | 1.10876E-26 | 4.38237E-24 |
| Skin_cancer | 2.69121E-26 | 8.50962E-24 |
| Breast_cancer | 8.01818E-17 | 2.11279E-14 |
| Acquired_metabolic_disease | 1.65991E-14 | 3.74902E-12 |
| Lung_cancer | 2.30061E-14 | 4.54659E-12 |
| Endometrial_cancer | 9.04405E-12 | 1.58874E-09 |
| Attention_deficit_hyperactivity_disorder | 4.39818E-11 | 6.95353E-09 |

#### Disease Perturbations from GEO down 2323 genes

| Term | P-value |
|---|---|
| dilated cardiomyopathy DOID-12930 human GSE42955 sample 678 | 0.00151 |
| idiopathic pulmonary fibrosis DOID-0050156 human GSE24206 sample 869 | 0.00452 |
| Bipolar Disorder C0005586 human GSE5389 sample 302 | 0.00559 |
| autism spectrum disorder DOID-0060041 human GSE62632 sample 1037 | 0.00941 |
| peripartum cardiomyopathy DOID-9997 human GSE42955 sample 817 | 0.01179 |
| Acne C0702166 human GSE10432 sample 297 | 0.01285 |
| cardiomyopathy DOID-0050700 human GSE9128 sample 781 | 0.01552 |
| idiopathic pulmonary fibrosis DOID-0050156 human GSE24206 sample 872 | 0.01657 |
| Down Syndrome C0013080 human GSE5390 sample 277 | 0.01875 |
| Down syndrome DOID-14250 human GSE16677 sample 1064 | 0.01994 |

#### Disease Perturbations from GEO down 4051 genes

| Term | P-value | Adjusted p-value |
|---|---|---|
| Bipolar Disorder C0005586 human GSE5389 sample 302 | 2.14561E-10 | 1.80017E-07 |
| autism spectrum disorder DOID-0060041 human GSE62632 sample 1037 | 8.52294E-10 | 3.57537E-07 |
| schizophrenia DOID-5419 human GSE25673 sample 892 | 3.04969E-09 | 8.52897E-07 |
| idiopathic pulmonary fibrosis DOID-0050156 human GSE44723 sample 850 | 3.50643E-08 | 6.19354E-06 |
| Crohn's disease DOID-8778 human GSE6731 sample 757 | 3.69102E-08 | 6.19354E-06 |
| schizophrenia DOID-5419 human GSE25673 sample 891 | 1.45048E-07 | 2.02826E-05 |
| ulcerative colitis DOID-8577 human GSE6731 sample 759 | 3.44284E-07 | 4.12649E-05 |
| Crohn's disease DOID-8778 human GSE6731 sample 758 | 2.50335E-06 | 0.000262539 |
| Alzheimer's disease DOID-10652 human GSE36980 sample 520 | 7.29851E-06 | 0.000663404 |
| Nephroblastoma C0027708 human GSE2712 sample 418 | 8.00156E-06 | 0.000663404 |

#### 4,637 human-specific TE-encoded loci expressed in human DLPFC

##### Disease Perturbations from GEO up 2323 genes

| Term | P-value |
|---|---|
| Spinal Muscular Atrophy C0026847 mouse GSE10599 sample 235 | 0.00102 |
| oligodendroglioma DOID-3181 human GSE15824 sample 858 | 0.00129 |
| Hypoxia C0242184 human GSE4483 sample 440 | 0.00265 |
| familial combined hyperlipidemia DOID-13809 human GSE11393 sample 773 | 0.00343 |
| Familial combined hyperlipidaemia C0020474 human GSE11393 sample 267 | 0.00477 |
| Acute Lung Injury C0242488 human GSE10474 sample 168 | 0.00495 |
| astrocytoma DOID-3069 human GSE15824 sample 861 | 0.00581 |
| oligodendroglioma DOID-3181 human GSE15824 sample 859 | 0.00725 |
| fragile X syndrome DOID-14261 human GSE7329 sample 809 | 0.00738 |
| Huntington's disease DOID-12858 human GSE1751 sample 795 | 0.00813 |

##### Disease Perturbations from GEO up 4051 genes

| Term | P-value | Adjusted p-value |
|---|---|---|
| Spinal Muscular Atrophy C0026847 mouse GSE10599 sample 235 | 1.30588E-20 | 1.09564E-17 |
| adrenoleukodystrophy DOID-10588 human GSE34309 sample 864 | 1.45115E-11 | 6.08757E-09 |
| morbid obesity DOID-11981 human GSE48964 sample 583 | 4.21321E-08 | 1.17829E-05 |
| schizophrenia DOID-5419 human GSE25673 sample 892 | 7.40939E-08 | 1.55412E-05 |
| schizophrenia DOID-5419 human GSE25673 sample 891 | 1.09191E-07 | 1.83222E-05 |
| idiopathic pulmonary fibrosis DOID-0050156 human GSE44723 sample 851 | 2.54256E-07 | 3.55534E-05 |
| idiopathic pulmonary fibrosis DOID-0050156 human GSE44723 sample 850 | 4.62071E-07 | 5.53825E-05 |
| adrenoleukodystrophy DOID-10588 human GSE34308 sample 709 | 7.11277E-07 | 7.45952E-05 |
| smoldering myeloma DOID-9551 human GSE47552 sample 562 | 8.43186E-06 | 0.000786037 |
| melanoma DOID-1909 human GSE6887 sample 948 | 1.68944E-05 | 0.001417438 |

##### ENCODE and ChEA Consensus TFs from ChIP-X 2323 genes

| Term | P-value | Adjusted p-value |
|---|---|---|
| AR_CHEA | 2.35459E-06 | 0.000245 |
| UBTF_ENCODE | 8.26486E-05 | 0.004298 |
| NFE2L2_CHEA | 0.00051872 | 0.017982 |
| CREB1_CHEA | 0.000838763 | 0.021808 |
| SMAD4_CHEA | 0.00141901 | 0.029515 |
| REST_CHEA | 0.001930935 | 0.033470 |
| PPARD_CHEA | 0.005210793 | 0.077417 |
| E2F1_CHEA | 0.007865845 | 0.102256 |
| TCF3_ENCODE | 0.00921029 | 0.106430 |
| BHLHE40_ENCODE | 0.016264119 | 0.154696 |

##### ENCODE and ChEA Consensus TFs from ChIP-X 4051 genes

| Term | P-value | Adjusted p-value |
|---|---|---|
| AR_CHEA | 1.64997E-45 | 1.71596E-43 |
| SMAD4_CHEA | 8.28178E-35 | 4.30653E-33 |
| NFE2L2_CHEA | 1.04996E-28 | 3.63986E-27 |
| SUZ12_CHEA | 1.75619E-28 | 4.56609E-27 |
| REST_CHEA | 7.0257E-19 | 1.46134E-17 |
| TRIM28_CHEA | 1.70459E-09 | 2.95463E-08 |
| STAT3_CHEA | 1.13034E-07 | 1.47386E-06 |
| GATA1_CHEA | 1.13374E-07 | 1.47386E-06 |
| TCF3_CHEA | 3.29897E-07 | 3.81215E-06 |
| UBTF_ENCODE | 8.55045E-07 | 8.89247E-06 |

#### 4,637 human-specific TE-encoded loci expressed in human DLPFC

##### GTEx Tissue Sample Gene Expression Profiles up 2323 genes

| | P-value | Adjusted p-value |
|---|---|---|
| GTEX-QDT8-0011-R10A-SM-32PKG_brain_female_30-39_years | 7.1111E-12 | 2.07502E-08 |
| GTEX-PVOW-0011-R3A-SM-32PKX_brain_male_40-49_years | 9.49921E-10 | 9.23956E-07 |
| GTEX-QVJO-0011-R10A-SM-2S1QJ_brain_female_60-69_years | 7.11092E-10 | 9.23956E-07 |
| GTEX-WVLH-0011-R10A-SM-3MJFM_brain_male_50-59_years | 2.68224E-09 | 1.79484E-06 |
| GTEX-T2IS-0011-R5A-SM-32QP4_brain_female_20-29_years | 3.69056E-09 | 1.79484E-06 |
| GTEX-QMR6-0011-R10A-SM-32PKO_brain_male_50-59_years | 3.36753E-09 | 1.79484E-06 |
| GTEX-QMR6-0011-R8A-SM-32PKJ_brain_male_50-59_years | 2.36488E-08 | 9.85818E-06 |
| GTEX-WL46-0011-R10A-SM-3MJFQ_brain_male_50-59_years | 8.65217E-08 | 3.15588E-05 |
| GTEX-N7MS-0011-R3a-SM-33HC6_brain_male_60-69_years | 1.69843E-07 | 5.50669E-05 |
| GTEX-NL3H-0011-R3a-SM-2I3GL_brain_male_60-69_years | 2.34416E-07 | 6.84027E-05 |

##### GTEx Tissue Sample Gene Expression Profiles up 4051 genes

| | P-value | Adjusted p-value |
|---|---|---|
| GTEX-QDT8-0011-R10A-SM-32PKG_brain_female_30-39_years | 4.63841E-42 | 1.35349E-38 |
| GTEX-PVOW-0011-R3A-SM-32PKX_brain_male_40-49_years | 4.30355E-36 | 6.27888E-33 |
| GTEX-T2IS-0011-R5A-SM-32QP4_brain_female_20-29_years | 4.51564E-33 | 4.39221E-30 |
| GTEX-PVOW-0011-R5A-SM-32PL7_brain_male_40-49_years | 2.56809E-31 | 1.87342E-28 |
| GTEX-QMR6-0011-R10A-SM-32PKO_brain_male_50-59_years | 1.2302E-30 | 7.17944E-28 |
| GTEX-QMR6-0011-R8A-SM-32PKJ_brain_male_50-59_years | 2.1578E-28 | 1.04941E-25 |
| GTEX-N7MS-0011-R3a-SM-33HC6_brain_male_60-69_years | 1.01871E-26 | 4.24658E-24 |
| GTEX-WVLH-0011-R10A-SM-3MJFM_brain_male_50-59_years | 1.71691E-26 | 6.26244E-24 |
| GTEX-QVJO-0011-R10A-SM-2S1QJ_brain_female_60-69_years | 4.26311E-26 | 1.3822E-23 |
| GTEX-PVOW-2526-SM-2XCF7_brain_male_40-49_years | 5.24644E-26 | 1.53091E-23 |

##### GTEx Tissue Sample Gene Expression Profiles down 2323 genes

| | P-value | Adjusted p-value |
|---|---|---|
| GTEX-XQ3S-0006-SM-4BOQ4_blood_male_20-29_years | 1.57265E-10 | 4.58427E-07 |
| GTEX-XOT4-0005-SM-4B64S_blood_female_60-69_years | 7.49373E-09 | 1.09221E-05 |
| GTEX-WHSE-0006-SM-3NMBW_blood_male_20-29_years | 1.18638E-08 | 1.15277E-05 |
| GTEX-WZTO-0006-SM-3NM9T_blood_male_40-49_years | 2.0799E-07 | 0.000152 |
| GTEX-U3ZG-0326-SM-47JXN_muscle_male_50-59_years | 5.52956E-07 | 0.000288 |
| GTEX-SJXC-0005-SM-2XCE7_blood_male_60-69_years | 5.91986E-07 | 0.000288 |
| GTEX-X638-0005-SM-47JX6_blood_female_70-79_years | 1.2182E-06 | 0.000503 |
| GTEX-PVOW-0006-SM-3NMB8_blood_male_40-49_years | 1.37946E-06 | 0.000503 |
| GTEX-NPJ7-0006-SM-3GACR_blood_female_60-69_years | 1.6573E-06 | 0.000537 |
| GTEX-T6MN-0005-SM-32PLJ_blood_male_50-59_years | 2.45953E-06 | 0.000717 |

##### GTEx Tissue Sample Gene Expression Profiles down 4051 genes

| | P-value | Adjusted p-value |
|---|---|---|
| GTEX-X638-0005-SM-47JX6_blood_female_70-79_years | 4.65379E-32 | 1.35798E-28 |
| GTEX-XQ3S-0006-SM-4BOQ4_blood_male_20-29_years | 8.89061E-30 | 1.29714E-26 |
| GTEX-XMK1-0005-SM-4B665_blood_male_40-49_years | 9.61904E-26 | 9.35612E-23 |
| GTEX-NPJ7-0006-SM-3GACR_blood_female_60-69_years | 4.2827E-24 | 3.12423E-21 |
| GTEX-X88G-0006-SM-47JX5_blood_male_30-39_years | 1.85727E-22 | 1.0839E-19 |
| GTEX-UJHI-0006-SM-3DB8H_blood_female_50-59_years | 1.04583E-21 | 5.08622E-19 |
| GTEX-XLM4-0005-SM-4AT4P_blood_male_60-69_years | 1.99878E-21 | 8.33207E-19 |
| GTEX-XMD3-0006-SM-4AT5X_blood_female_50-59_years | 1.7854E-20 | 6.51223E-18 |
| GTEX-WCDI-0005-SM-3NB2M_blood_male_50-59_years | 3.80474E-20 | 1.23358E-17 |
| GTEX-OIZF-0006-SM-2I5GQ_blood_male_60-69_years | 4.35816E-20 | 1.27171E-17 |

**4,637 human-specific TE-encoded loci expressed in human DLPFC**

Tissue Protein Expression from Human Proteome Map 4051 genes

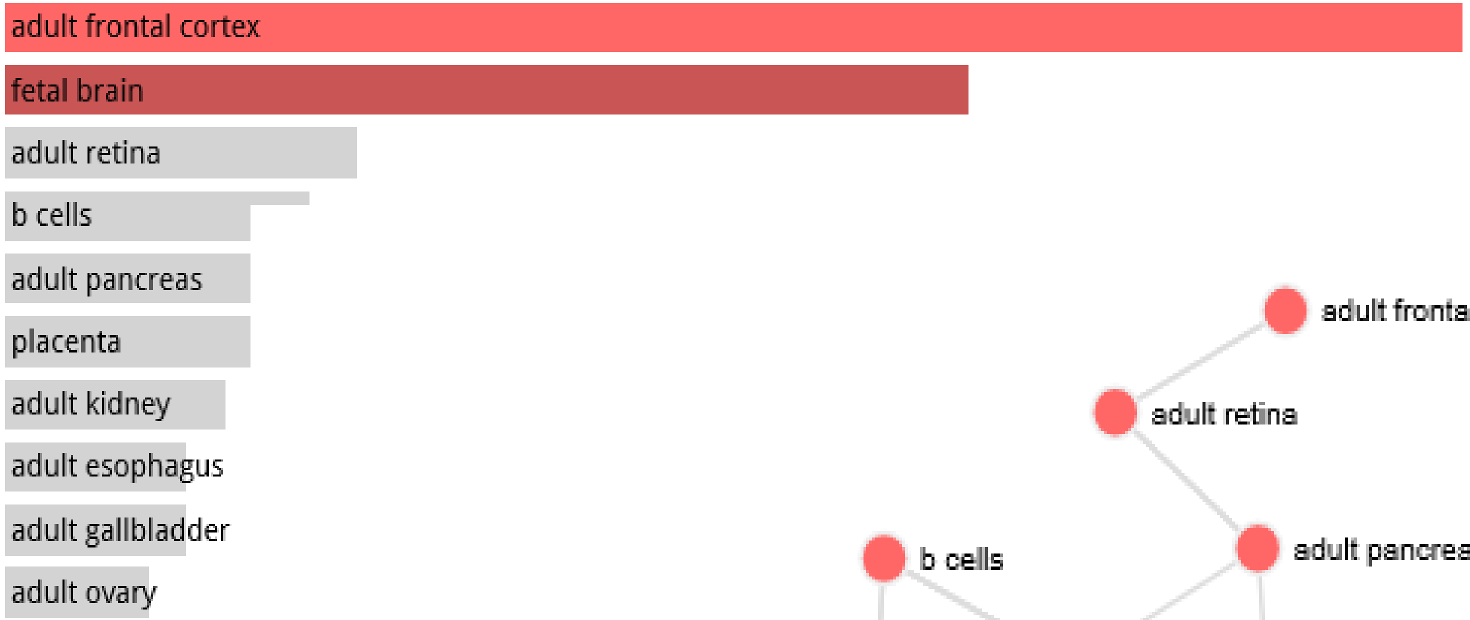


**p-value: 0.00006954 Adjusted p-value: 0.002086**

**p-value: 0.001310; Adjusted p-value: 0.01966**

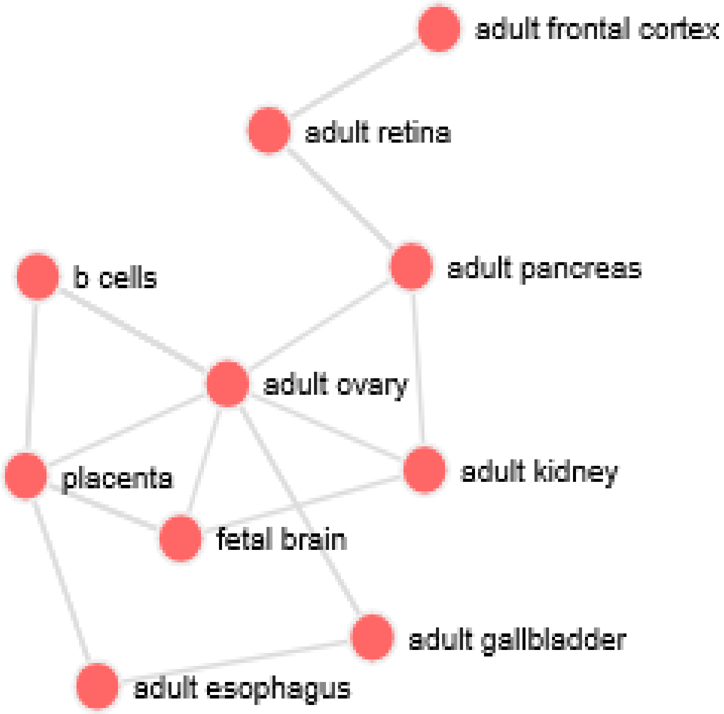

#### 4,637 human-specific TE-encoded loci expressed in human DLPFC

Allen Brain Atlas up 4051 genes

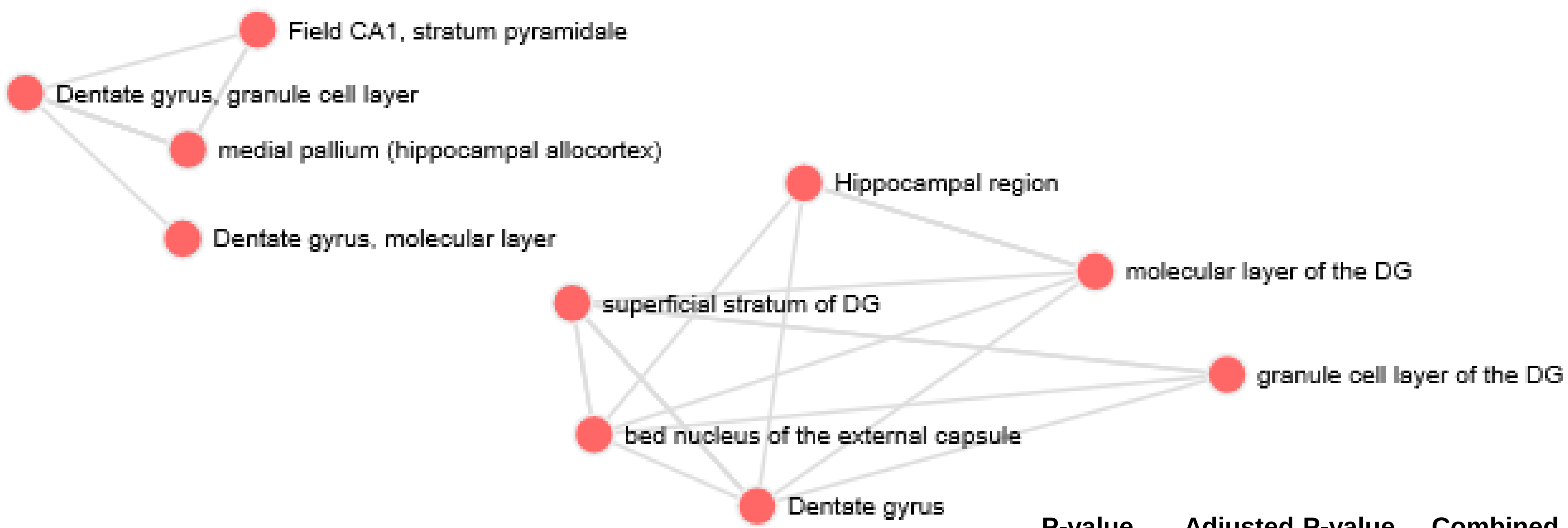


| | P-value | Adjusted P-value | Combined Score |
|---|---|---|---|
| Dentate gyrus | 2.06994E-09 | 1.5124E-06 | 36.78 |
| Dentate gyrus, molecular layer | 1.91586E-09 | 1.5124E-06 | 35.25 |
| granule cell layer of the DG | 1.83231E-08 | 8.0328E-06 | 33.31 |
| molecular layer of the DG | 1.91586E-09 | 1.5124E-06 | 32.40 |
| Hippocampal region | 3.77717E-08 | 1.1828E-05 | 31.40 |
| Dentate gyrus, granule cell layer | 3.77717E-08 | 1.1828E-05 | 31.15 |
| bed nucleus of the external capsule | 7.67266E-08 | 1.6818E-05 | 31.02 |
| Field CA1, stratum pyramidale | 7.67266E-08 | 1.6818E-05 | 30.60 |
| medial pallium (hippocampal allocortex) | 7.67266E-08 | 1.6818E-05 | 29.96 |
| superficial stratum of DG | 1.13777E-08 | 6.235E-06 | 29.70 |

sorted by combined score ranking

**4,637 human-specific TE-encoded loci expressed in human DLPFC**

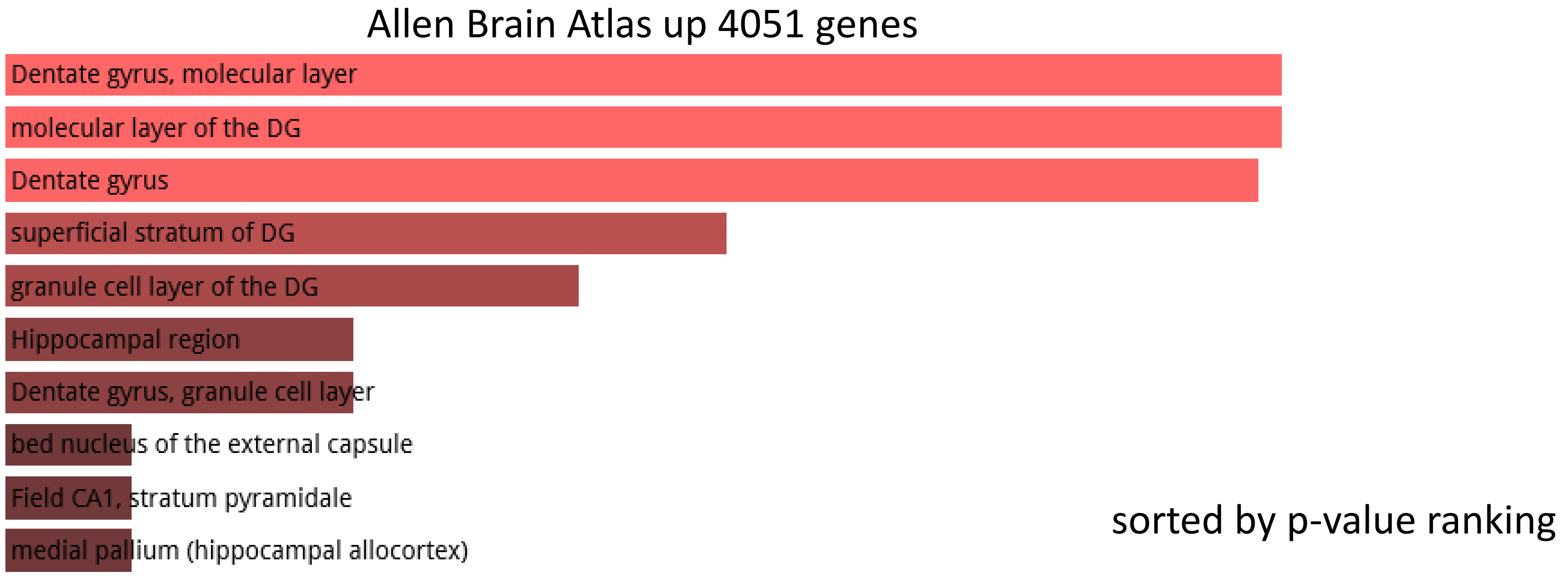


#### TOP 15 CATEGORIES

| Term | Overlap | P-value | Adjusted P-value |
|---|---|---|---|
| Dentate gyrus | 110/320 | 2.07E-09 | 1.51E-06 |
| Dentate gyrus, molecular layer | 105/301 | 1.92E-09 | 1.51E-06 |
| molecular layer of the DG | 105/301 | 1.92E-09 | 1.51E-06 |
| superficial stratum of DG | 179/601 | 1.14E-08 | 6.24E-06 |
| granule cell layer of the DG | 102/301 | 1.83E-08 | 8.03E-06 |
| Hippocampal region | 101/301 | 3.78E-08 | 1.18E-05 |
| Dentate gyrus, granule cell layer | 101/301 | 3.78E-08 | 1.18E-05 |
| bed nucleus of the external capsule | 100/301 | 7.67E-08 | 1.68E-05 |
| Field CA1, stratum pyramidale | 100/301 | 7.67E-08 | 1.68E-05 |
| medial pallium (hippocampal allocortex) | 100/301 | 7.67E-08 | 1.68E-05 |
| mantle zone of DG | 174/599 | 1.22E-07 | 2.4E-05 |
| Subiculum, dorsal part, molecular layer | 99/301 | 1.54E-07 | 2.4E-05 |
| hilus of the DG | 99/301 | 1.54E-07 | 2.4E-05 |
| Dentate gyrus, polymorph layer | 99/301 | 1.54E-07 | 2.4E-05 |
| hippocampus (cortex Ammonis) | 98/301 | 3.03E-07 | 2.88E-05 |

**4,637 human-specific TE-encoded loci expressed in human DLPFC**

Allen Brain Atlas up 4051 genes

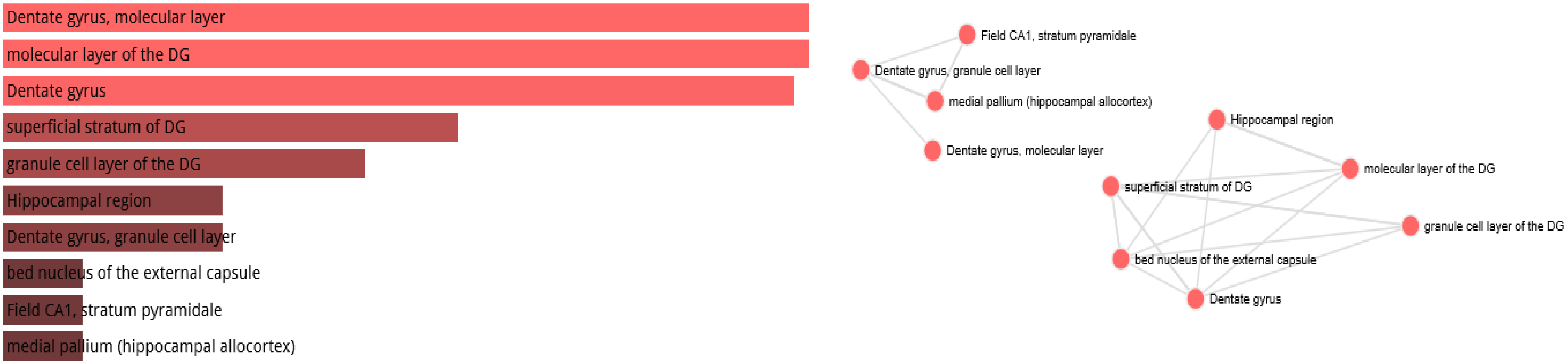


Allen Brain Atlas up 2323 genes

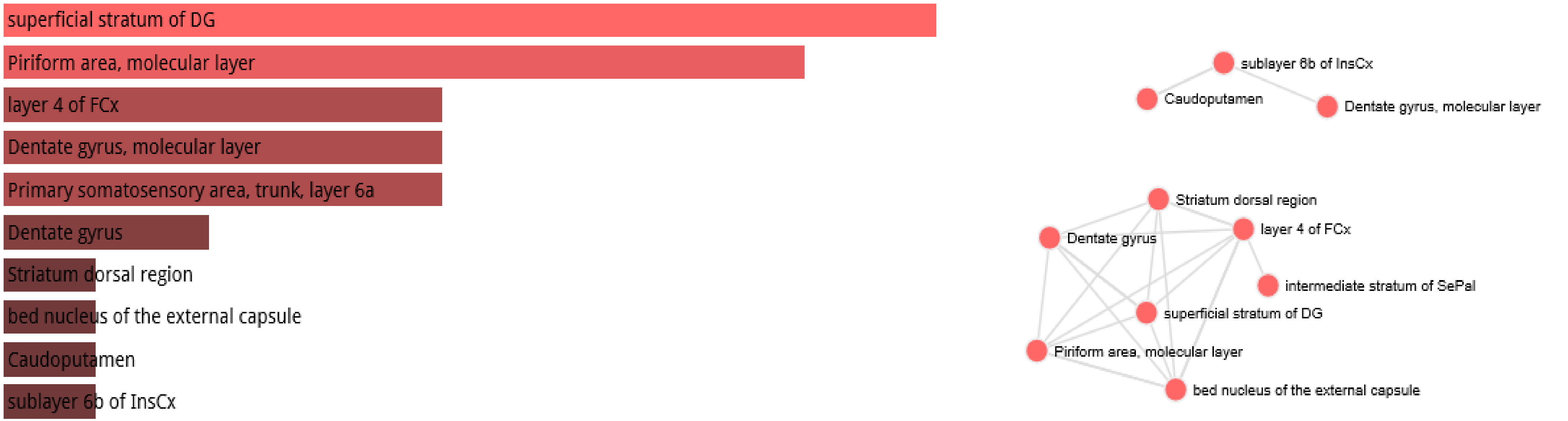

### Comparisons of putative regulatory target genes associated with

### 11,878 fixed human-specific insertions

### and

### 4,637 human-specific TE - encoded loci expressed in human dorsolateral prefrontal cortex (DLPFC)

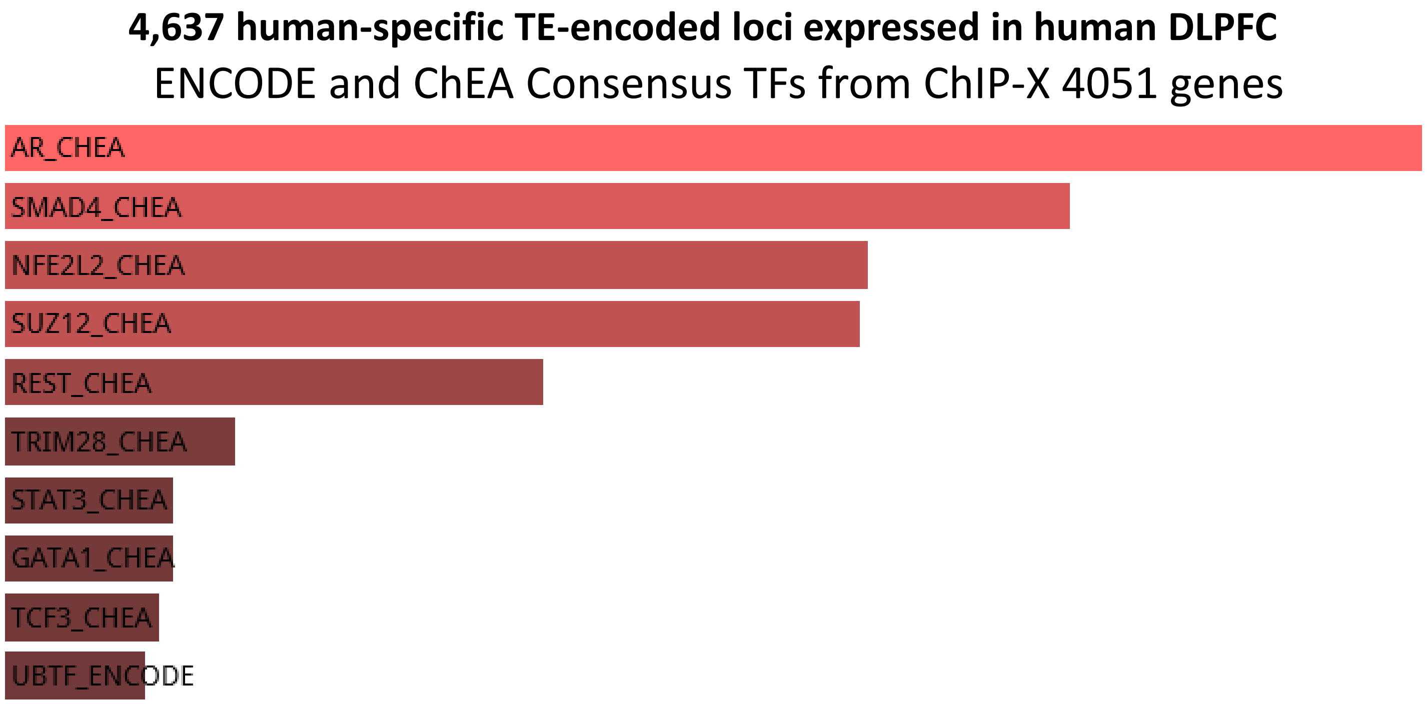
4,637 human-specific TE-encoded loci expressed in human DLPFC
ENCODE and ChEA Consensus TFs from ChIP-X 4051 genes
AR_CHEA
SMAD4_CHEA
NFE2L2_CHEA
SUZ12_CHEA
REST_CHEA
TRIM28_CHEA
STAT3_CHEA
GATA1_CHEA
TCF3_CHEA
UBTF_ENCODE

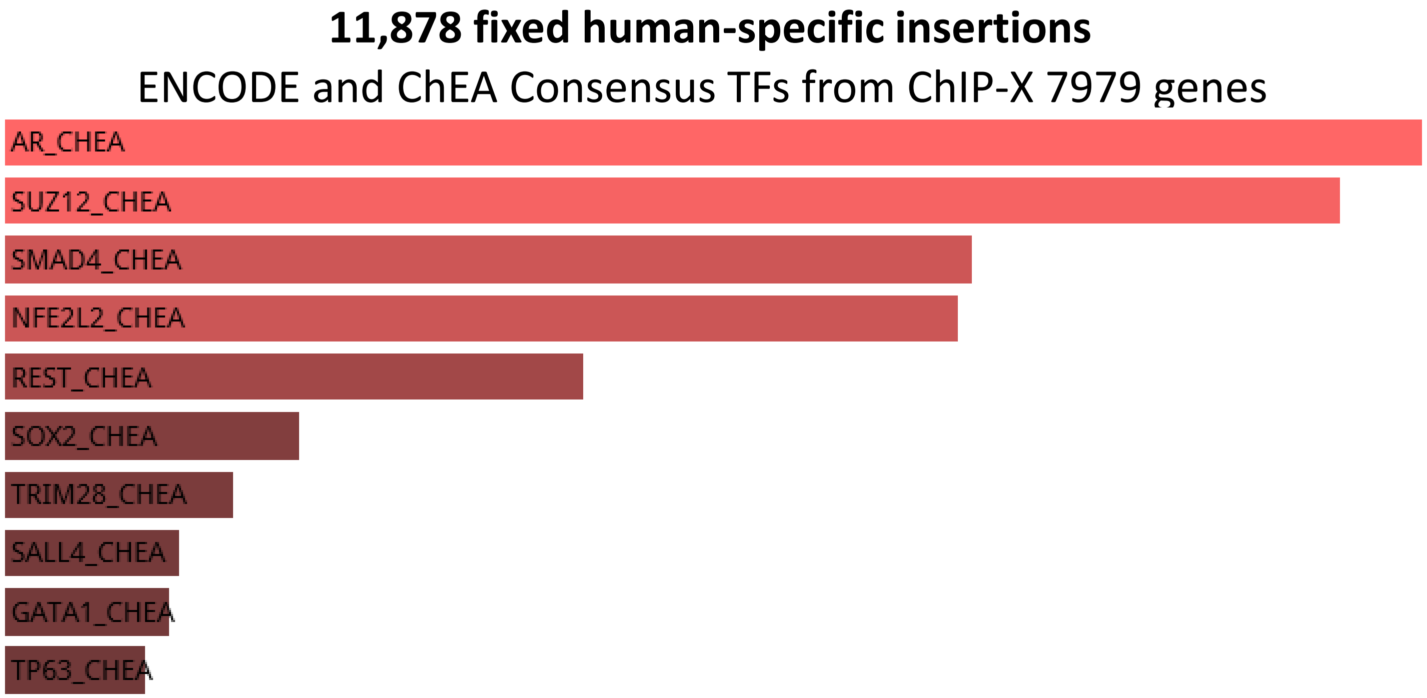
11,878 fixed human-specific insertions
ENCODE and ChEA Consensus TFs from ChIP-X 7979 genes
AR_CHEA
SUZ12_CHEA
SMAD4_CHEA
NFE2L2_CHEA
REST_CHEA
SOX2_CHEA
TRIM28_CHEA
SALL4_CHEA
GATA1_CHEA
TP63_CHEA

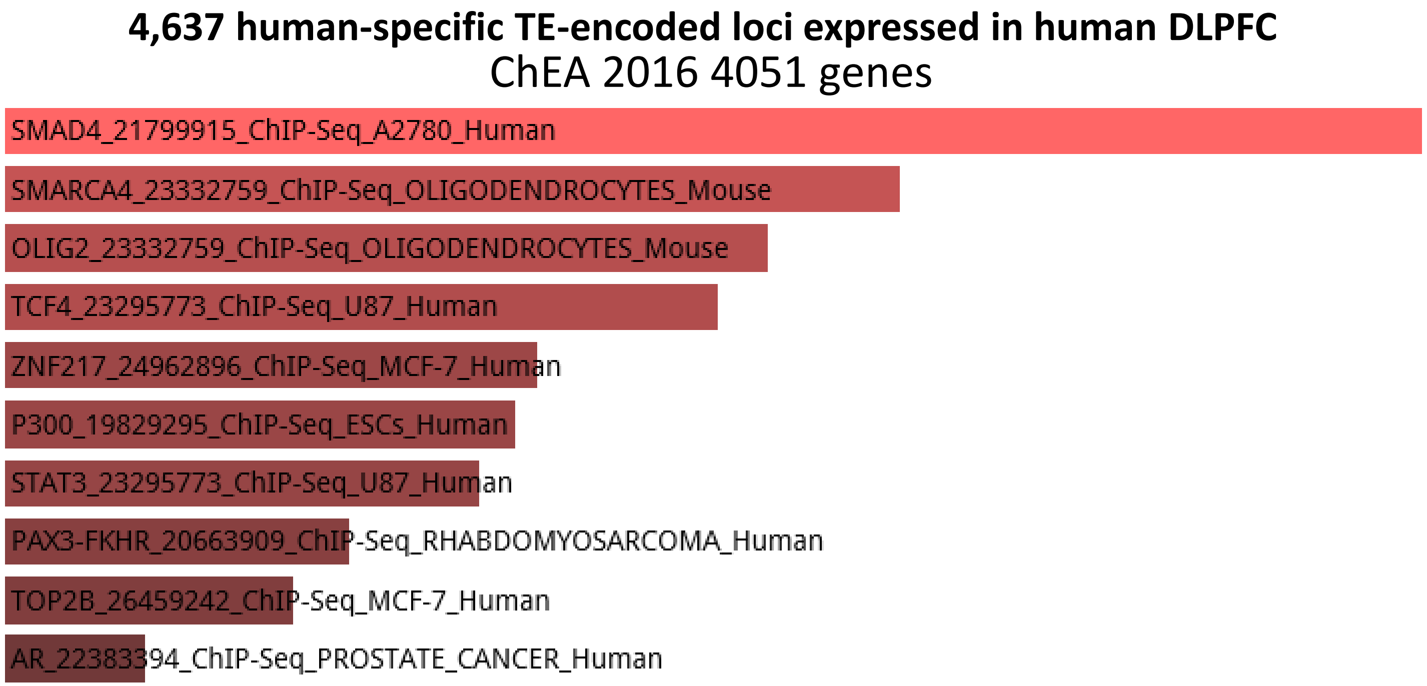
4,637 human-specific TE-encoded loci expressed in human DLPFC
ChEA 2016 4051 genes
SMAD4_21799915_ChIP-Seq_A2780_Human
SMARCA4_23332759_ChIP-Seq_OLIGODENDROCYTES_Mouse
OLIG2_23332759_ChIP-Seq_OLIGODENDROCYTES_Mouse
TCF4_23295773_ChIP-Seq_U87_Human
ZNF217_24962896_ChIP-Seq_MCF-7_Human
P300_19829295_ChIP-Seq_ESCs_Human
STAT3_23295773_ChIP-Seq_U87_Human
PAX3-FKHR_20663909_ChIP-Seq_RHABDOMYOSARCOMA_Human
TOP2B_26459242_ChIP-Seq_MCF-7_Human
AR_22383394_ChIP-Seq_PROSTATE_CANCER_Human

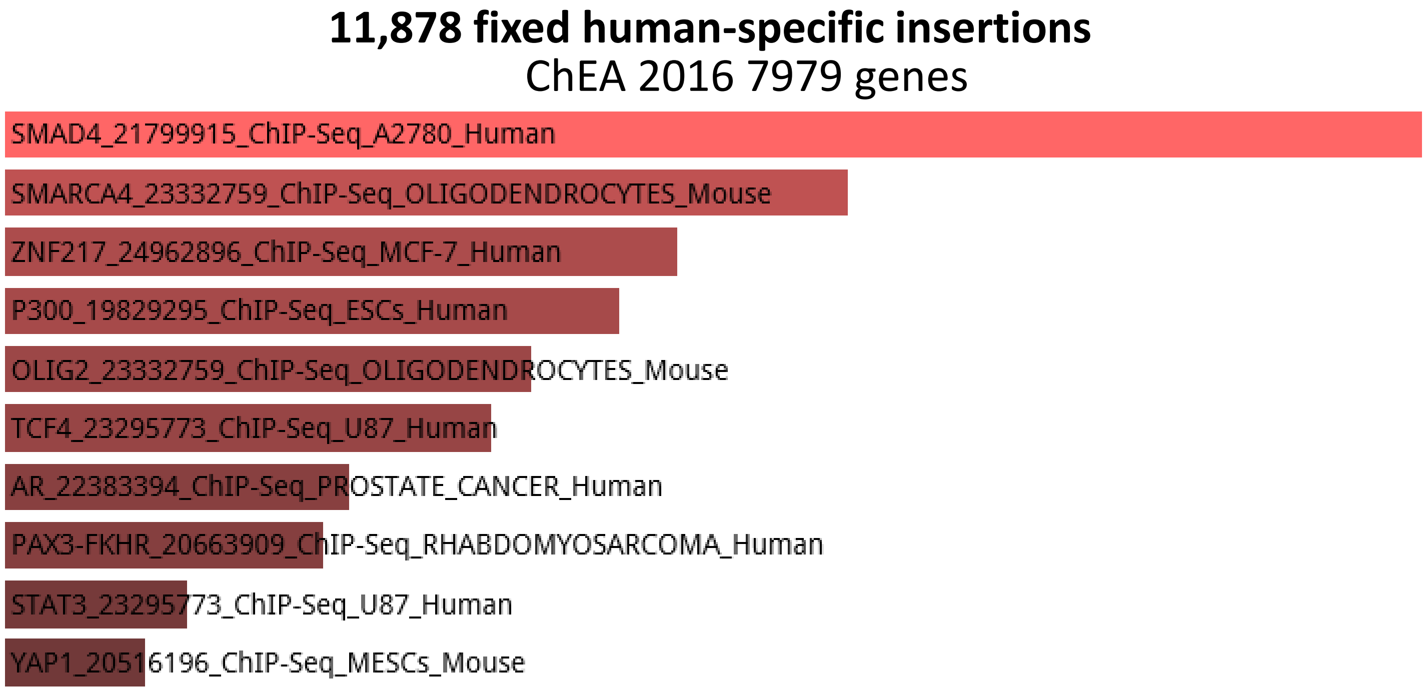
11,878 fixed human-specific insertions
ChEA 2016 7979 genes
SMAD4_21799915_ChIP-Seq_A2780_Human
SMARCA4_23332759_ChIP-Seq_OLIGODENDROCYTES_Mouse
ZNF217_24962896_ChIP-Seq_MCF-7_Human
P300_19829295_ChIP-Seq_ESCs_Human
OLIG2_23332759_ChIP-Seq_OLIGODENDROCYTES_Mouse
TCF4_23295773_ChIP-Seq_U87_Human
AR_22383394_ChIP-Seq_PROSTATE_CANCER_Human
PAX3-FKHR_20663909_ChIP-Seq_RHABDOMYOSARCOMA_Human
STAT3_23295773_ChIP-Seq_U87_Human
YAP1_20516196_ChIP-Seq_MESCs_Mouse

**4,637 human-specific TE-encoded loci expressed in human DLPFC**

TF Perturbations Followed by Expression 4051 genes

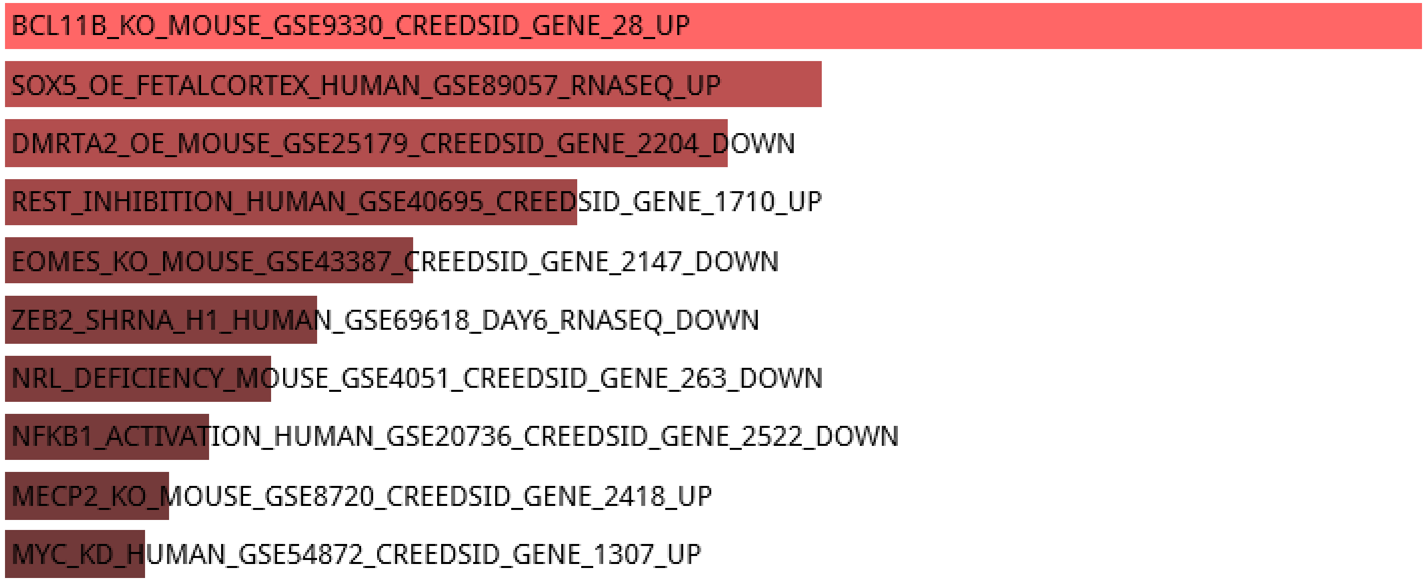


**11,878 fixed human-specific insertions**

TF Perturbations Followed by Expression 7979 genes

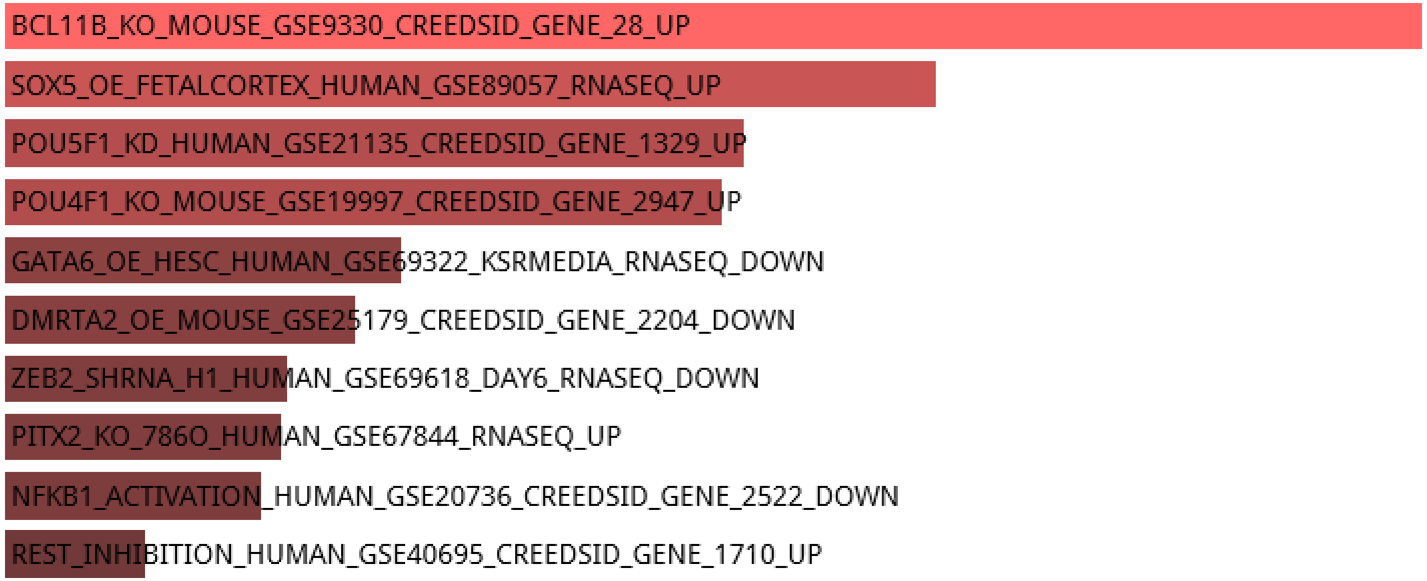

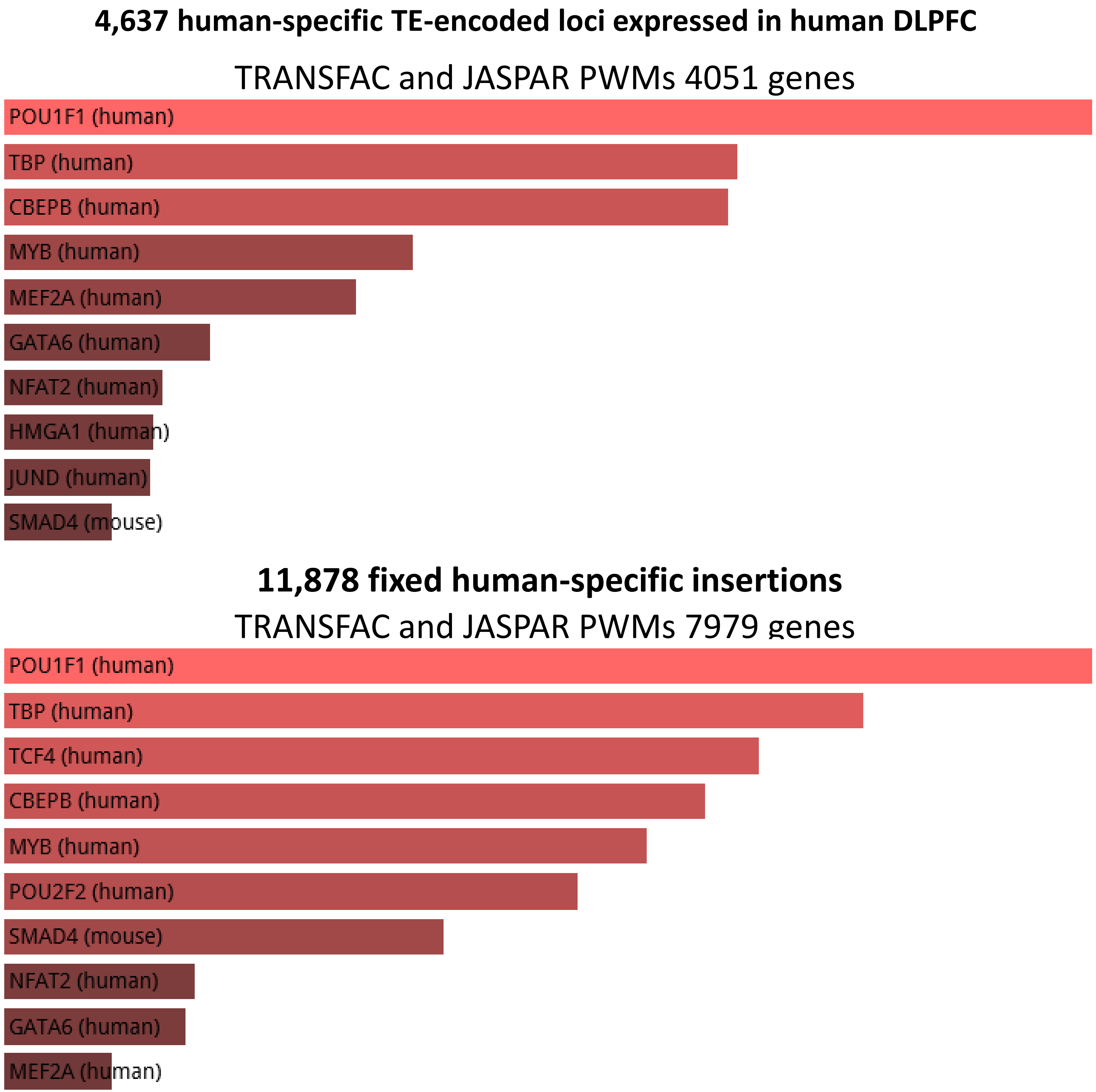
4,637 human-specific TE-encoded loci expressed in human DLPFC
TRANSFAC and JASPAR PWMs 4051 genes
POU1F1 (human)
TBP (human)
CBEPB (human)
MYB (human)
MEF2A (human)
GATA6 (human)
NFAT2 (human)
HMGA1 (human)
JUND (human)
SMAD4 (mouse)
11,878 fixed human-specific insertions
TRANSFAC and JASPAR PWMs 7979 genes
POU1F1 (human)
TBP (human)
TCF4 (human)
CBEPB (human)
MYB (human)
POU2F2 (human)
SMAD4 (mouse)
NFAT2 (human)
GATA6 (human)
MEF2A (human)

### RNA-Seq Expression Data from GTEx (53 Tissues, 570 Donors)

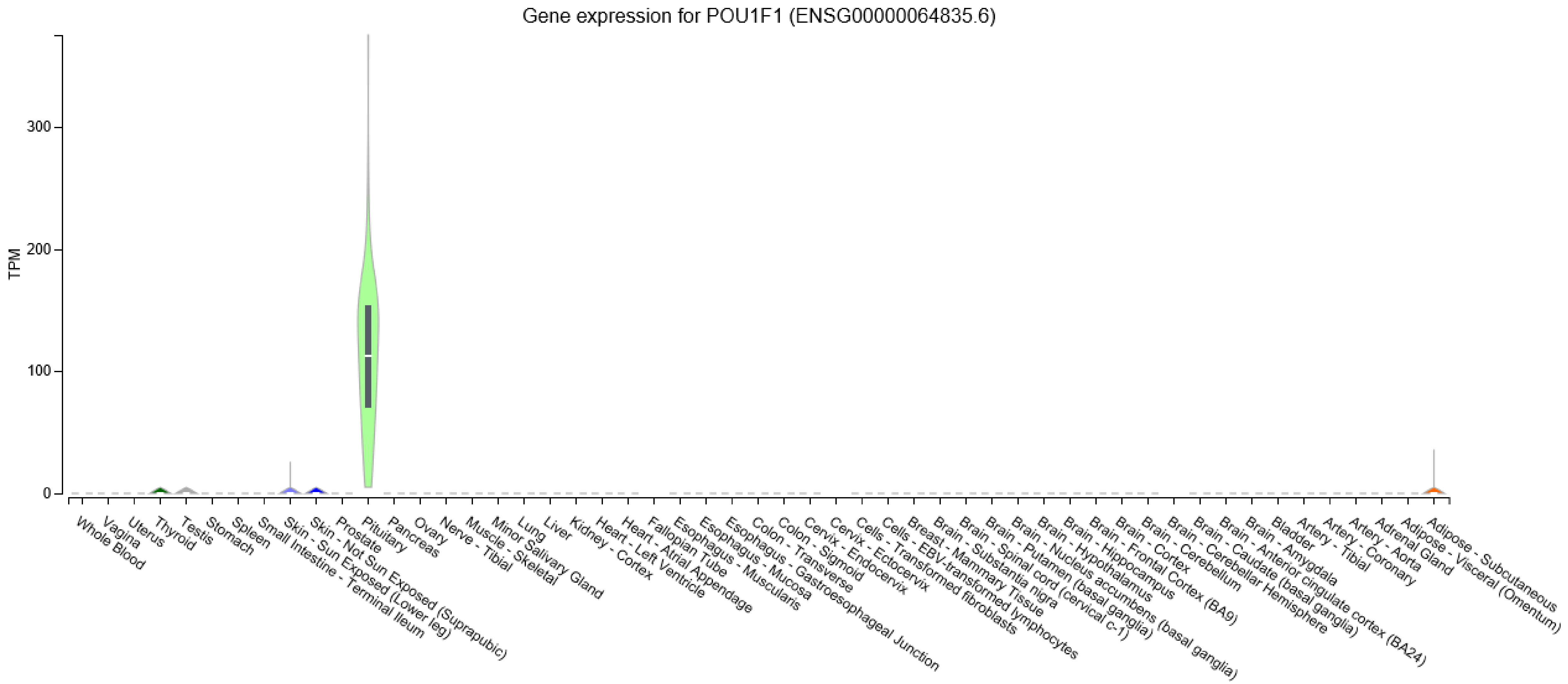

#### 4,637 human-specific TE-encoded loci expressed in human DLPFC

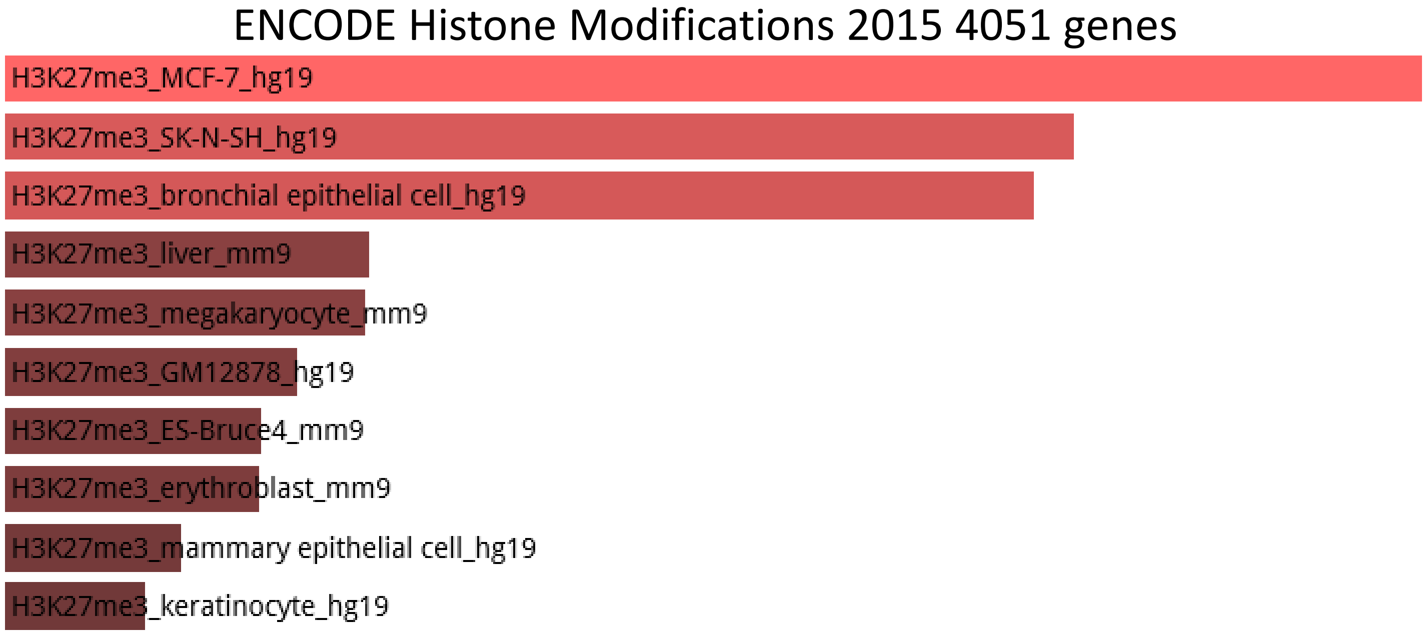


#### 11,878 fixed human-specific insertions

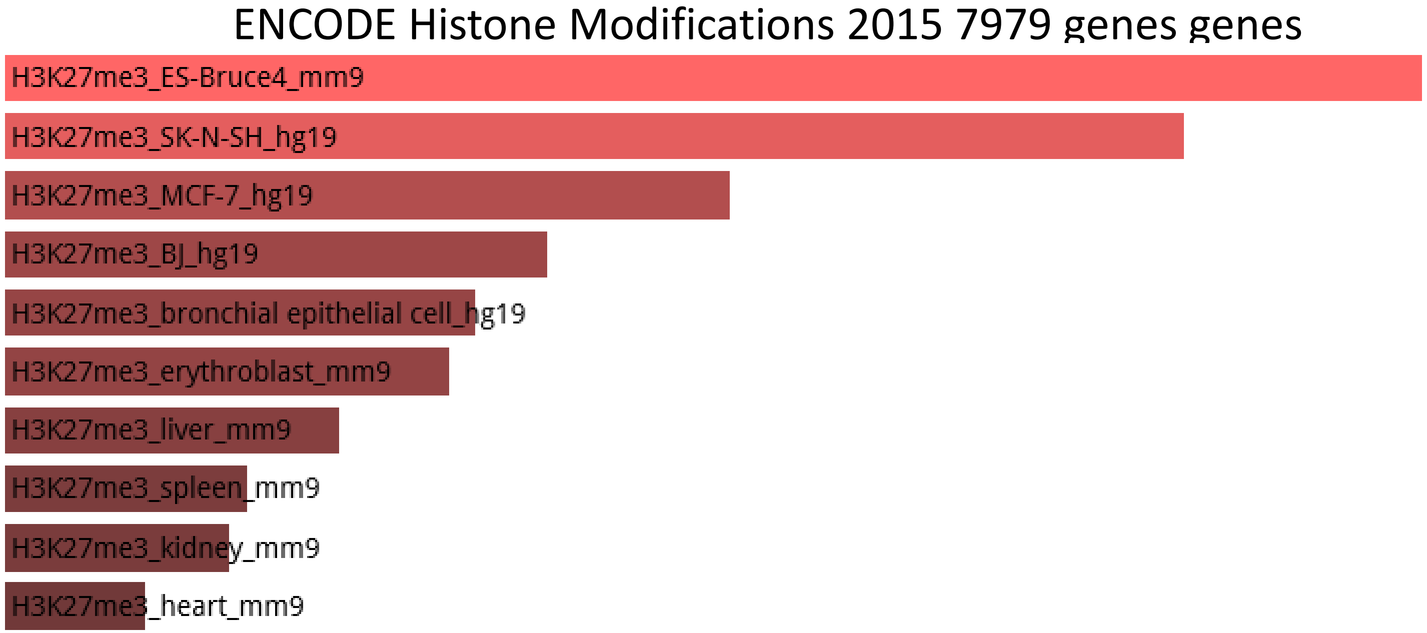

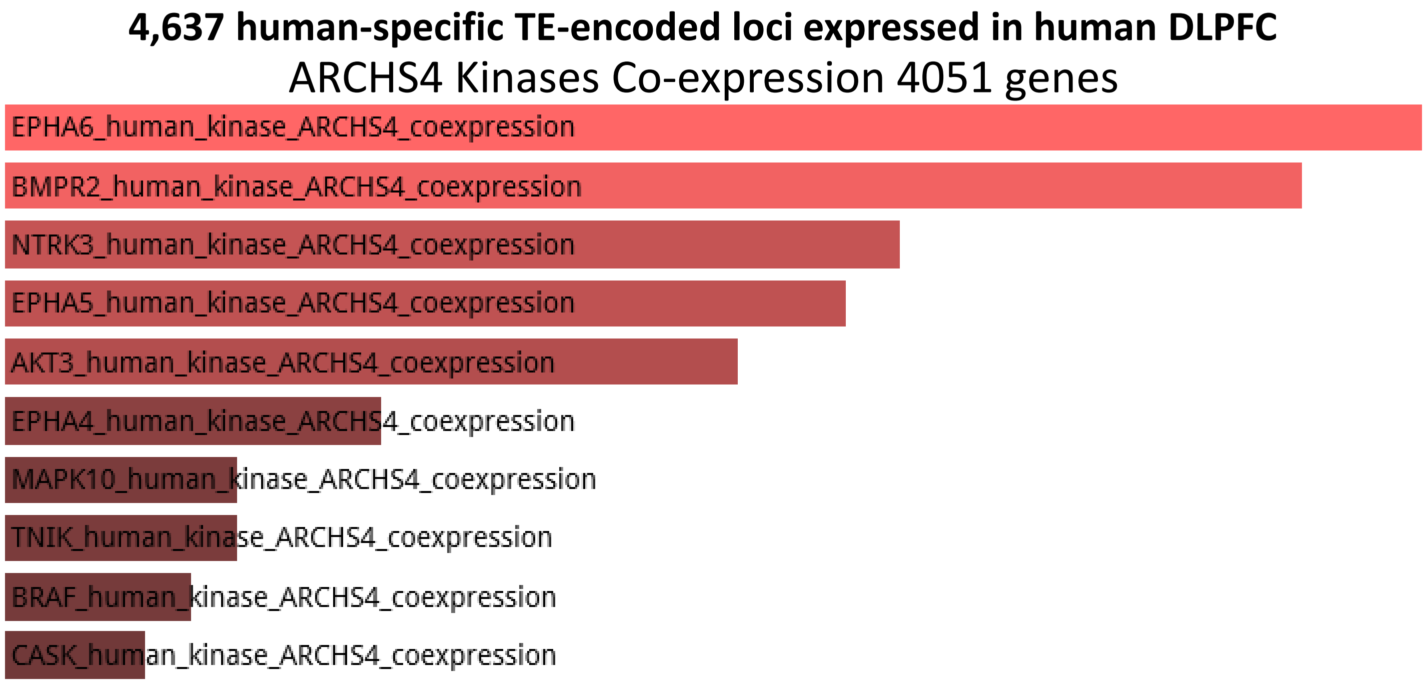
4,637 human-specific TE-encoded loci expressed in human DLPFC
ARCHS4 Kinases Co-expression 4051 genes
EPHA6_human_kinase_ARCHS4_coexpression
BMPR2_human_kinase_ARCHS4_coexpression
NTRK3_human_kinase_ARCHS4_coexpression
EPHA5_human_kinase_ARCHS4_coexpression
AKT3_human_kinase_ARCHS4_coexpression
EPHA4_human_kinase_ARCHS4_coexpression
MAPK10_human_kinase_ARCHS4_coexpression
TNIK_human_kinase_ARCHS4_coexpression
BRAF_human_kinase_ARCHS4_coexpression
CASK_human_kinase_ARCHS4_coexpression

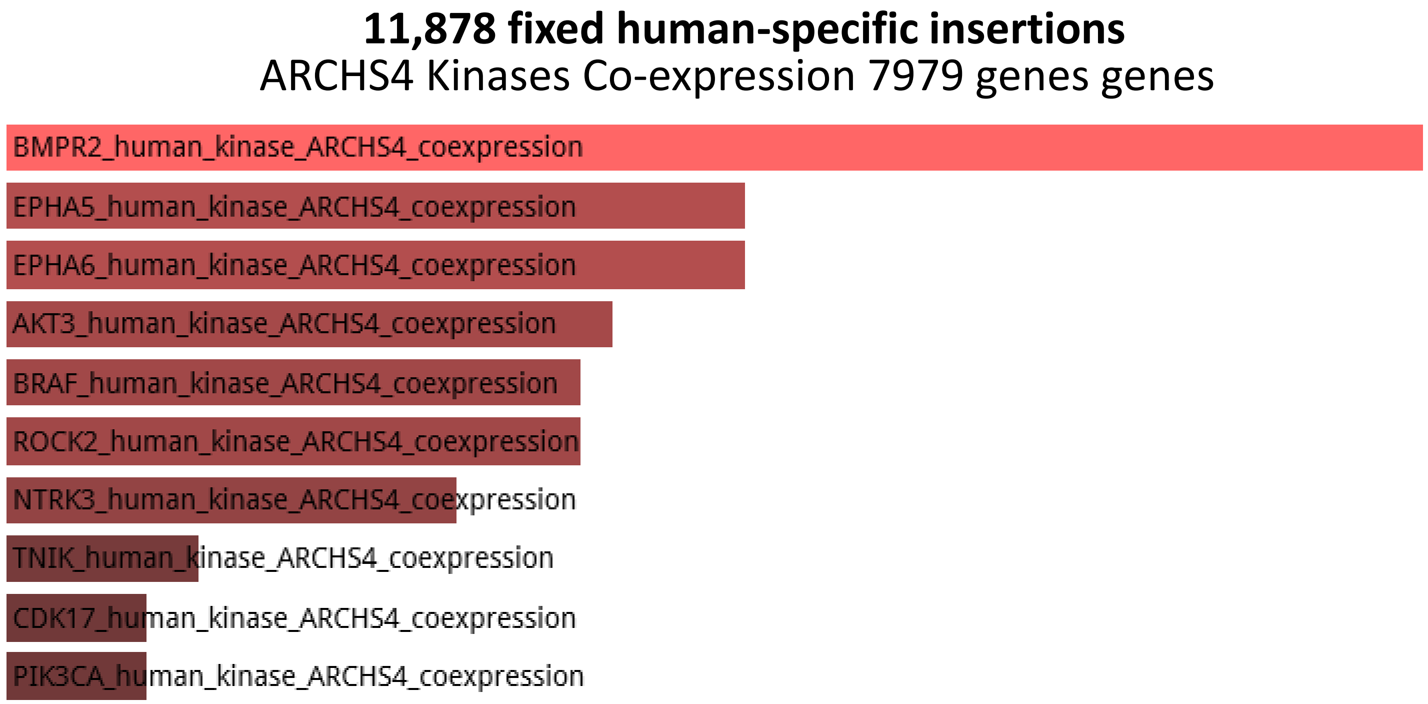
11,878 fixed human-specific insertions
ARCHS4 Kinases Co-expression 7979 genes genes
BMPR2_human_kinase_ARCHS4_coexpression
EPHA5_human_kinase_ARCHS4_coexpression
EPHA6_human_kinase_ARCHS4_coexpression
AKT3_human_kinase_ARCHS4_coexpression
BRAF_human_kinase_ARCHS4_coexpression
ROCK2_human_kinase_ARCHS4_coexpression
NTRK3_human_kinase_ARCHS4_coexpression
TNIK_human_kinase_ARCHS4_coexpression
CDK17_human_kinase_ARCHS4_coexpression
PIK3CA_human_kinase_ARCHS4_coexpression

**4,637 human-specific TE-encoded loci expressed in human DLPFC**

Transcription Factor PPIs 4051 genes

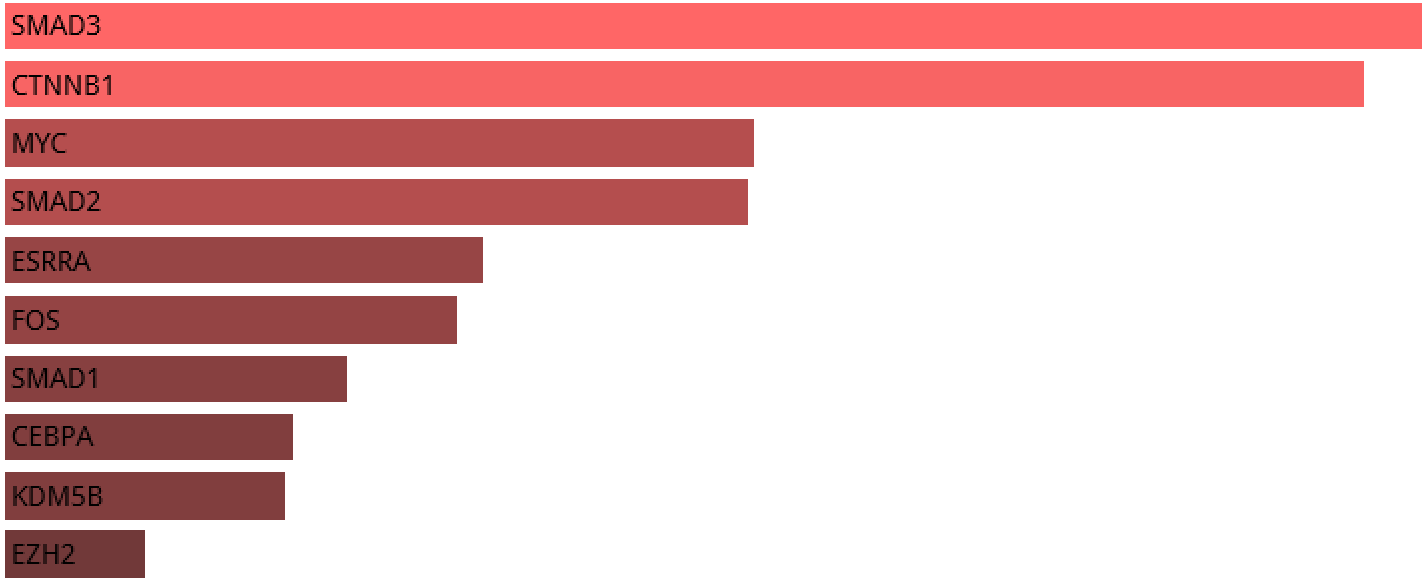


**11,878 fixed human-specific insertions**

Transcription Factor PPIs 7979 genes

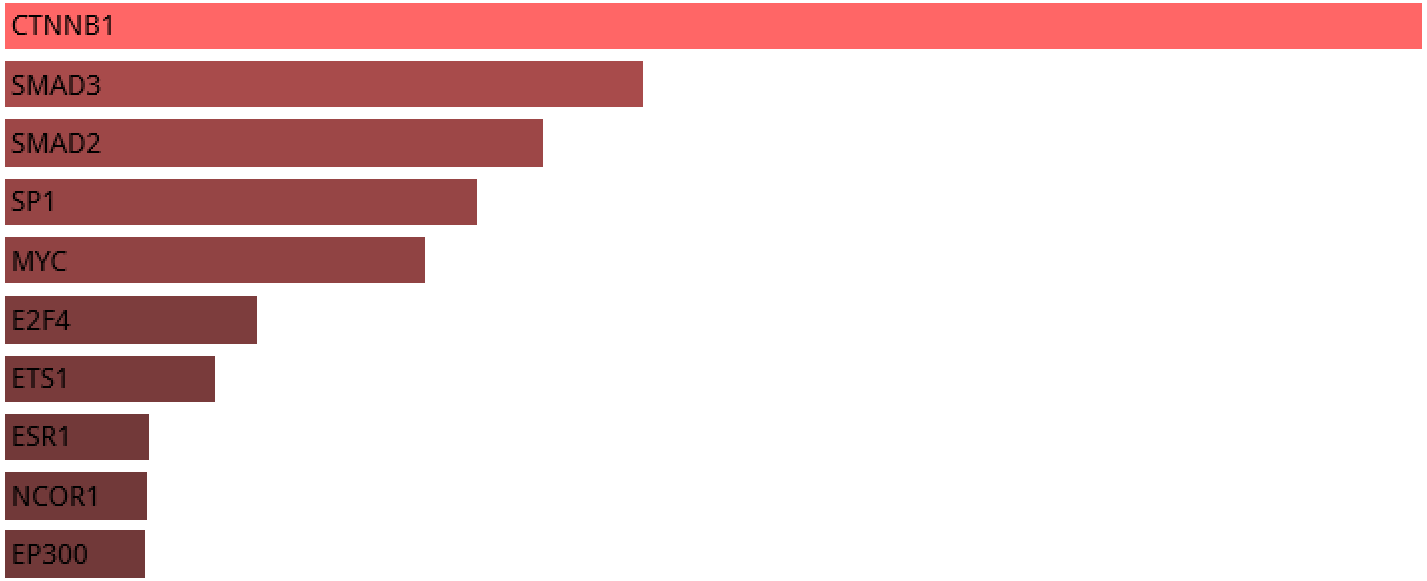

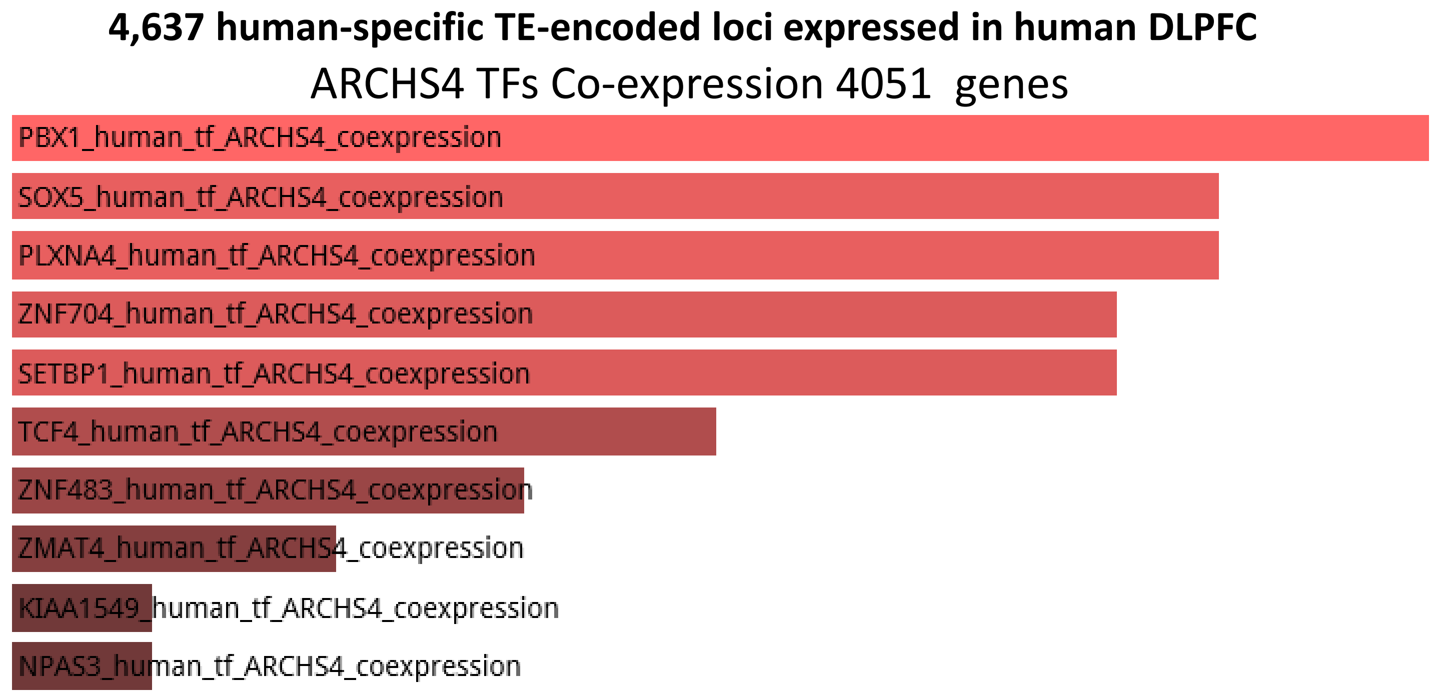
4,637 human-specific TE-encoded loci expressed in human DLPFC
ARCHS4 TFs Co-expression 4051 genes
PBX1_human_tf_ARCHS4_coexpression
SOX5_human_tf_ARCHS4_coexpression
PLXNA4_human_tf_ARCHS4_coexpression
ZNF704_human_tf_ARCHS4_coexpression
SETBP1_human_tf_ARCHS4_coexpression
TCF4_human_tf_ARCHS4_coexpression
ZNF483_human_tf_ARCHS4_coexpression
ZMAT4_human_tf_ARCHS4_coexpression
KIAA1549_human_tf_ARCHS4_coexpression
NPAS3_human_tf_ARCHS4_coexpression

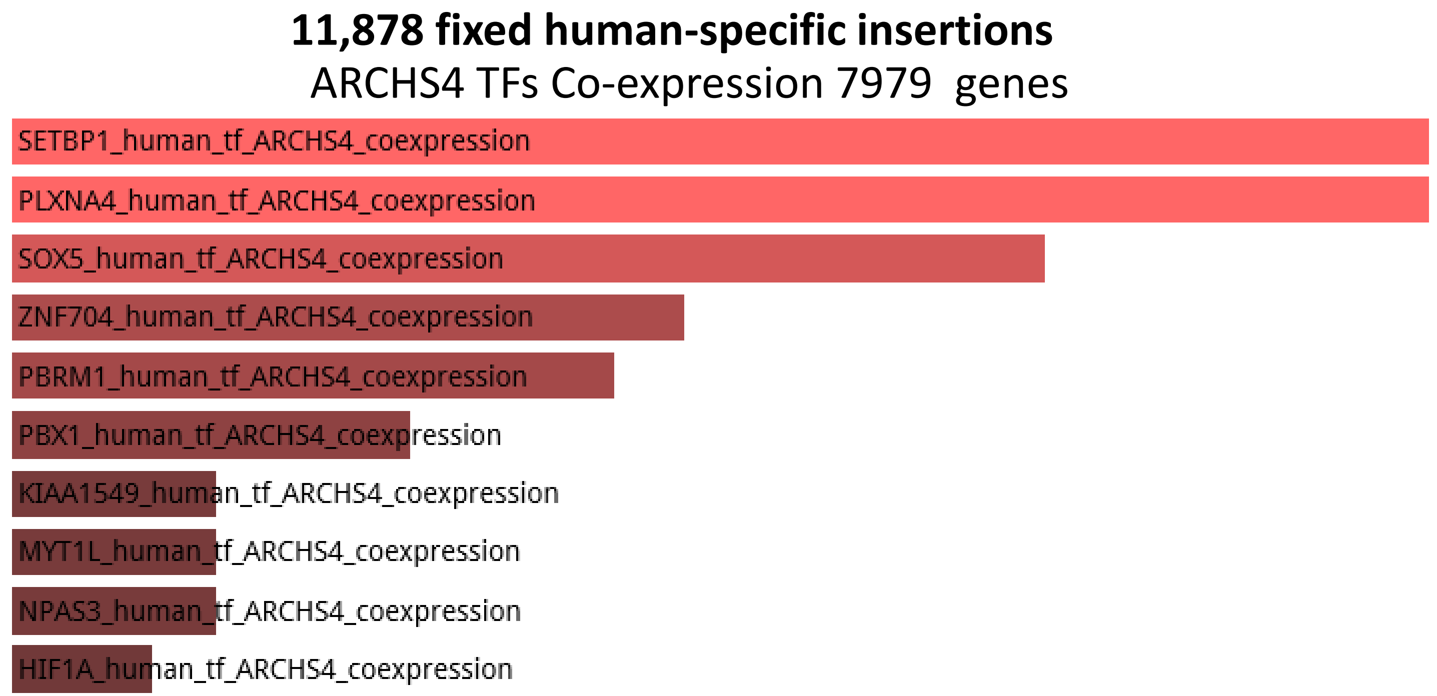
11,878 fixed human-specific insertions
ARCHS4 TFs Co-expression 7979 genes
SETBP1_human_tf_ARCHS4_coexpression
PLXNA4_human_tf_ARCHS4_coexpression
SOX5_human_tf_ARCHS4_coexpression
ZNF704_human_tf_ARCHS4_coexpression
PBRM1_human_tf_ARCHS4_coexpression
PBX1_human_tf_ARCHS4_coexpression
KIAA1549_human_tf_ARCHS4_coexpression
MYT1L_human_tf_ARCHS4_coexpression
NPAS3_human_tf_ARCHS4_coexpression
HIF1A_human_tf_ARCHS4_coexpression

#### 4,637 human-specific TE-encoded loci expressed in human DLPFC

ESCAPE 4051 genes

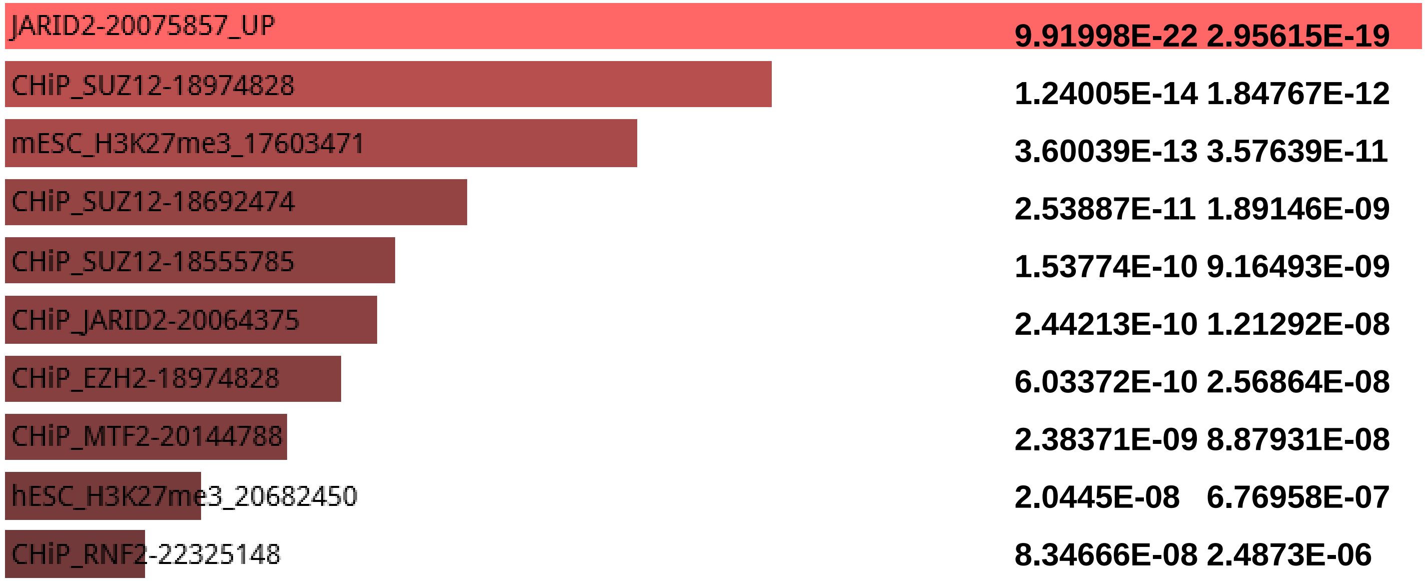


#### 11,878 fixed human-specific insertions

ESCAPE 7979 genes

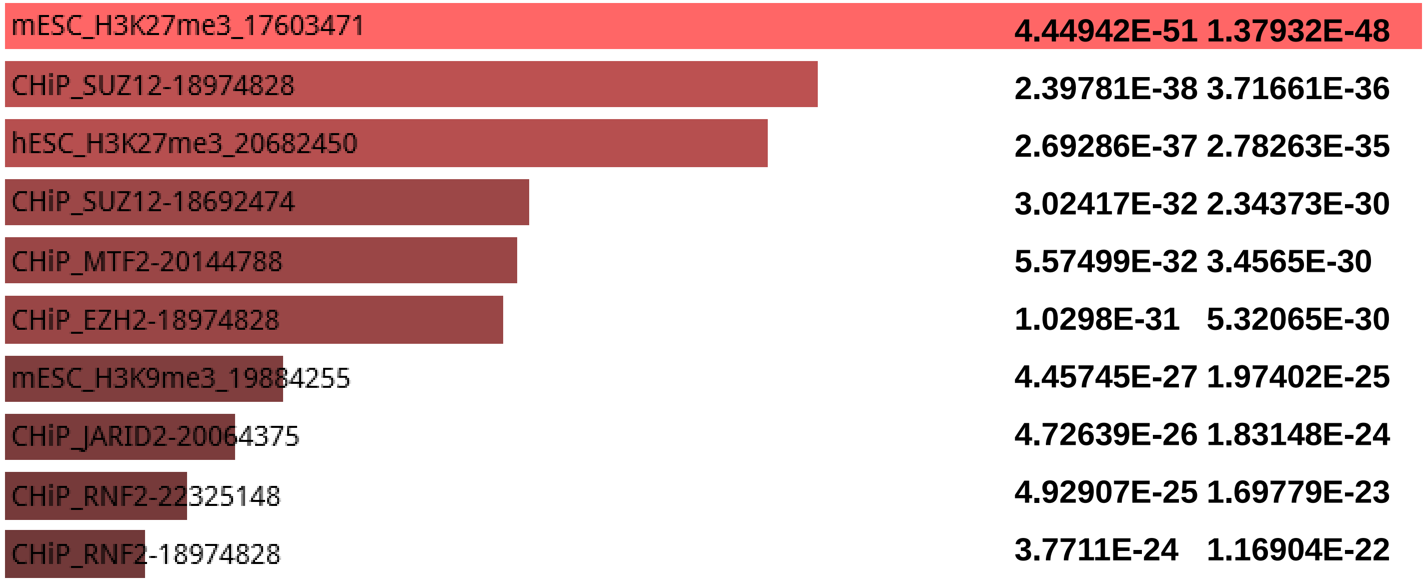

**7,599 duplicated regions in the human genome**

### 7,599 duplicated regions in the human genome

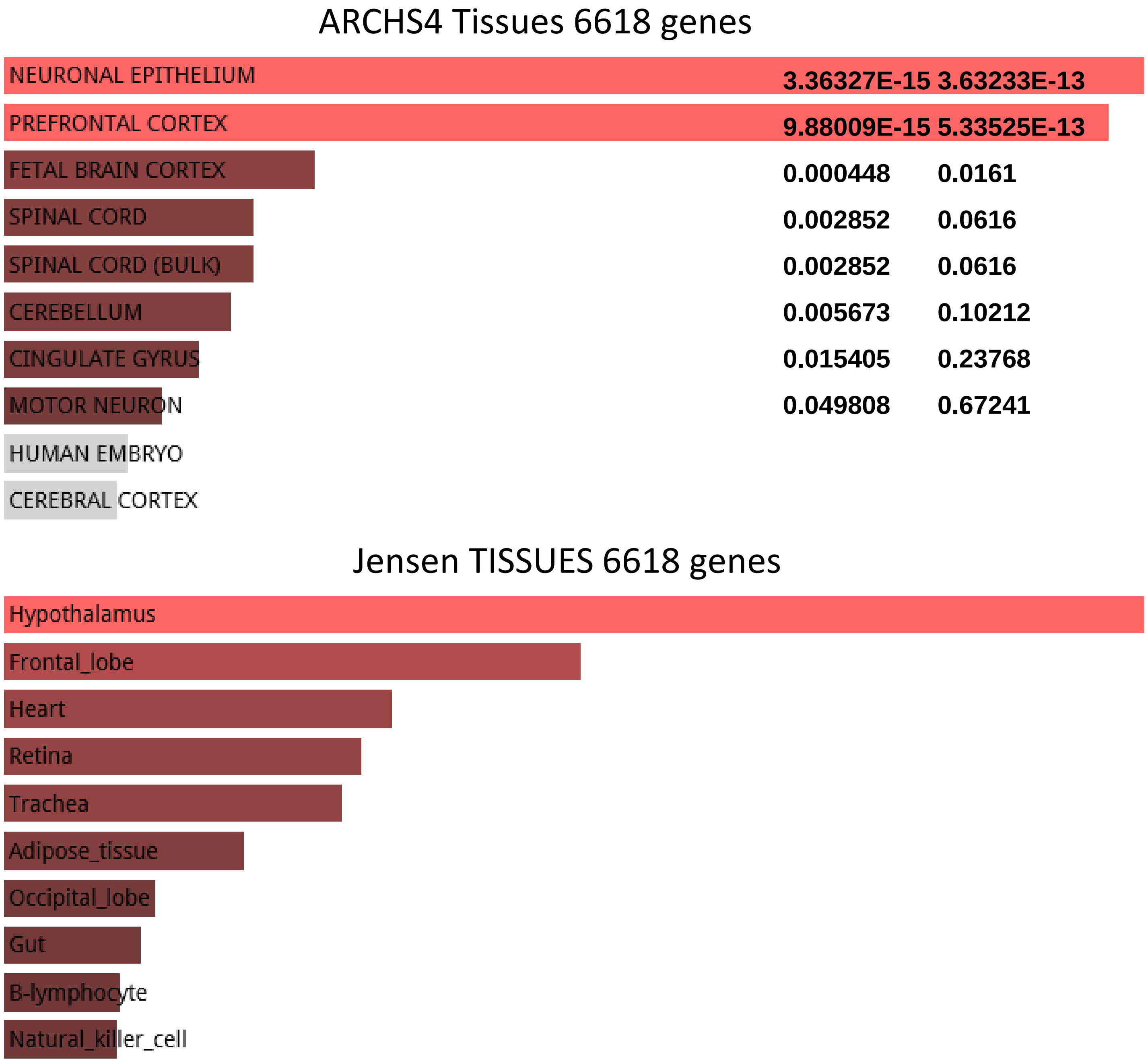

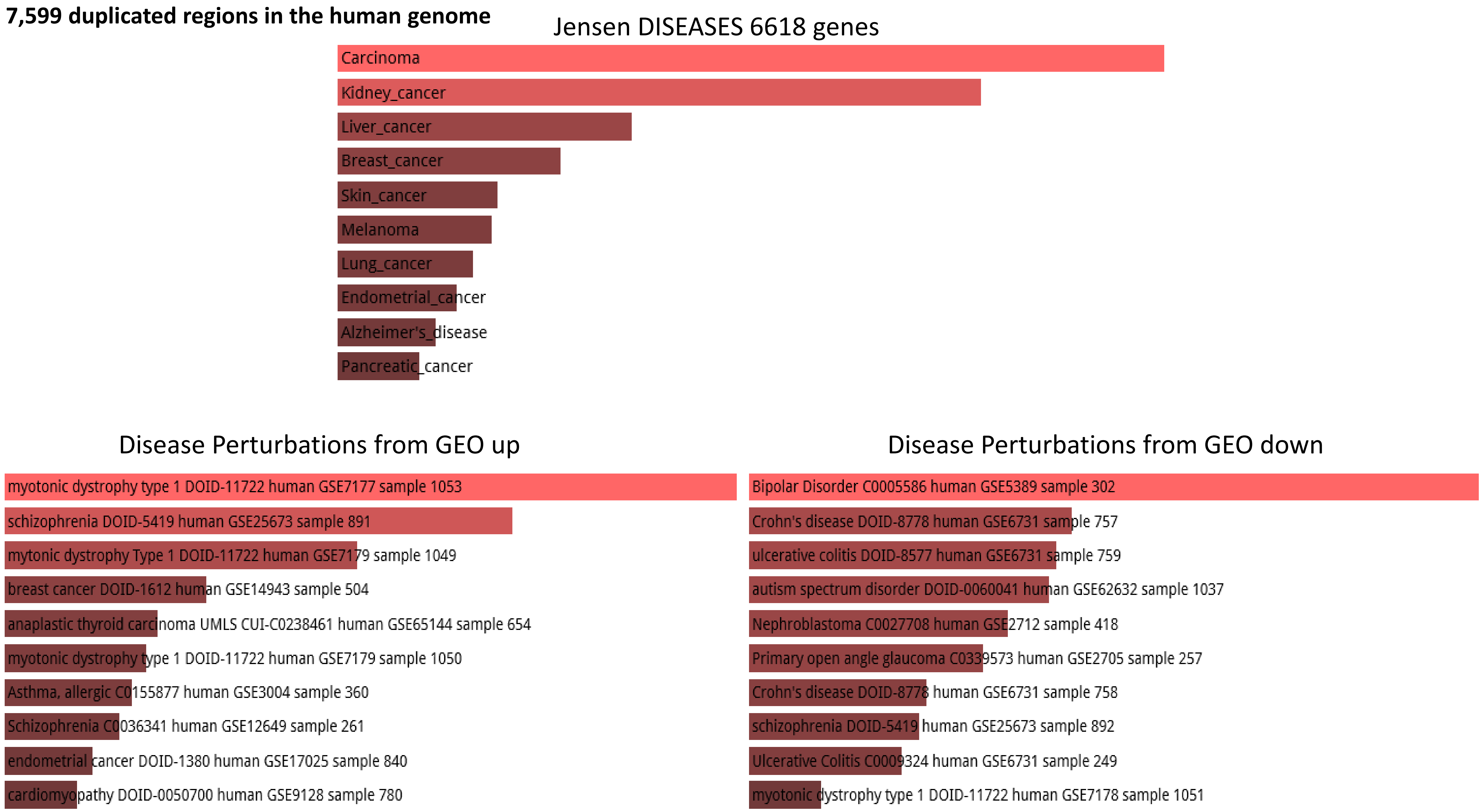
7,599 duplicated regions in the human genome
Jensen DISEASES 6618 genes
Carcinoma
Kidney_cancer
Liver_cancer
Breast_cancer
Skin_cancer
Melanoma
Lung_cancer
Endometrial_cancer
Alzheimer's_disease
Pancreatic_cancer
Disease Perturbations from GEO up
myotonic dystrophy type 1 DOID-11722 human GSE7177 sample 1053
schizophrenia DOID-5419 human GSE25673 sample 891
mytonic dystrophy Type 1 DOID-11722 human GSE7179 sample 1049
breast cancer DOID-1612 human GSE14943 sample 504
anaplastic thyroid carcinoma UMLS CUI-C0238461 human GSE65144 sample 654
myotonic dystrophy type 1 DOID-11722 human GSE7179 sample 1050
Asthma, allergic C0155877 human GSE3004 sample 360
Schizophrenia C0036341 human GSE12649 sample 261
endometrial cancer DOID-1380 human GSE17025 sample 840
cardiomyopathy DOID-0050700 human GSE9128 sample 780
Disease Perturbations from GEO down
Bipolar Disorder C0005586 human GSE5389 sample 302
Crohn's disease DOID-8778 human GSE6731 sample 757
ulcerative colitis DOID-8577 human GSE6731 sample 759
autism spectrum disorder DOID-0060041 human GSE62632 sample 1037
Nephroblastoma C0027708 human GSE2712 sample 418
Primary open angle glaucoma C0339573 human GSE2705 sample 257
Crohn's disease DOID-8778 human GSE6731 sample 758
schizophrenia DOID-5419 human GSE25673 sample 892
Ulcerative Colitis C0009324 human GSE6731 sample 249
myotonic dystrophy type 1 DOID-11722 human GSE7178 sample 1051

### 5,883 fixed human-specific deletions

### 5,883 fixed human-specific deletions

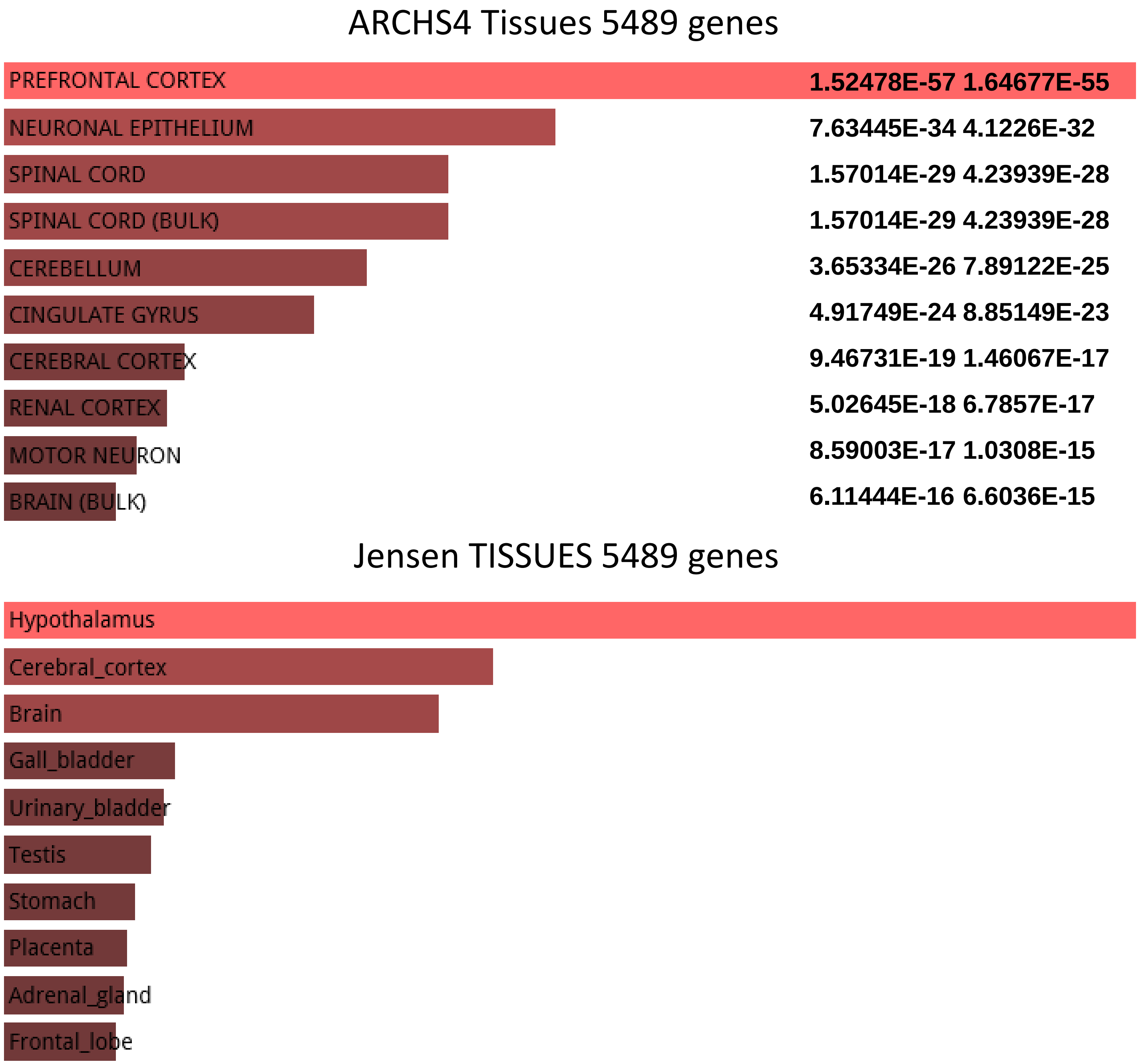

#### 5,883 fixed human-specific deletions

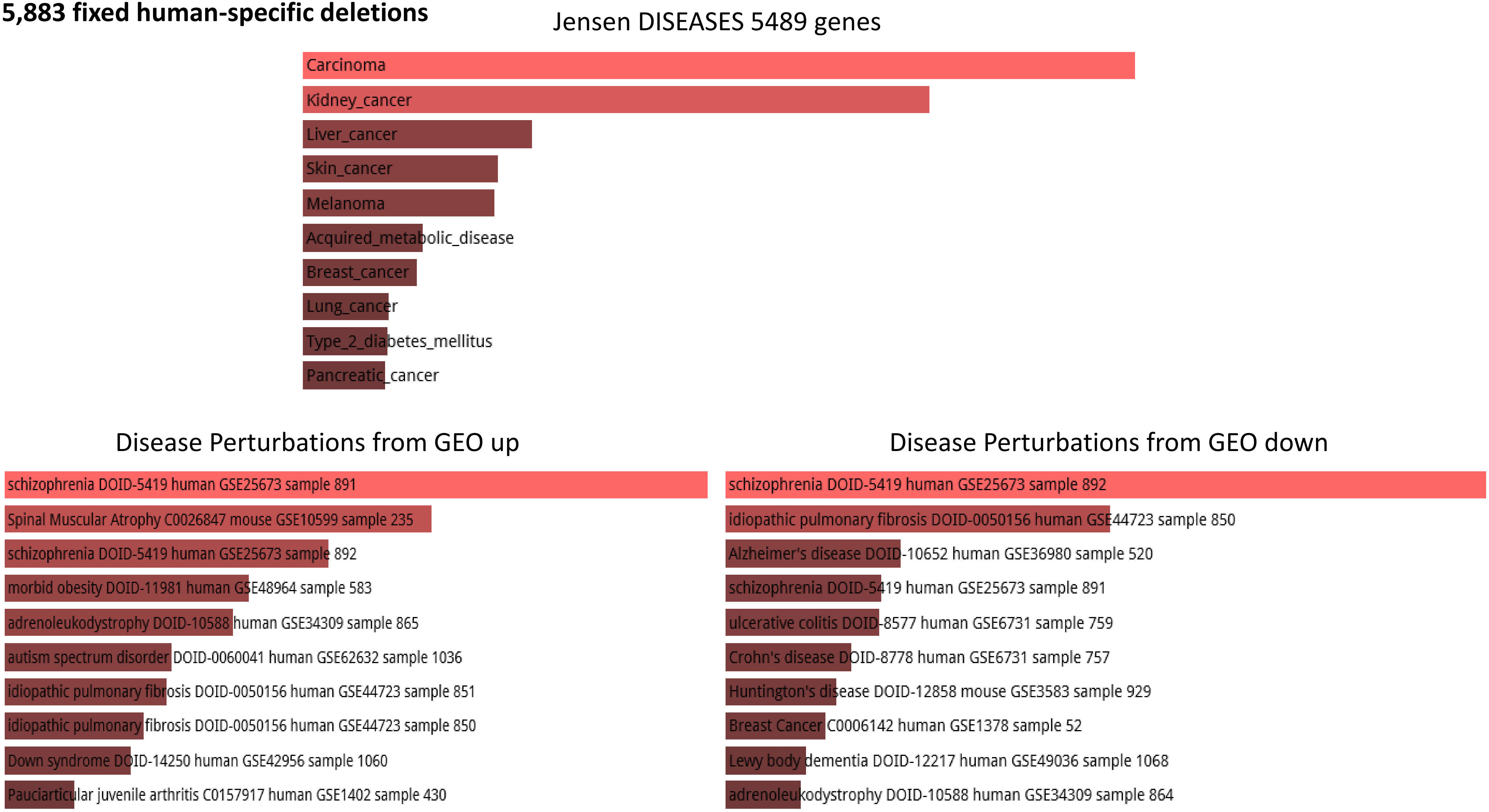

### 4,875 human-specific STR expansions

#### 4,875 human-specific STR expansions

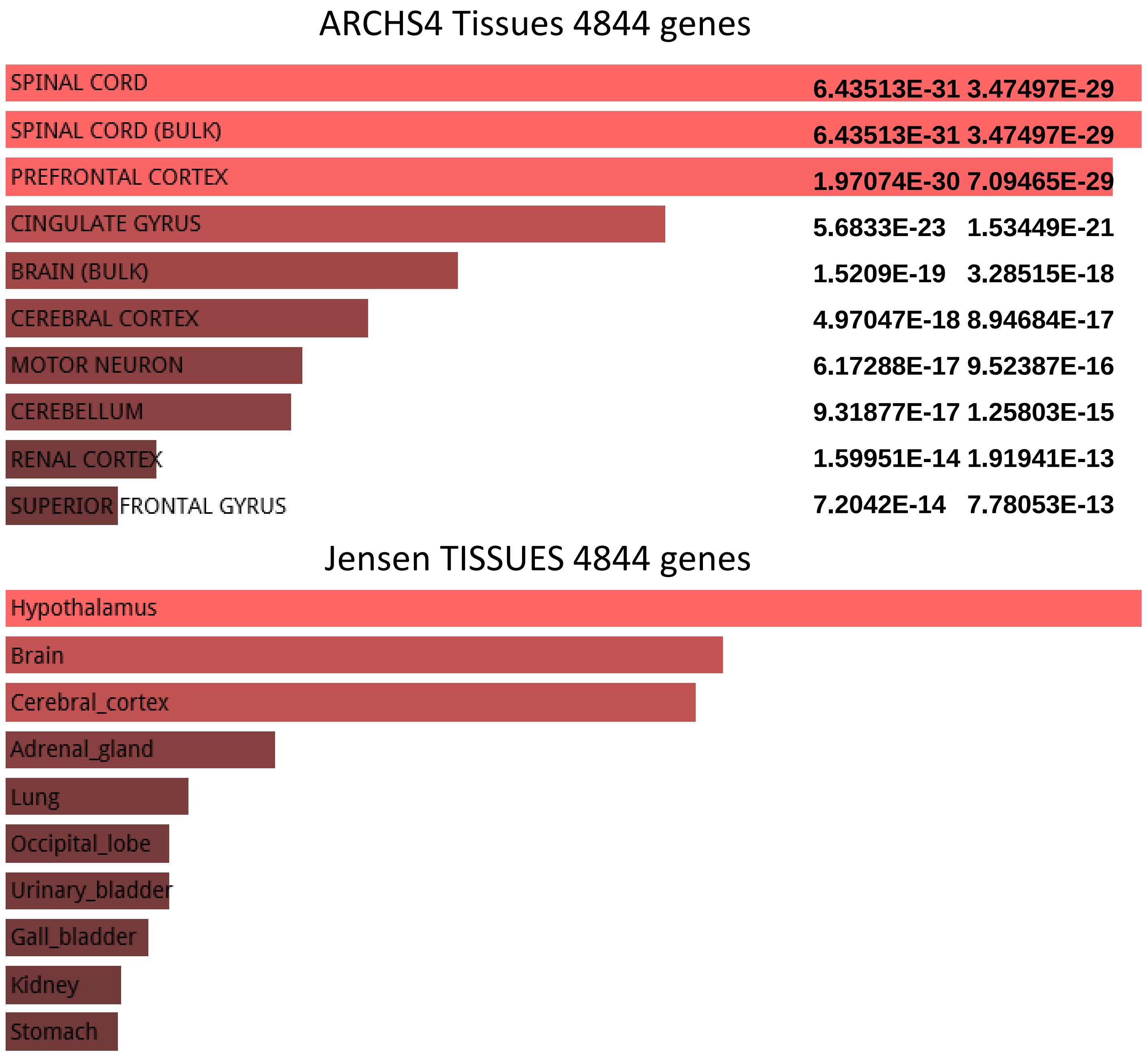

#### 4,875 human-specific STR expansions

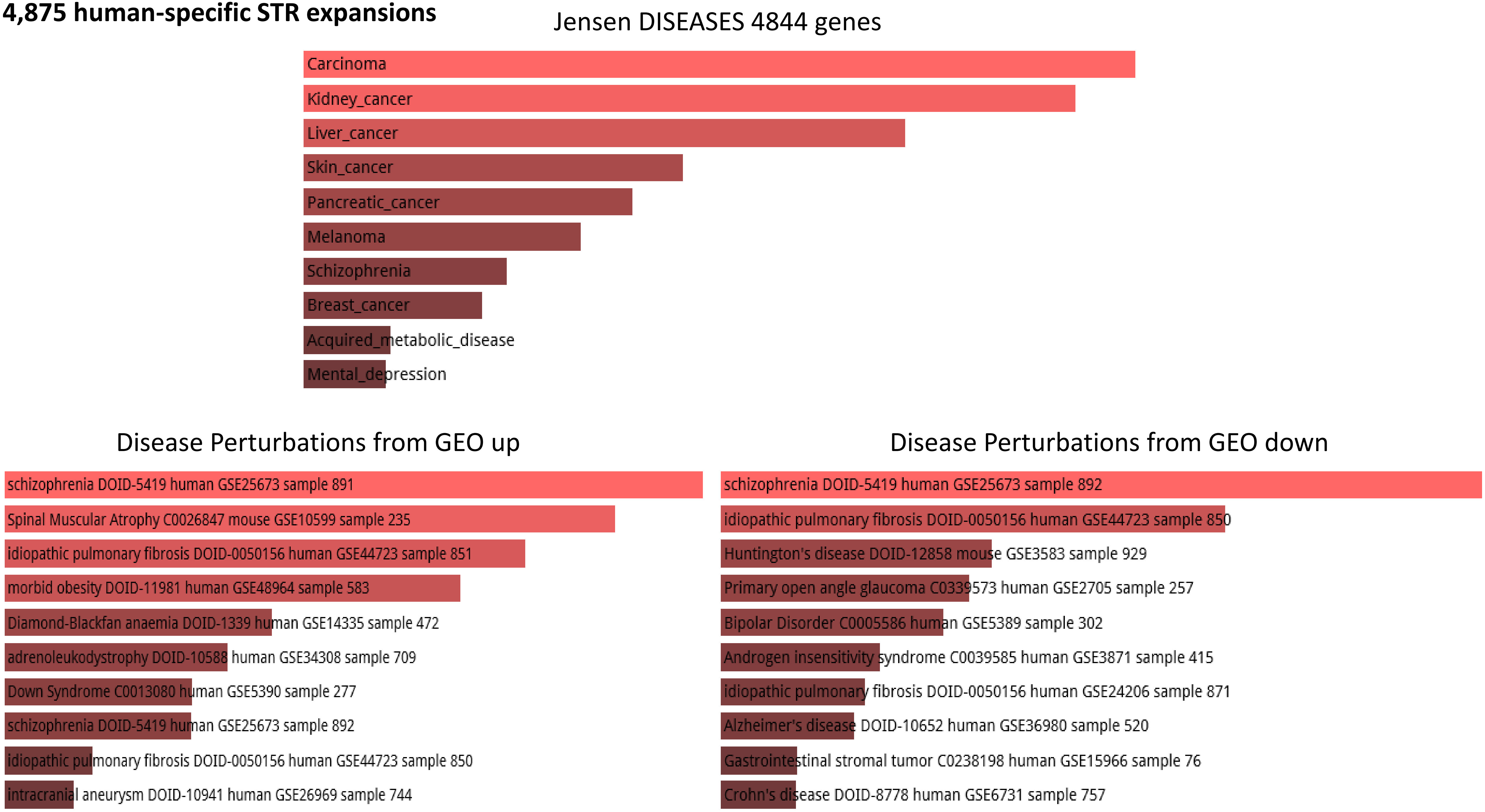

**3,538 human-specific ace-DHS**

**3,538 human-specific ace-DHS**

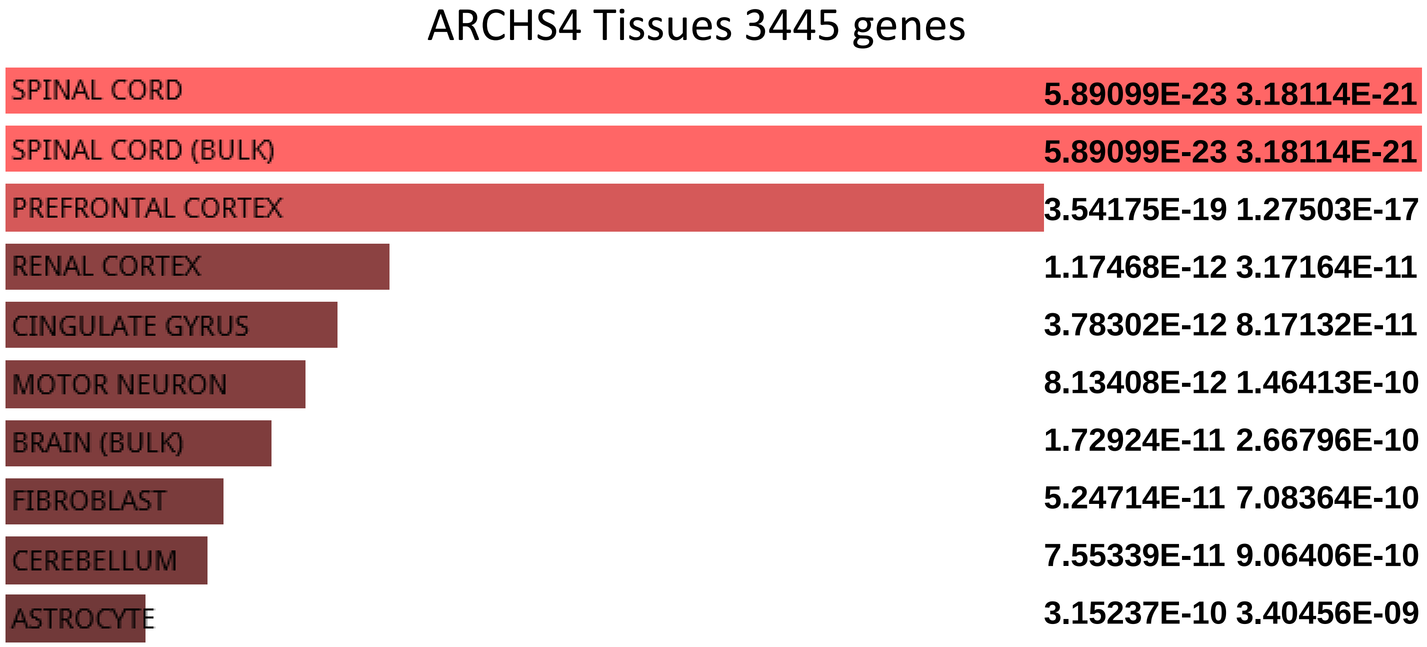


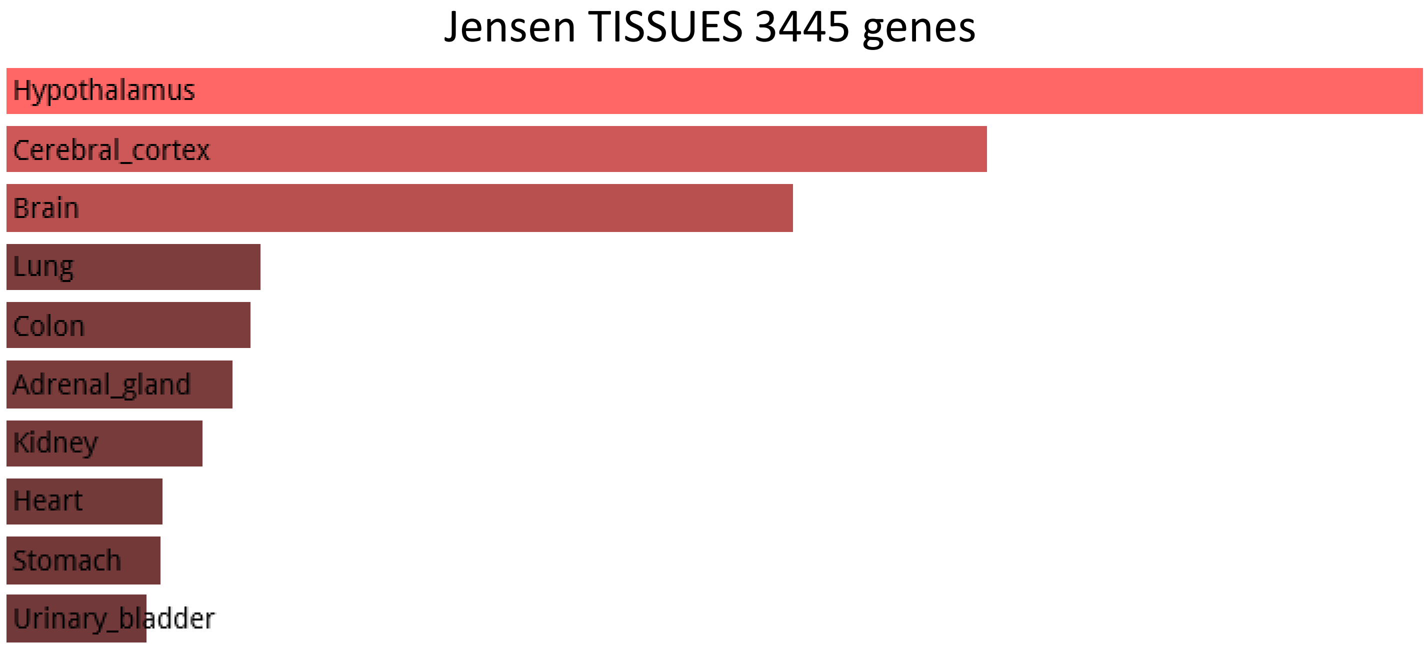

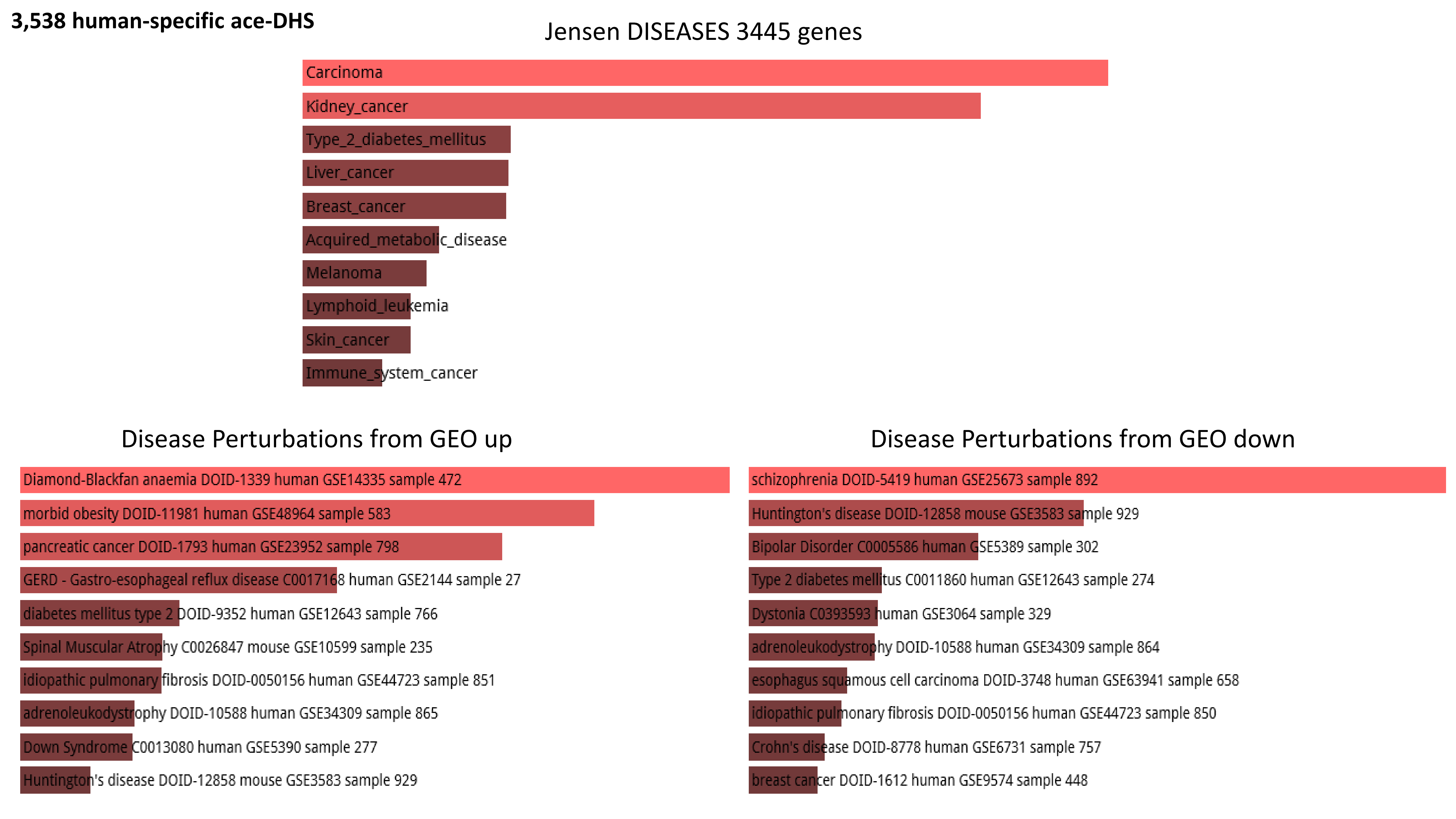
3,538 human-specific ace-DHS
Jensen DISEASES 3445 genes
Carcinoma
Kidney_cancer
Type_2_diabetes_mellitus
Liver_cancer
Breast_cancer
Acquired_metabolic_disease
Melanoma
Lymphoid_leukemia
Skin_cancer
Immune_system_cancer
Disease Perturbations from GEO up
Diamond-Blackfan anaemia DOID-1339 human GSE14335 sample 472
morbid obesity DOID-11981 human GSE48964 sample 583
pancreatic cancer DOID-1793 human GSE23952 sample 798
GERD - Gastro-esophageal reflux disease C0017168 human GSE2144 sample 27
diabetes mellitus type 2 DOID-9352 human GSE12643 sample 766
Spinal Muscular Atrophy C0026847 mouse GSE10599 sample 235
idiopathic pulmonary fibrosis DOID-0050156 human GSE44723 sample 851
adrenoleukodystrophy DOID-10588 human GSE34309 sample 865
Down Syndrome C0013080 human GSE5390 sample 277
Huntington's disease DOID-12858 mouse GSE3583 sample 929
Disease Perturbations from GEO down
schizophrenia DOID-5419 human GSE25673 sample 892
Huntington's disease DOID-12858 mouse GSE3583 sample 929
Bipolar Disorder C0005586 human GSE5389 sample 302
Type 2 diabetes mellitus C0011860 human GSE12643 sample 274
Dystonia C0393593 human GSE3064 sample 329
adrenoleukodystrophy DOID-10588 human GSE34309 sample 864
esophagus squamous cell carcinoma DOID-3748 human GSE63941 sample 658
idiopathic pulmonary fibrosis DOID-0050156 human GSE44723 sample 850
Crohn's disease DOID-8778 human GSE6731 sample 757
breast cancer DOID-1612 human GSE9574 sample 448

**3,538 human-specific ace-DHS**

#### Allen Brain Atlas up 3445 genes

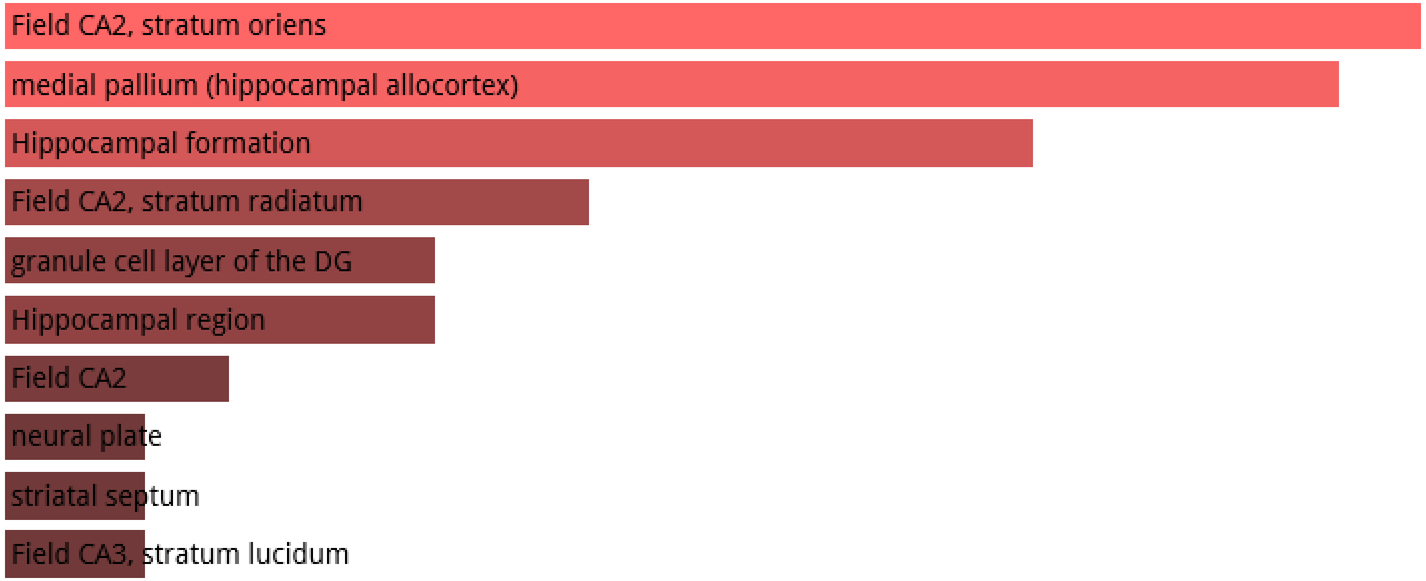


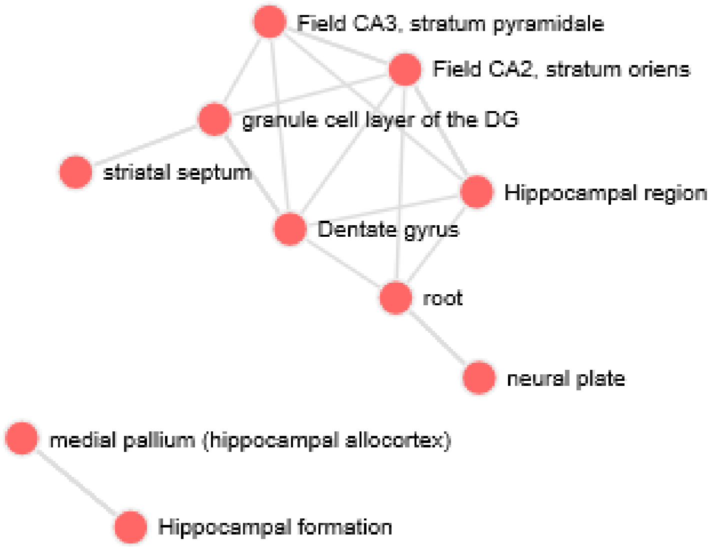

### 3,803 human-specific TFBS in hESC

**3,803 human-specific TFBS in hESC**

ARCHS4 Tissues 1087 genes

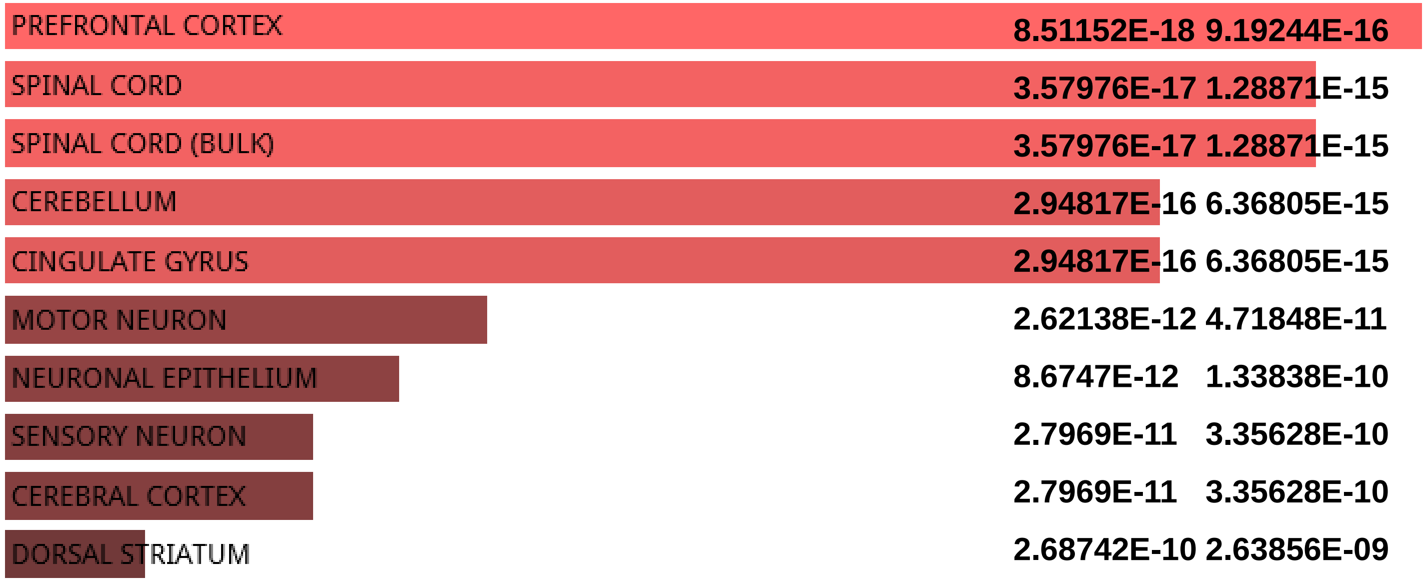


Jensen TISSUES 1087 genes

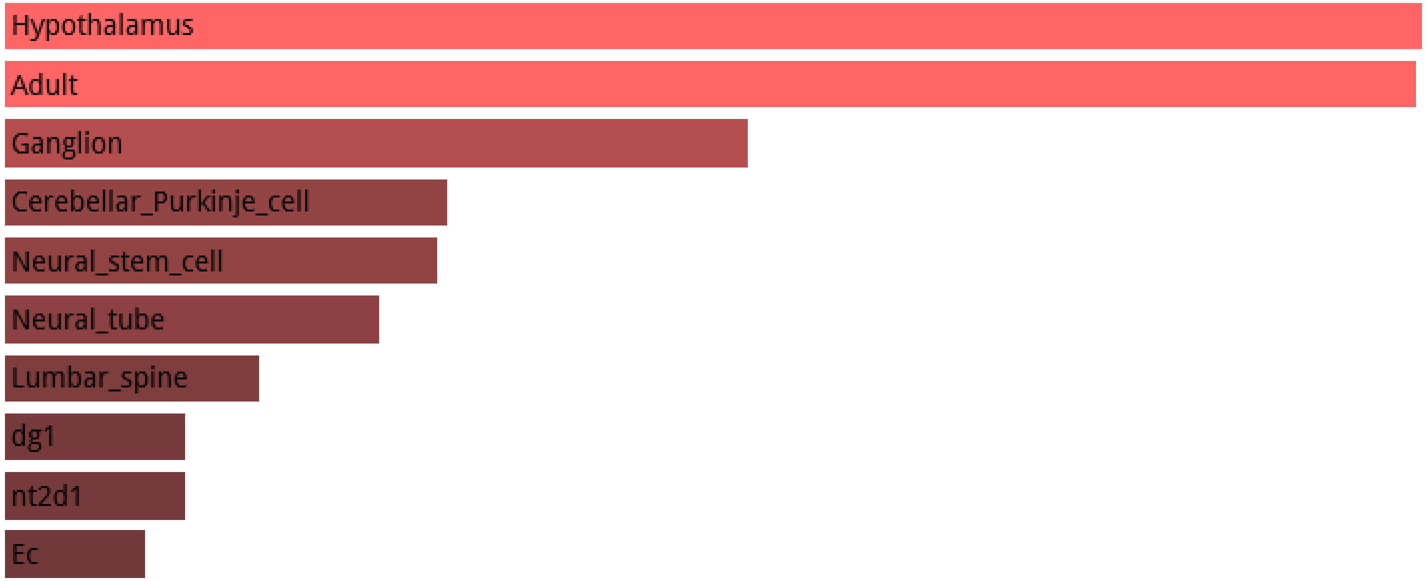

### 3,803 human-specific TFBS in hESC

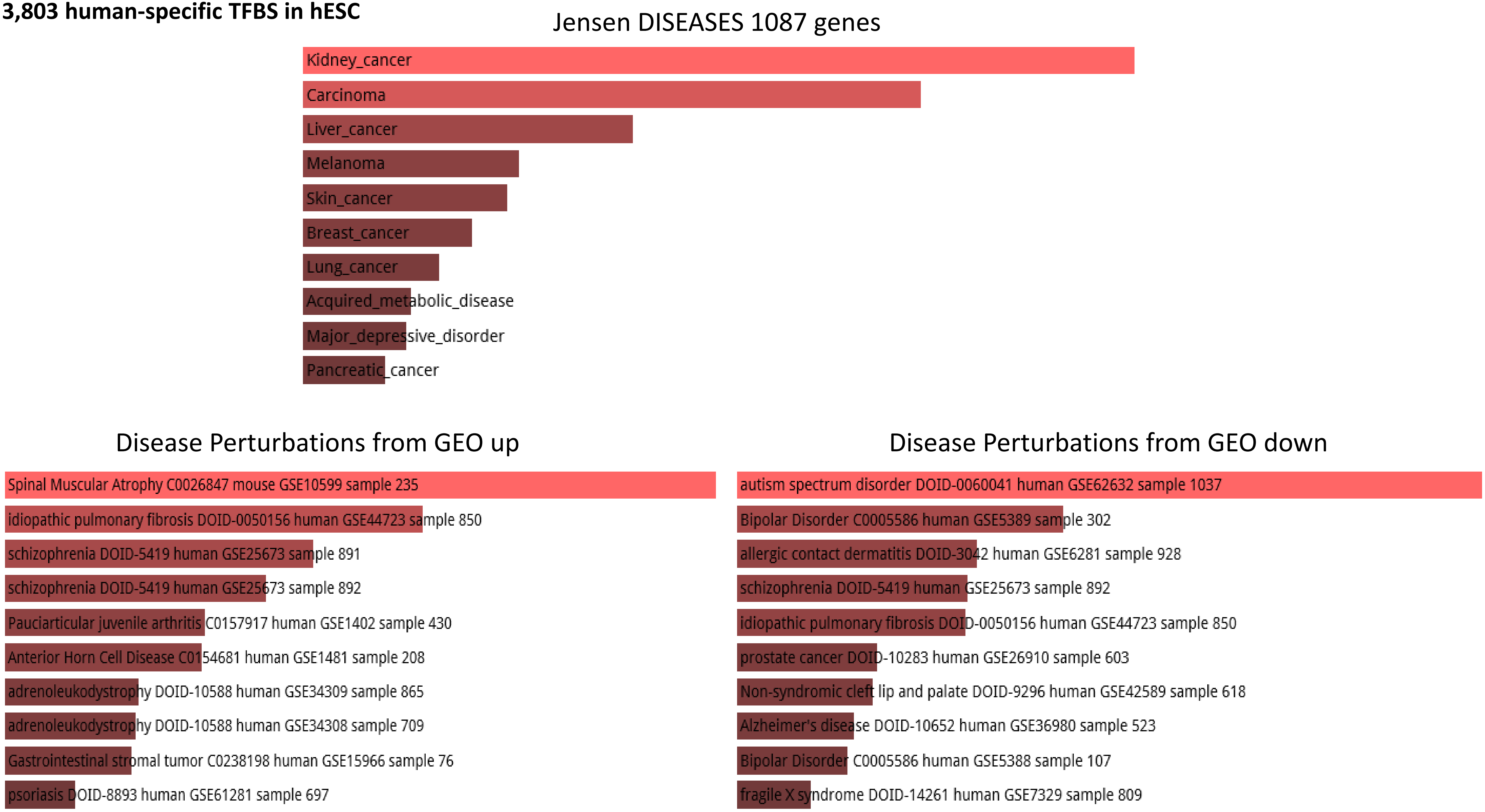

**4,249 fixed human-specific regulatory loci**

### 4,249 fixed human-specific regulatory loci

#### ARCHS4 Tissues 2810 genes

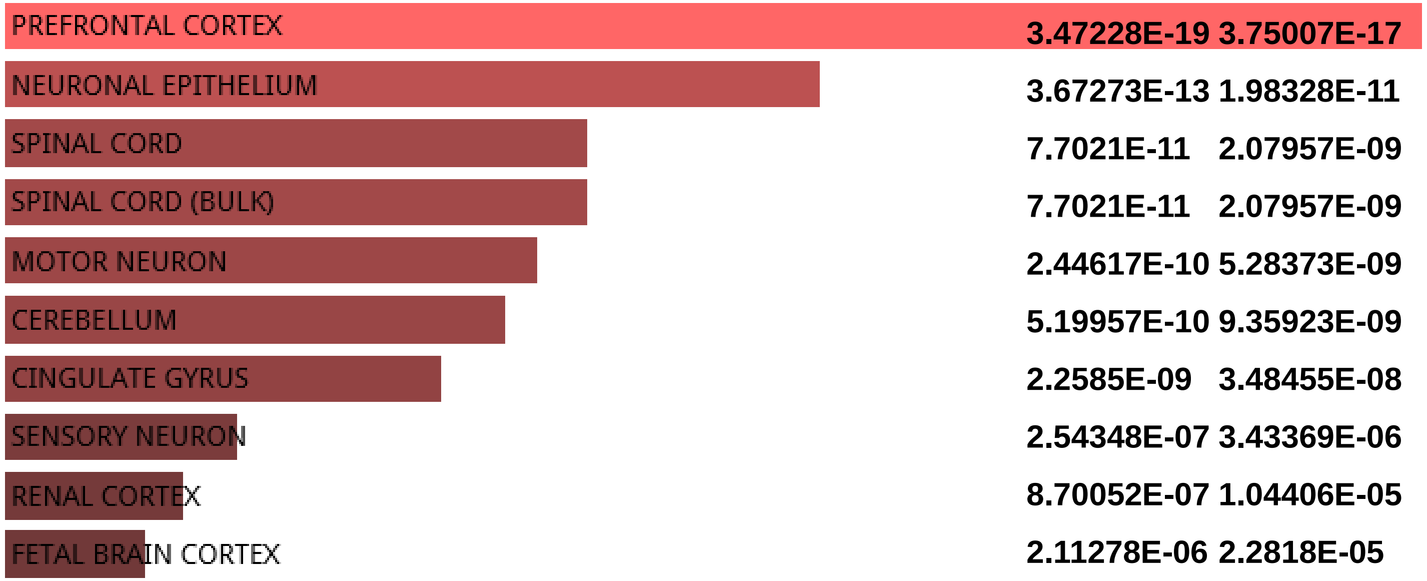


#### Jensen TISSUES 2810 genes

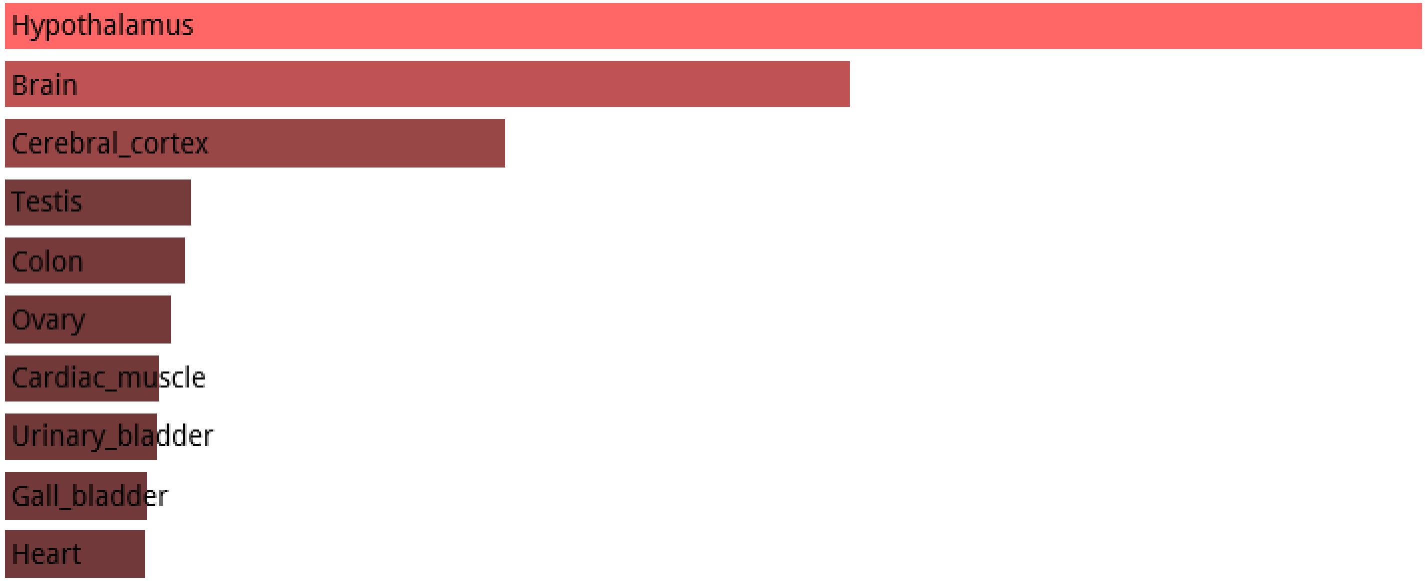

**4,249 fixed human-specific regulatory loci**

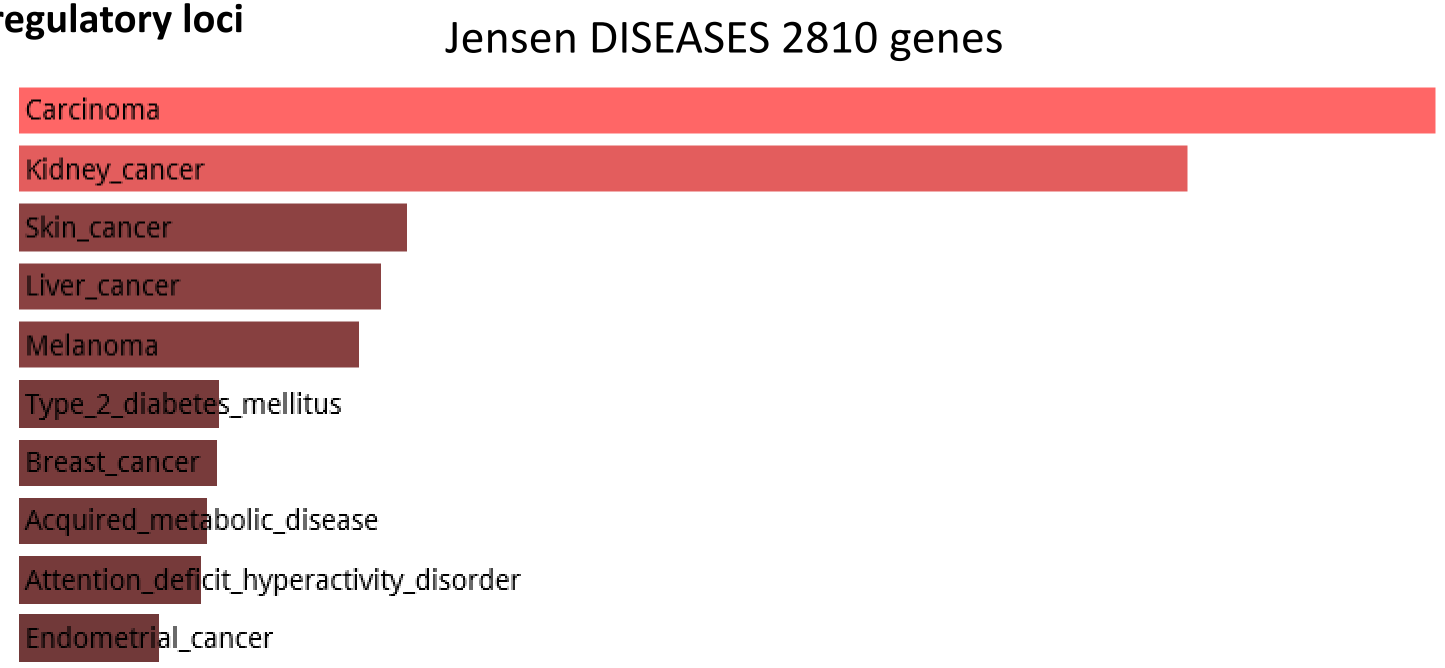


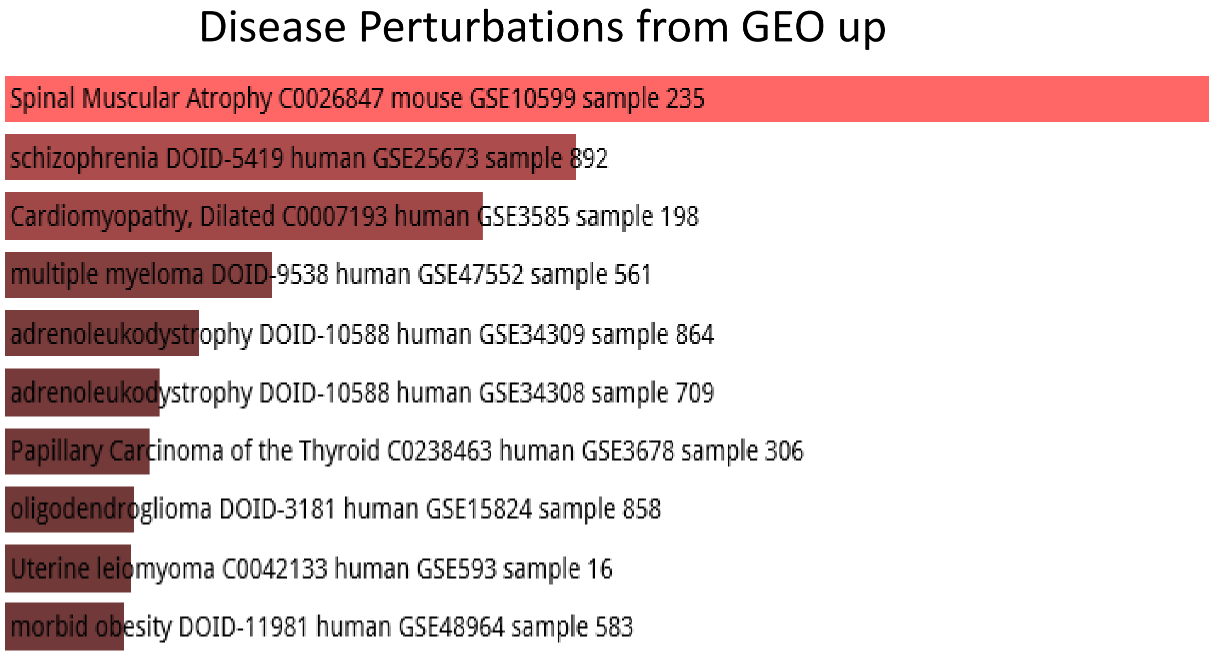


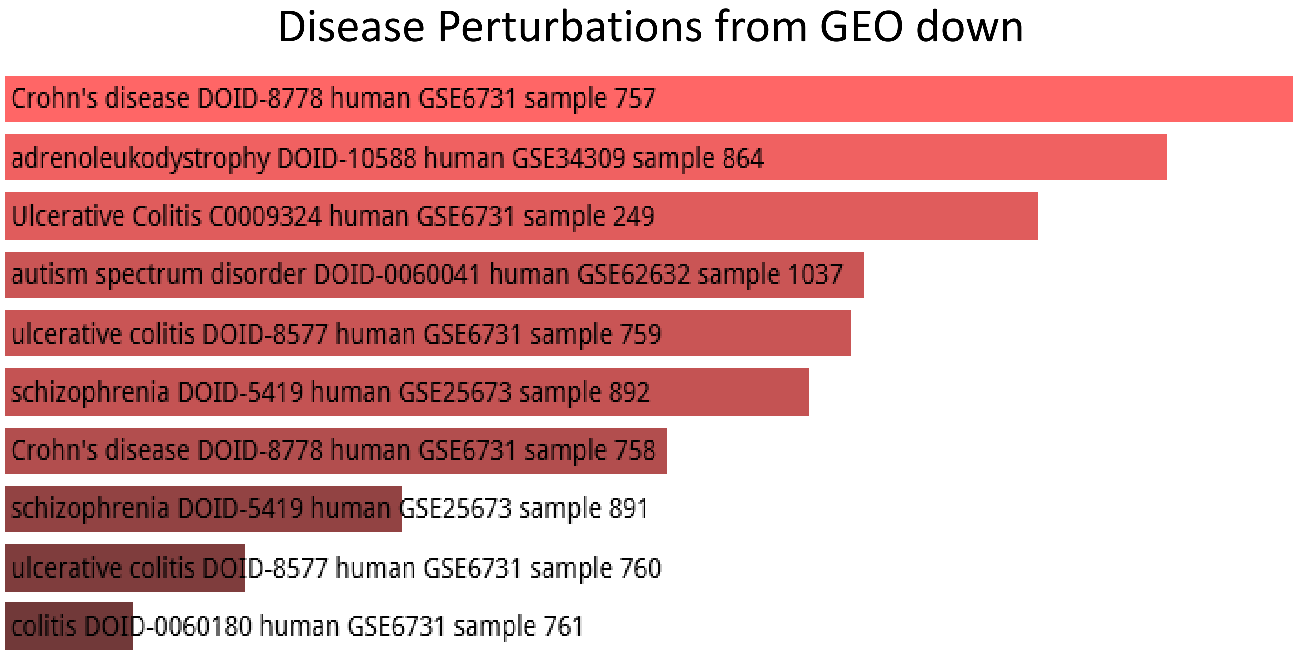

### 2,745 human accelerated regions

#### 2,745 human accelerated regions

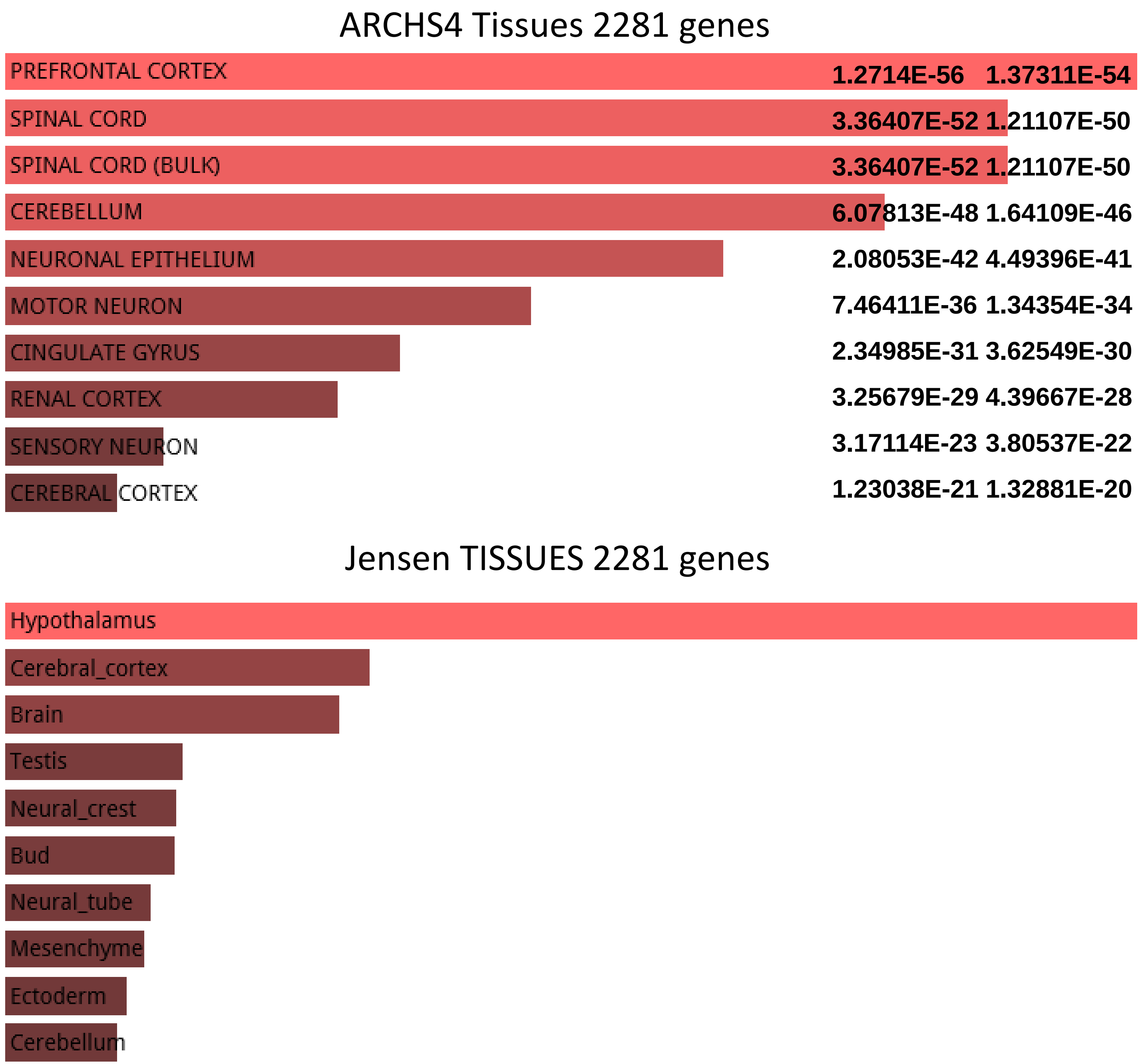

**2,745 human accelerated regions**

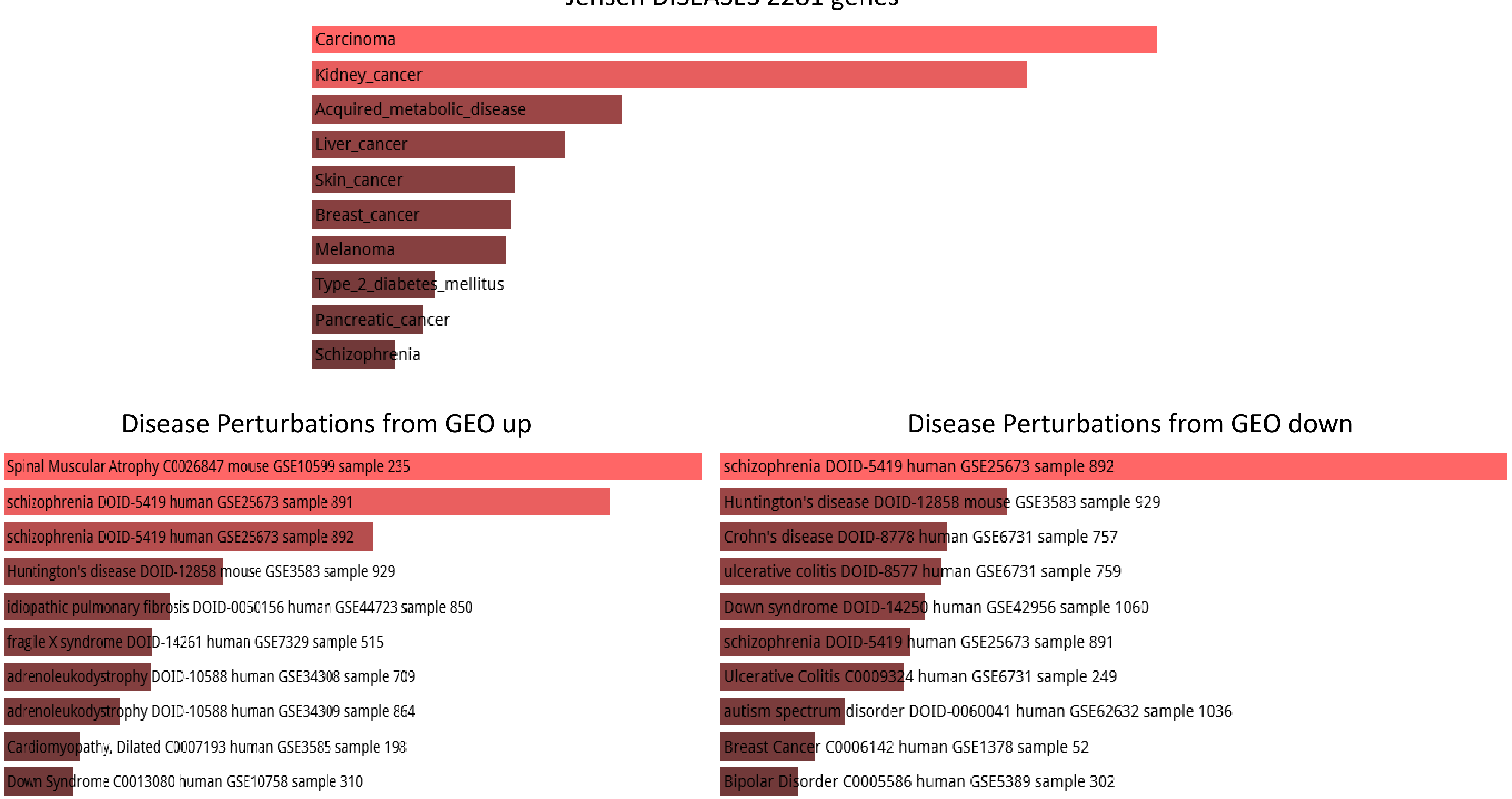

**1,932 fixed human-specific regulatory regions (hESC DHS)**

### 1,932 fixed human-specific regulatory regions (hESC DHS)

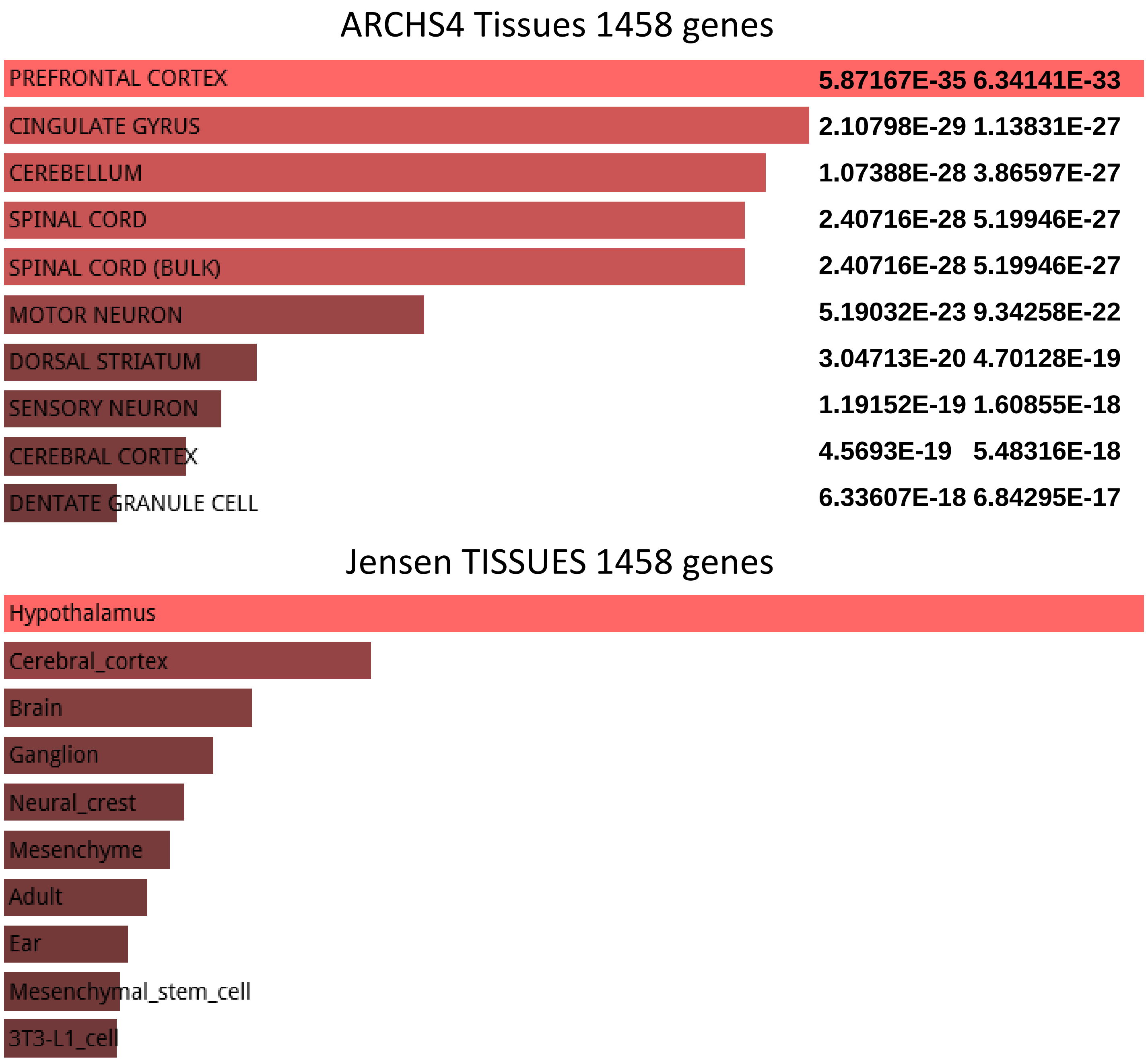

1,932 fixed human-specific regulatory regions (hESC DHS)

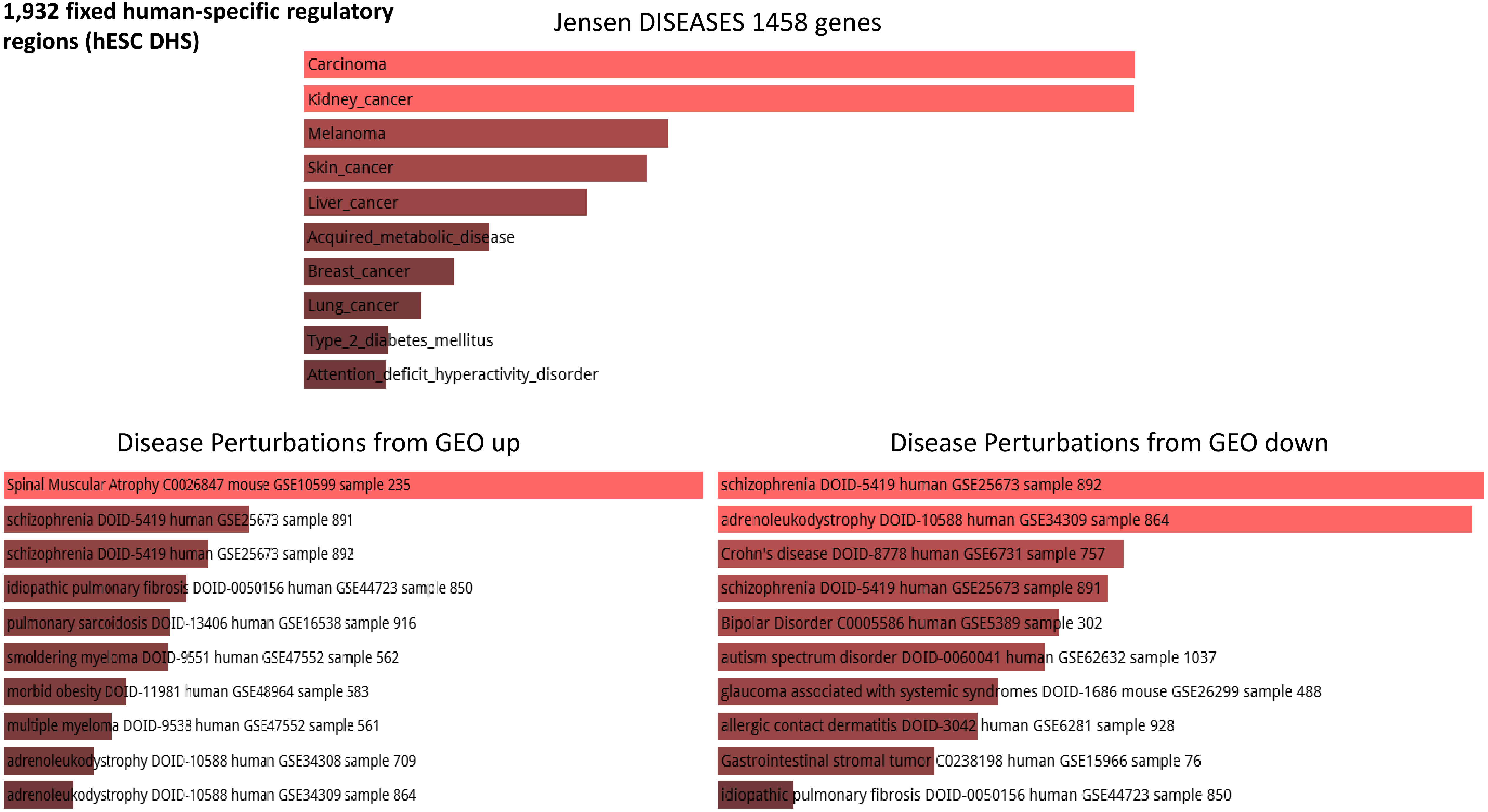

**1,000 human-biased cranial neural crest cells (CNCCs) enhancers**

### 1,000 human-biased cranial neural crest cells (CNCCs) enhancers

#### ARCHS4 Tissues 1439 genes

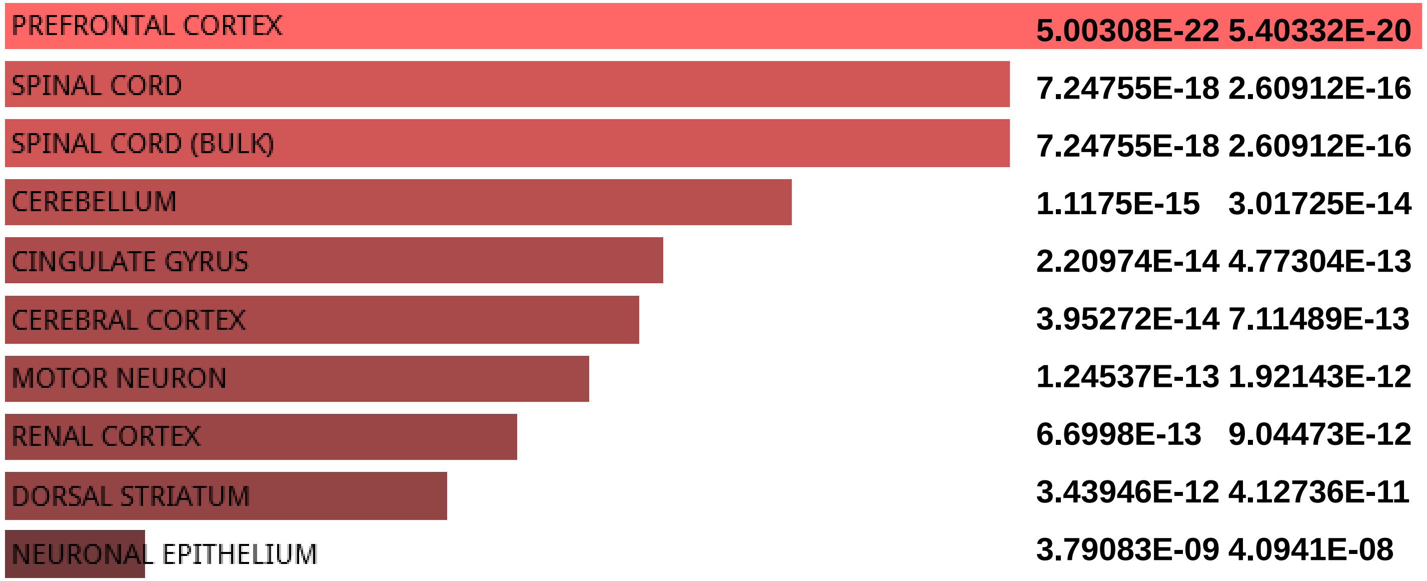


#### Jensen TISSUES 1439 genes

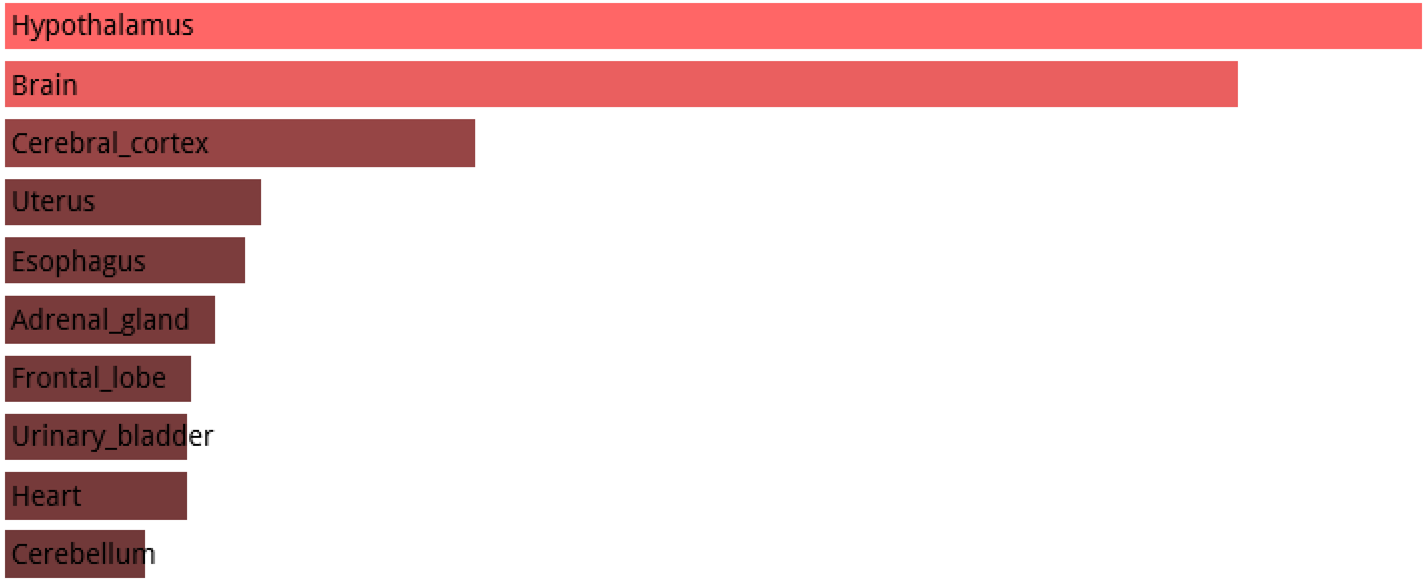

**1,000 human-biased cranial neural crest cells (CNCCs) enhancers**

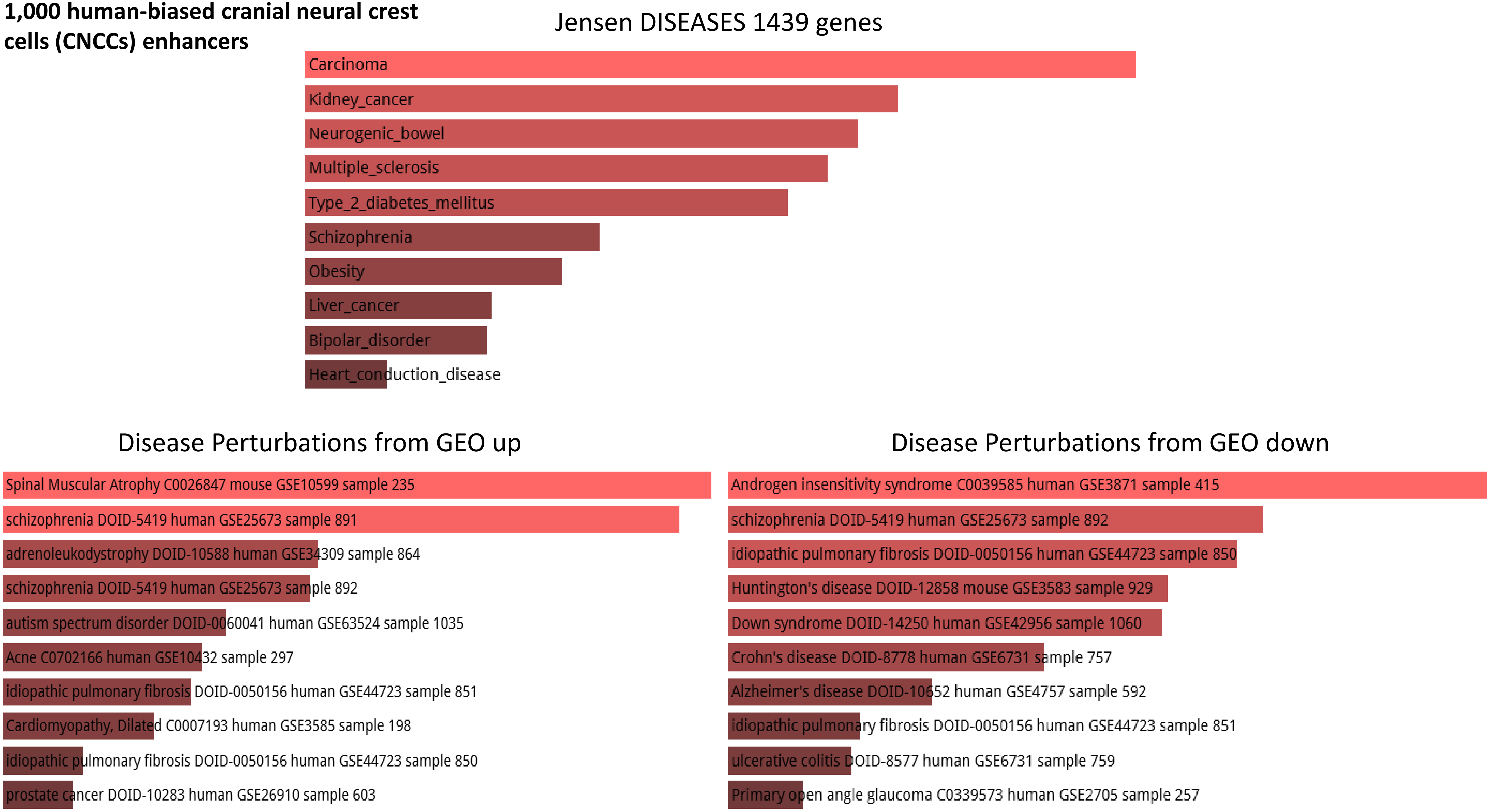

**1,000 chimp-biased cranial neural crest cells (CNCCs) enhancers**

**1,000 chimp-biased cranial neural crest cells (CNCCs) enhancers**

ARCHS4 Tissues 1445 genes

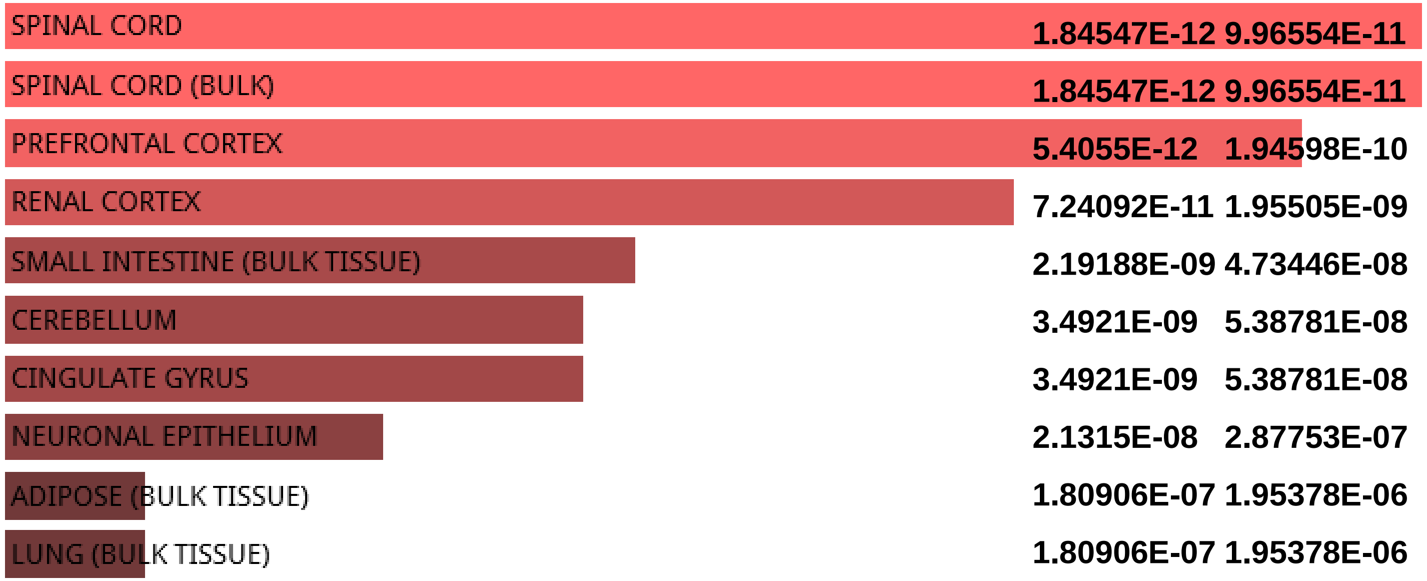


Jensen TISSUES 1445 genes

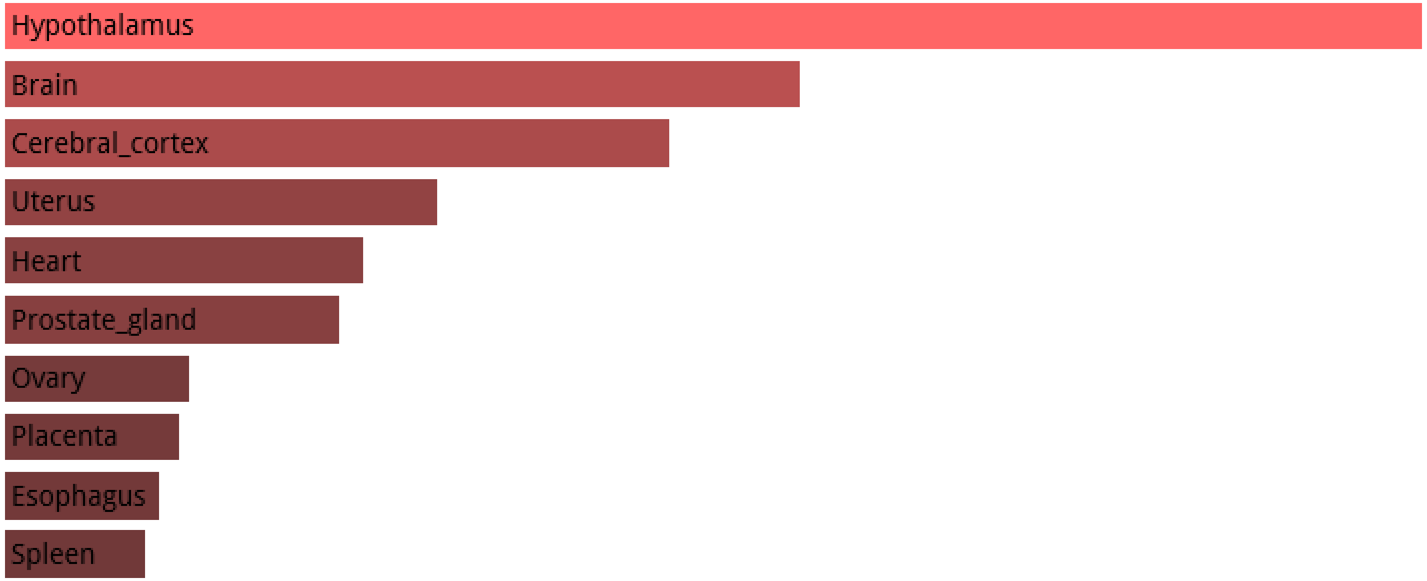

**1,000 chimp-biased cranial neural crest cells (CNCCs) enhancers**

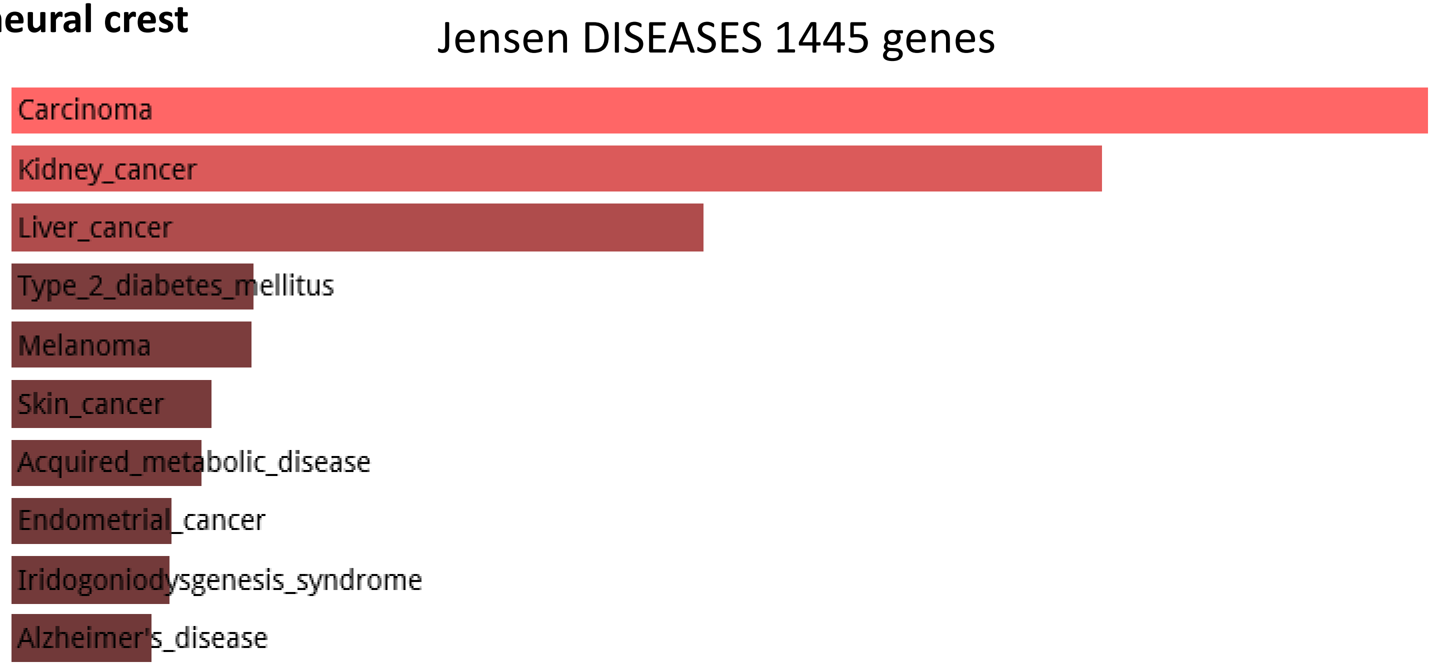


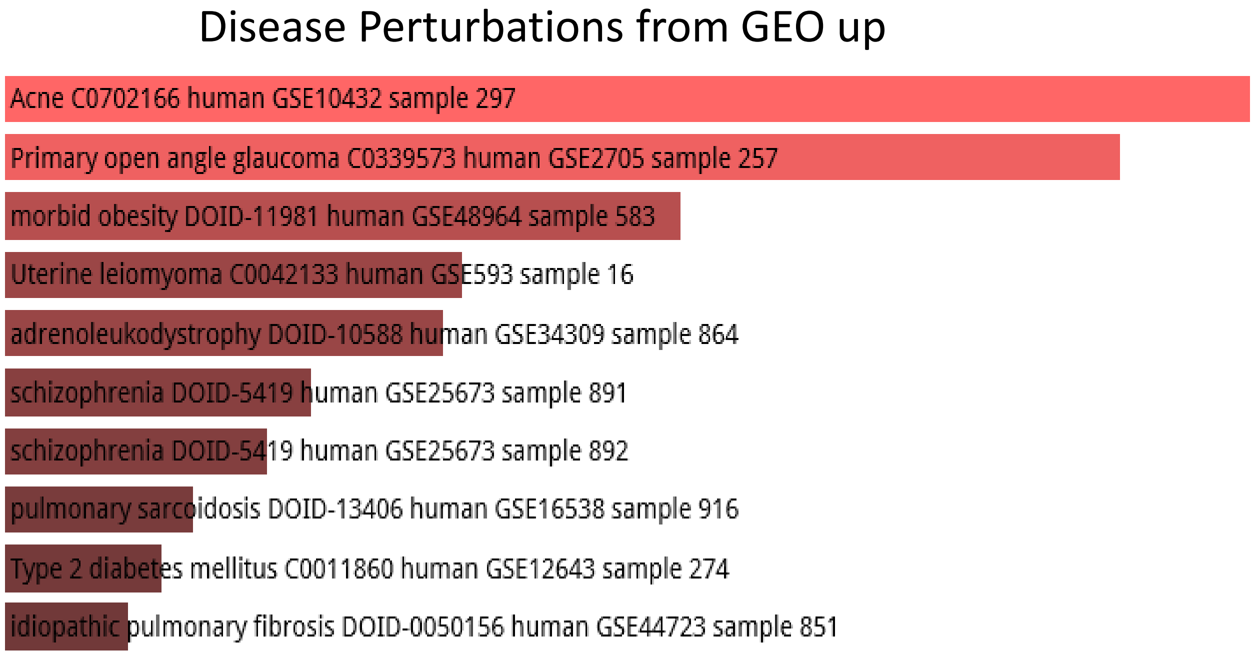


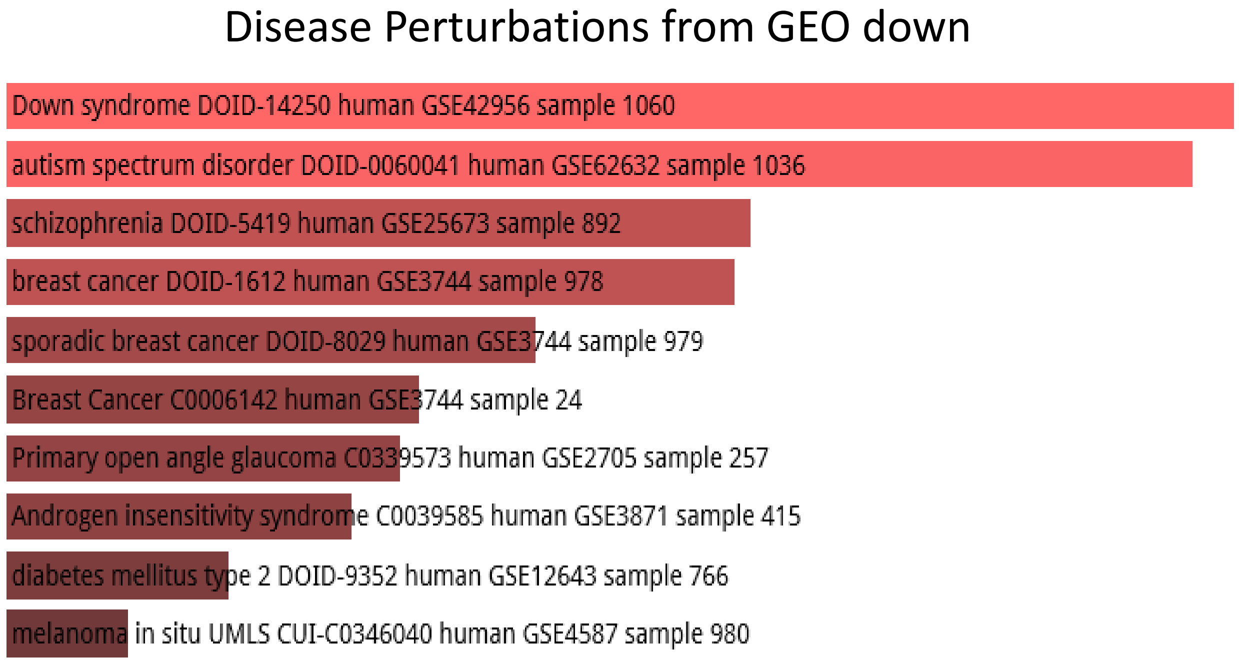

### 1,619 human-specific functional hESC enhancers

1,619 human-specific functional hESC enhancers

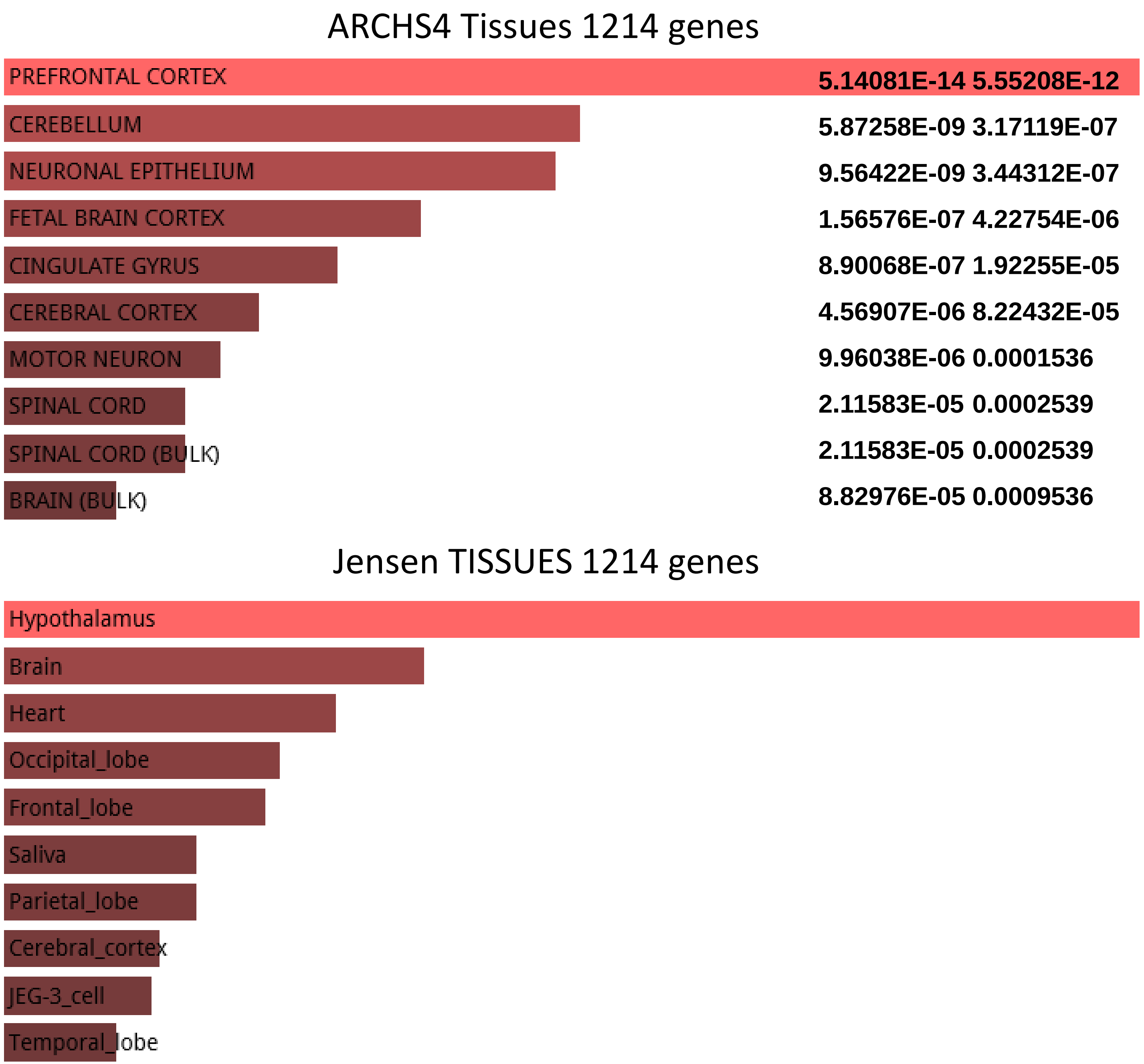

**1,619 human-specific functional hESC enhancers**

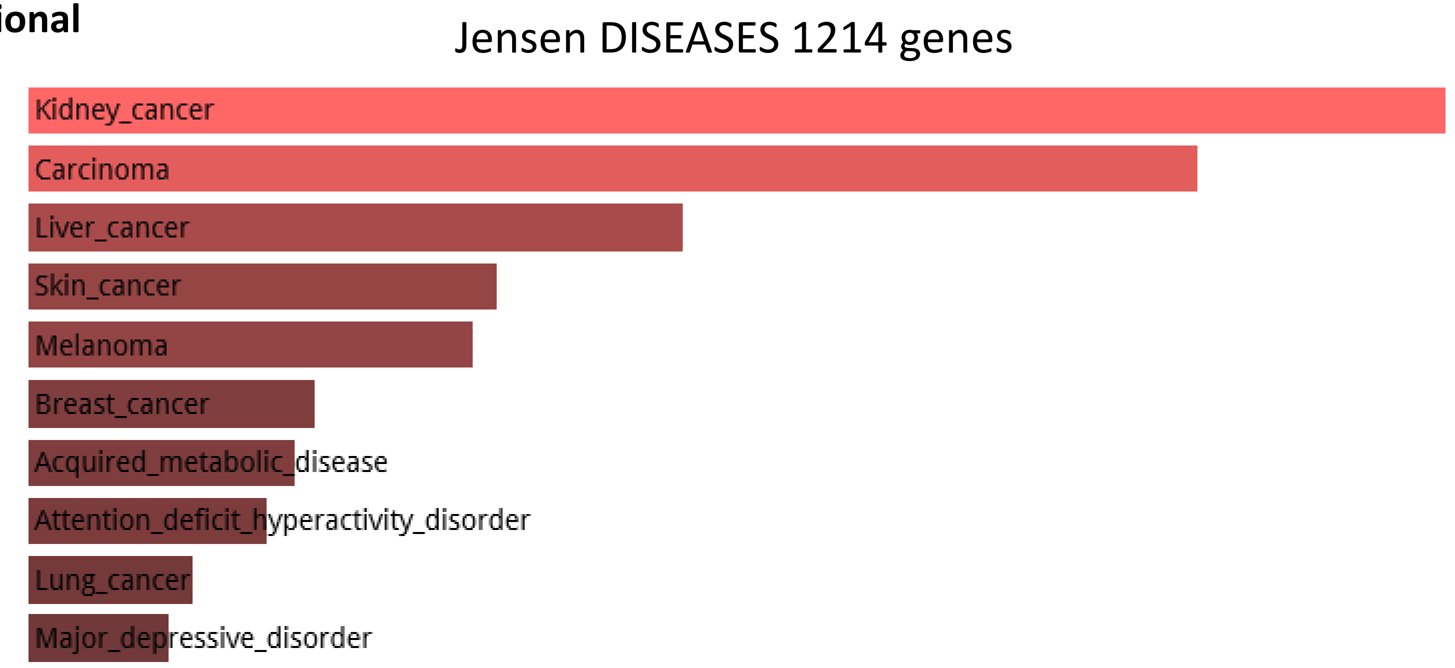


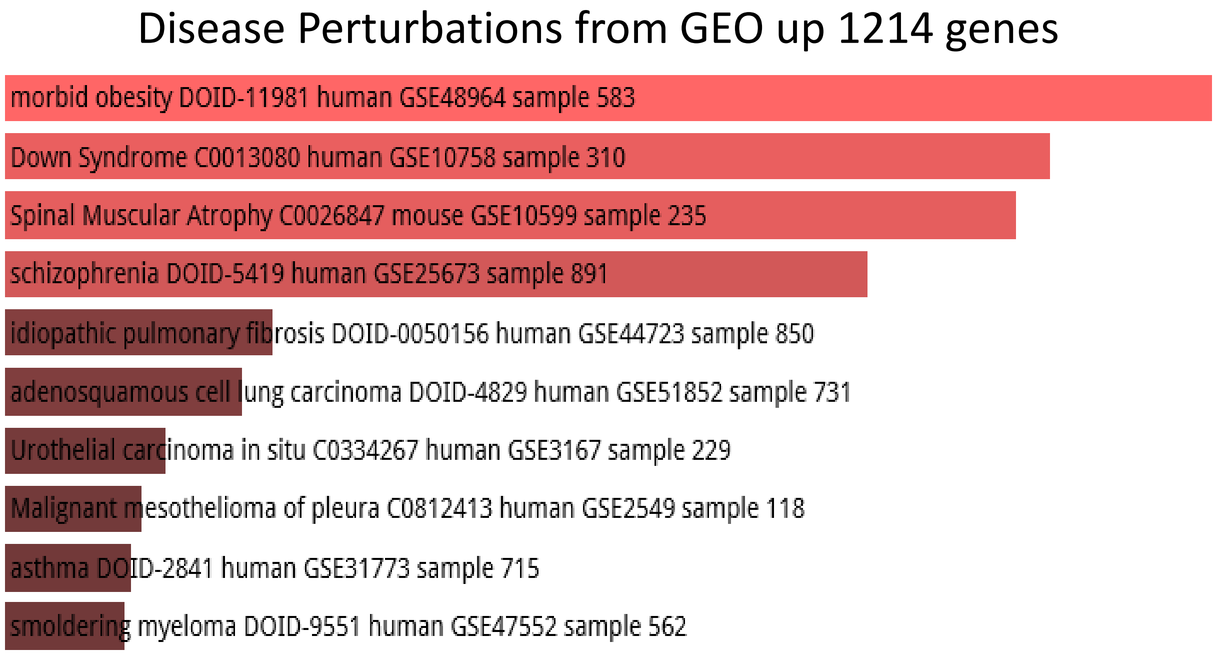


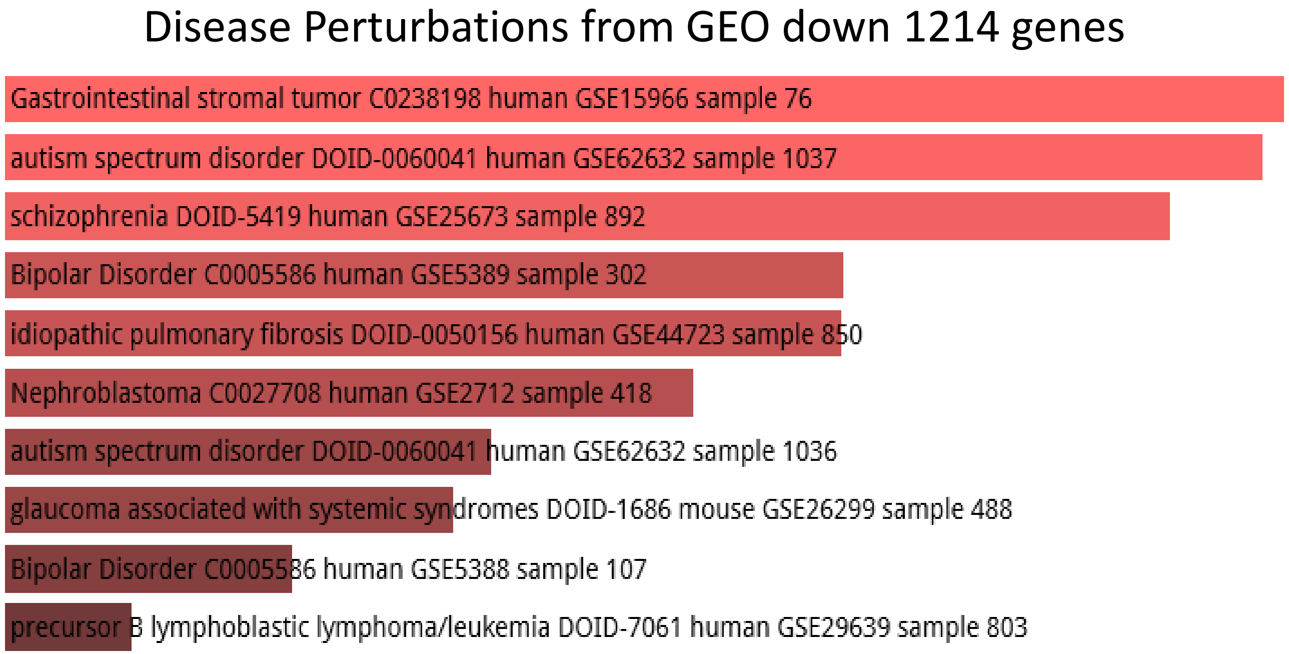

**1,619 human-specific functional hESC enhancers**

Allen Brain Atlas up 1214 genes

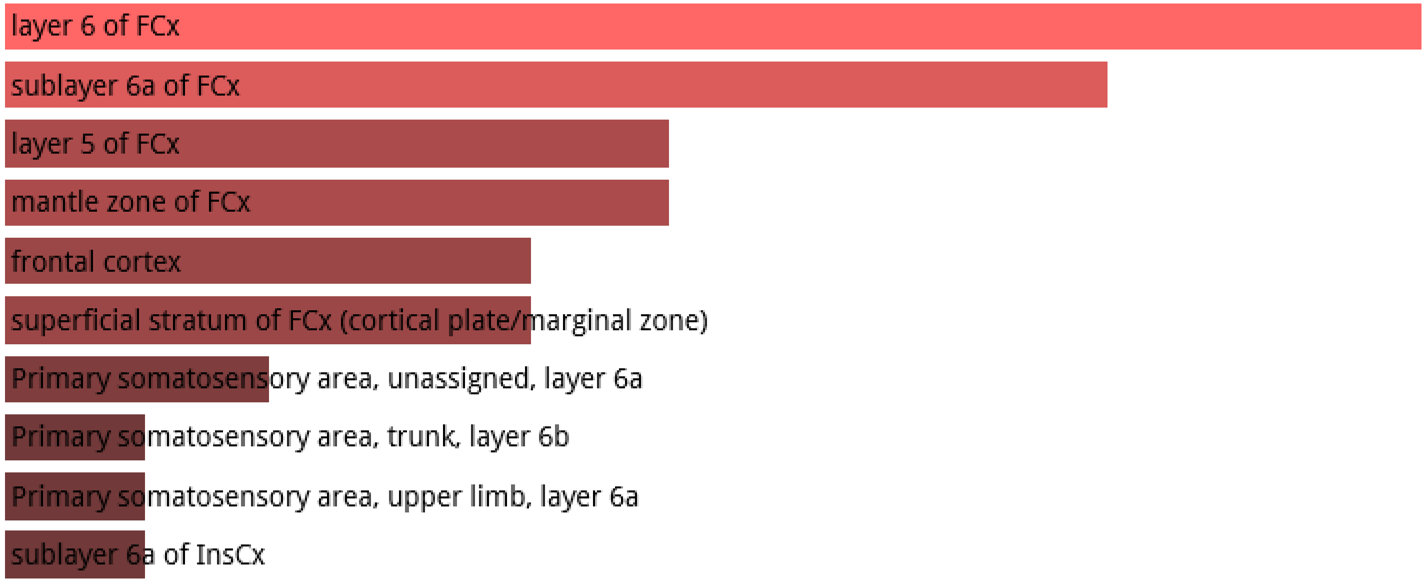


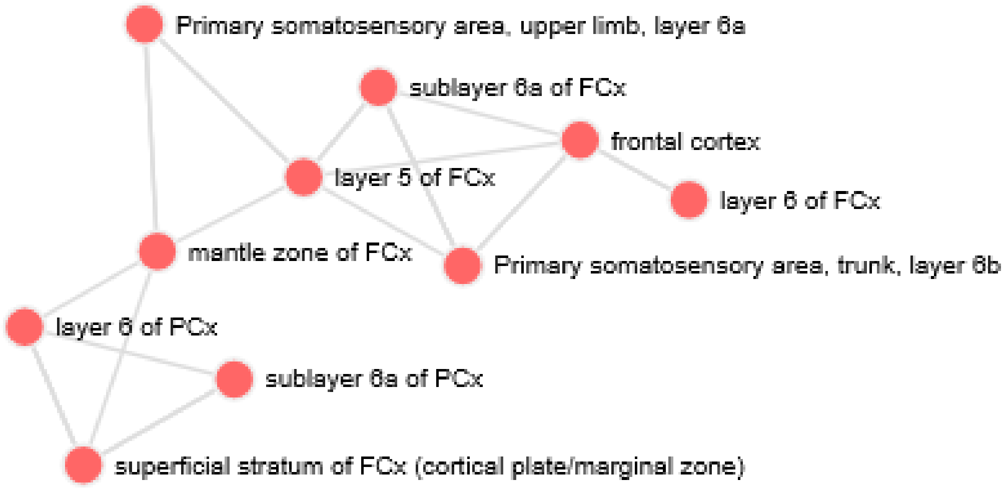

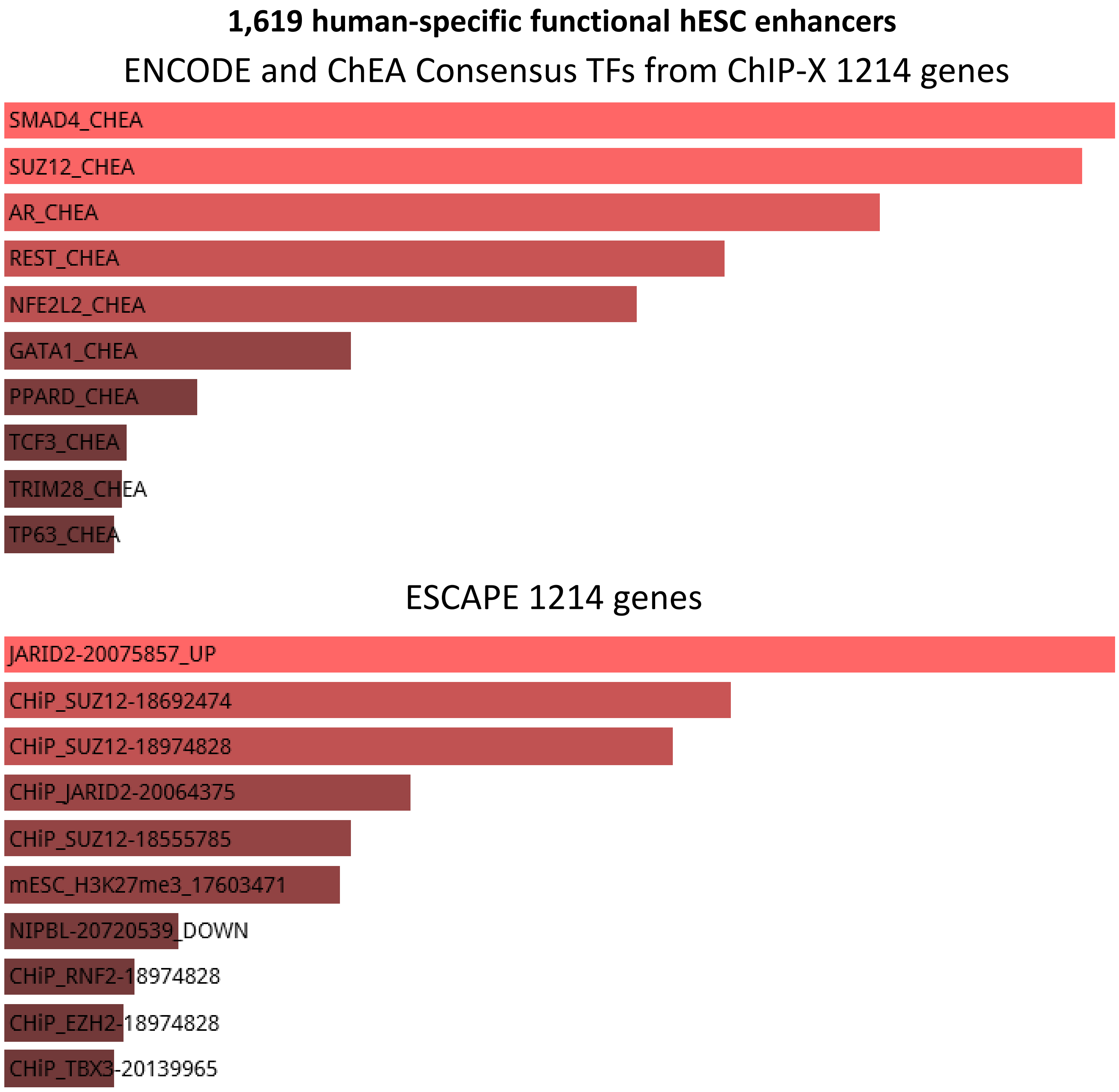
1,619 human-specific functional hESC enhancers
ENCODE and ChEA Consensus TFs from ChIP-X 1214 genes
SMAD4_CHEA
SUZ12_CHEA
AR_CHEA
REST_CHEA
NFE2L2_CHEA
GATA1_CHEA
PPARD_CHEA
TCF3_CHEA
TRIM28_CHEA
TP63_CHEA
ESCAPE 1214 genes
JARID2-20075857_UP
CHiP_SUZ12-18692474
CHiP_SUZ12-18974828
CHiP_JARID2-20064375
CHiP_SUZ12-18555785
mESC_H3K27me3_17603471
NIPBL-20720539_DOWN
CHiP_RNF2-18974828
CHiP_EZH2-18974828
CHiP_TBX3-20139965

**1,619 human-specific functional hESC enhancers**

GTEx Tissue Sample Gene Expression Profiles up 1214 genes

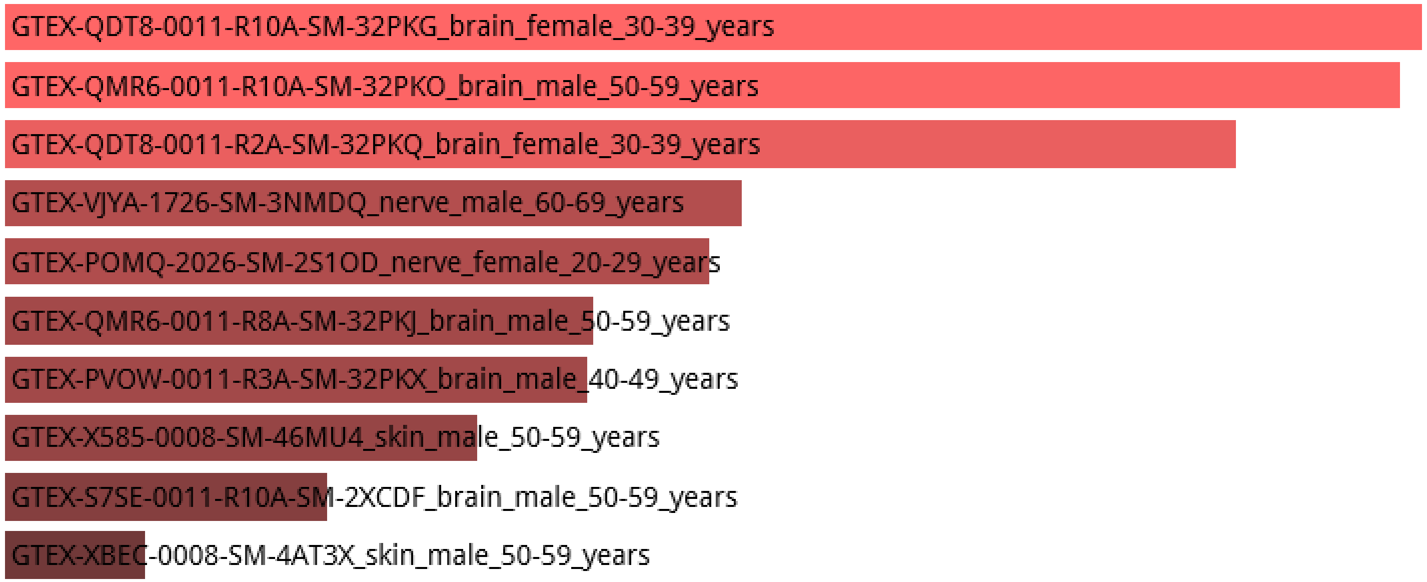


GTEx Tissue Sample Gene Expression Profiles down 1214 genes

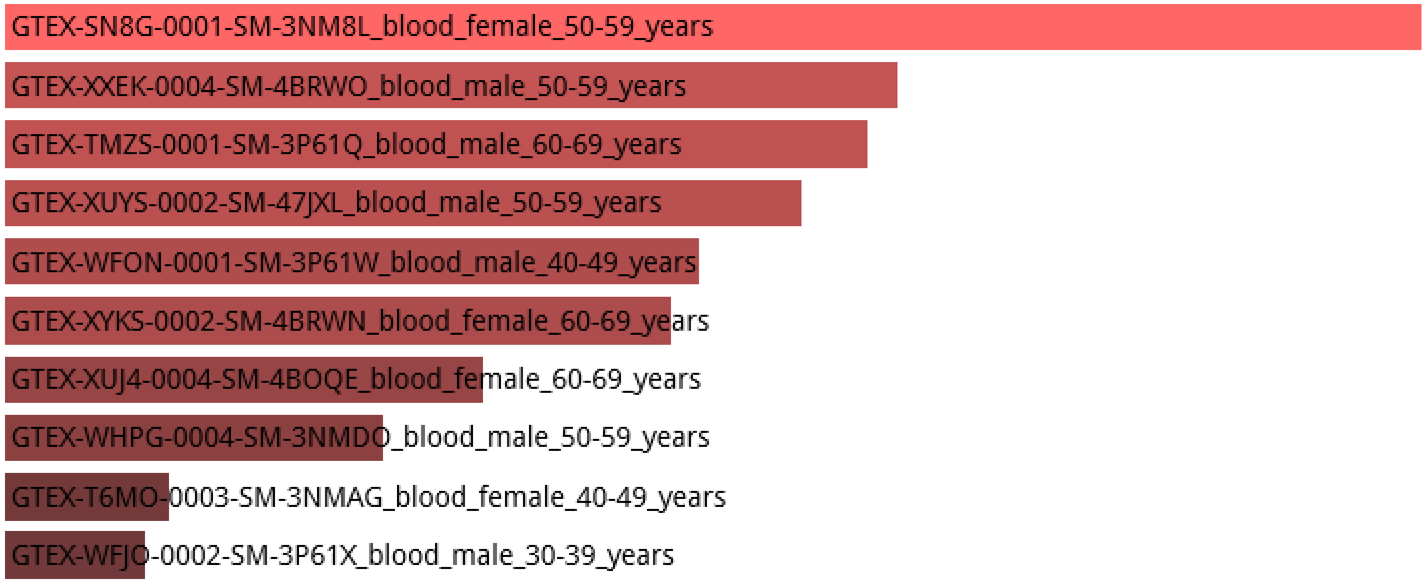

### 524 human-specific DHS

### 524 human-specific DHS

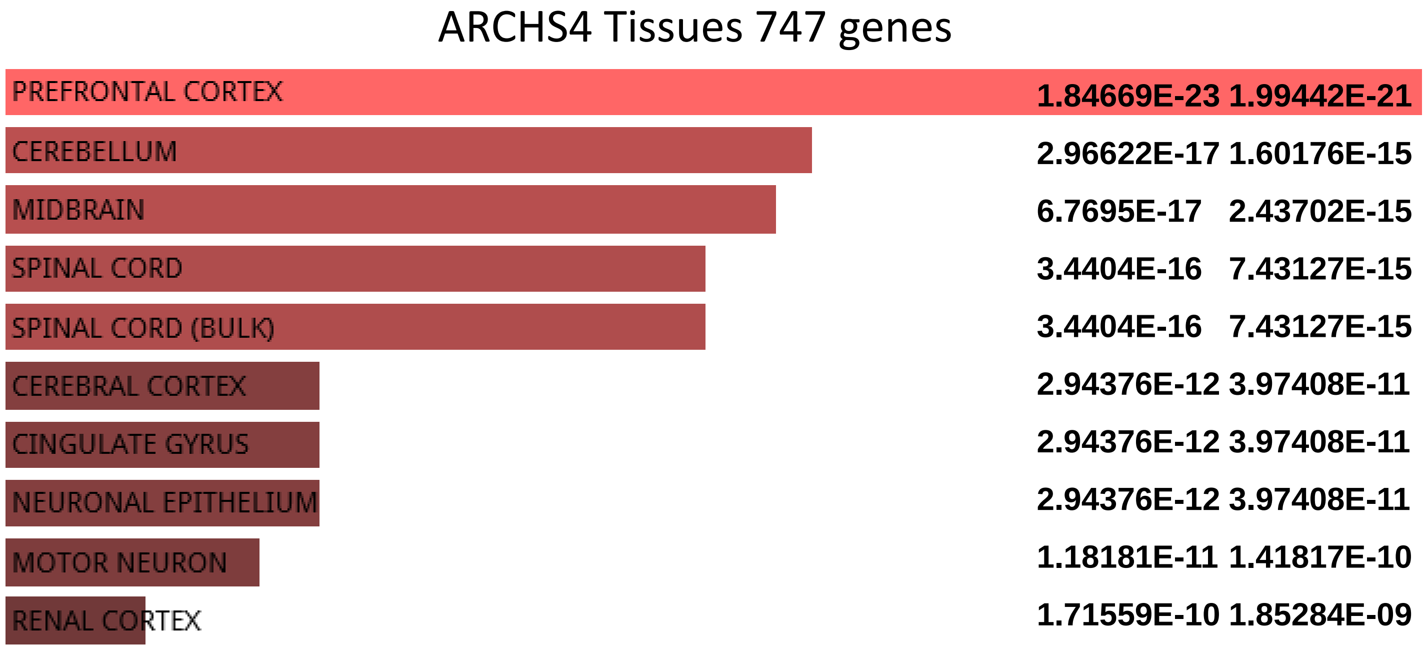


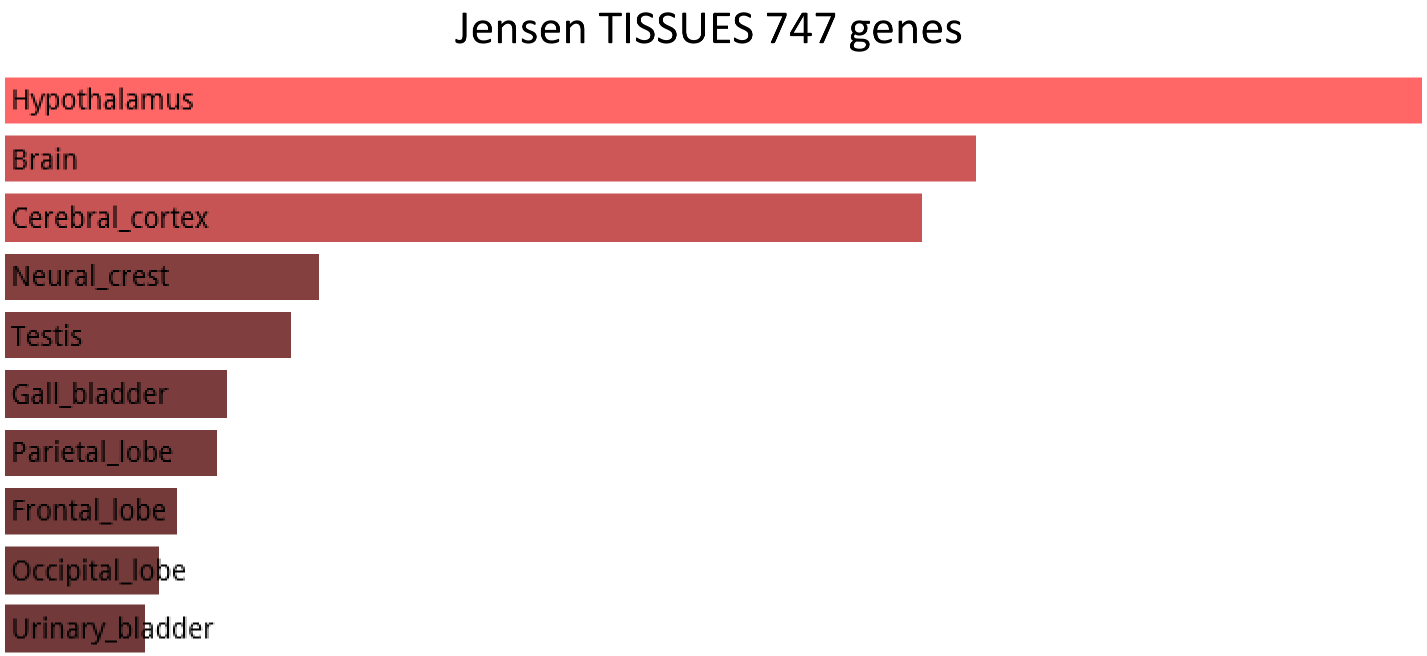

**524 human-specific DHS**

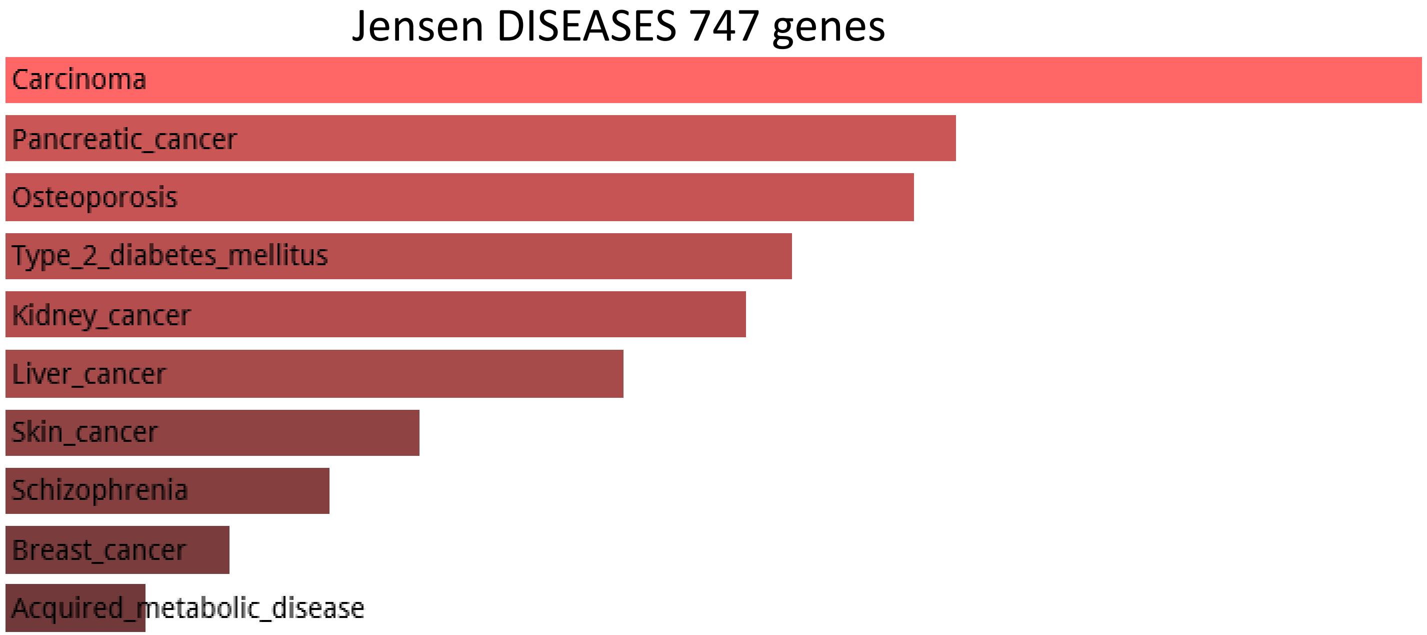


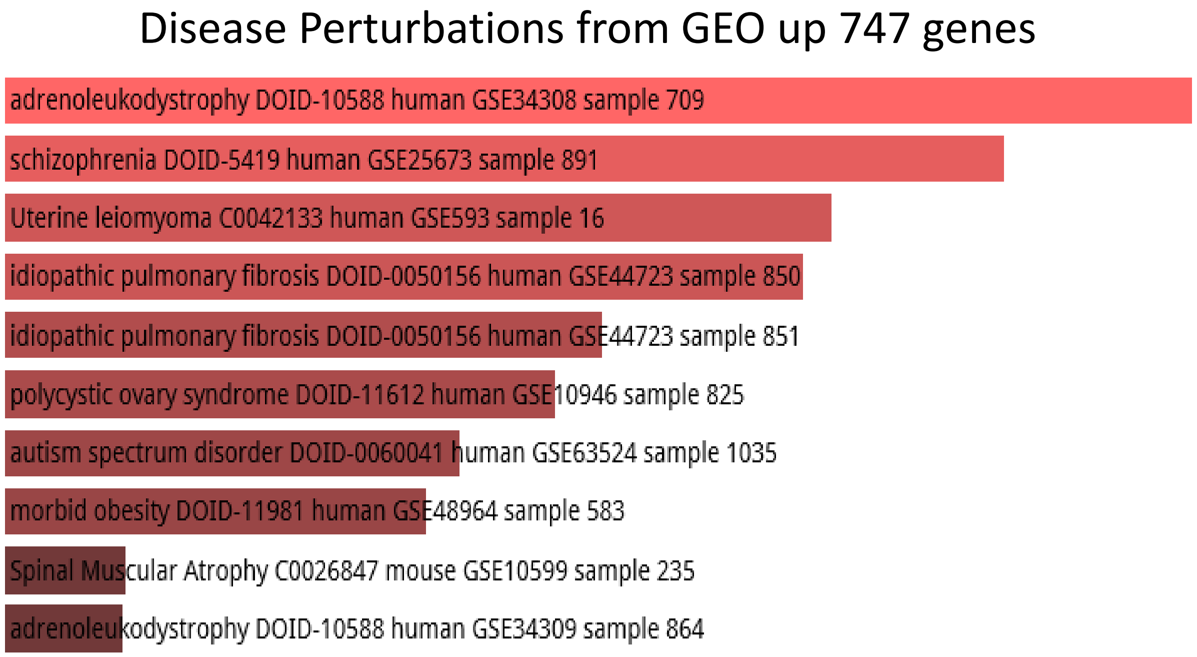


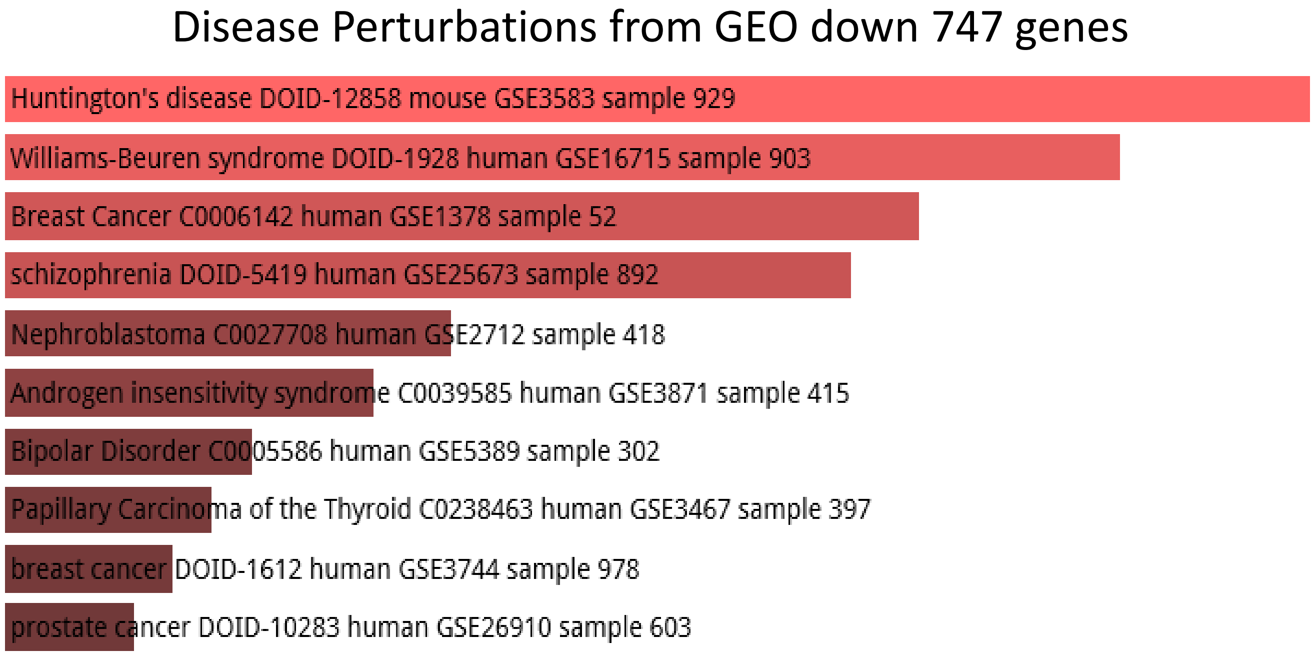

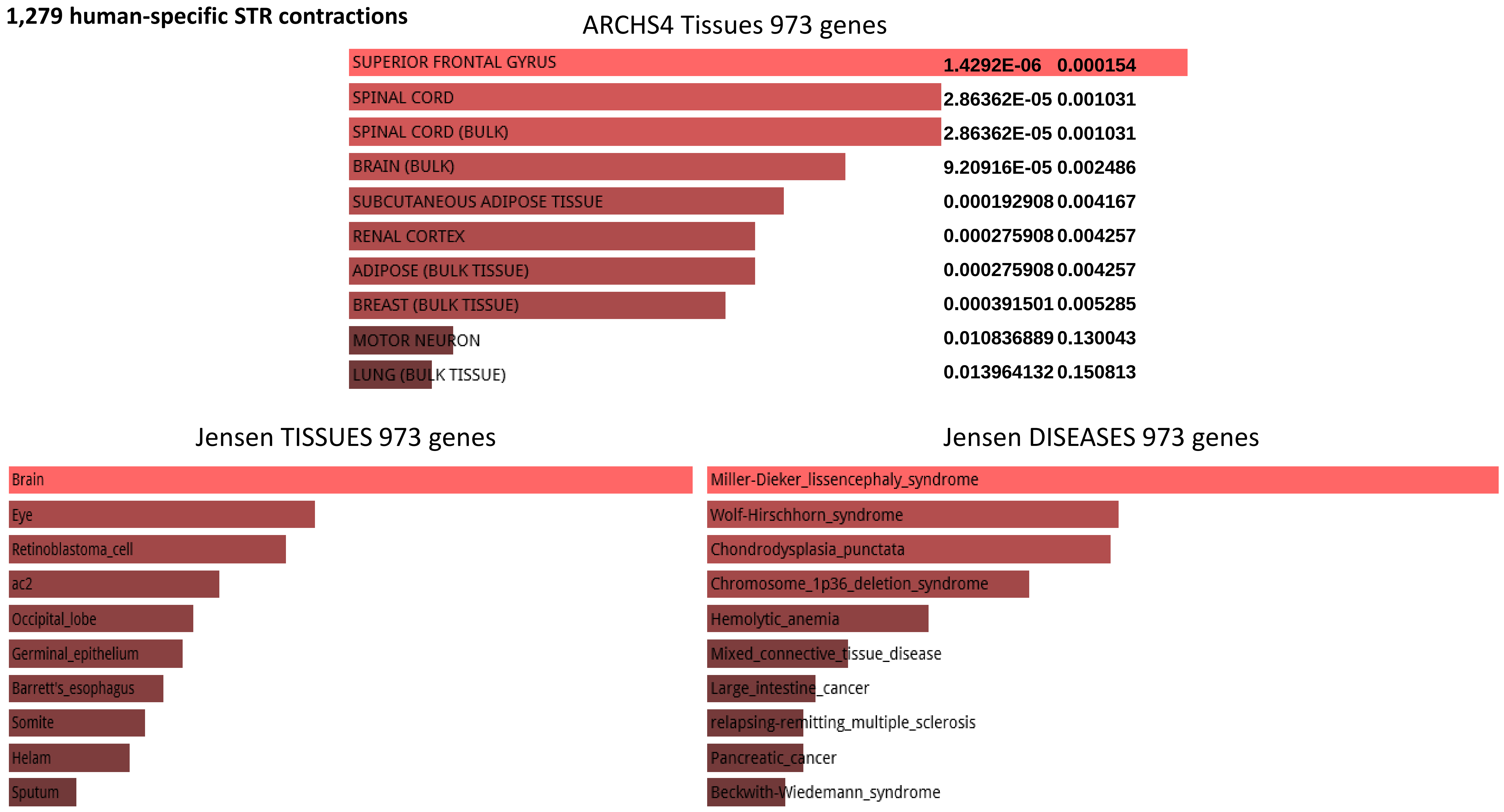
1,279 human-specific STR contractions
ARCHS4 Tissues 973 genes
SUPERIOR FRONTAL GYRUS 1.4292E-06 0.000154
SPINAL CORD 2.86362E-05 0.001031
SPINAL CORD (BULK) 2.86362E-05 0.001031
BRAIN (BULK) 9.20916E-05 0.002486
SUBCUTANEOUS ADIPOSE TISSUE 0.000192908 0.004167
RENAL CORTEX 0.000275908 0.004257
ADIPOSE (BULK TISSUE) 0.000275908 0.004257
BREAST (BULK TISSUE) 0.000391501 0.005285
MOTOR NEURON 0.010836889 0.130043
LUNG (BULK TISSUE) 0.013964132 0.150813
Jensen TISSUES 973 genes
Brain
Eye
Retinoblastoma_cell
ac2
Occipital_lobe
Germinal_epithelium
Barrett's_esophagus
Somite
Helam
Sputum
Jensen DISEASES 973 genes
Miller-Dieker_lissencephaly_syndrome
Wolf-Hirschhorn_syndrome
Chondrodysplasia_punctata
Chromosome_1p36_deletion_syndrome
Hemolytic_anemia
Mixed_connective_tissue_disease
Large_intestine_cancer
relapsing-remitting_multiple_sclerosis
Pancreatic_cancer
Beckwith-Wiedemann_syndrome

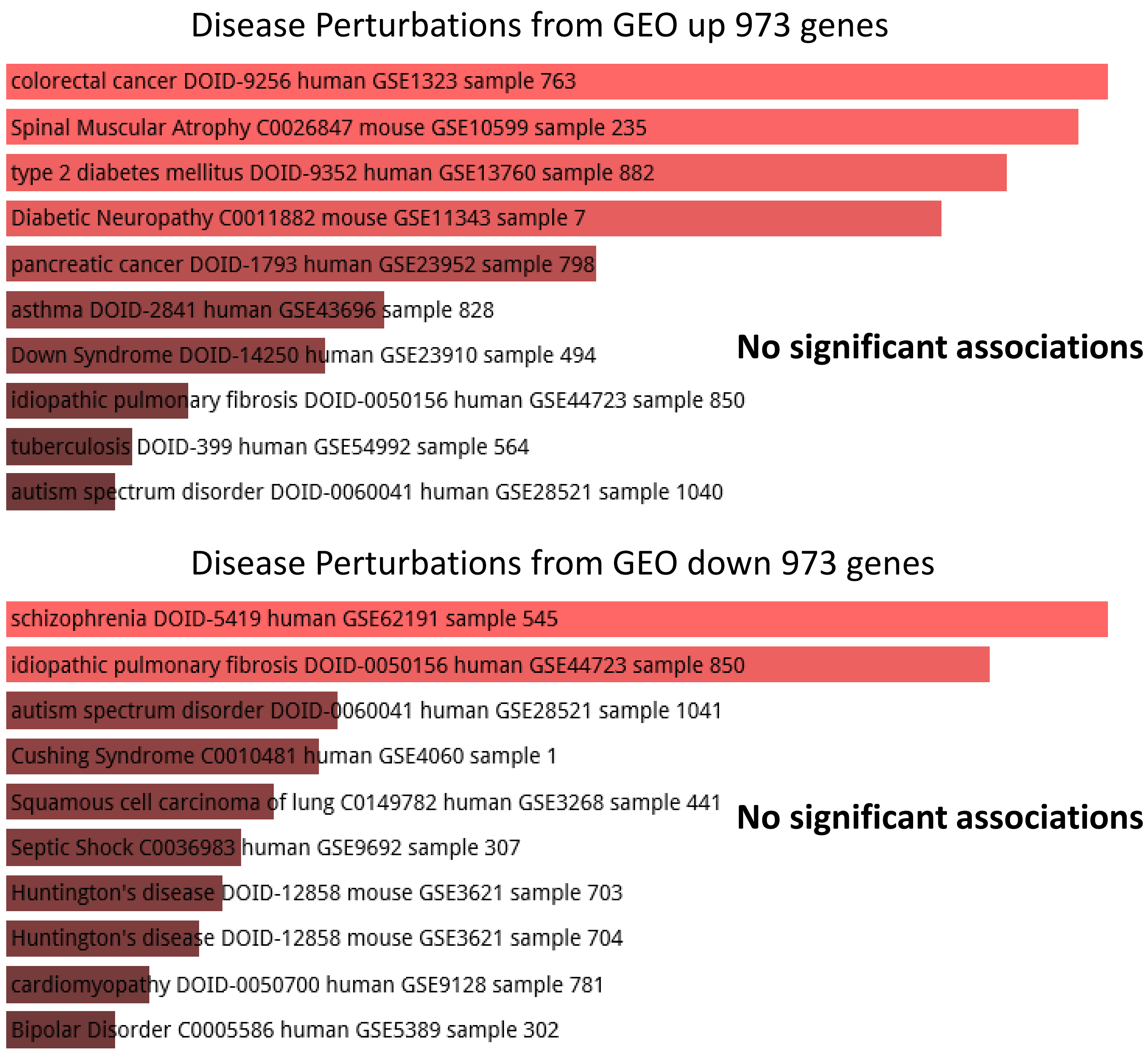
Disease Perturbations from GEO up 973 genes
colorectal cancer DOID-9256 human GSE1323 sample 763
Spinal Muscular Atrophy C0026847 mouse GSE10599 sample 235
type 2 diabetes mellitus DOID-9352 human GSE13760 sample 882
Diabetic Neuropathy C0011882 mouse GSE11343 sample 7
pancreatic cancer DOID-1793 human GSE23952 sample 798
asthma DOID-2841 human GSE43696 sample 828
Down Syndrome DOID-14250 human GSE23910 sample 494
idiopathic pulmonary fibrosis DOID-0050156 human GSE44723 sample 850
tuberculosis DOID-399 human GSE54992 sample 564
autism spectrum disorder DOID-0060041 human GSE28521 sample 1040
No significant associations
Disease Perturbations from GEO down 973 genes
schizophrenia DOID-5419 human GSE62191 sample 545
idiopathic pulmonary fibrosis DOID-0050156 human GSE44723 sample 850
autism spectrum disorder DOID-0060041 human GSE28521 sample 1041
Cushing Syndrome C0010481 human GSE4060 sample 1
Squamous cell carcinoma of lung C0149782 human GSE3268 sample 441
Septic Shock C0036983 human GSE9692 sample 307
Huntington's disease DOID-12858 mouse GSE3621 sample 703
Huntington's disease DOID-12858 mouse GSE3621 sample 704
cardiomyopathy DOID-0050700 human GSE9128 sample 781
Bipolar Disorder C0005586 human GSE5389 sample 302
No significant associations

**2,118 fixed human-specific regulatory regions (non-hESC DHS)**

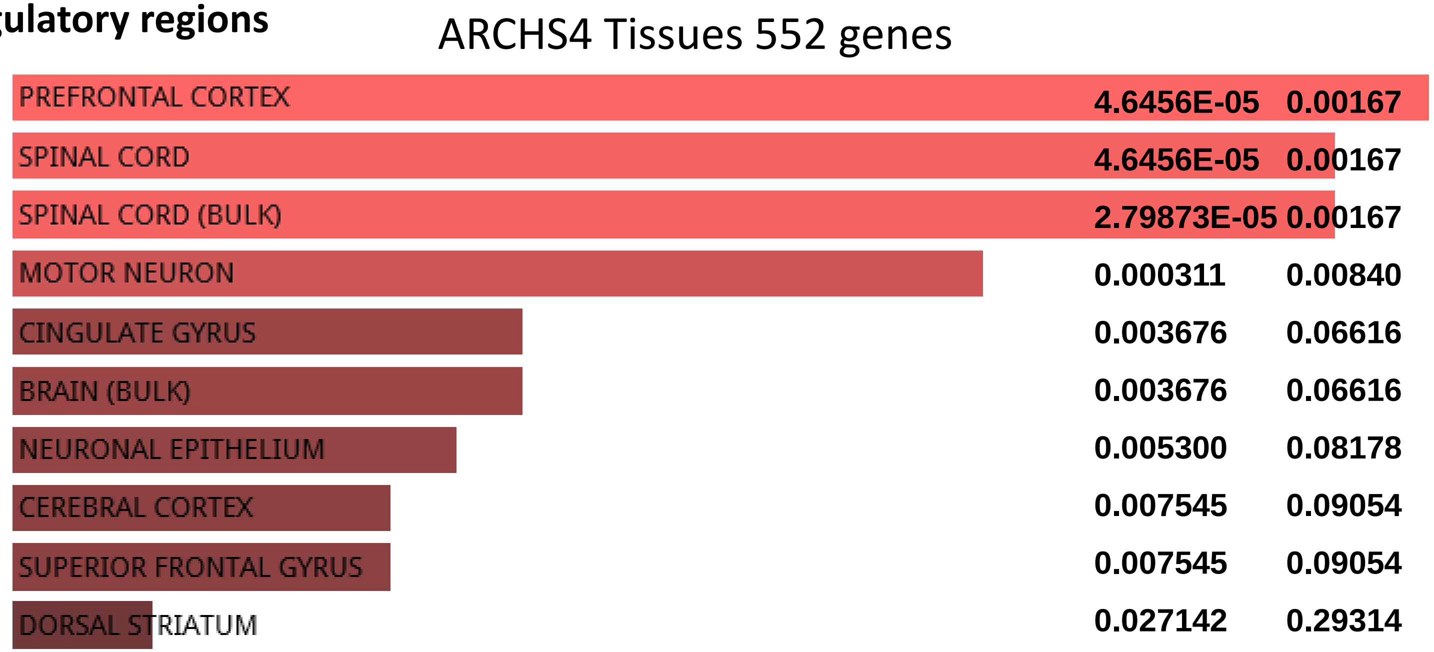


**410 H3K4me3 sites with human-specific enrichment in prefrontal cortex neurons**

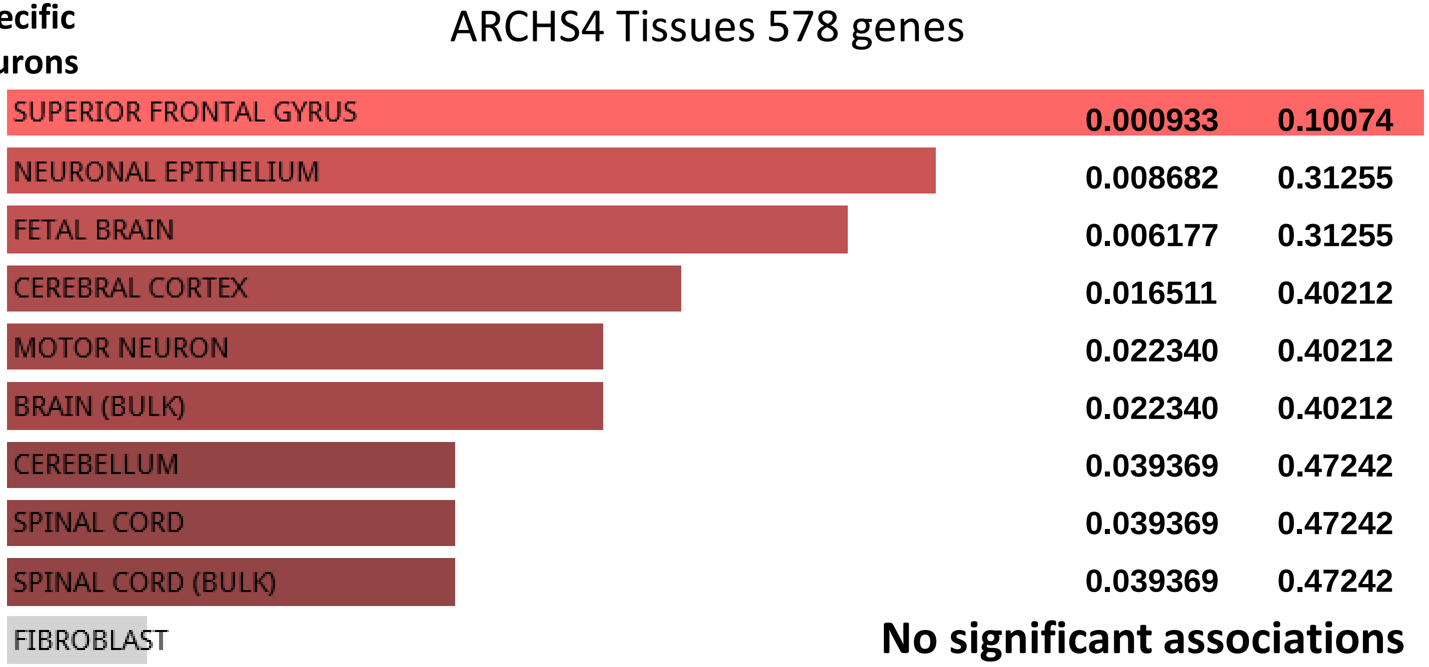

**42,847 human genes not linked by the GREAT algorithm with HSRS were randomly split into 21 control gene sets of various sizes ranging from 2,847 to 6,847 genes and subjected to the GSEA**

### GSEA OF TWENTY-ONE CONTROL GENE SETS NOT ASSOCIATED WITH HUMAN-SPECIFIC GENOMIC REGULATORY LOCI

**Random Set 1**

ARCHS4 Tissues random 6000 genes

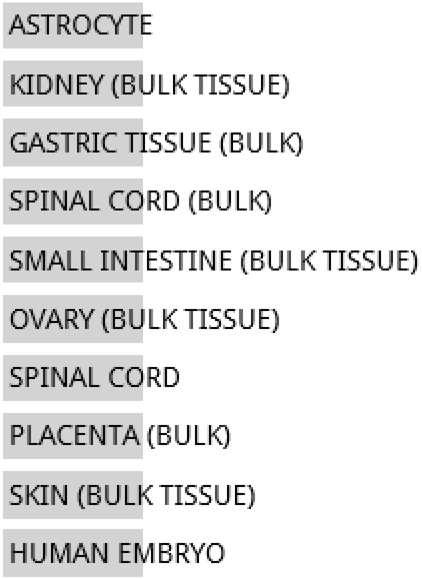


**NO SIGNIFICANT ENRICHMENT**

Jensen TISSUES random 6847 genes

**NO DATA AVAILABLE**

**Random Set 1**

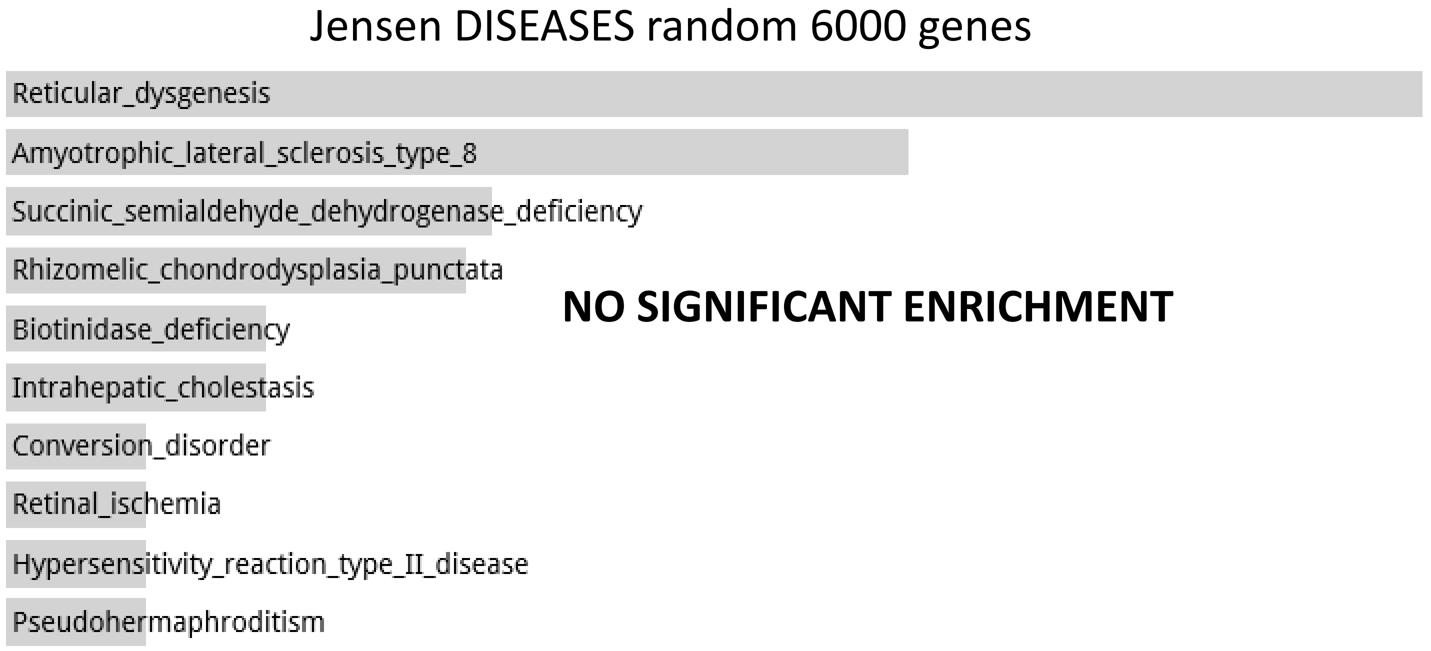
Jensen DISEASES random 6000 genes
Reticular_dysgenesis
Amyotrophic_lateral_sclerosis_type_8
Succinic_semialdehyde_dehydrogenase_deficiency
Rhizomelic_chondrodysplasia_punctata
Biotinidase_deficiency
Intrahepatic_cholestasis
Conversion_disorder
Retinal_ischemia
Hypersensitivity_reaction_type_II_disease
Pseudohermaphroditism
NO SIGNIFICANT ENRICHMENT

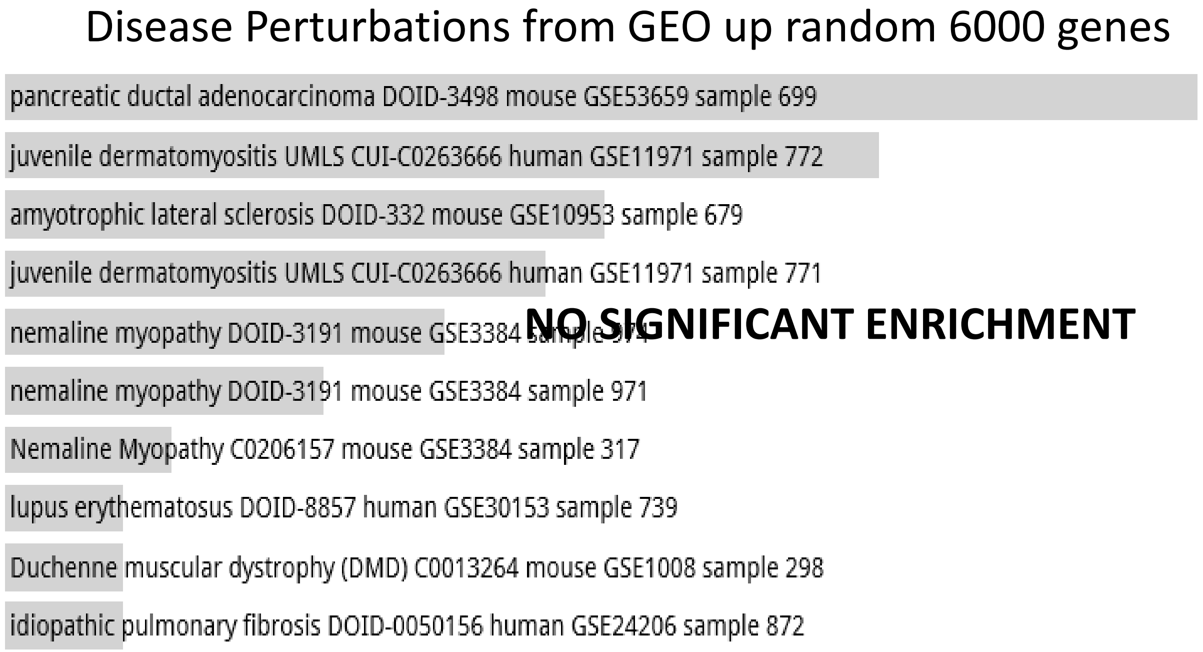
Disease Perturbations from GEO up random 6000 genes
pancreatic ductal adenocarcinoma DOID-3498 mouse GSE53659 sample 699
juvenile dermatomyositis UMLS CUI-C0263666 human GSE11971 sample 772
amyotrophic lateral sclerosis DOID-332 mouse GSE10953 sample 679
juvenile dermatomyositis UMLS CUI-C0263666 human GSE11971 sample 771
nemaline myopathy DOID-3191 mouse GSE3384
nemaline myopathy DOID-3191 mouse GSE3384 sample 971
Nemaline Myopathy C0206157 mouse GSE3384 sample 317
lupus erythematosus DOID-8857 human GSE30153 sample 739
Duchenne muscular dystrophy (DMD) C0013264 mouse GSE1008 sample 298
idiopathic pulmonary fibrosis DOID-0050156 human GSE24206 sample 872
NO SIGNIFICANT ENRICHMENT

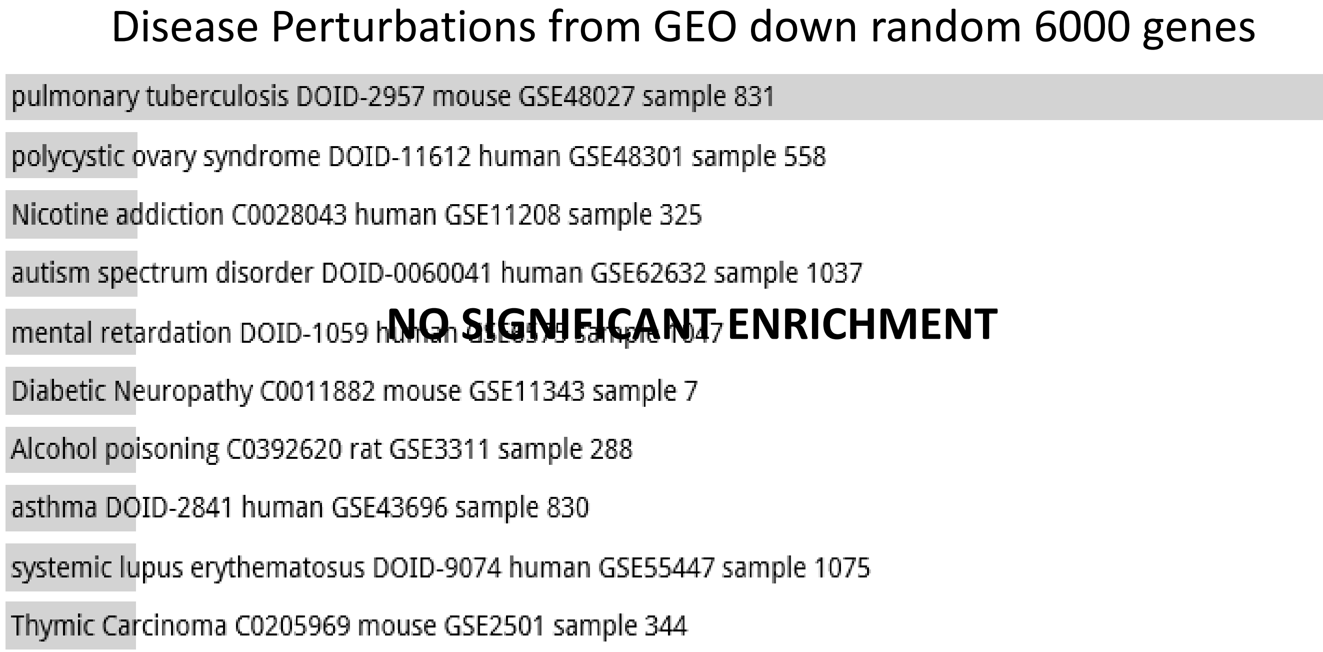
Disease Perturbations from GEO down random 6000 genes
pulmonary tuberculosis DOID-2957 mouse GSE48027 sample 831
polycystic ovary syndrome DOID-11612 human GSE48301 sample 558
Nicotine addiction C0028043 human GSE11208 sample 325
autism spectrum disorder DOID-0060041 human GSE62632 sample 1037
mental retardation DOID-1059
Diabetic Neuropathy C0011882 mouse GSE11343 sample 7
Alcohol poisoning C0392620 rat GSE3311 sample 288
asthma DOID-2841 human GSE43696 sample 830
systemic lupus erythematosus DOID-9074 human GSE55447 sample 1075
Thymic Carcinoma C0205969 mouse GSE2501 sample 344
NO SIGNIFICANT ENRICHMENT

**Random Set 1**

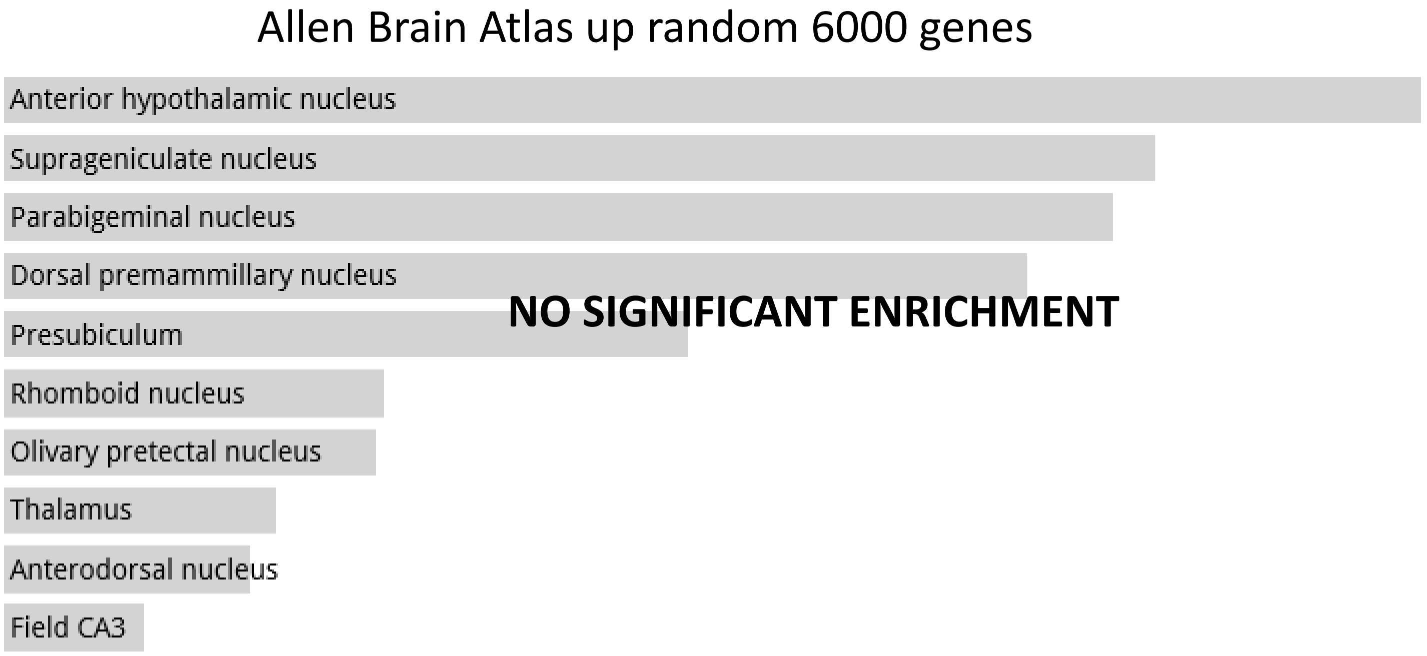

Allen Brain Atlas up random 6000 genes
Anterior hypothalamic nucleus
Suprageniculate nucleus
Parabigeminal nucleus
Dorsal premammillary nucleus
Presubiculum
Rhomboid nucleus
Olivary pretectal nucleus
Thalamus
Anterodorsal nucleus
Field CA3
NO SIGNIFICANT ENRICHMENT

**Random Set 2**

ARCHS4 Tissues random 6000 genes

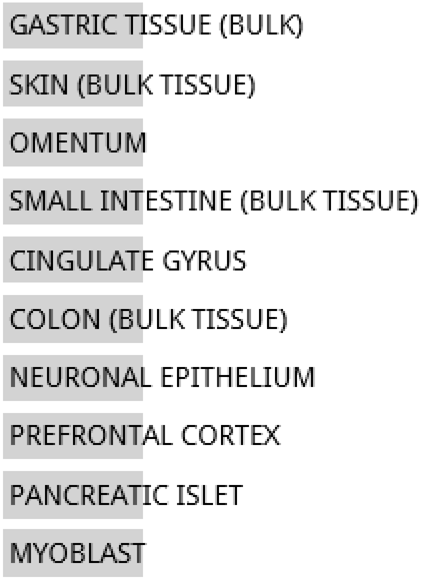


**NO SIGNIFICANT ENRICHMENT**

Jensen TISSUES random 6847 genes

**NO DATA AVAILABLE**

**Random Set 2**

#### Jensen DISEASES random 6000 genes

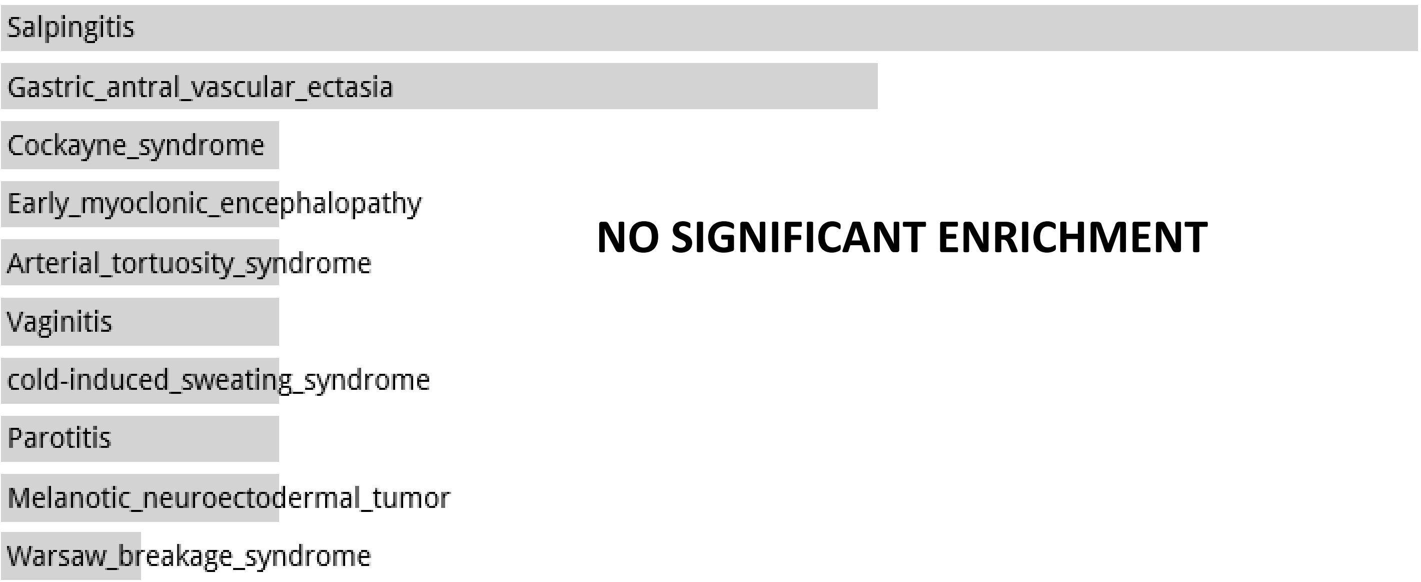


#### Disease Perturbations from GEO up random 6000 genes

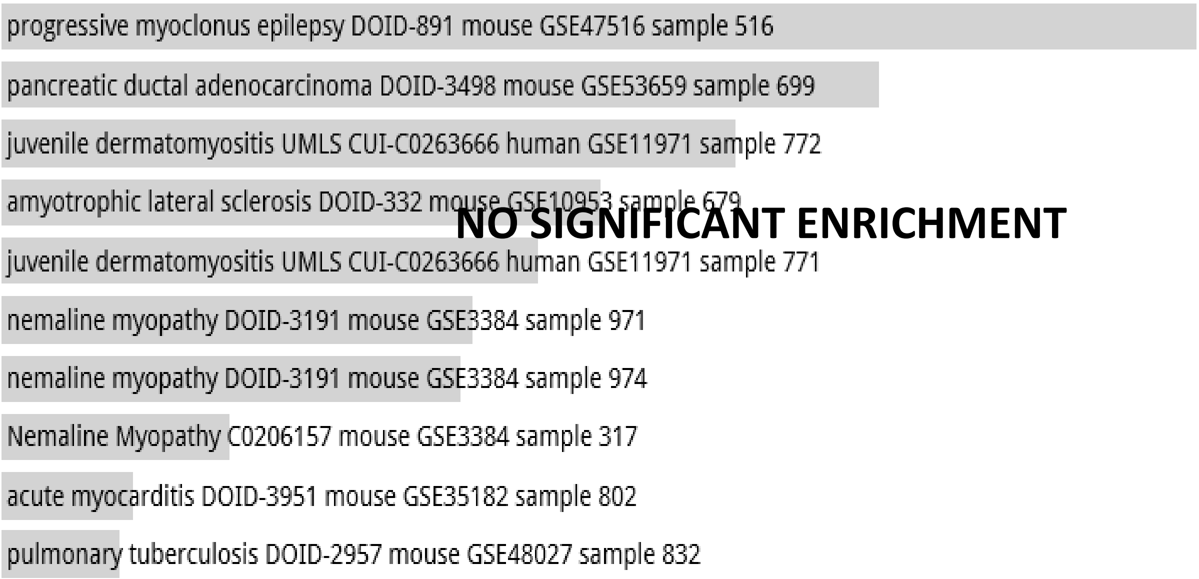


#### Disease Perturbations from GEO down random 6000 genes

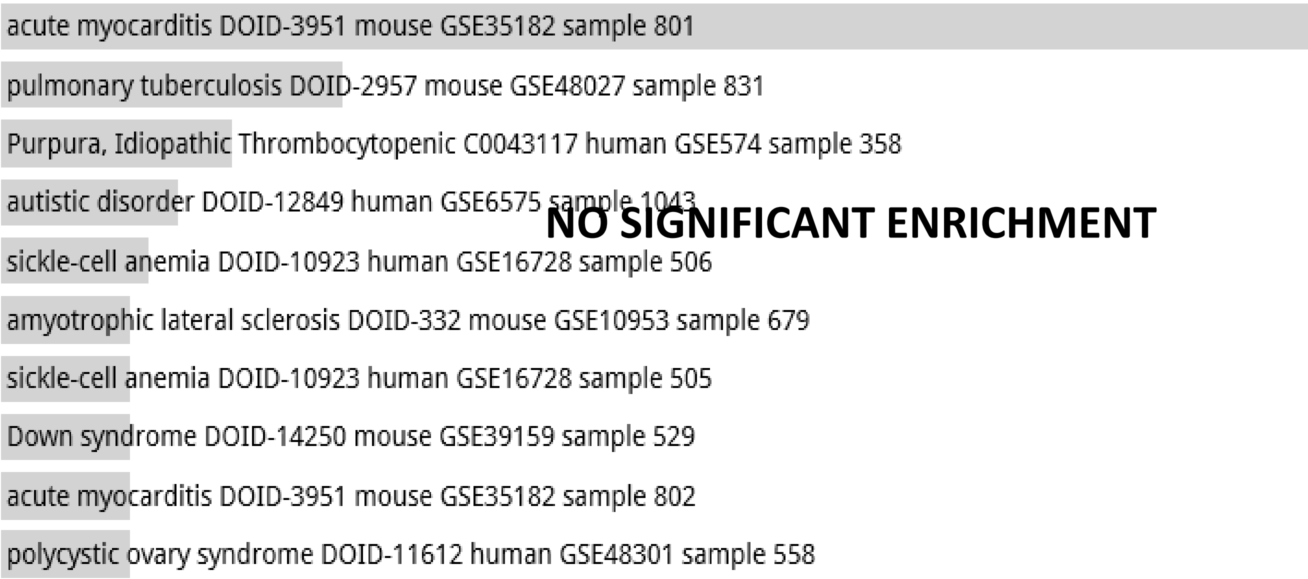

**Random Set 2**

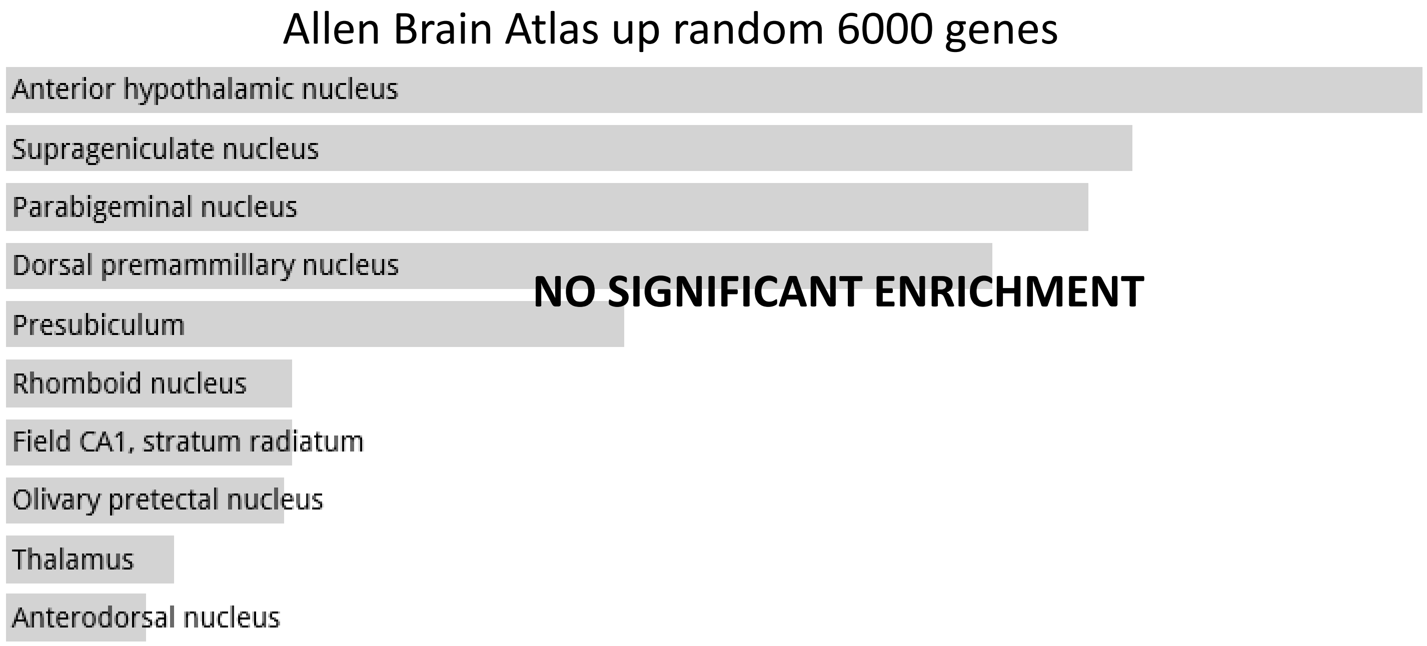
Allen Brain Atlas up random 6000 genes
Anterior hypothalamic nucleus
Suprageniculate nucleus
Parabigeminal nucleus
Dorsal premammillary nucleus
Presubiculum
Rhomboid nucleus
Field CA1, stratum radiatum
Olivary pretectal nucleus
Thalamus
Anterodorsal nucleus
NO SIGNIFICANT ENRICHMENT

**Random Set 3**

ARCHS4 Tissues random 6000 genes

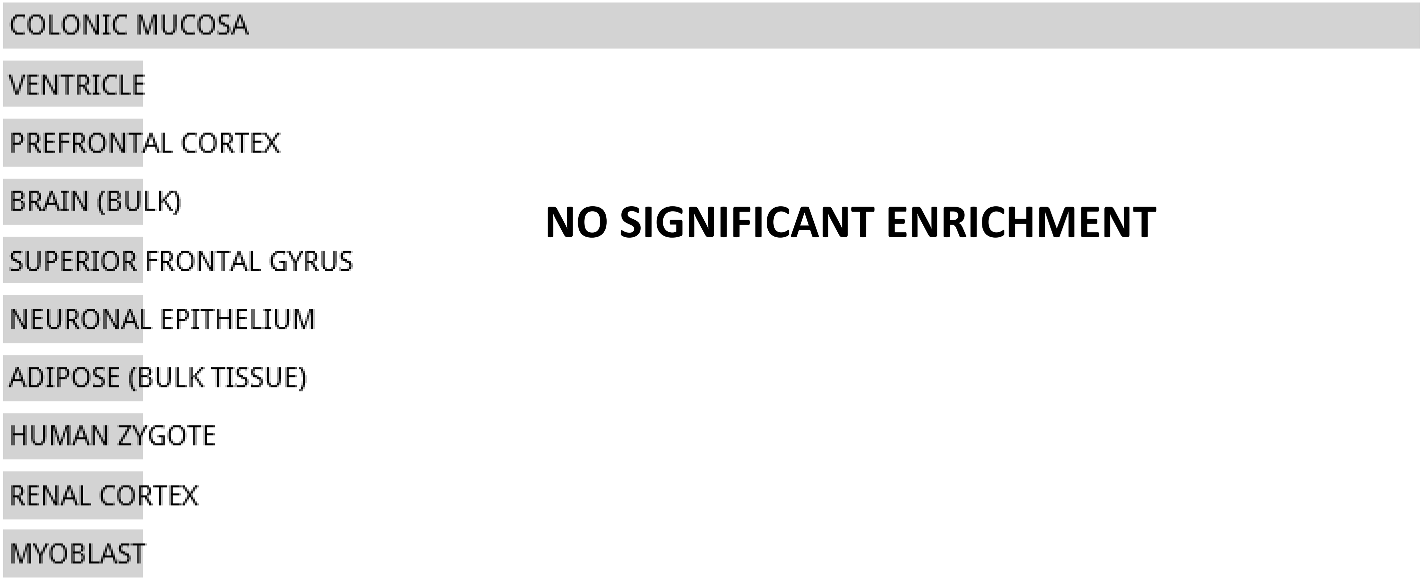


**NO SIGNIFICANT ENRICHMENT**

Jensen TISSUES random 6847 genes

**NO DATA AVAILABLE**

#### Random Set 3

Jensen DISEASES random 6000 genes

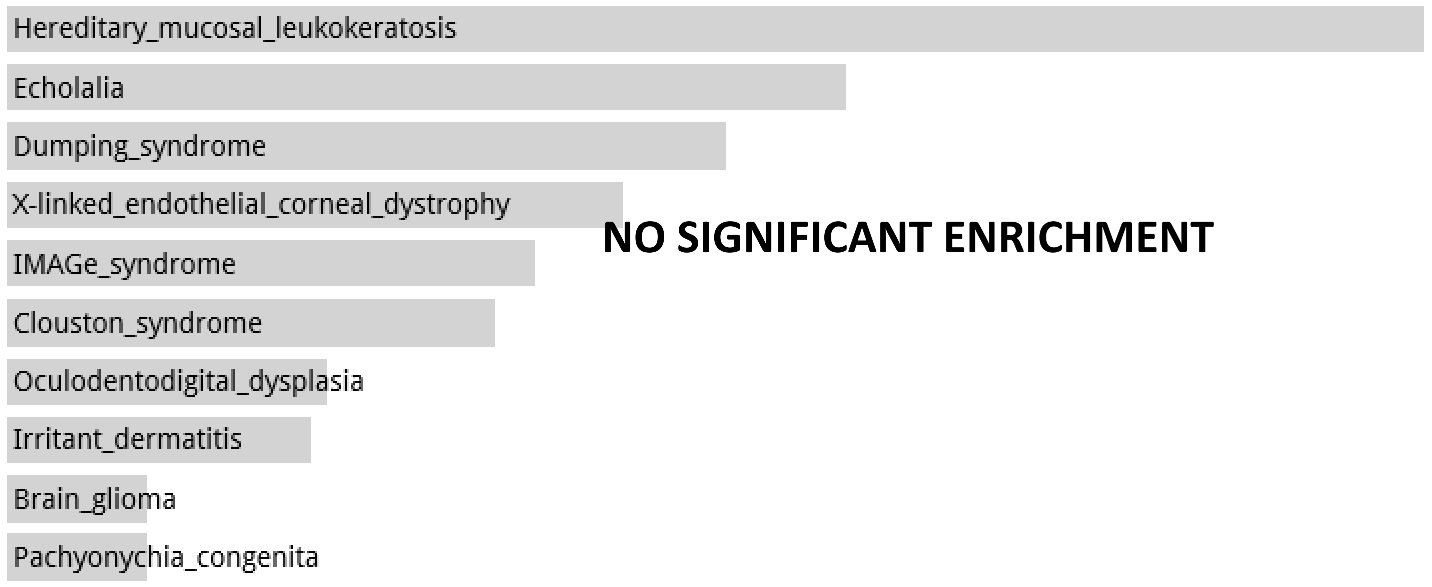


Disease Perturbations from GEO up random 6000 genes

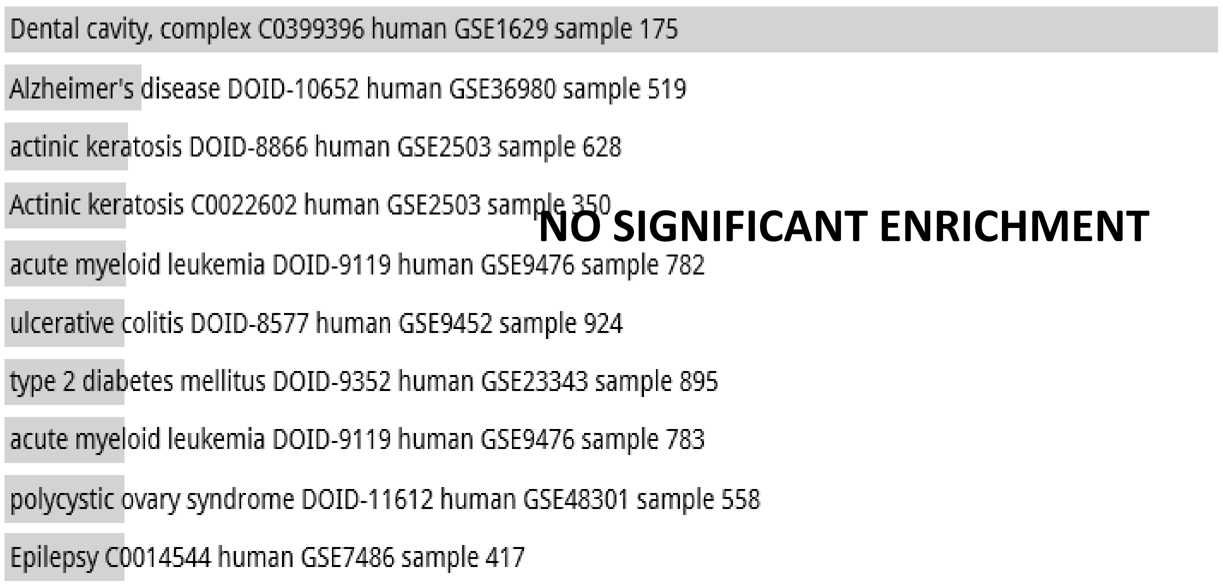


Disease Perturbations from GEO down random 6000 genes

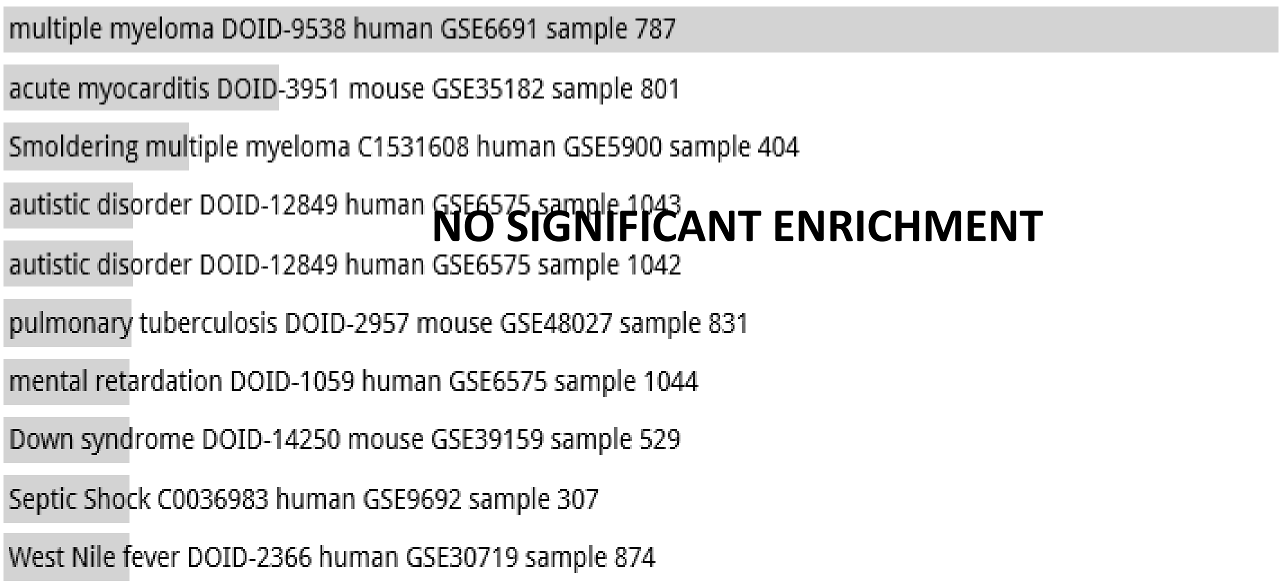

**Random Set 3**

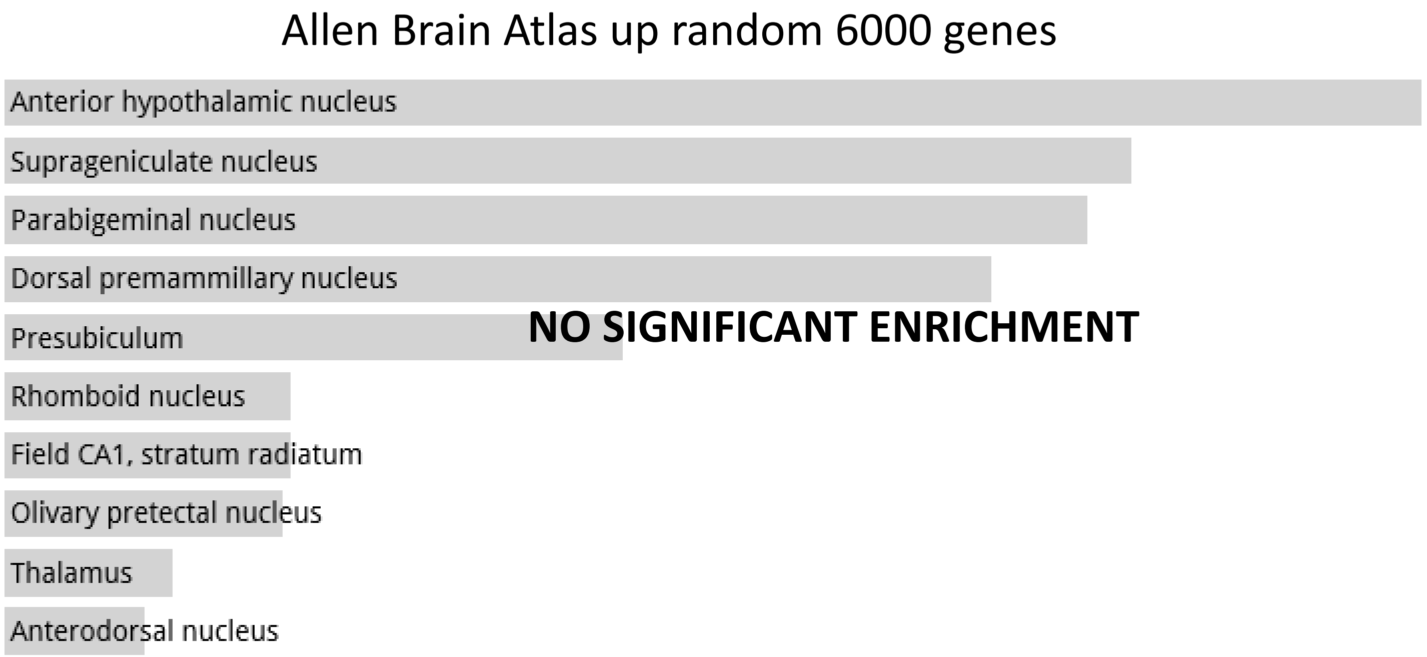
Allen Brain Atlas up random 6000 genes
Anterior hypothalamic nucleus
Suprageniculate nucleus
Parabigeminal nucleus
Dorsal premammillary nucleus
Presubiculum
Rhomboid nucleus
Field CA1, stratum radiatum
Olivary pretectal nucleus
Thalamus
Anterodorsal nucleus
NO SIGNIFICANT ENRICHMENT

**Random Set 4**

ARCHS4 Tissues random 6000 genes

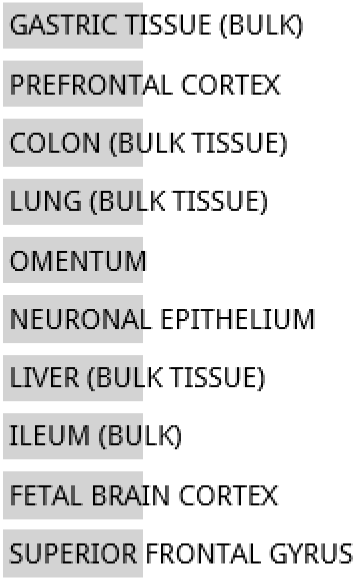


**NO SIGNIFICANT ENRICHMENT**

Jensen TISSUES random 6847 genes

**NO DATA AVAILABLE**

**Random Set 4**

Jensen DISEASES random 6000 genes

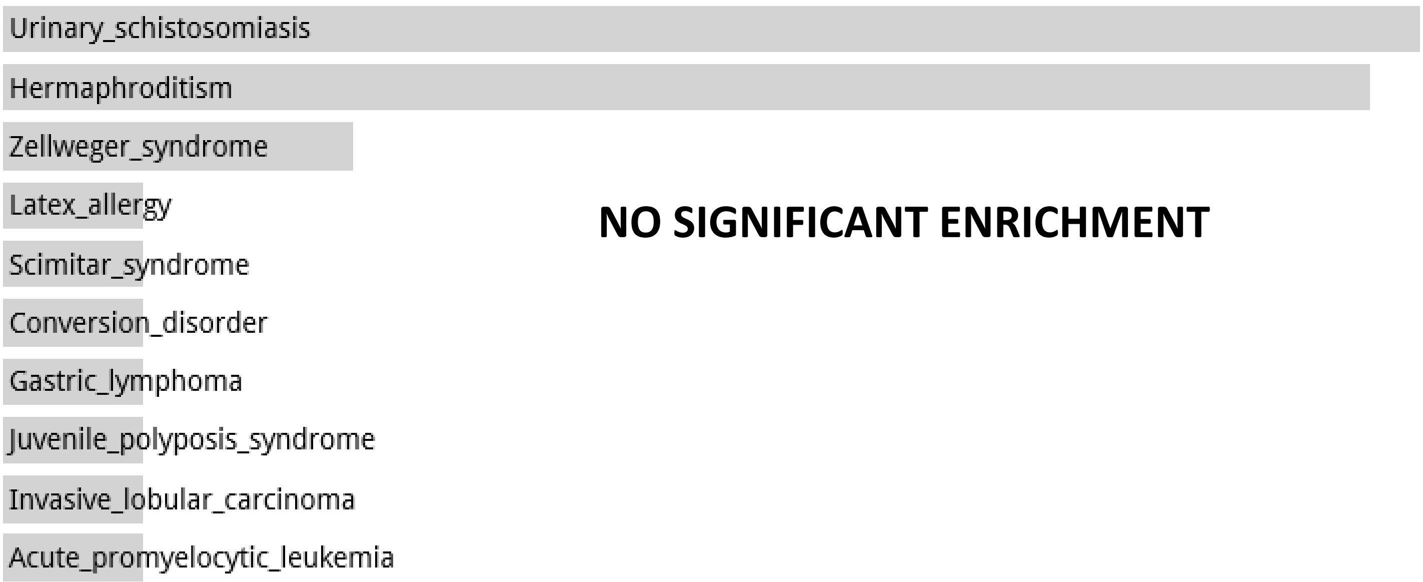


Disease Perturbations from GEO up random 6000 genes

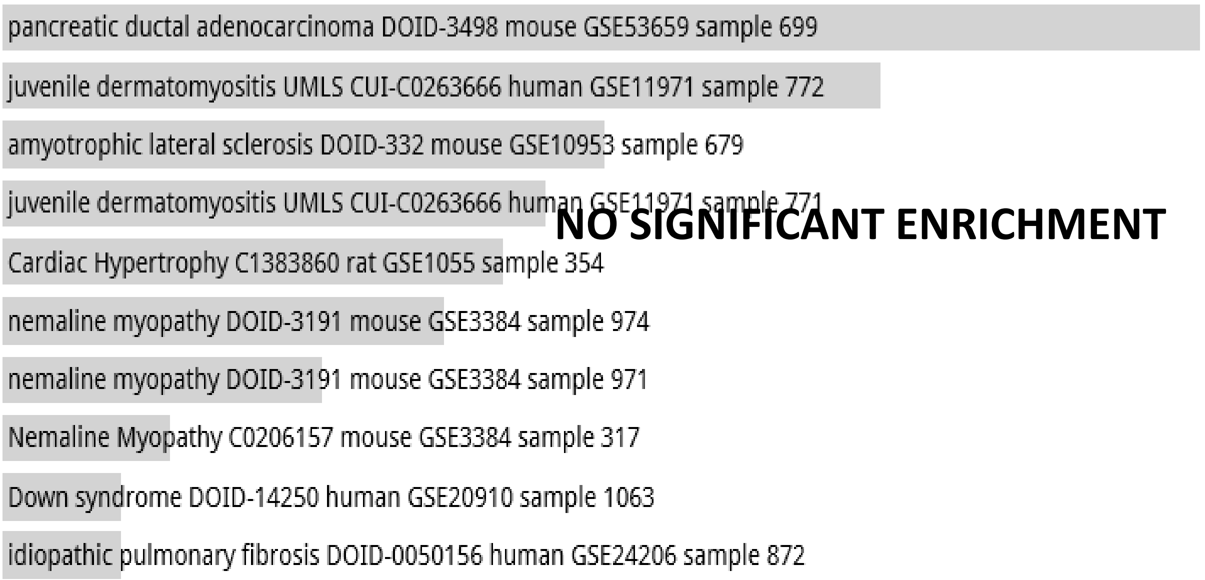


Disease Perturbations from GEO down random 6000 genes

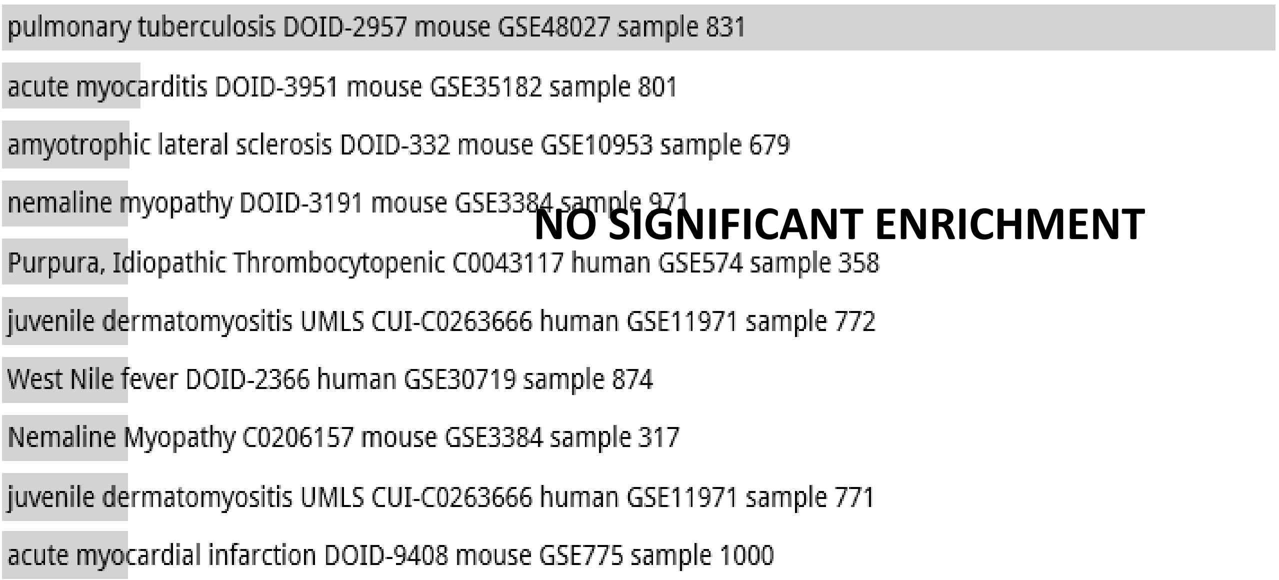

**Random Set 4**

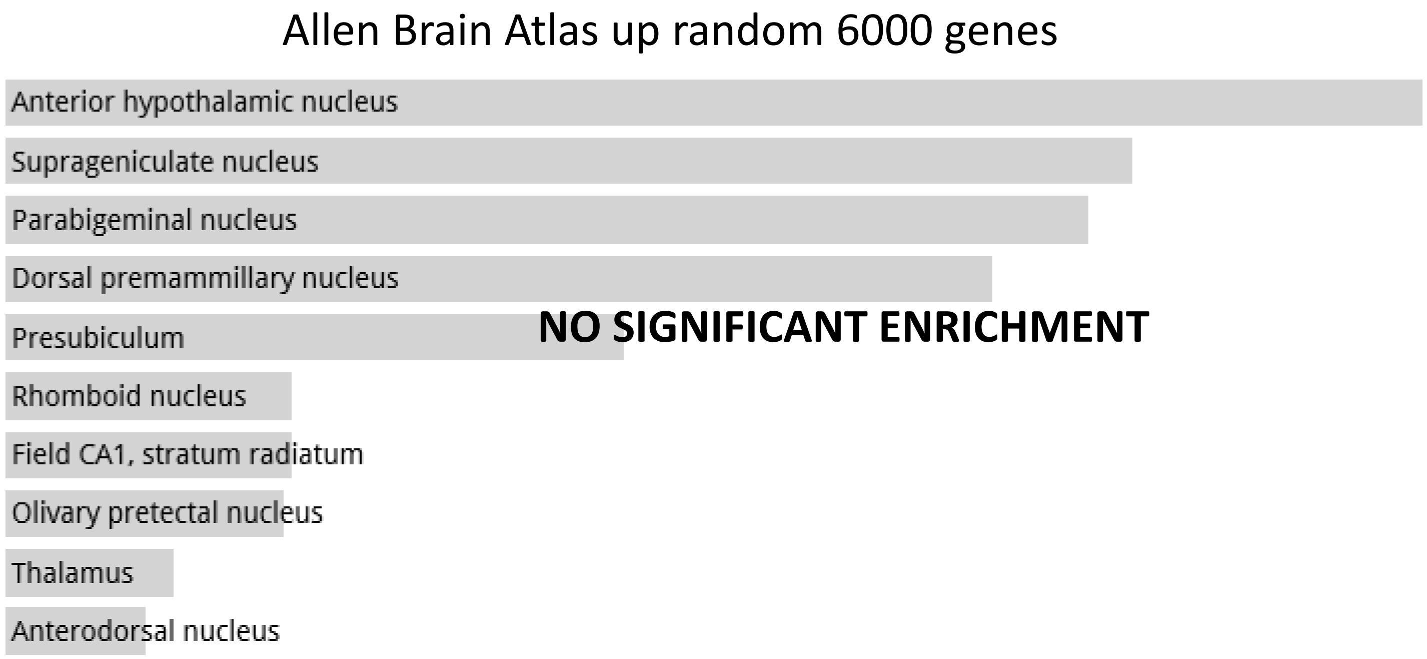
Allen Brain Atlas up random 6000 genes
Anterior hypothalamic nucleus
Suprageniculate nucleus
Parabigeminal nucleus
Dorsal premammillary nucleus
Presubiculum
Rhomboid nucleus
Field CA1, stratum radiatum
Olivary pretectal nucleus
Thalamus
Anterodorsal nucleus
NO SIGNIFICANT ENRICHMENT

**Random Set 5**

#### ARCHS4 Tissues random 6000 genes

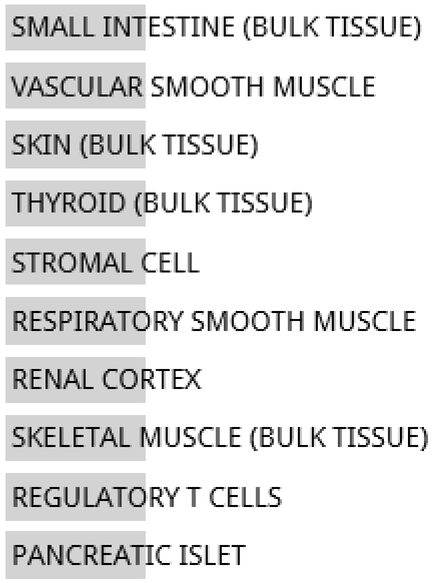


**NO SIGNIFICANT ENRICHMENT**

#### Jensen TISSUES random 6847 genes

**NO DATA AVAILABLE**

#### Random Set 5

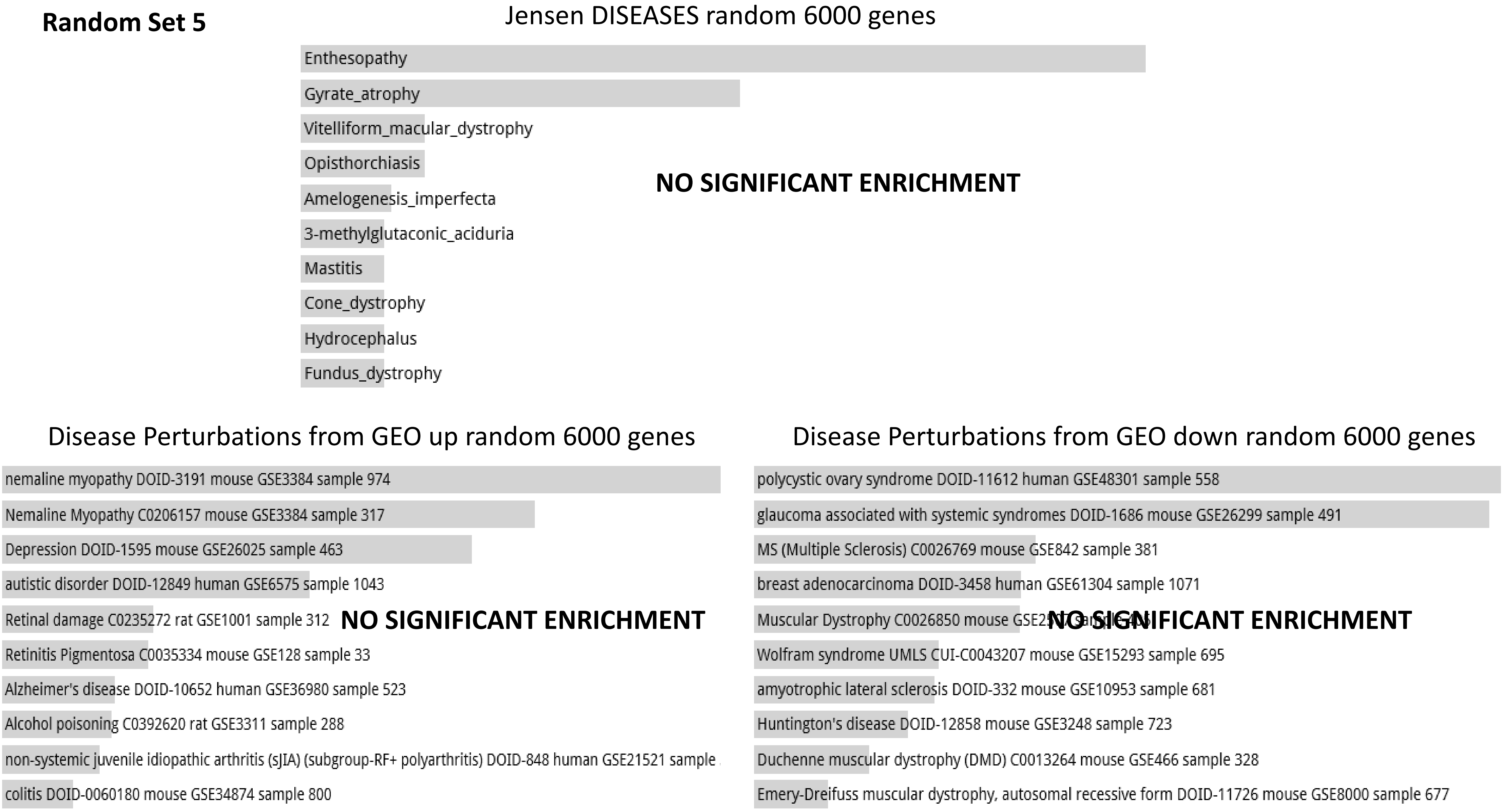

Jensen DISEASES random 6000 genes
Enthesopathy
Gyrate_atrophy
Vitelliform_macular_dystrophy
Opisthorchiasis
Amelogenesis_imperfecta
3-methylglutaconic_aciduria
Mastitis
Cone_dystrophy
Hydrocephalus
Fundus_dystrophy
NO SIGNIFICANT ENRICHMENT
Disease Perturbations from GEO up random 6000 genes
nemaline myopathy DOID-3191 mouse GSE3384 sample 974
Nemaline Myopathy C0206157 mouse GSE3384 sample 317
Depression DOID-1595 mouse GSE26025 sample 463
autistic disorder DOID-12849 human GSE6575 sample 1043
Retinal damage C0235272 rat GSE1001 sample 312
Retinitis Pigmentosa C0035334 mouse GSE128 sample 33
Alzheimer's disease DOID-10652 human GSE36980 sample 523
Alcohol poisoning C0392620 rat GSE3311 sample 288
non-systemic juvenile idiopathic arthritis (sJIA) (subgroup-RF+ polyarthritis) DOID-848 human GSE21521 sample
colitis DOID-0060180 mouse GSE34874 sample 800
NO SIGNIFICANT ENRICHMENT
Disease Perturbations from GEO down random 6000 genes
polycystic ovary syndrome DOID-11612 human GSE48301 sample 558
glaucoma associated with systemic syndromes DOID-1686 mouse GSE26299 sample 491
MS (Multiple Sclerosis) C0026769 mouse GSE842 sample 381
breast adenocarcinoma DOID-3458 human GSE61304 sample 1071
Muscular Dystrophy C0026850 mouse
Wolfram syndrome UMLS CUI-C0043207 mouse GSE15293 sample 695
amyotrophic lateral sclerosis DOID-332 mouse GSE10953 sample 681
Huntington's disease DOID-12858 mouse GSE3248 sample 723
Duchenne muscular dystrophy (DMD) C0013264 mouse GSE466 sample 328
Emery-Dreifuss muscular dystrophy, autosomal recessive form DOID-11726 mouse GSE8000 sample 677
NO SIGNIFICANT ENRICHMENT

**Random Set 5**

#### Allen Brain Atlas up random 6000 genes

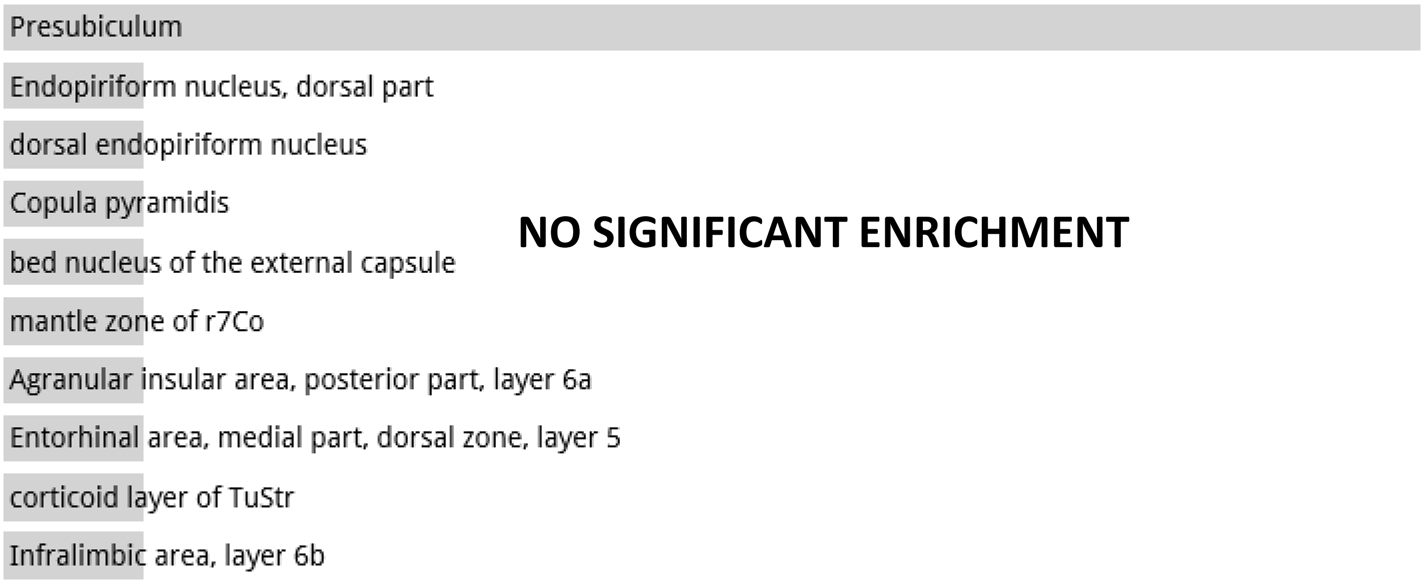

**Random Set 6**

ARCHS4 Tissues random 6000 genes

**NO DATA AVAILABLE**

Jensen TISSUES random 6000 genes

**NO DATA AVAILABLE**

**Random Set 6**

Jensen DISEASES random 6000 genes

**NO DATA AVAILABLE**

Disease Perturbations from GEO up random 6000 genes

**NO DATA AVAILABLE**

Disease Perturbations from GEO down random 6000 genes

Systemic lupus erythematosus (SLE) C0024141 human GSE12374 sample 123

**NO SIGNIFICANT ENRICHMENT**

**Random Set 6**

Allen Brain Atlas up random 6000 genes

**NO DATA AVAILABLE**

**Random Set 7**

ARCHS4 Tissues random 6847 genes

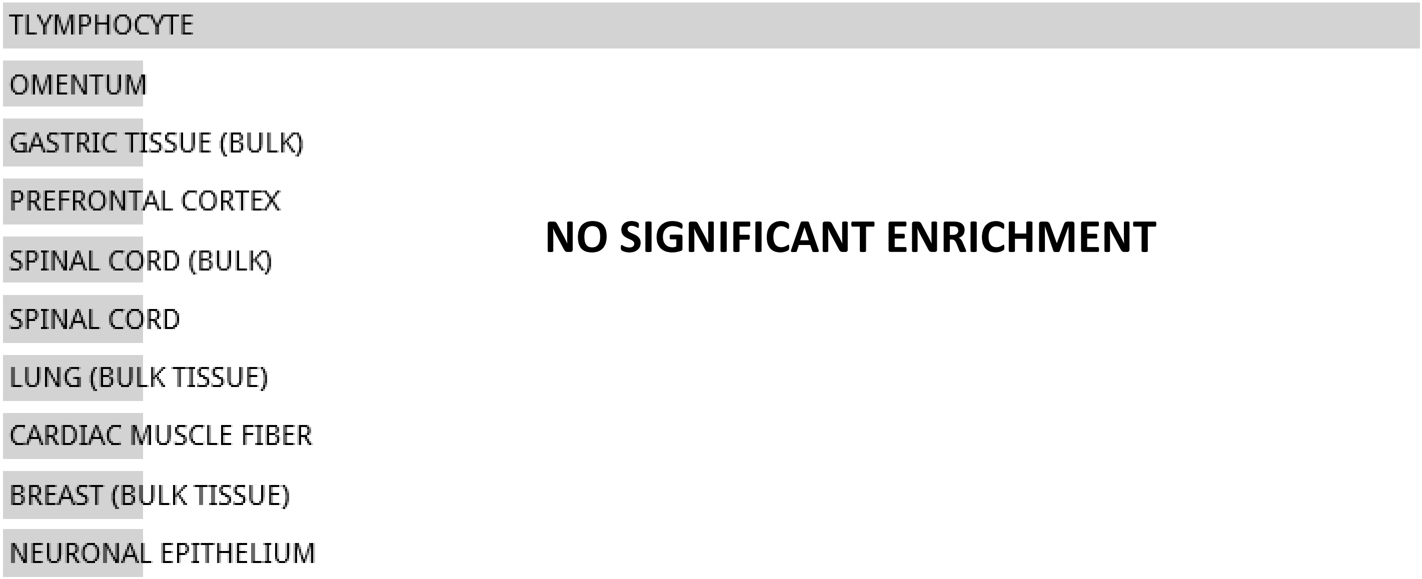


**NO SIGNIFICANT ENRICHMENT**

Jensen TISSUES random 6847 genes

**NO DATA AVAILABLE**

**Random Set 7**

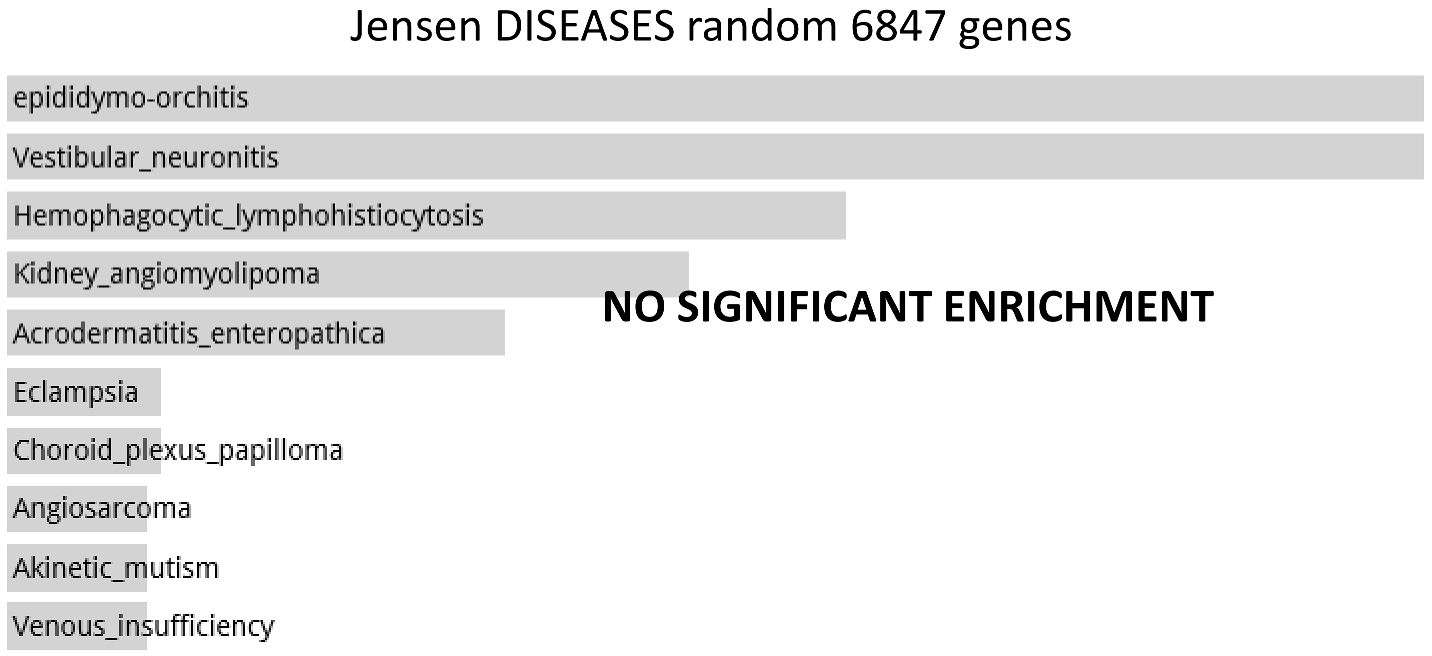
Jensen DISEASES random 6847 genes
epididymo-orchitis
Vestibular_neuronitis
Hemophagocytic_lymphohistiocytosis
Kidney_angiomyolipoma
Acrodermatitis_enteropathica
Eclampsia
Choroid_plexus_papilloma
Angiosarcoma
Akinetic_mutism
Venous_insufficiency
NO SIGNIFICANT ENRICHMENT

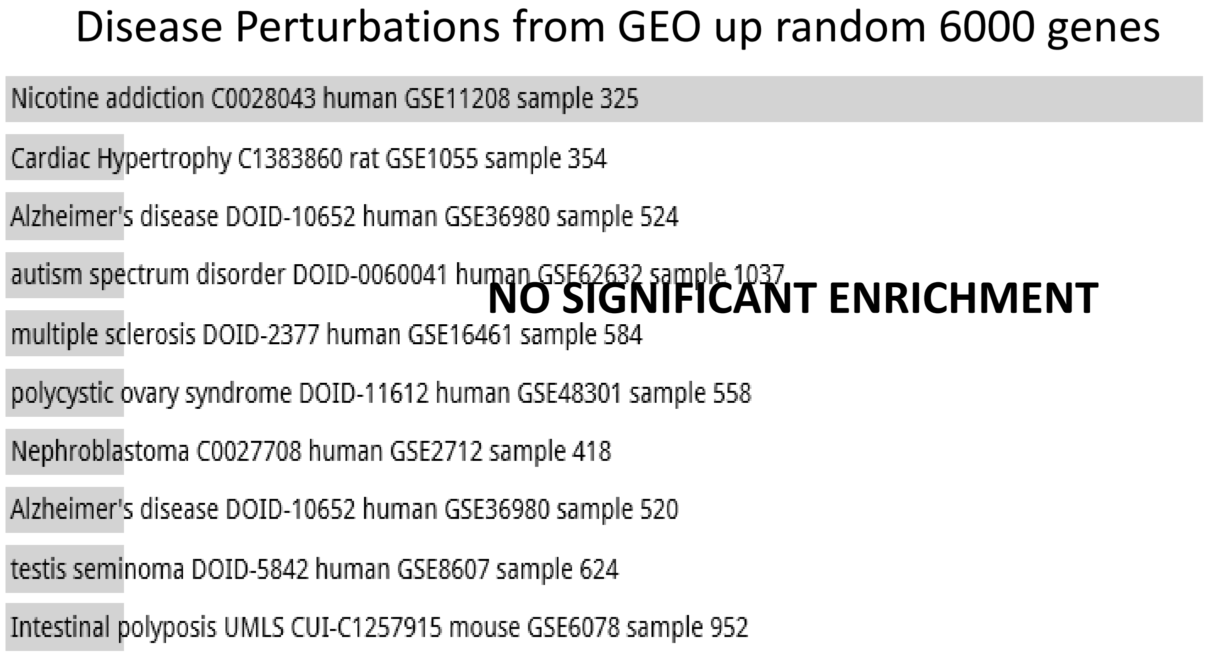
Disease Perturbations from GEO up random 6000 genes
Nicotine addiction C0028043 human GSE11208 sample 325
Cardiac Hypertrophy C1383860 rat GSE1055 sample 354
Alzheimer's disease DOID-10652 human GSE36980 sample 524
autism spectrum disorder DOID-0060041 human GSE62632 sample 1037
multiple sclerosis DOID-2377 human GSE16461 sample 584
polycystic ovary syndrome DOID-11612 human GSE48301 sample 558
Nephroblastoma C0027708 human GSE2712 sample 418
Alzheimer's disease DOID-10652 human GSE36980 sample 520
testis seminoma DOID-5842 human GSE8607 sample 624
Intestinal polyposis UMLS CUI-C1257915 mouse GSE6078 sample 952
NO SIGNIFICANT ENRICHMENT

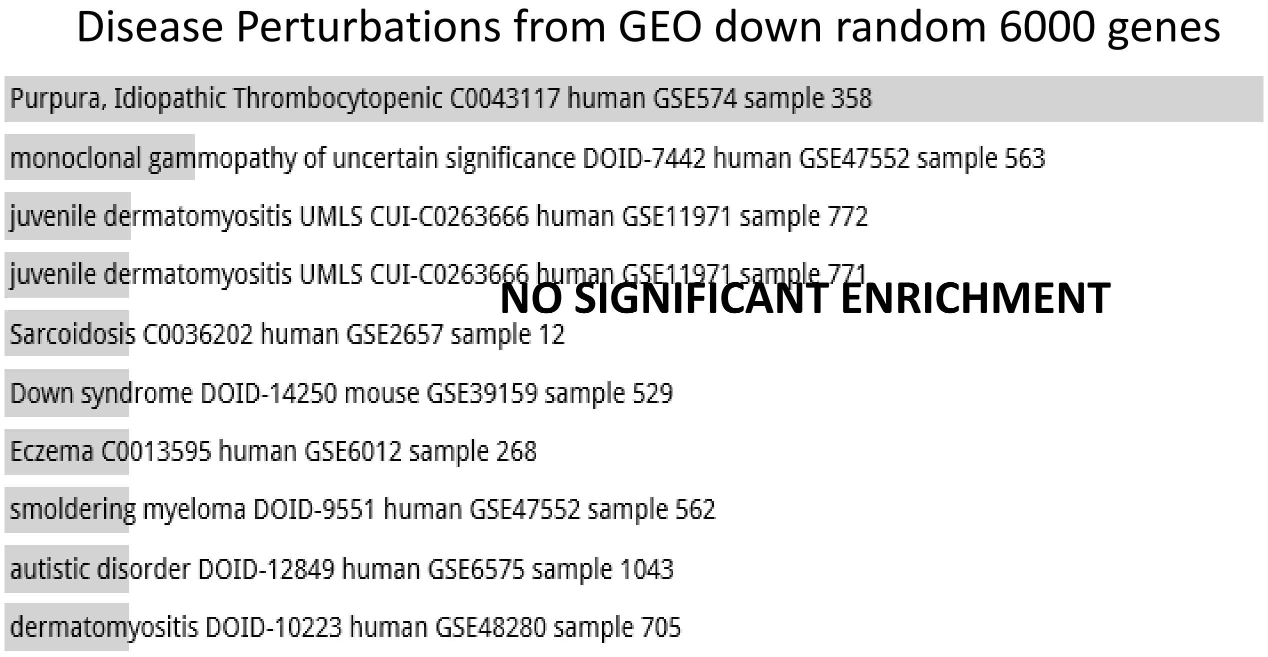
Disease Perturbations from GEO down random 6000 genes
Purpura, Idiopathic Thrombocytopenic C0043117 human GSE574 sample 358
monoclonal gammopathy of uncertain significance DOID-7442 human GSE47552 sample 563
juvenile dermatomyositis UMLS CUI-C0263666 human GSE11971 sample 772
juvenile dermatomyositis UMLS CUI-C0263666 human GSE11971 sample 771
Sarcoidosis C0036202 human GSE2657 sample 12
Down syndrome DOID-14250 mouse GSE39159 sample 529
Eczema C0013595 human GSE6012 sample 268
smoldering myeloma DOID-9551 human GSE47552 sample 562
autistic disorder DOID-12849 human GSE6575 sample 1043
dermatomyositis DOID-10223 human GSE48280 sample 705
NO SIGNIFICANT ENRICHMENT

**Random Set 7**

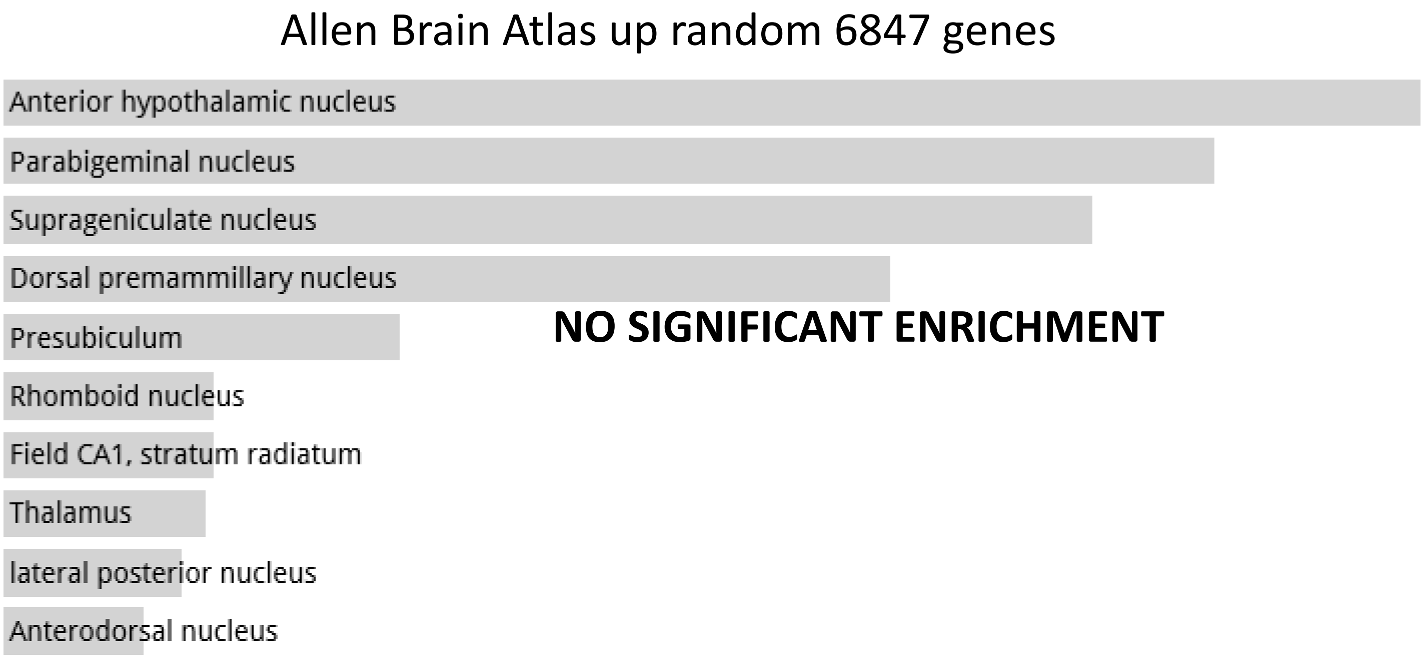
Allen Brain Atlas up random 6847 genes
Anterior hypothalamic nucleus
Parabigeminal nucleus
Suprageniculate nucleus
Dorsal premammillary nucleus
Presubiculum
Rhomboid nucleus
Field CA1, stratum radiatum
Thalamus
lateral posterior nucleus
Anterodorsal nucleus
NO SIGNIFICANT ENRICHMENT

**Random Set 8**

ARCHS4 Tissues random 6001 genes

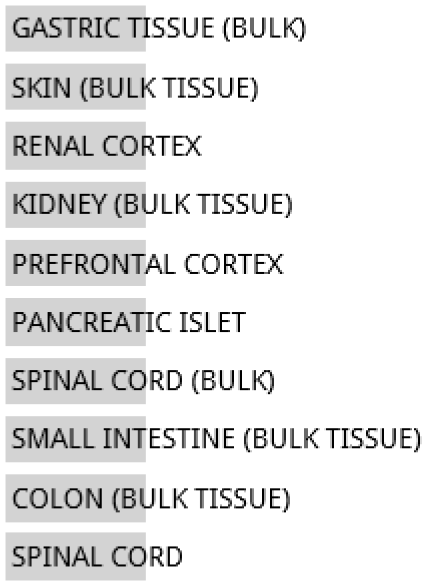


**NO SIGNIFICANT ENRICHMENT**

Jensen TISSUES random 6001 genes

**NO DATA AVAILABLE**

#### Random Set 8

Jensen DISEASES random 6001 genes

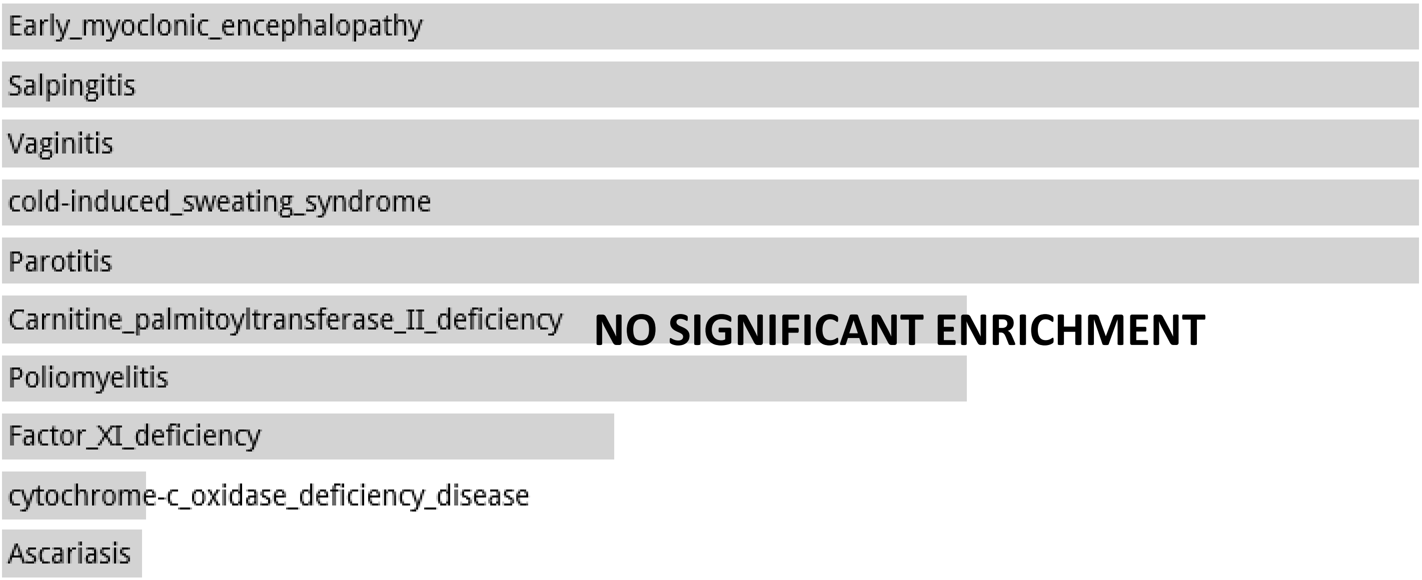


Disease Perturbations from GEO up random 6000 genes

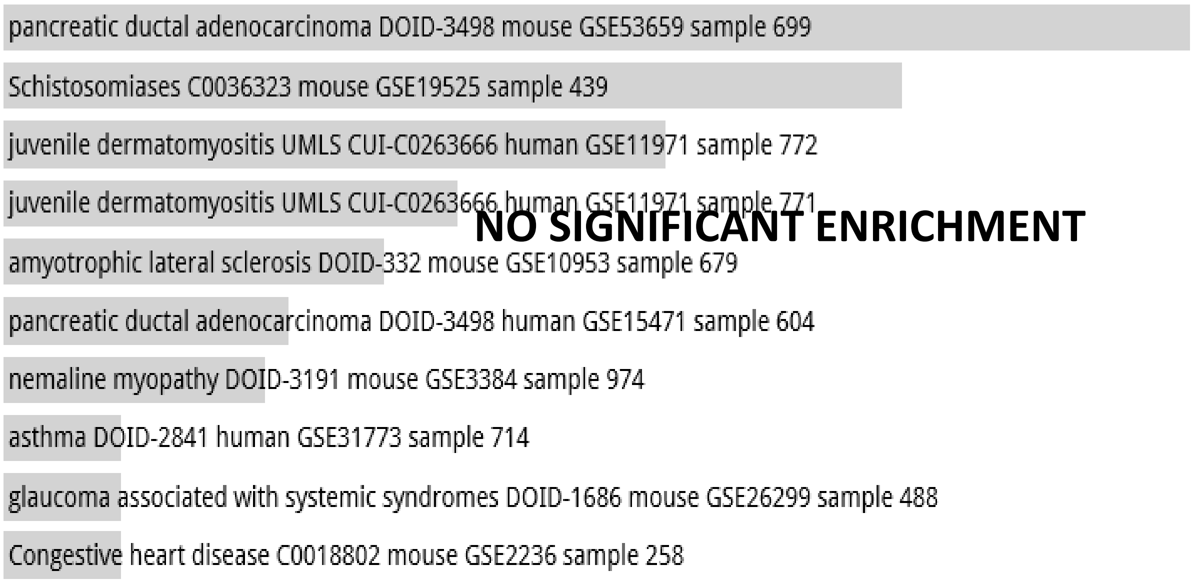


Disease Perturbations from GEO down random 6000 genes

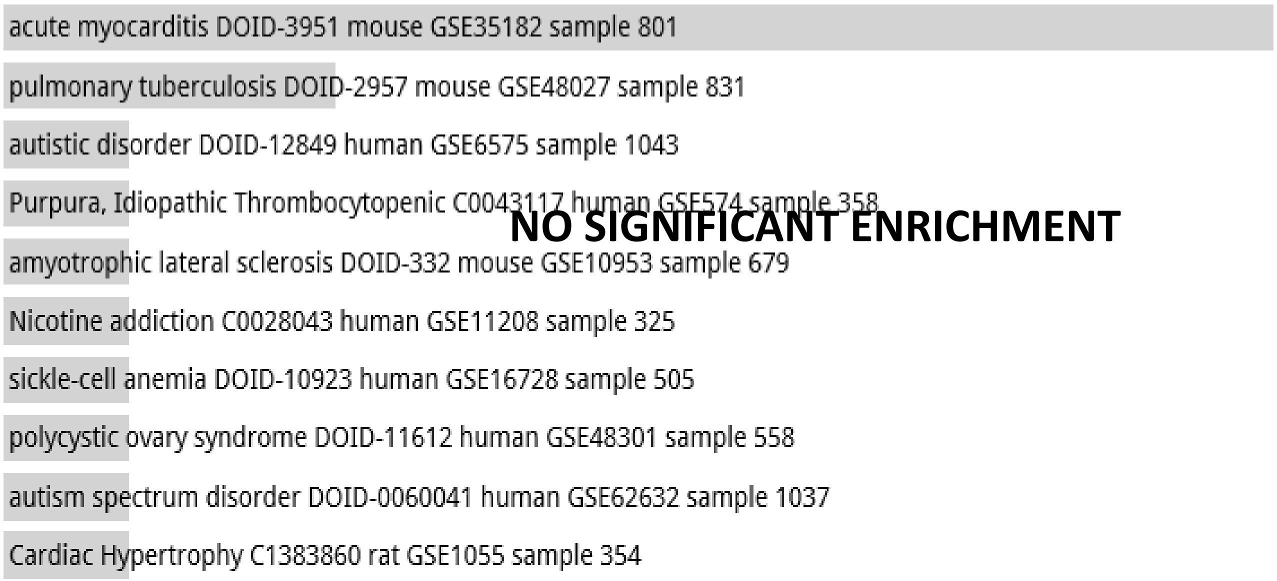

**Random Set 8**

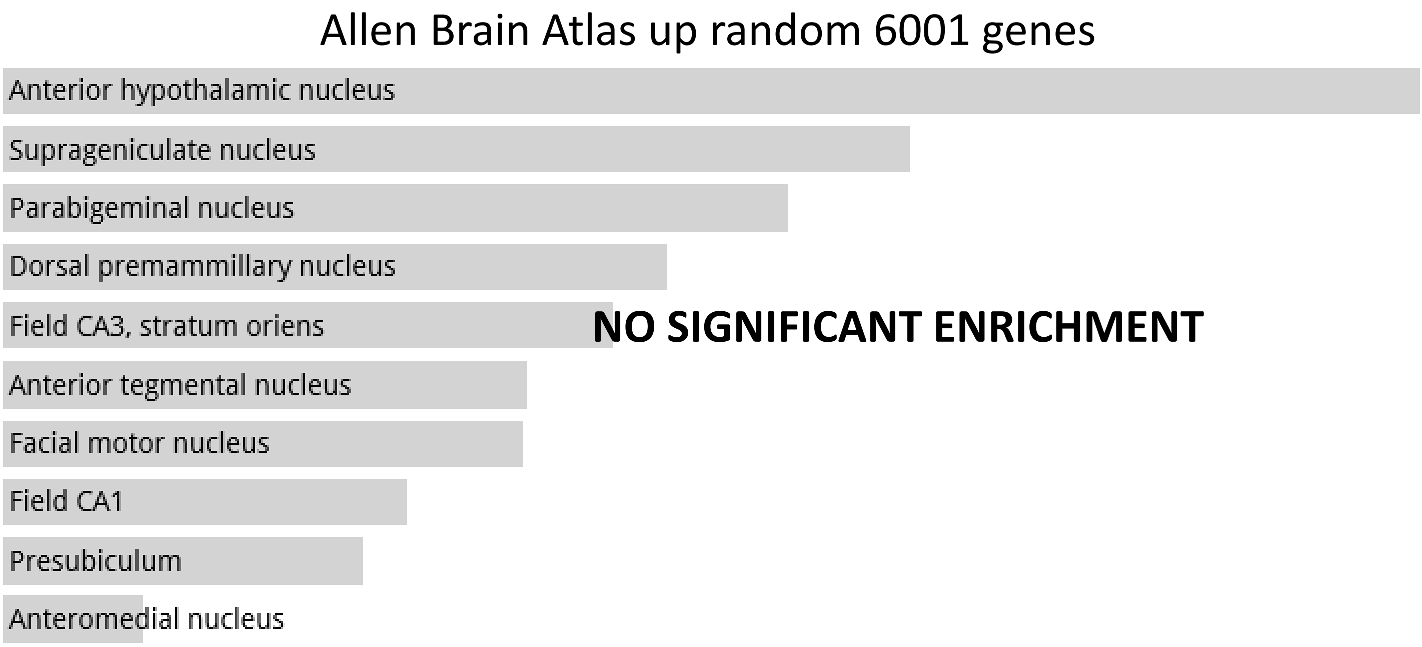
Allen Brain Atlas up random 6001 genes
Anterior hypothalamic nucleus
Suprageniculate nucleus
Parabigeminal nucleus
Dorsal premammillary nucleus
Field CA3, stratum oriens
Anterior tegmental nucleus
Facial motor nucleus
Field CA1
Presubiculum
Anteromedial nucleus
NO SIGNIFICANT ENRICHMENT

**Random Set 9**

ARCHS4 Tissues random 6000 genes

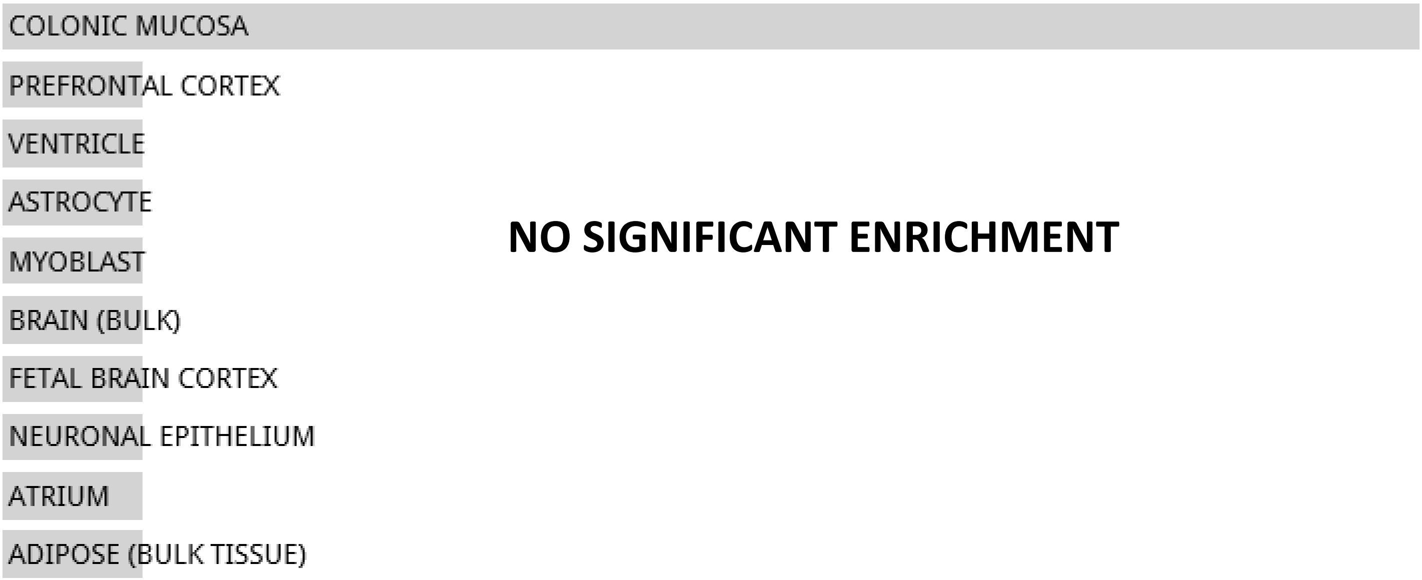


**NO SIGNIFICANT ENRICHMENT**

Jensen TISSUES random 6000 genes

**NO DATA AVAILABLE**

**Random Set 9**

#### Jensen DISEASES random 6000 genes

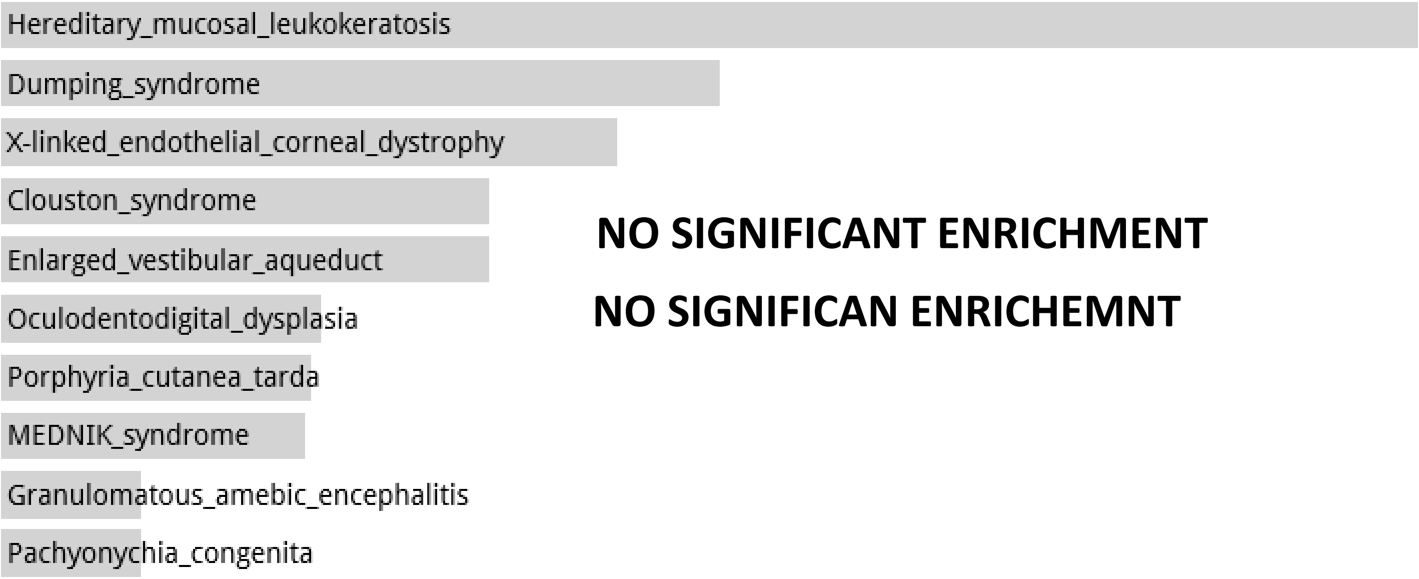


NO SIGNIFICANT ENRICHMENT

NO SIGNIFICAN ENRICHEMNT

#### Disease Perturbations from GEO up random 6000 genes

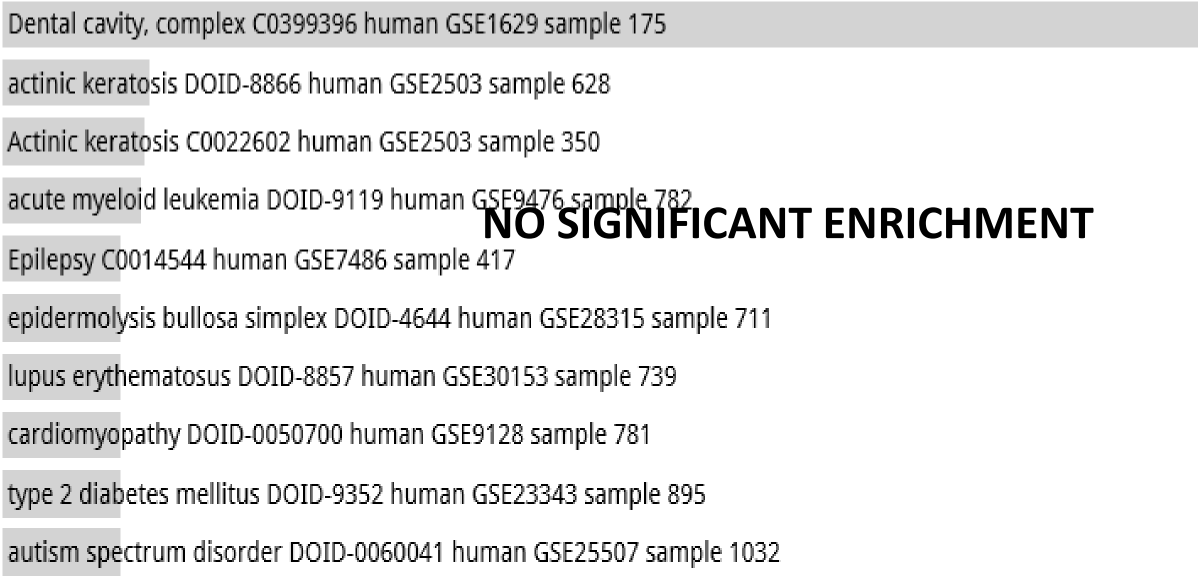


NO SIGNIFICANT ENRICHMENT

#### Disease Perturbations from GEO down random 6000 genes

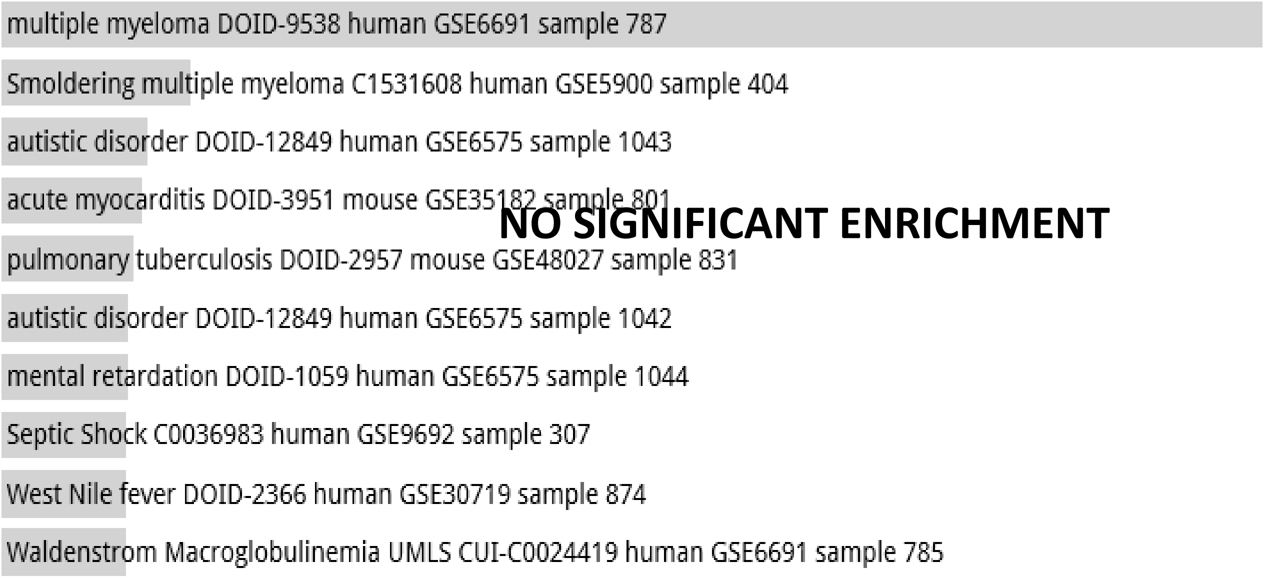


NO SIGNIFICANT ENRICHMENT

**Random Set 9**

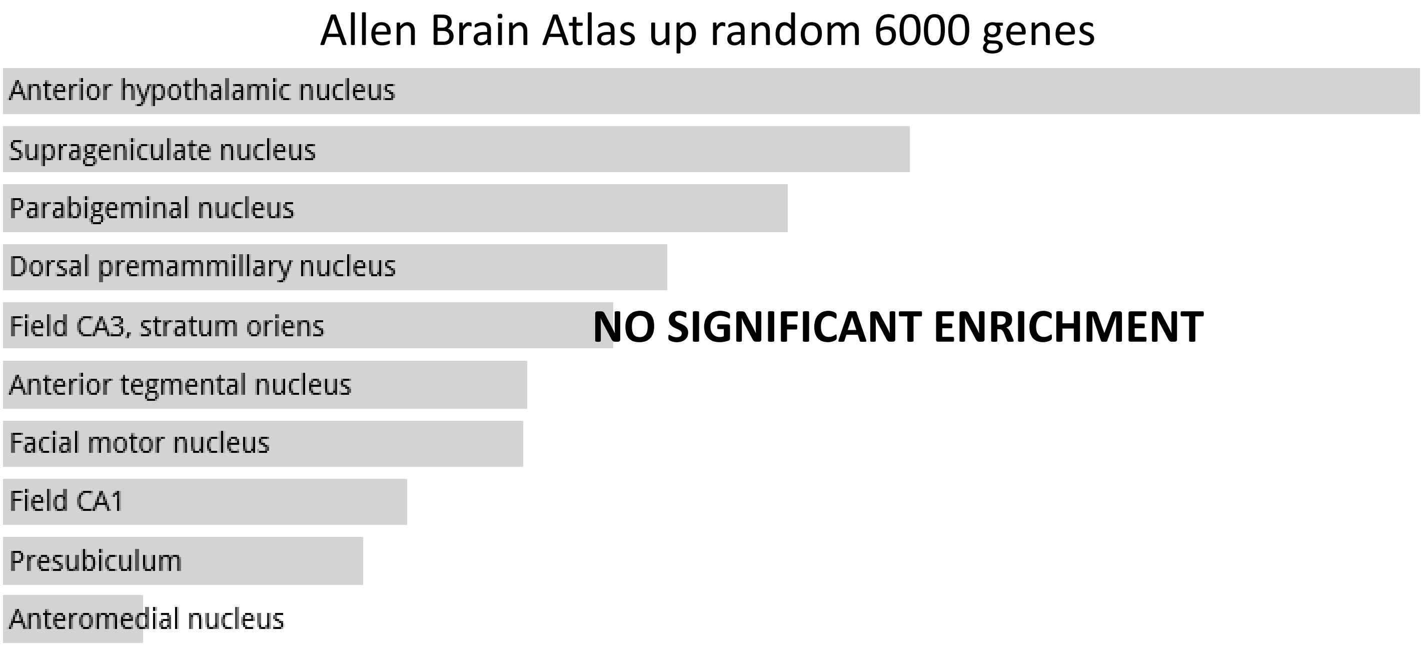
Allen Brain Atlas up random 6000 genes
Anterior hypothalamic nucleus
Suprageniculate nucleus
Parabigeminal nucleus
Dorsal premammillary nucleus
Field CA3, stratum oriens
Anterior tegmental nucleus
Facial motor nucleus
Field CA1
Presubiculum
Anteromedial nucleus
NO SIGNIFICANT ENRICHMENT

**Random Set 10**

ARCHS4 Tissues random 6000 genes

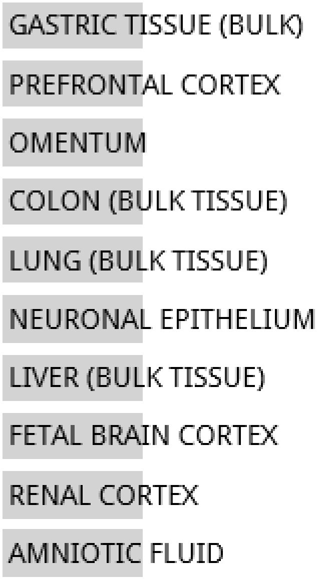


**NO SIGNIFICANT ENRICHMENT**

Jensen TISSUES random 6000 genes

**NO DATA AVAILABLE**

**Random Set 10**

Jensen DISEASES random 6000 genes

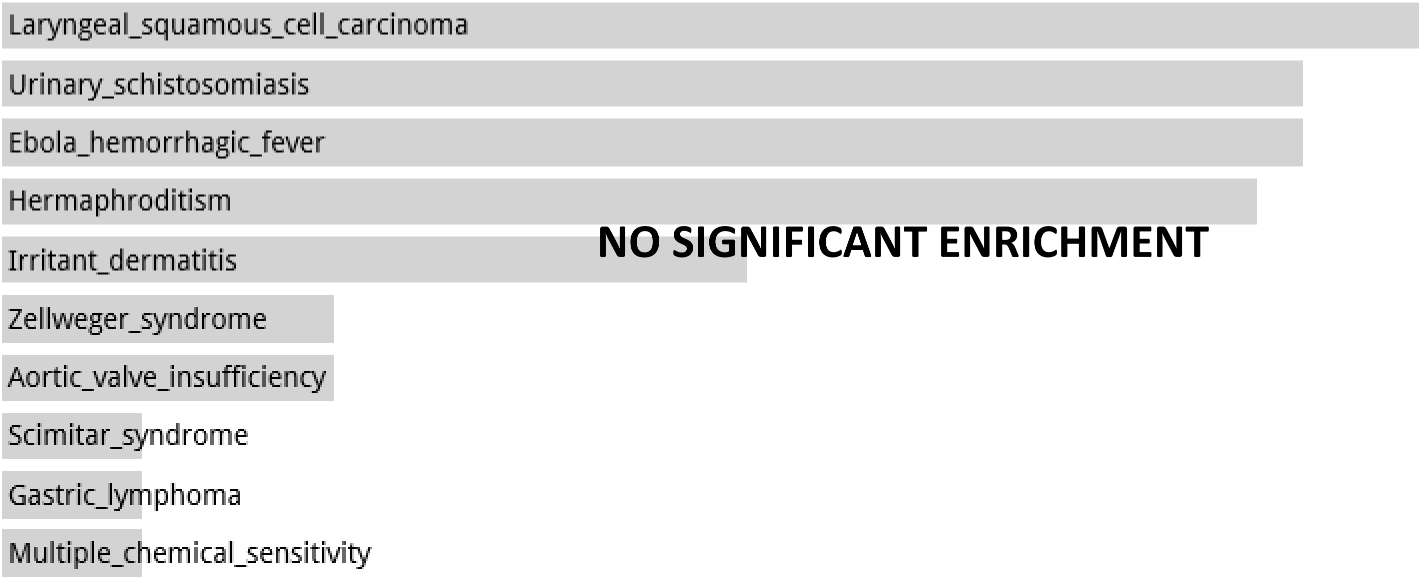


Disease Perturbations from GEO up random 6000 genes

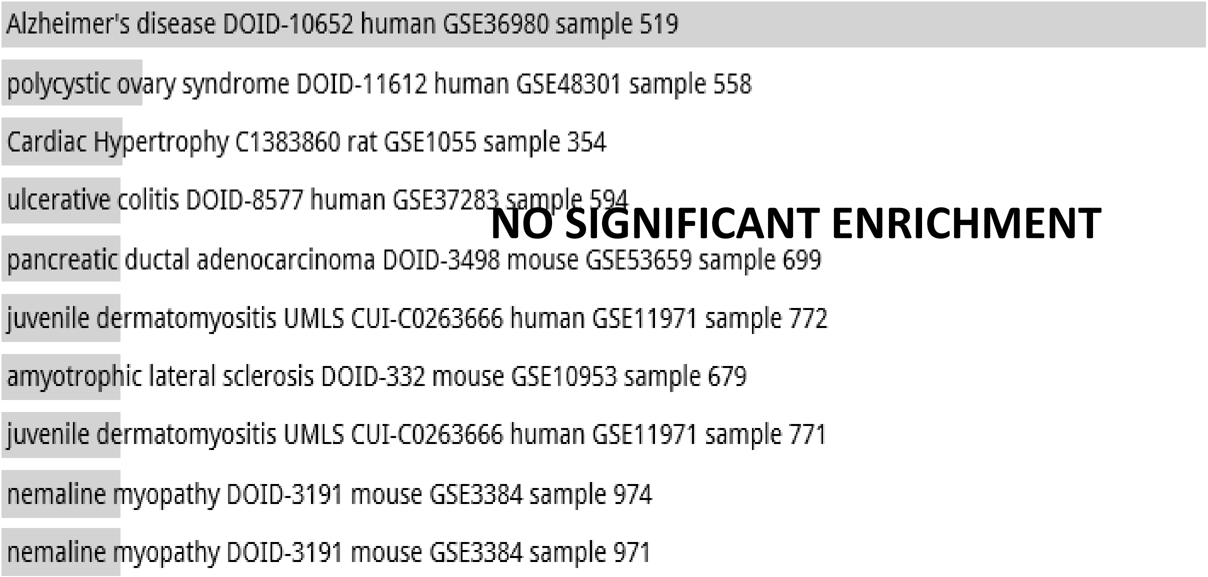


Disease Perturbations from GEO down random 6000 genes

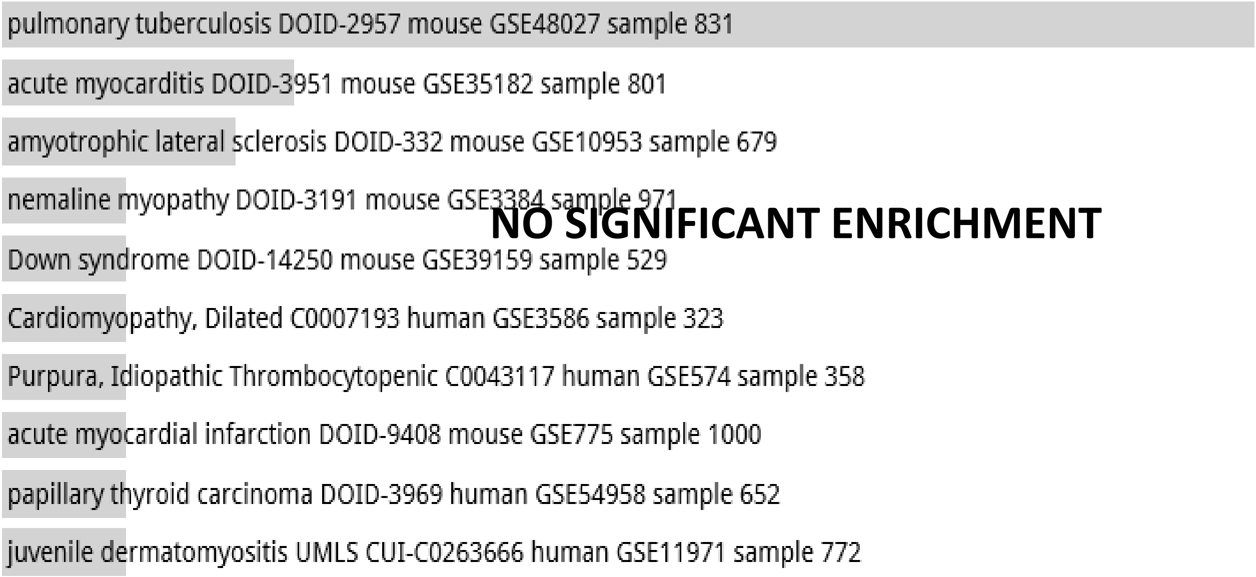

**Random Set 10**

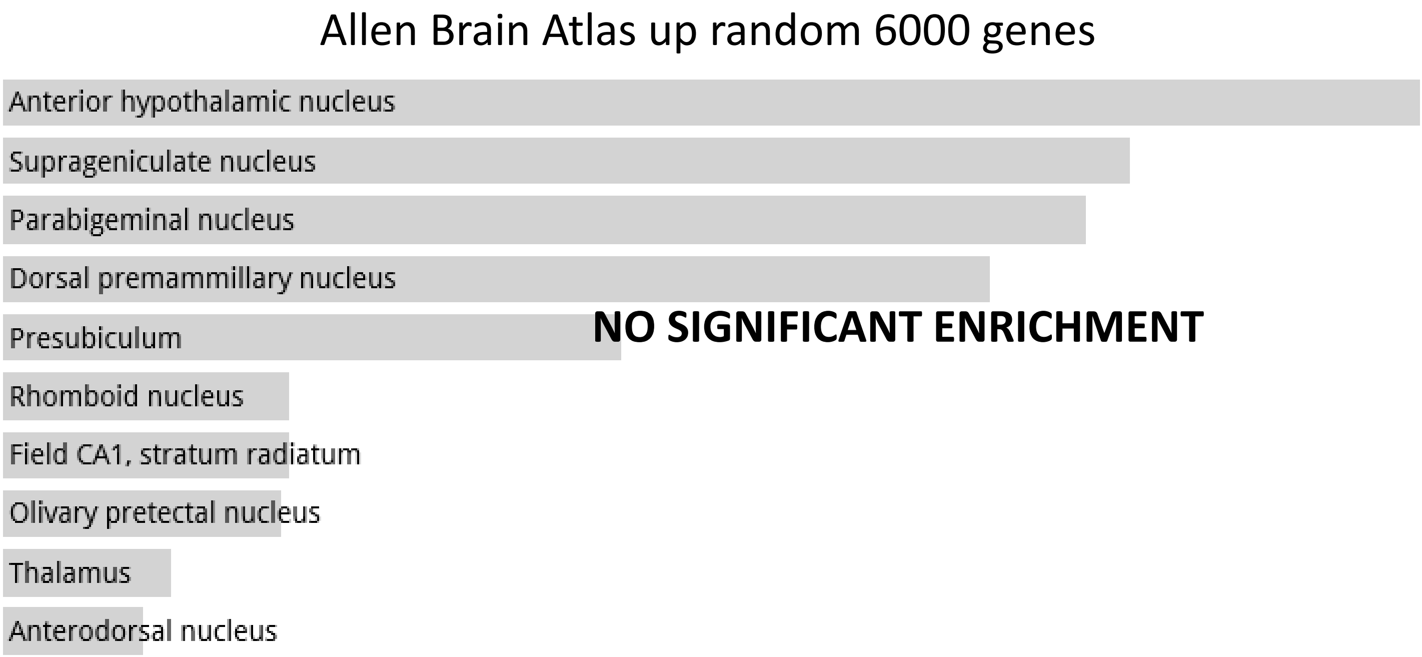

**Random Set 11**

ARCHS4 Tissues random 4000 genes

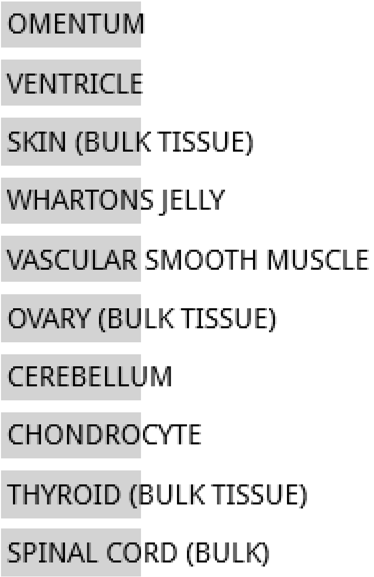


**NO SIGNIFICANT ENRICHMENT**

Jensen TISSUES random 4000 genes

**NO DATA AVAILABLE**

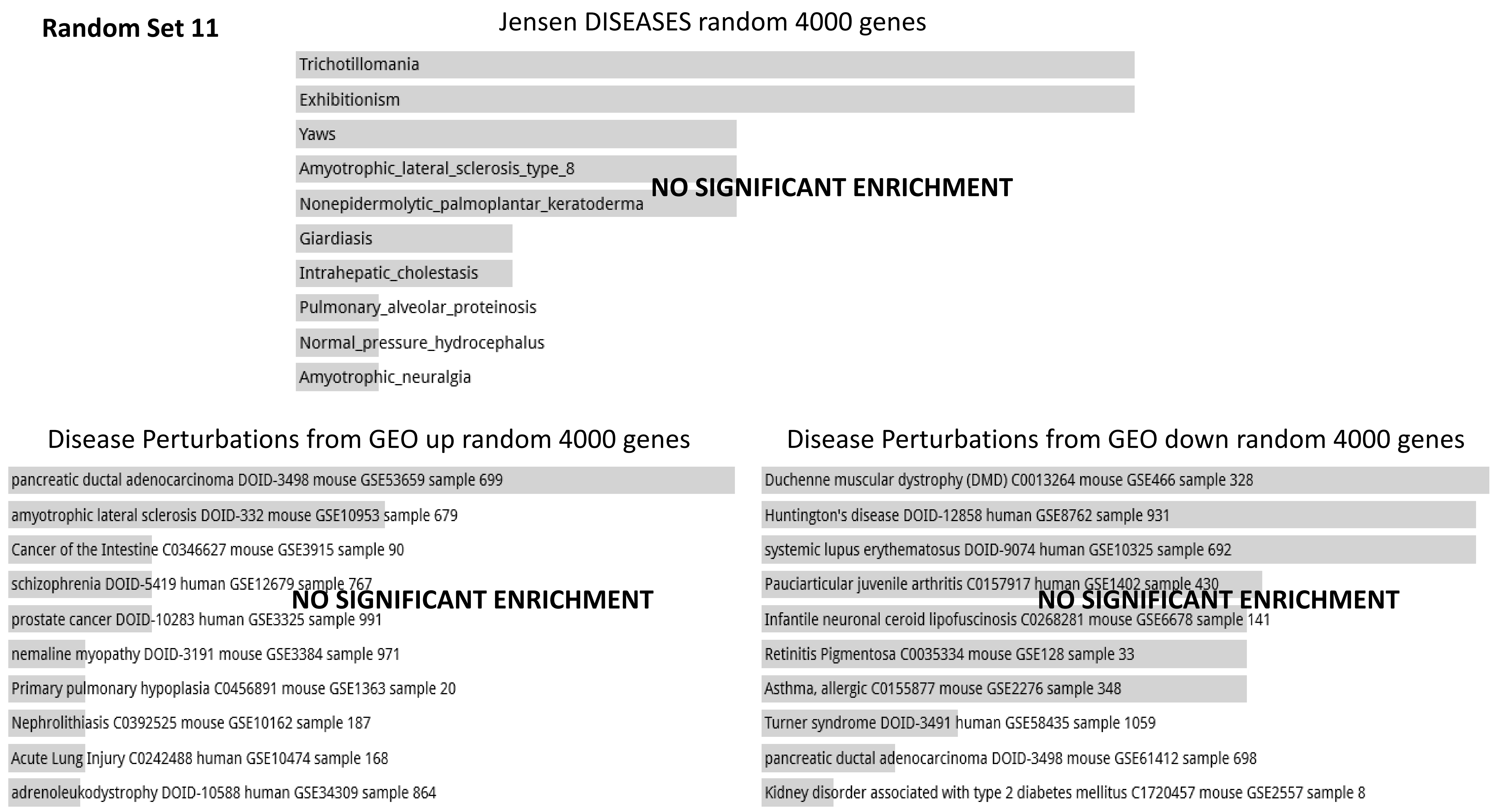
Random Set 11
Jensen DISEASES random 4000 genes
Trichotillomania
Exhibitionism
Yaws
Amyotrophic_lateral_sclerosis_type_8
Nonepidermolytic_palmoplantar_keratoderma
Giardiasis
Intrahepatic_cholestasis
Pulmonary_alveolar_proteinosis
Normal_pressure_hydrocephalus
Amyotrophic_neuralgia
NO SIGNIFICANT ENRICHMENT
Disease Perturbations from GEO up random 4000 genes
pancreatic ductal adenocarcinoma DOID-3498 mouse GSE53659 sample 699
amyotrophic lateral sclerosis DOID-332 mouse GSE10953 sample 679
Cancer of the Intestine C0346627 mouse GSE3915 sample 90
schizophrenia DOID-5419 human GSE12679 sample 767
prostate cancer DOID-10283 human GSE3325 sample 991
nemaline myopathy DOID-3191 mouse GSE3384 sample 971
Primary pulmonary hypoplasia C0456891 mouse GSE1363 sample 20
Nephrolithiasis C0392525 mouse GSE10162 sample 187
Acute Lung Injury C0242488 human GSE10474 sample 168
adrenoleukodystrophy DOID-10588 human GSE34309 sample 864
NO SIGNIFICANT ENRICHMENT
Disease Perturbations from GEO down random 4000 genes
Duchenne muscular dystrophy (DMD) C0013264 mouse GSE466 sample 328
Huntington's disease DOID-12858 human GSE8762 sample 931
systemic lupus erythematosus DOID-9074 human GSE10325 sample 692
Pauciarticular juvenile arthritis C0157917 human GSE1402 sample 430
Infantile neuronal ceroid lipofuscinosis C0268281 mouse GSE6678 sample 141
Retinitis Pigmentosa C0035334 mouse GSE128 sample 33
Asthma, allergic C0155877 mouse GSE2276 sample 348
Turner syndrome DOID-3491 human GSE58435 sample 1059
pancreatic ductal adenocarcinoma DOID-3498 mouse GSE61412 sample 698
Kidney disorder associated with type 2 diabetes mellitus C1720457 mouse GSE2557 sample 8
NO SIGNIFICANT ENRICHMENT

**Random Set 11**

Allen Brain Atlas up random 4000 genes

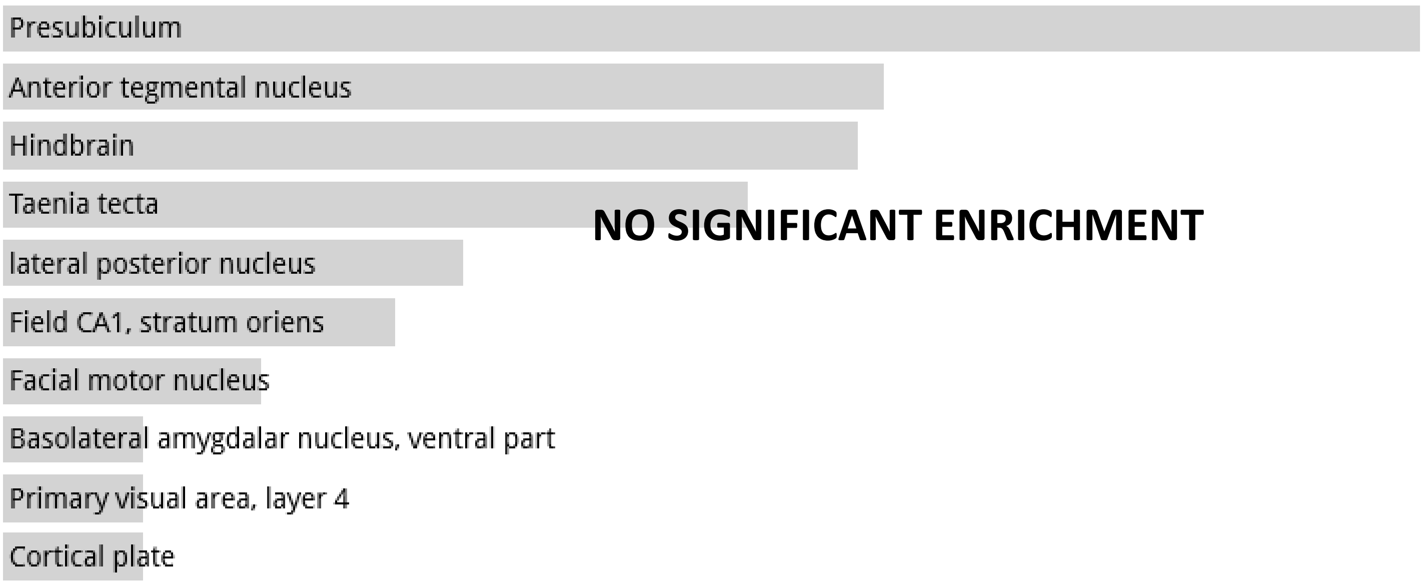

**Random Set 12**

ARCHS4 Tissues random 4000 genes

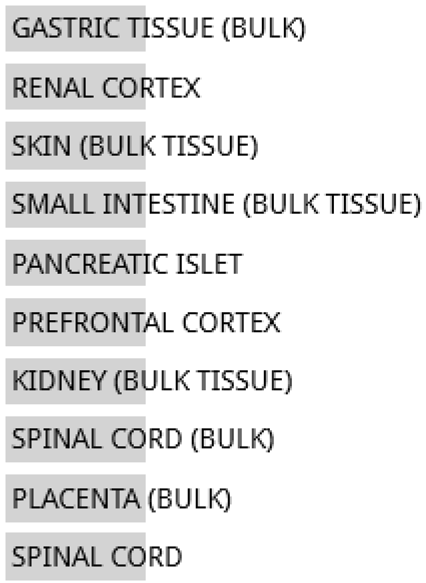


**NO SIGNIFICANT ENRICHMENT**

Jensen TISSUES random 4000 genes

**NO DATA AVAILABLE**

**Random Set 12**

#### Jensen DISEASES random 4000 genes

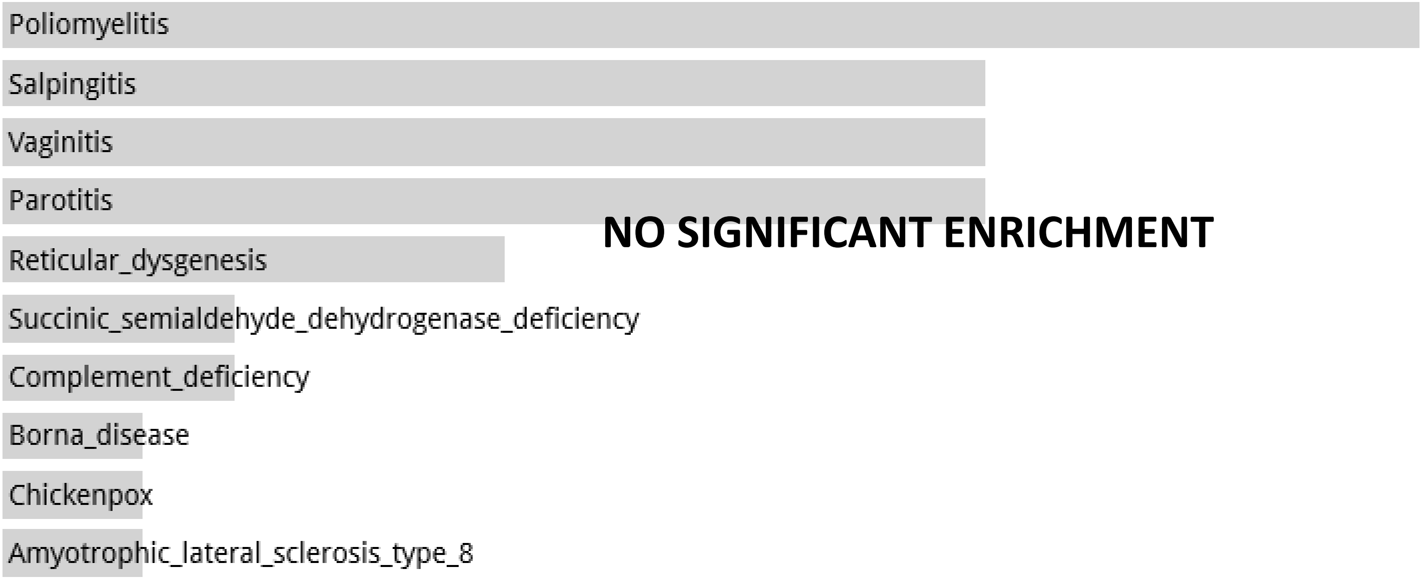


#### Disease Perturbations from GEO up random 4000 genes

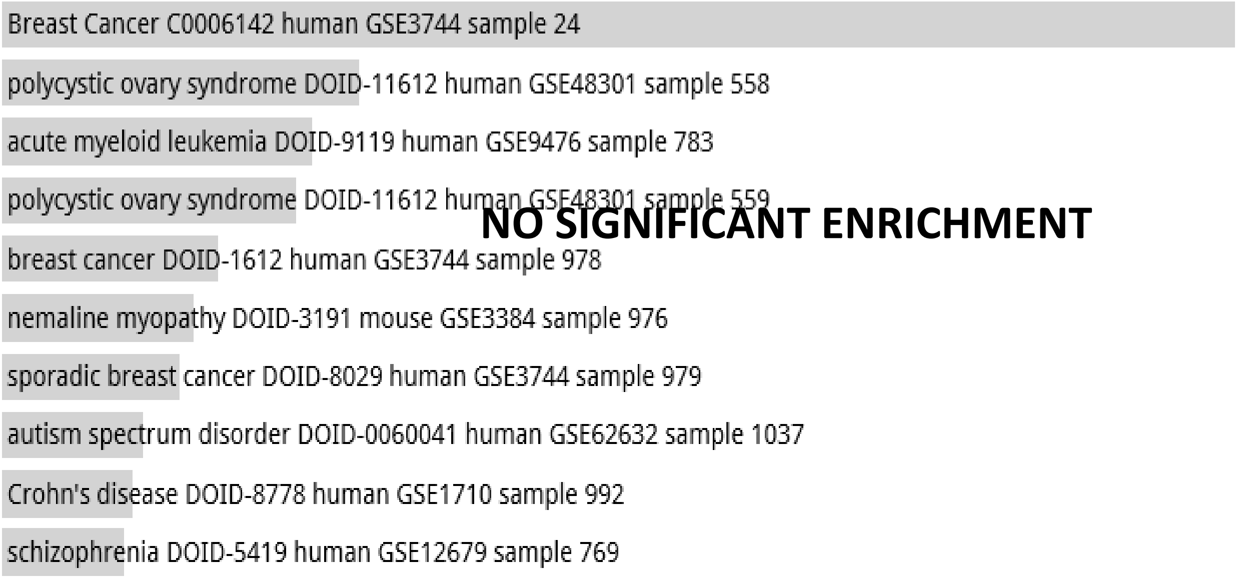


#### Disease Perturbations from GEO down random 4000 genes

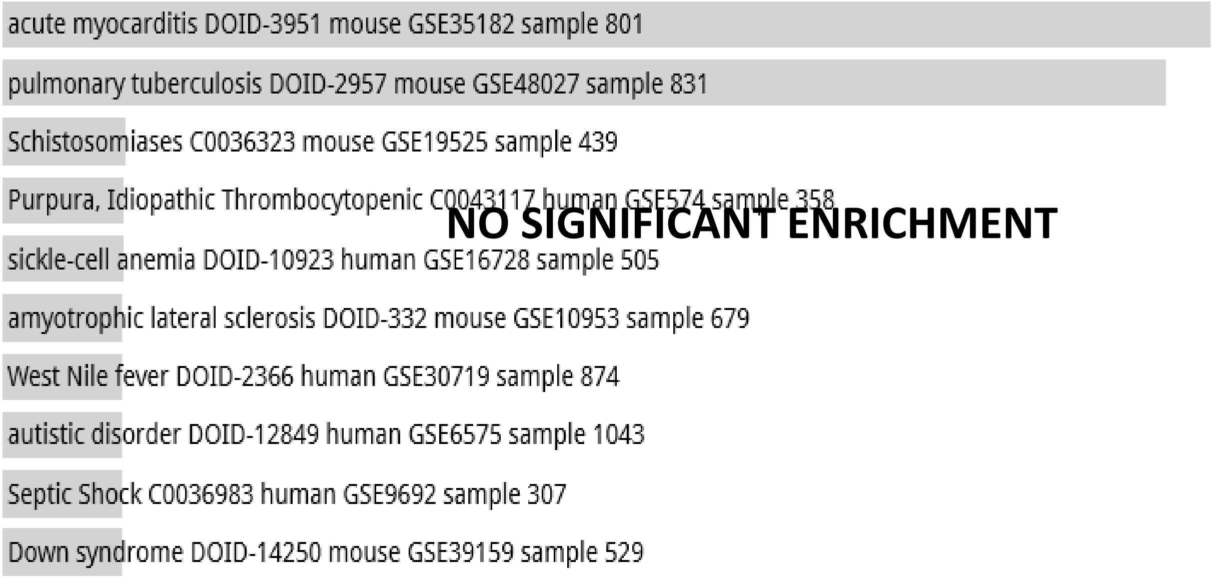

**Random Set 12**

### Allen Brain Atlas up random 4000 genes

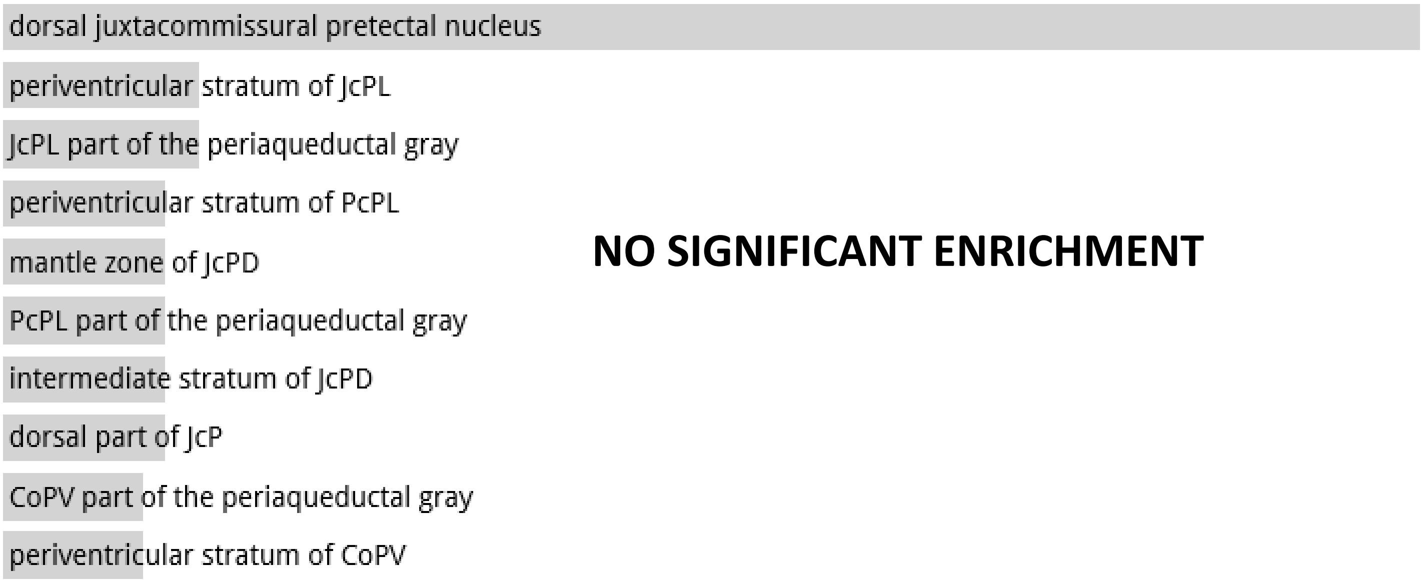


**NO SIGNIFICANT ENRICHMENT**

**Random Set 13**

ARCHS4 Tissues random 4000 genes

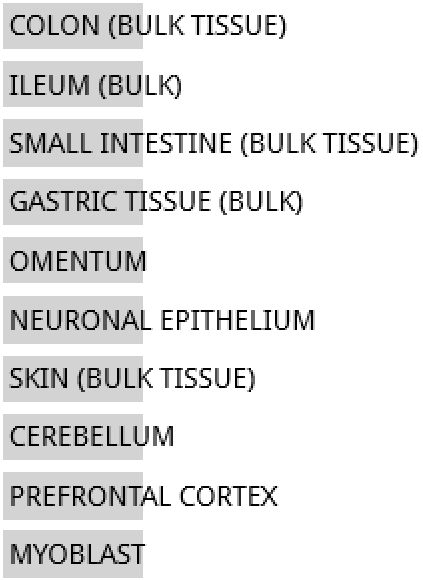


**NO SIGNIFICANT ENRICHMENT**

Jensen TISSUES random 4000 genes

**NO DATA AVAILABLE**

**Random Set 13**

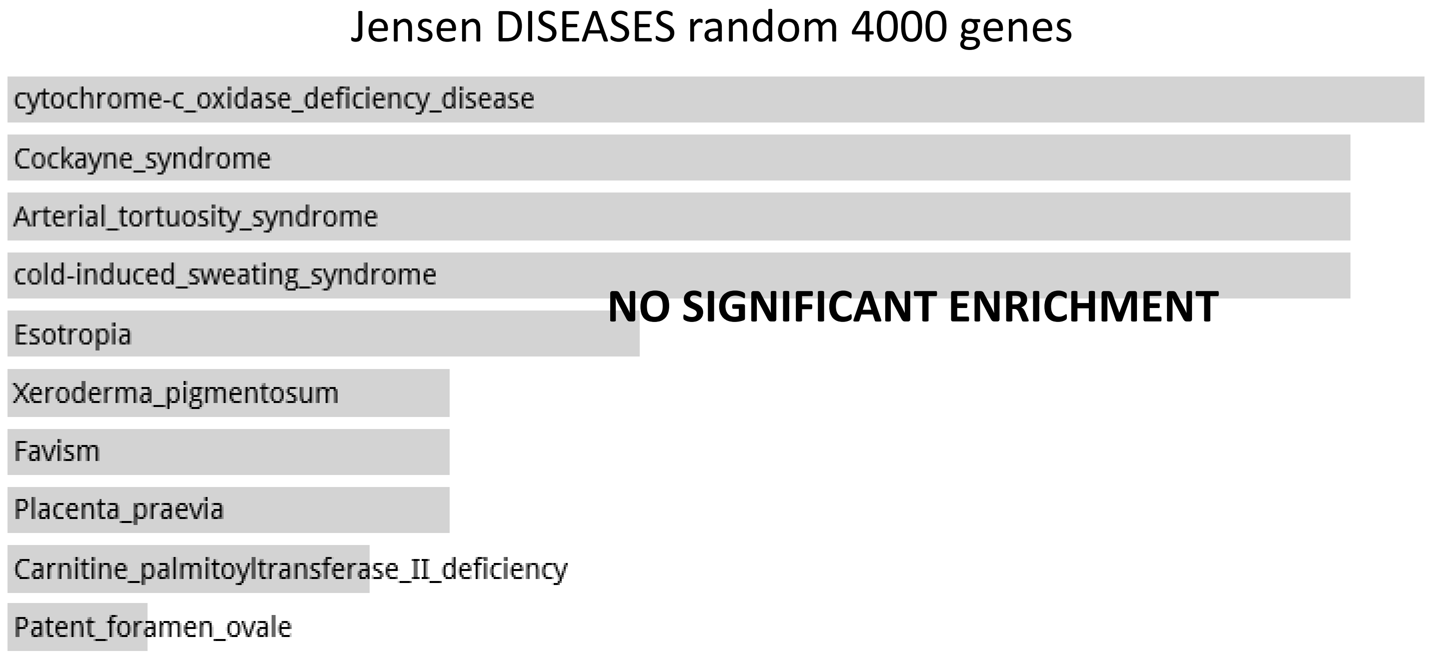


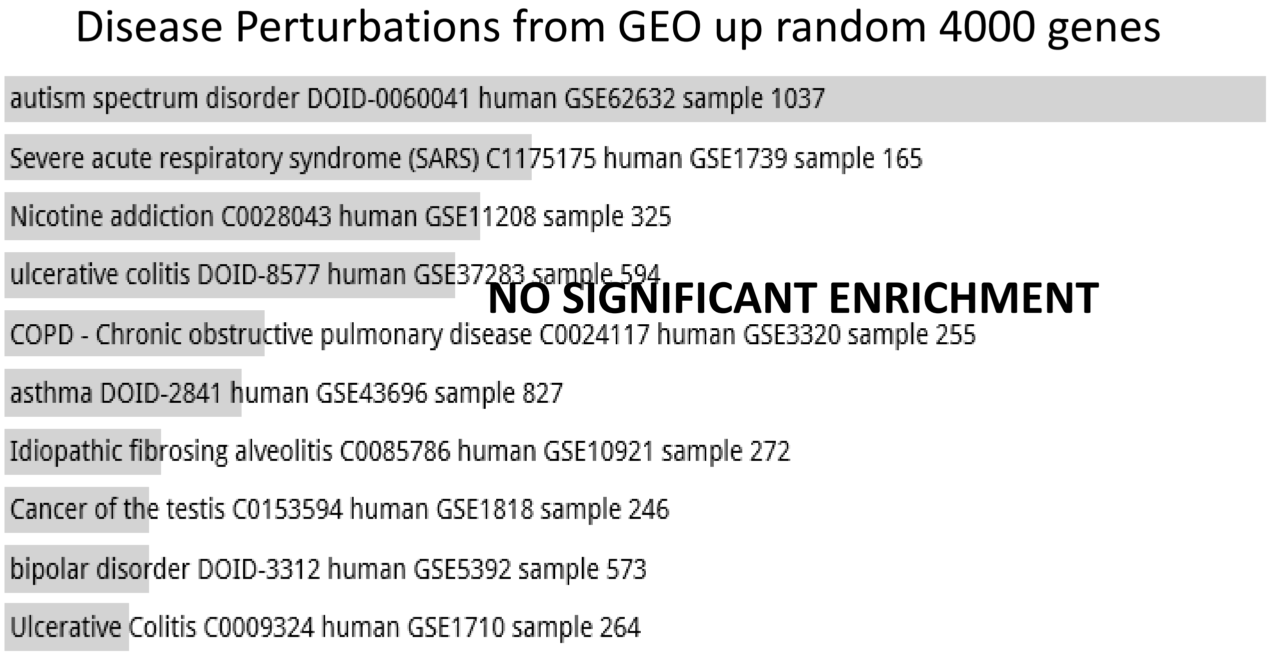


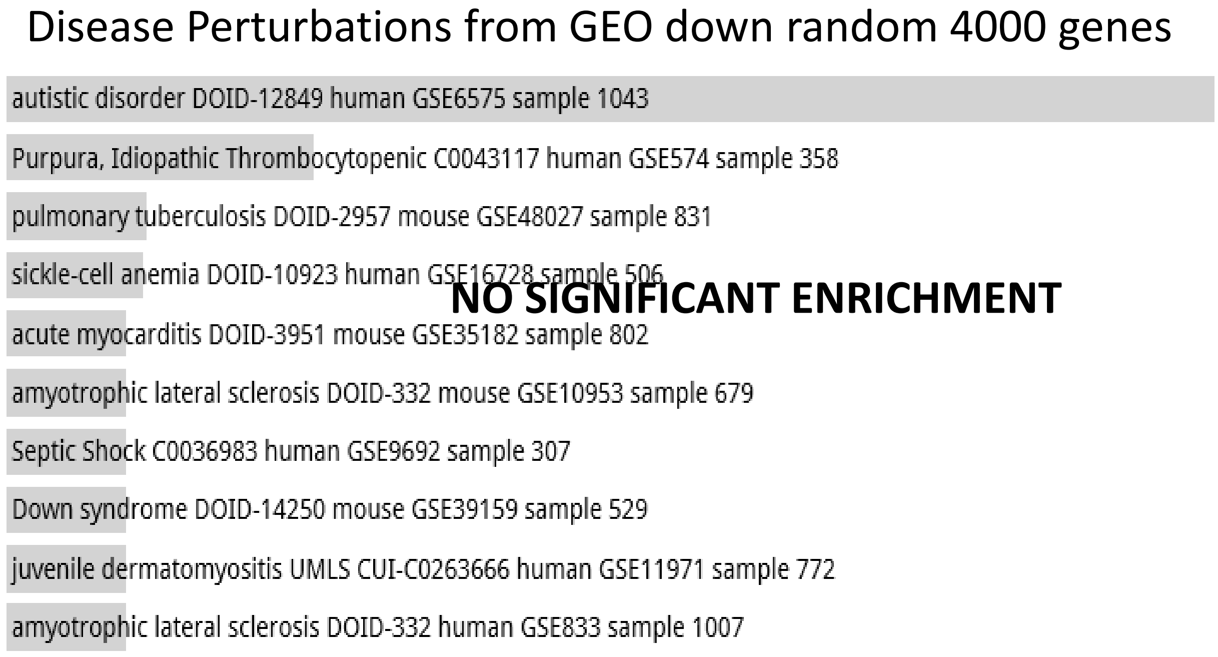

**Random Set 13**

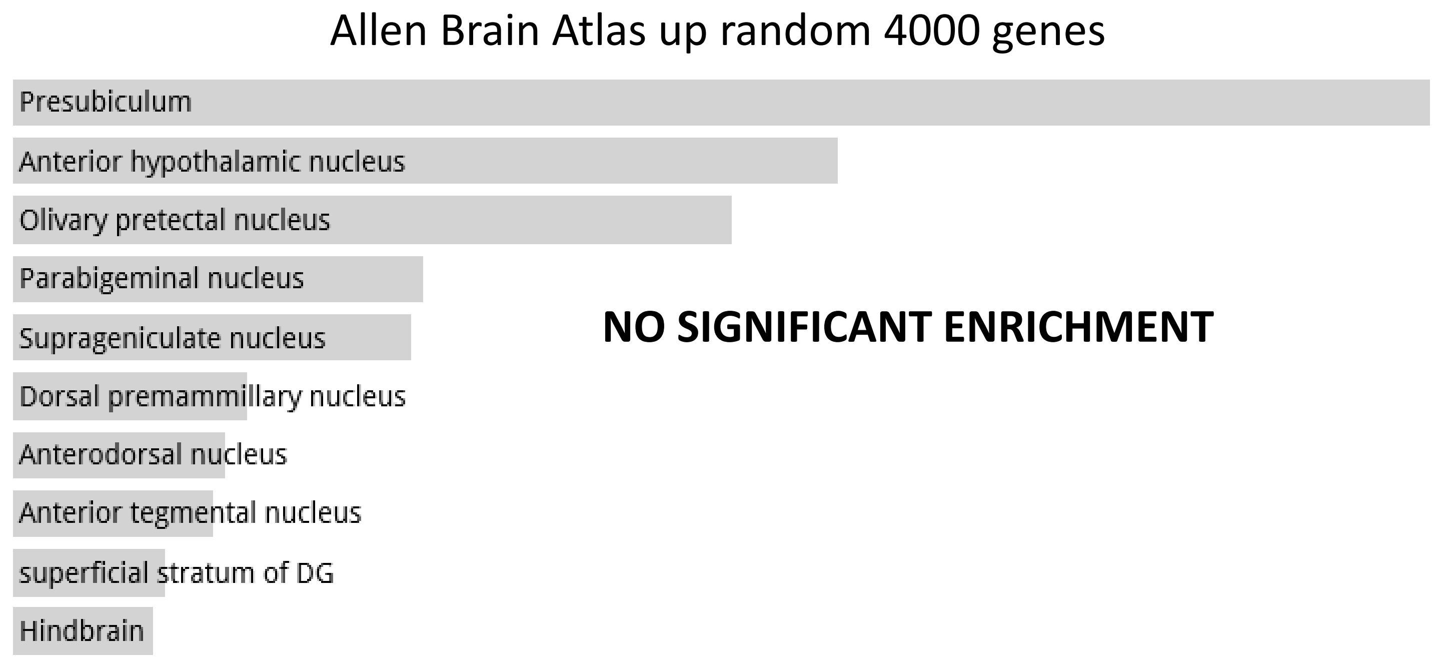

**Random Set 14**

ARCHS4 Tissues random 4000 genes

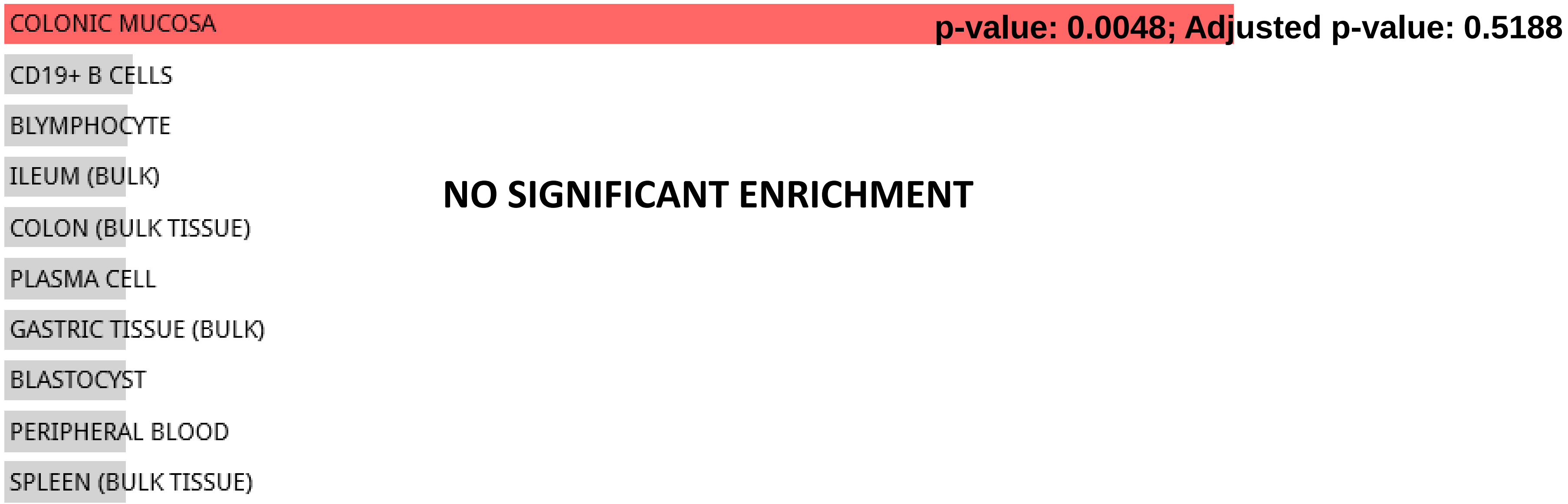


Jensen TISSUES random 4000 genes

**NO DATA AVAILABLE**

**Random Set 14**

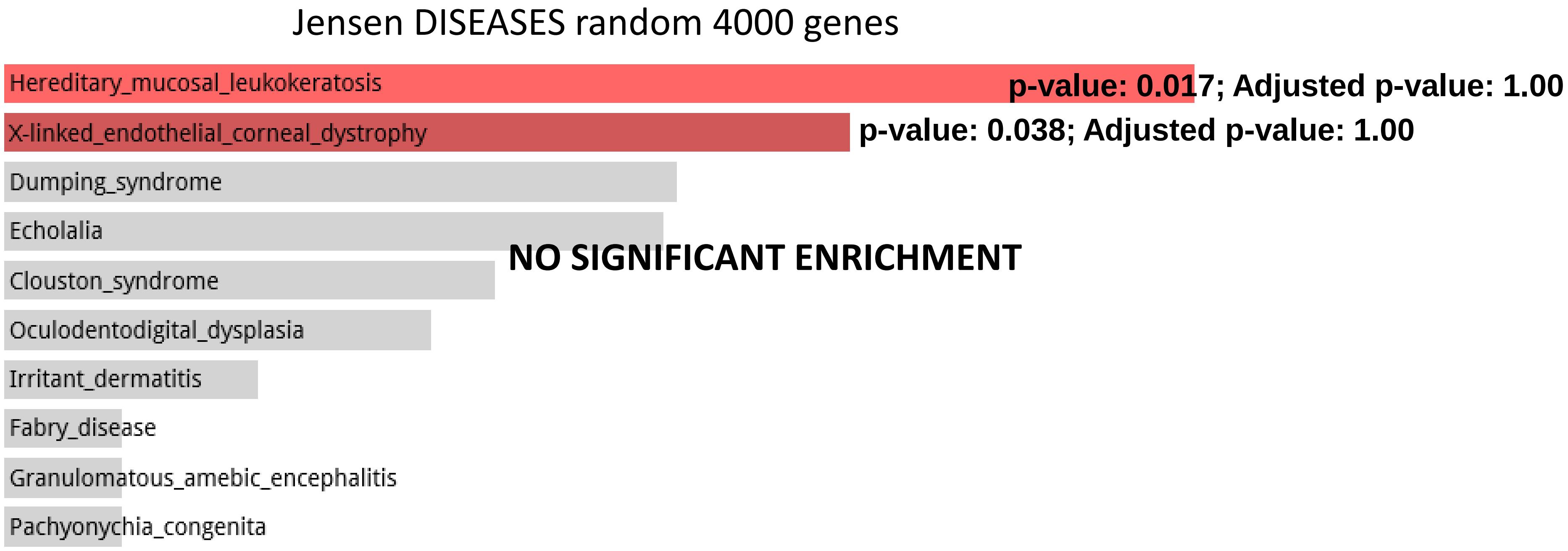


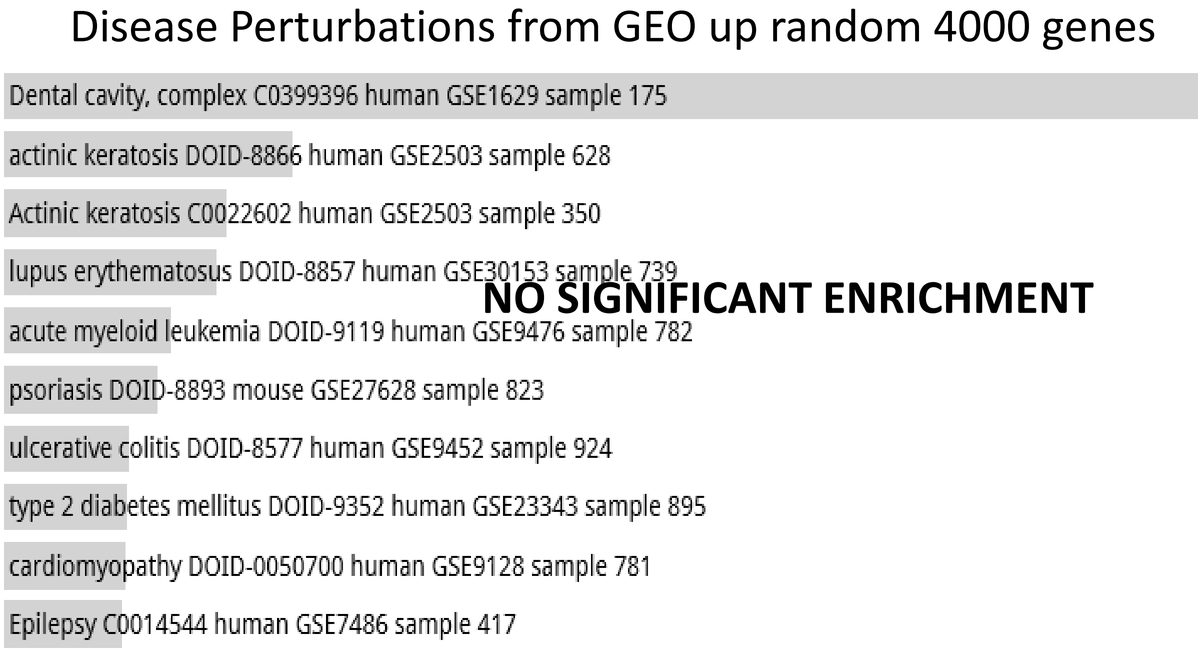


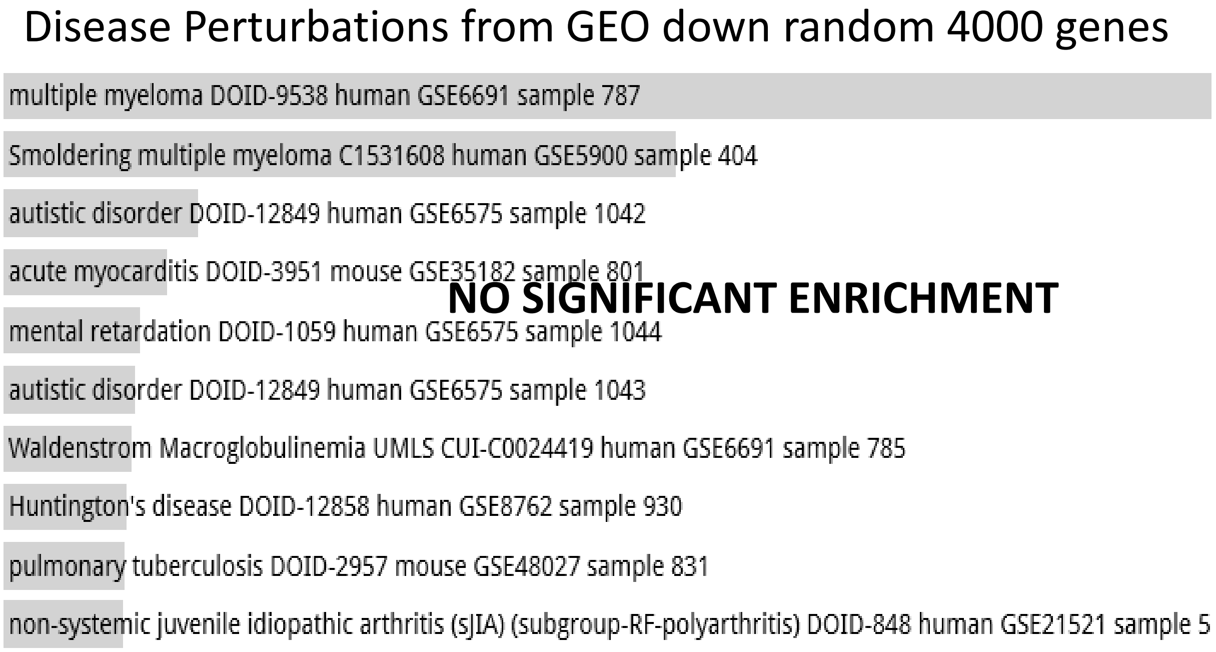

**Random Set 14**

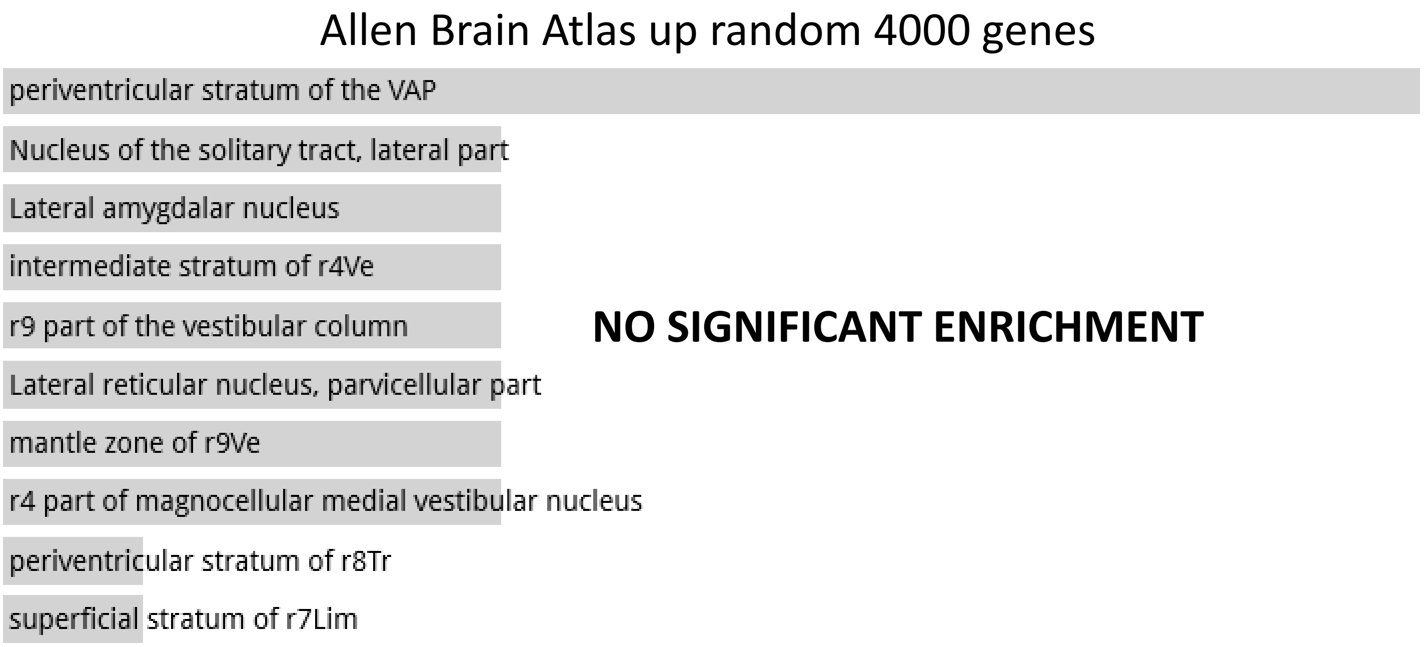

**Random Set 15**

ARCHS4 Tissues random 4000 genes

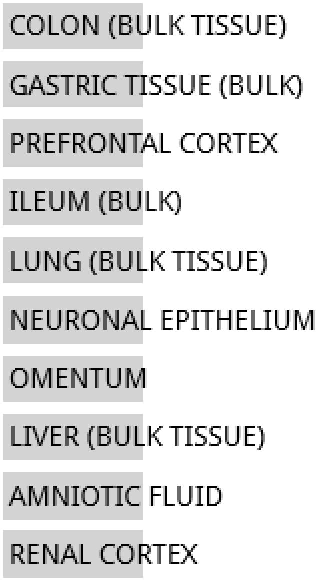


**NO SIGNIFICANT ENRICHMENT**

Jensen TISSUES random 4000 genes

**NO DATA AVAILABLE**

**Random Set 15**

Jensen DISEASES random 4000 genes

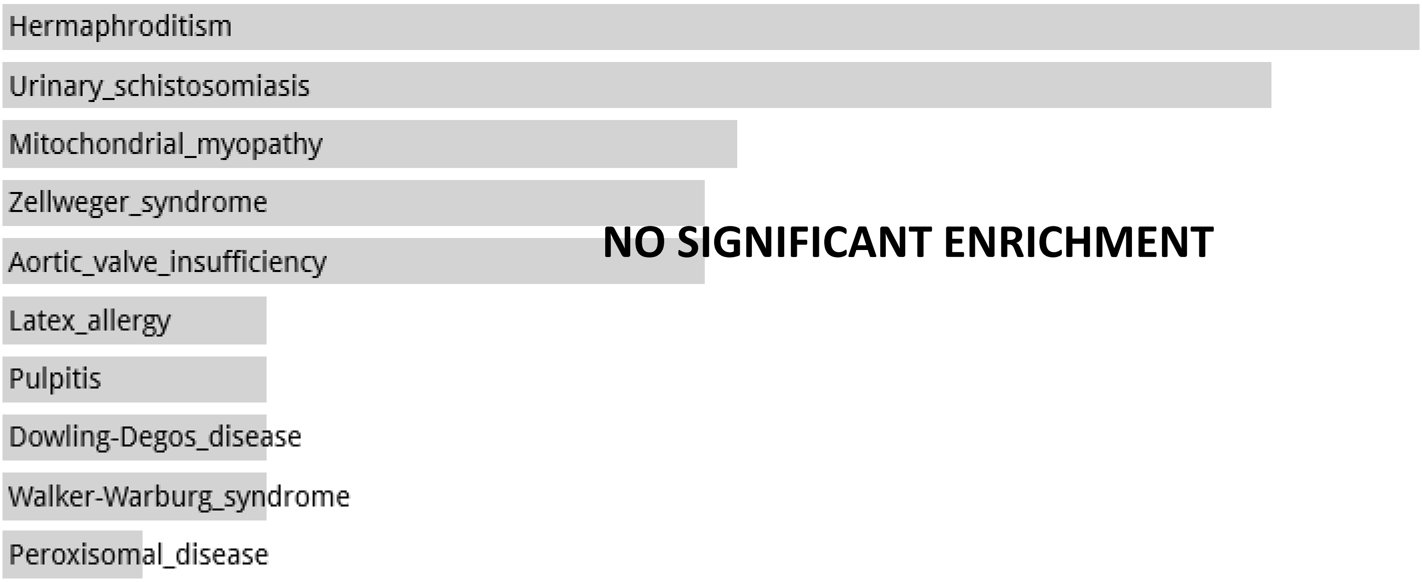


Disease Perturbations from GEO up random 4000 genes

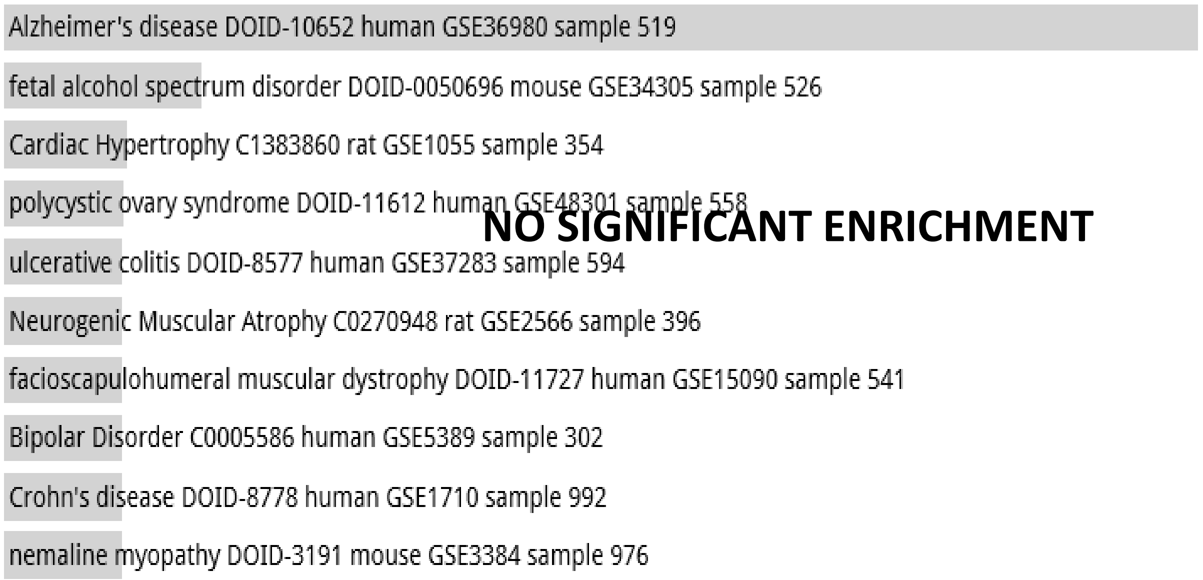


Disease Perturbations from GEO down random 4000 genes

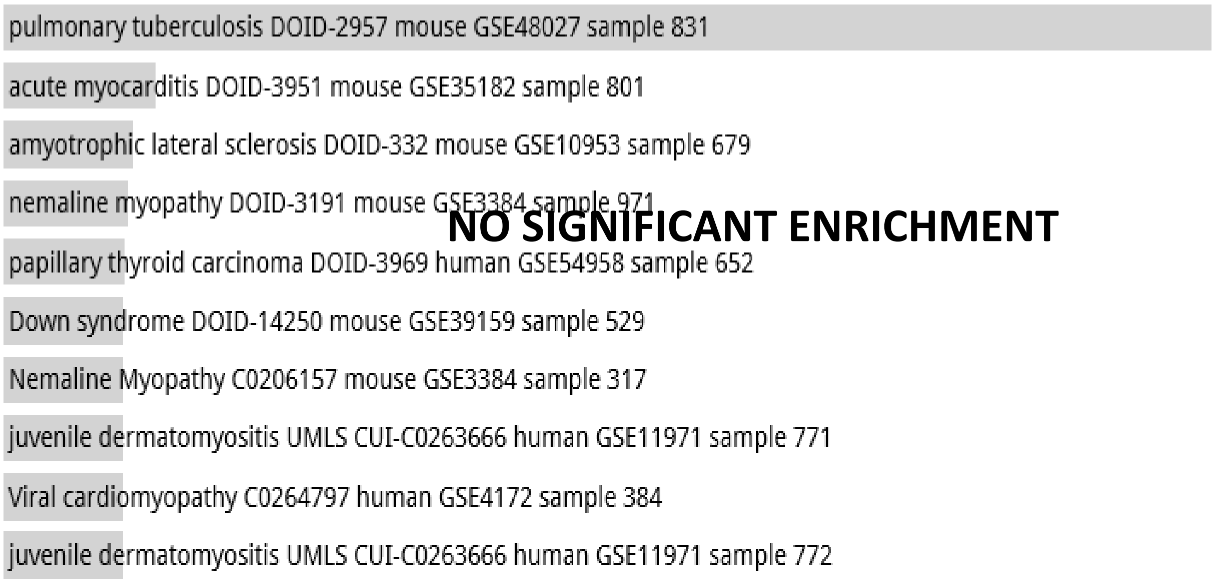

**Random Set 15**

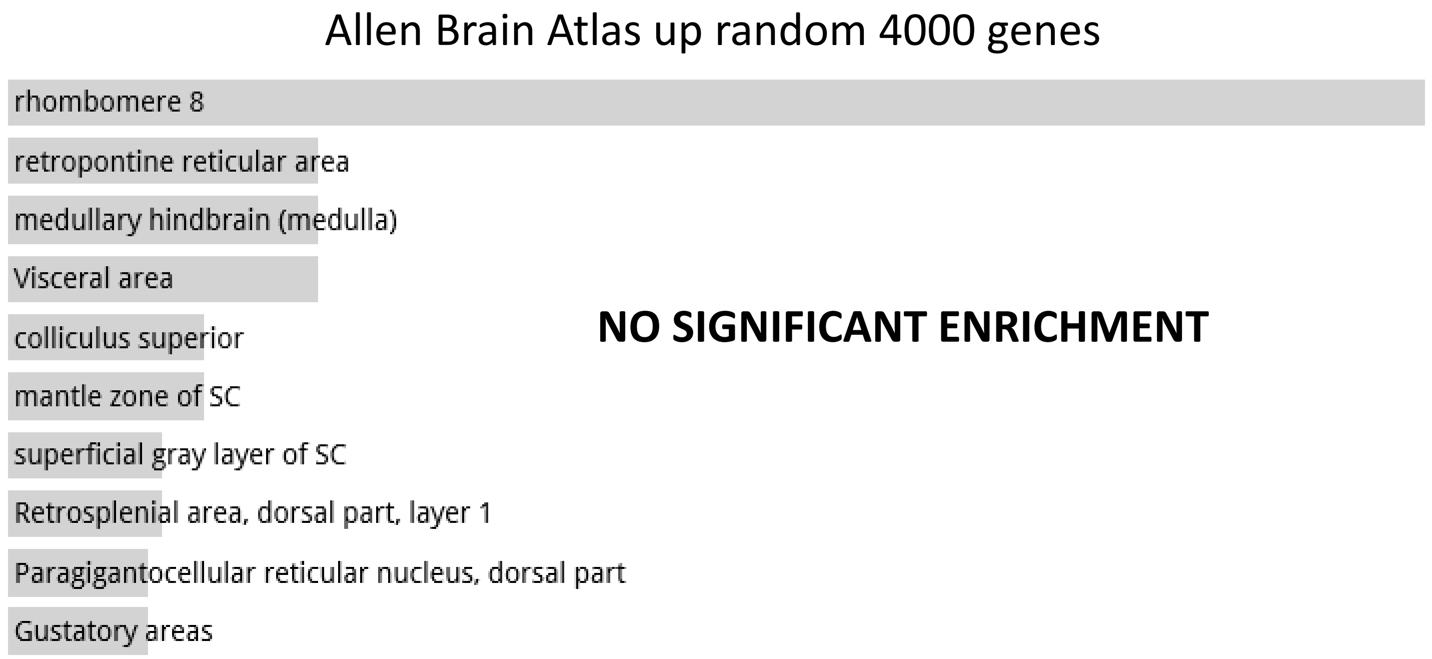


**NO SIGNIFICANT ENRICHMENT**

**Random Set 16**

ARCHS4 Tissues random 4000 genes

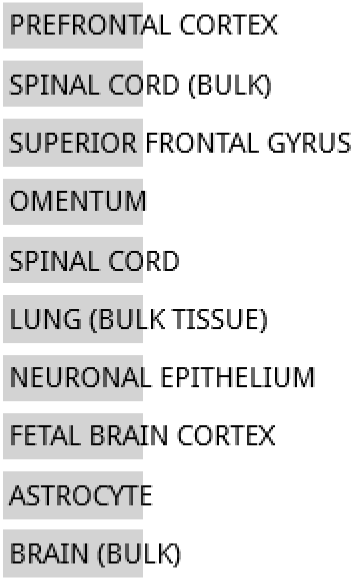


**NO SIGNIFICANT ENRICHMENT**

Jensen TISSUES random 4000 genes

**NO DATA AVAILABLE**

**Random Set 16**

#### Jensen DISEASES random 4000 genes

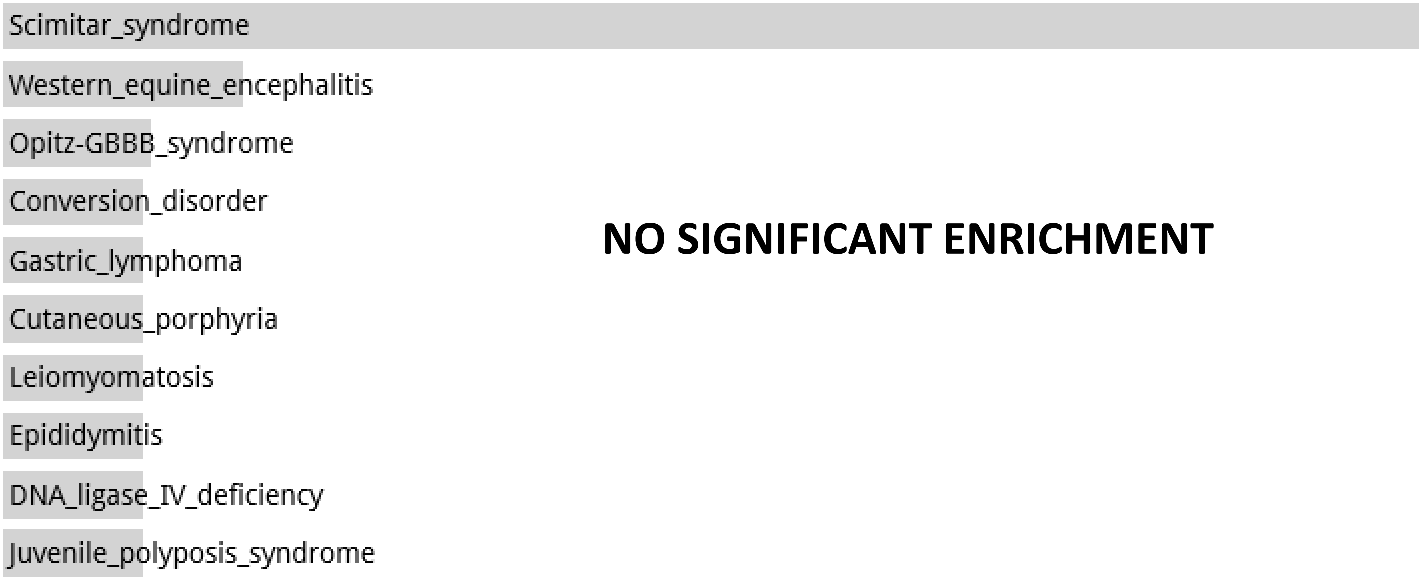


#### Disease Perturbations from GEO up random 4000 genes

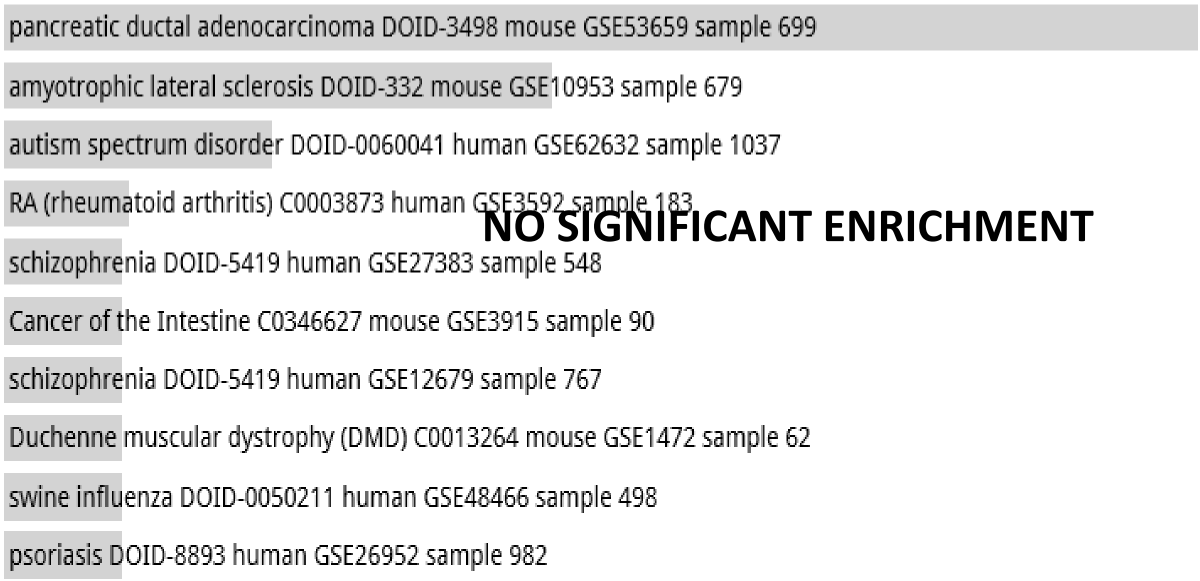


#### Disease Perturbations from GEO down random 4000 genes

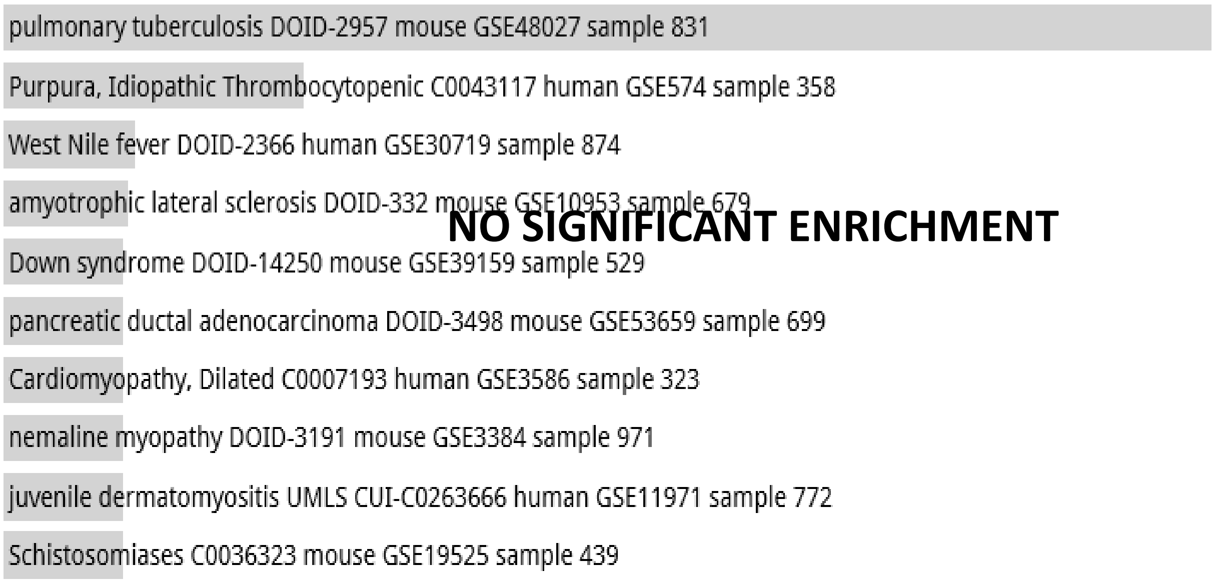

**Random Set 16**

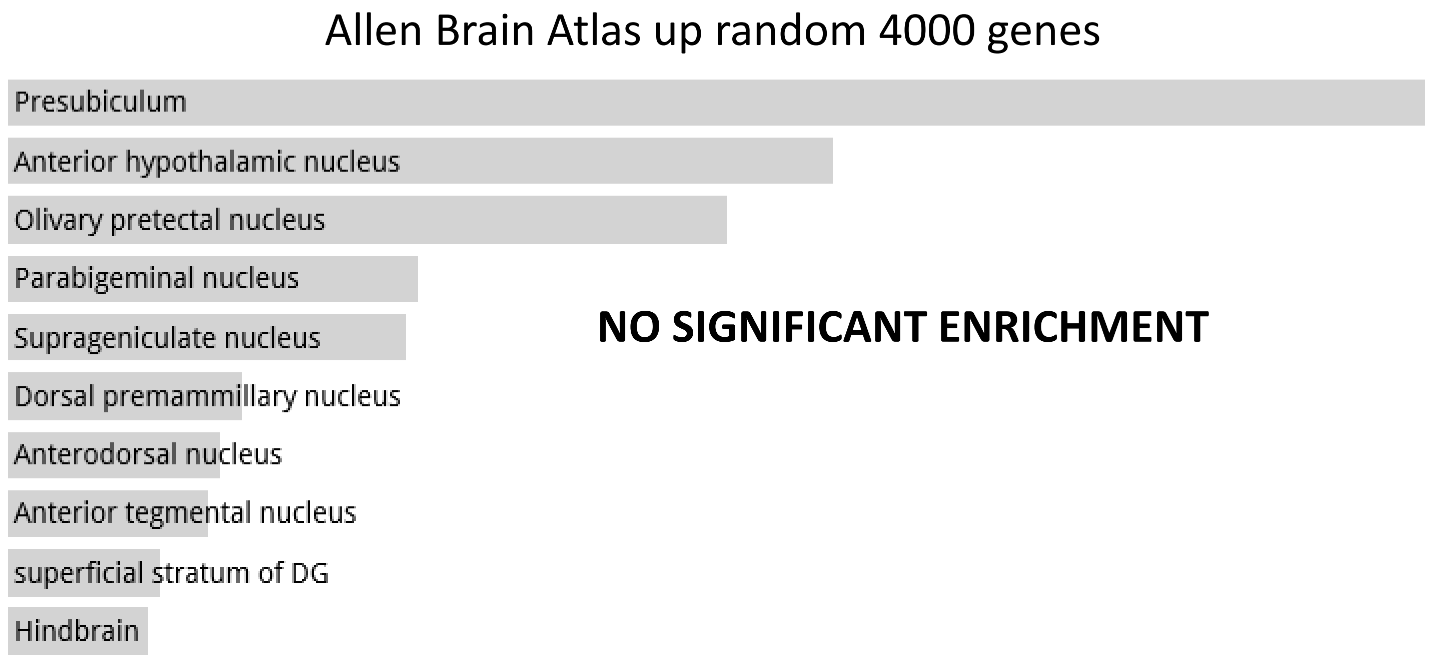

**Random Set 17**

ARCHS4 Tissues random 4000 genes

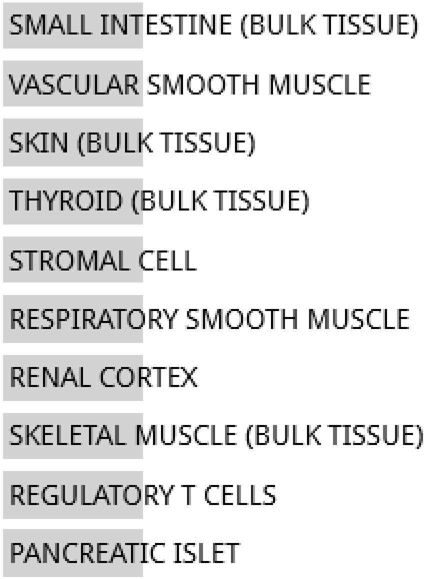


**NO SIGNIFICANT ENRICHMENT**

Jensen TISSUES random 4000 genes

**NO DATA AVAILABLE**

**Random Set 17**

Jensen DISEASES random 4000 genes

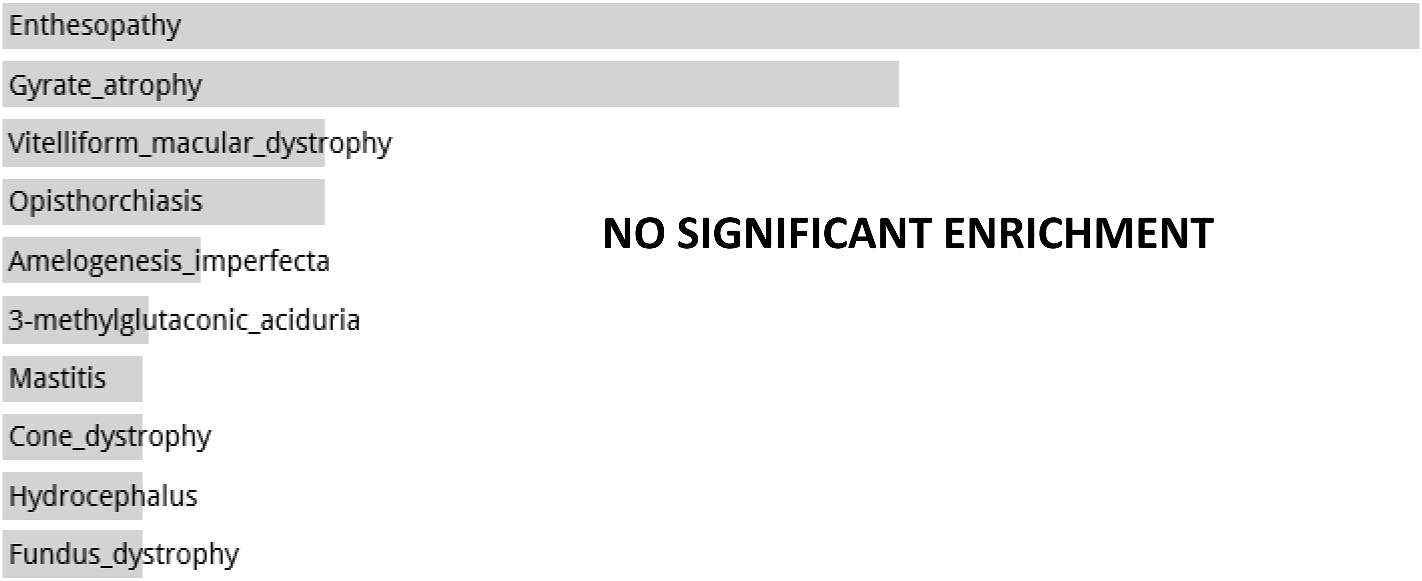


Disease Perturbations from GEO up random 4000 genes

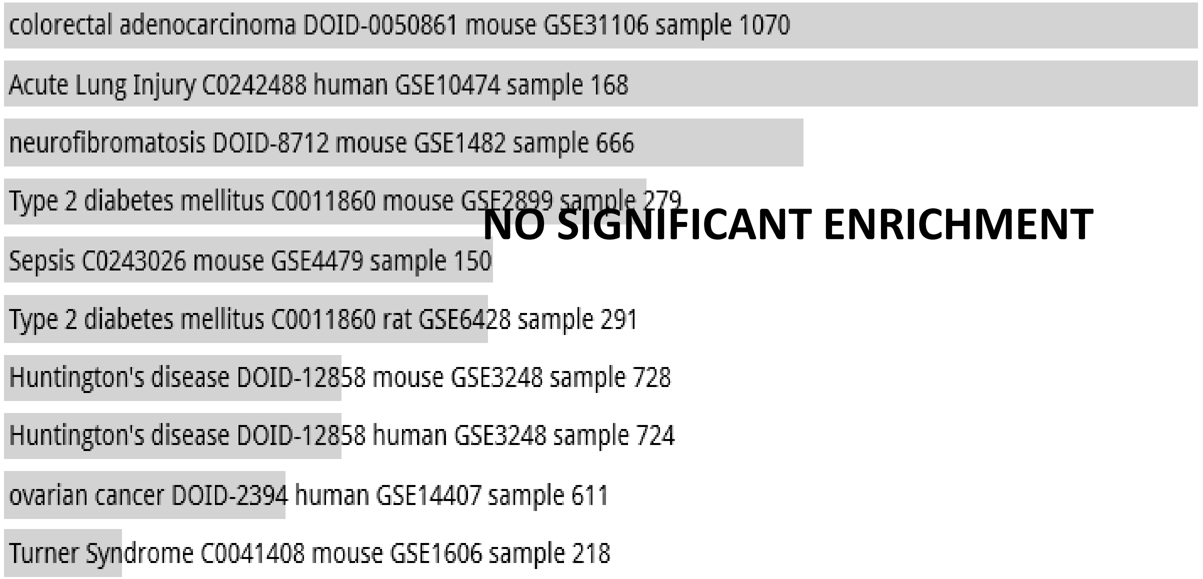


Disease Perturbations from GEO down random 4000 genes

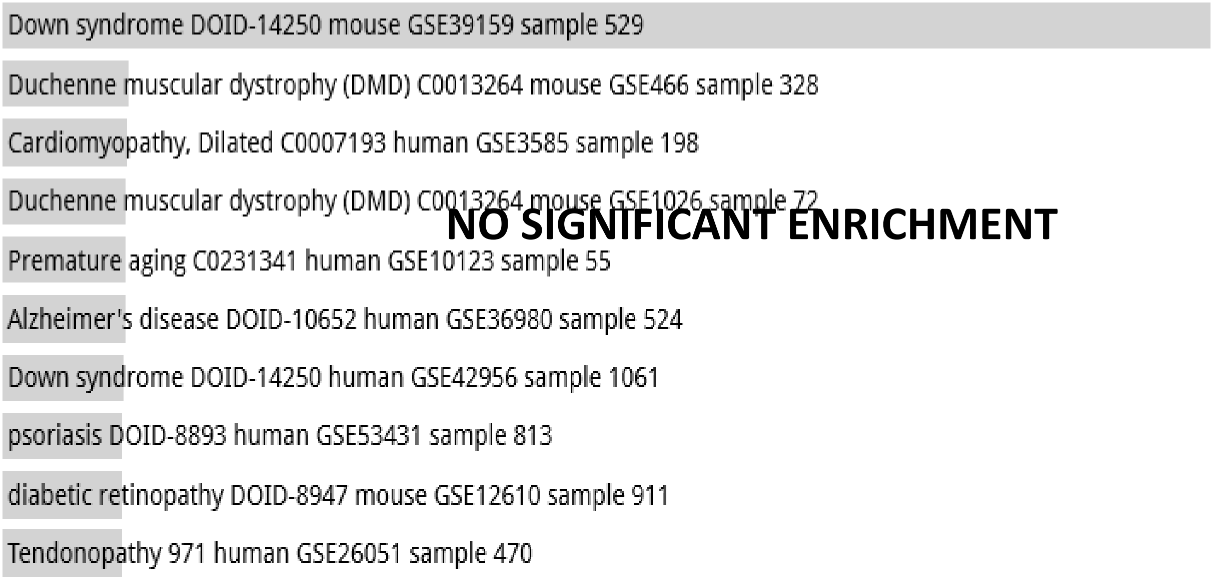

**Random Set 17**

### Allen Brain Atlas up random 4000 genes

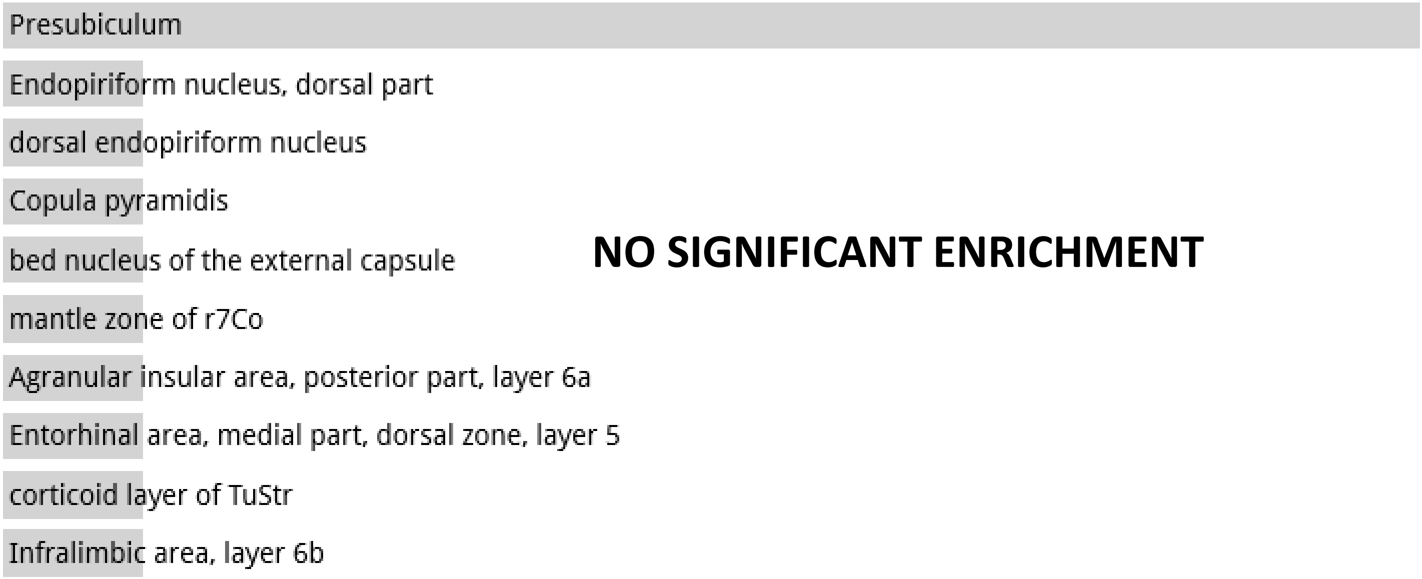


**NO SIGNIFICANT ENRICHMENT**

**Random Set 18**

ARCHS4 Tissues random 4000 genes

**NO DATA AVAILABLE**

Jensen TISSUES random 4000 genes

**NO DATA AVAILABLE**

**Random Set 18**

Jensen DISEASES random 4000 genes

**NO DATA AVAILABLE**

Disease Perturbations from GEO up random 4000 genes

**NO DATA AVAILABLE**

Disease Perturbations from GEO down random 4000 genes

**NO DATA AVAILABLE**

**Random Set 18**

Allen Brain Atlas up random 4000 genes

**NO DATA AVAILABLE**

**Random Set 19**

ARCHS4 Tissues random 4000 genes

**NO DATA AVIALABLE**

Jensen TISSUES random 4000 genes

**NO DATA AVAILABLE**

**Random Set 19**

Jensen DISEASES random 4000 genes

**NO DATA AVAILABLE**

Disease Perturbations from GEO up random 4000 genes

**NO DATA AVAILABLE**

Disease Perturbations from GEO down random 4000 genes

**NO DATA AVAILABLE**

**Random Set 19**

Allen Brain Atlas up random 4000 genes

**NO DATA AVAILABLE**

**Random Set 20**

ARCHS4 Tissues random 4000 genes

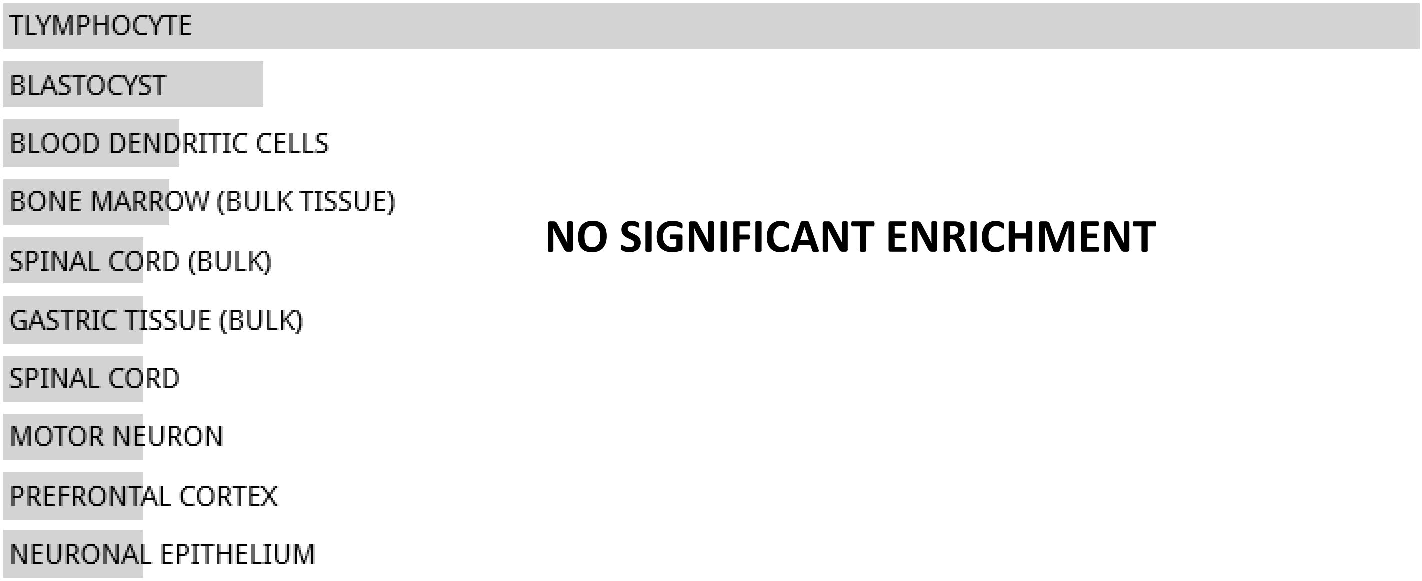


**NO SIGNIFICANT ENRICHMENT**

Jensen TISSUES random 4000 genes

**NO DATA AVAILABLE**

**Random Set 20**

Jensen DISEASES random 4000 genes

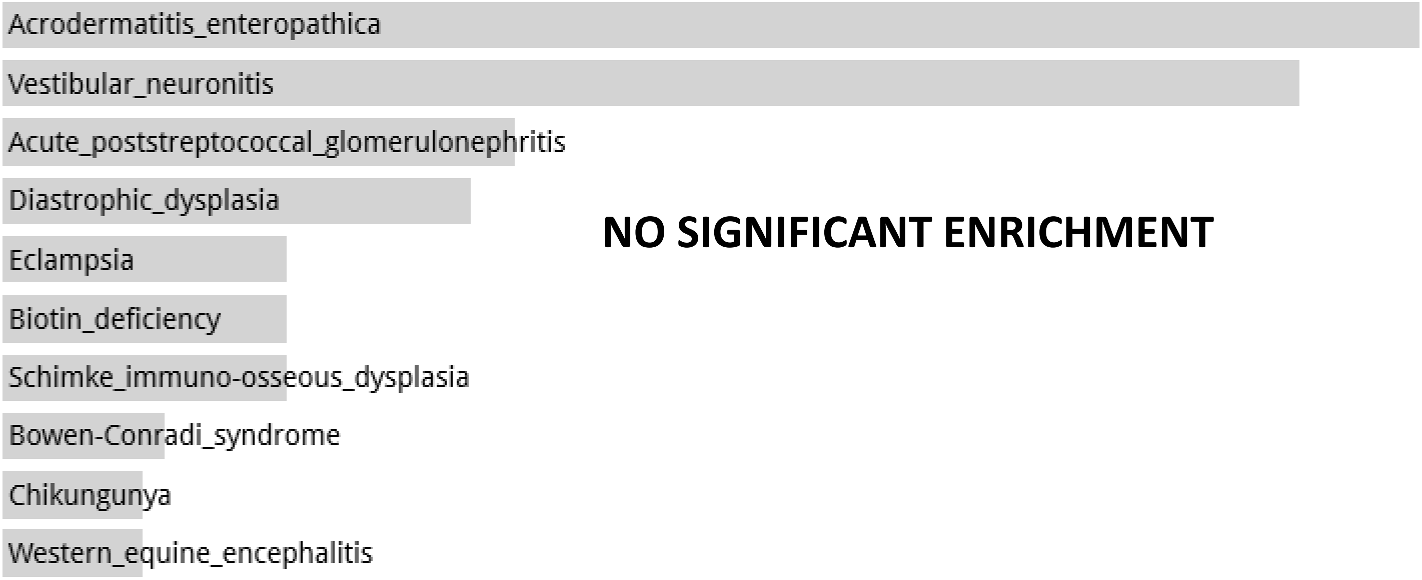


Disease Perturbations from GEO up random 4000 genes

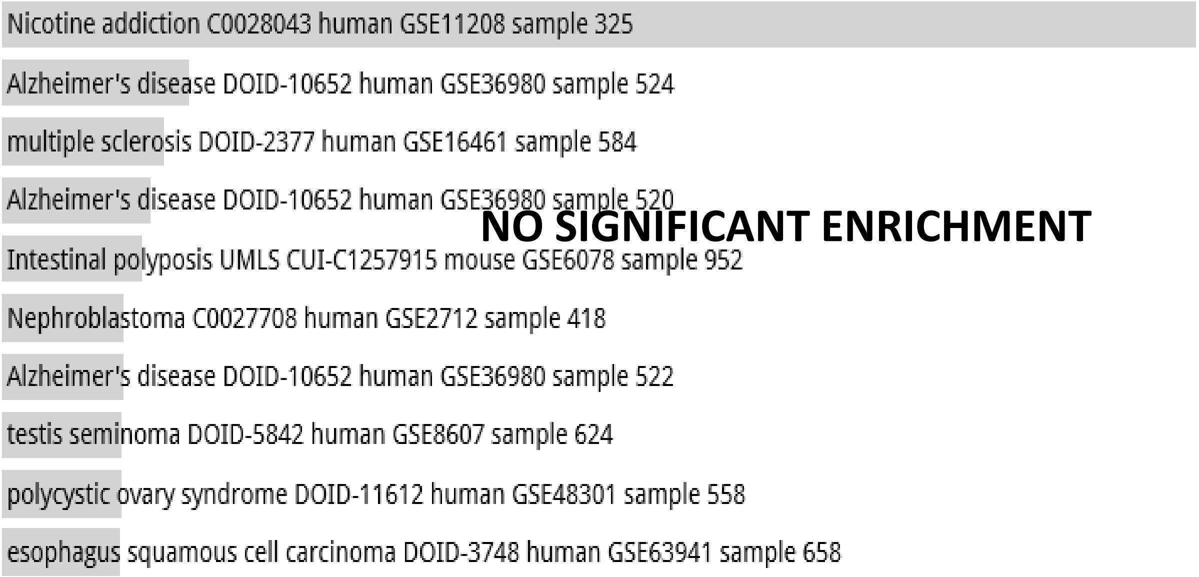


Disease Perturbations from GEO down random 4000 genes

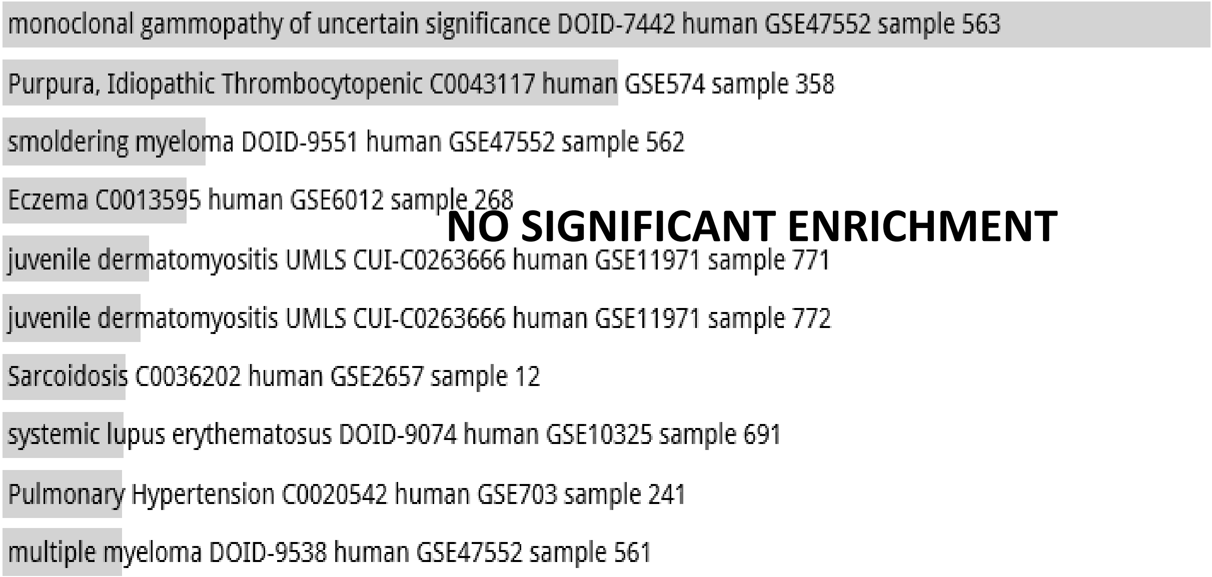

**Random Set 20**

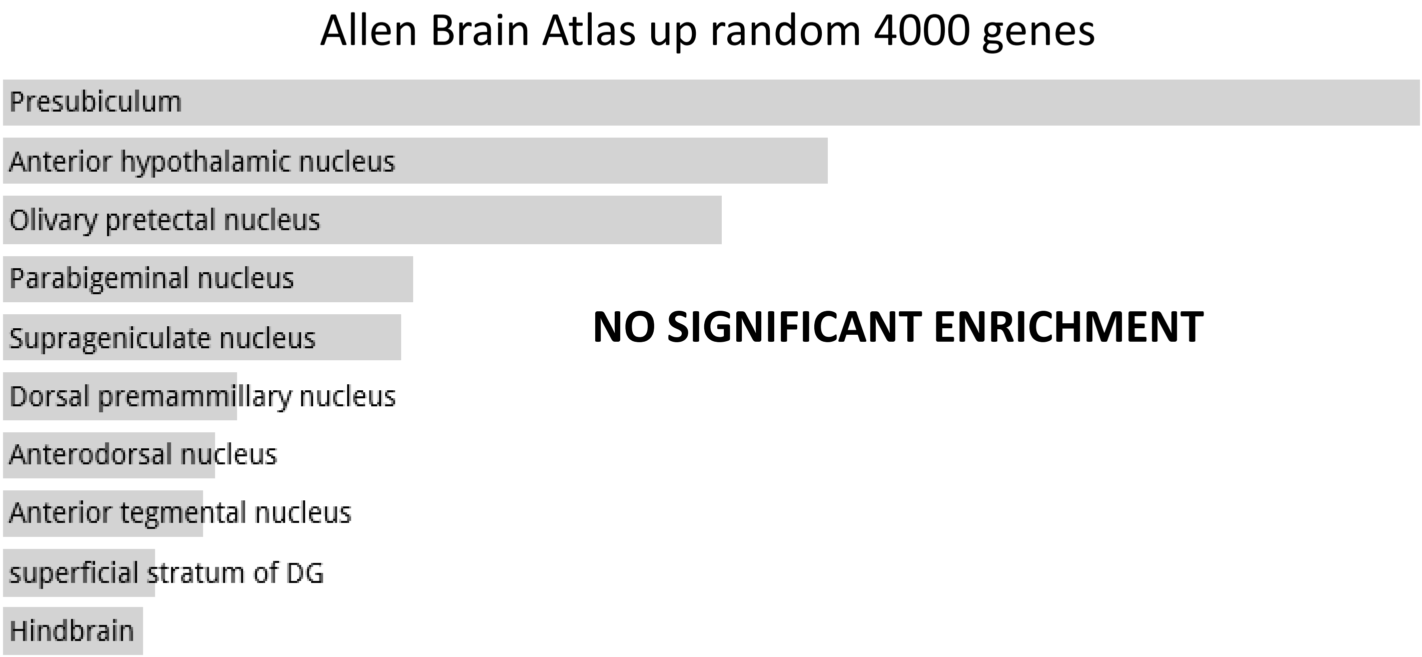

**Random Set 21**

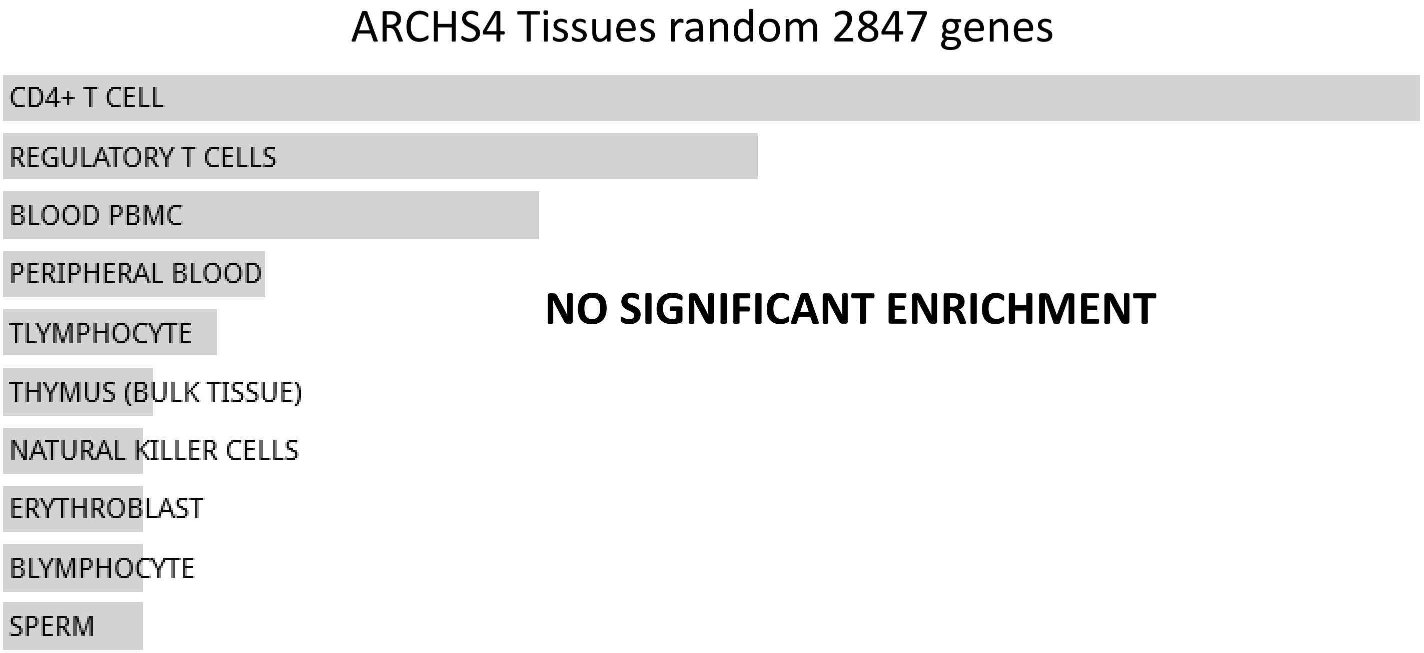


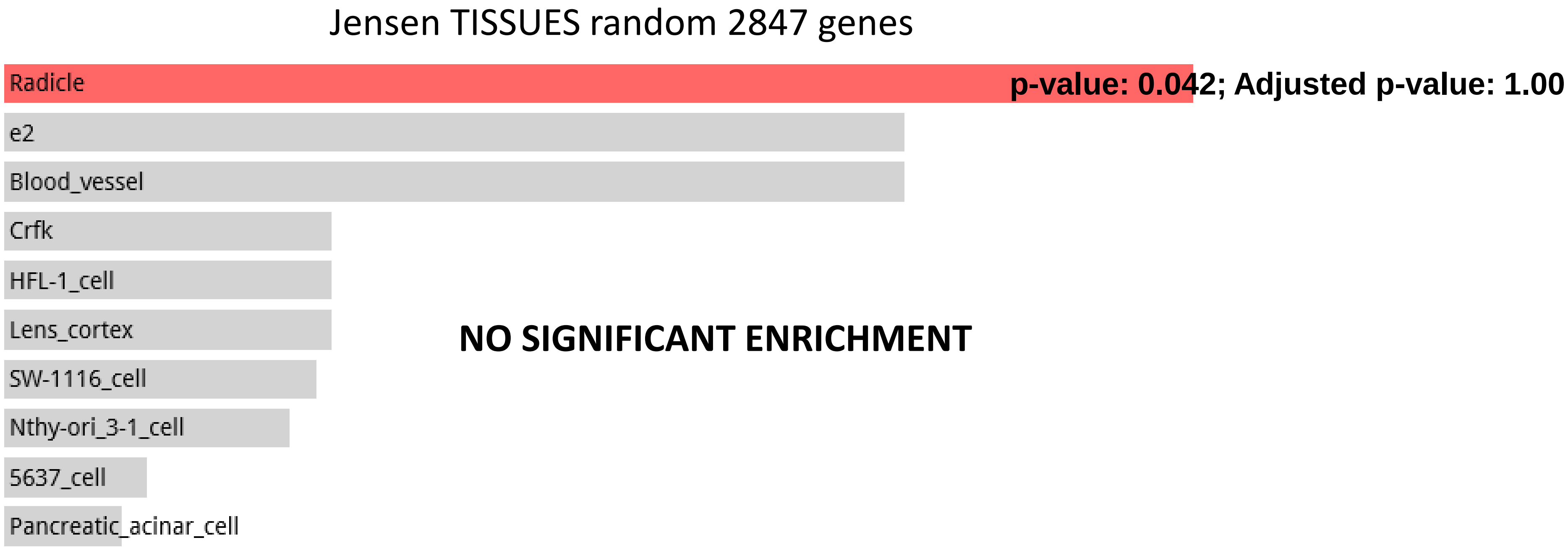

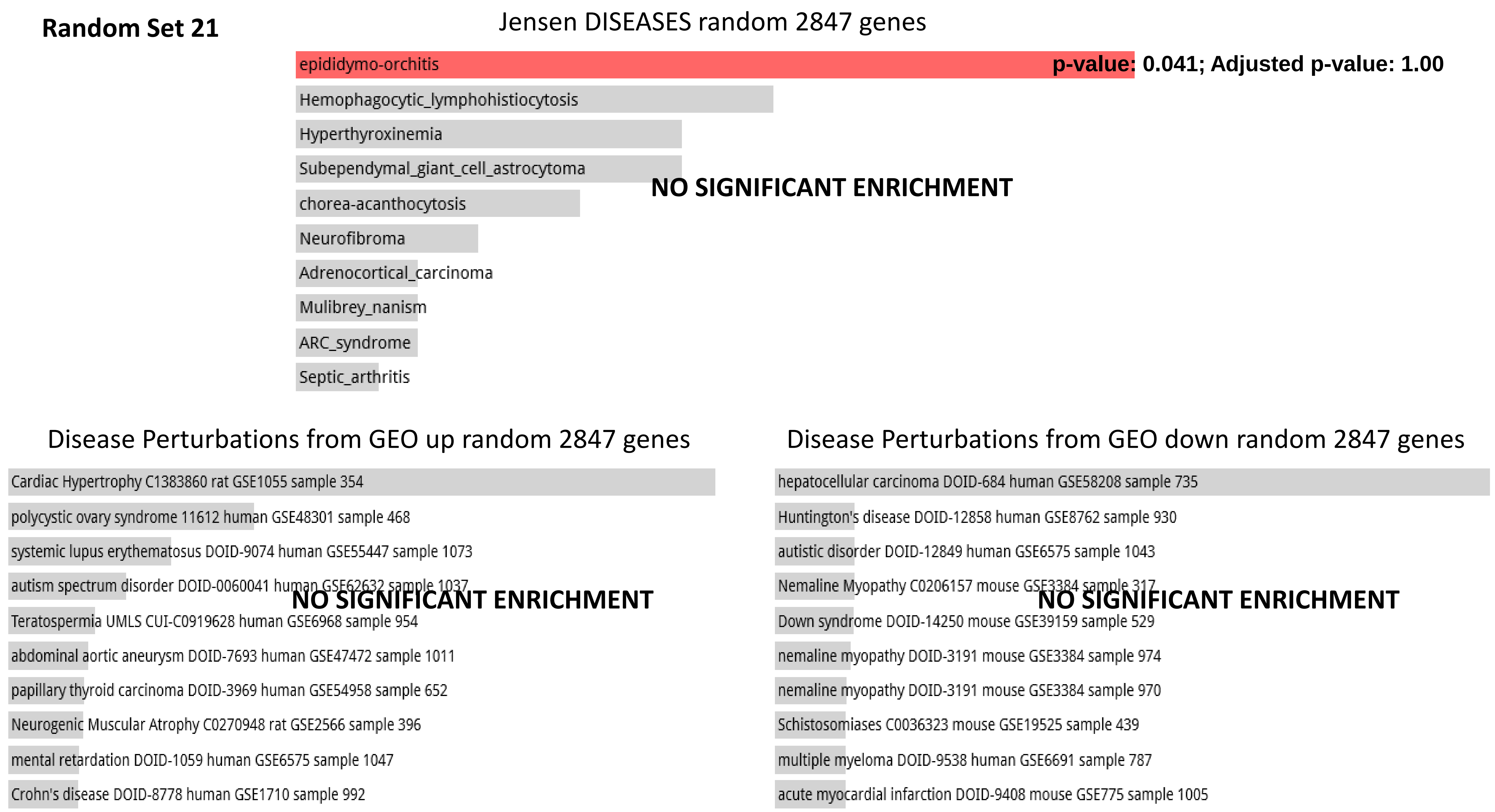
Random Set 21
Jensen DISEASES random 2847 genes
epididymo-orchitis
p-value: 0.041; Adjusted p-value: 1.00
Hemophagocytic_lymphohistiocytosis
Hyperthyroxinemia
Subependymal_giant_cell_astrocytoma
chorea-acanthocytosis
Neurofibroma
Adrenocortical_carcinoma
Mulibrey_nanism
ARC_syndrome
Septic_arthritis
NO SIGNIFICANT ENRICHMENT
Disease Perturbations from GEO up random 2847 genes
Cardiac Hypertrophy C1383860 rat GSE1055 sample 354
polycystic ovary syndrome 11612 human GSE48301 sample 468
systemic lupus erythematosus DOID-9074 human GSE55447 sample 1073
autism spectrum disorder DOID-0060041 human GSE62632 sample 1037
Teratospermia UMLS CUI-C0919628 human GSE6968 sample 954
abdominal aortic aneurysm DOID-7693 human GSE47472 sample 1011
papillary thyroid carcinoma DOID-3969 human GSE54958 sample 652
Neurogenic Muscular Atrophy C0270948 rat GSE2566 sample 396
mental retardation DOID-1059 human GSE6575 sample 1047
Crohn's disease DOID-8778 human GSE1710 sample 992
NO SIGNIFICANT ENRICHMENT
Disease Perturbations from GEO down random 2847 genes
hepatocellular carcinoma DOID-684 human GSE58208 sample 735
Huntington's disease DOID-12858 human GSE8762 sample 930
autistic disorder DOID-12849 human GSE6575 sample 1043
Nemaline Myopathy C0206157 mouse GSE3384 sample 317
Down syndrome DOID-14250 mouse GSE39159 sample 529
nemaline myopathy DOID-3191 mouse GSE3384 sample 974
nemaline myopathy DOID-3191 mouse GSE3384 sample 970
Schistosomiases C0036323 mouse GSE19525 sample 439
multiple myeloma DOID-9538 human GSE6691 sample 787
acute myocardial infarction DOID-9408 mouse GSE775 sample 1005
NO SIGNIFICANT ENRICHMENT

#### Random Set 21

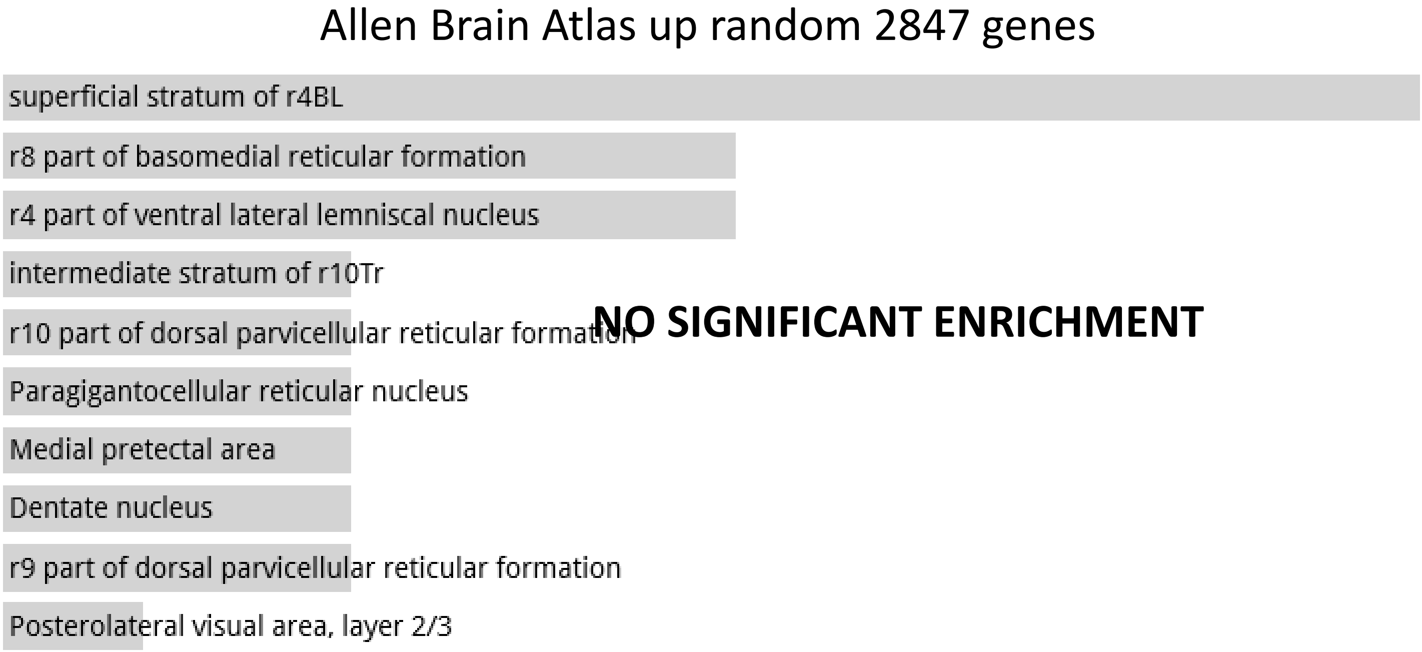


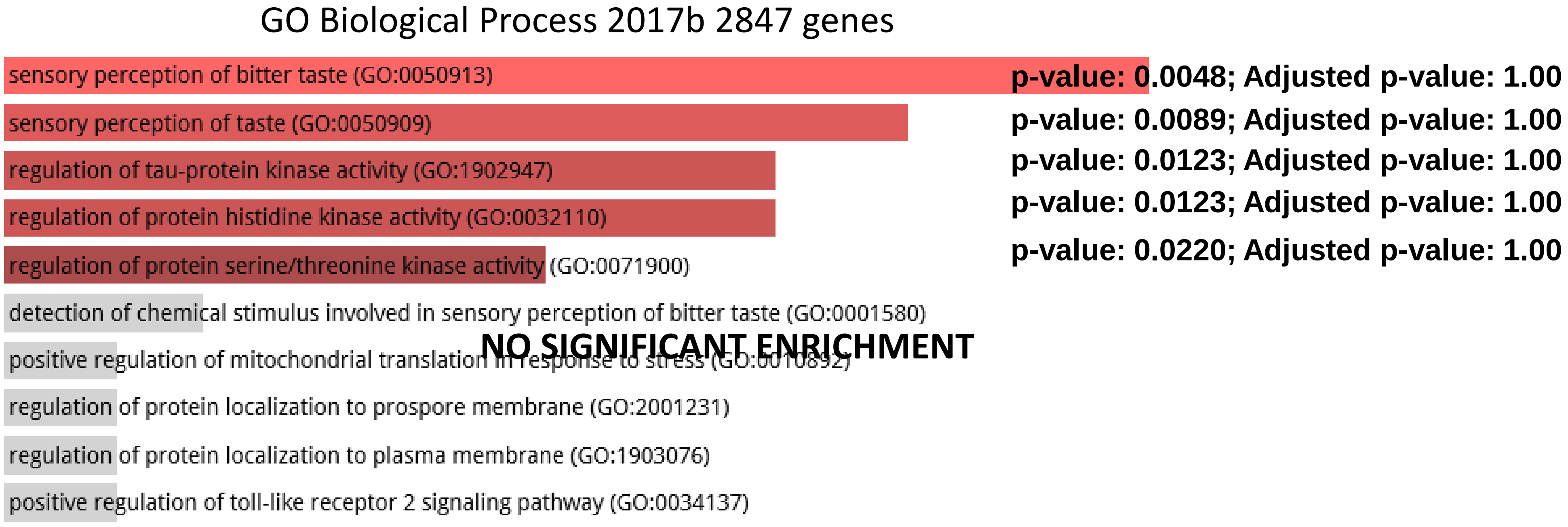

#### Random Set 21

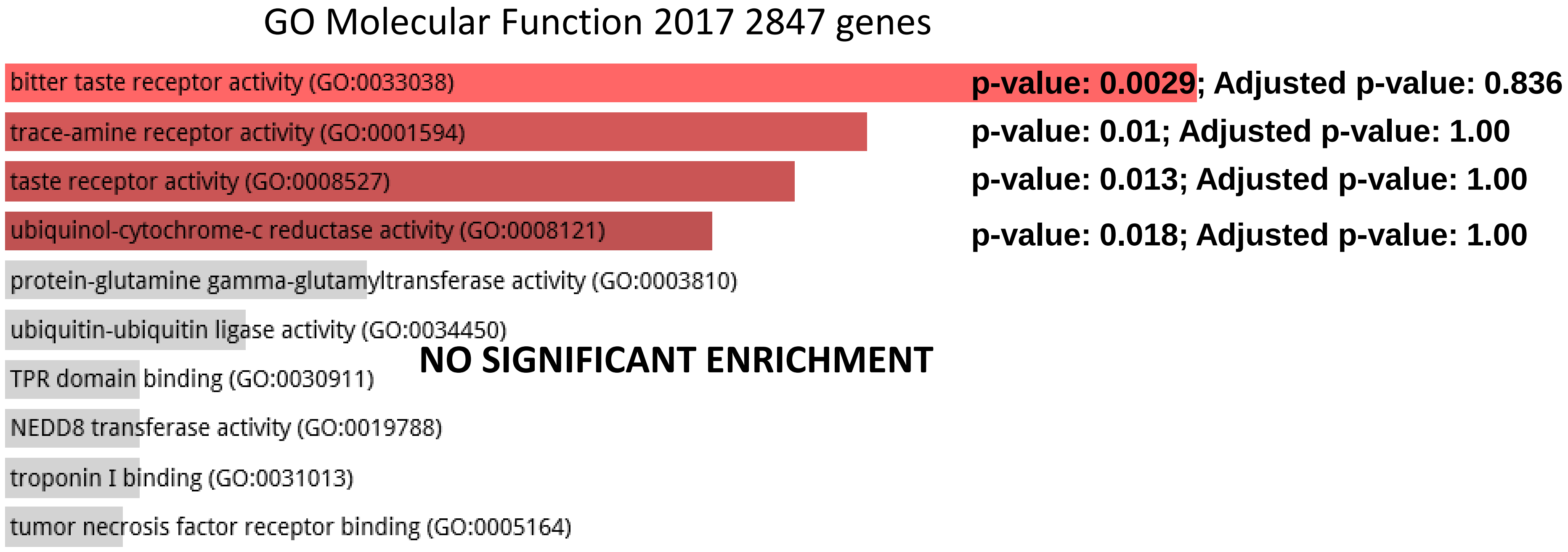


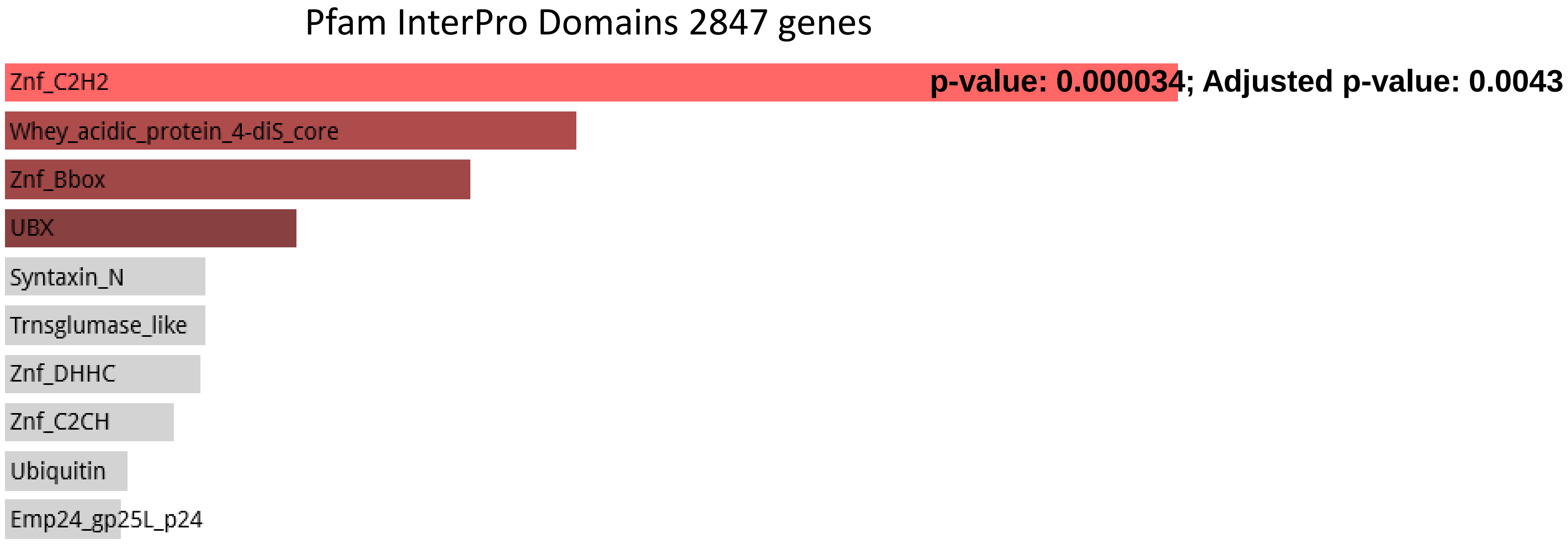

#### Random Set 21

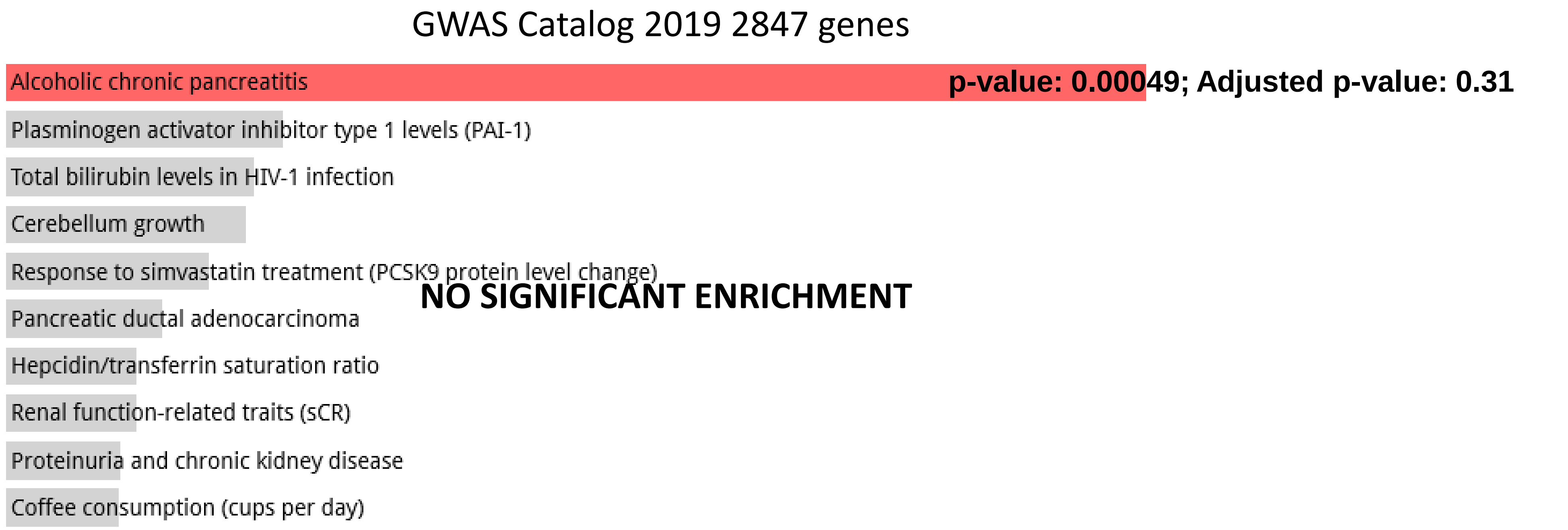


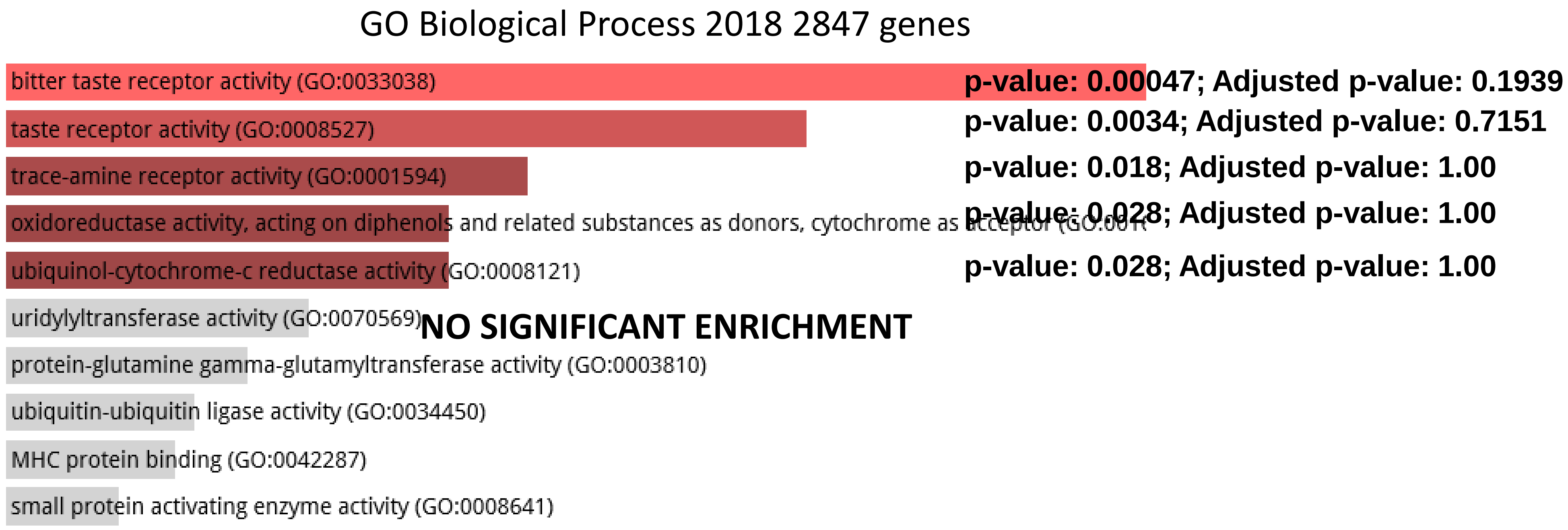

#### Random Set 21

CORUM 2847 genes

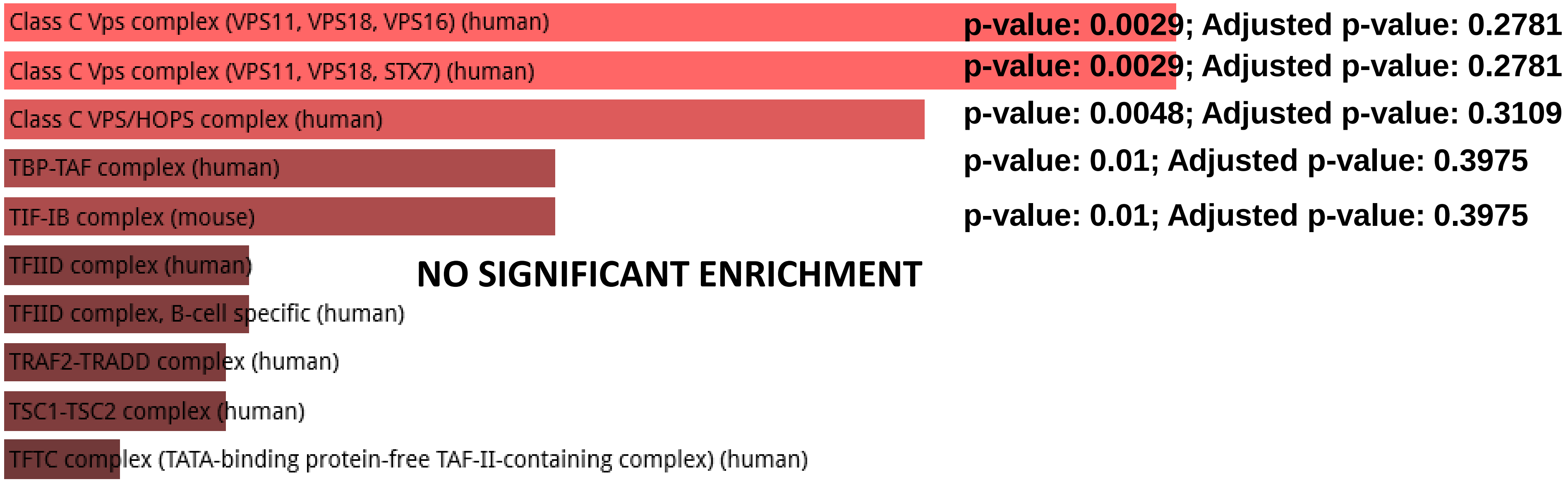


**CORUM database of protein complexes identified by mass spectrometry**

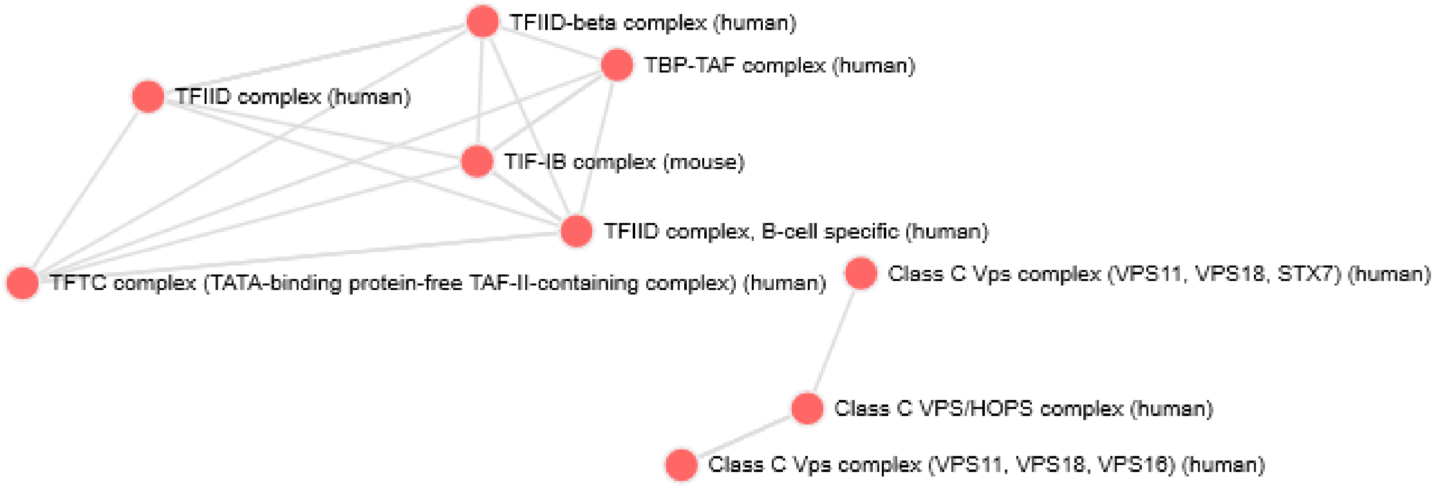
