## Supplemental Note for "Impacts of genomic networks governed by human-specific regulatory sequences and genetic loci harboring fixed human-specific neuro-regulatory single nucleotide mutations on phenotypic traits of Modern Humans"

The principal element of our analytical approach is to define specific sets of genomic regulatory loci
WITHOUT ANY PRIOR KNOWLEDGE
of what genes (if any) they may (or may not) control.

This is in striking contrast with the approaches that being utilized in the prior art: to identify genes that are differentially regulated between different states and/or conditions, thus introducing the confounders and biases associated with the multiple hypothesis testing, overfitting, and random co-occurrence due to the very large excess of analyzed features (genes) over the relatively small numbers of samples (100K range or more of analyzed features versus just a few hundred analyzed samples at best). Attempts to address these issues using statistical approaches (corrections for multiple hypothesis testing) and experimental design improvements (multiple replications, validation in independent data sets and using independent analytical techniques) did not adequately resolve these problems.

Importantly, our analyses revealed that this approach identifies not only genes altered expression of which implicated in the broad spectrum of human physiological and pathological conditions. It also identifies genes defined by the independent studies as genetic loci comprising mutation signatures associated with development and progression of multiple common human disorders, including cancer, neurodevelopmental, neuropsychiatric, and neurodegenerative disorders, as well autoimmune and immuno-inflammatory diseases.

Supporting the concept that many major human disorders are driven by aberrant functions of primate-specific genomic regulatory networks with prominent unique-to-human (human-specific) components, this approach identifies genetic loci implicated in inter-individual genetic mosaicism (somatic mosaicism) of cells, tissues and organs in the human body.

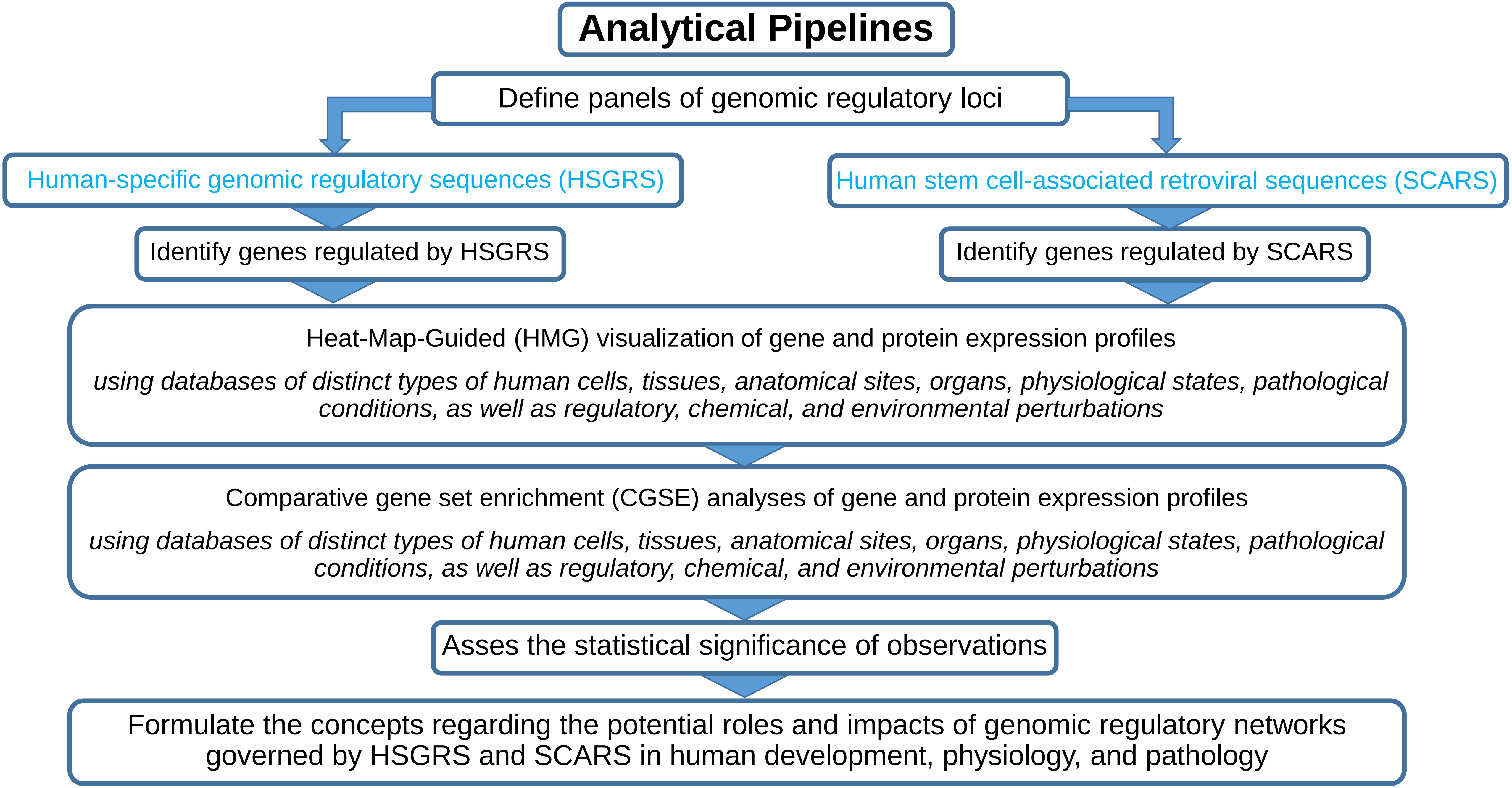
Analytical Pipelines
Define panels of genomic regulatory loci
Human-specific genomic regulatory sequences (HSGRS)
Human stem cell-associated retroviral sequences (SCARS)
Identify genes regulated by HSGRS
Identify genes regulated by SCARS
Heat-Map-Guided (HMG) visualization of gene and protein expression profiles
using databases of distinct types of human cells, tissues, anatomical sites, organs, physiological states, pathological conditions, as well as regulatory, chemical, and environmental perturbations
Comparative gene set enrichment (CGSE) analyses of gene and protein expression profiles
using databases of distinct types of human cells, tissues, anatomical sites, organs, physiological states, pathological conditions, as well as regulatory, chemical, and environmental perturbations
Asses the statistical significance of observations
Formulate the concepts regarding the potential roles and impacts of genomic regulatory networks governed by HSGRS and SCARS in human development, physiology, and pathology

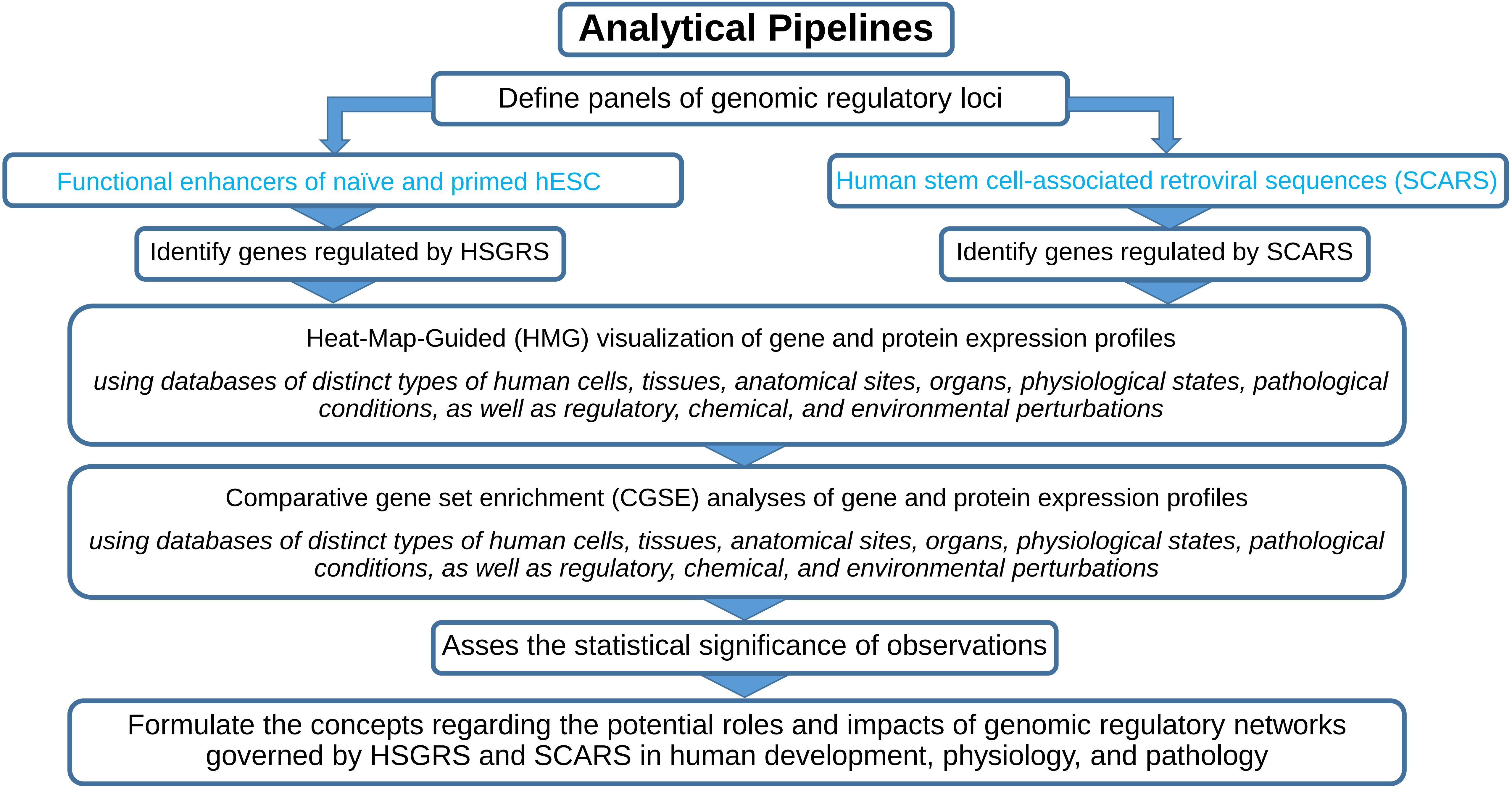
Analytical Pipelines
Define panels of genomic regulatory loci
Functional enhancers of naïve and primed hESC
Human stem cell-associated retroviral sequences (SCARS)
Identify genes regulated by HSGRS
Identify genes regulated by SCARS
Heat-Map-Guided (HMG) visualization of gene and protein expression profiles
using databases of distinct types of human cells, tissues, anatomical sites, organs, physiological states, pathological conditions, as well as regulatory, chemical, and environmental perturbations
Comparative gene set enrichment (CGSE) analyses of gene and protein expression profiles
using databases of distinct types of human cells, tissues, anatomical sites, organs, physiological states, pathological conditions, as well as regulatory, chemical, and environmental perturbations
Asses the statistical significance of observations
Formulate the concepts regarding the potential roles and impacts of genomic regulatory networks governed by HSGRS and SCARS in human development, physiology, and pathology

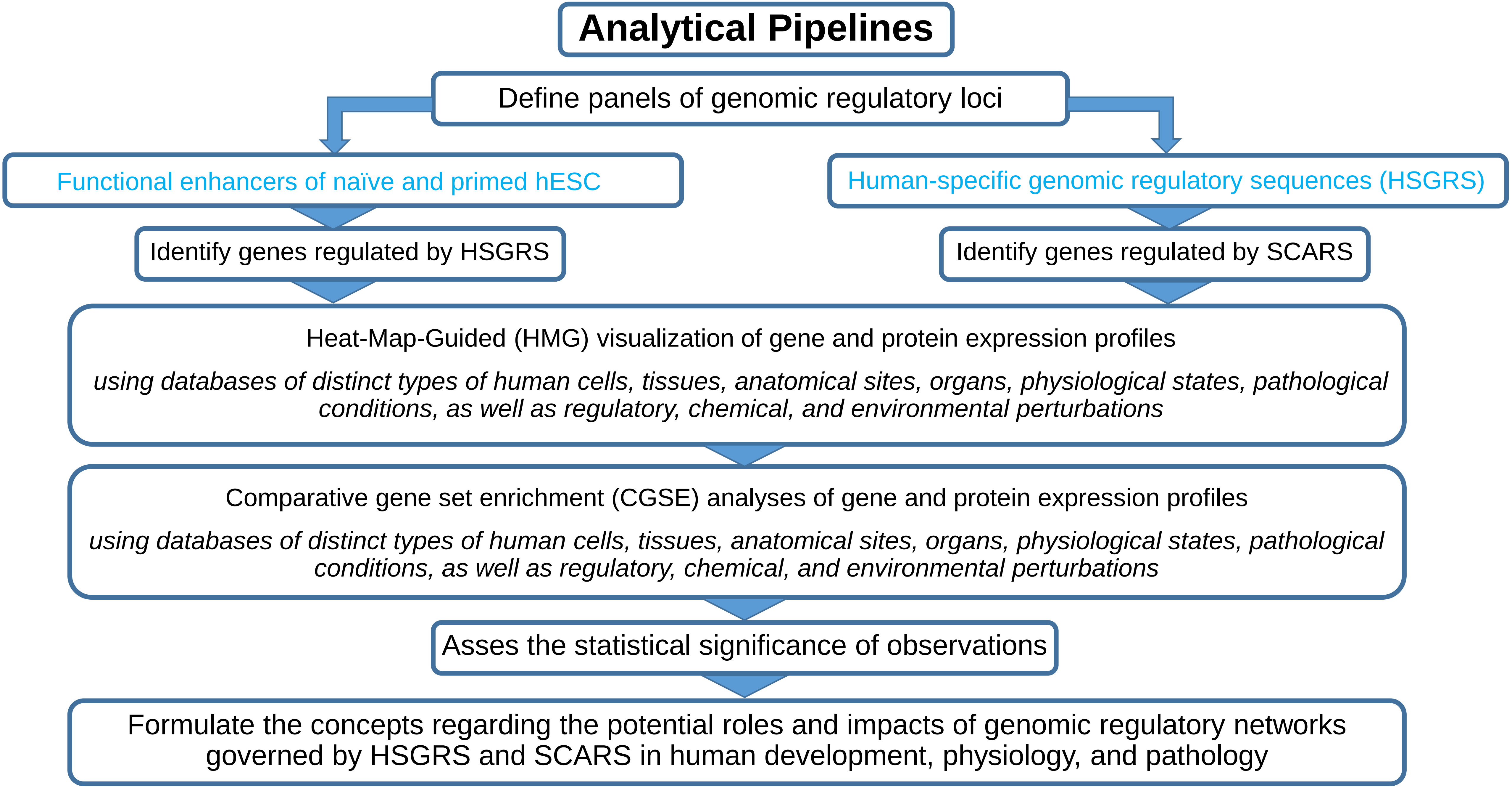
Analytical Pipelines
Define panels of genomic regulatory loci
Functional enhancers of naïve and primed hESC
Human-specific genomic regulatory sequences (HSGRS)
Identify genes regulated by HSGRS
Identify genes regulated by SCARS
Heat-Map-Guided (HMG) visualization of gene and protein expression profiles
using databases of distinct types of human cells, tissues, anatomical sites, organs, physiological states, pathological conditions, as well as regulatory, chemical, and environmental perturbations
Comparative gene set enrichment (CGSE) analyses of gene and protein expression profiles
using databases of distinct types of human cells, tissues, anatomical sites, organs, physiological states, pathological conditions, as well as regulatory, chemical, and environmental perturbations
Asses the statistical significance of observations
Formulate the concepts regarding the potential roles and impacts of genomic regulatory networks governed by HSGRS and SCARS in human development, physiology, and pathology

### ~59K human-specific genomic regulatory sequences

13,824 genes are putative regulatory targets of HSGRS

**Structurally, functionally, and evolutionary distinct families of human-specific regulatory sequences (HSRS) and associated putative regulatory target genes defined by the GREAT algorithm.**

| Classification category/Reference database | Number of records (hg19) | Associated genes |
|---|---|---|
| Fixed human-specific insertions. | 11878 | 7979 |
| Human-specific TE loci expressed in human dorsolateral prefrontal cortex | 4637 | 4051 |
| Set of duplicated regions in GRCh38 space | 7599 | 6618 |
| Fixed human-specific deletions | 5883 | 5489 |
| Human-specific STR expansions | 4875 | 4844 |
| hsTFBS | 3803 | 1087 |
| ace-DHS | 3538 | 3445 |
| FHSRR | 4249 | 2810 |
| Human-specific STR contractions | 1279 | 973 |
| hESC_FHSRR_DHS | 1932 | 1458 |
| DHS_FHSRR (non-hESC) | 2118 | 552 |
| HARs | 2745 | 2281 |
| haDHS | 524 | 747 |
| Human-biased CNCC enhances | 1000 | 1439 |
| Chimp-biased CNCC enhances | 1000 | 1445 |
| H3K4me3 peaks with human-specific enrichment in prefrontal neurons | 410 | 578 |
| Human-specific hESC functional enhancers | 1619 | 1214 |
| All HSRS | 59089 | 13824 |

Legend: Definitions of structurally, functionally, and evolutionary distinct families of human-specific regulatory sequences (HSRS) can be found in Glinsky (2020);

#### Association of human cancer survival predictor genes with human-specific genomic regulatory sequences (HSGRS)

| Cancer type | All genes | Associated with HSGRS | Percent |
|---|---|---|---|
| Thyroid | 347 | 269 | 77.52 |
| Glioma | 271 | 206 | 76.01 |
| Melanoma | 205 | 153 | 74.63 |
| Head and neck | 808 | 597 | 73.89 |
| Colorectal | 603 | 440 | 72.97 |
| Renal | 6070 | 4418 | 72.78 |
| Ovarian | 504 | 366 | 72.62 |
| Liver | 2892 | 2086 | 72.13 |
| Lung | 662 | 477 | 72.05 |
| Breast | 582 | 414 | 71.13 |
| Urothelial | 1101 | 783 | 71.12 |
| Stomach | 307 | 218 | 71.01 |
| Prostate | 161 | 114 | 70.81 |
| Endometrial | 1631 | 1153 | 70.69 |
| Cervical | 717 | 505 | 70.43 |
| Pancreatic | 1549 | 1075 | 69.40 |
| All human cancer survival genes | 10713 | 7738 | 72.23 |

**11,878 fixed human-specific insertions**

ARCHS4 Tissues 7979 genes

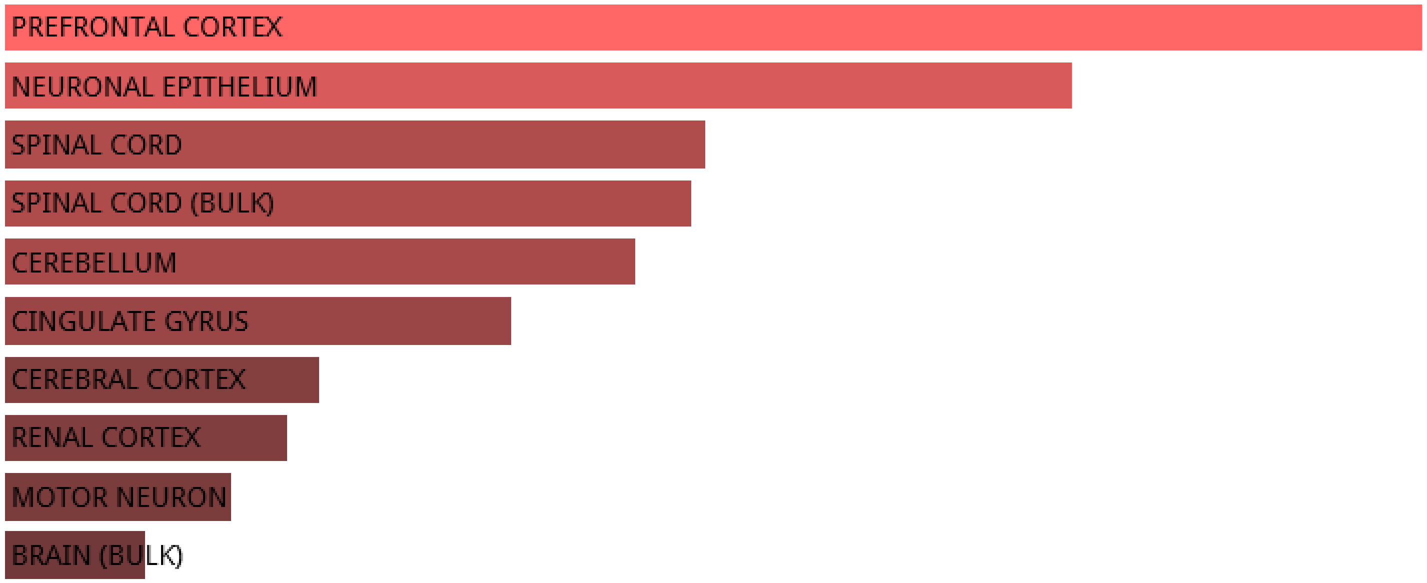


Jensen TISSUES 7979 genes

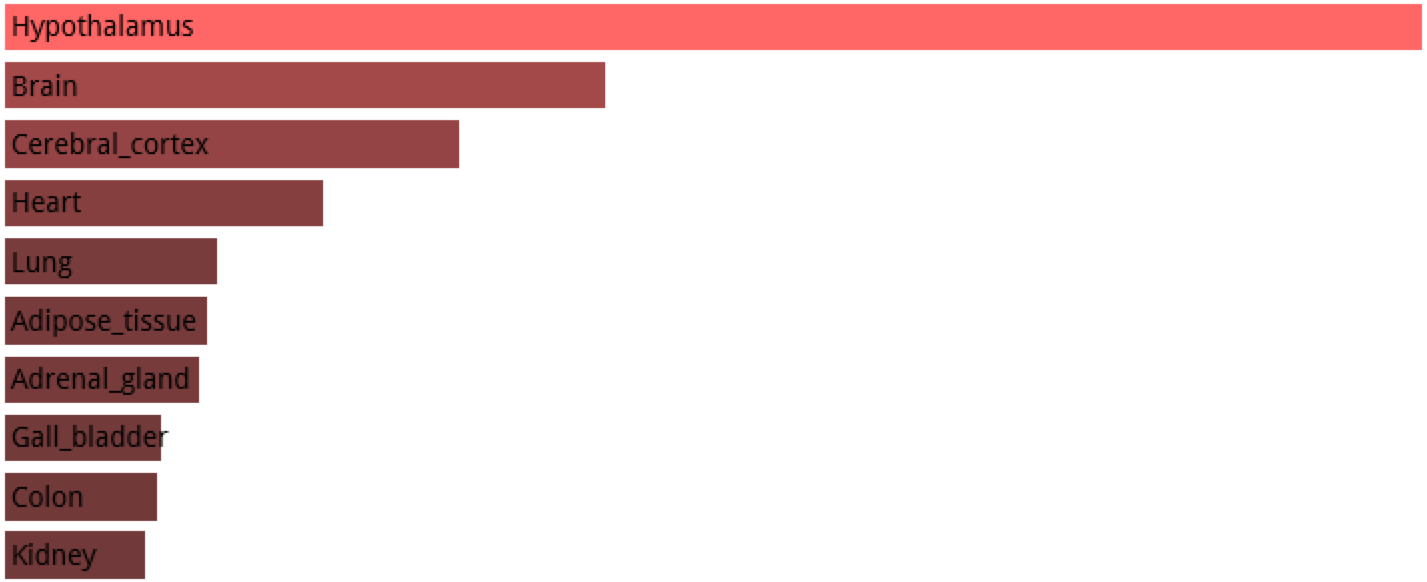

**11,878 fixed human-specific insertions**

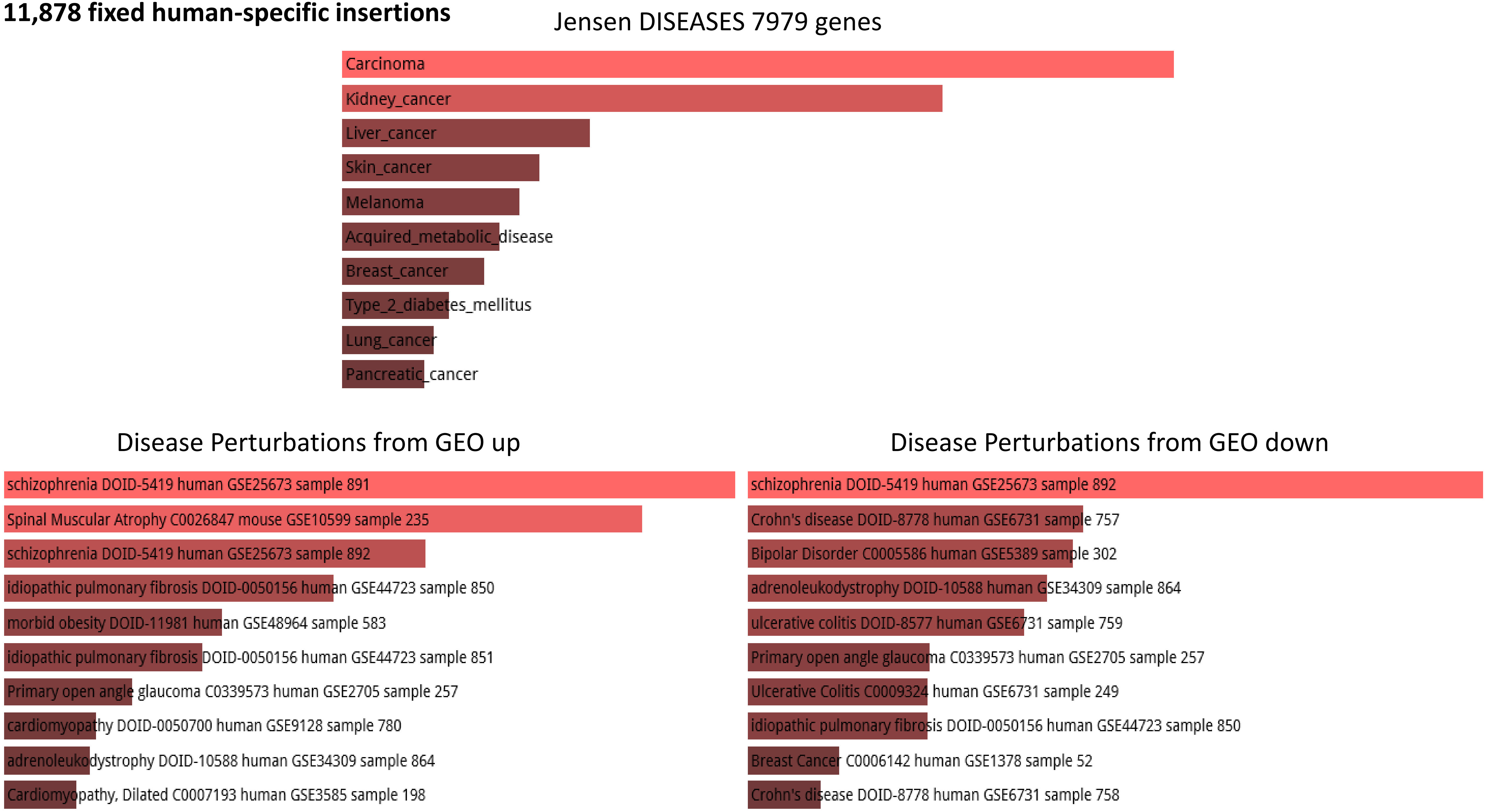

**7,599 duplicated regions in the human genome**

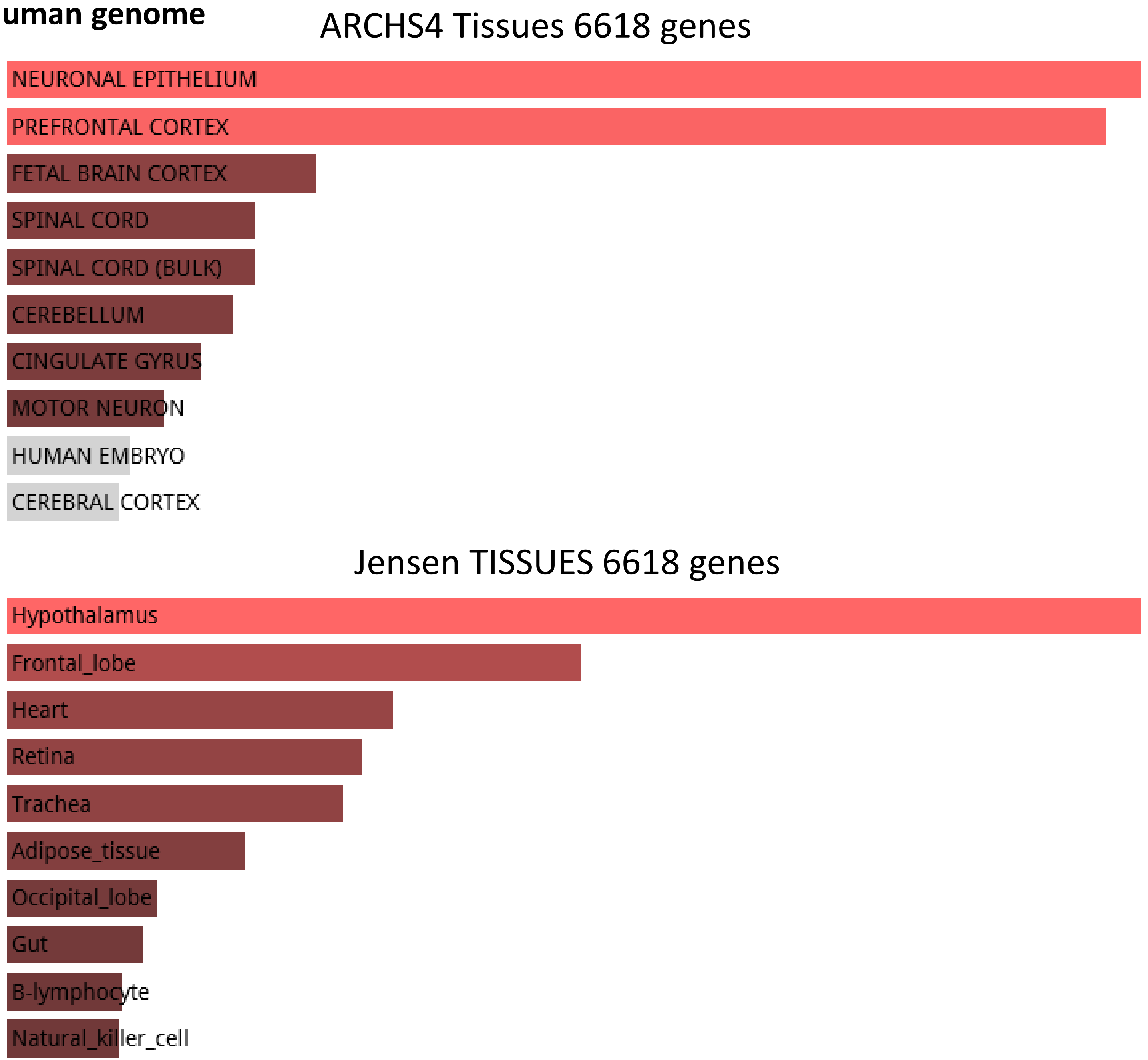

**7,599 duplicated regions in the human genome**

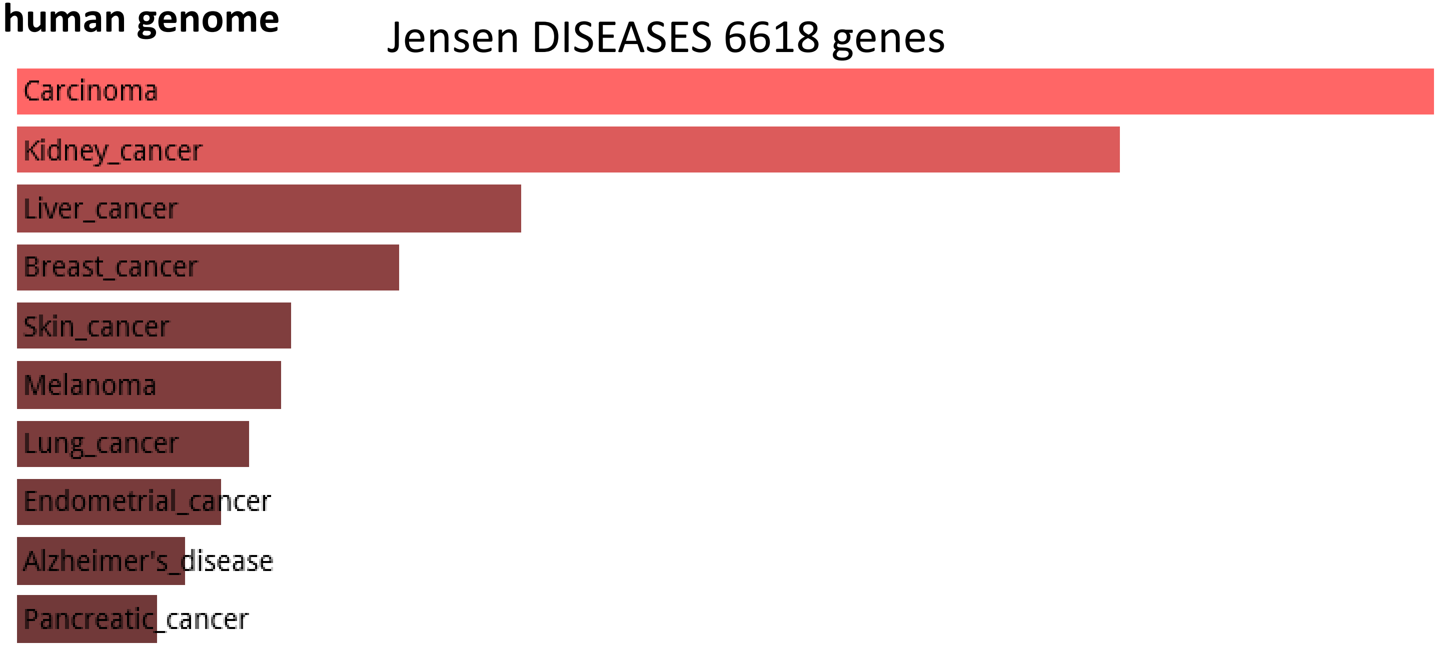


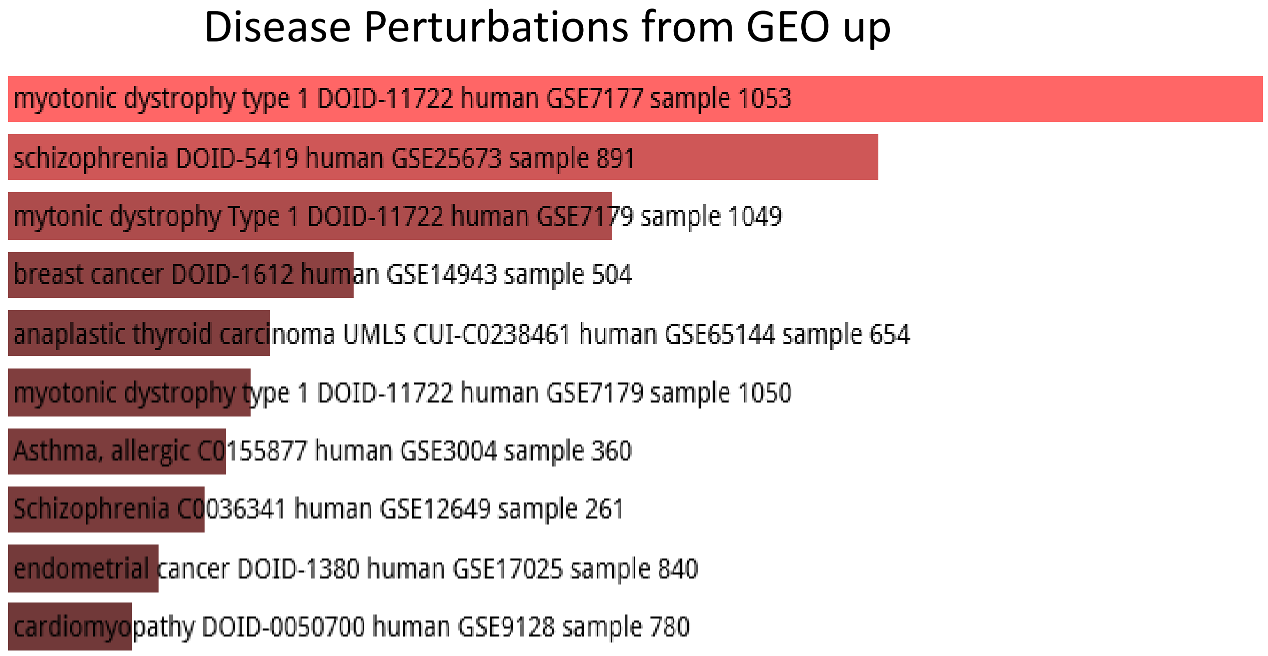


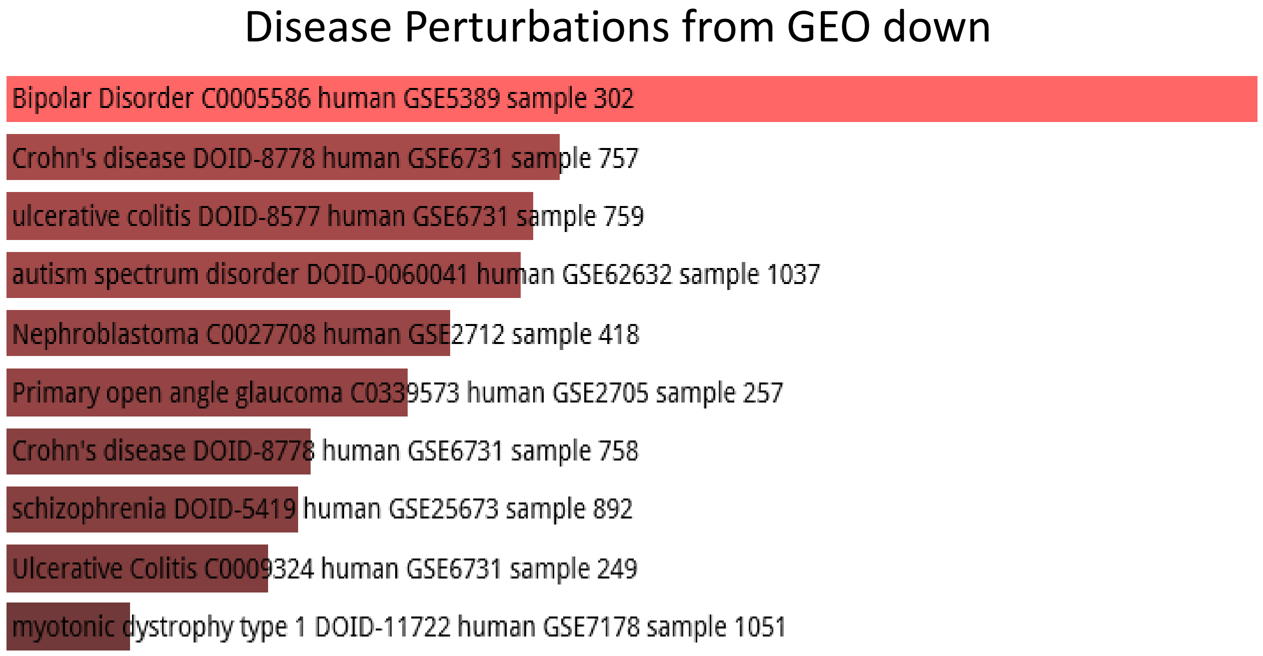

**5,883 fixed human-specific deletions**

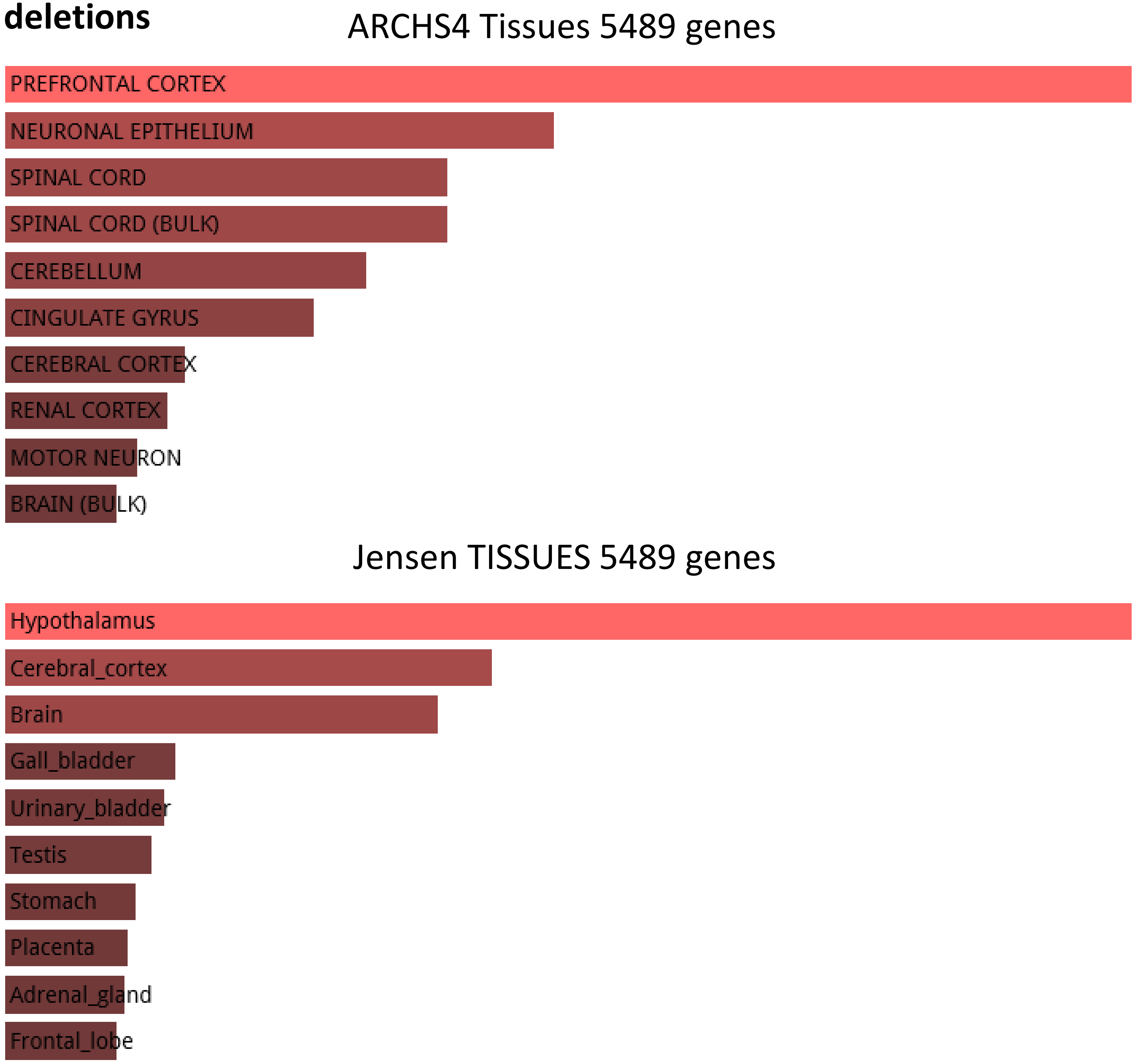

#### 5,883 fixed human-specific deletions

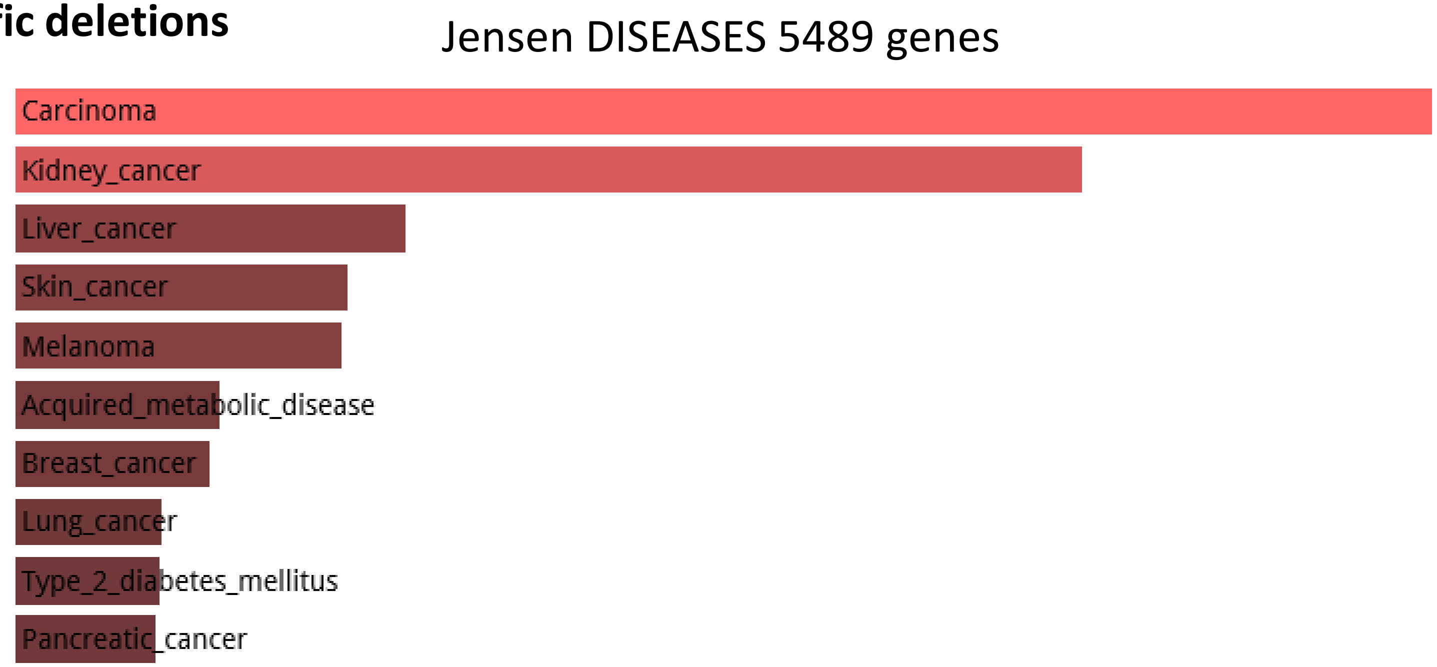


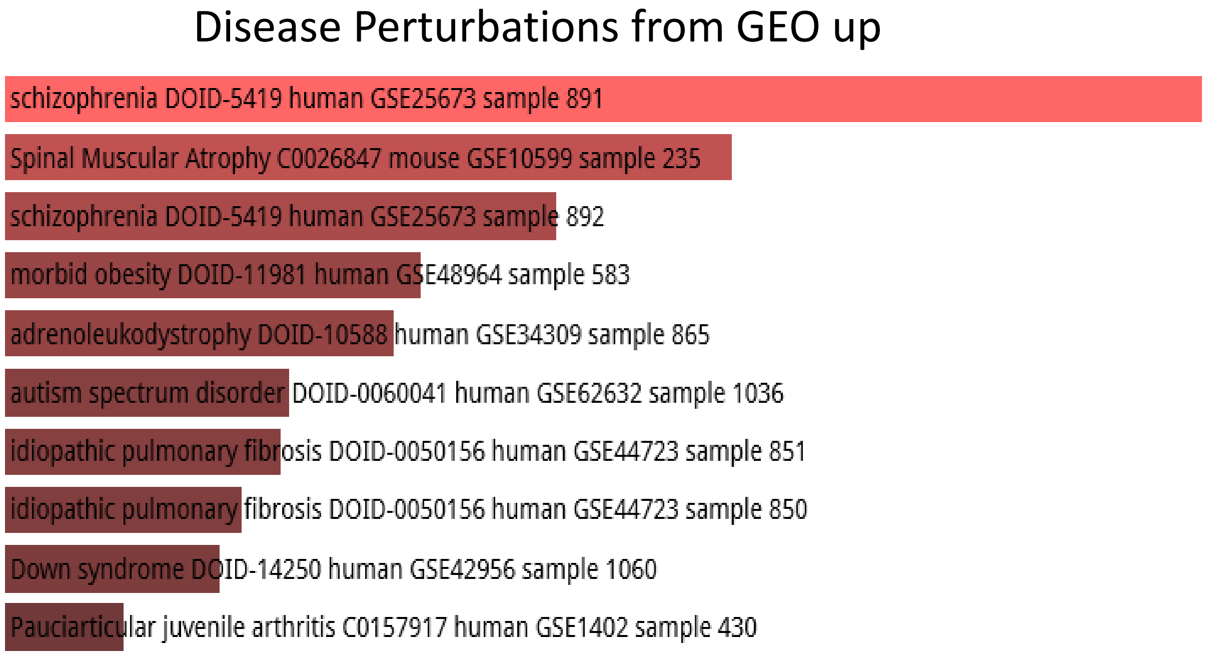


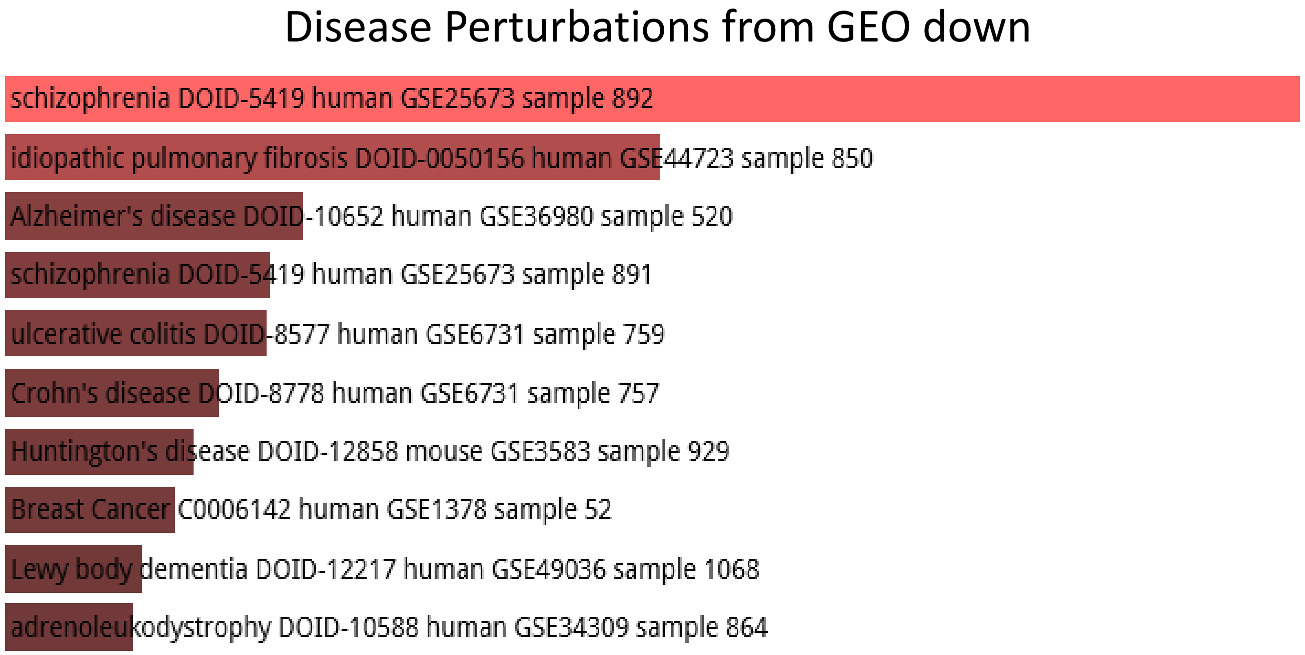

**4,875 human-specific STR expansions**

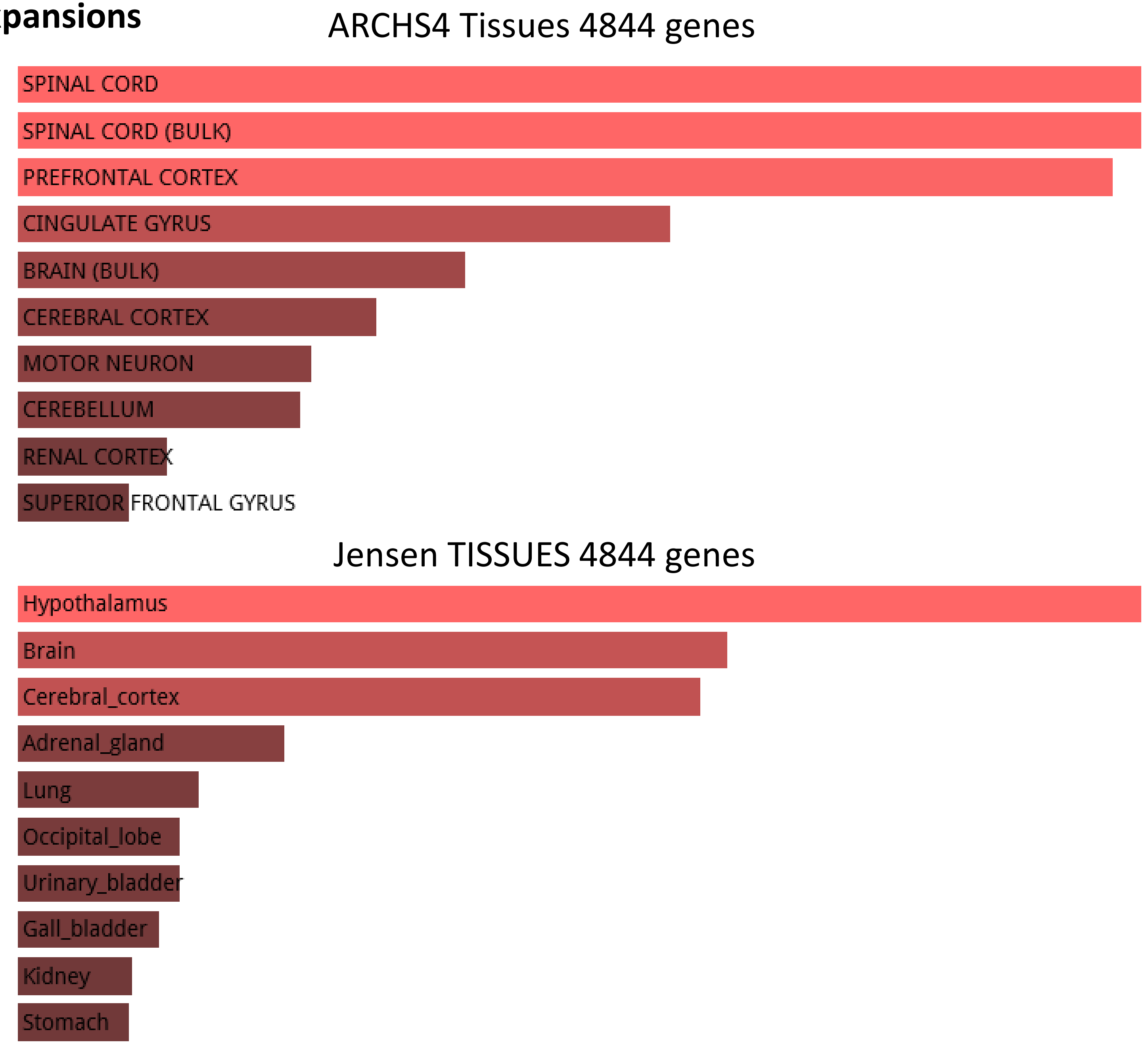

**4,875 human-specific STR expansions**

#### Jensen DISEASES 4844 genes

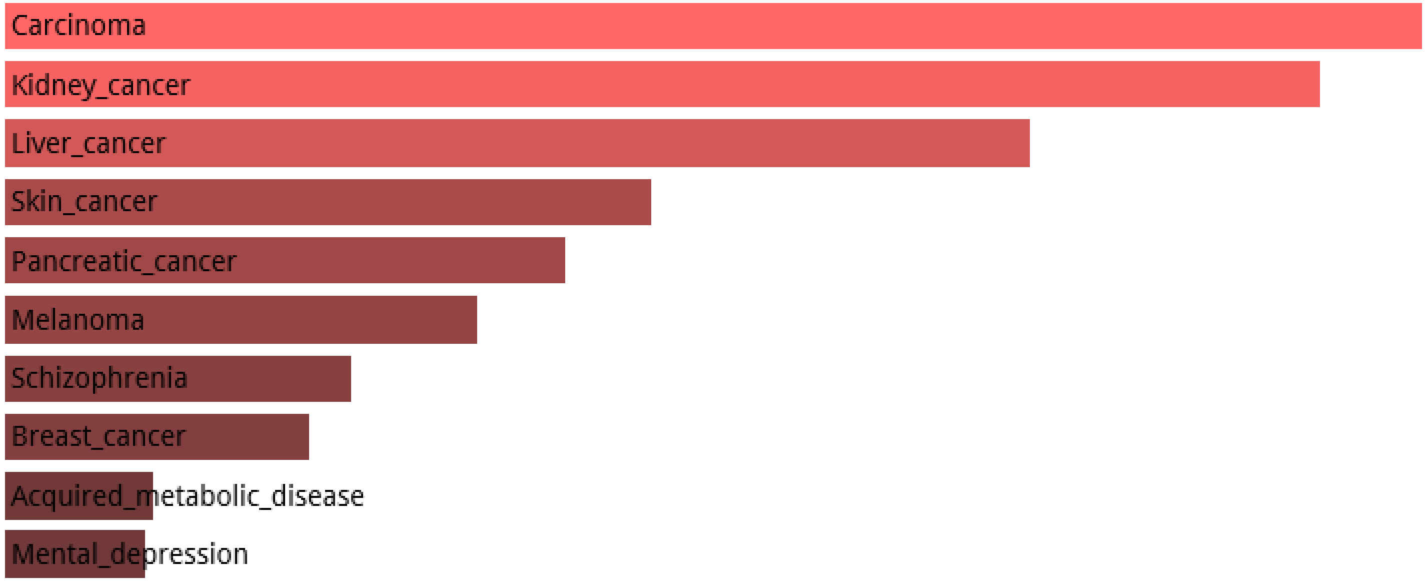


#### Disease Perturbations from GEO up

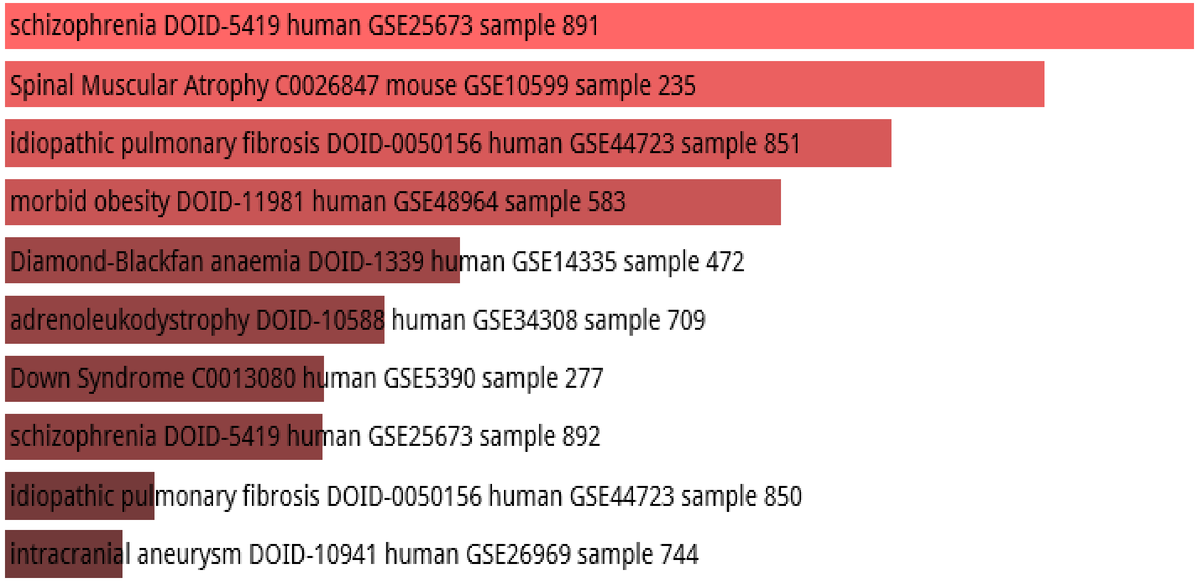


#### Disease Perturbations from GEO down

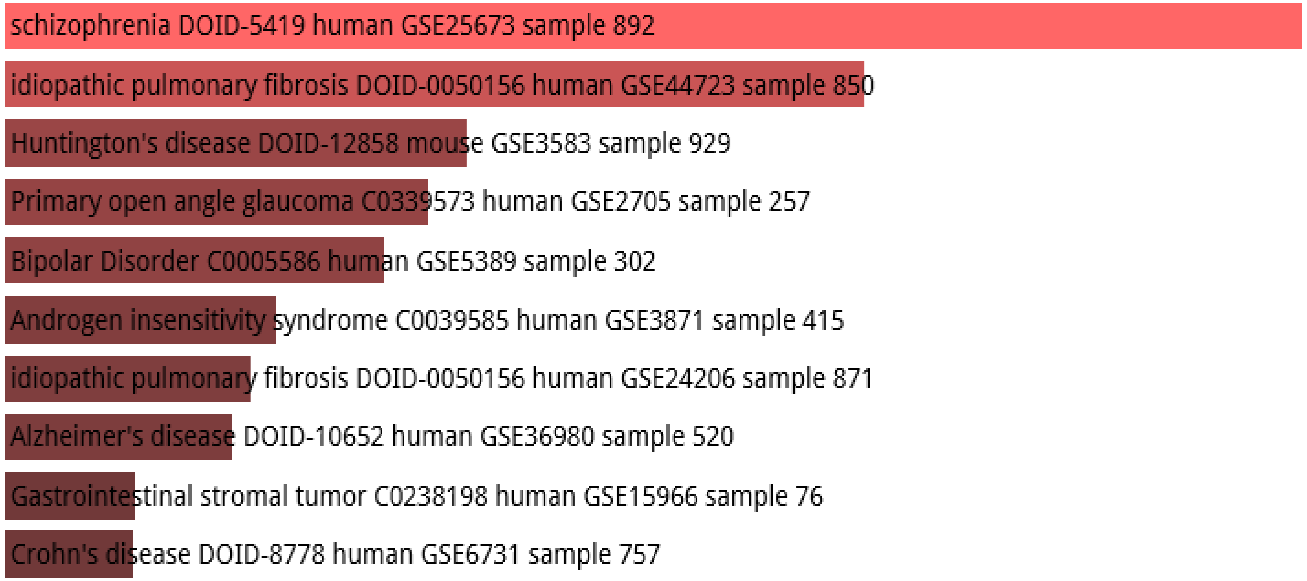

**4,637 human-specific TE-encoded loci expressed in human DLPFC**

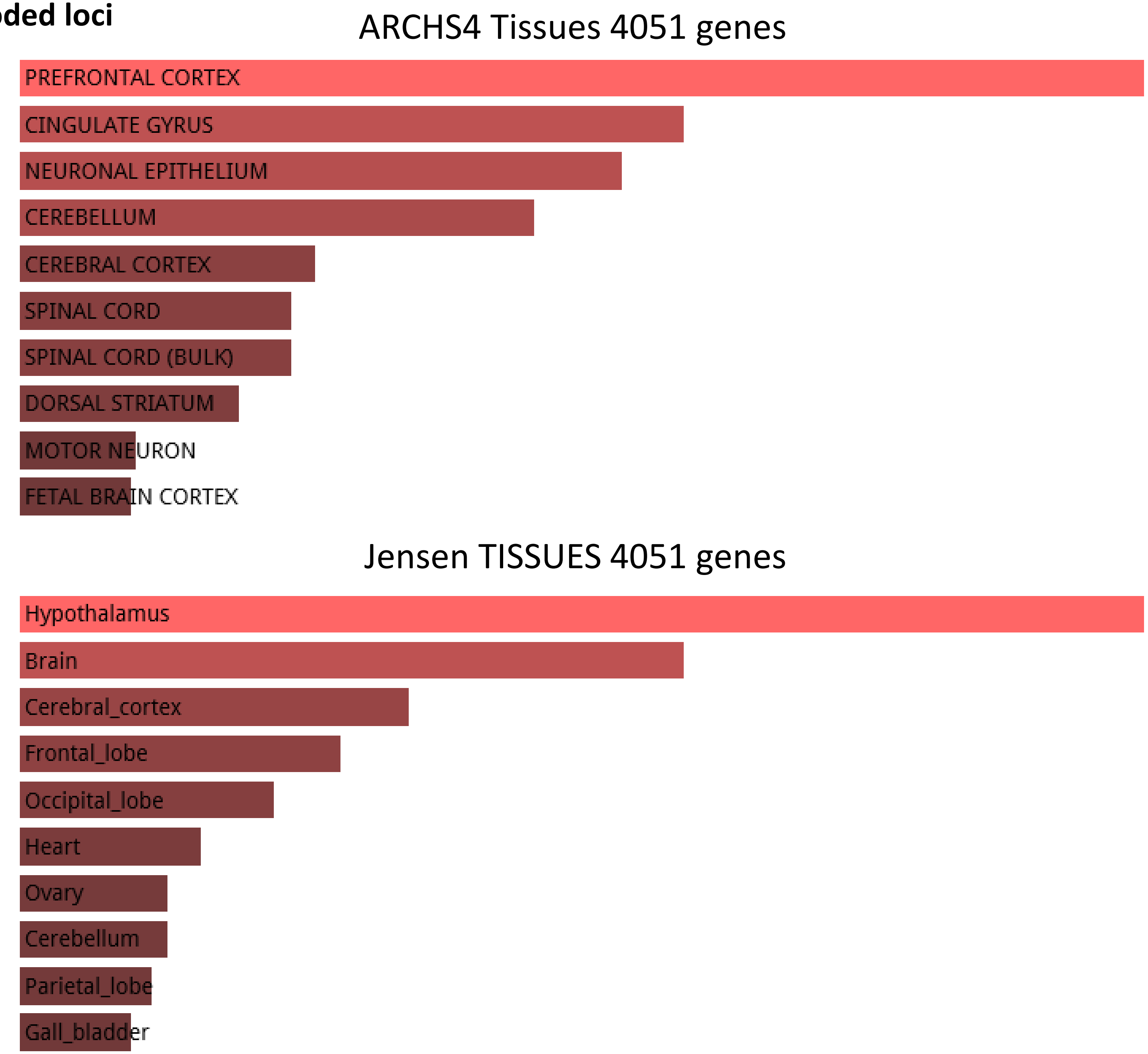

**4,637 human-specific TE-encoded loci expressed in human DLPFC**

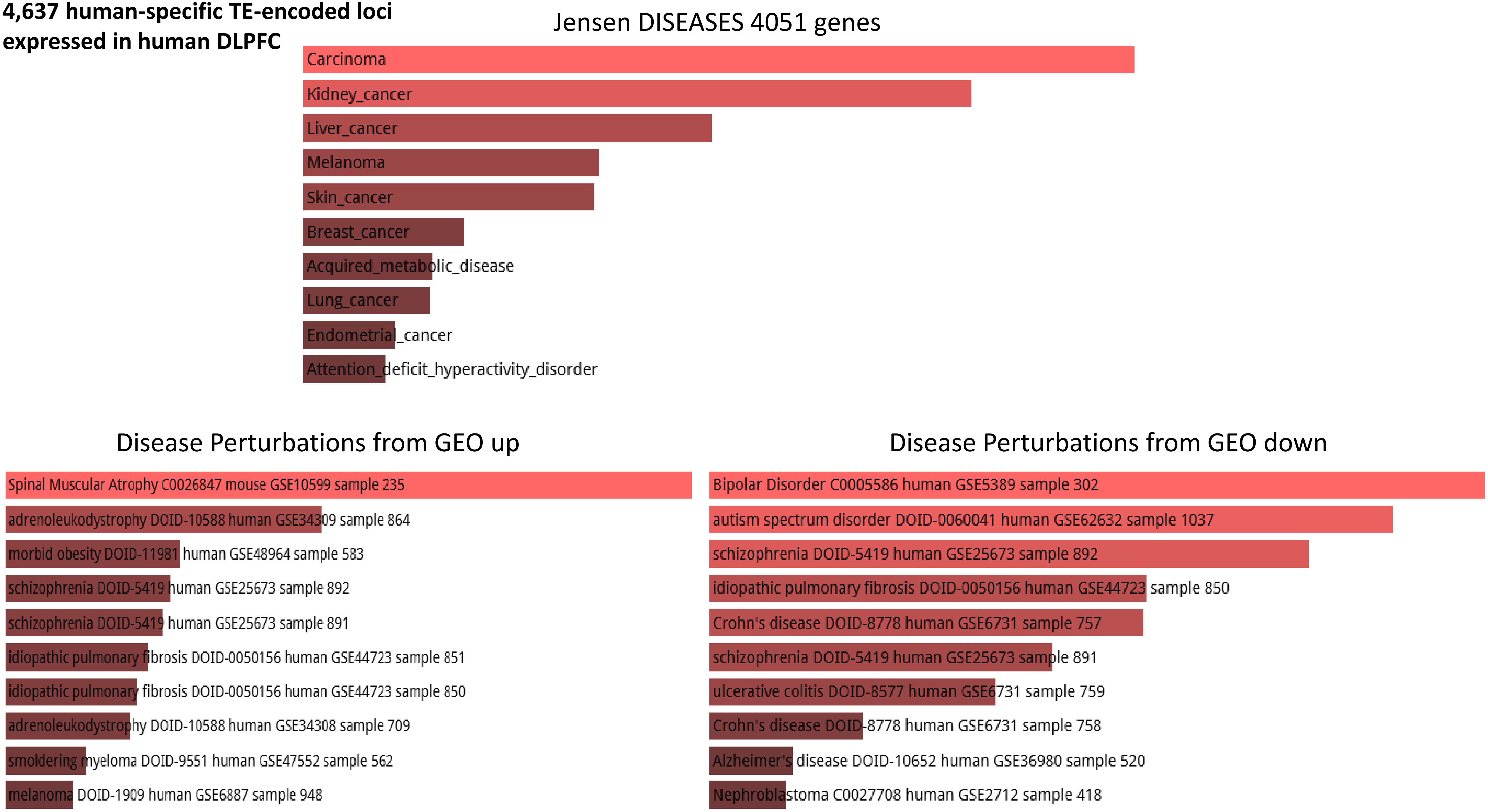

**4,637 human-specific TE-encoded loci expressed in human DLPFC**

Allen Brain Atlas up 4051 genes

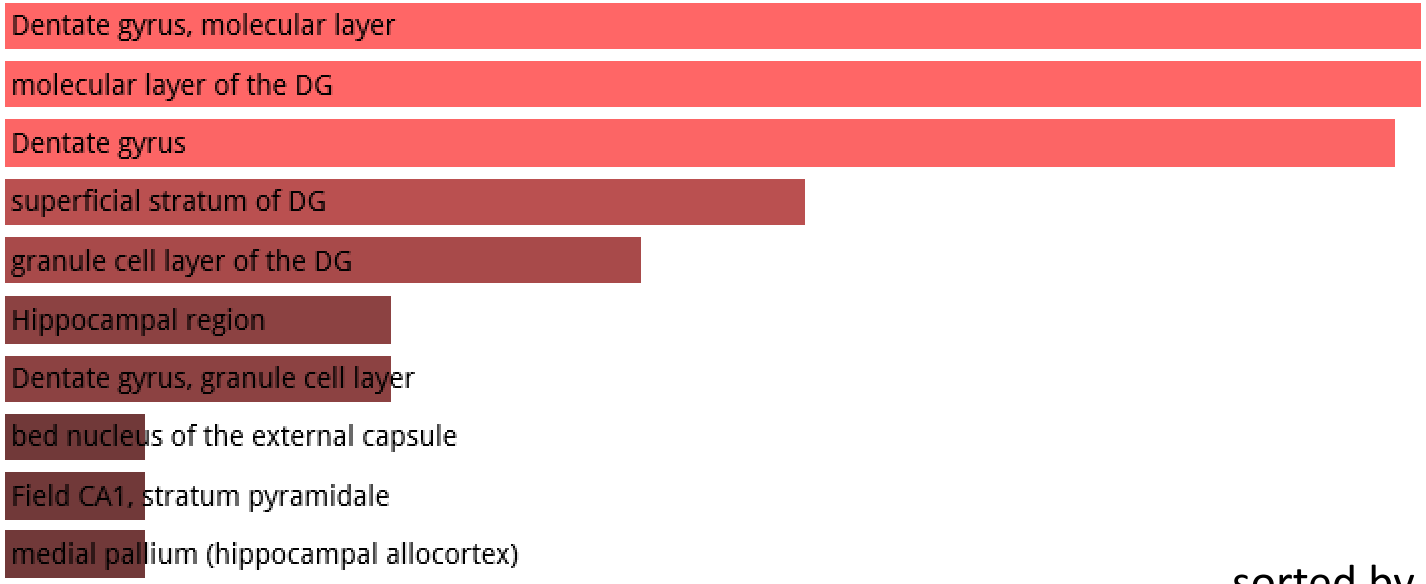


sorted by p-value ranking

#### TOP 15 CATEGORIES

| Term | Overlap | P-value | Adjusted P-value |
|---|---|---|---|
| Dentate gyrus | 110/320 | 2.07E-09 | 1.51E-06 |
| Dentate gyrus, molecular layer | 105/301 | 1.92E-09 | 1.51E-06 |
| molecular layer of the DG | 105/301 | 1.92E-09 | 1.51E-06 |
| superficial stratum of DG | 179/601 | 1.14E-08 | 6.24E-06 |
| granule cell layer of the DG | 102/301 | 1.83E-08 | 8.03E-06 |
| Hippocampal region | 101/301 | 3.78E-08 | 1.18E-05 |
| Dentate gyrus, granule cell layer | 101/301 | 3.78E-08 | 1.18E-05 |
| bed nucleus of the external capsule | 100/301 | 7.67E-08 | 1.68E-05 |
| Field CA1, stratum pyramidale | 100/301 | 7.67E-08 | 1.68E-05 |
| medial pallium (hippocampal allocortex) | 100/301 | 7.67E-08 | 1.68E-05 |
| mantle zone of DG | 174/599 | 1.22E-07 | 2.4E-05 |
| Subiculum, dorsal part, molecular layer | 99/301 | 1.54E-07 | 2.4E-05 |
| hilus of the DG | 99/301 | 1.54E-07 | 2.4E-05 |
| Dentate gyrus, polymorph layer | 99/301 | 1.54E-07 | 2.4E-05 |
| hippocampus (cortex Ammonis) | 98/301 | 3.03E-07 | 2.88E-05 |

**4,637 human-specific TE-encoded loci expressed in human DLPFC**

Allen Brain Atlas up 4051 genes

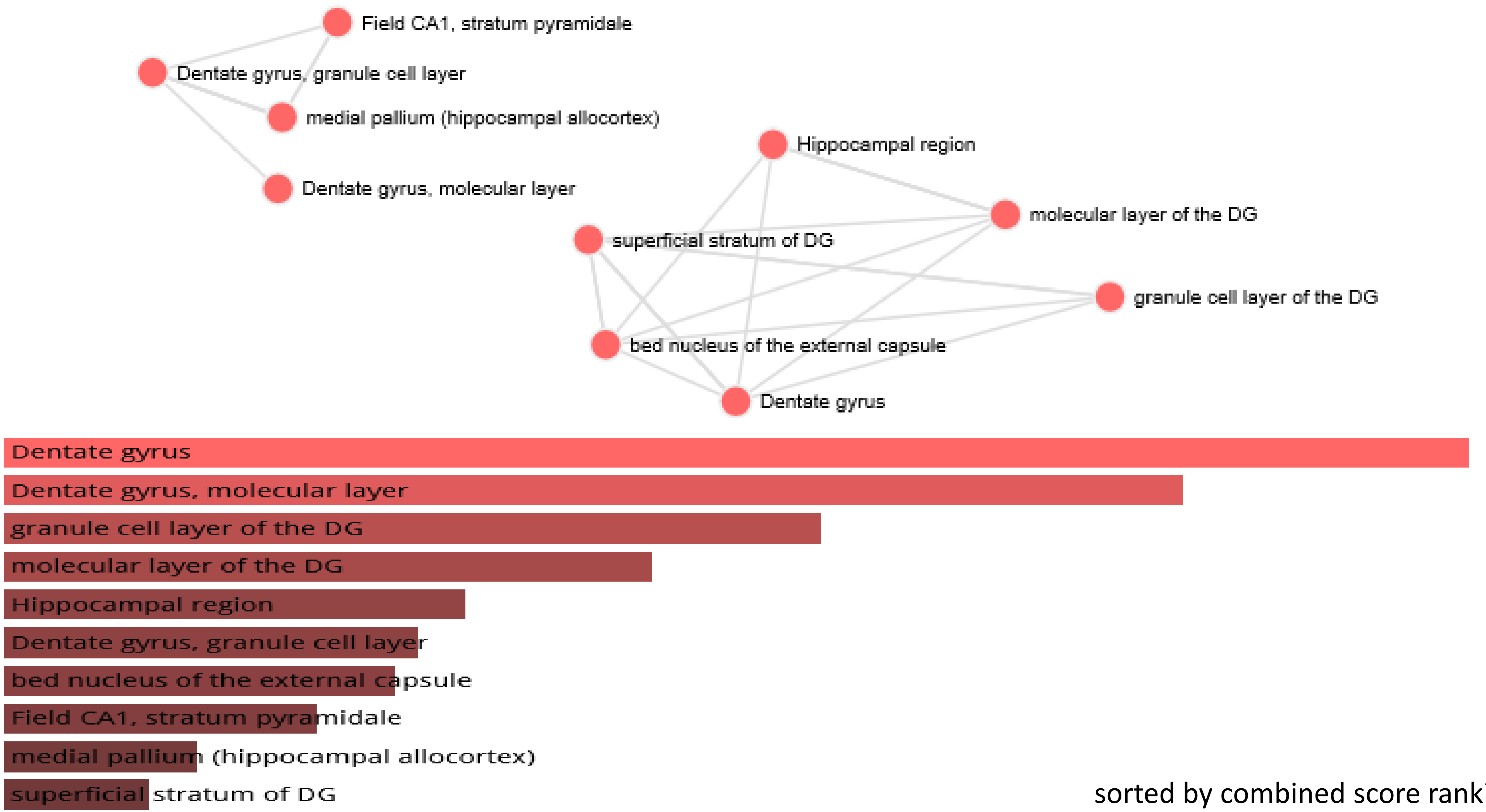


sorted by combined score ranking

**4,637 human-specific TE-encoded loci expressed in human DLPFC**

Tissue Protein Expression from Human Proteome Map: 4051 genes

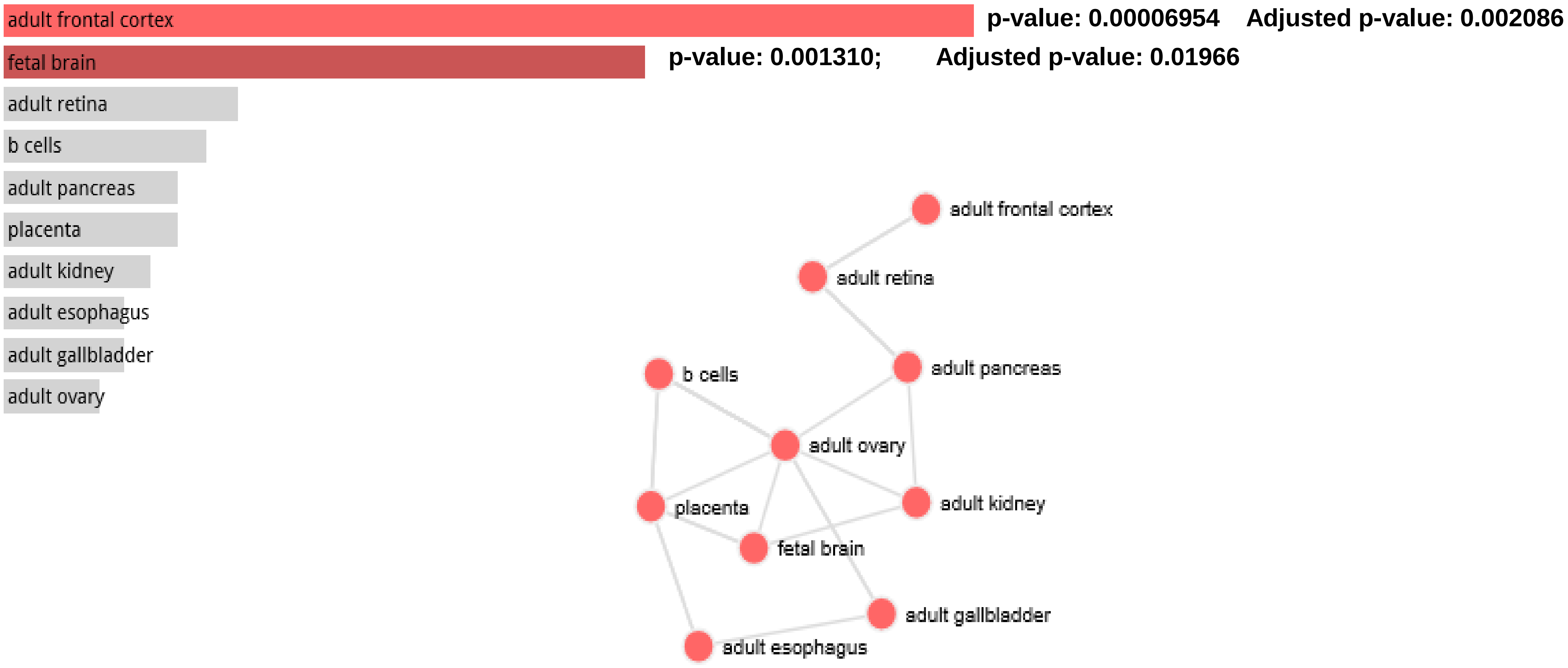

3,538 human-specific ace-DHS

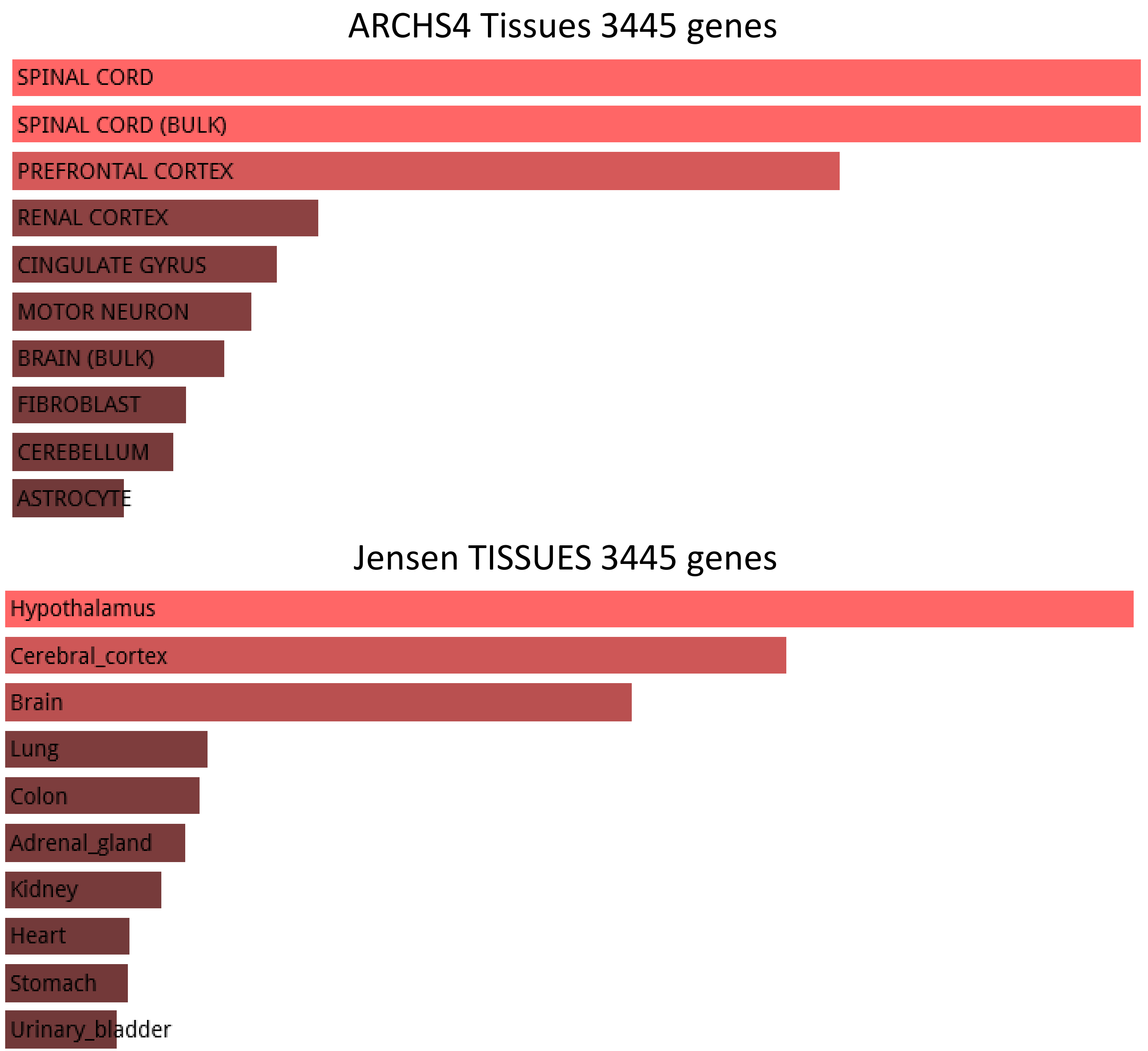

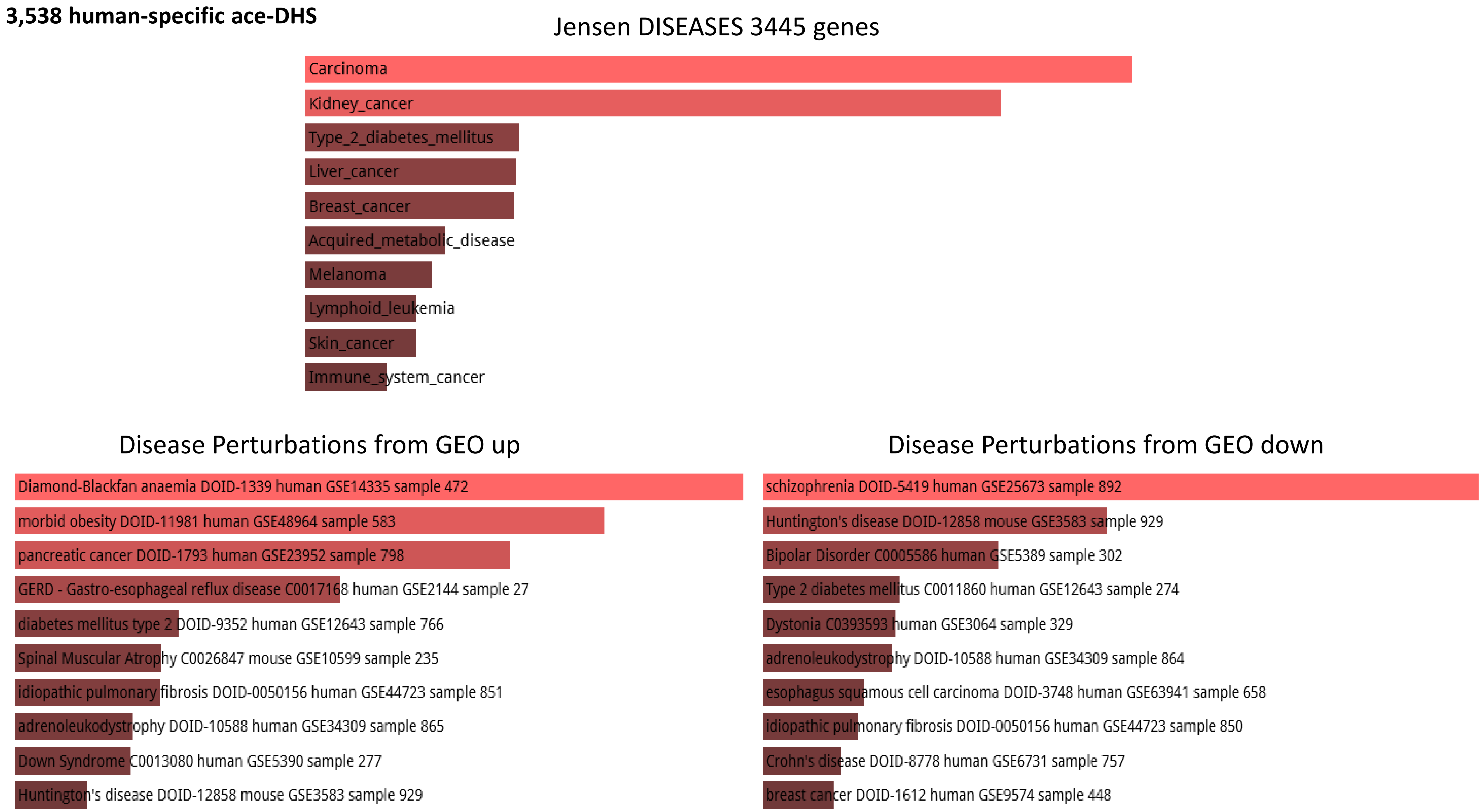
3,538 human-specific ace-DHS
Jensen DISEASES 3445 genes
Carcinoma
Kidney_cancer
Type_2_diabetes_mellitus
Liver_cancer
Breast_cancer
Acquired_metabolic_disease
Melanoma
Lymphoid_leukemia
Skin_cancer
Immune_system_cancer
Disease Perturbations from GEO up
Diamond-Blackfan anaemia DOID-1339 human GSE14335 sample 472
morbid obesity DOID-11981 human GSE48964 sample 583
pancreatic cancer DOID-1793 human GSE23952 sample 798
GERD - Gastro-esophageal reflux disease C0017168 human GSE2144 sample 27
diabetes mellitus type 2 DOID-9352 human GSE12643 sample 766
Spinal Muscular Atrophy C0026847 mouse GSE10599 sample 235
idiopathic pulmonary fibrosis DOID-0050156 human GSE44723 sample 851
adrenoleukodystrophy DOID-10588 human GSE34309 sample 865
Down Syndrome C0013080 human GSE5390 sample 277
Huntington's disease DOID-12858 mouse GSE3583 sample 929
Disease Perturbations from GEO down
schizophrenia DOID-5419 human GSE25673 sample 892
Huntington's disease DOID-12858 mouse GSE3583 sample 929
Bipolar Disorder C0005586 human GSE5389 sample 302
Type 2 diabetes mellitus C0011860 human GSE12643 sample 274
Dystonia C0393593 human GSE3064 sample 329
adrenoleukodystrophy DOID-10588 human GSE34309 sample 864
esophagus squamous cell carcinoma DOID-3748 human GSE63941 sample 658
idiopathic pulmonary fibrosis DOID-0050156 human GSE44723 sample 850
Crohn's disease DOID-8778 human GSE6731 sample 757
breast cancer DOID-1612 human GSE9574 sample 448

**3,538 human-specific ace-DHS**

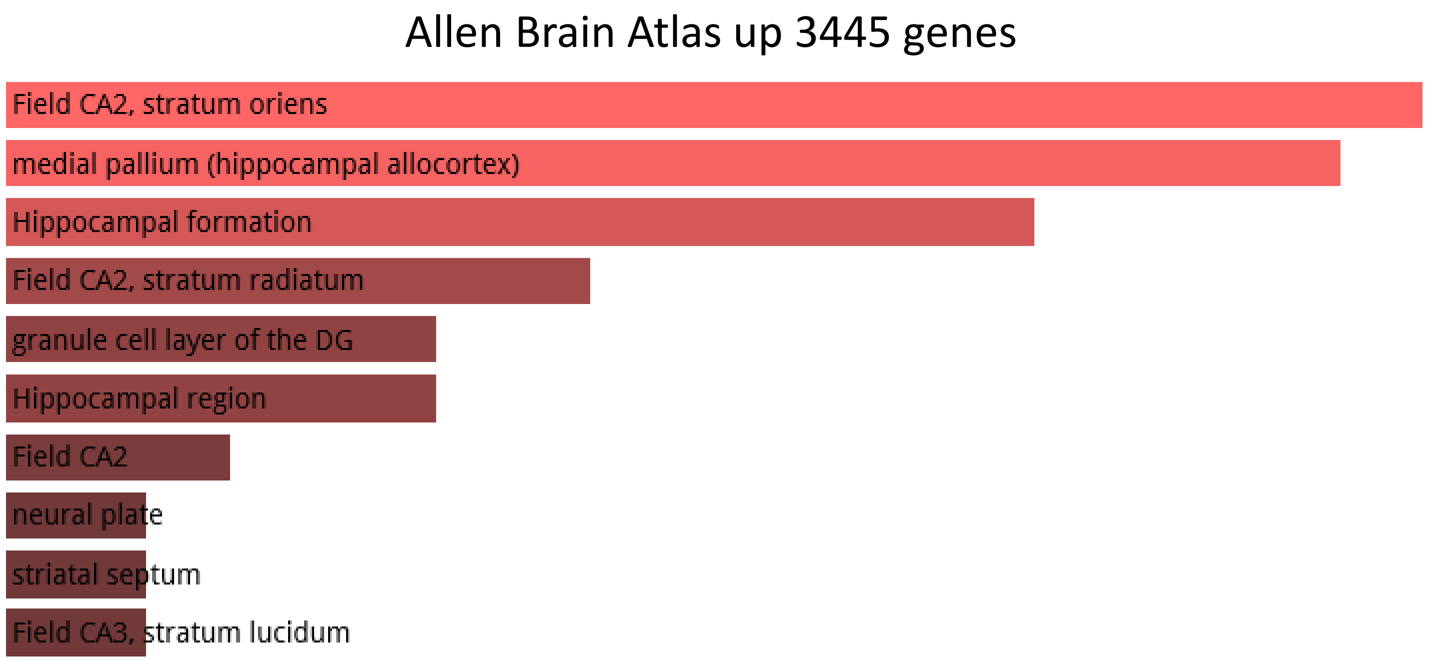


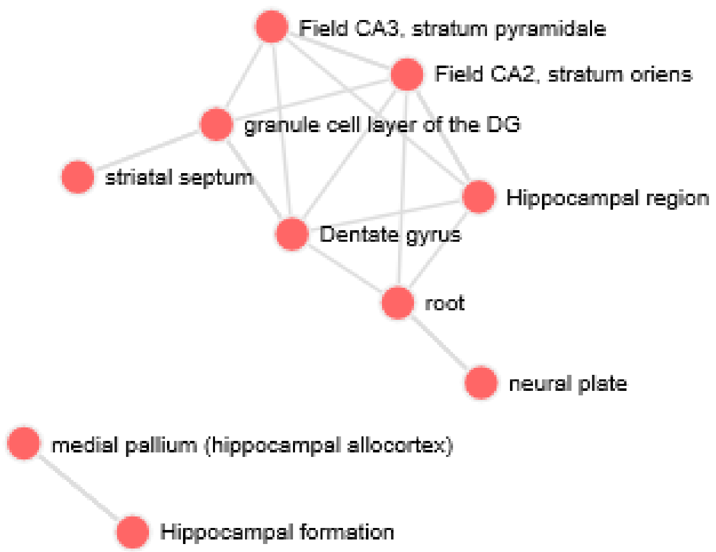

3,803 human-specific TFBS in hESC

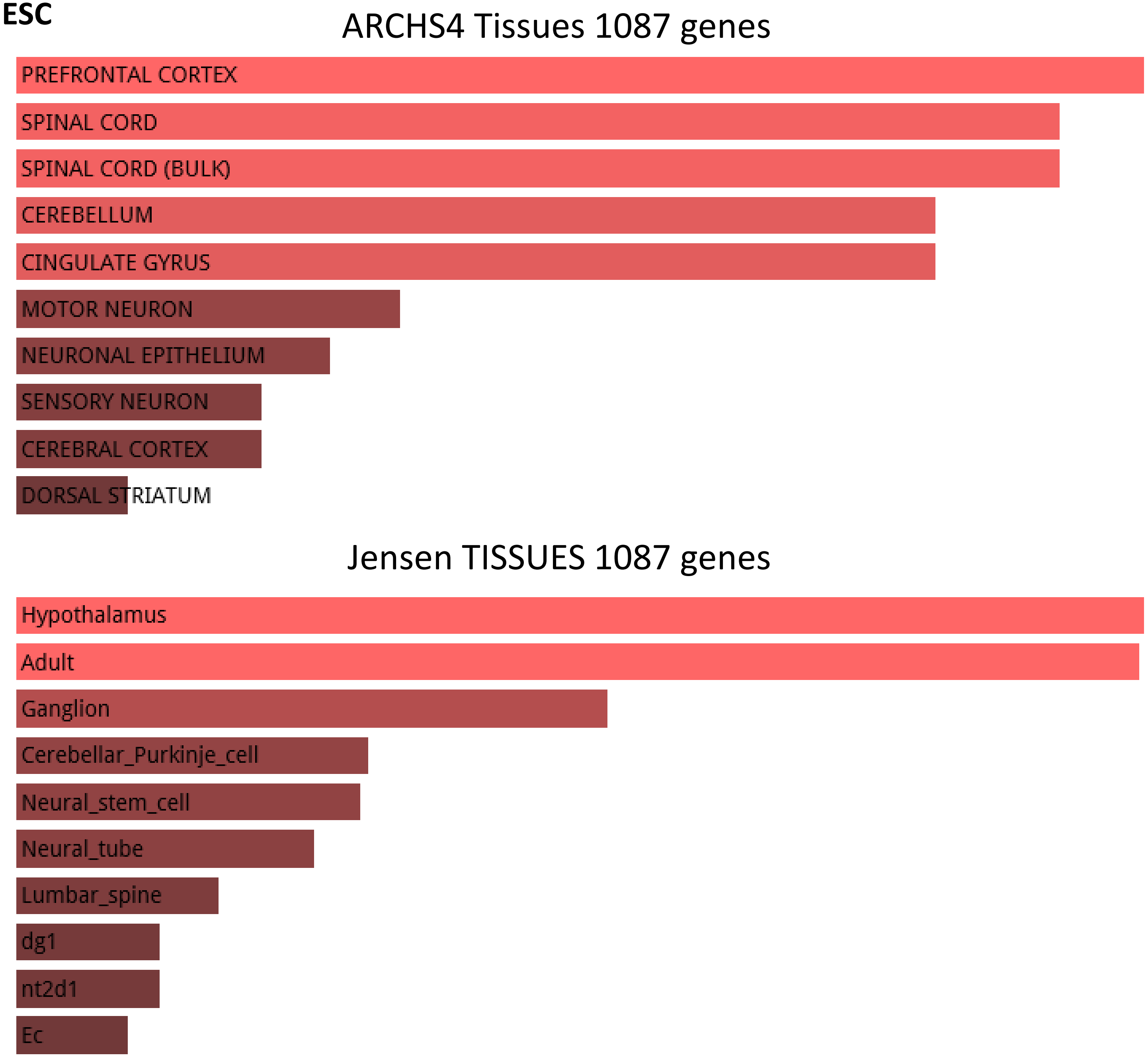

**3,803 human-specific TFBS in hESC**

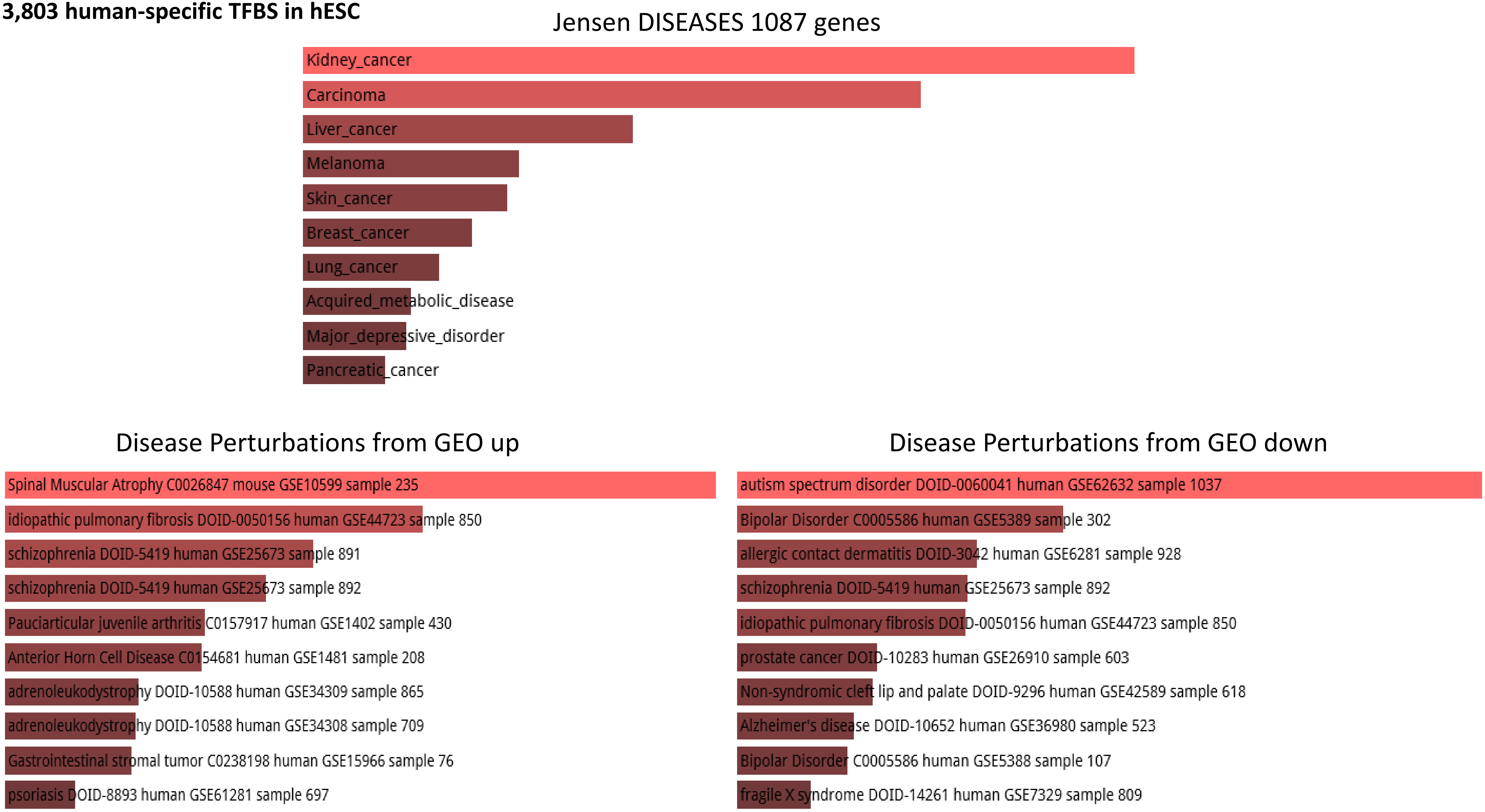

**4,249 fixed human-specific regulatory loci**

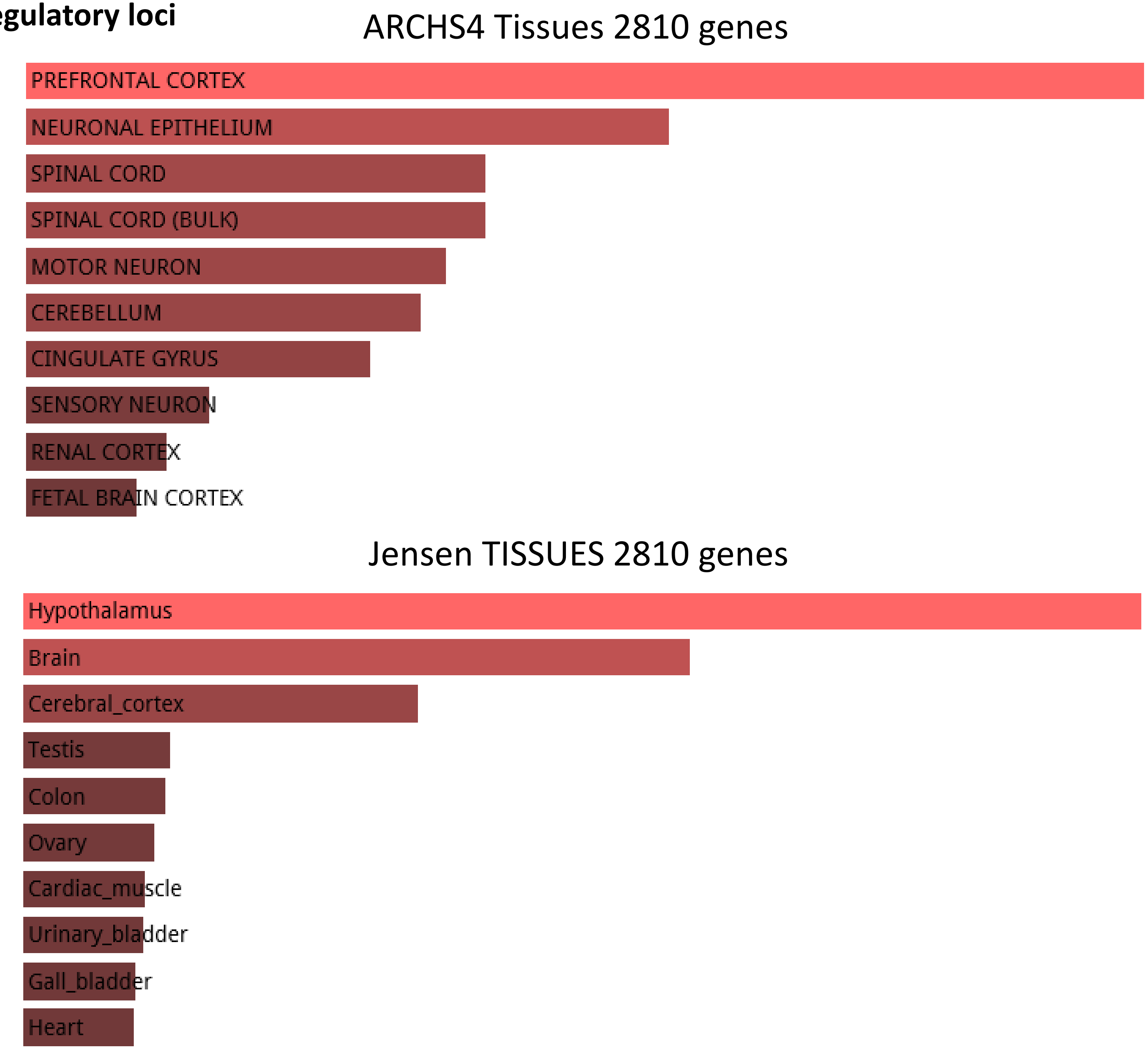

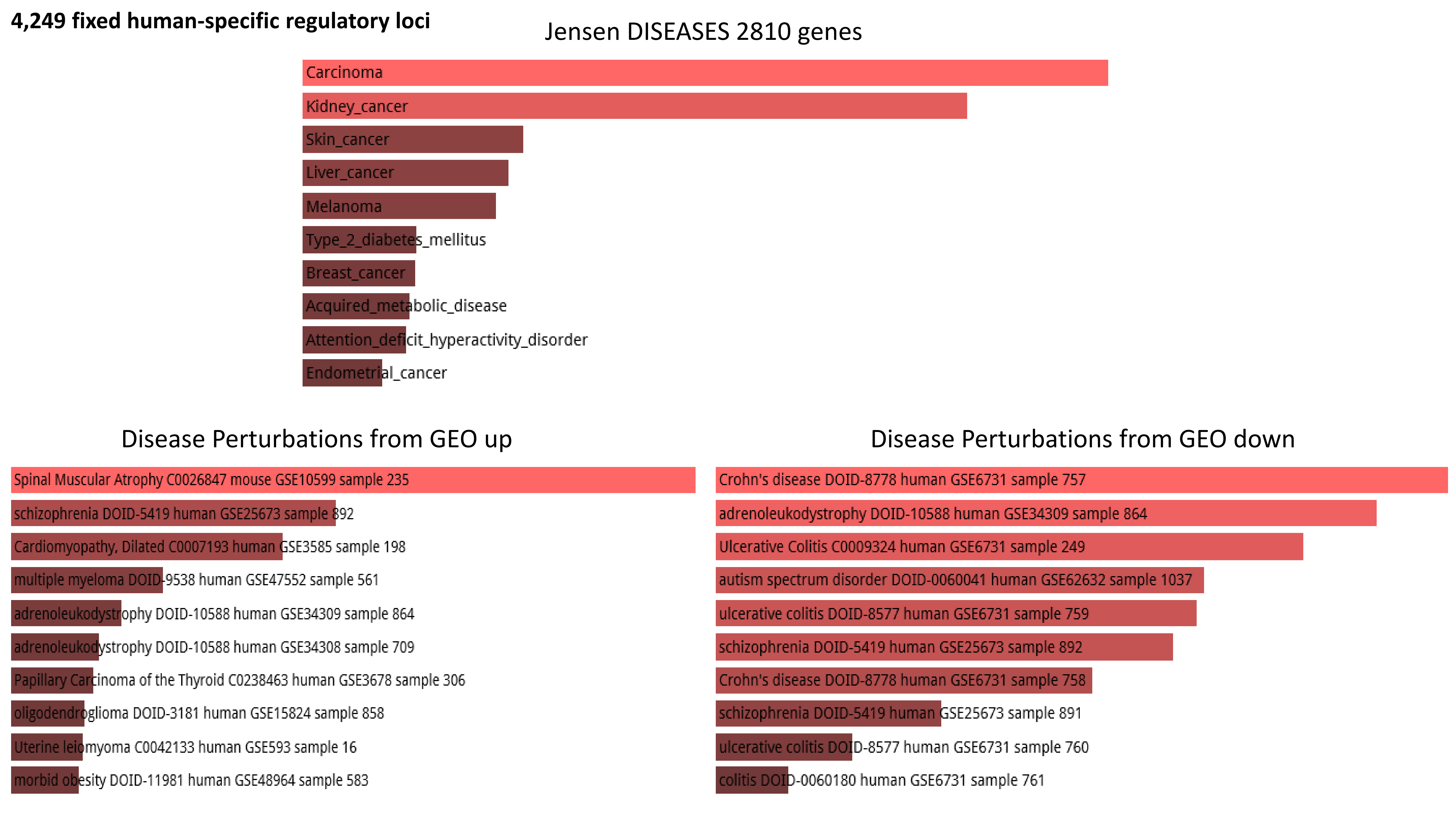
4,249 fixed human-specific regulatory loci
Jensen DISEASES 2810 genes
Carcinoma
Kidney_cancer
Skin_cancer
Liver_cancer
Melanoma
Type_2_diabetes_mellitus
Breast_cancer
Acquired_metabolic_disease
Attention_deficit_hyperactivity_disorder
Endometrial_cancer
Disease Perturbations from GEO up
Spinal Muscular Atrophy C0026847 mouse GSE10599 sample 235
schizophrenia DOID-5419 human GSE25673 sample 892
Cardiomyopathy, Dilated C0007193 human GSE3585 sample 198
multiple myeloma DOID-9538 human GSE47552 sample 561
adrenoleukodystrophy DOID-10588 human GSE34309 sample 864
adrenoleukodystrophy DOID-10588 human GSE34308 sample 709
Papillary Carcinoma of the Thyroid C0238463 human GSE3678 sample 306
oligodendroglioma DOID-3181 human GSE15824 sample 858
Uterine leiomyoma C0042133 human GSE593 sample 16
morbid obesity DOID-11981 human GSE48964 sample 583
Disease Perturbations from GEO down
Crohn's disease DOID-8778 human GSE6731 sample 757
adrenoleukodystrophy DOID-10588 human GSE34309 sample 864
Ulcerative Colitis C0009324 human GSE6731 sample 249
autism spectrum disorder DOID-0060041 human GSE62632 sample 1037
ulcerative colitis DOID-8577 human GSE6731 sample 759
schizophrenia DOID-5419 human GSE25673 sample 892
Crohn's disease DOID-8778 human GSE6731 sample 758
schizophrenia DOID-5419 human GSE25673 sample 891
ulcerative colitis DOID-8577 human GSE6731 sample 760
colitis DOID-0060180 human GSE6731 sample 761

**2,745 human accelerated regions**

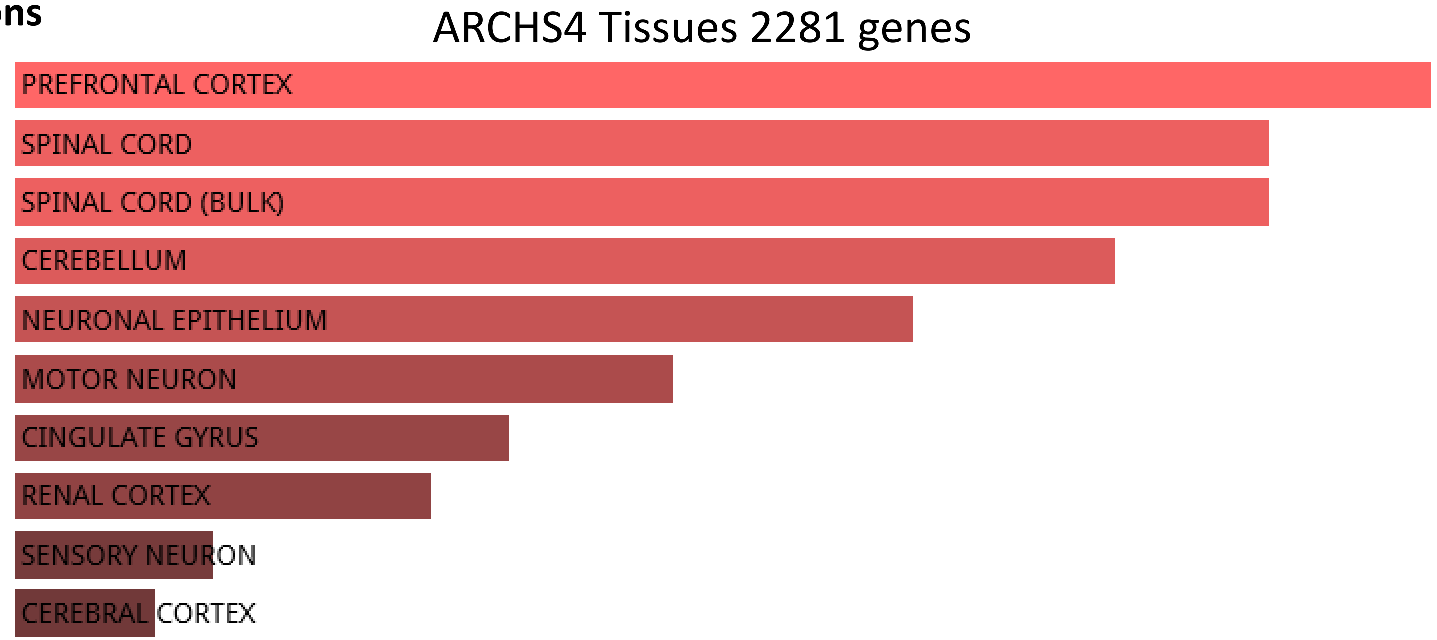


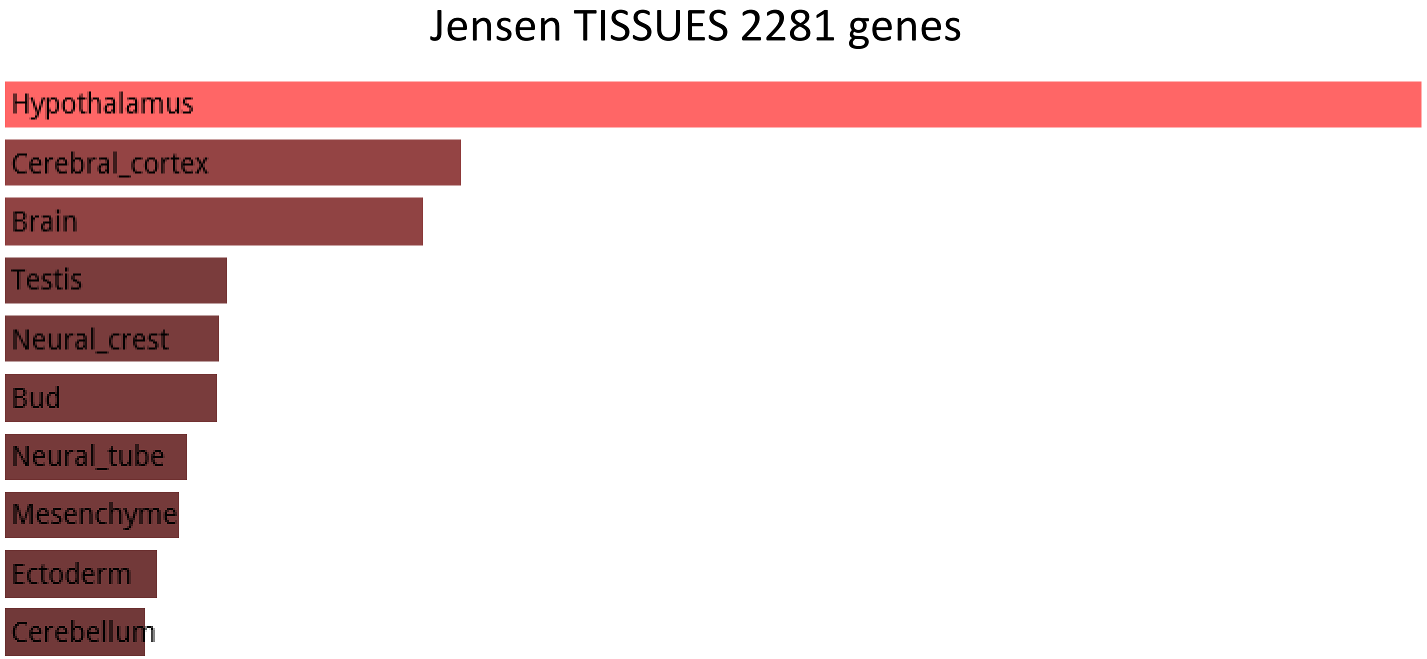

**2,745 human accelerated regions**

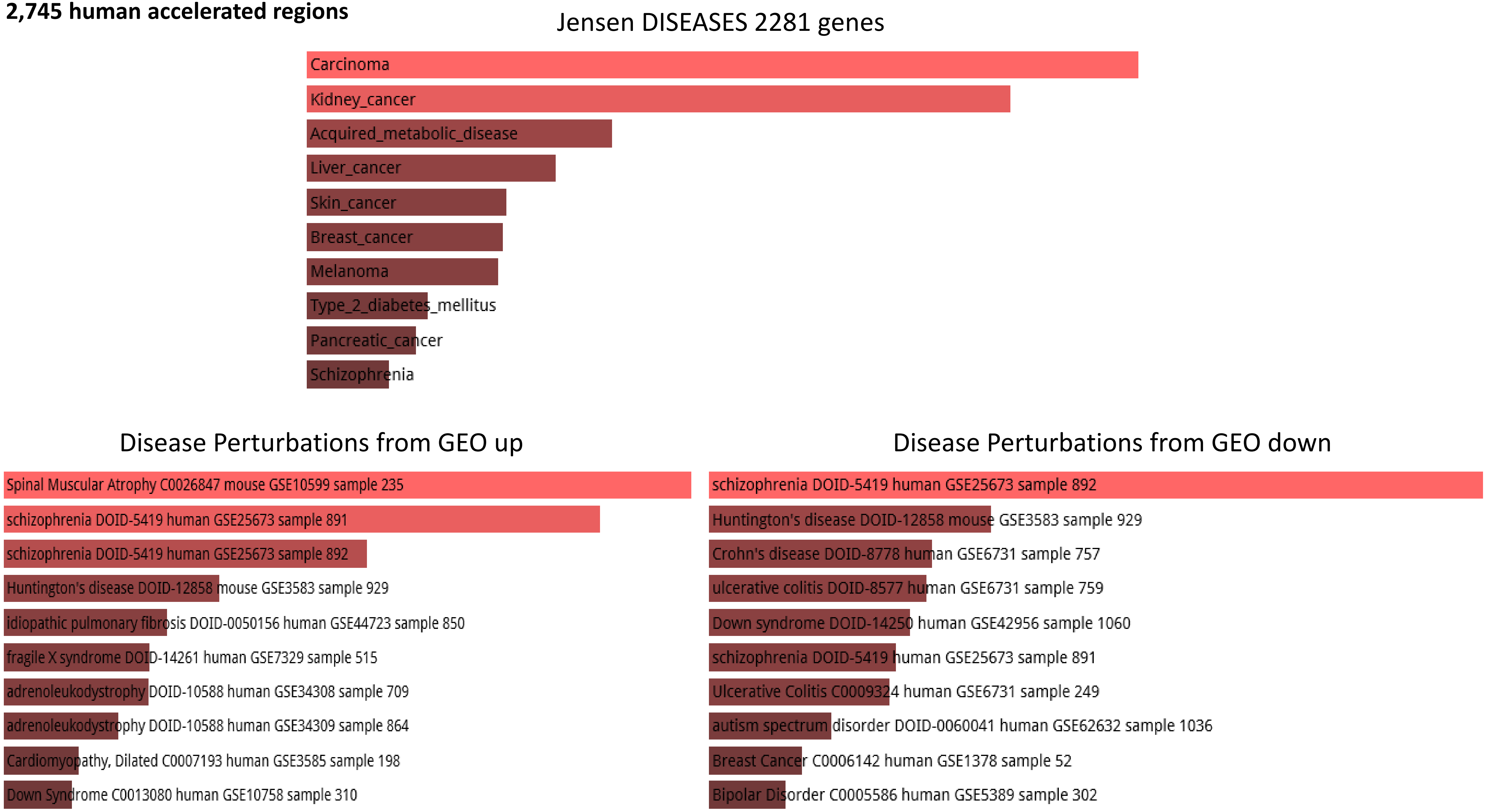

**1,932 fixed human-specific regulatory regions (hESC DHS)**

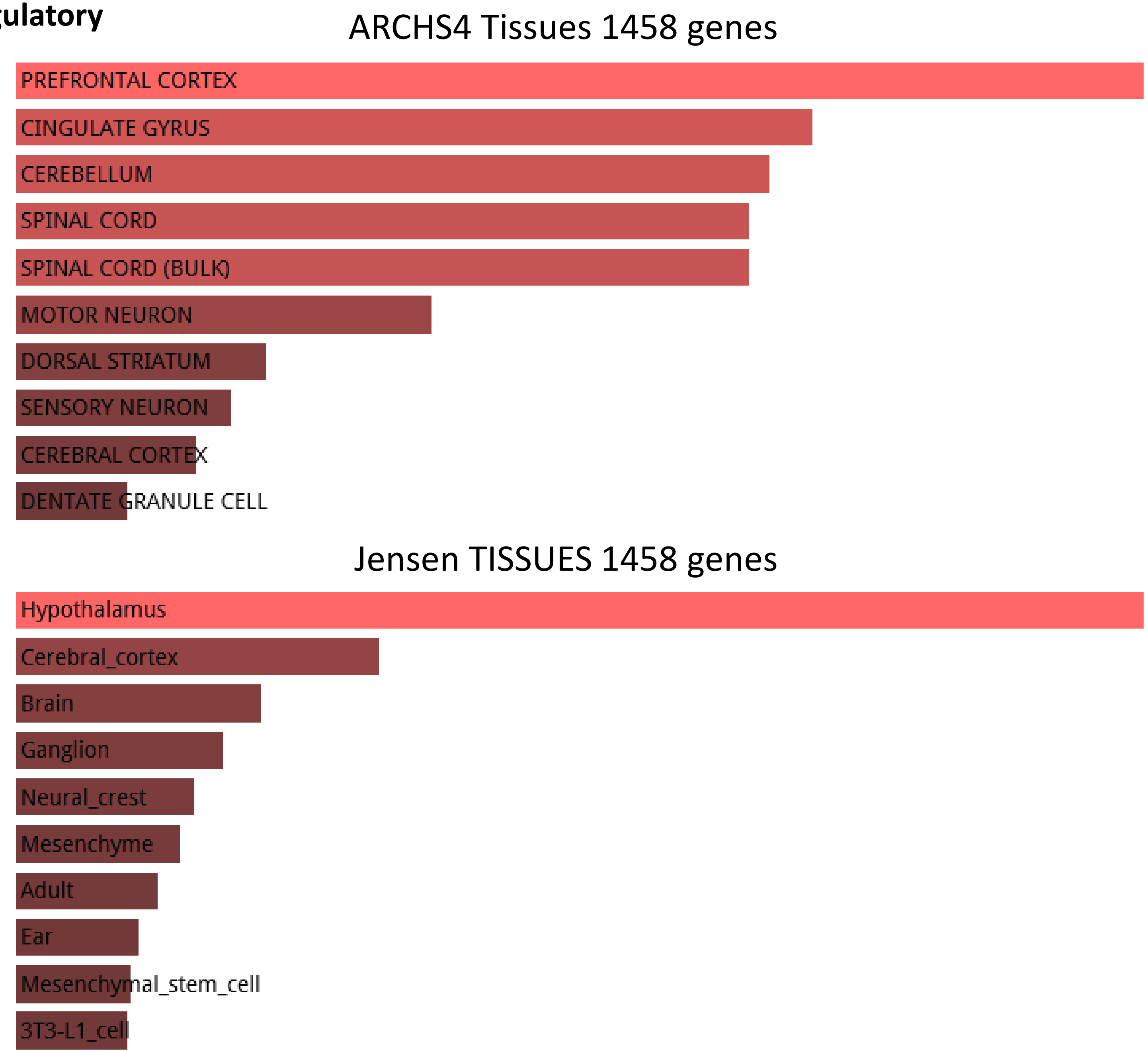

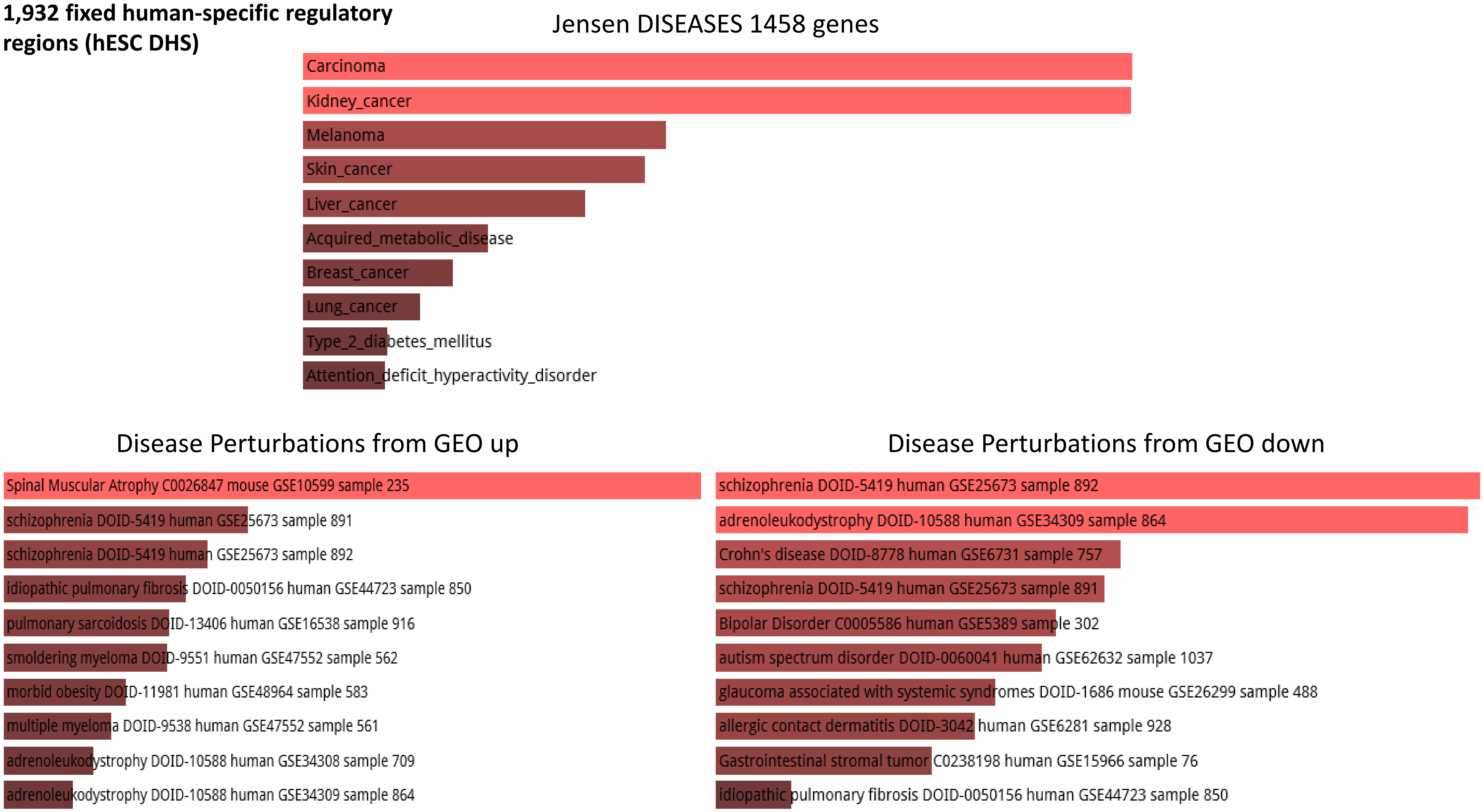
1,932 fixed human-specific regulatory regions (hESC DHS)
Jensen DISEASES 1458 genes
Carcinoma
Kidney_cancer
Melanoma
Skin_cancer
Liver_cancer
Acquired_metabolic_disease
Breast_cancer
Lung_cancer
Type_2_diabetes_mellitus
Attention_deficit_hyperactivity_disorder
Disease Perturbations from GEO up
Spinal Muscular Atrophy C0026847 mouse GSE10599 sample 235
schizophrenia DOID-5419 human GSE25673 sample 891
schizophrenia DOID-5419 human GSE25673 sample 892
idiopathic pulmonary fibrosis DOID-0050156 human GSE44723 sample 850
pulmonary sarcoidosis DOID-13406 human GSE16538 sample 916
smoldering myeloma DOID-9551 human GSE47552 sample 562
morbid obesity DOID-11981 human GSE48964 sample 583
multiple myeloma DOID-9538 human GSE47552 sample 561
adrenoleukodystrophy DOID-10588 human GSE34308 sample 709
adrenoleukodystrophy DOID-10588 human GSE34309 sample 864
Disease Perturbations from GEO down
schizophrenia DOID-5419 human GSE25673 sample 892
adrenoleukodystrophy DOID-10588 human GSE34309 sample 864
Crohn's disease DOID-8778 human GSE6731 sample 757
schizophrenia DOID-5419 human GSE25673 sample 891
Bipolar Disorder C0005586 human GSE5389 sample 302
autism spectrum disorder DOID-0060041 human GSE62632 sample 1037
glaucoma associated with systemic syndromes DOID-1686 mouse GSE26299 sample 488
allergic contact dermatitis DOID-3042 human GSE6281 sample 928
Gastrointestinal stromal tumor C0238198 human GSE15966 sample 76
idiopathic pulmonary fibrosis DOID-0050156 human GSE44723 sample 850

**1,000 human-biased cranial neural crest cells (CNCCs) enhancers**

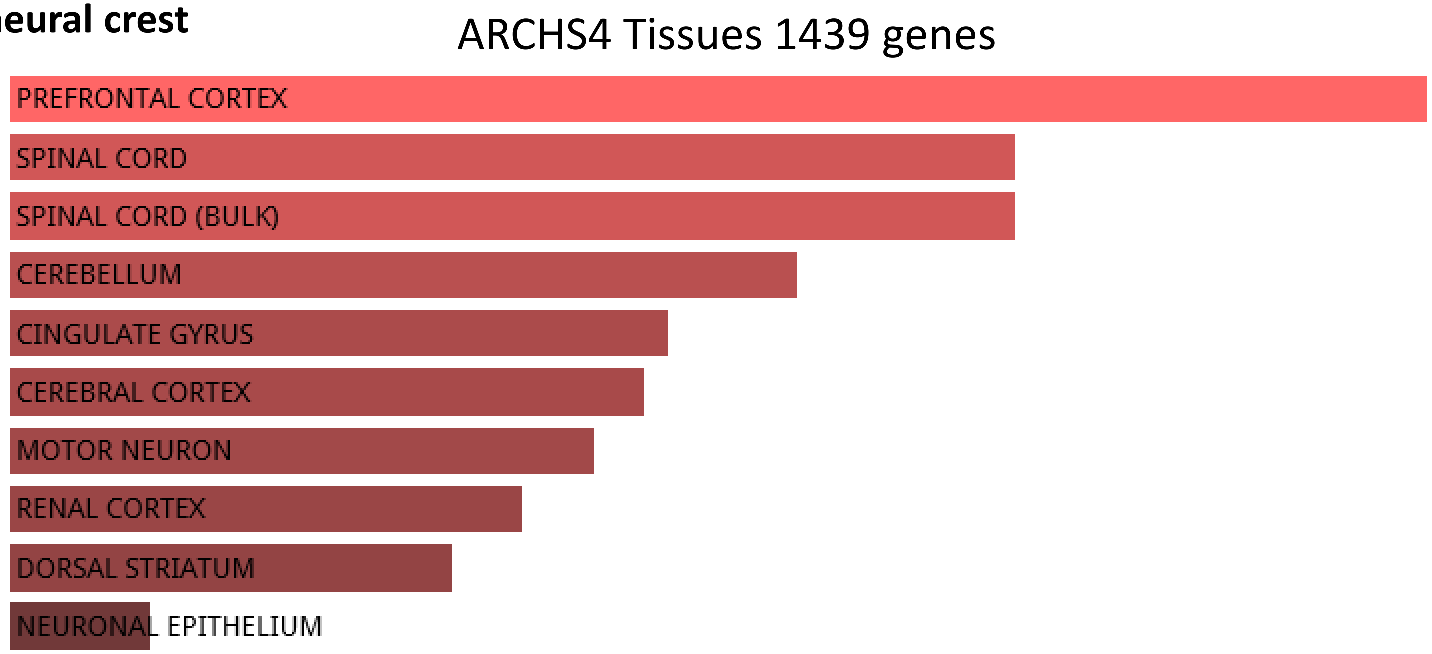


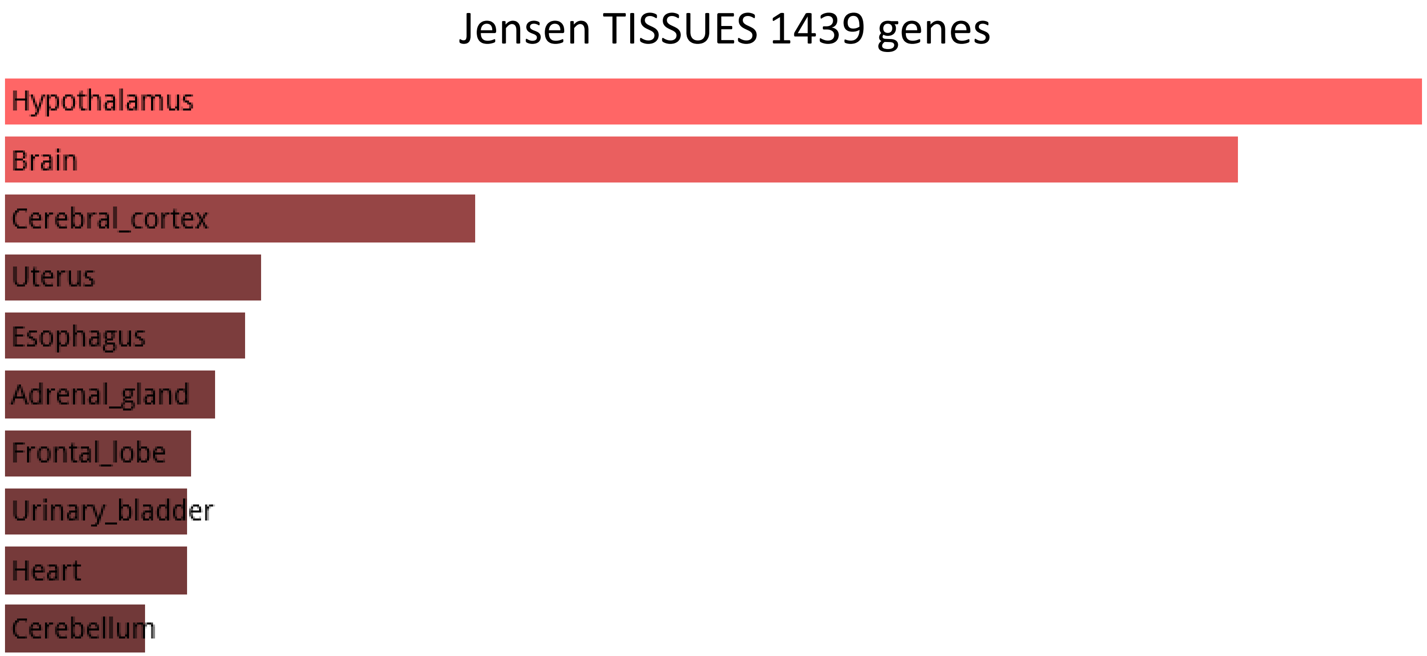

1,000 human-biased cranial neural crest cells (CNCCs) enhancers

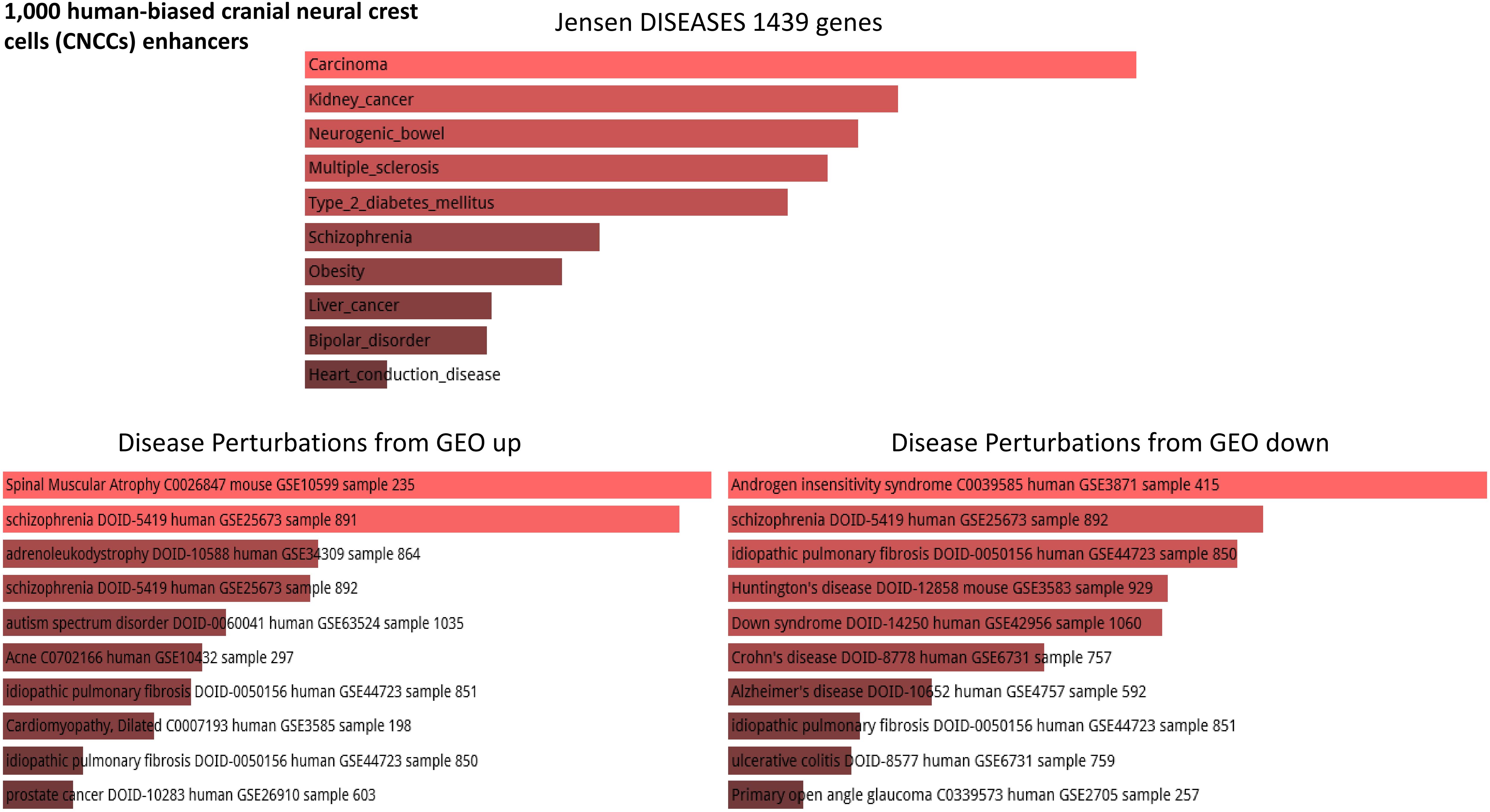

**1,000 chimp-biased cranial neural crest cells (CNCCs) enhancers**

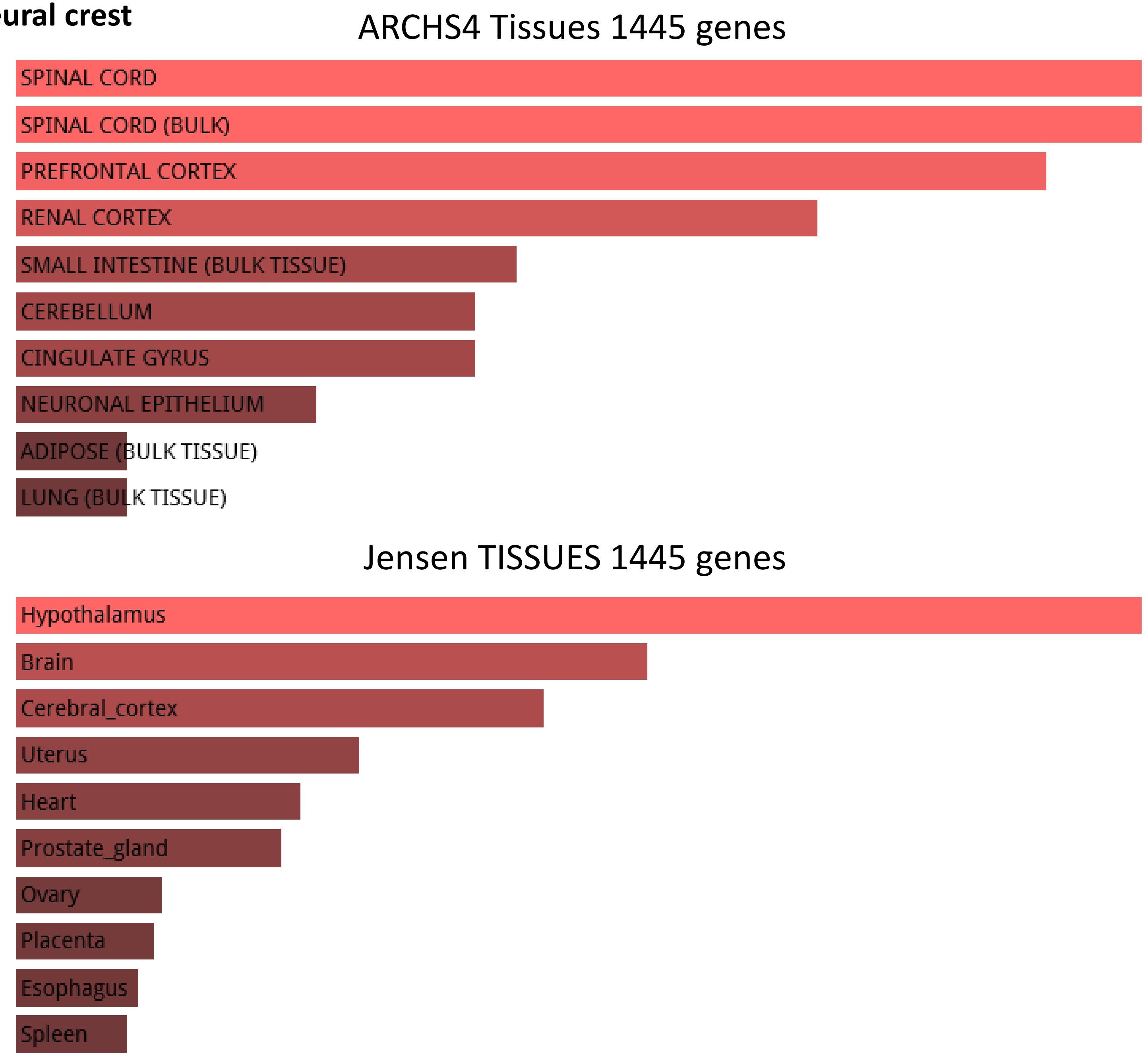

**1,000 chimp-biased cranial neural crest cells (CNCCs) enhancers**

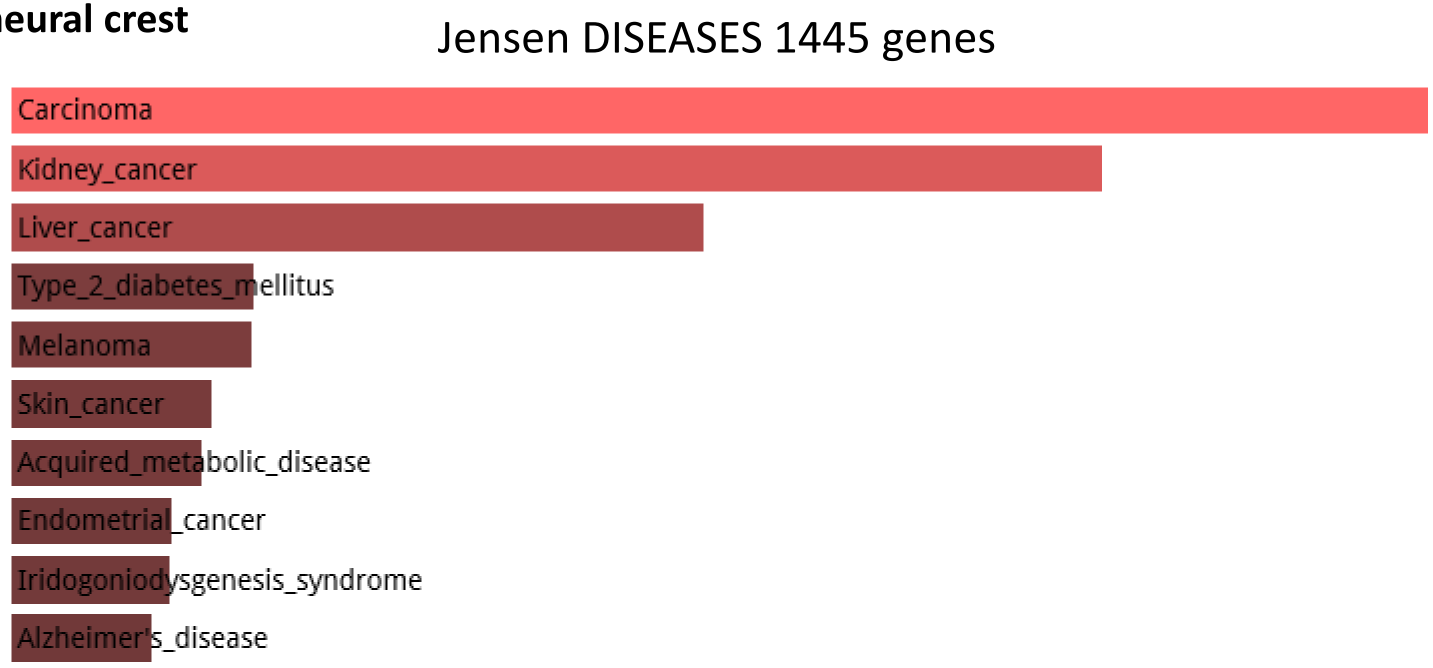


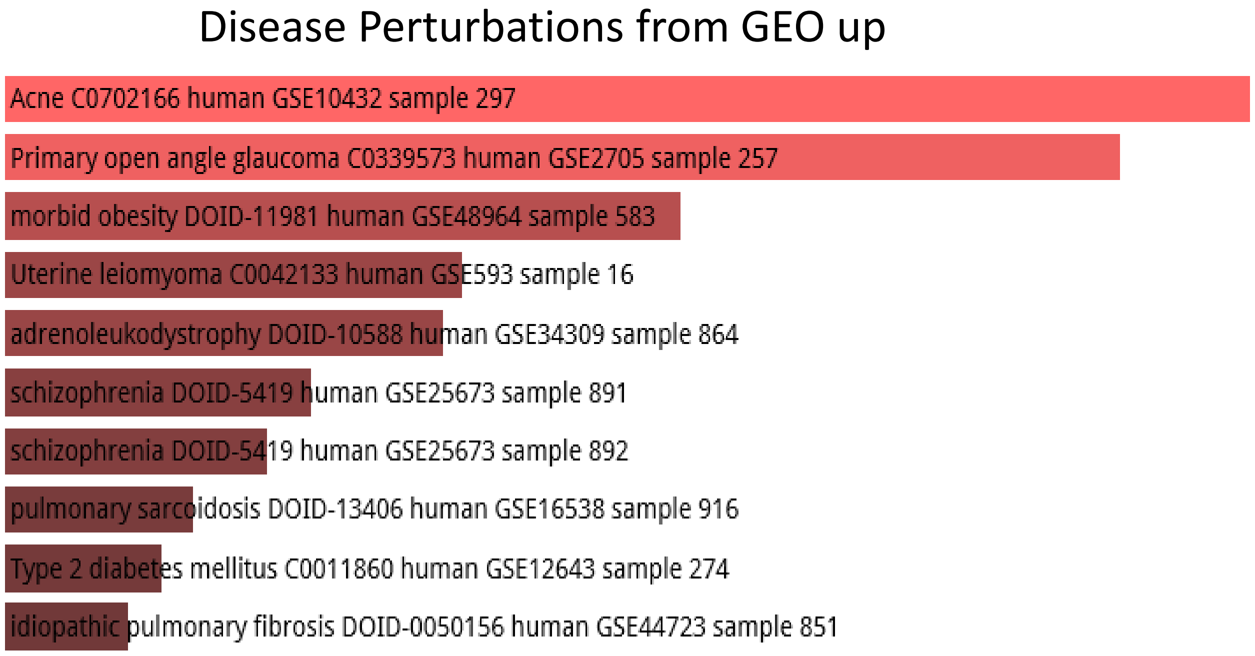


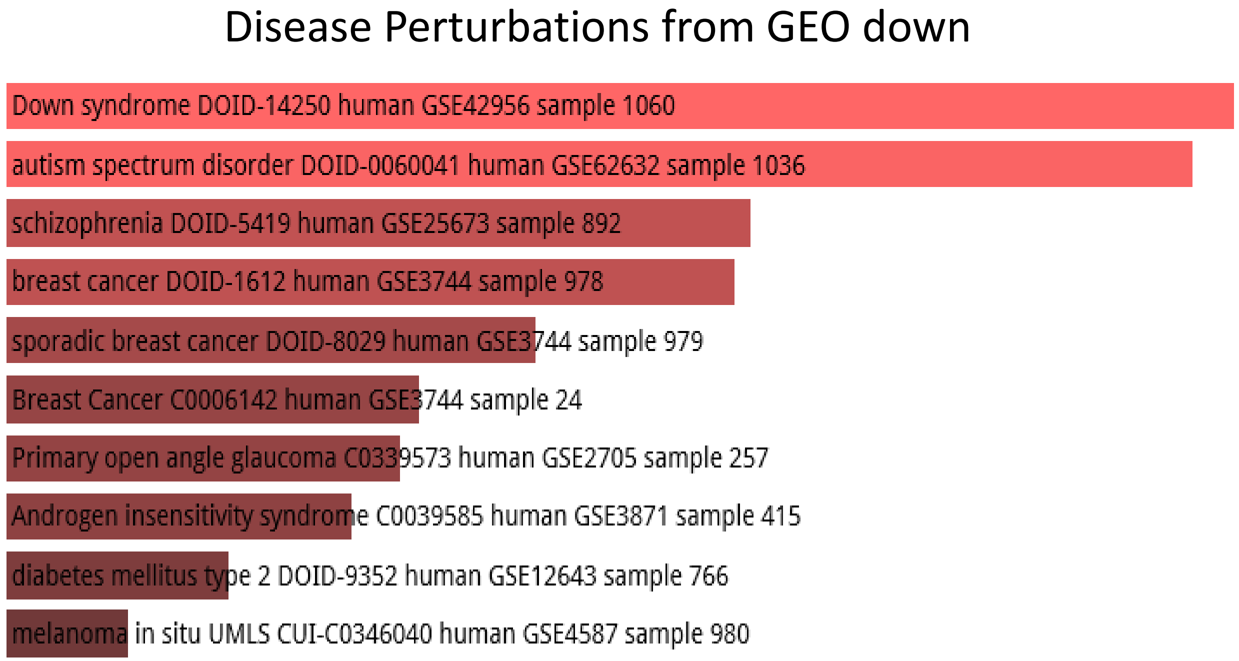

**1,619 human-specific functional hESC enhancers**

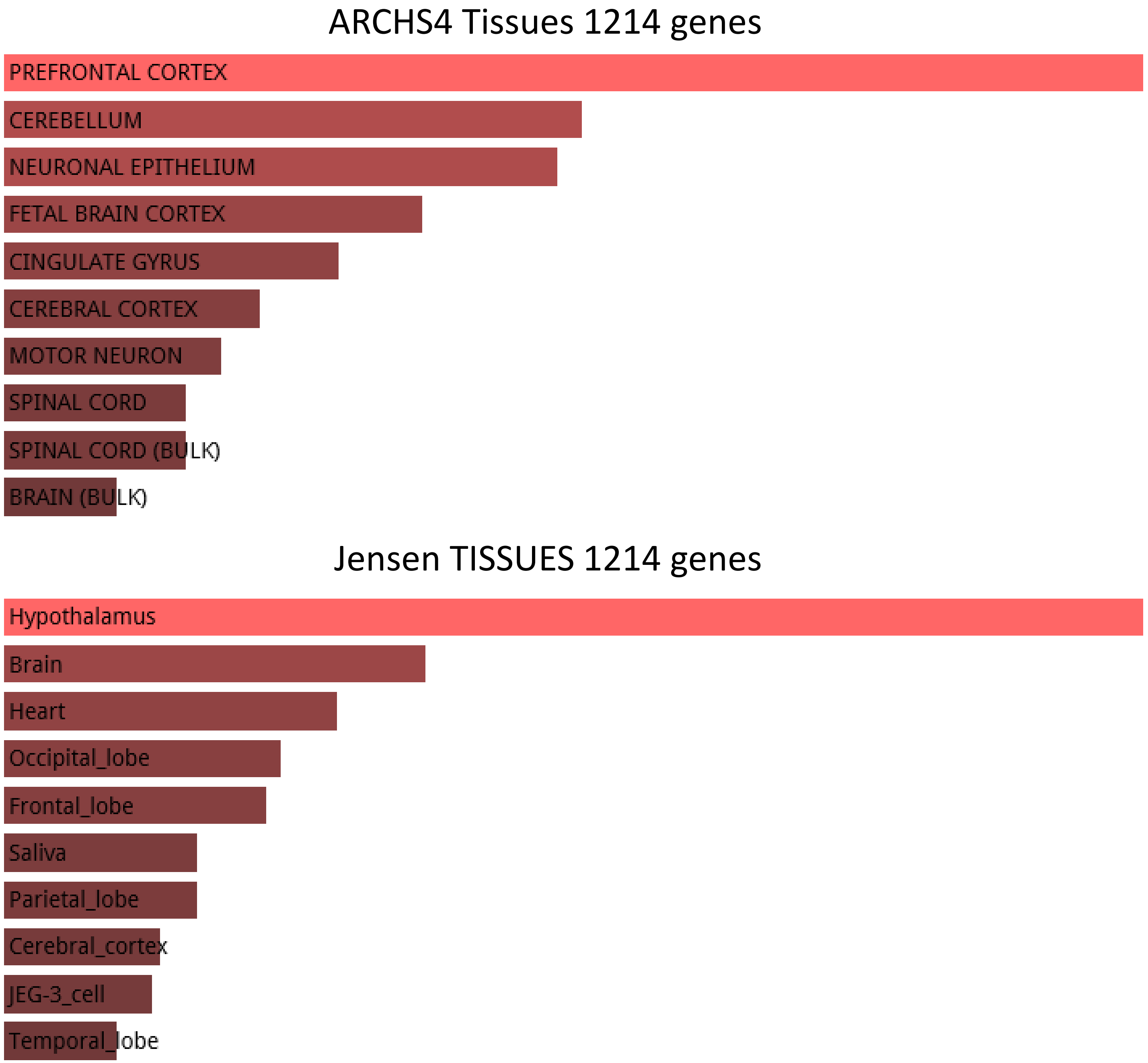

**1,619 human-specific functional hESC enhancers**

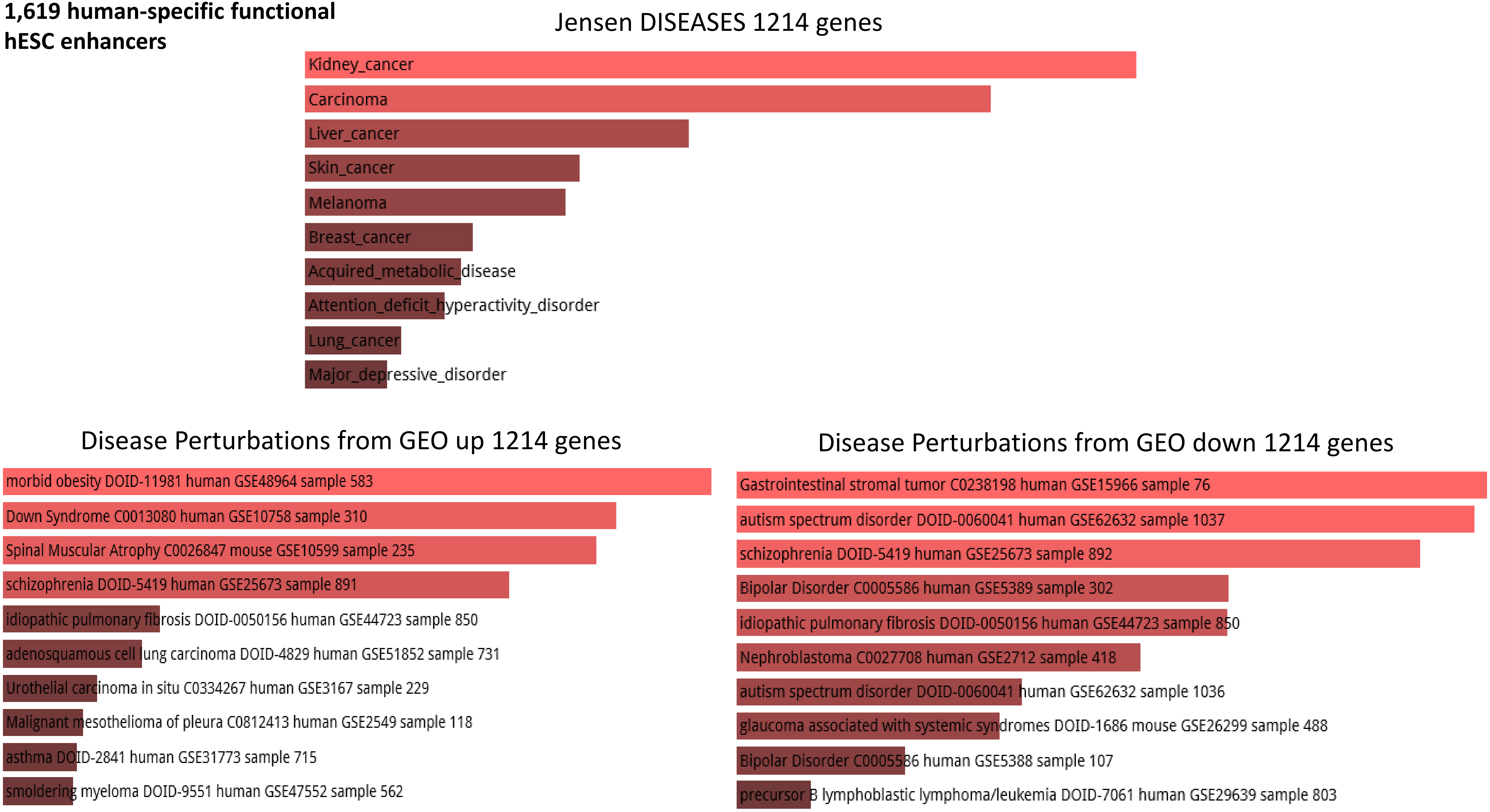

**1,619 human-specific functional hESC enhancers**

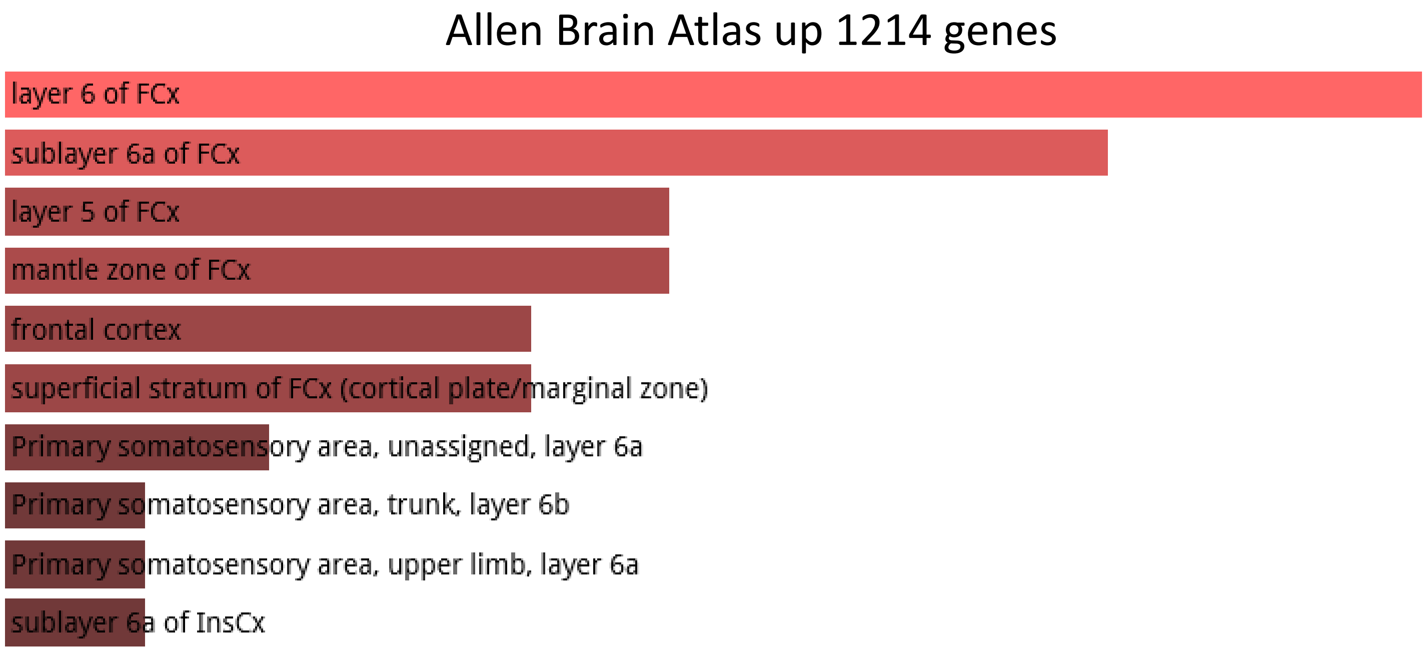


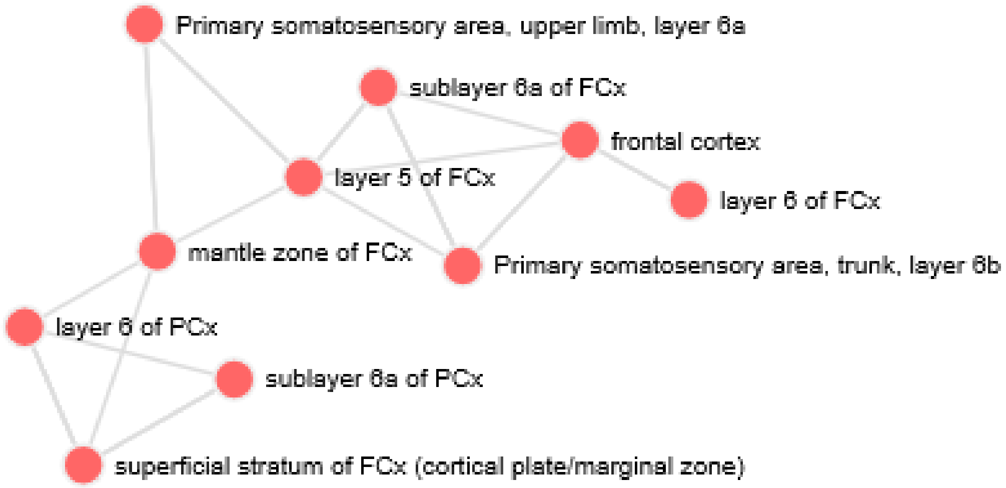

**524 human-specific DHS**

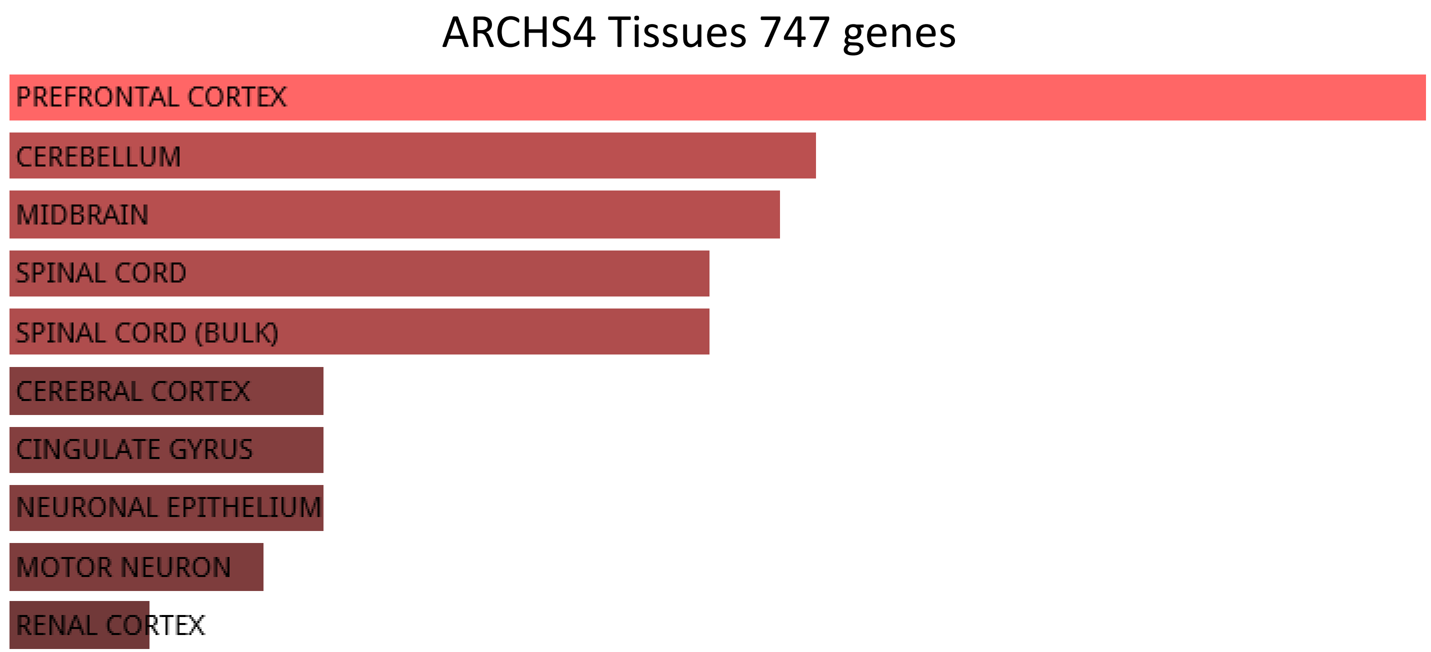


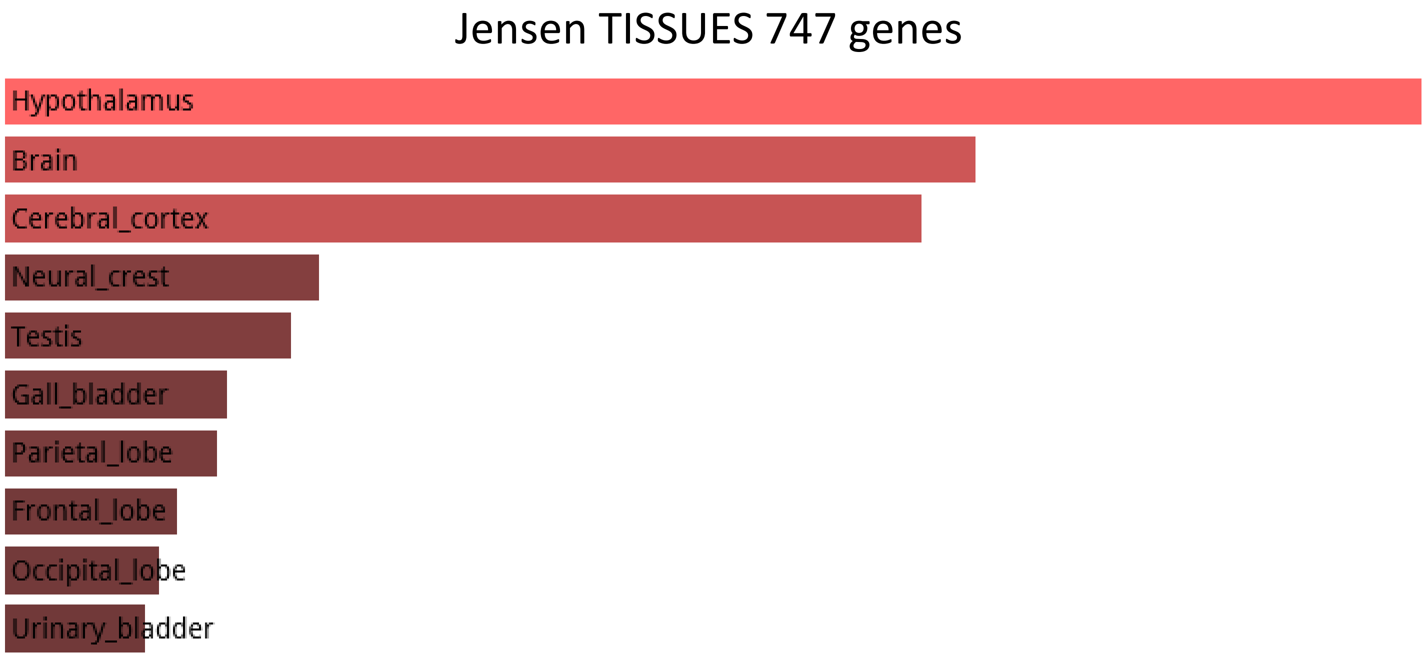

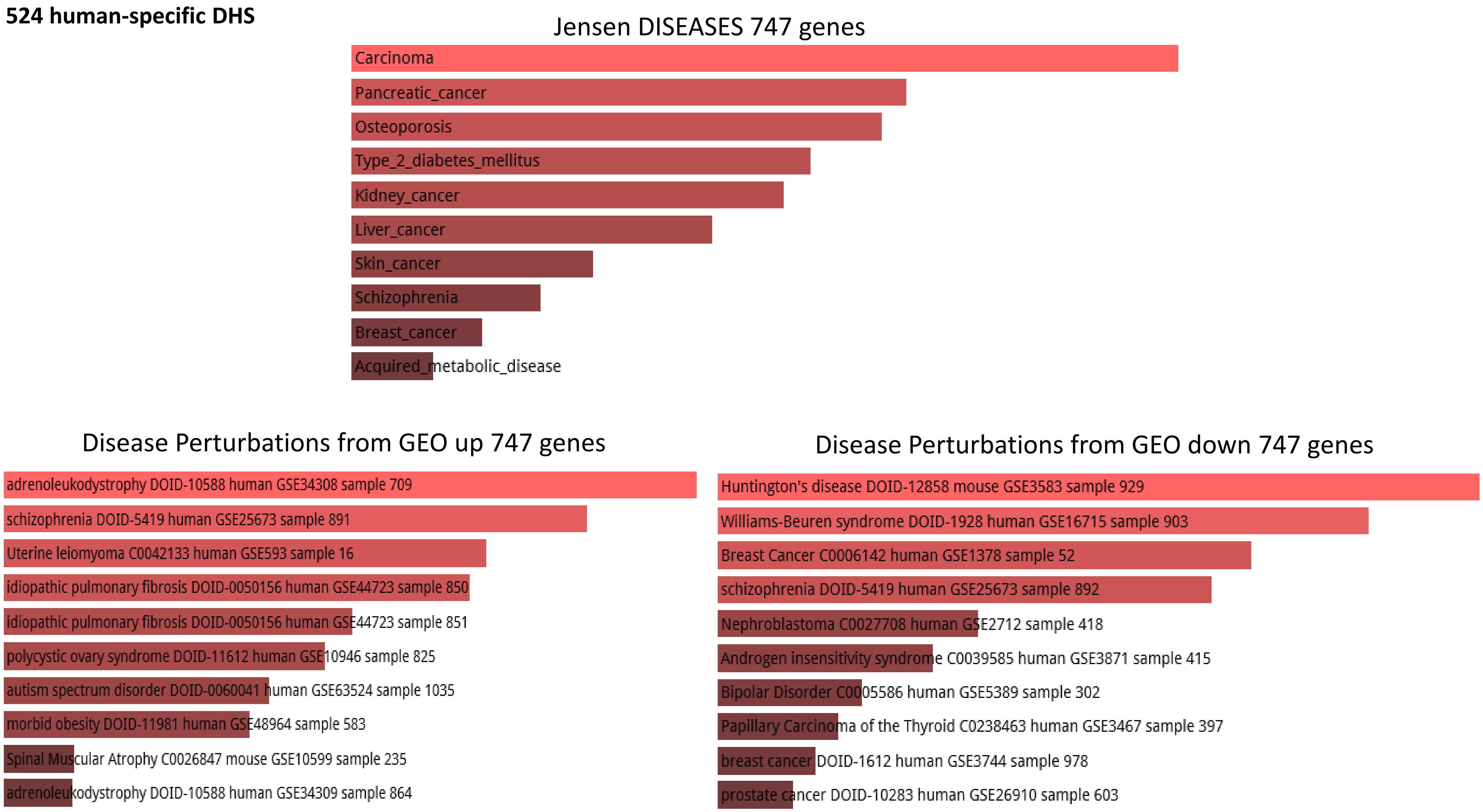
524 human-specific DHS
Jensen DISEASES 747 genes
Carcinoma
Pancreatic_cancer
Osteoporosis
Type_2_diabetes_mellitus
Kidney_cancer
Liver_cancer
Skin_cancer
Schizophrenia
Breast_cancer
Acquired_metabolic_disease
Disease Perturbations from GEO up 747 genes
adrenoleukodystrophy DOID-10588 human GSE34308 sample 709
schizophrenia DOID-5419 human GSE25673 sample 891
Uterine leiomyoma C0042133 human GSE593 sample 16
idiopathic pulmonary fibrosis DOID-0050156 human GSE44723 sample 850
idiopathic pulmonary fibrosis DOID-0050156 human GSE44723 sample 851
polycystic ovary syndrome DOID-11612 human GSE10946 sample 825
autism spectrum disorder DOID-0060041 human GSE63524 sample 1035
morbid obesity DOID-11981 human GSE48964 sample 583
Spinal Muscular Atrophy C0026847 mouse GSE10599 sample 235
adrenoleukodystrophy DOID-10588 human GSE34309 sample 864
Disease Perturbations from GEO down 747 genes
Huntington's disease DOID-12858 mouse GSE3583 sample 929
Williams-Beuren syndrome DOID-1928 human GSE16715 sample 903
Breast Cancer C0006142 human GSE1378 sample 52
schizophrenia DOID-5419 human GSE25673 sample 892
Nephroblastoma C0027708 human GSE2712 sample 418
Androgen insensitivity syndrome C0039585 human GSE3871 sample 415
Bipolar Disorder C0005586 human GSE5389 sample 302
Papillary Carcinoma of the Thyroid C0238463 human GSE3467 sample 397
breast cancer DOID-1612 human GSE3744 sample 978
prostate cancer DOID-10283 human GSE26910 sample 603

**42,847 human genes not linked by the GREAT algorithm with HSRS were randomly split into 21 control gene sets of various sizes ranging from 2,847 to 6,847 genes and subjected to the GSEA**

### GSEA OF TWENTY-ONE CONTROL GENE SETS NOT ASSOCIATED WITH HUMAN-SPECIFIC GENOMIC REGULATORY LOCI

**Random Set 1**

ARCHS4 Tissues random 6000 genes

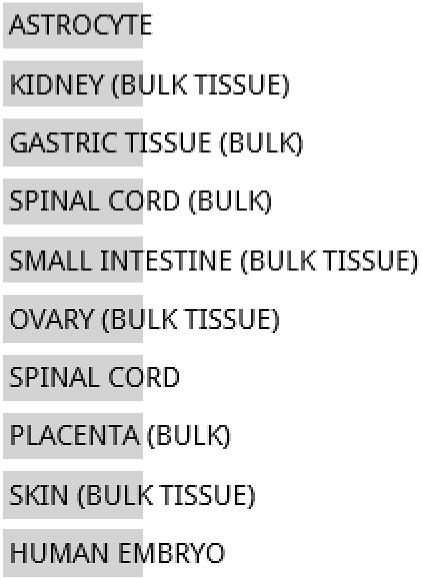


**NO SIGNIFICANT ENRICHMENT**

Jensen TISSUES random 6847 genes

**NO DATA AVAILABLE**

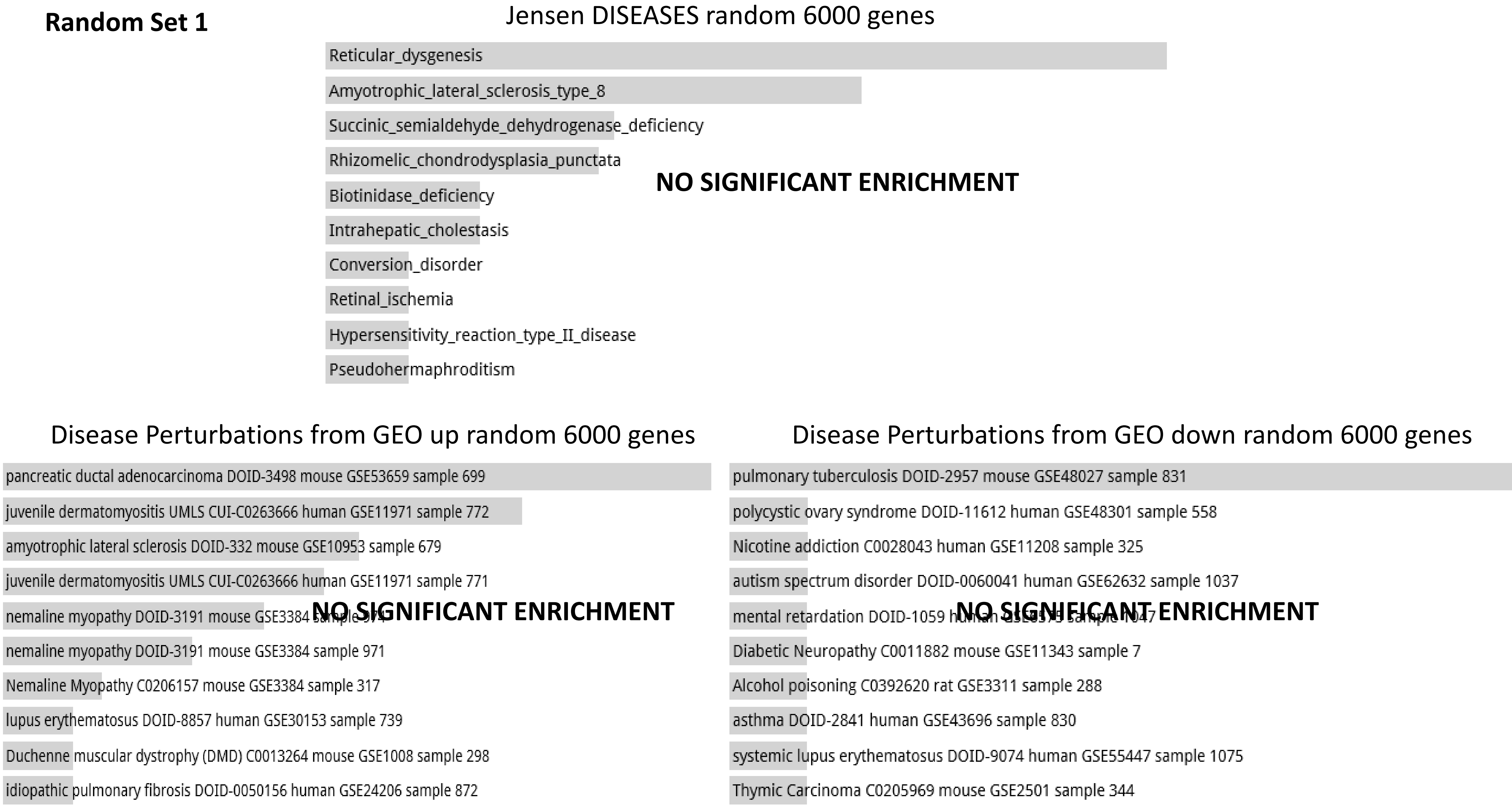

Random Set 1
Jensen DISEASES random 6000 genes
Reticular_dysgenesis
Amyotrophic_lateral_sclerosis_type_8
Succinic_semialdehyde_dehydrogenase_deficiency
Rhizomelic_chondrodysplasia_punctata
Biotinidase_deficiency
Intrahepatic_cholestasis
Conversion_disorder
Retinal_ischemia
Hypersensitivity_reaction_type_II_disease
Pseudohermaphroditism
NO SIGNIFICANT ENRICHMENT
Disease Perturbations from GEO up random 6000 genes
pancreatic ductal adenocarcinoma DOID-3498 mouse GSE53659 sample 699
juvenile dermatomyositis UMLS CUI-C0263666 human GSE11971 sample 772
amyotrophic lateral sclerosis DOID-332 mouse GSE10953 sample 679
juvenile dermatomyositis UMLS CUI-C0263666 human GSE11971 sample 771
nemaline myopathy DOID-3191 mouse GSE3384
nemaline myopathy DOID-3191 mouse GSE3384 sample 971
Nemaline Myopathy C0206157 mouse GSE3384 sample 317
lupus erythematosus DOID-8857 human GSE30153 sample 739
Duchenne muscular dystrophy (DMD) C0013264 mouse GSE1008 sample 298
idiopathic pulmonary fibrosis DOID-0050156 human GSE24206 sample 872
NO SIGNIFICANT ENRICHMENT
Disease Perturbations from GEO down random 6000 genes
pulmonary tuberculosis DOID-2957 mouse GSE48027 sample 831
polycystic ovary syndrome DOID-11612 human GSE48301 sample 558
Nicotine addiction C0028043 human GSE11208 sample 325
autism spectrum disorder DOID-0060041 human GSE62632 sample 1037
mental retardation DOID-1059
Diabetic Neuropathy C0011882 mouse GSE11343 sample 7
Alcohol poisoning C0392620 rat GSE3311 sample 288
asthma DOID-2841 human GSE43696 sample 830
systemic lupus erythematosus DOID-9074 human GSE55447 sample 1075
Thymic Carcinoma C0205969 mouse GSE2501 sample 344
NO SIGNIFICANT ENRICHMENT

**Random Set 1**

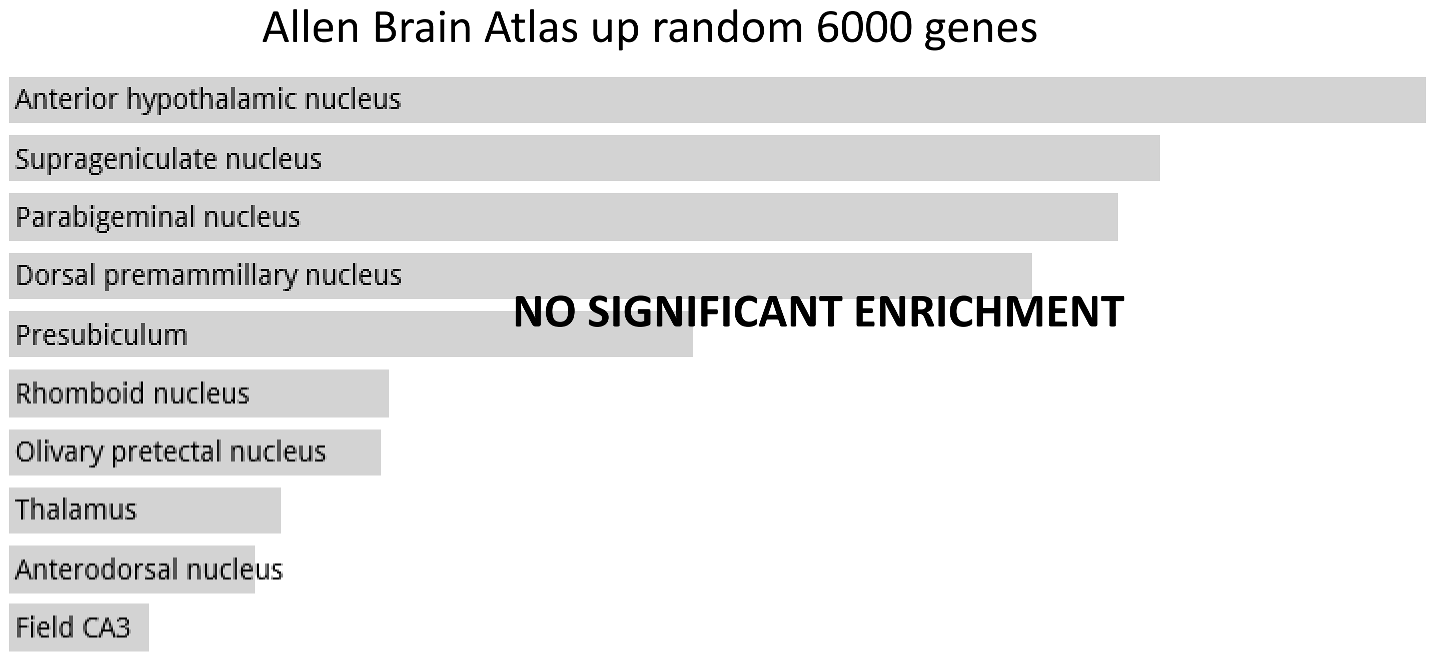
Allen Brain Atlas up random 6000 genes
Anterior hypothalamic nucleus
Suprageniculate nucleus
Parabigeminal nucleus
Dorsal premammillary nucleus
Presubiculum
Rhomboid nucleus
Olivary pretectal nucleus
Thalamus
Anterodorsal nucleus
Field CA3
NO SIGNIFICANT ENRICHMENT

**Random Set 2**

ARCHS4 Tissues random 6000 genes

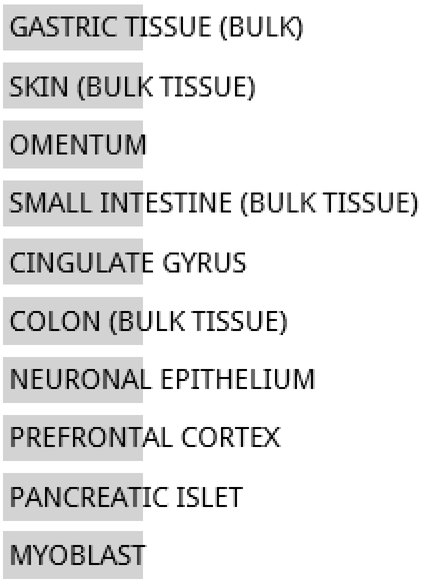


**NO SIGNIFICANT ENRICHMENT**

Jensen TISSUES random 6847 genes

**NO DATA AVAILABLE**

**Random Set 2**

#### Jensen DISEASES random 6000 genes

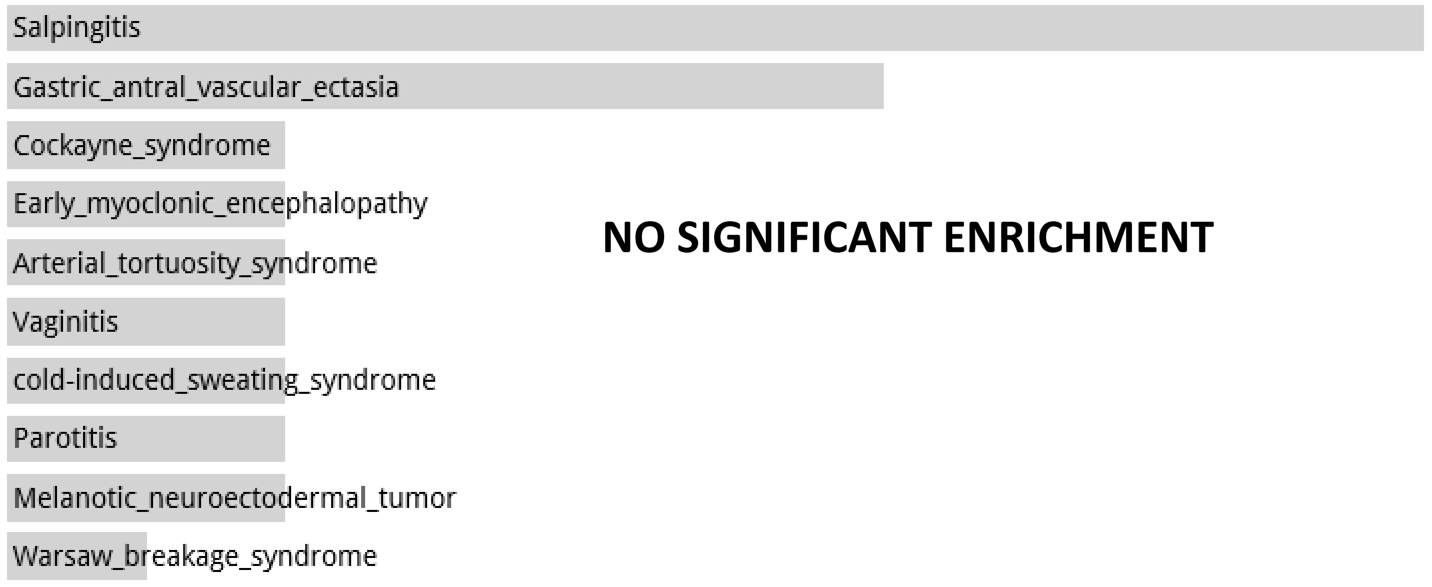


#### Disease Perturbations from GEO up random 6000 genes

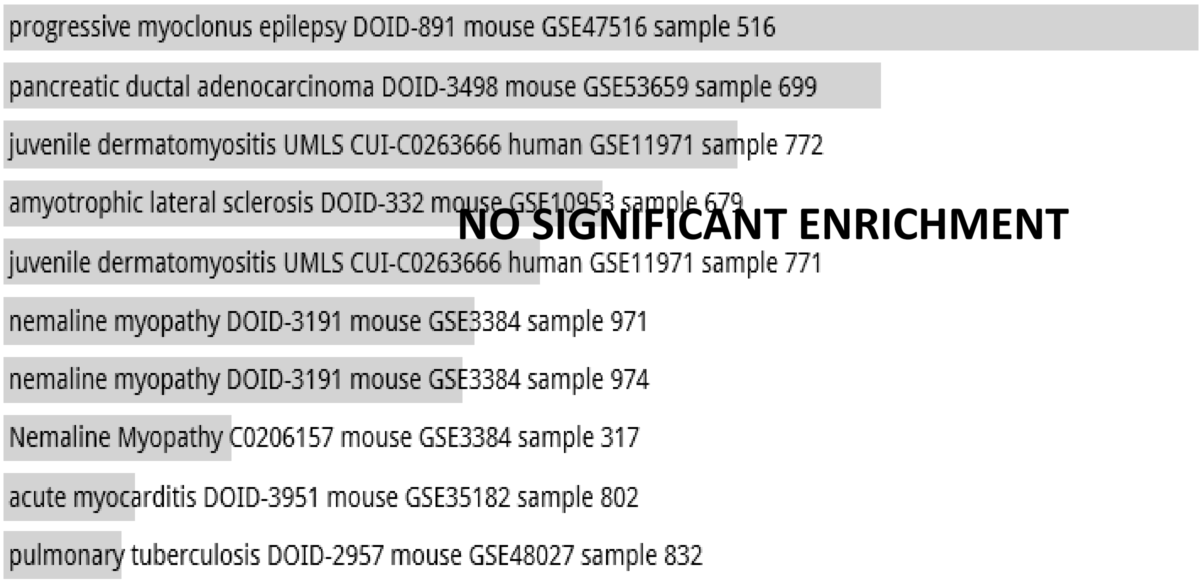


#### Disease Perturbations from GEO down random 6000 genes

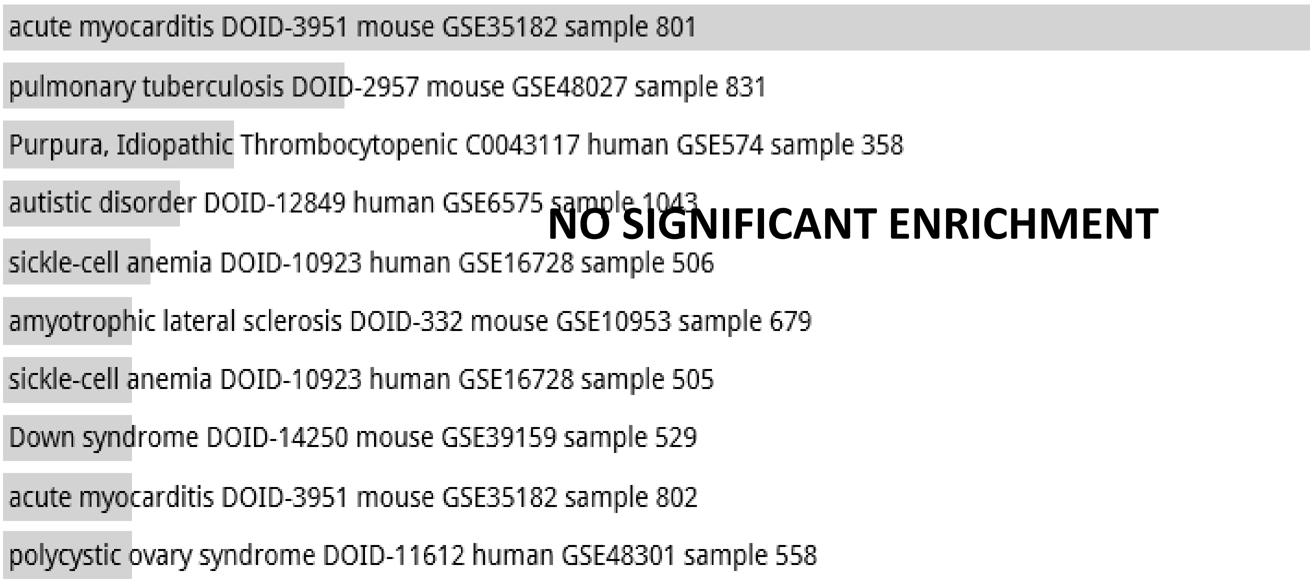

**Random Set 2**

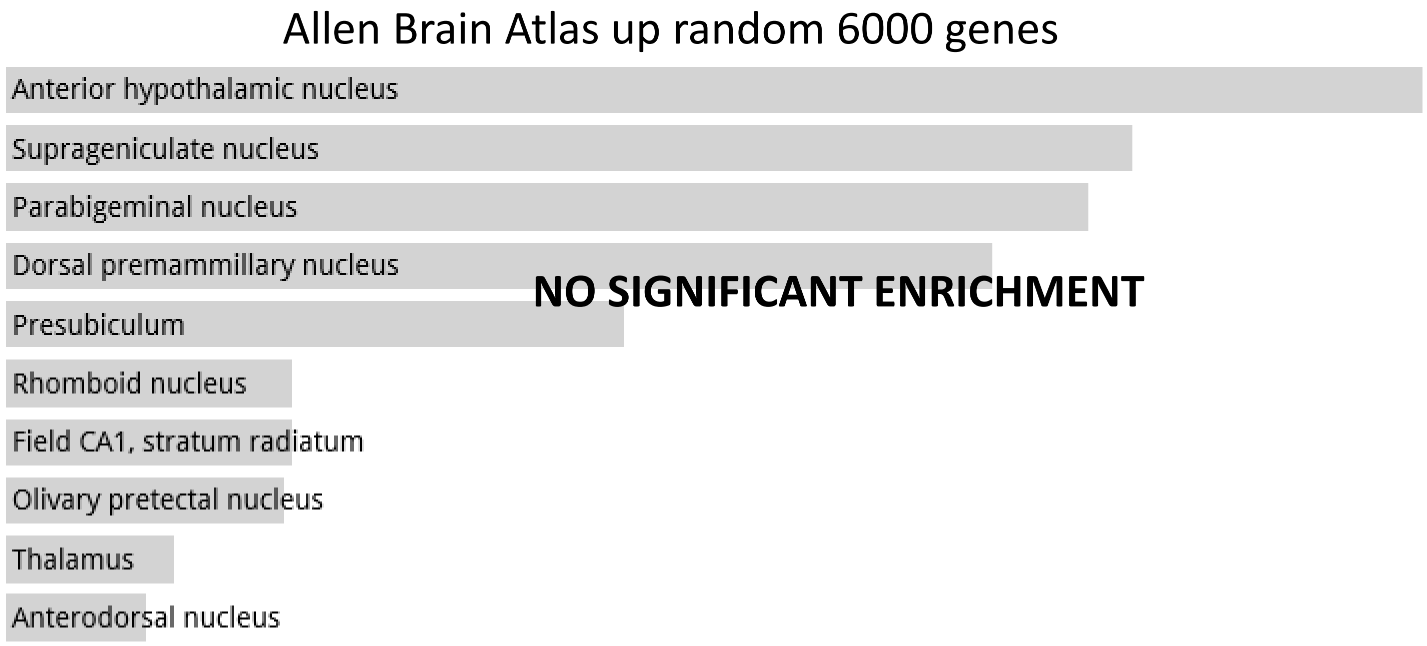

Allen Brain Atlas up random 6000 genes
Anterior hypothalamic nucleus
Suprageniculate nucleus
Parabigeminal nucleus
Dorsal premammillary nucleus
Presubiculum
Rhomboid nucleus
Field CA1, stratum radiatum
Olivary pretectal nucleus
Thalamus
Anterodorsal nucleus
NO SIGNIFICANT ENRICHMENT

**Random Set 3**

#### ARCHS4 Tissues random 6000 genes

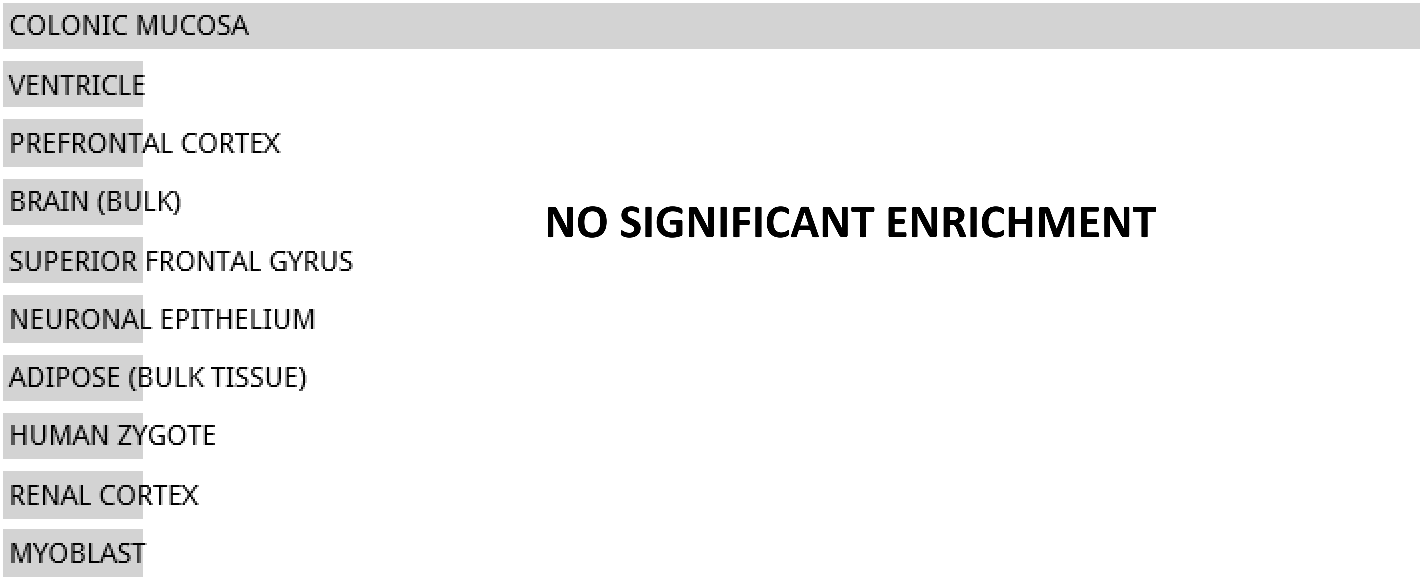


**NO SIGNIFICANT ENRICHMENT**

#### Jensen TISSUES random 6847 genes

**NO DATA AVAILABLE**

**Random Set 3**

Jensen DISEASES random 6000 genes

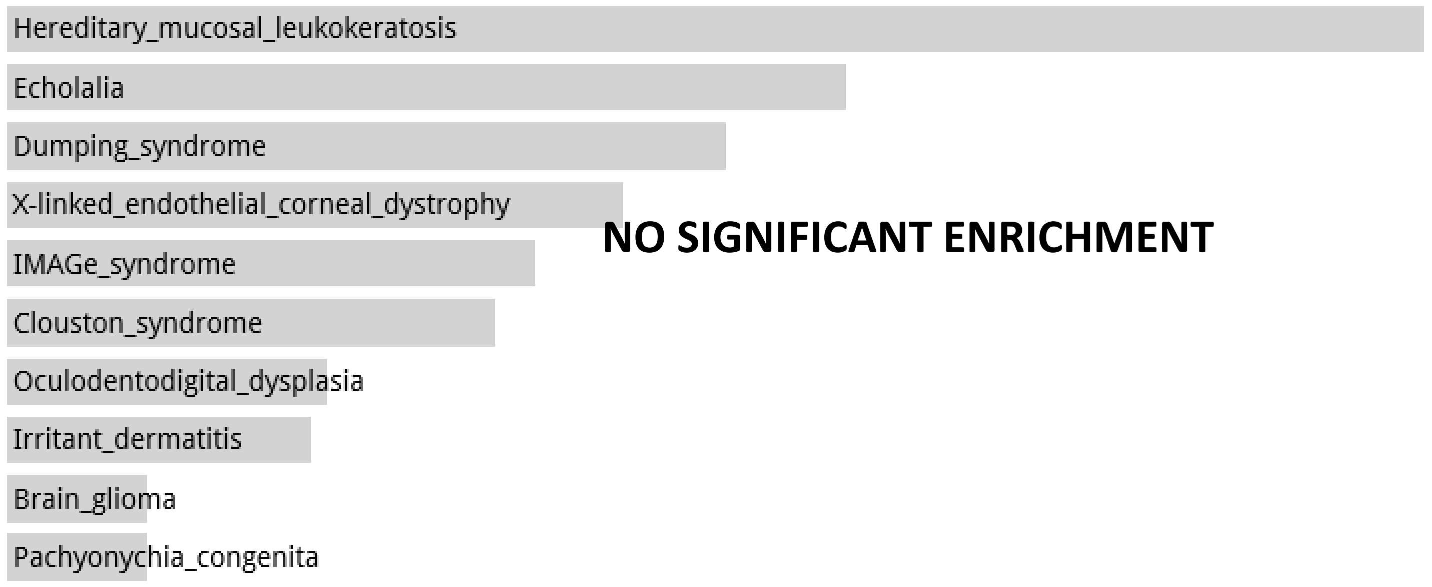


Disease Perturbations from GEO up random 6000 genes

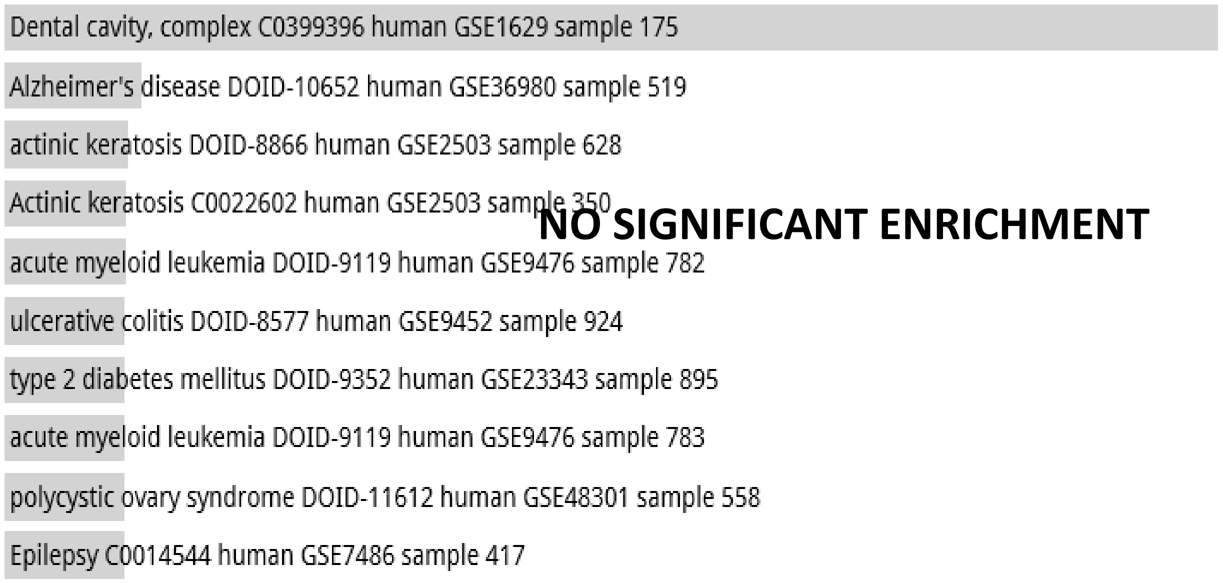


Disease Perturbations from GEO down random 6000 genes

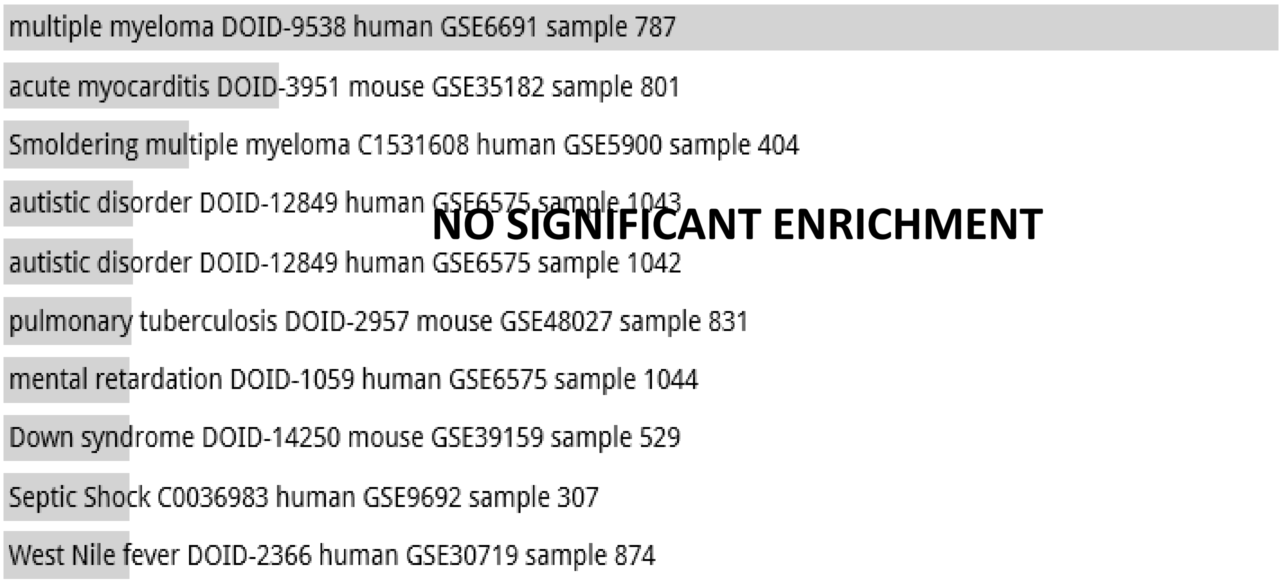

**Random Set 3**

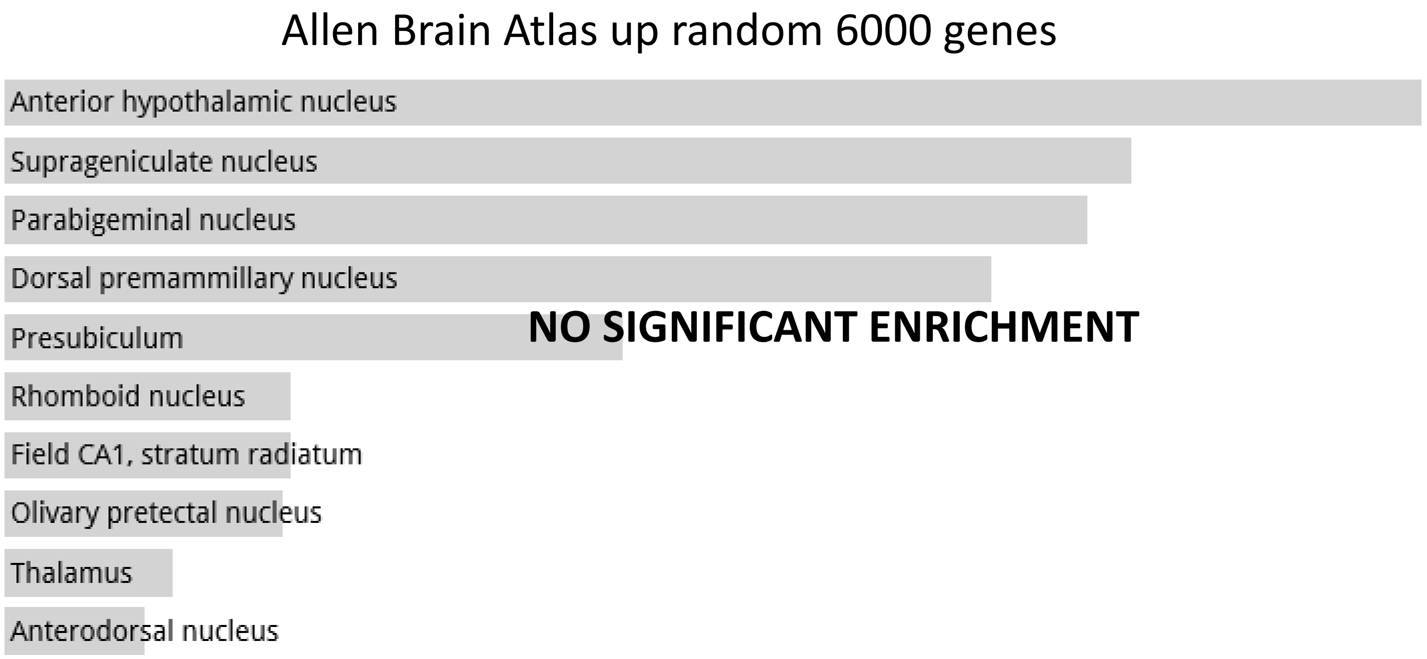

Allen Brain Atlas up random 6000 genes
Anterior hypothalamic nucleus
Suprageniculate nucleus
Parabigeminal nucleus
Dorsal premammillary nucleus
Presubiculum
Rhomboid nucleus
Field CA1, stratum radiatum
Olivary pretectal nucleus
Thalamus
Anterodorsal nucleus
NO SIGNIFICANT ENRICHMENT

**Random Set 4**

ARCHS4 Tissues random 6000 genes

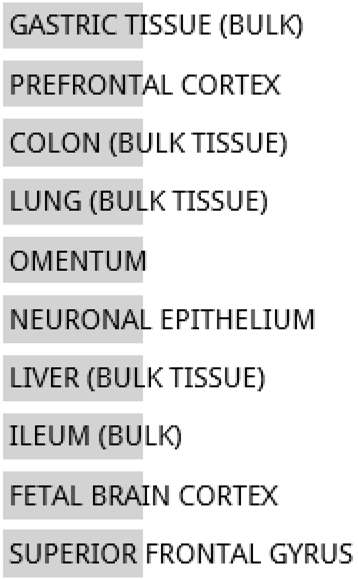


**NO SIGNIFICANT ENRICHMENT**

Jensen TISSUES random 6847 genes

**NO DATA AVAILABLE**

**Random Set 4**

#### Jensen DISEASES random 6000 genes

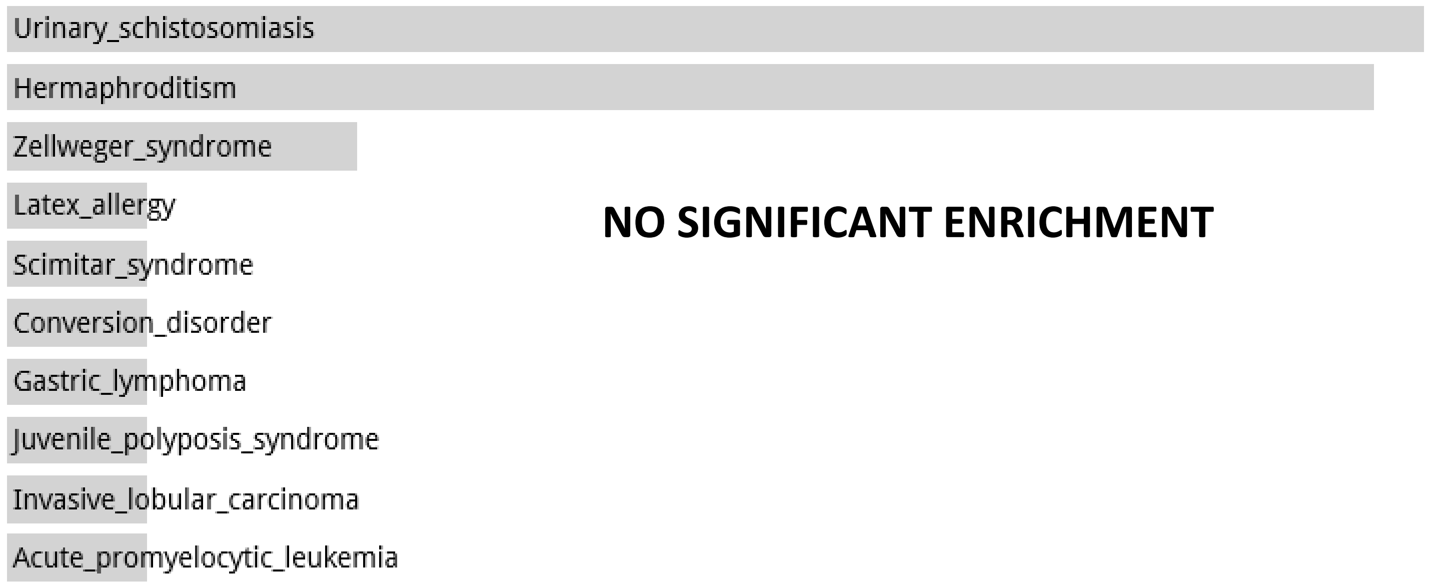


#### Disease Perturbations from GEO up random 6000 genes

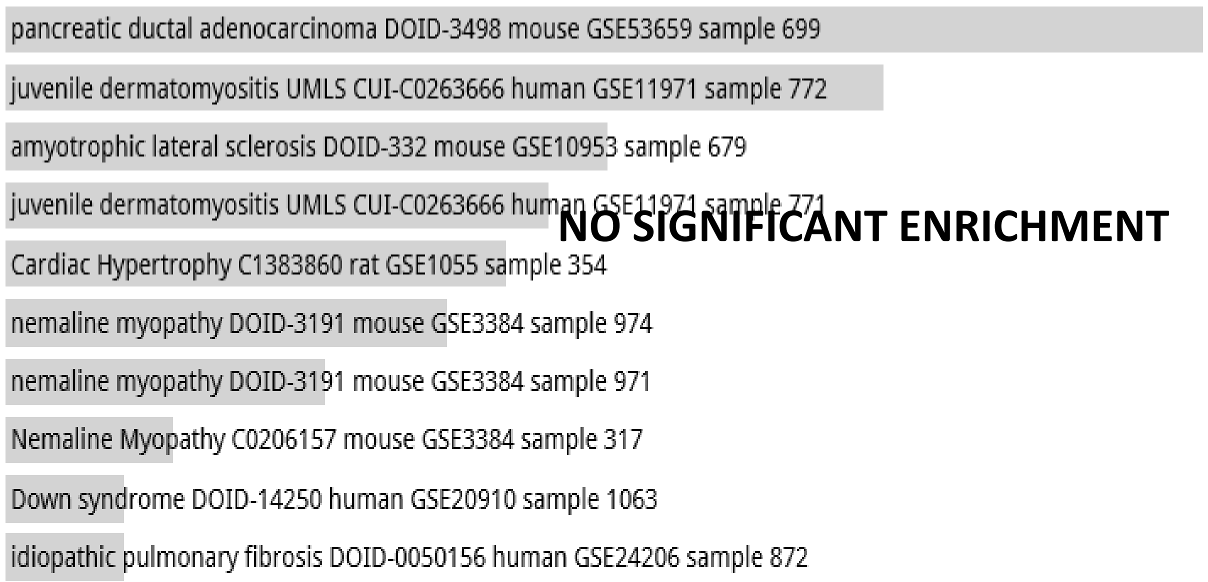


#### Disease Perturbations from GEO down random 6000 genes

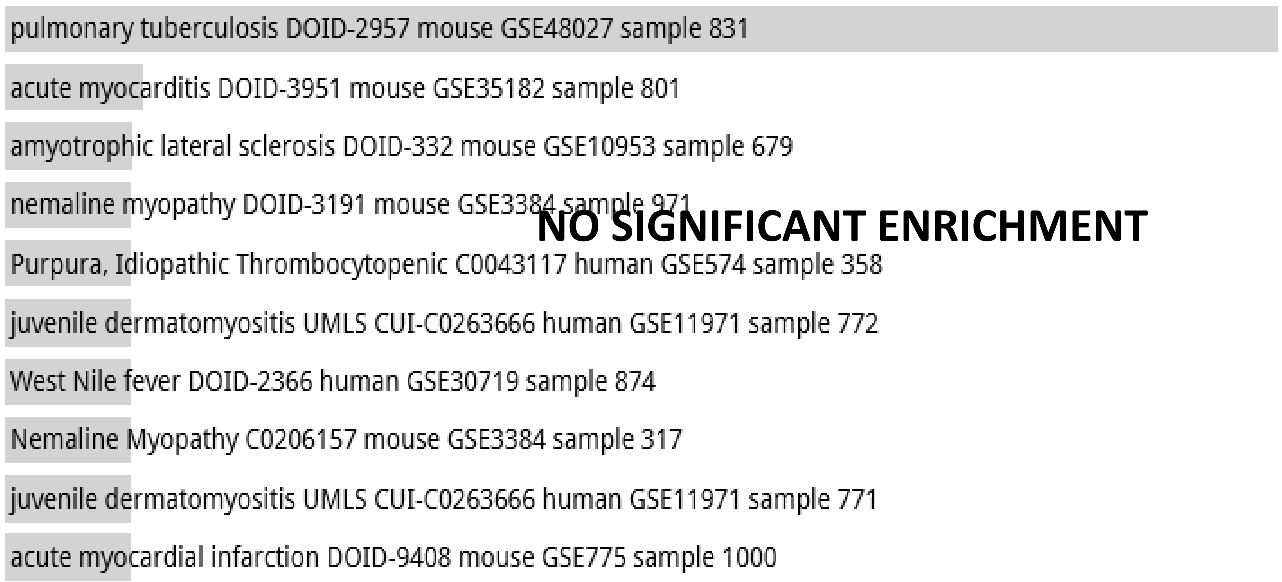

**Random Set 4**

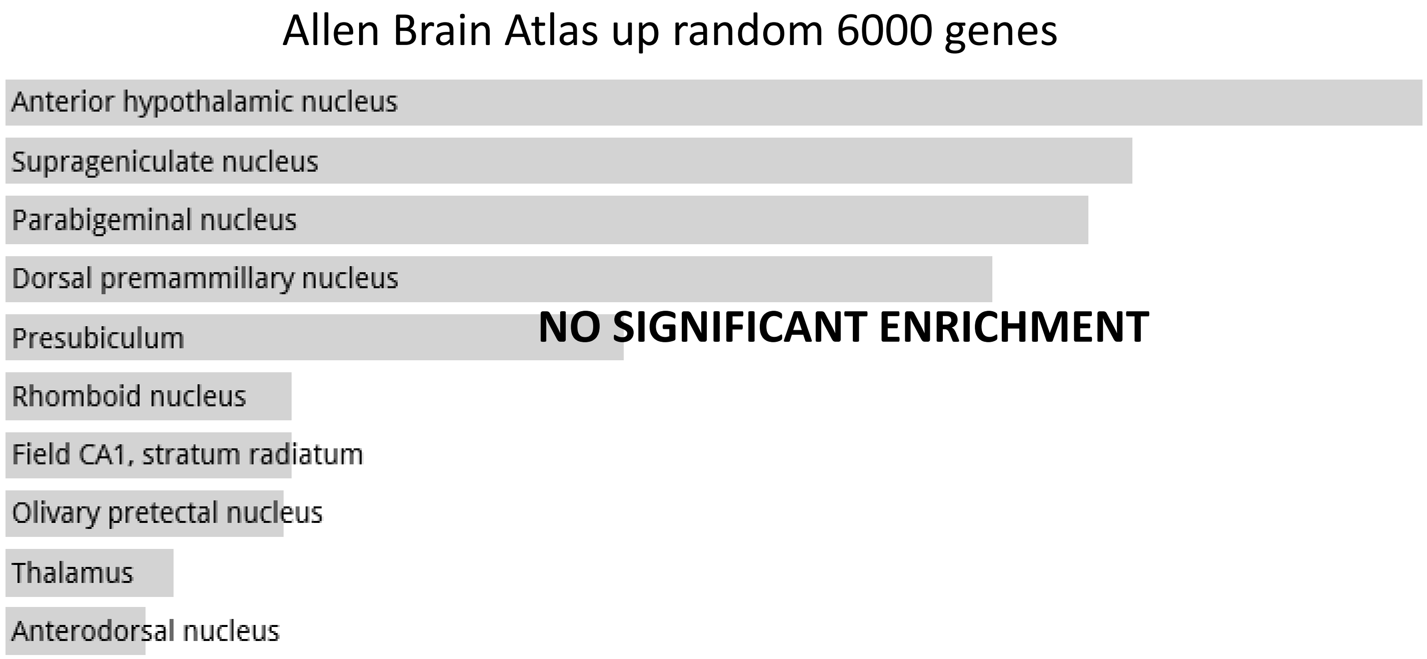
Allen Brain Atlas up random 6000 genes
Anterior hypothalamic nucleus
Suprageniculate nucleus
Parabigeminal nucleus
Dorsal premammillary nucleus
Presubiculum
Rhomboid nucleus
Field CA1, stratum radiatum
Olivary pretectal nucleus
Thalamus
Anterodorsal nucleus
NO SIGNIFICANT ENRICHMENT

**Random Set 5**

#### ARCHS4 Tissues random 6000 genes

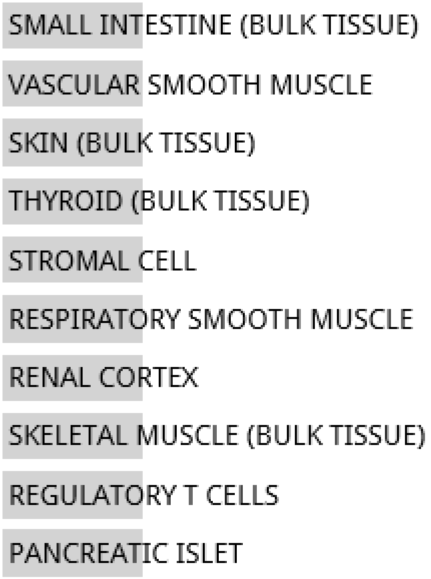


**NO SIGNIFICANT ENRICHMENT**

#### Jensen TISSUES random 6847 genes

**NO DATA AVAILABLE**

**Random Set 5**

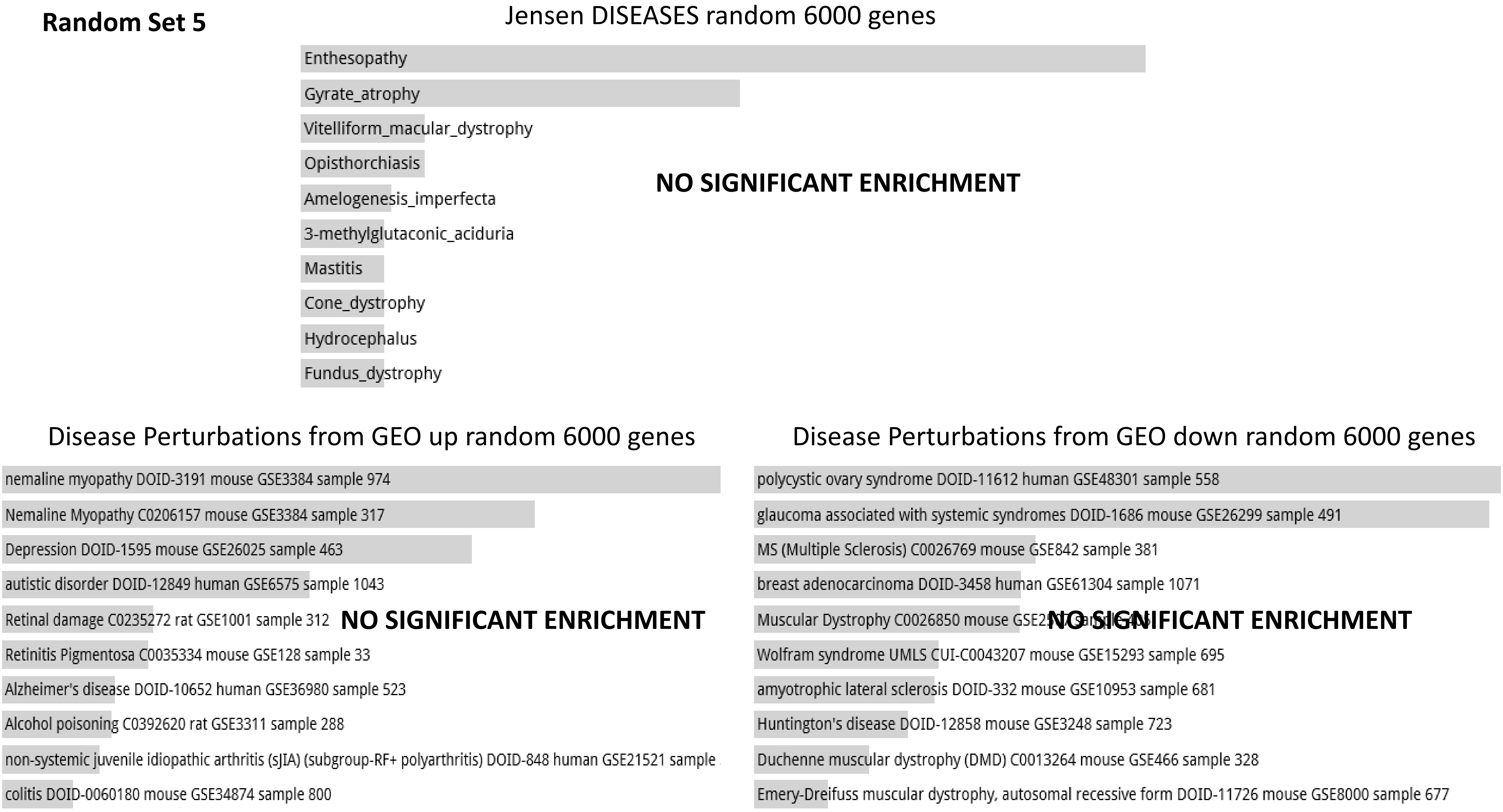

Jensen DISEASES random 6000 genes
Enthesopathy
Gyrate_atrophy
Vitelliform_macular_dystrophy
Opisthorchiasis
Amelogenesis_imperfecta
3-methylglutaconic_aciduria
Mastitis
Cone_dystrophy
Hydrocephalus
Fundus_dystrophy
NO SIGNIFICANT ENRICHMENT
Disease Perturbations from GEO up random 6000 genes
nemaline myopathy DOID-3191 mouse GSE3384 sample 974
Nemaline Myopathy C0206157 mouse GSE3384 sample 317
Depression DOID-1595 mouse GSE26025 sample 463
autistic disorder DOID-12849 human GSE6575 sample 1043
Retinal damage C0235272 rat GSE1001 sample 312
Retinitis Pigmentosa C0035334 mouse GSE128 sample 33
Alzheimer's disease DOID-10652 human GSE36980 sample 523
Alcohol poisoning C0392620 rat GSE3311 sample 288
non-systemic juvenile idiopathic arthritis (sJIA) (subgroup-RF+ polyarthritis) DOID-848 human GSE21521 sample
colitis DOID-0060180 mouse GSE34874 sample 800
NO SIGNIFICANT ENRICHMENT
Disease Perturbations from GEO down random 6000 genes
polycystic ovary syndrome DOID-11612 human GSE48301 sample 558
glaucoma associated with systemic syndromes DOID-1686 mouse GSE26299 sample 491
MS (Multiple Sclerosis) C0026769 mouse GSE842 sample 381
breast adenocarcinoma DOID-3458 human GSE61304 sample 1071
Muscular Dystrophy C0026850 mouse GSE2
Wolfram syndrome UMLS CUI-C0043207 mouse GSE15293 sample 695
amyotrophic lateral sclerosis DOID-332 mouse GSE10953 sample 681
Huntington's disease DOID-12858 mouse GSE3248 sample 723
Duchenne muscular dystrophy (DMD) C0013264 mouse GSE466 sample 328
Emery-Dreifuss muscular dystrophy, autosomal recessive form DOID-11726 mouse GSE8000 sample 677
NO SIGNIFICANT ENRICHMENT

#### Random Set 5

Allen Brain Atlas up random 6000 genes

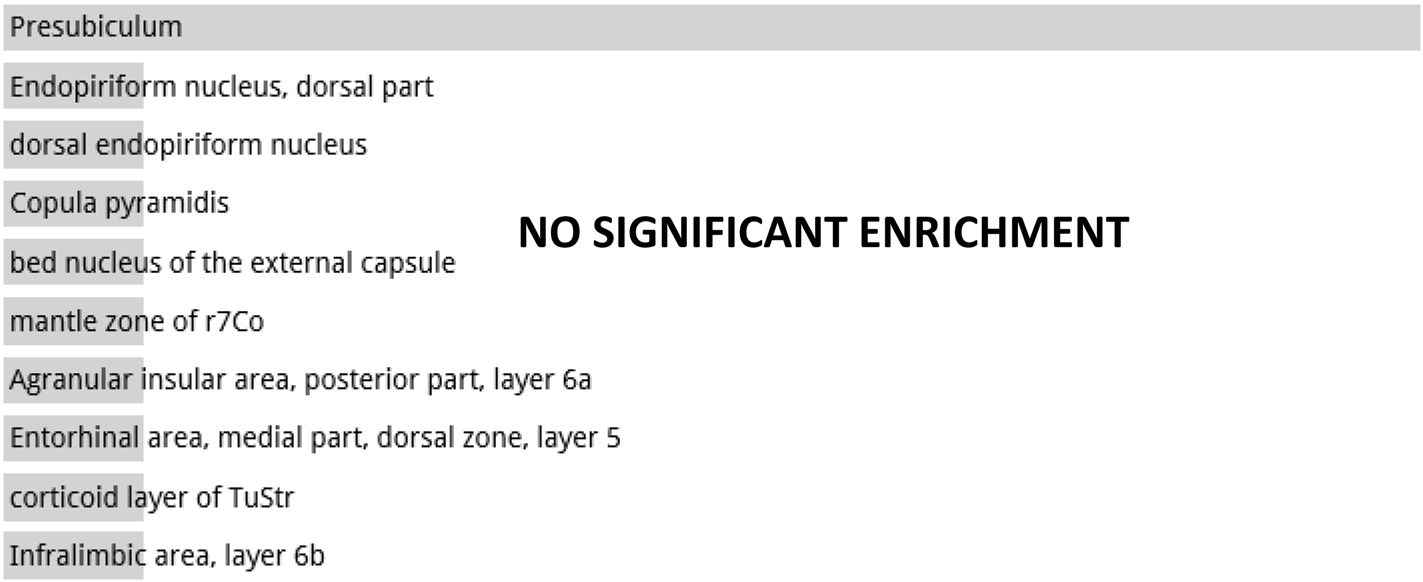


**NO SIGNIFICANT ENRICHMENT**

**Random Set 6**

ARCHS4 Tissues random 6000 genes

**NO DATA AVAILABLE**

Jensen TISSUES random 6000 genes

**NO DATA AVAILABLE**

**Random Set 6**

Jensen DISEASES random 6000 genes

**NO DATA AVAILABLE**

Disease Perturbations from GEO up random 6000 genes

**NO DATA AVAILABLE**

Disease Perturbations from GEO down random 6000 genes

Systemic lupus erythematosus (SLE) C0024141 human GSE12374 sample 123

**NO SIGNIFICANT ENRICHMENT**

**Random Set 6**

Allen Brain Atlas up random 6000 genes

**NO DATA AVAILABLE**

**Random Set 7**

ARCHS4 Tissues random 6847 genes

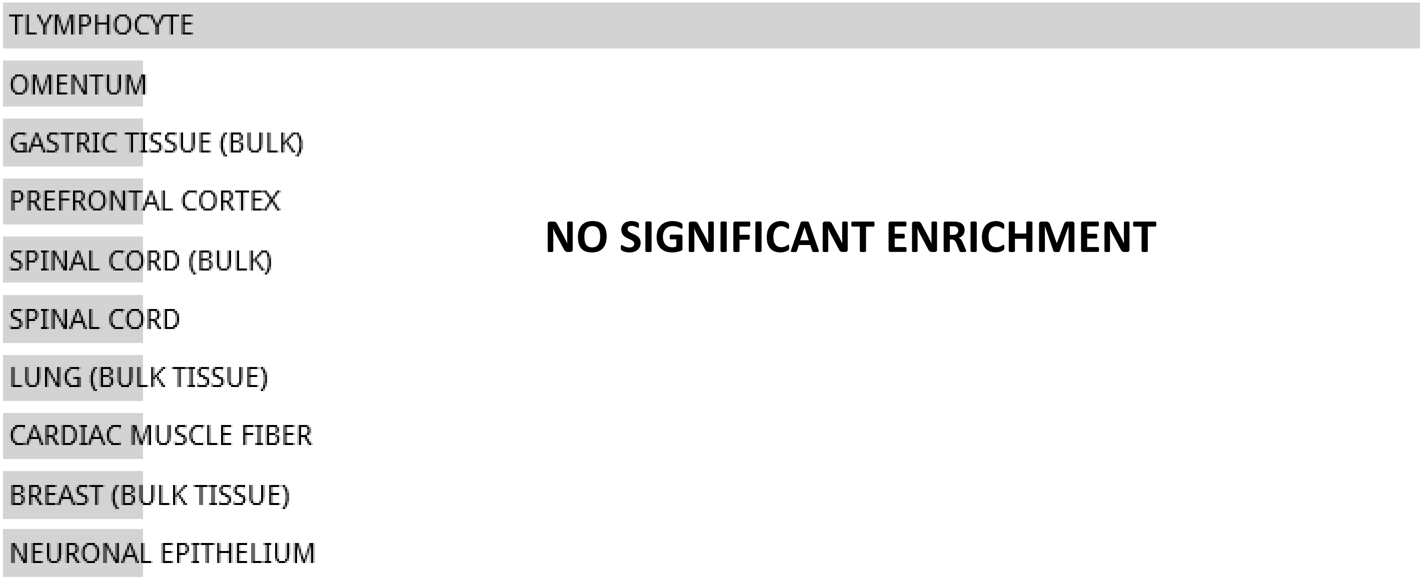


**NO SIGNIFICANT ENRICHMENT**

Jensen TISSUES random 6847 genes

**NO DATA AVAILABLE**

**Random Set 7**

Jensen DISEASES random 6847 genes

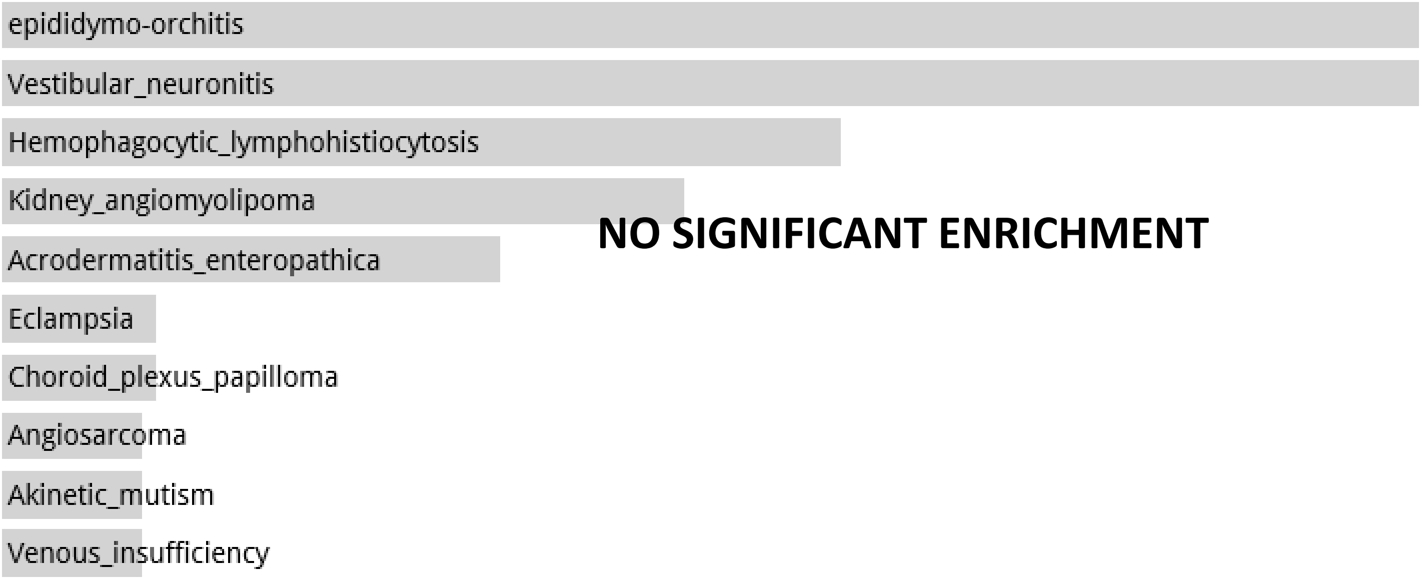


Disease Perturbations from GEO up random 6000 genes

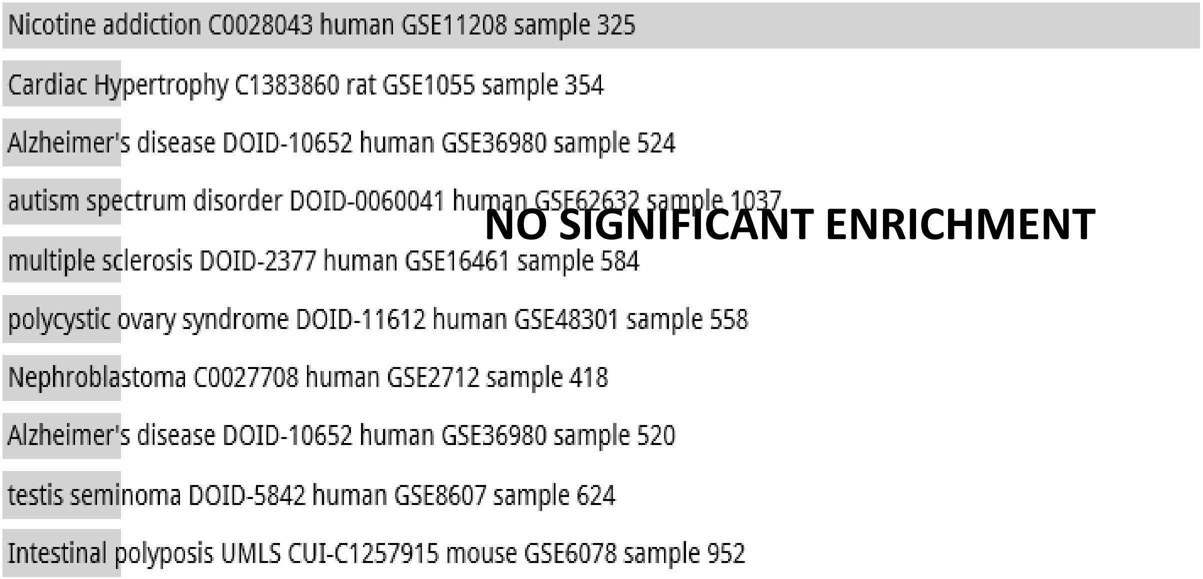


Disease Perturbations from GEO down random 6000 genes

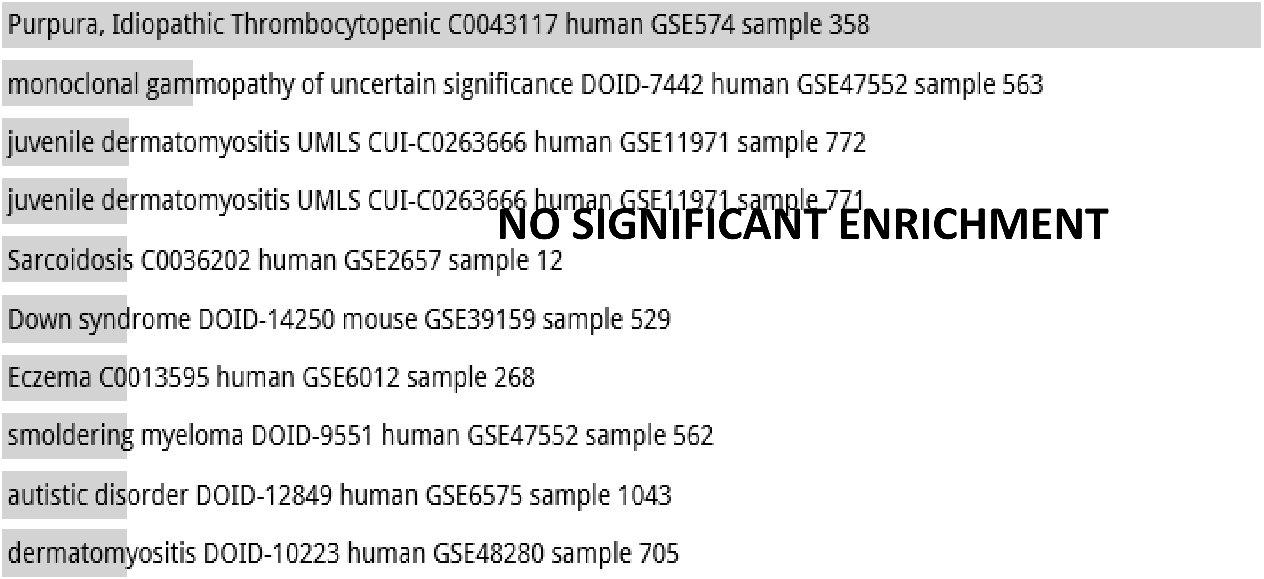

**Random Set 7**

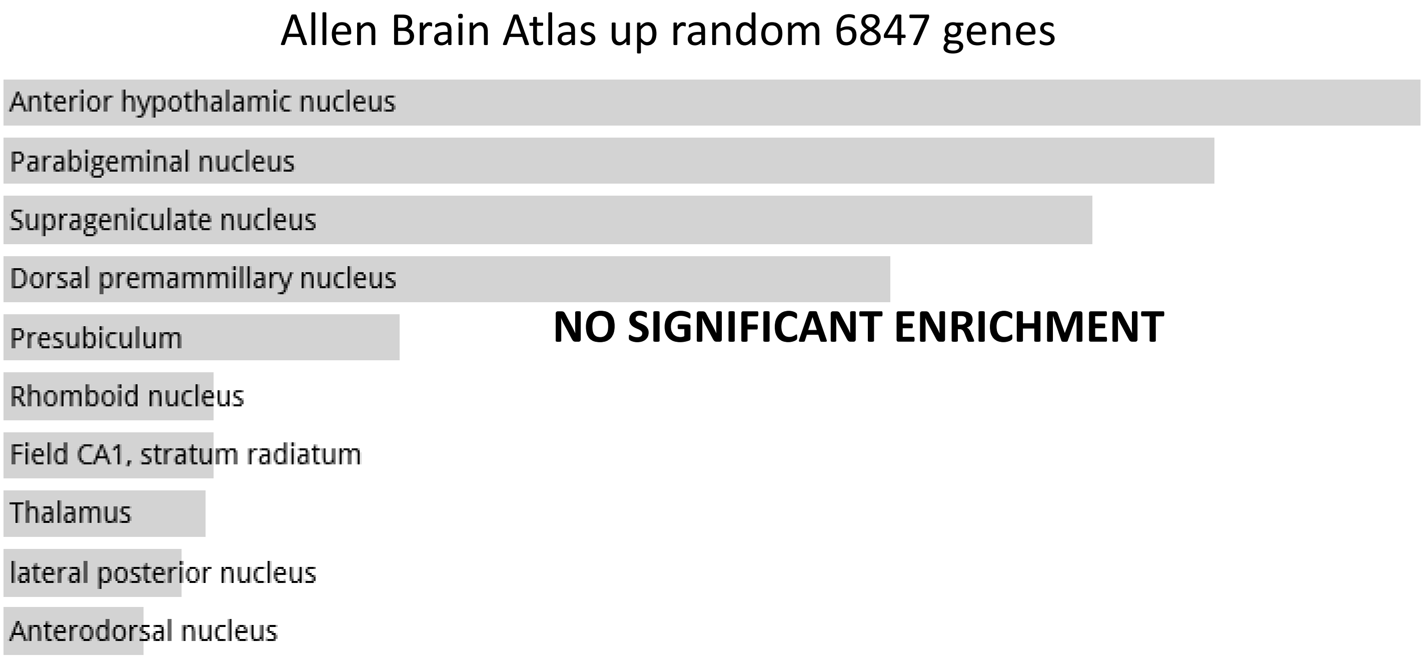
Allen Brain Atlas up random 6847 genes
Anterior hypothalamic nucleus
Parabigeminal nucleus
Suprageniculate nucleus
Dorsal premammillary nucleus
Presubiculum
Rhomboid nucleus
Field CA1, stratum radiatum
Thalamus
lateral posterior nucleus
Anterodorsal nucleus
NO SIGNIFICANT ENRICHMENT

**Random Set 8**

ARCHS4 Tissues random 6001 genes

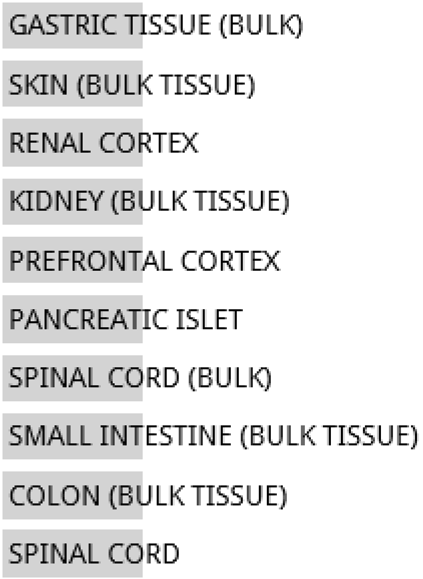


**NO SIGNIFICANT ENRICHMENT**

Jensen TISSUES random 6001 genes

**NO DATA AVAILABLE**

**Random Set 8**

Jensen DISEASES random 6001 genes

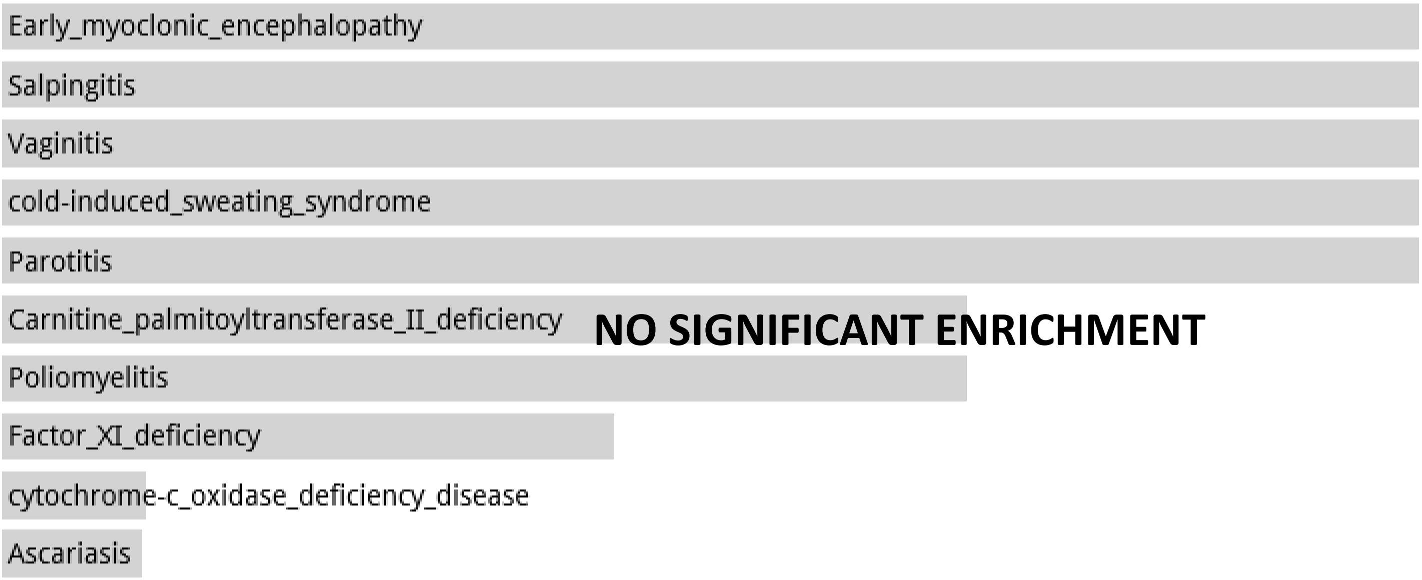


Disease Perturbations from GEO up random 6000 genes

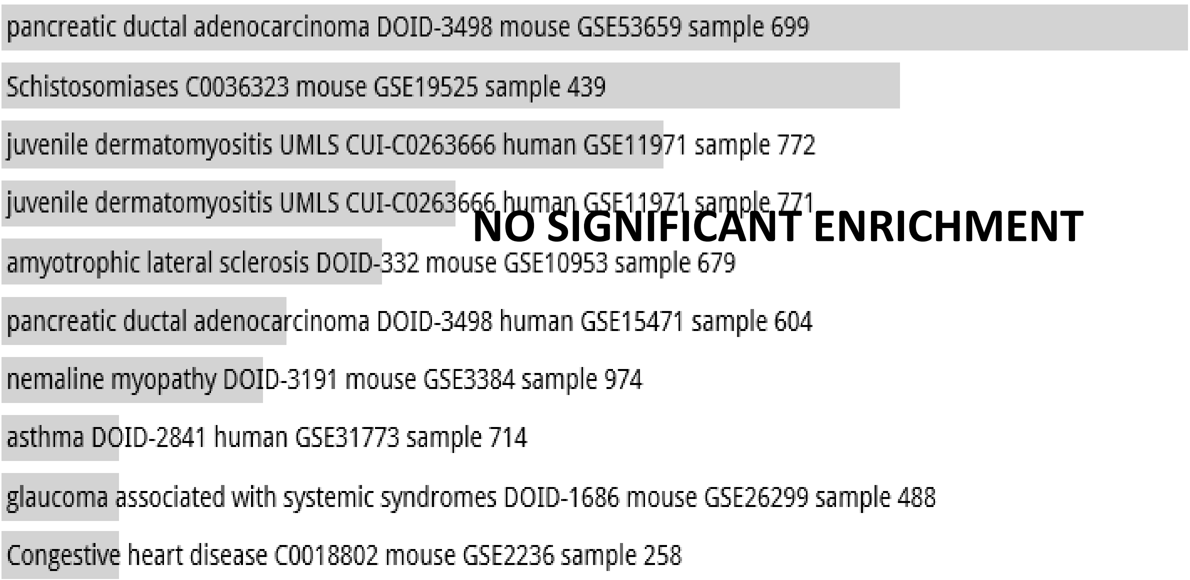


Disease Perturbations from GEO down random 6000 genes

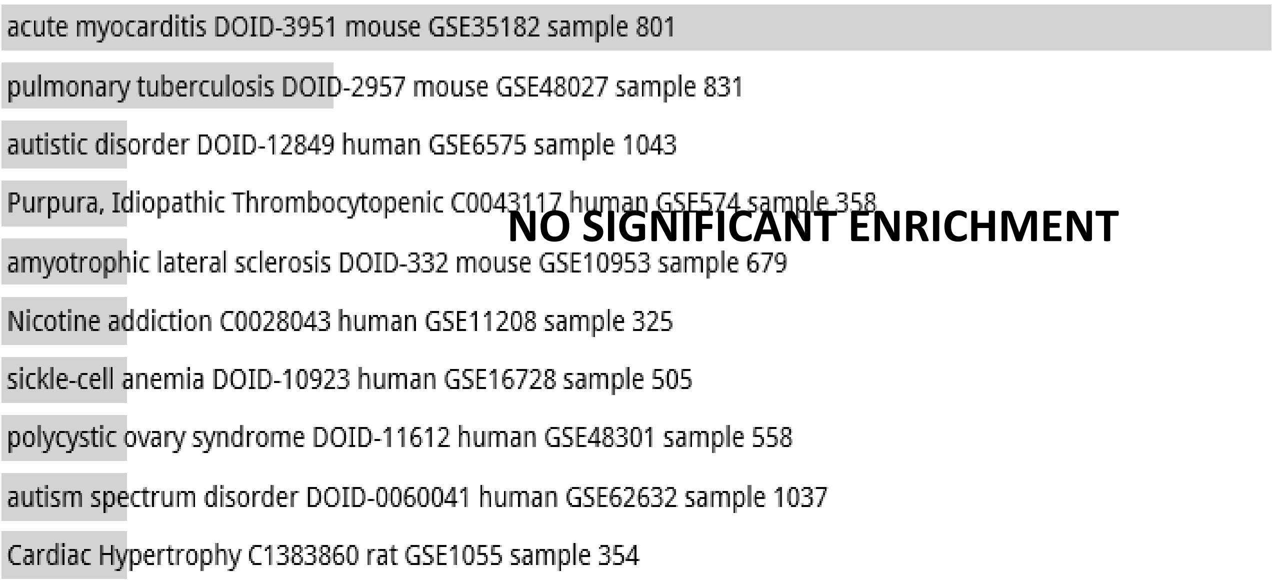

**Random Set 8**

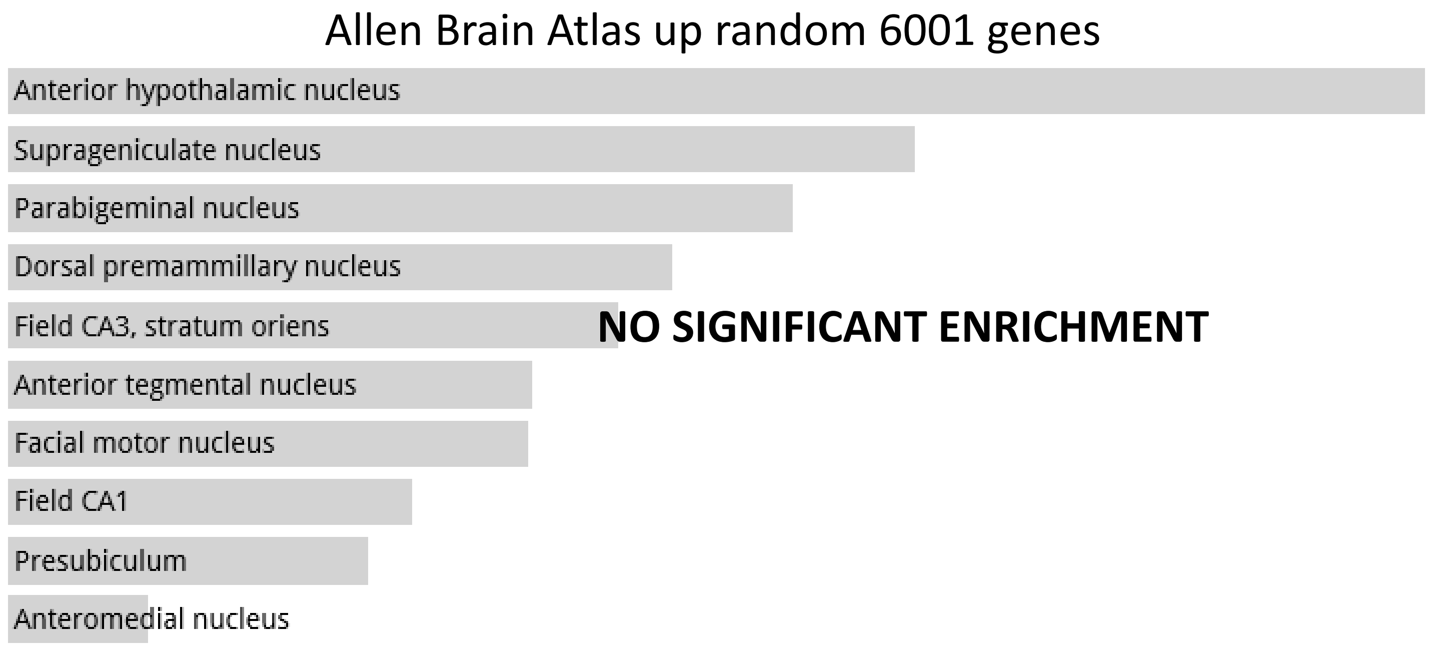
Allen Brain Atlas up random 6001 genes
Anterior hypothalamic nucleus
Suprageniculate nucleus
Parabigeminal nucleus
Dorsal premammillary nucleus
Field CA3, stratum oriens
Anterior tegmental nucleus
Facial motor nucleus
Field CA1
Presubiculum
Anteromedial nucleus
NO SIGNIFICANT ENRICHMENT

**Random Set 9**

ARCHS4 Tissues random 6000 genes

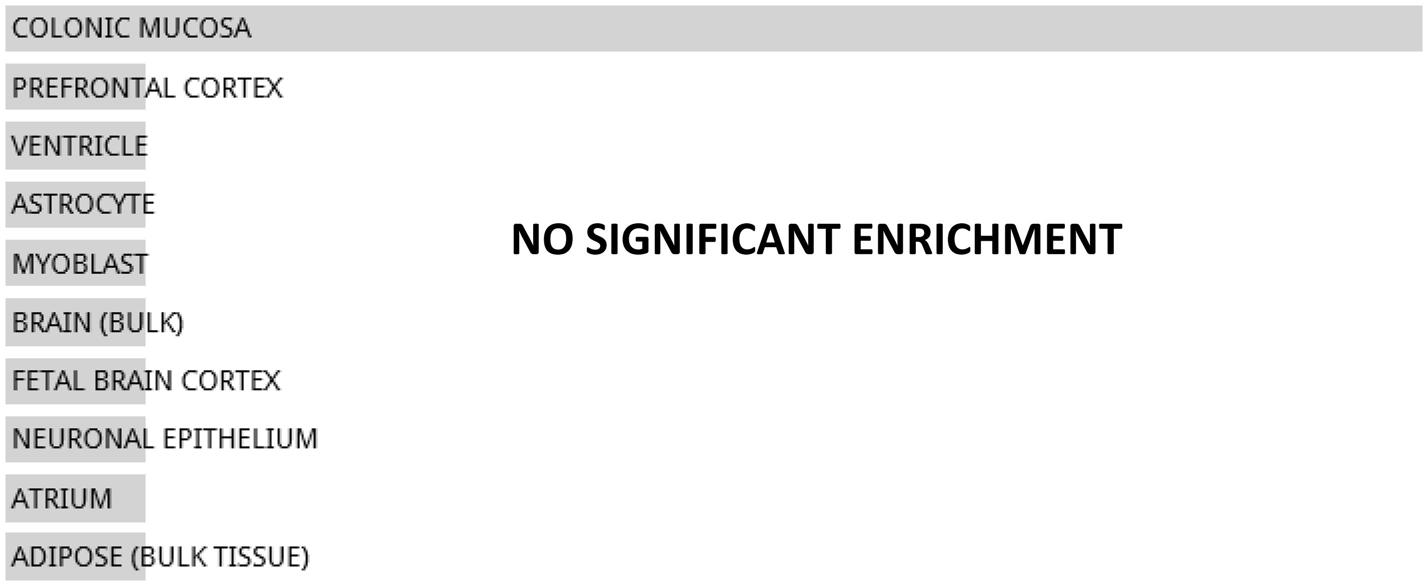


**NO SIGNIFICANT ENRICHMENT**

Jensen TISSUES random 6000 genes

**NO DATA AVAILABLE**

**Random Set 9**

#### Jensen DISEASES random 6000 genes

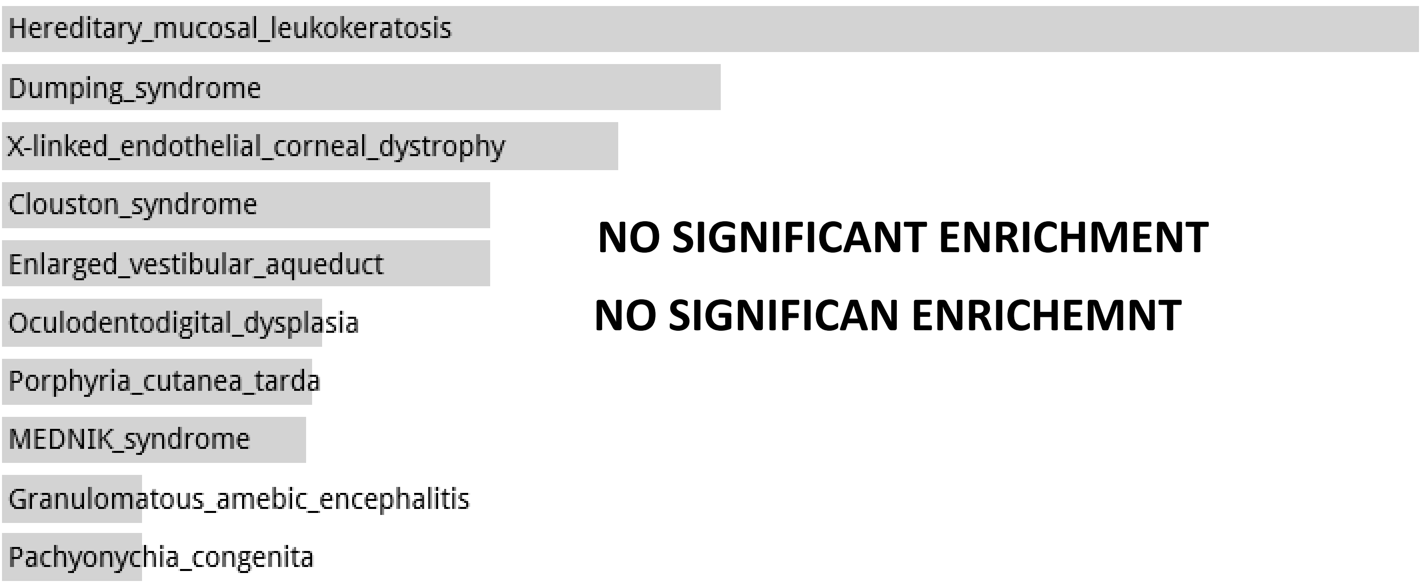


#### Disease Perturbations from GEO up random 6000 genes

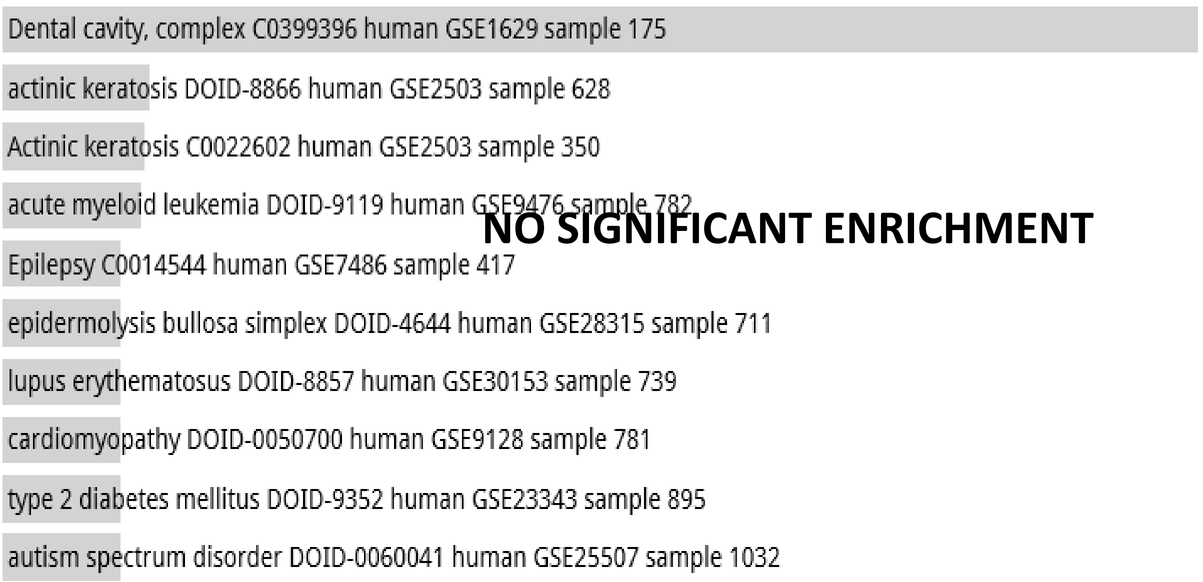


#### Disease Perturbations from GEO down random 6000 genes

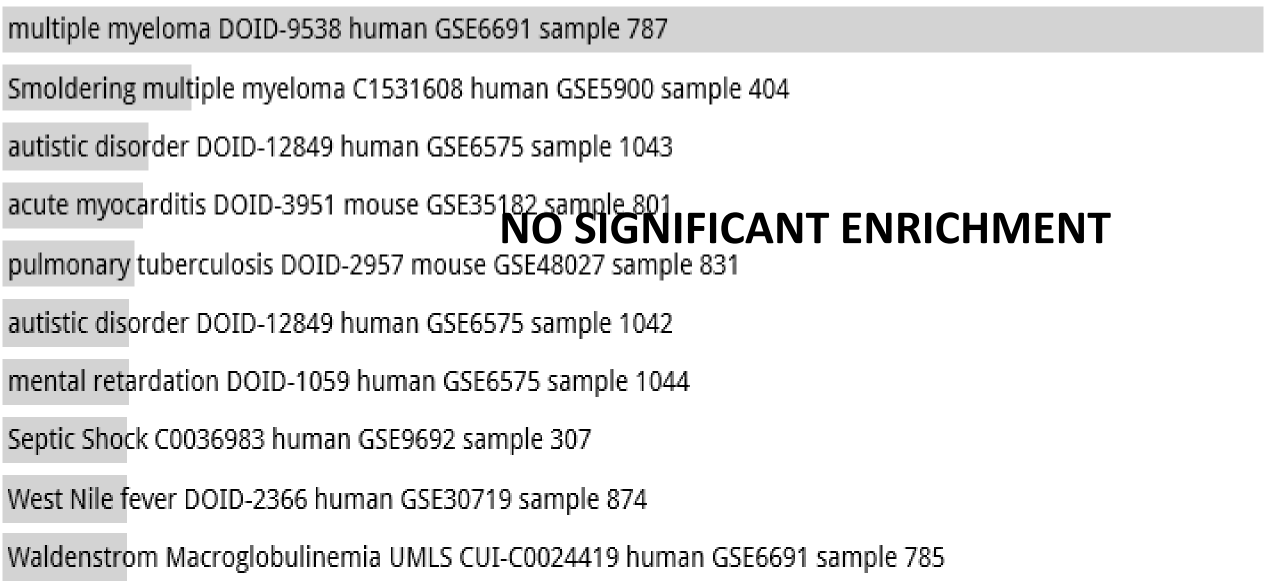

**Random Set 9**

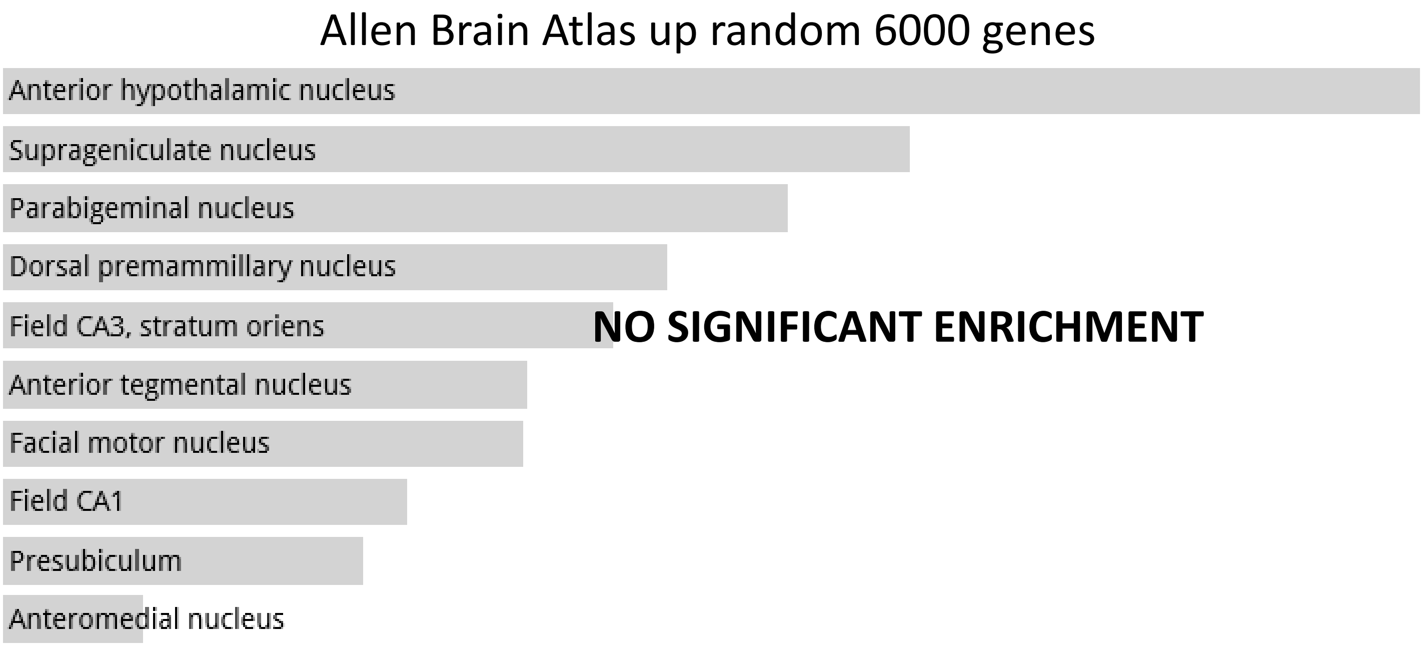
Allen Brain Atlas up random 6000 genes
Anterior hypothalamic nucleus
Suprageniculate nucleus
Parabigeminal nucleus
Dorsal premammillary nucleus
Field CA3, stratum oriens
Anterior tegmental nucleus
Facial motor nucleus
Field CA1
Presubiculum
Anteromedial nucleus
NO SIGNIFICANT ENRICHMENT

**Random Set 10**

ARCHS4 Tissues random 6000 genes

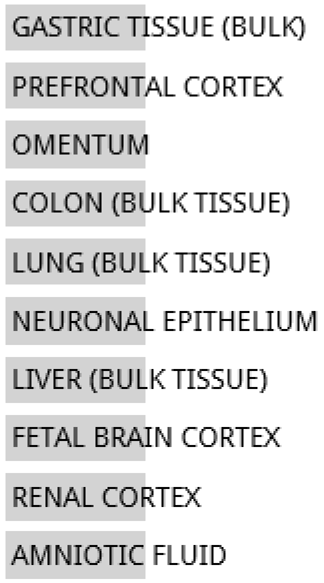


**NO SIGNIFICANT ENRICHMENT**

Jensen TISSUES random 6000 genes

**NO DATA AVAILABLE**

#### Random Set 10

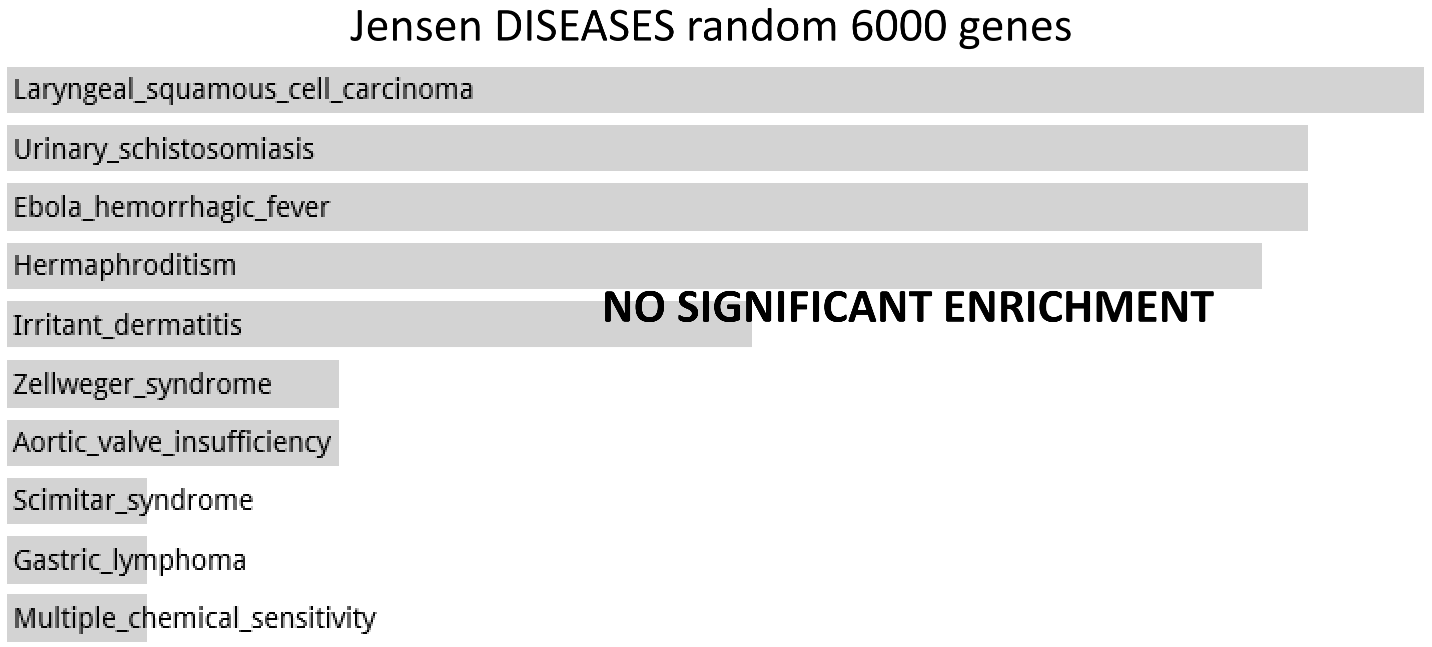


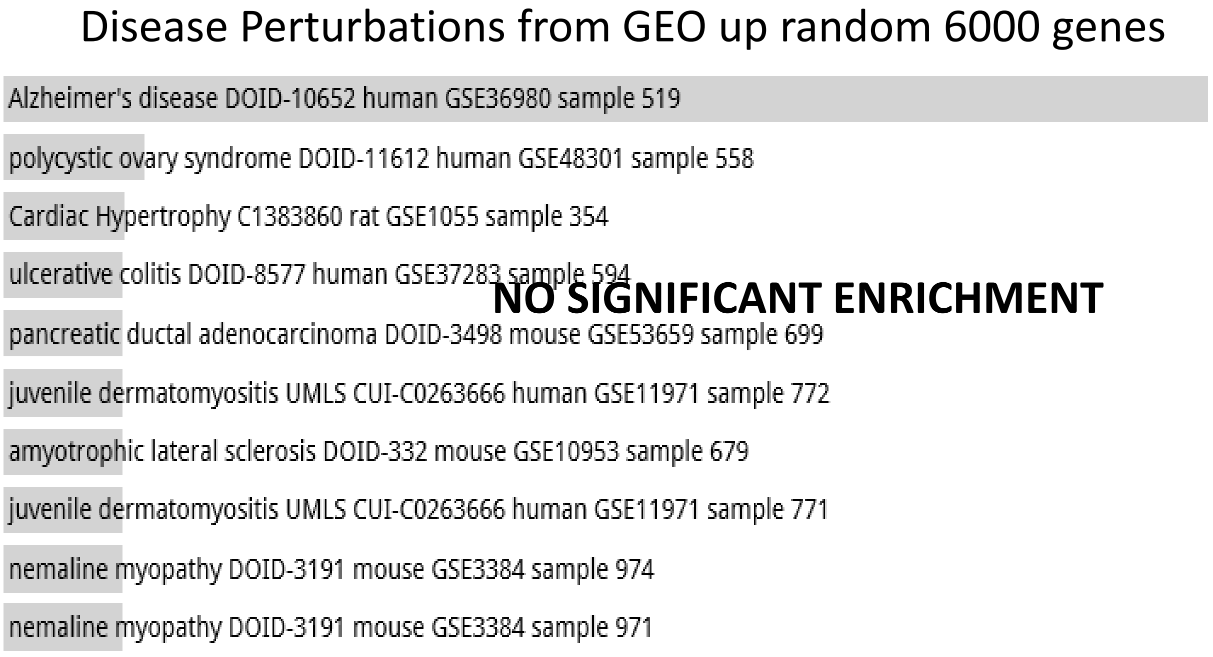


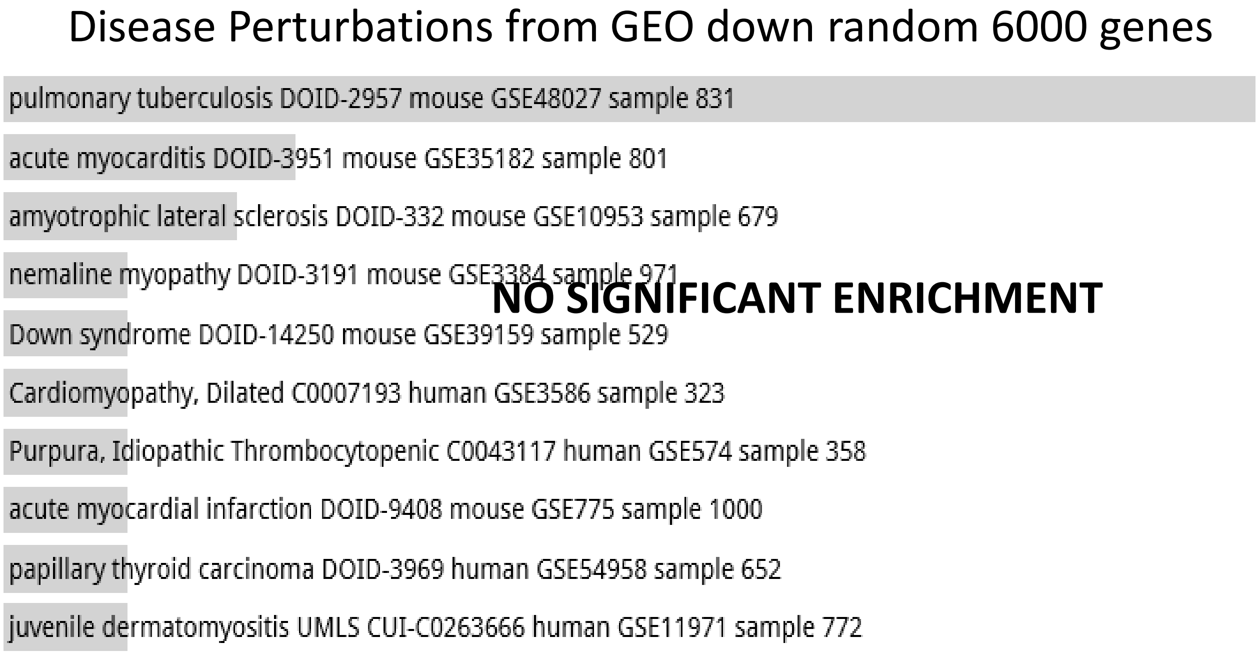

#### Random Set 10

Allen Brain Atlas up random 6000 genes

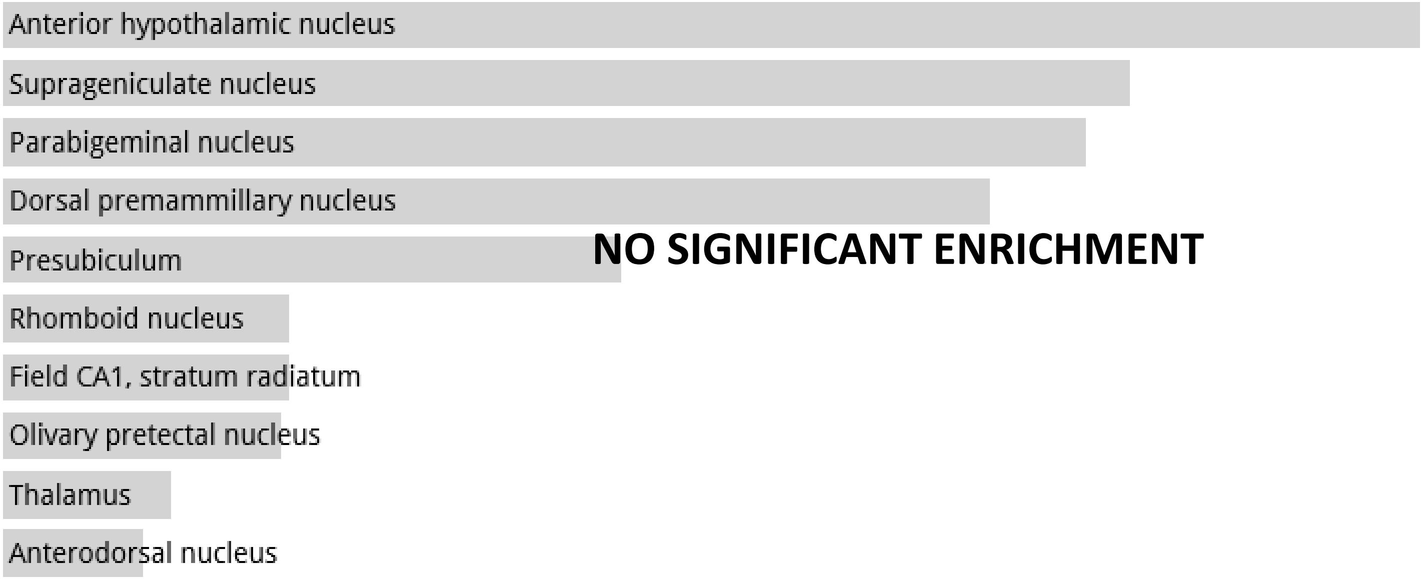

**Random Set 11**

ARCHS4 Tissues random 4000 genes

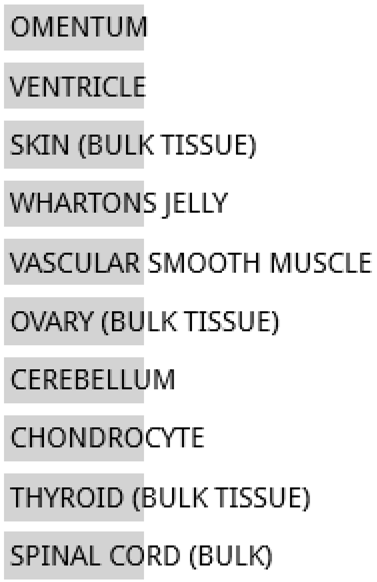


**NO SIGNIFICANT ENRICHMENT**

Jensen TISSUES random 4000 genes

**NO DATA AVAILABLE**

**Random Set 11**

Jensen DISEASES random 4000 genes

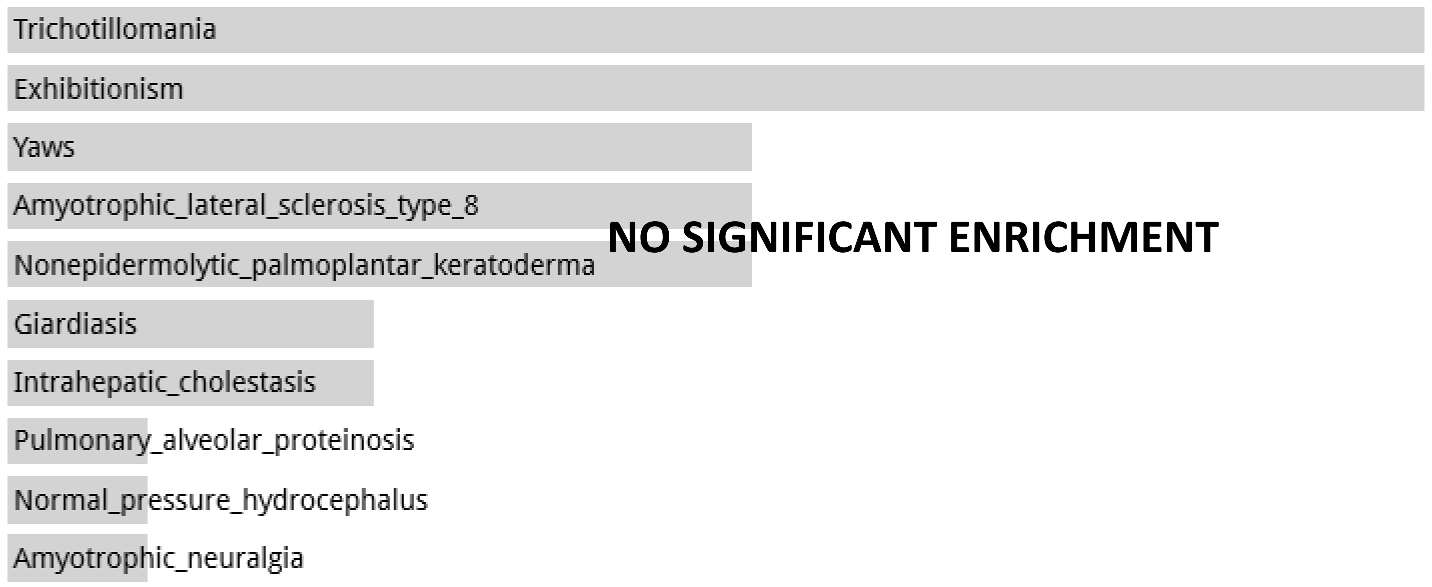


Disease Perturbations from GEO up random 4000 genes

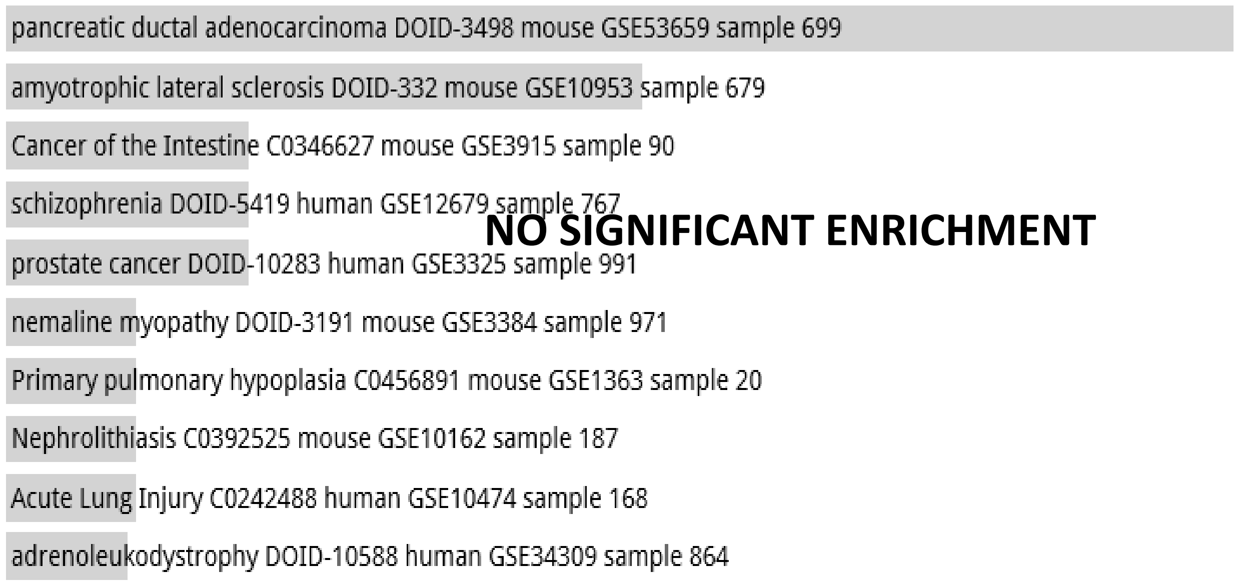


Disease Perturbations from GEO down random 4000 genes

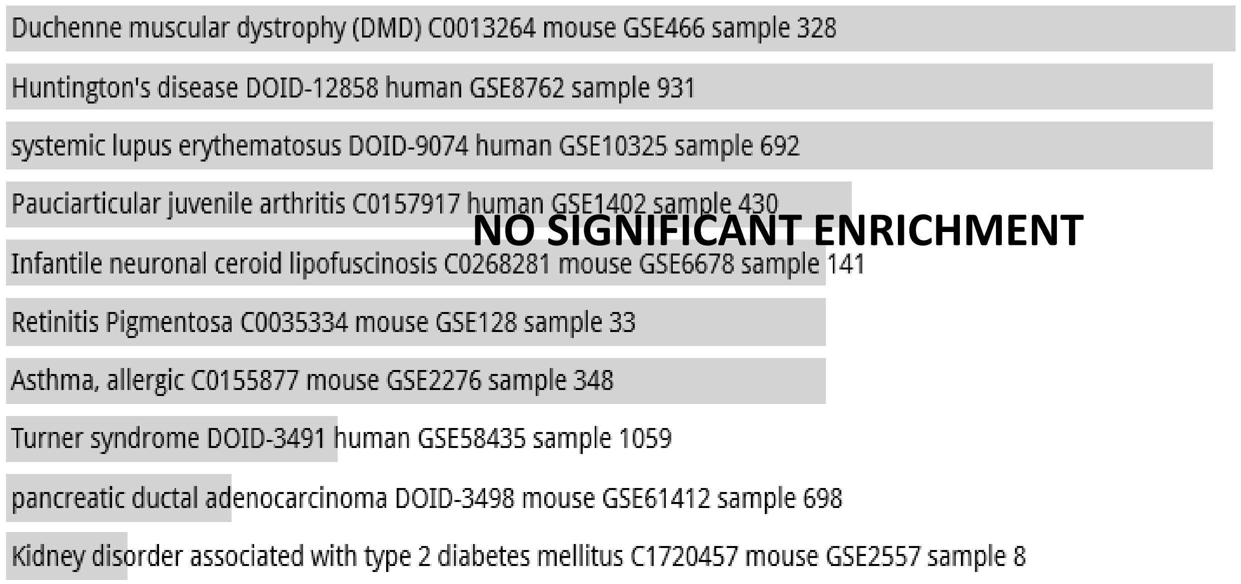

**Random Set 11**

Allen Brain Atlas up random 4000 genes

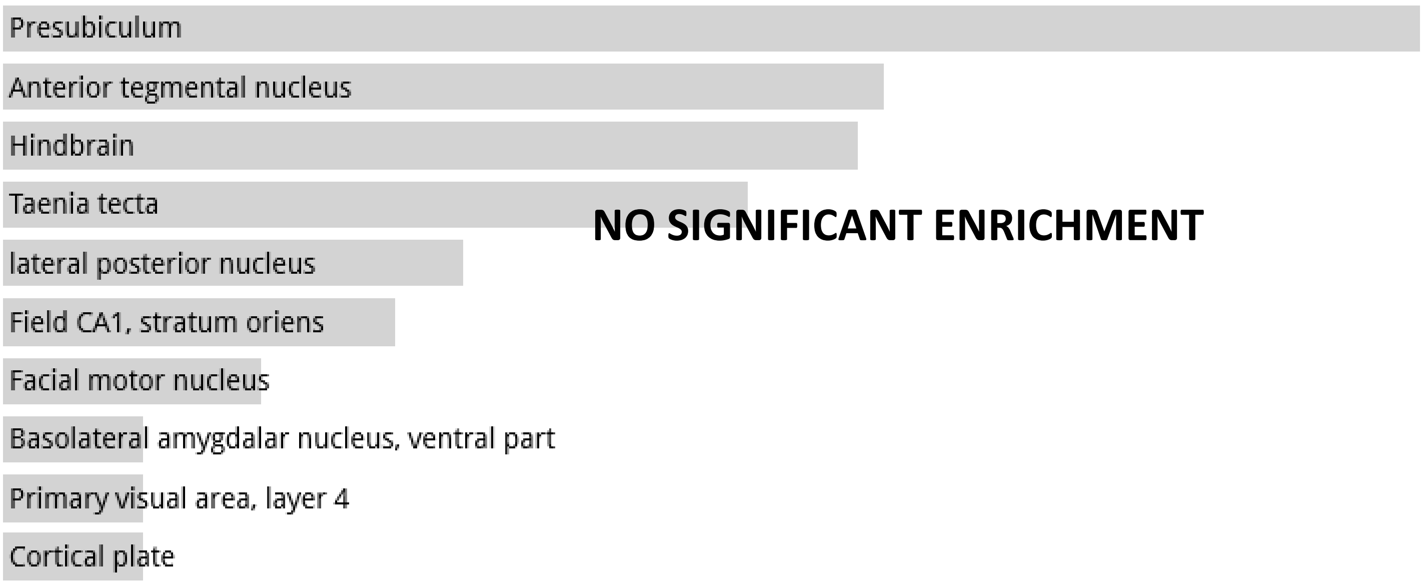

**Random Set 12**

ARCHS4 Tissues random 4000 genes

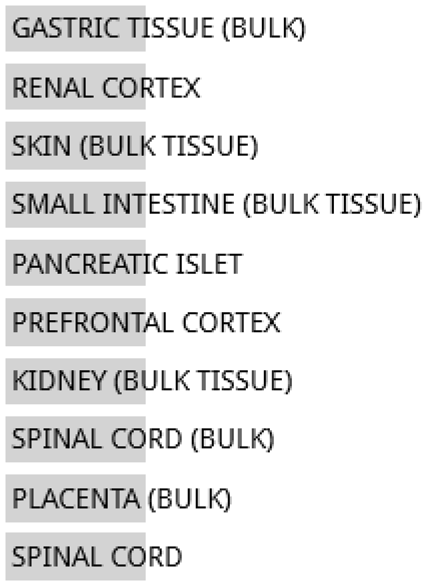


**NO SIGNIFICANT ENRICHMENT**

Jensen TISSUES random 4000 genes

**NO DATA AVAILABLE**

#### Random Set 12

Jensen DISEASES random 4000 genes

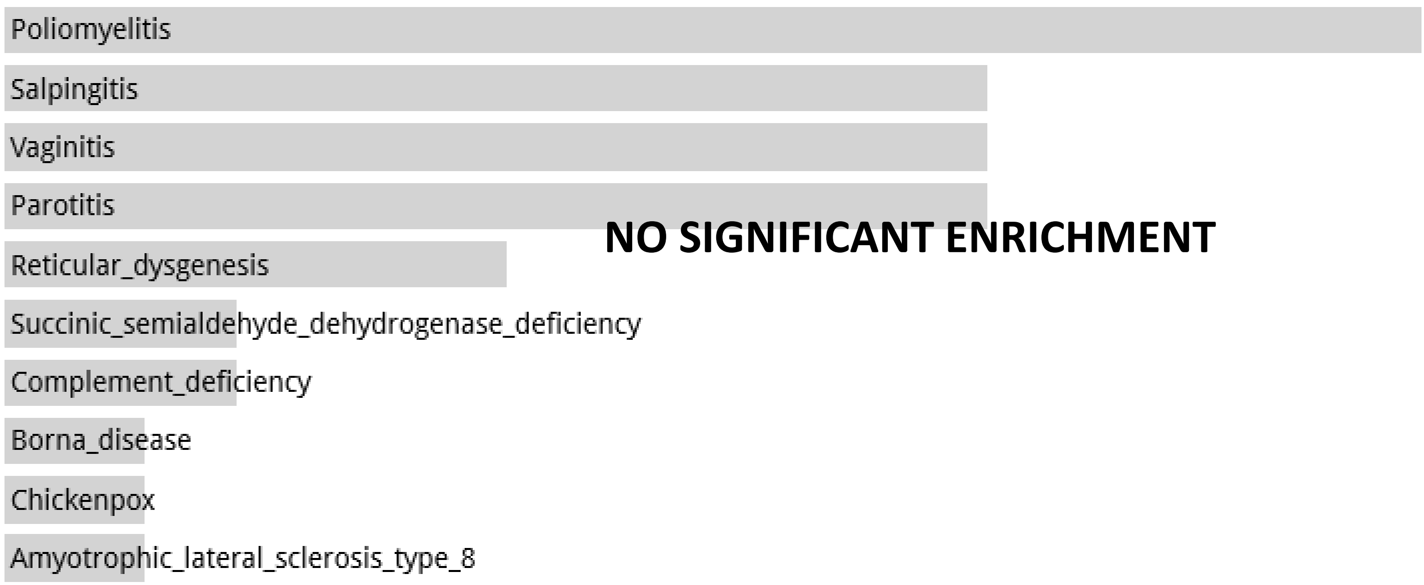


Disease Perturbations from GEO up random 4000 genes

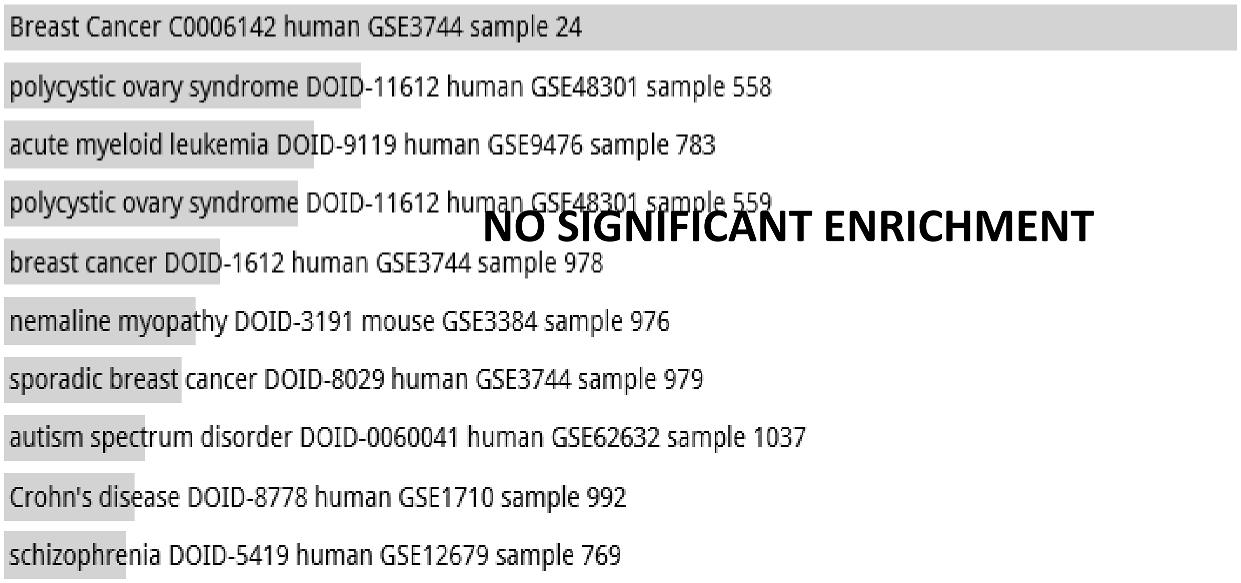


Disease Perturbations from GEO down random 4000 genes

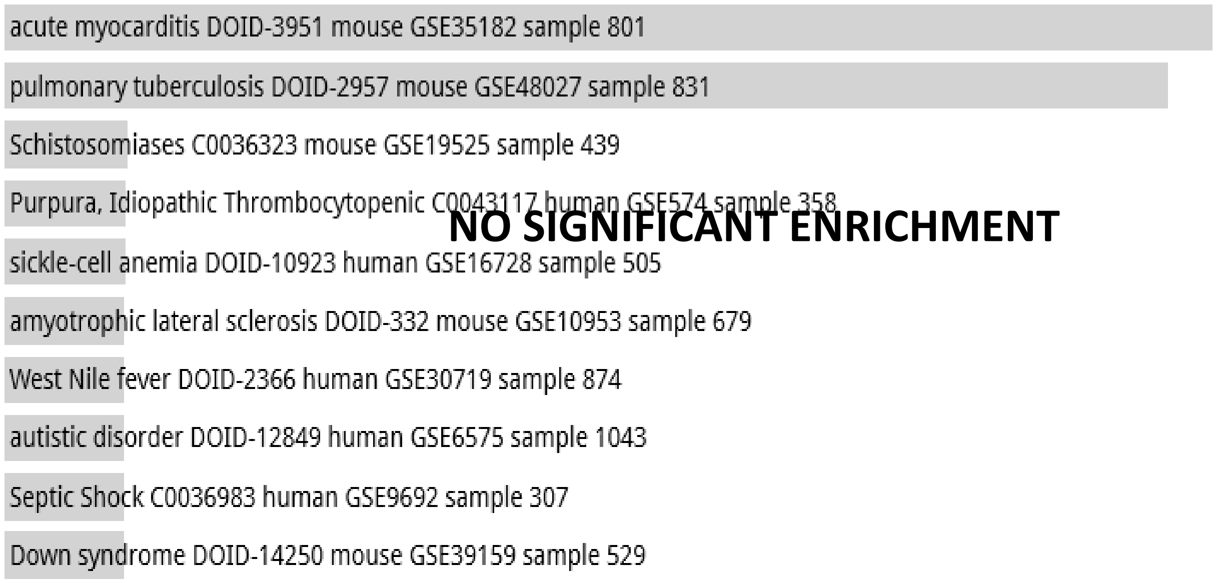

**Random Set 12**

#### Allen Brain Atlas up random 4000 genes

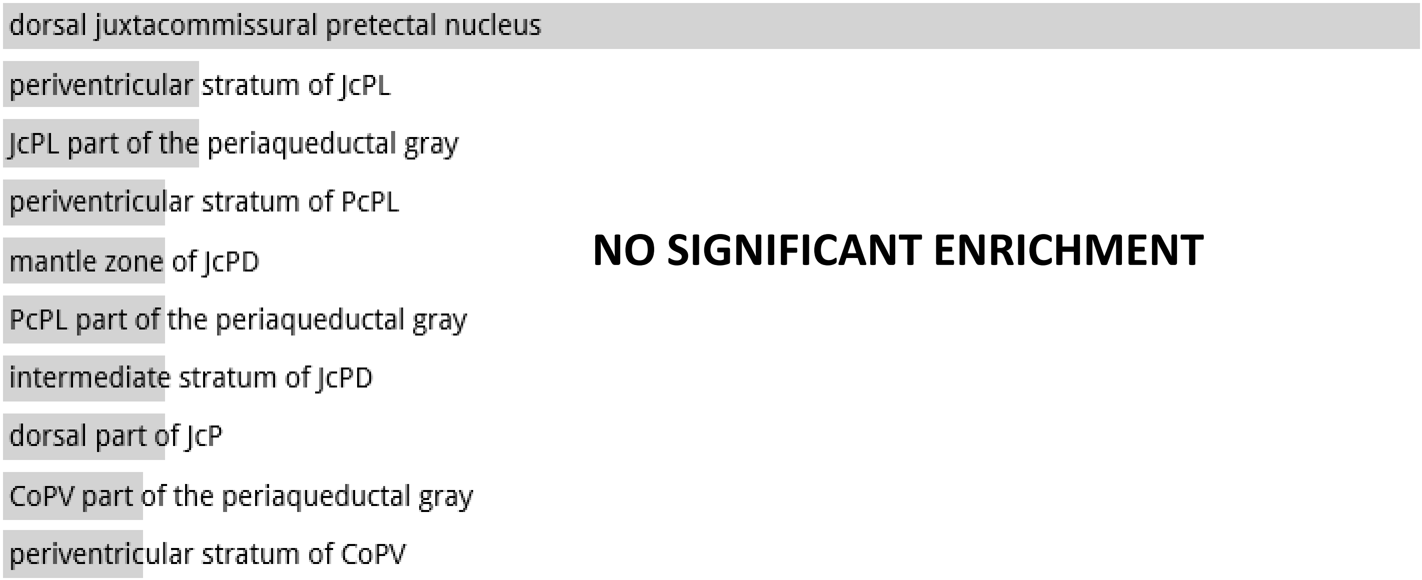


**NO SIGNIFICANT ENRICHMENT**

**Random Set 13**

#### ARCHS4 Tissues random 4000 genes

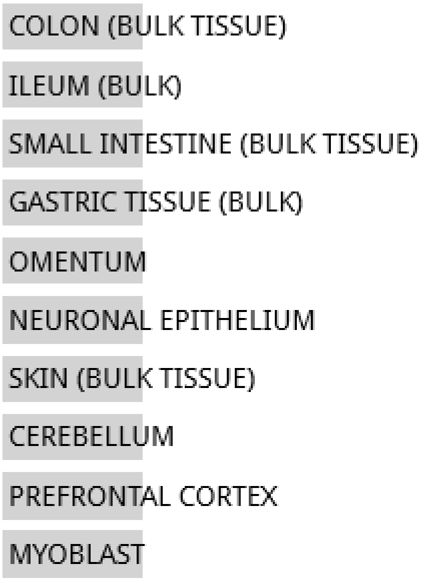


**NO SIGNIFICANT ENRICHMENT**

#### Jensen TISSUES random 4000 genes

**NO DATA AVAILABLE**

**Random Set 13**

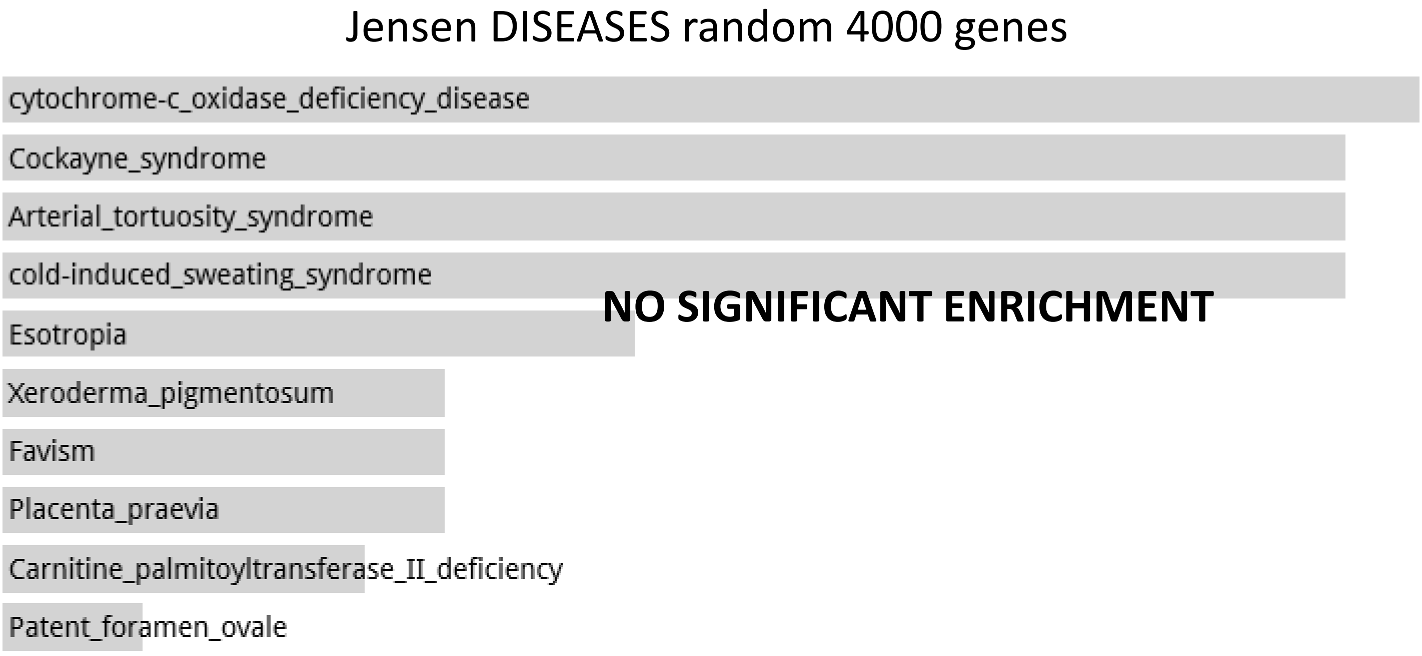


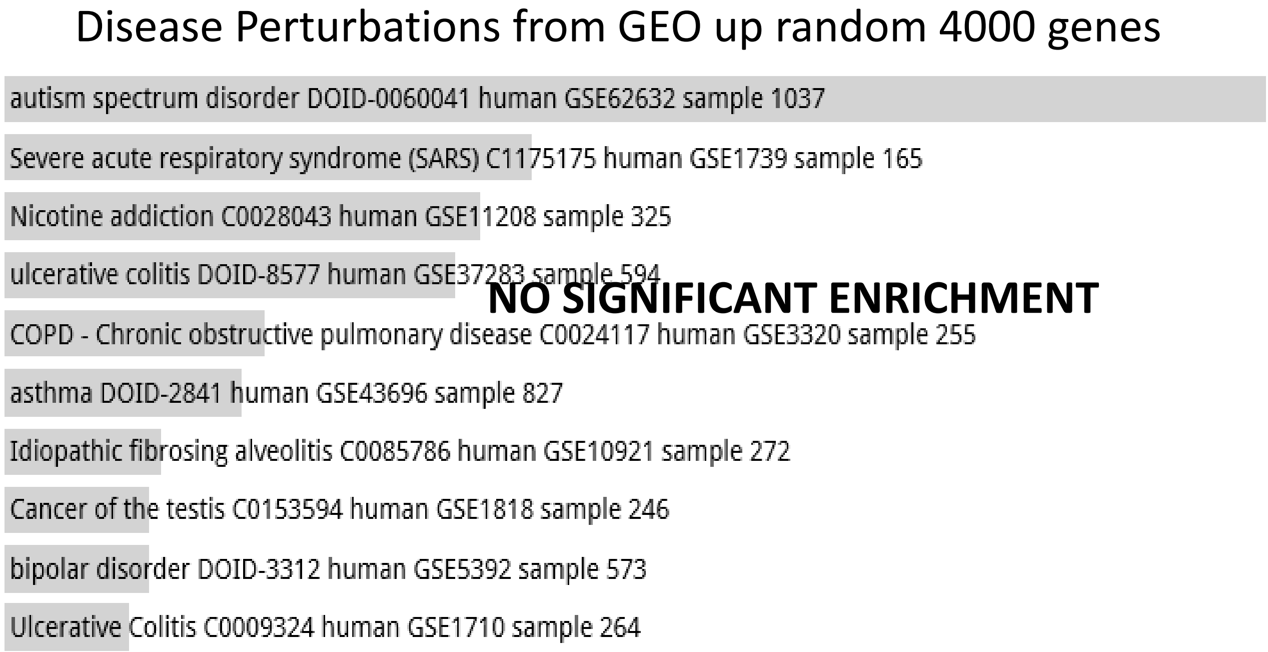


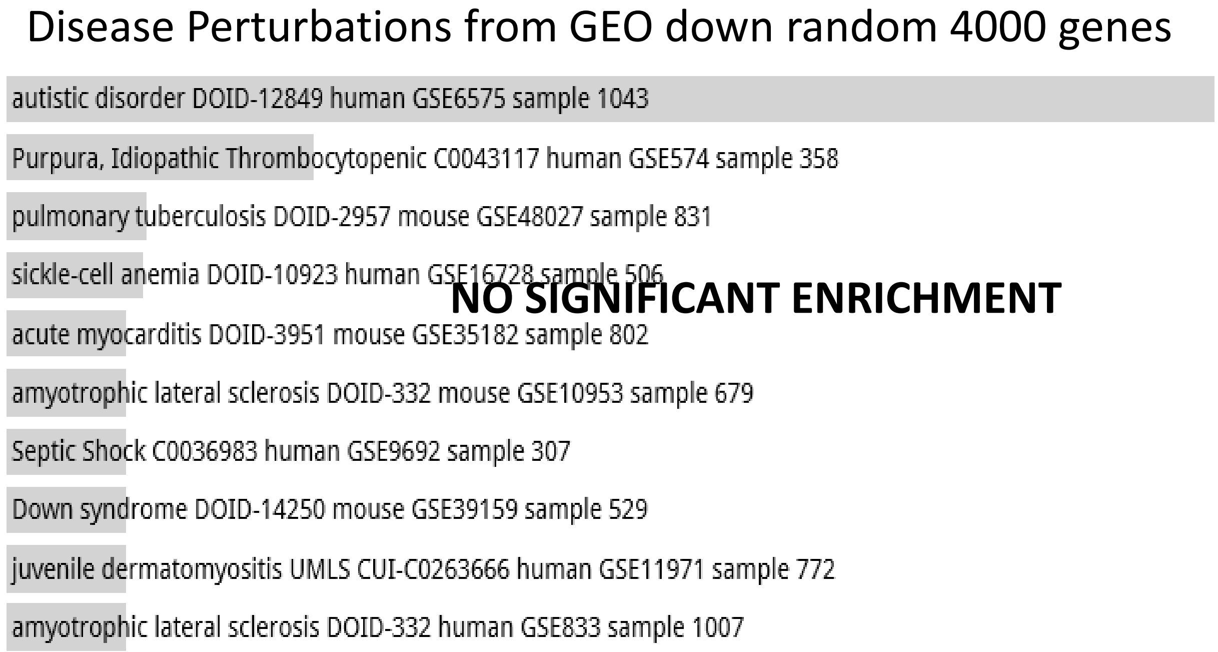

**Random Set 13**

Allen Brain Atlas up random 4000 genes

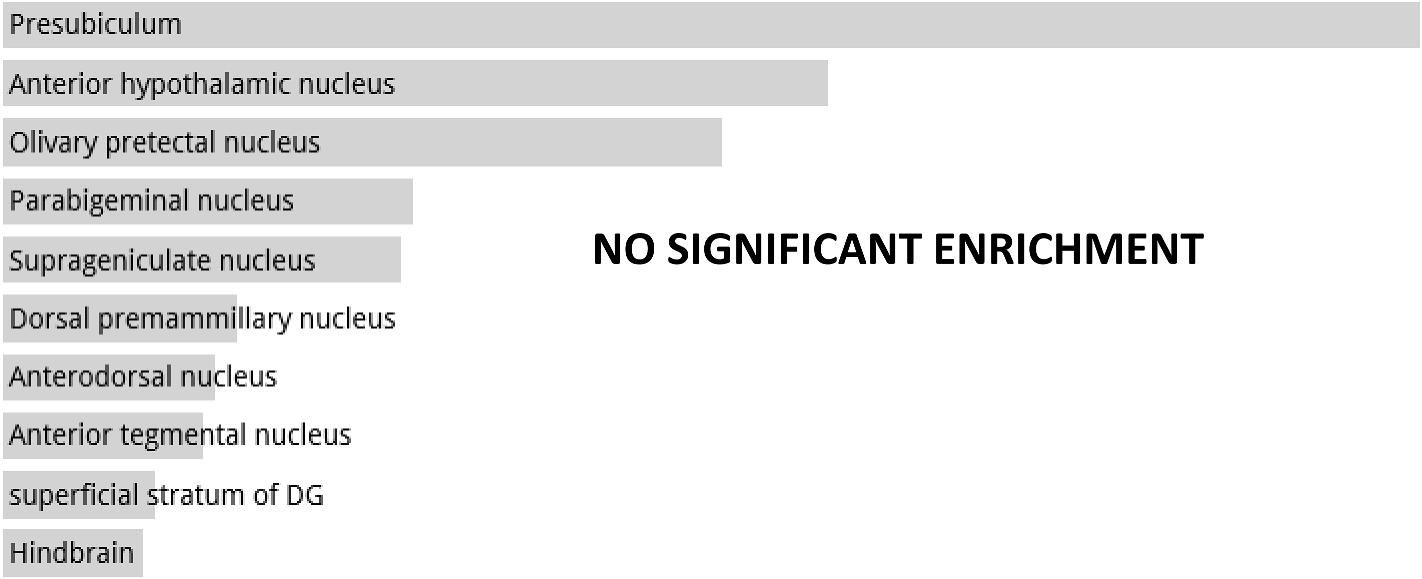


**NO SIGNIFICANT ENRICHMENT**

**Random Set 14**

ARCHS4 Tissues random 4000 genes

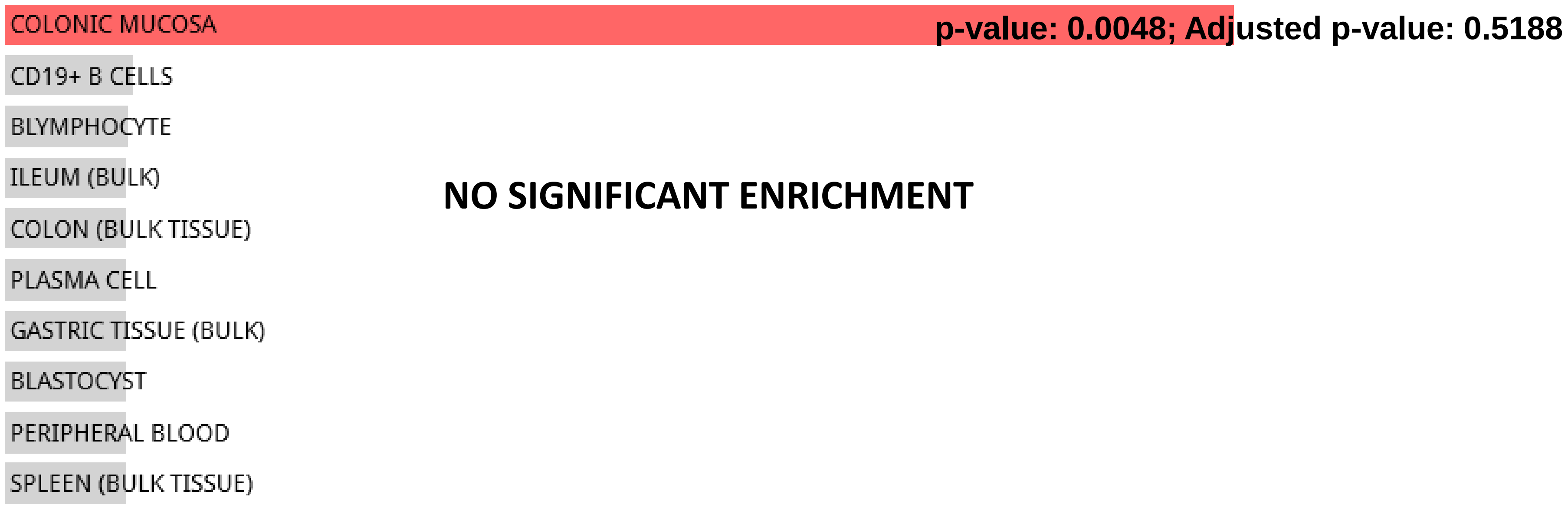


**NO SIGNIFICANT ENRICHMENT**

Jensen TISSUES random 4000 genes

**NO DATA AVAILABLE**

**Random Set 14**

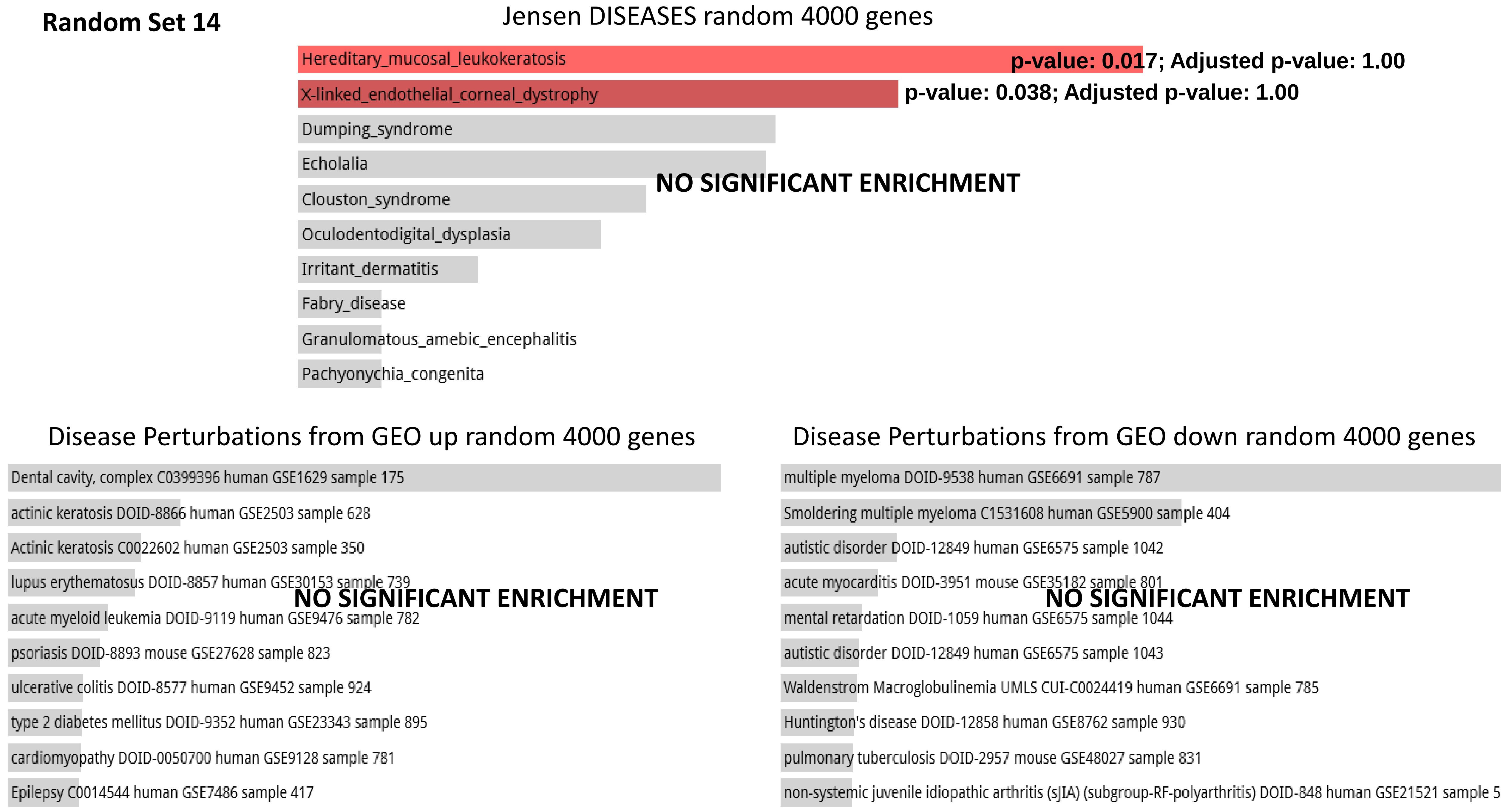

#### Random Set 14

Allen Brain Atlas up random 4000 genes

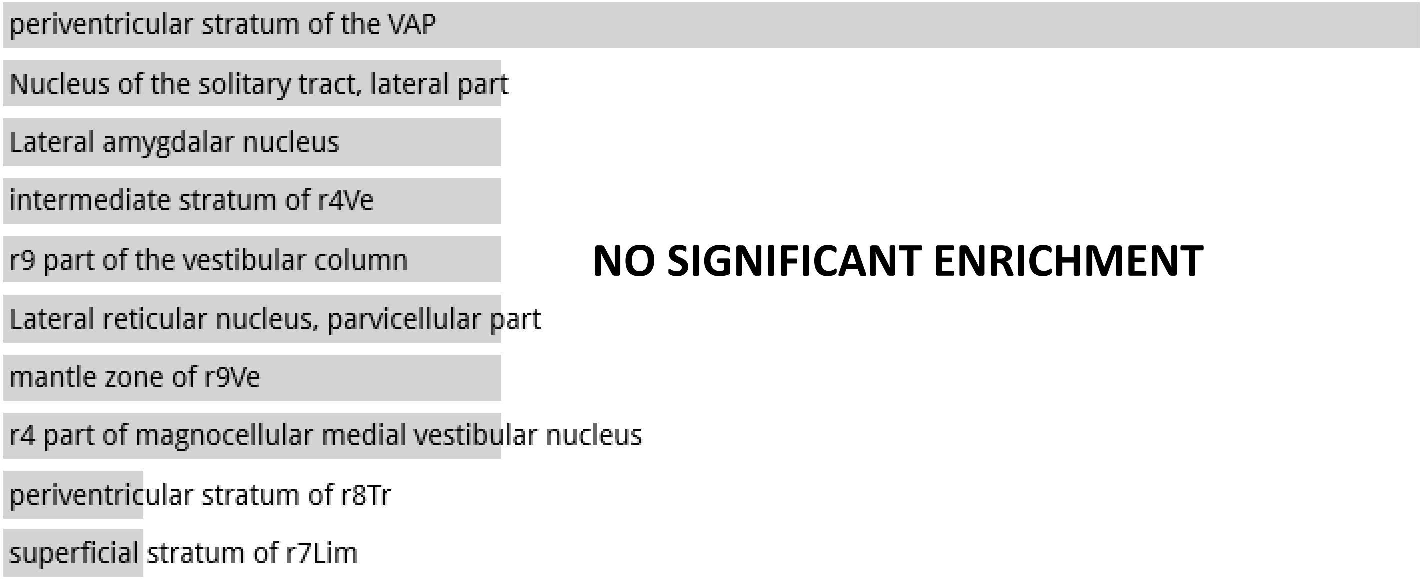


**NO SIGNIFICANT ENRICHMENT**

**Random Set 15**

ARCHS4 Tissues random 4000 genes

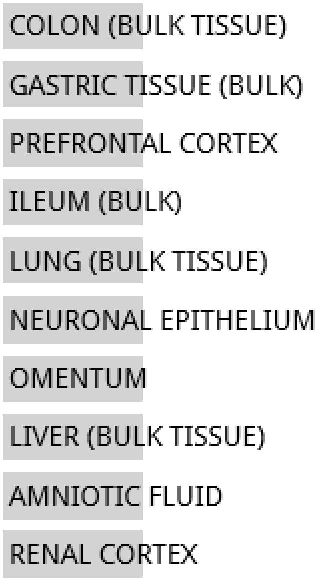


**NO SIGNIFICANT ENRICHMENT**

Jensen TISSUES random 4000 genes

**NO DATA AVAILABLE**

**Random Set 15**

Jensen DISEASES random 4000 genes

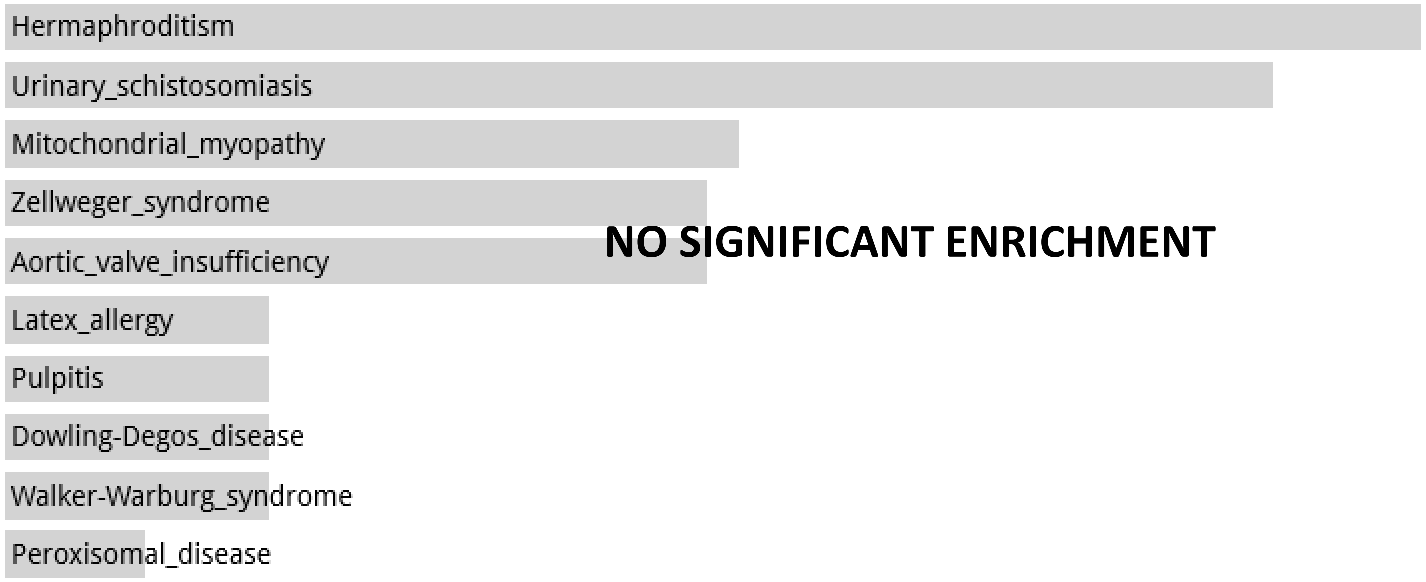


Disease Perturbations from GEO up random 4000 genes

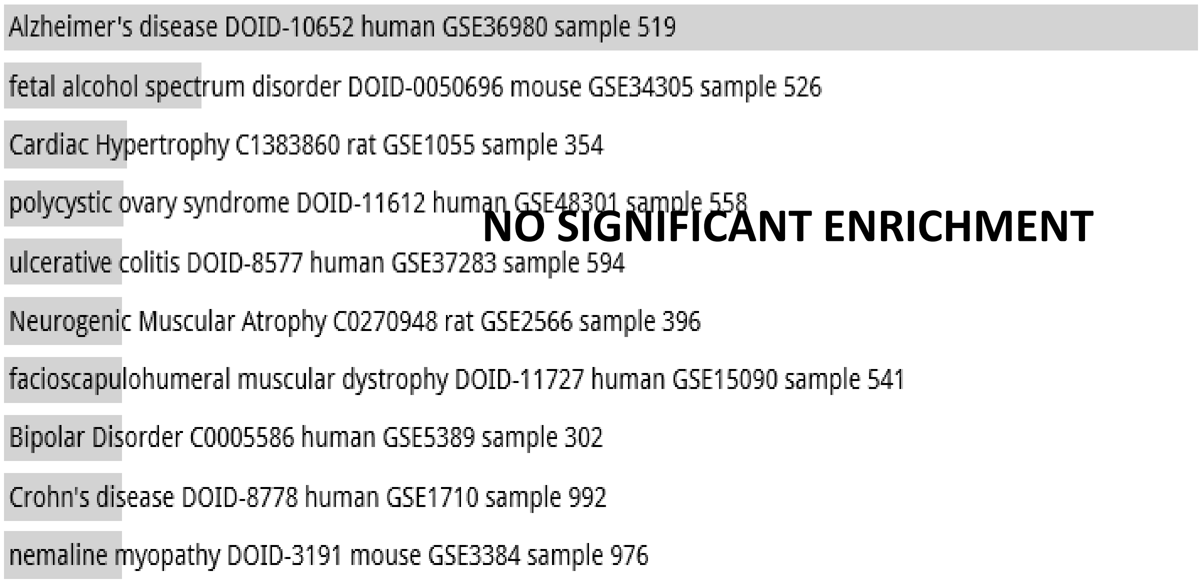


Disease Perturbations from GEO down random 4000 genes

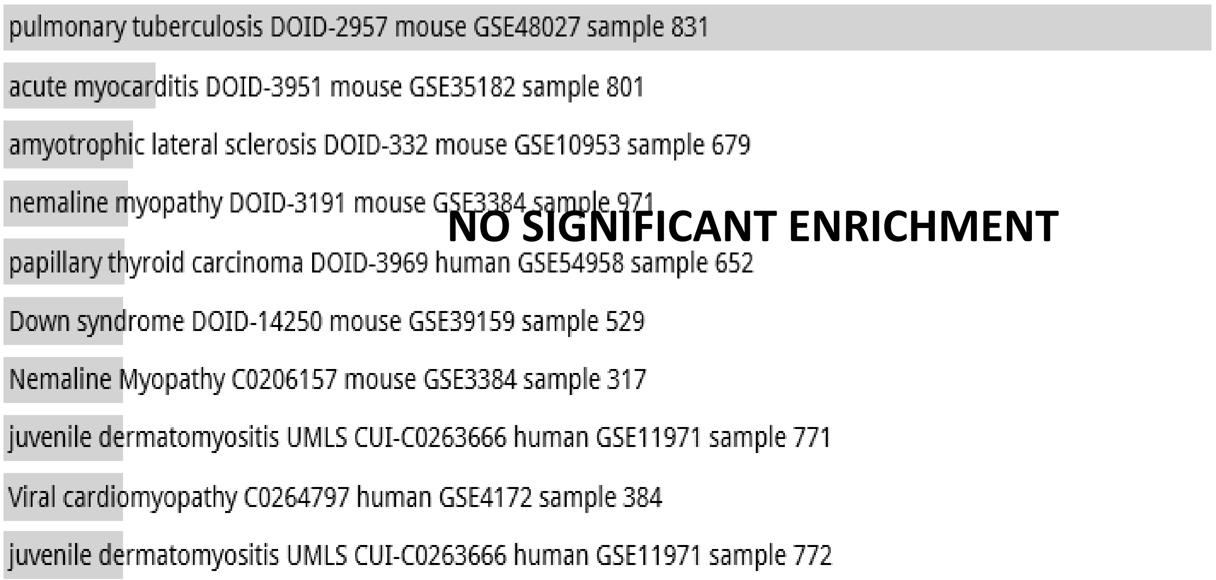

**Random Set 15**

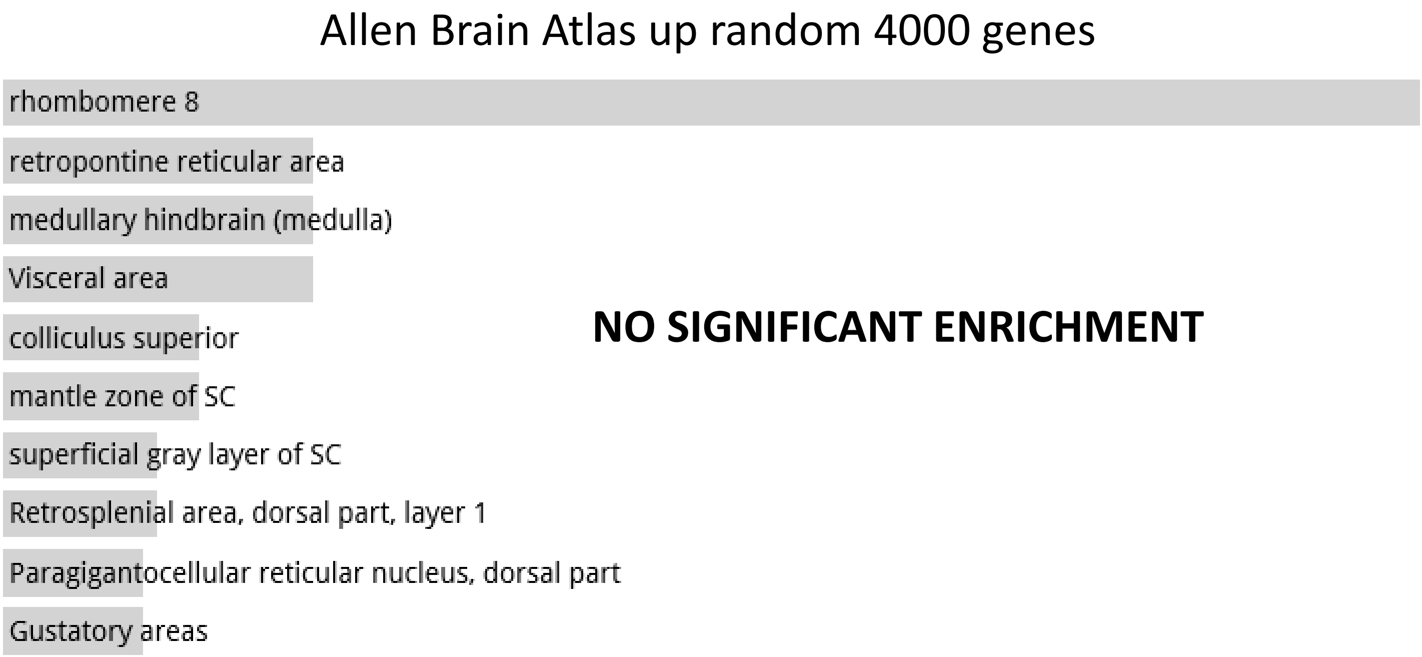

**Random Set 16**

ARCHS4 Tissues random 4000 genes

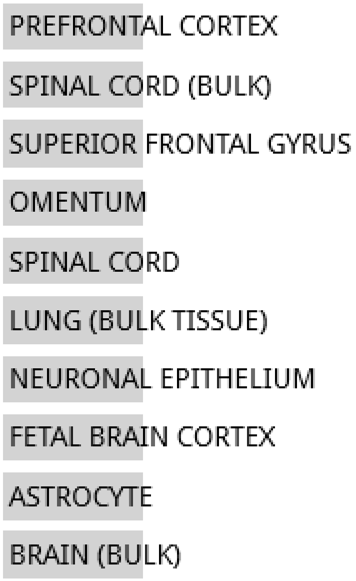


**NO SIGNIFICANT ENRICHMENT**

Jensen TISSUES random 4000 genes

**NO DATA AVAILABLE**

**Random Set 16**

#### Jensen DISEASES random 4000 genes

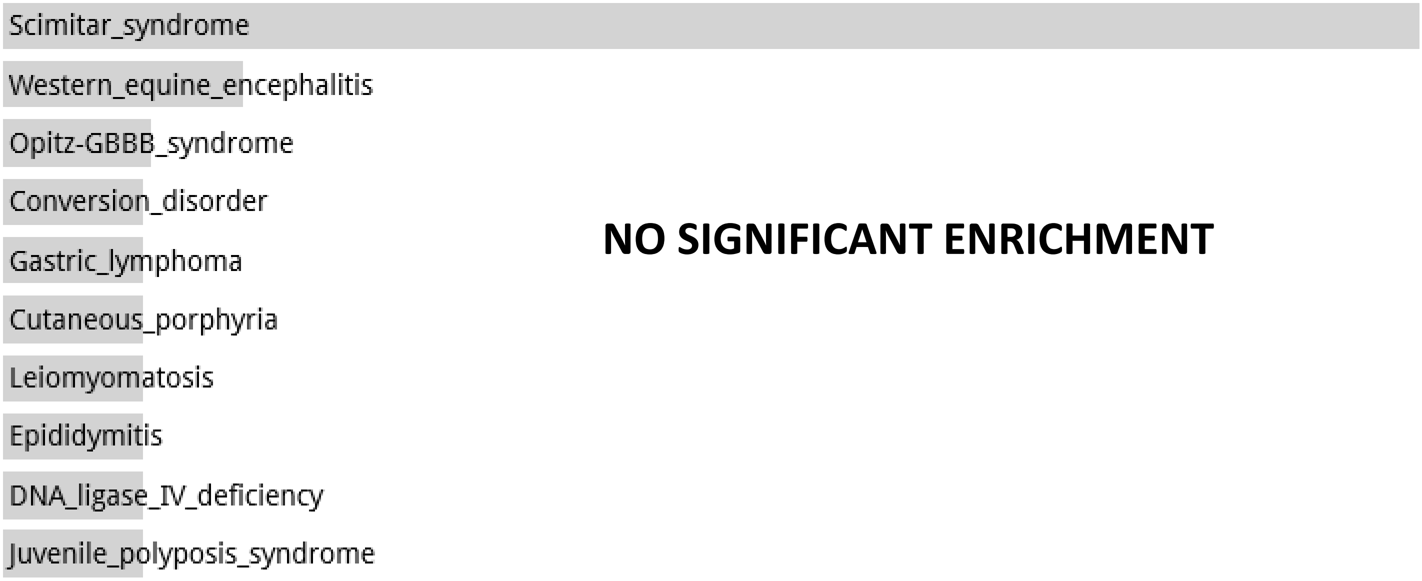


#### Disease Perturbations from GEO up random 4000 genes

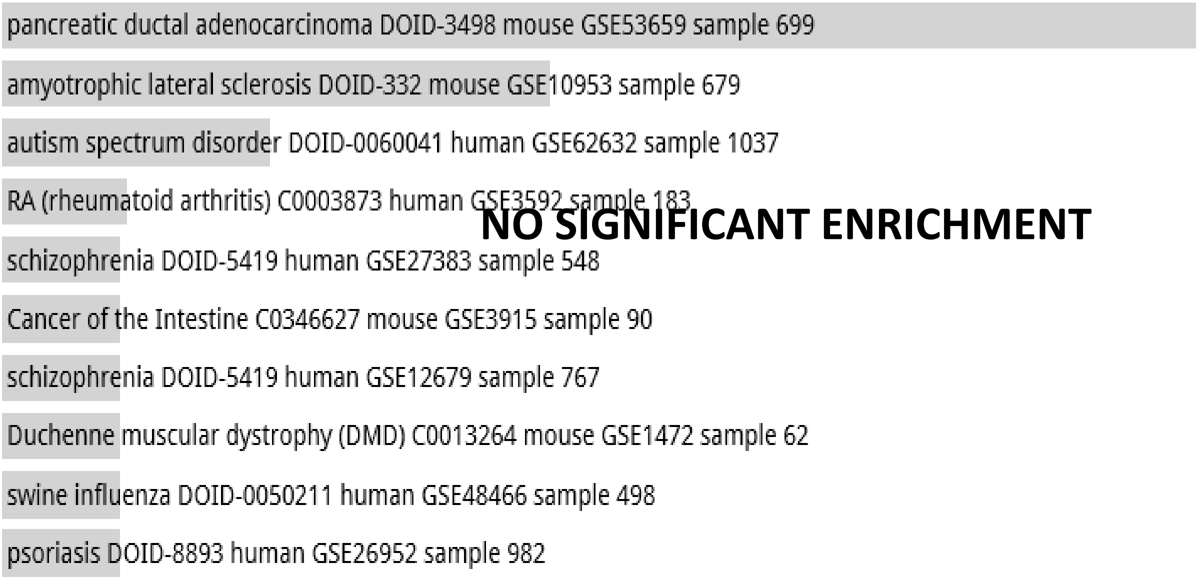


#### Disease Perturbations from GEO down random 4000 genes

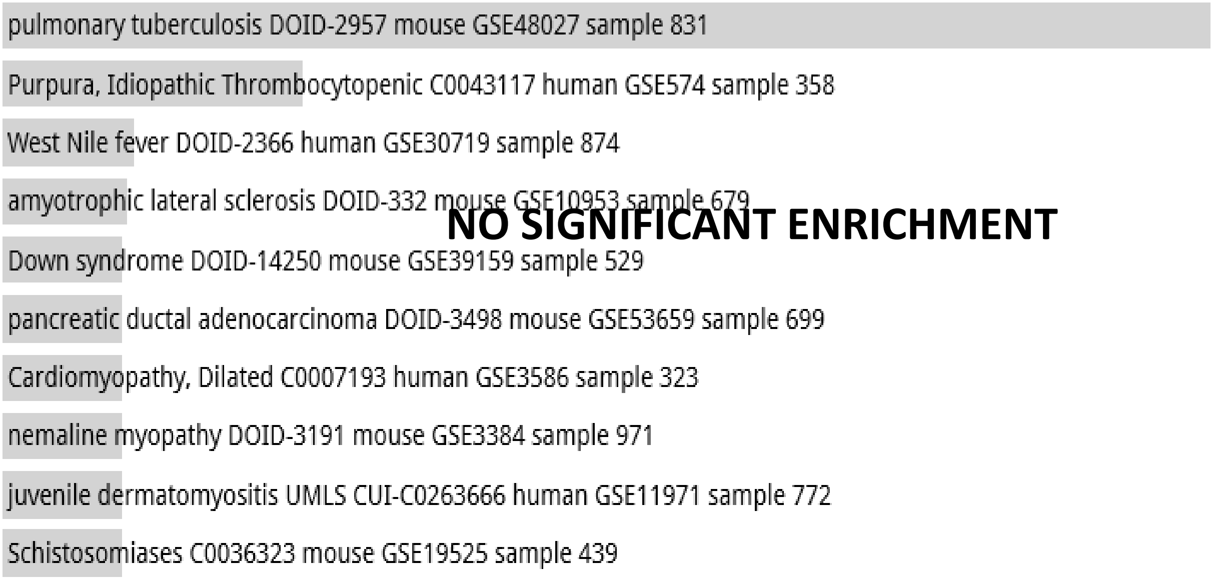

**Random Set 16**

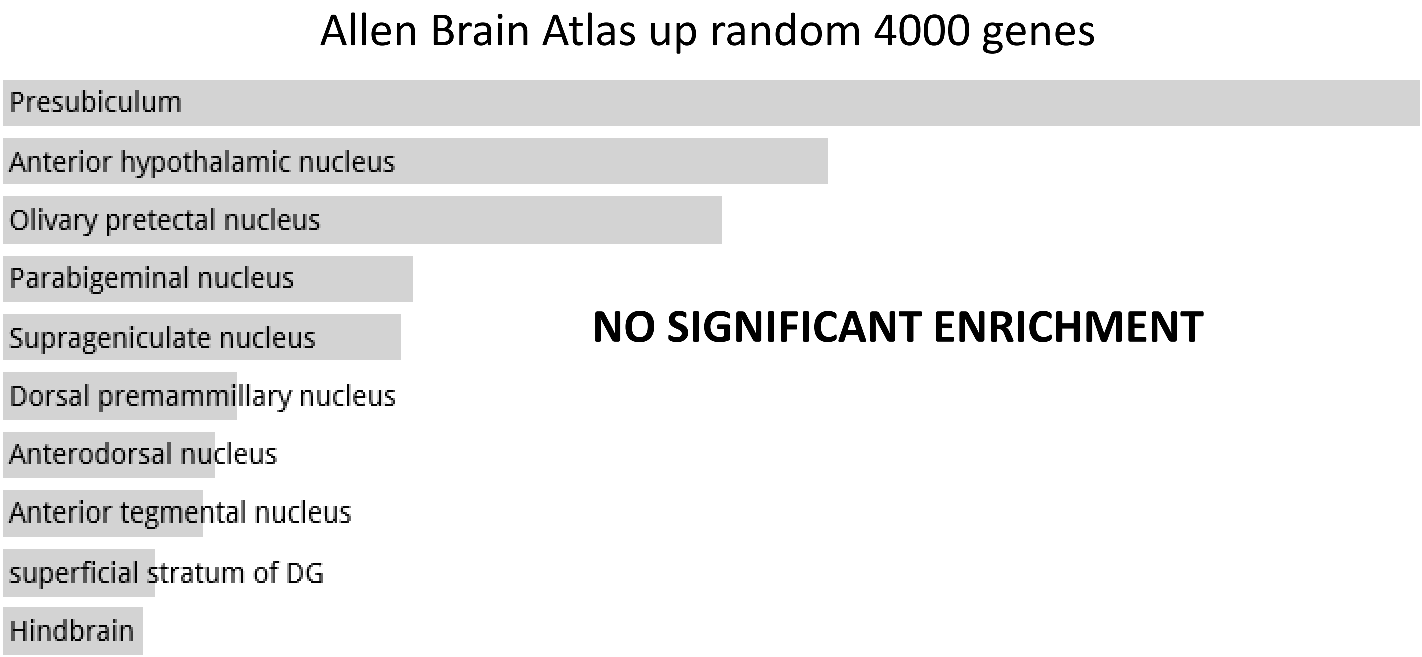

**Random Set 17**

ARCHS4 Tissues random 4000 genes

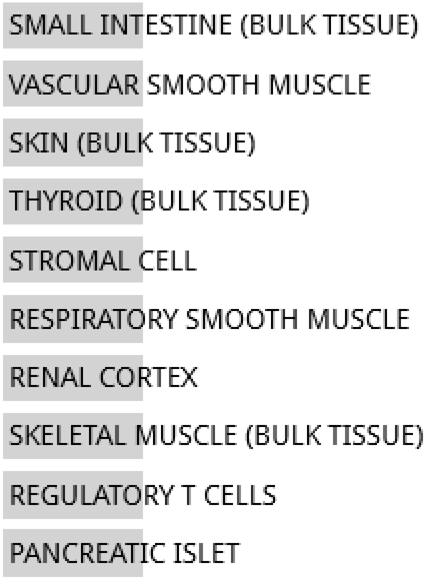


**NO SIGNIFICANT ENRICHMENT**

Jensen TISSUES random 4000 genes

**NO DATA AVAILABLE**

**Random Set 17**

Jensen DISEASES random 4000 genes

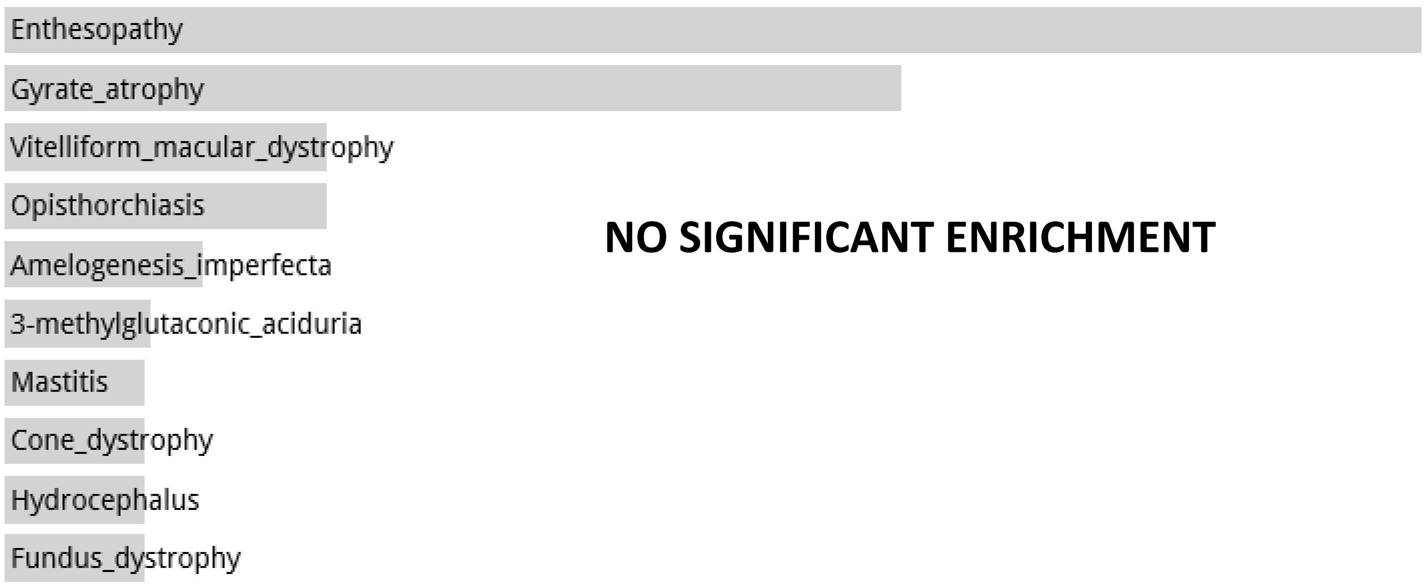


Disease Perturbations from GEO up random 4000 genes

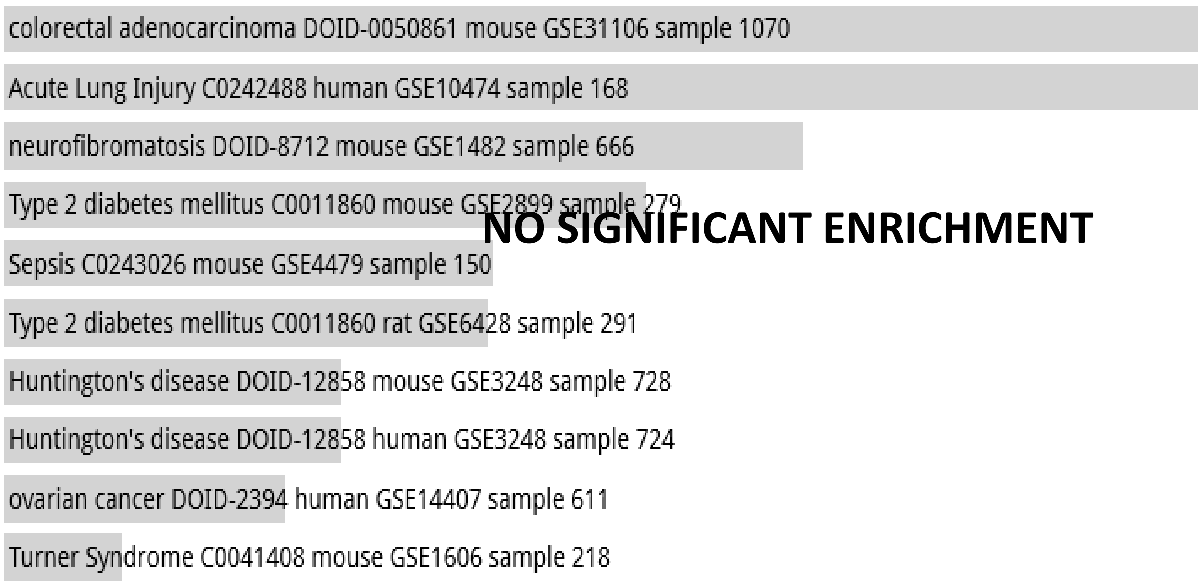


Disease Perturbations from GEO down random 4000 genes

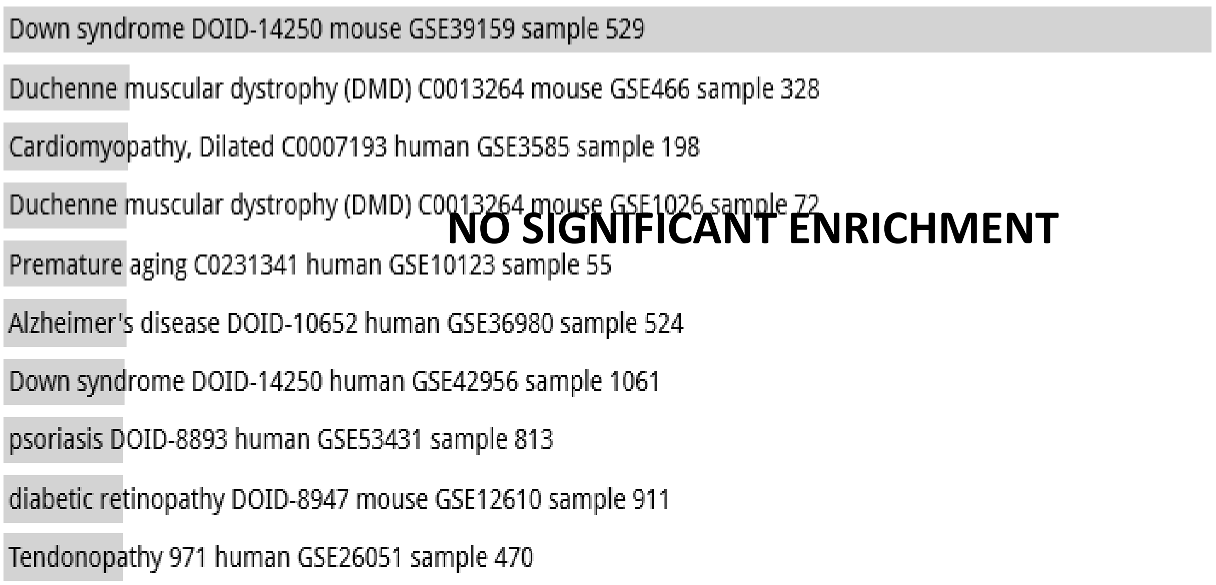

#### Random Set 17

### Allen Brain Atlas up random 4000 genes

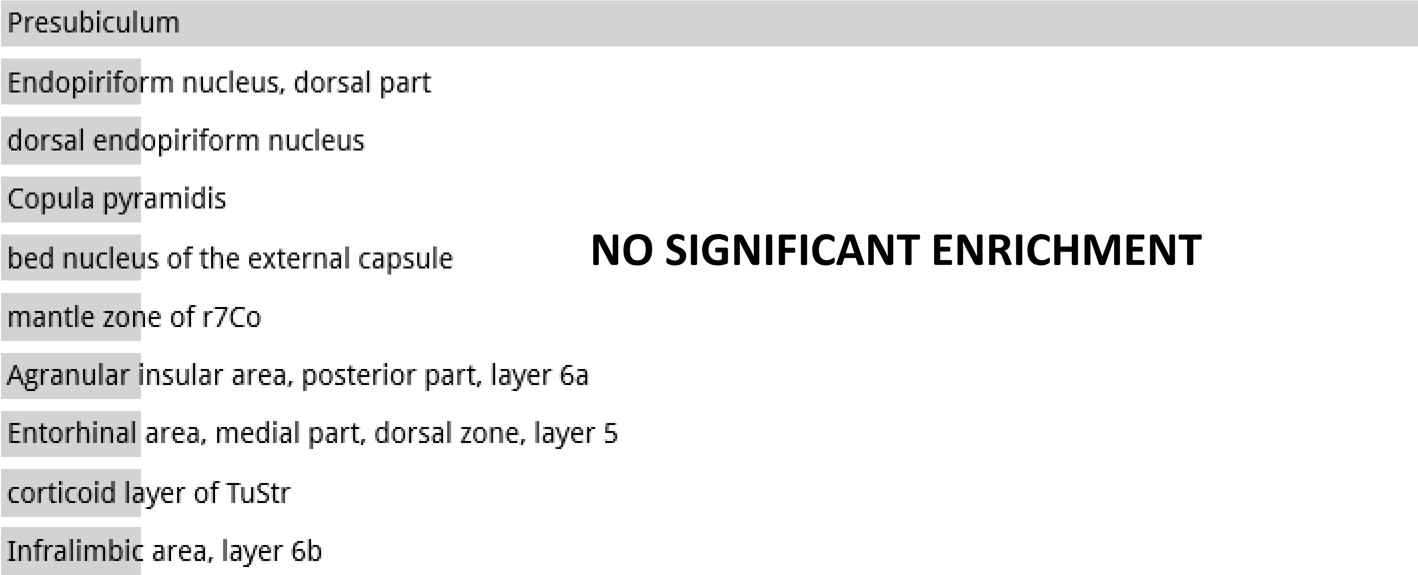


**NO SIGNIFICANT ENRICHMENT**

**Random Set 18**

ARCHS4 Tissues random 4000 genes

**NO DATA AVAILABLE**

Jensen TISSUES random 4000 genes

**NO DATA AVAILABLE**

**Random Set 18**

Jensen DISEASES random 4000 genes

**NO DATA AVAILABLE**

Disease Perturbations from GEO up random 4000 genes

**NO DATA AVAILABLE**

Disease Perturbations from GEO down random 4000 genes

**NO DATA AVAILABLE**

**Random Set 18**

Allen Brain Atlas up random 4000 genes

**NO DATA AVAILABLE**

**Random Set 19**

ARCHS4 Tissues random 4000 genes

**NO DATA AVIALABLE**

Jensen TISSUES random 4000 genes

**NO DATA AVAILABLE**

**Random Set 19**

Jensen DISEASES random 4000 genes

**NO DATA AVAILABLE**

Disease Perturbations from GEO up random 4000 genes

**NO DATA AVAILABLE**

Disease Perturbations from GEO down random 4000 genes

**NO DATA AVAILABLE**

**Random Set 19**

Allen Brain Atlas up random 4000 genes

**NO DATA AVAILABLE**

**Random Set 20**

ARCHS4 Tissues random 4000 genes

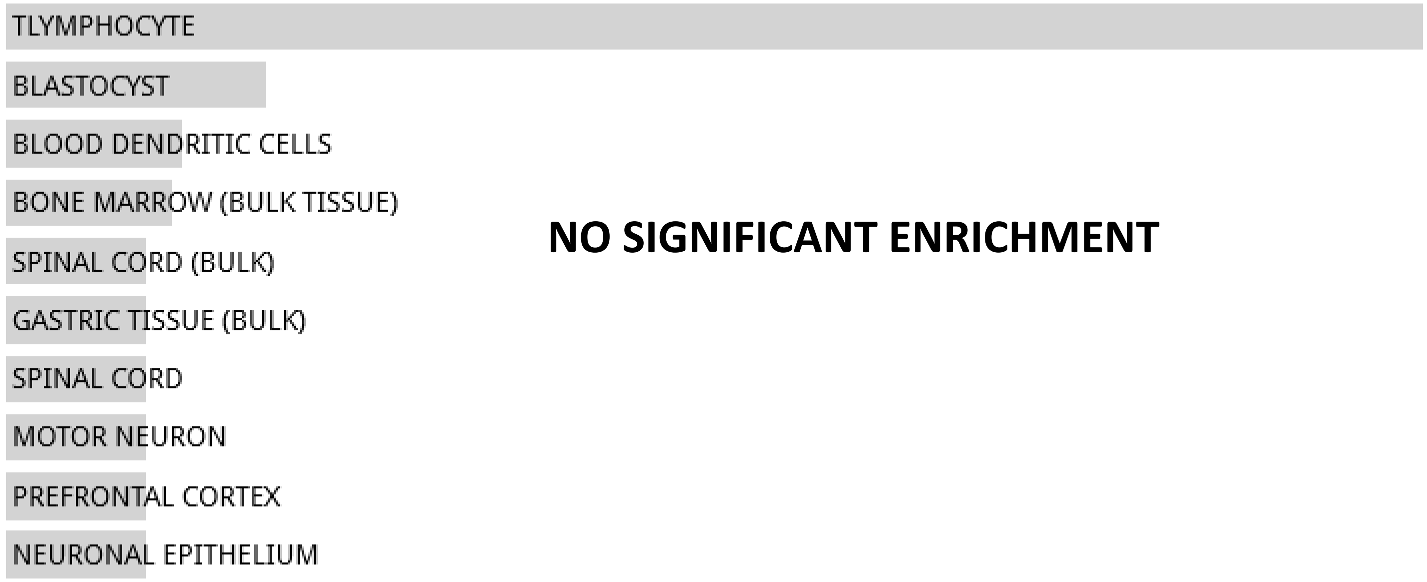


**NO SIGNIFICANT ENRICHMENT**

Jensen TISSUES random 4000 genes

**NO DATA AVAILABLE**

**Random Set 20**

Jensen DISEASES random 4000 genes

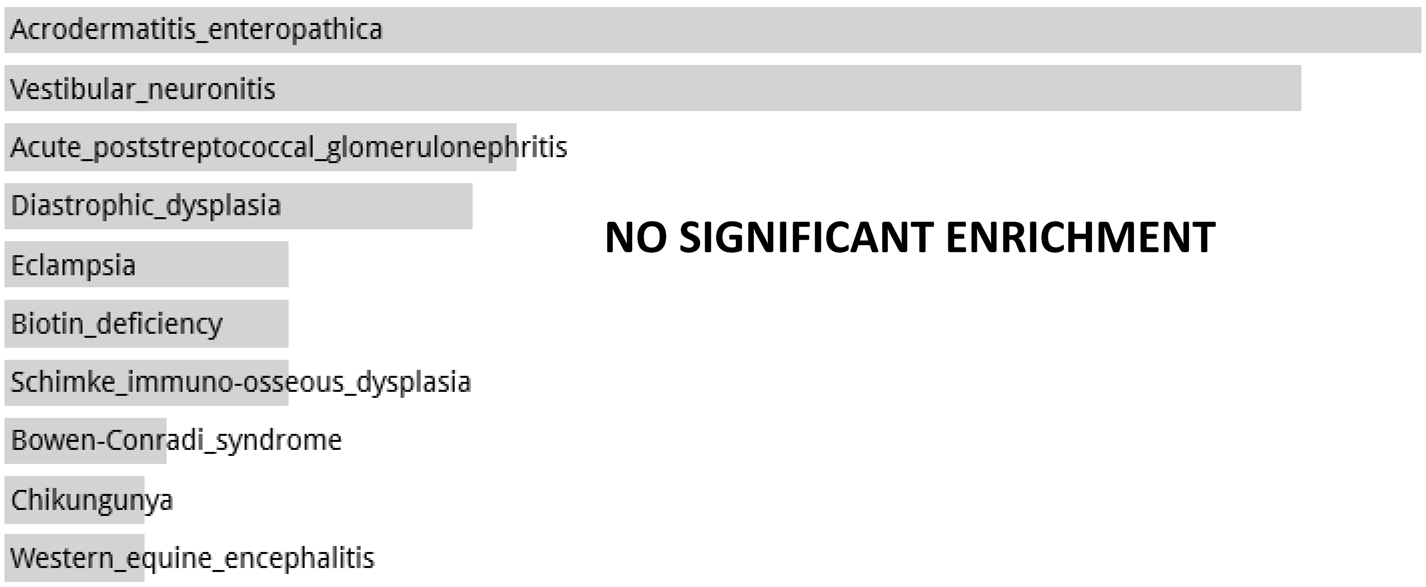


Disease Perturbations from GEO up random 4000 genes

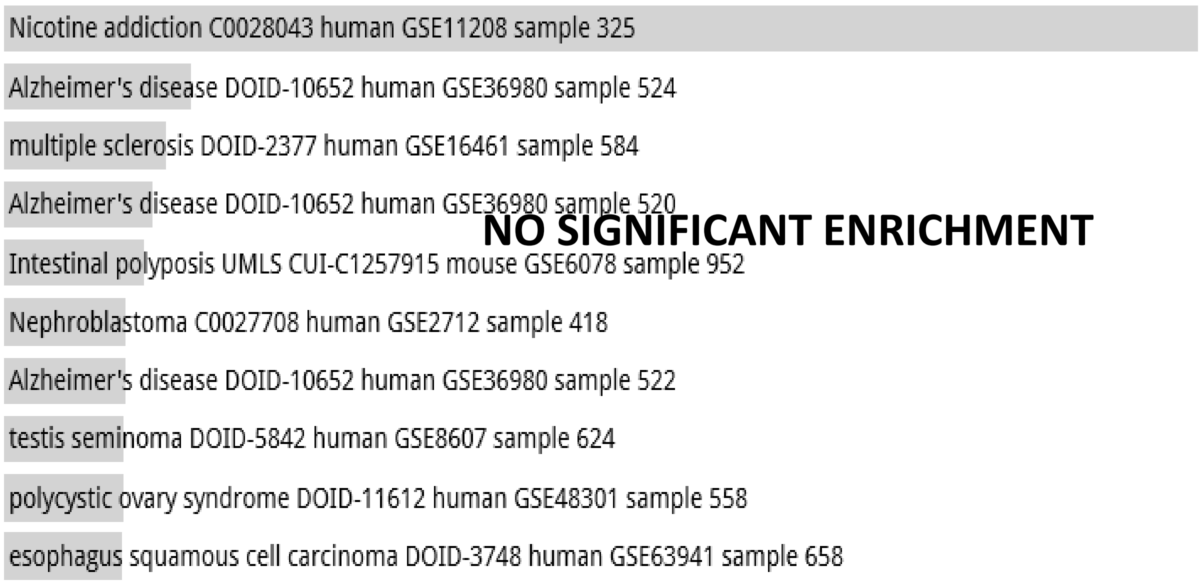


Disease Perturbations from GEO down random 4000 genes

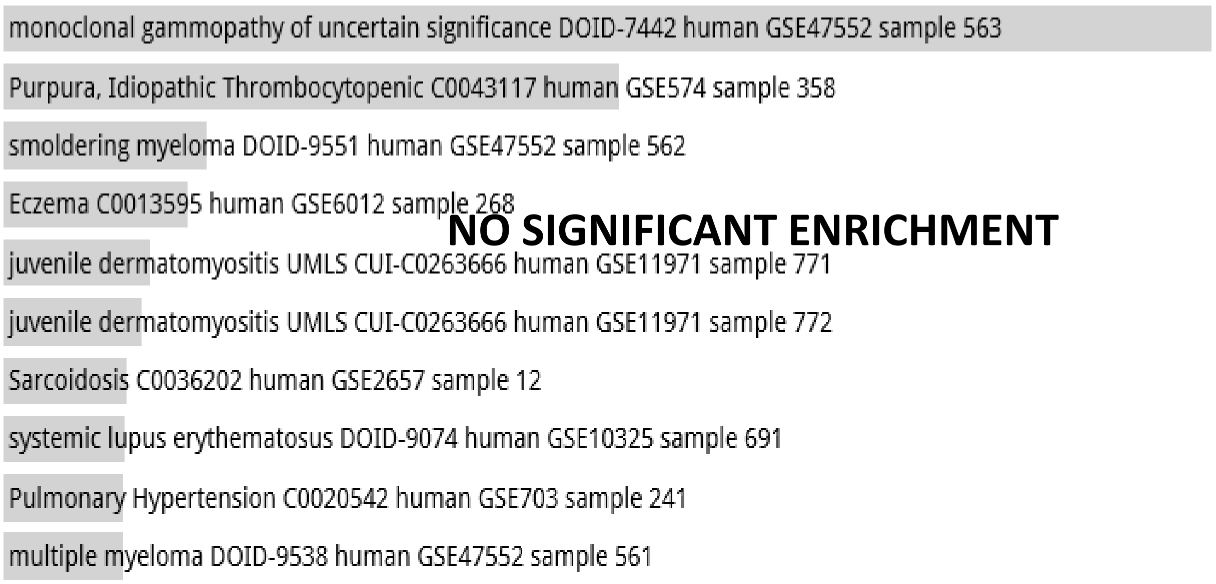

**Random Set 20**

Allen Brain Atlas up random 4000 genes

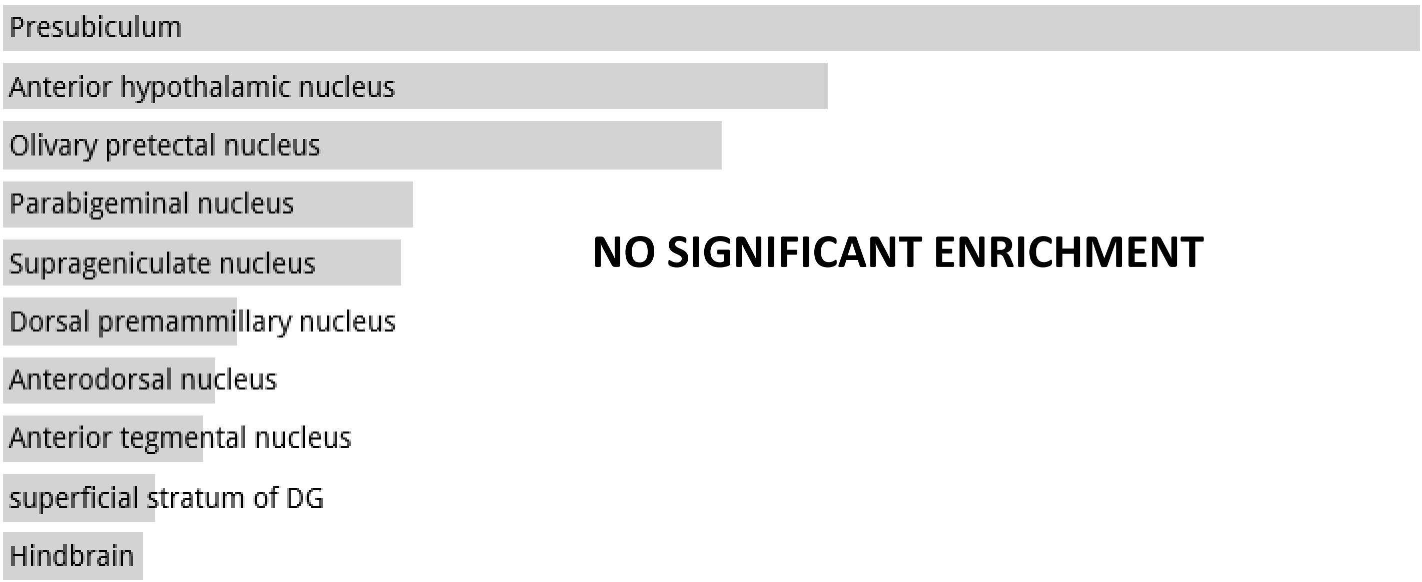


**NO SIGNIFICANT ENRICHMENT**

**Random Set 21**

ARCHS4 Tissues random 2847 genes

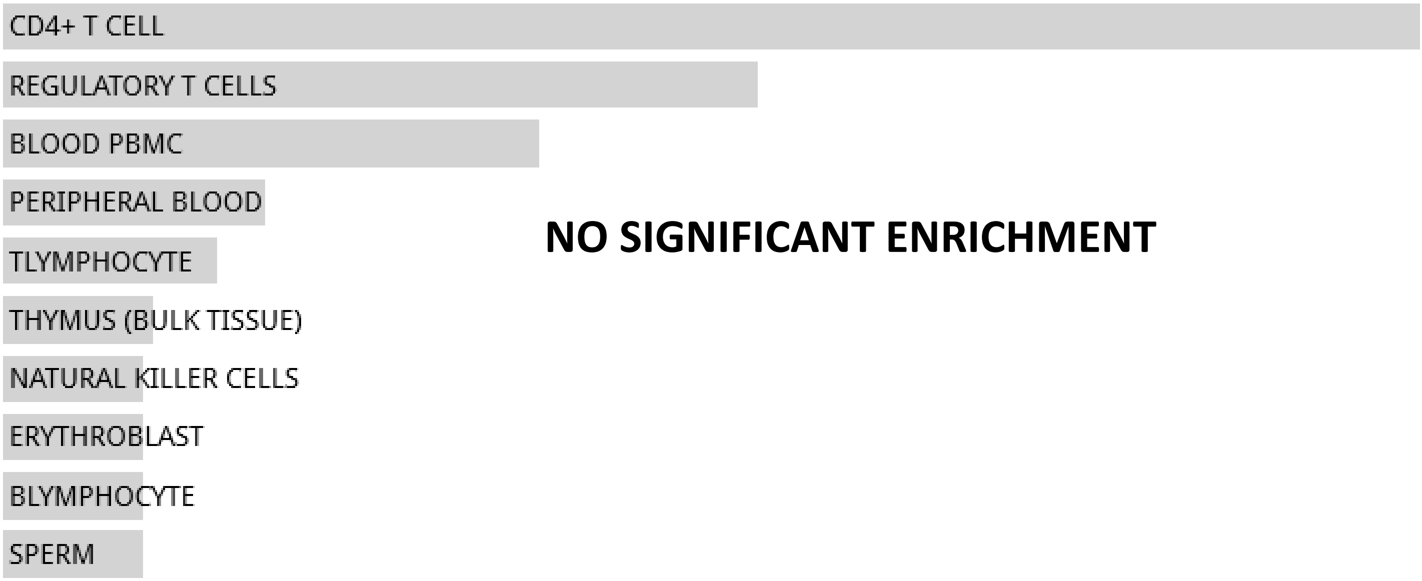


**NO SIGNIFICANT ENRICHMENT**

Jensen TISSUES random 2847 genes

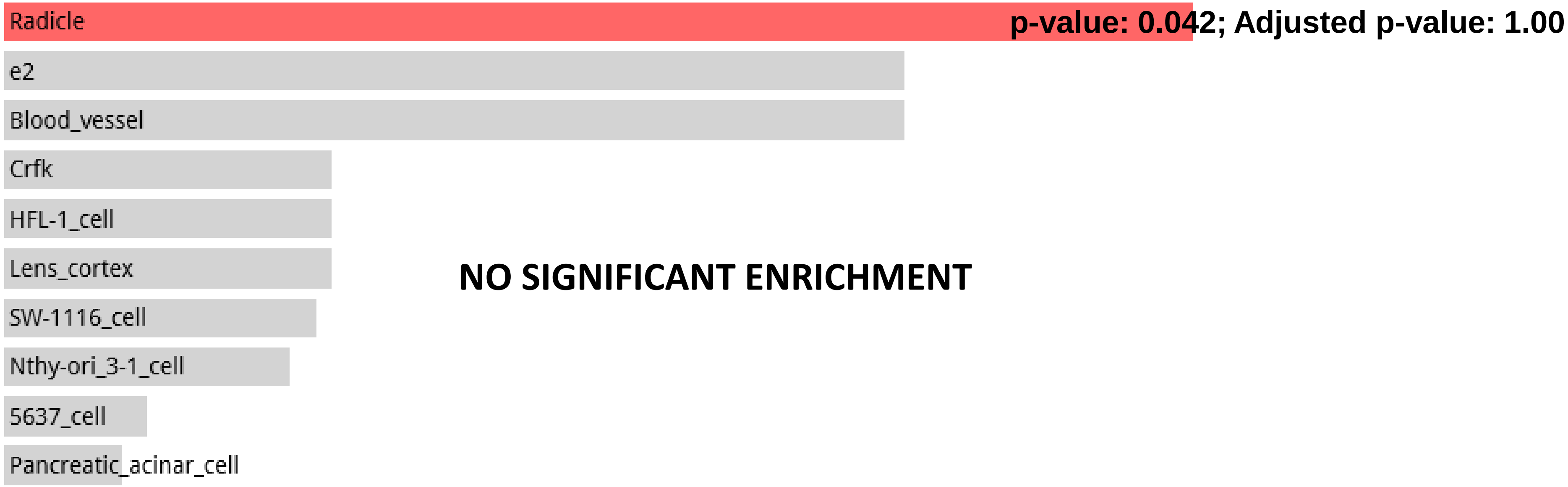


**NO SIGNIFICANT ENRICHMENT**

#### Random Set 21

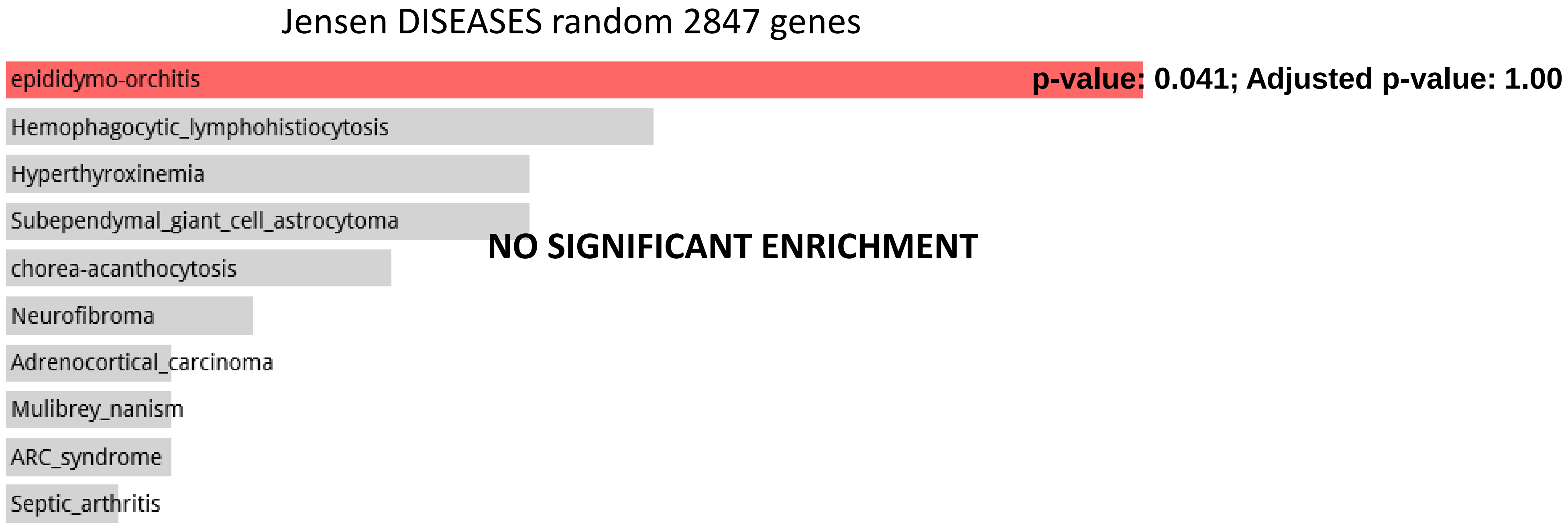


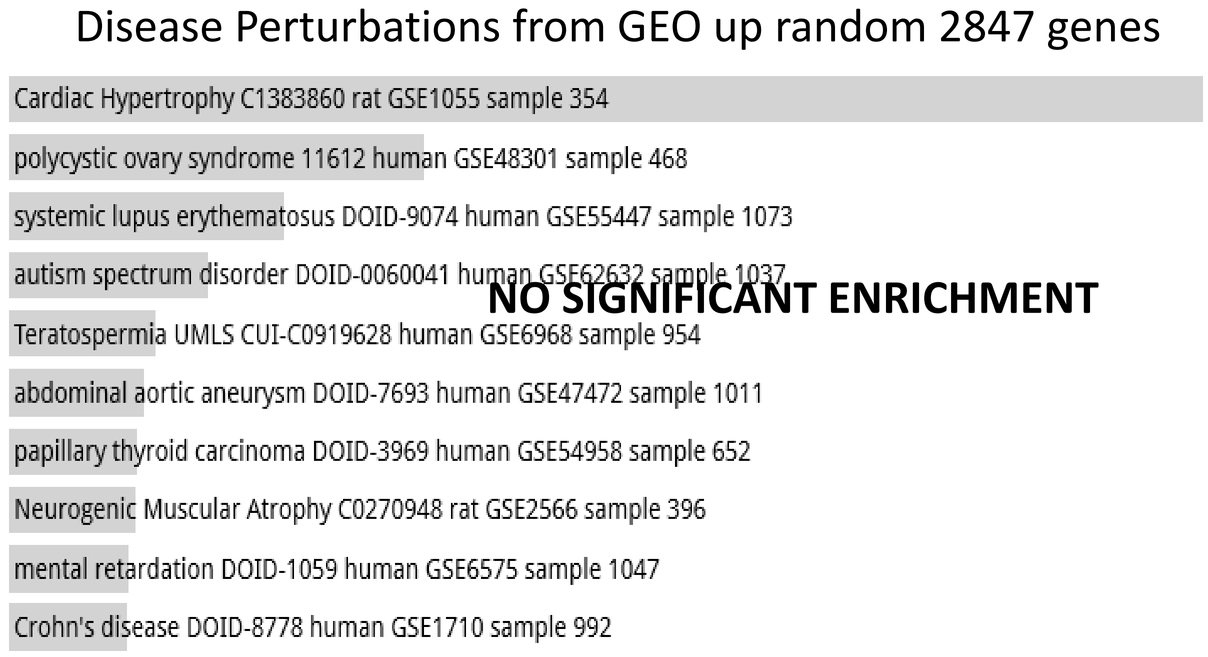


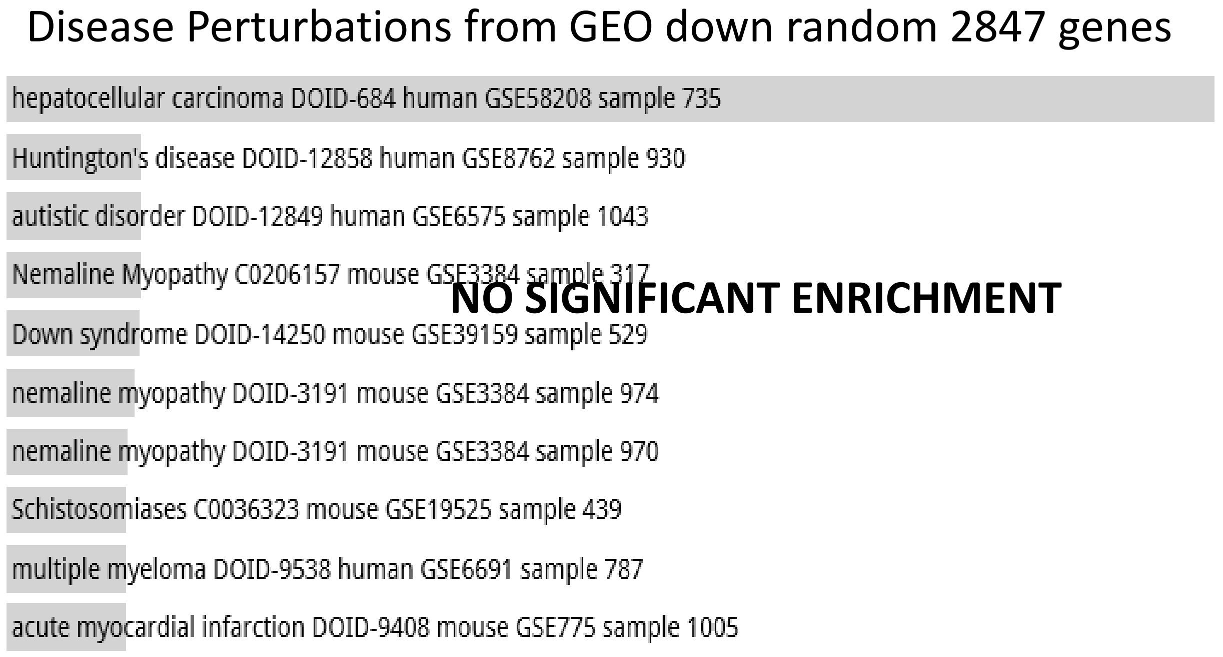

**Random Set 21**

#### Allen Brain Atlas up random 2847 genes

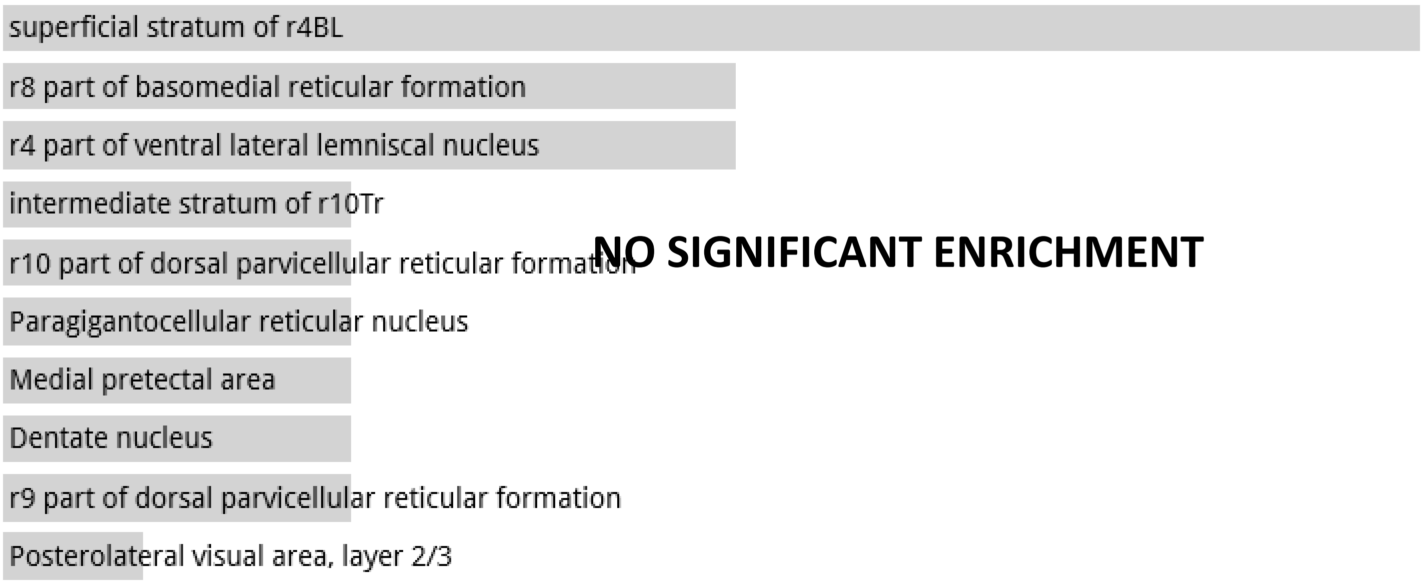


#### GO Biological Process 2017b 2847 genes

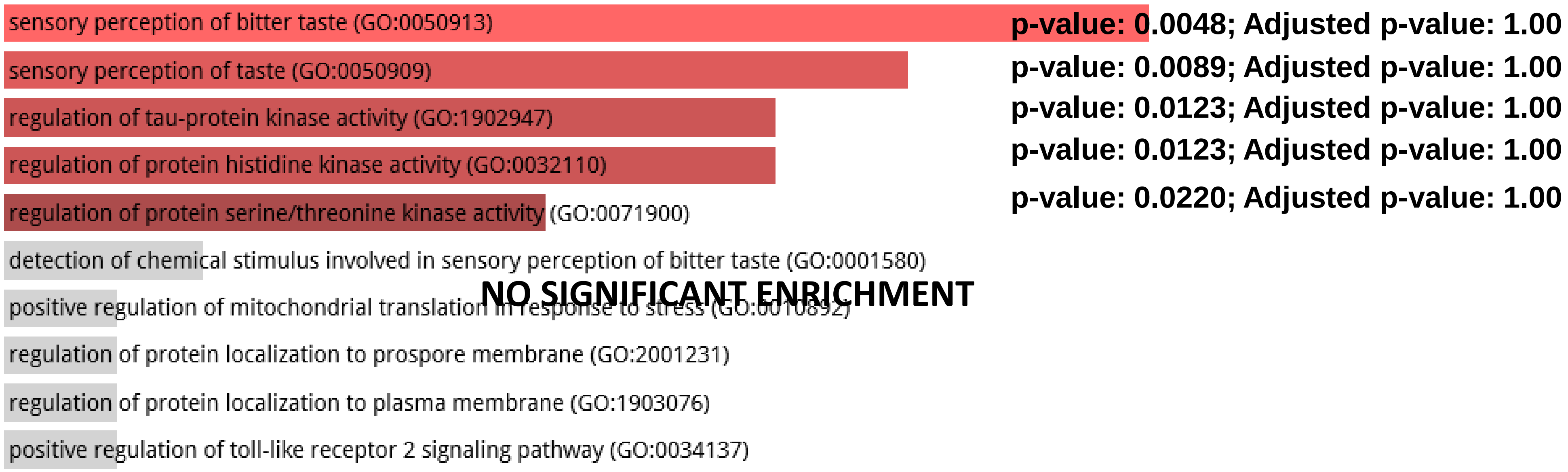

**Random Set 21**

GO Molecular Function 2017 2847 genes

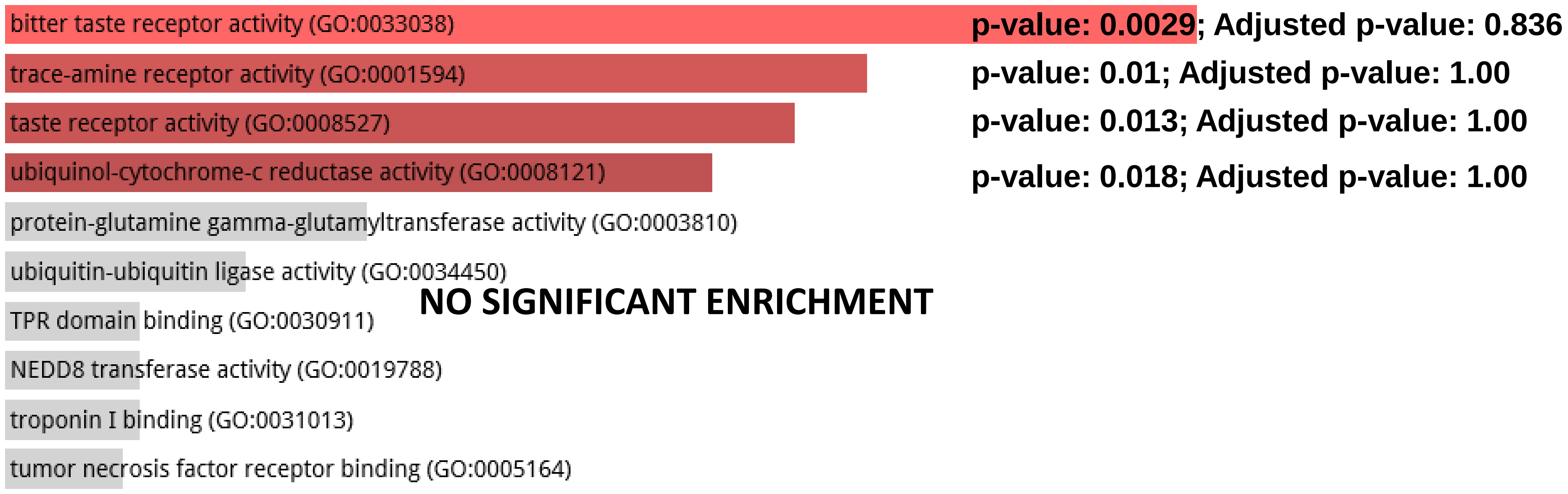


Pfam InterPro Domains 2847 genes

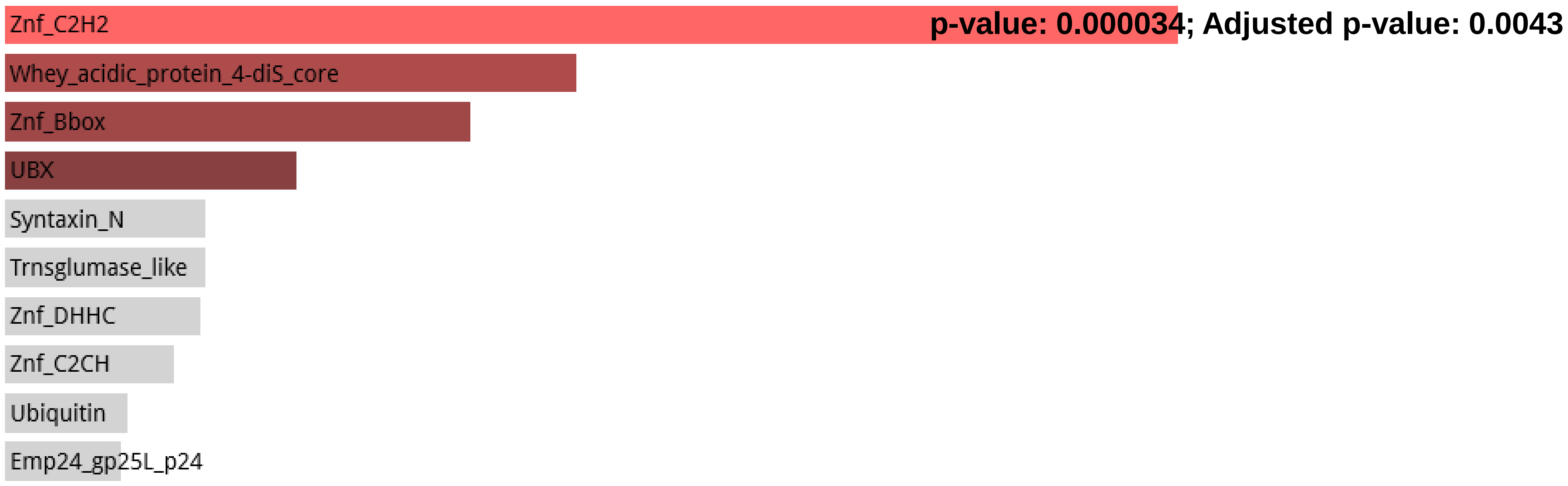

#### Random Set 21

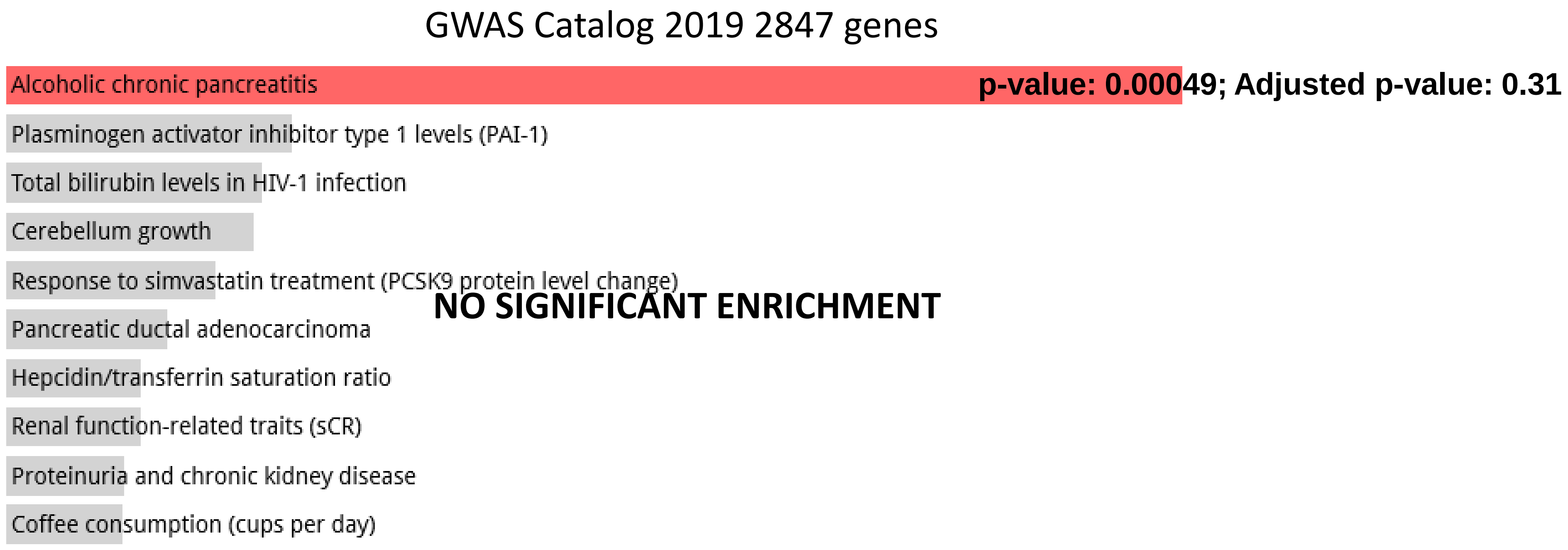


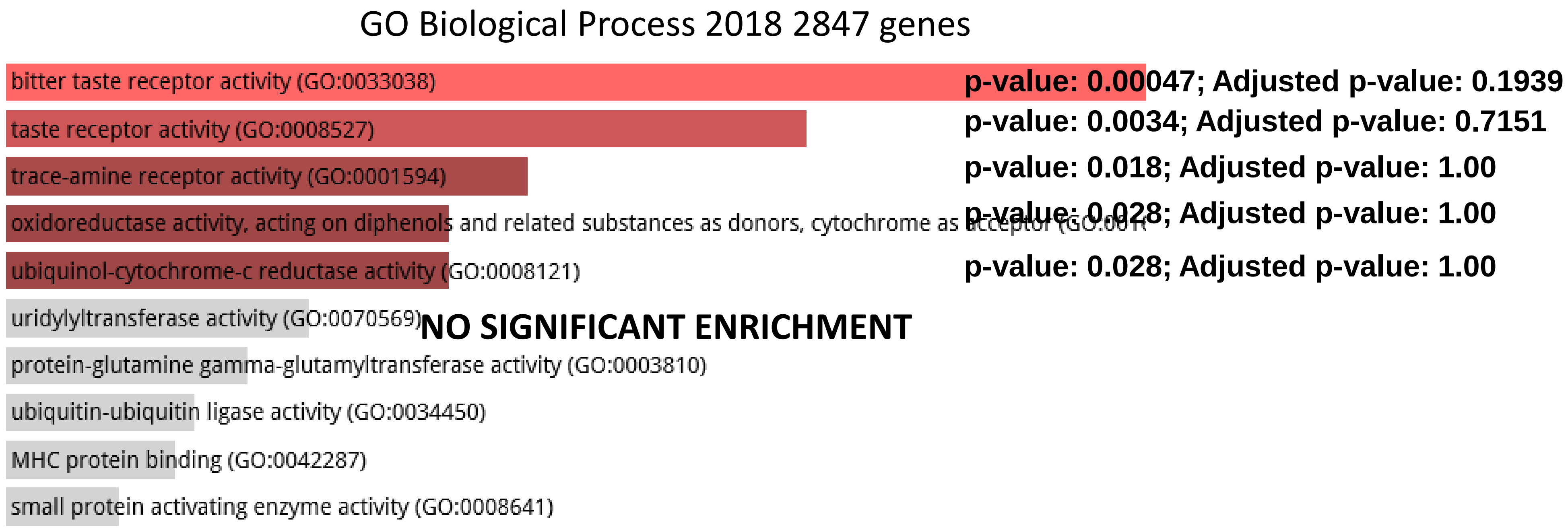

#### Random Set 21

CORUM 2847 genes

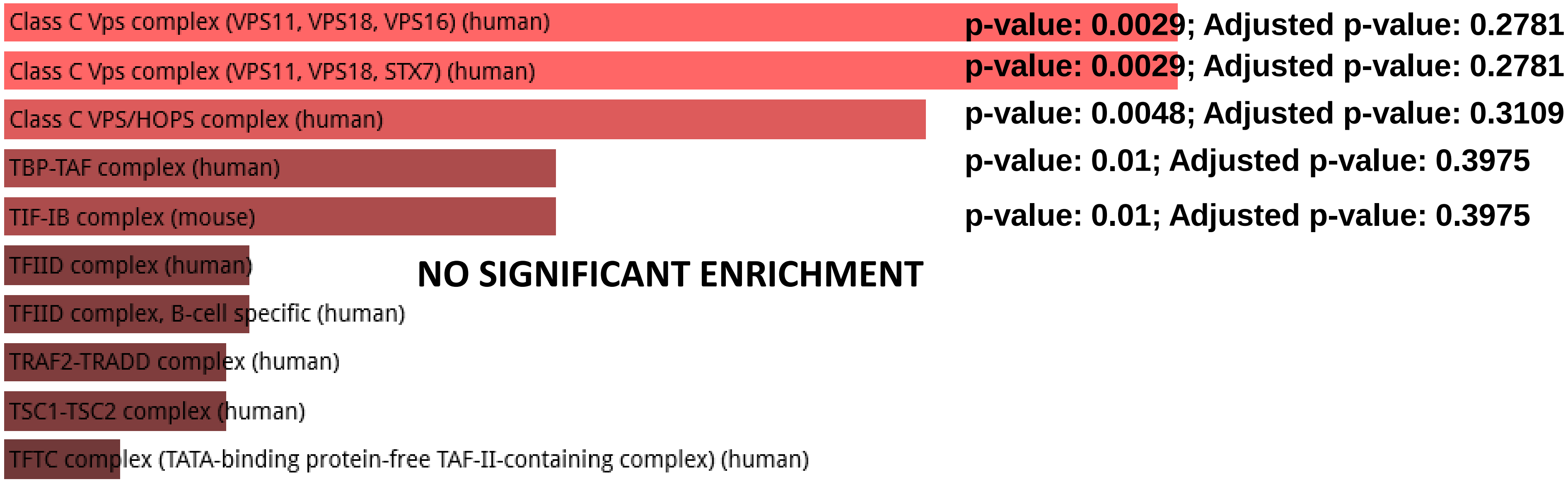


**CORUM database of protein complexes identified by mass spectrometry**

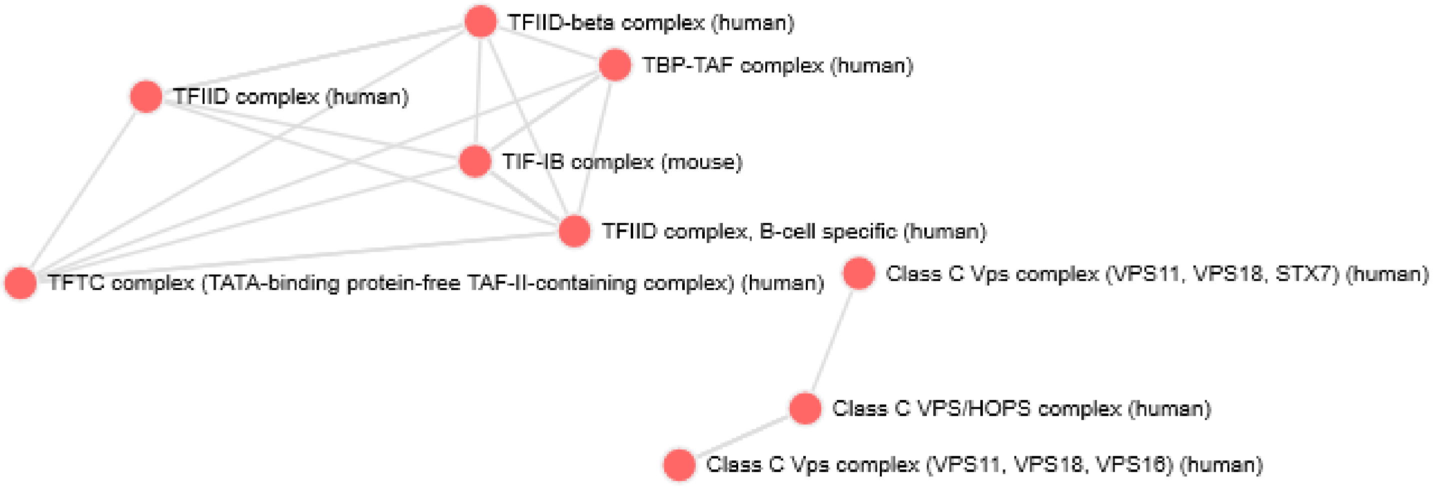

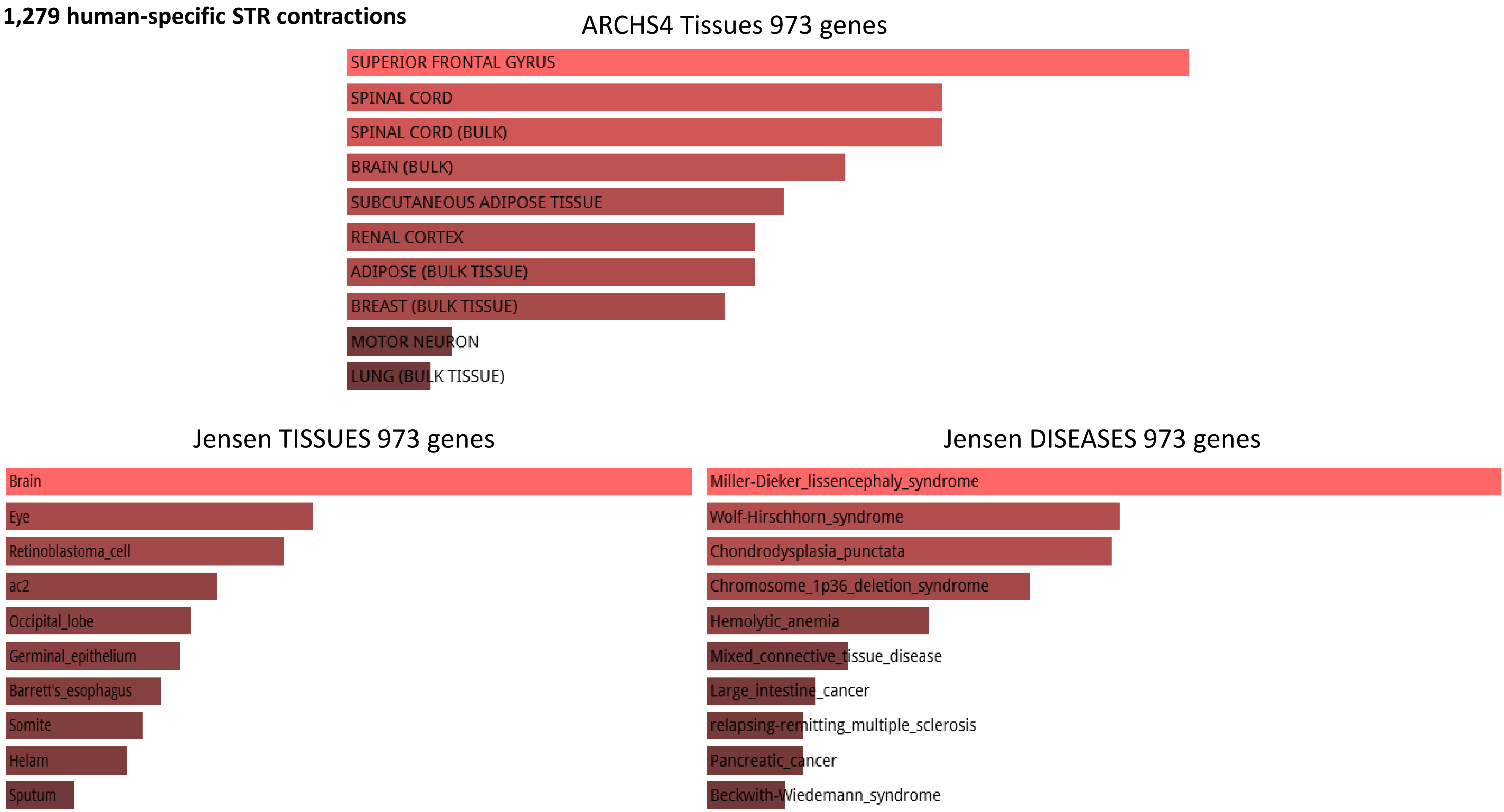
1,279 human-specific STR contractions
ARCHS4 Tissues 973 genes
SUPERIOR FRONTAL GYRUS
SPINAL CORD
SPINAL CORD (BULK)
BRAIN (BULK)
SUBCUTANEOUS ADIPOSE TISSUE
RENAL CORTEX
ADIPOSE (BULK TISSUE)
BREAST (BULK TISSUE)
MOTOR NEURON
LUNG (BULK TISSUE)
Jensen TISSUES 973 genes
Brain
Eye
Retinoblastoma_cell
ac2
Occipital_lobe
Germinal_epithelium
Barrett's_esophagus
Somite
Helam
Sputum
Jensen DISEASES 973 genes
Miller-Dieker_lissencephaly_syndrome
Wolf-Hirschhorn_syndrome
Chondrodysplasia_punctata
Chromosome_1p36_deletion_syndrome
Hemolytic_anemia
Mixed_connective_tissue_disease
Large_intestine_cancer
relapsing-remitting_multiple_sclerosis
Pancreatic_cancer
Beckwith-Wiedemann_syndrome

**2,118 fixed human-specific regulatory regions (non-hESC DHS)**

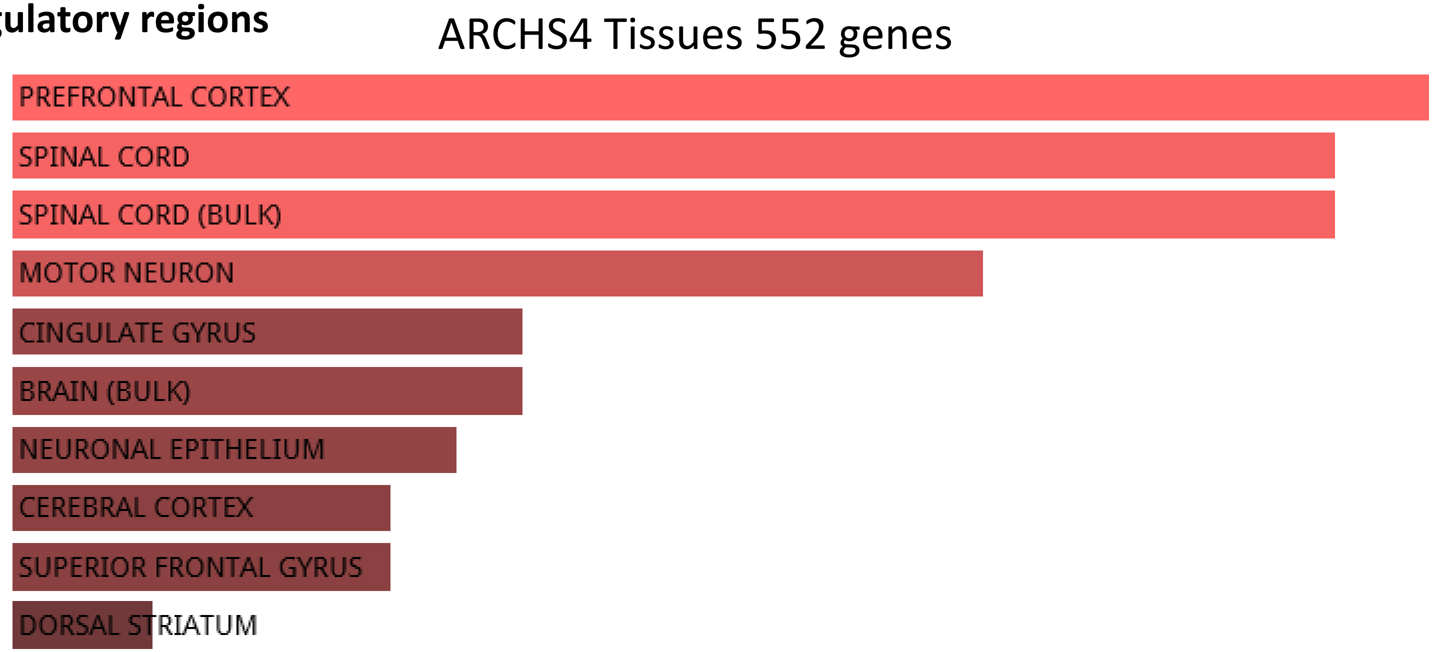


**410 H3K4me3 sites with human-specific enrichment in prefrontal cortex neurons**

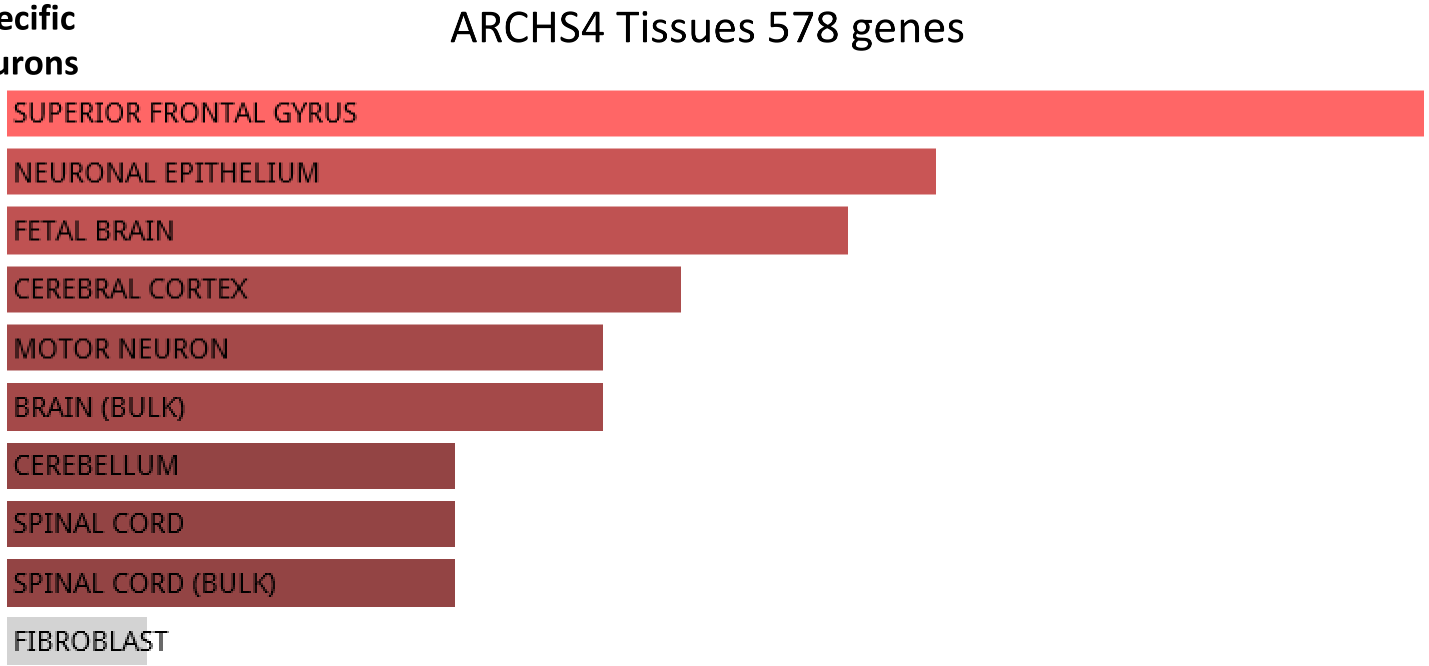
